# Mechanistic description of spatial processes using integrative modelling of noise-corrupted imaging data

**DOI:** 10.1101/284547

**Authors:** Sabrina Hross, Fabian J. Theis, Michael Sixt, Jan Hasenauer

## Abstract

Spatial patterns are ubiquitous on the subcellular, cellular and tissue level, and can be studied using imaging techniques such as light and fluorescence microscopy. Imaging data provide quantitative information about biological systems, however, mechanisms causing spatial patterning often remain illusive. In recent years, spatio-temporal mathematical modelling helped to overcome this problem. Yet, outliers and structured noise limit modelling of whole imaging data, and models often consider spatial summary statistics. Here, we introduce an integrated data-driven modelling approach that can cope with measurement artefacts and whole imaging data. Our approach combines mechanistic models of the biological processes with robust statistical models of the measurement process. The parameters of the integrated model are calibrated using a maximum likelihood approach. We used this integrated modelling approach to study *in vivo* gradients of the chemokine (C-C motif) ligand 21 (CCL21). CCL21 gradients guide dendritic cells and are important in the adaptive immune response. Using artificial data, we verified that the integrated modelling approach provides reliable parameter estimates in the presence of measurement noise and that bias and variance of these estimates are reduced compared to conventional approaches. The application to experimental data allowed the parameterisation and subsequent refinement of the model using additional mechanisms. Among others, model-based hypothesis testing predicted lymphatic vessel dependent concentration of heparan sulfate, the binding partner of CCL21. The selected model provided an accurate description of the experimental data and was partially validated using published data. Our findings demonstrate that integrated statistical modelling of whole imaging data is computationally feasible and can provide novel biological insights.

## Introduction

In the past decades, our understanding of biological processes has been revolutionised by imaging technologies. Nowadays, super-resolved fluorescence microscopy (Huang *et al.*, 2009), light sheet fluorescence microscopy (Santi, 2011), cryo electron microscopy (Al-Amoudi *et al.*, 2004), and other technologies provide information about cell and tissue structures over a broad range of scales. Multiplexed information about intracellular processes is, among others, provided by matrix-assisted laser desorption/ionization (MALDI) imaging mass spectrometry (Cornett *et al.*, 2007) and mass cytometry (Giesen *et al.*, 2014). These imaging data are analysed using tailored image processing pipelines to quantify properties of interest (see (Chenouard *et al.*, 2014) and references therein). This provides detailed information about the imaged system, e.g., biological tissues. Yet, mechanisms often remain elusive, for instance, it is usually not evident from imaging data how the observed spatial patterns are established and controlled. However, such insights are necessary to improve the understanding of complex biological systems (Turing, 1952; Gurdon & Bourillot, 2001).

Model-based approaches have been introduced to unravel the mechanisms underlying the spatio-temporal organisation of tissues (Iber *et al.*, 2015; Uzkudun *et al.*, 2015). Partial differential equation (PDE) models and agent-based models which capture static and dynamic properties of tissue-scale images have been developed (see (Menshykau *et al.*, 2014; Uzkudun *et al.*, 2015; Hersch *et al.*, 2015; Jagiella *et al.*, 2017) and references therein). These models can describe the underlying biological mechanism and allow for the evaluation of competing biological hypotheses.

Modeling and hypothesis testing, however, mostly employ qualitative information (Uzkudun *et al.*, 2015) or summary statistics (Hock *et al.*, 2013b; Hersch *et al.*, 2015; Hross *et al.*, 2016; Jagiella *et al.*, 2017). Qualitative information is used due to limited image quality caused, among others, by limitations of labelling methods. Summary statistics are considered as they are easy to assess using available processing pipelines. Although qualitative abstractions and summary statistics provide only a fraction of the information encoded in the images, they are widely used. A key reason is the use of sequential analysis approaches (Figure 1A) which exploit established image processing pipelines.

**Figure 1:**
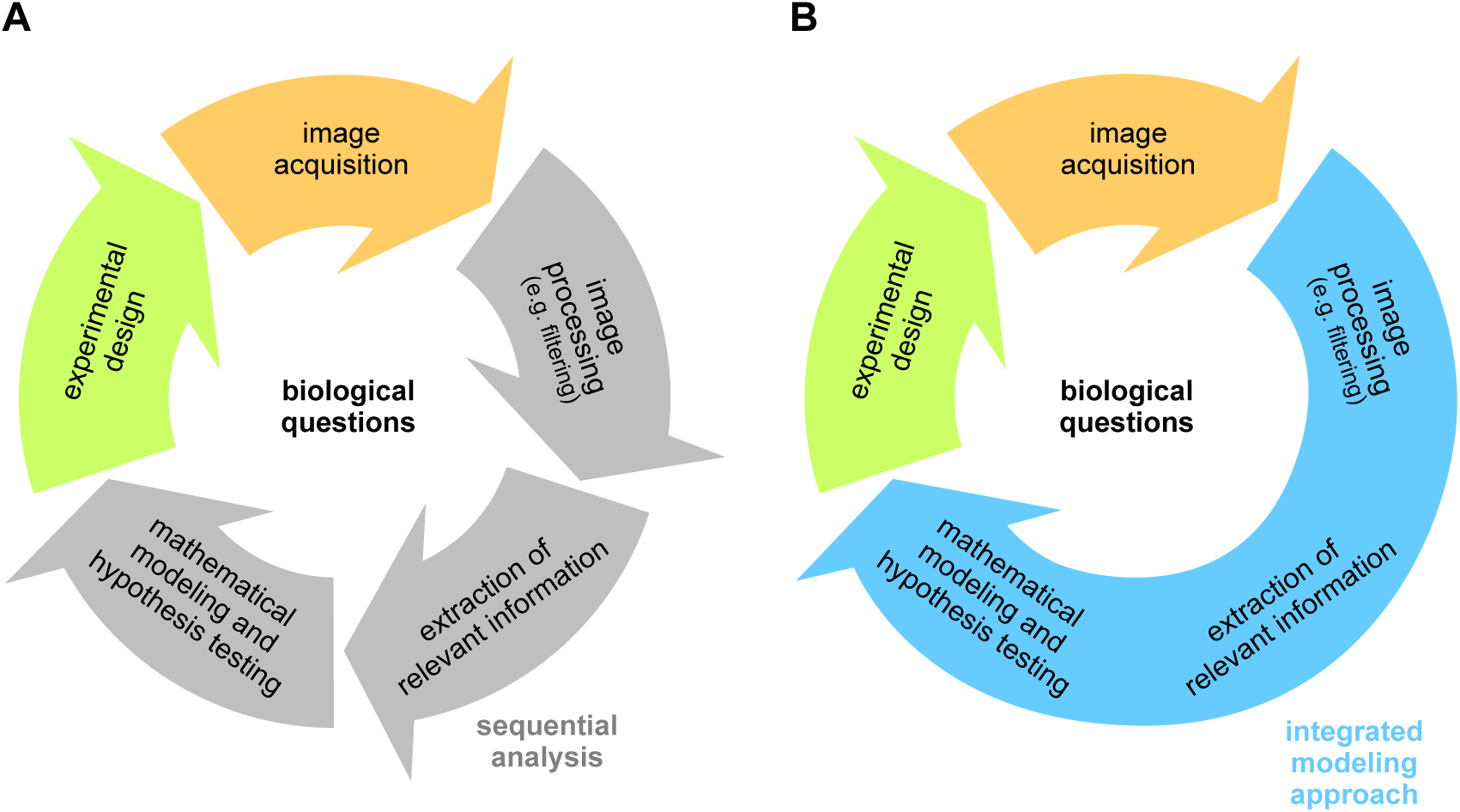
Illustration of data-driven modelling in image-based systems biology. (A) Sequential analysis relying on image processing and extracted features. (B) Integrated modelling approach combining image processing, information retrieval and modelling in a single step.

In this manuscript, we propose an integrated modelling approach for imaging data (Figure 1B). The proposed framework combines image processing with the mechanistic description of the biochemical process using PDE models, instead of performing these steps sequentially. To account for outliers and structured measurement noise, e.g., signals generated by biological processes not considered in the model, we employ concepts from robust regression (Lange *et al.*, 1989; Peel & McLachlan, 2000). The integrated modelling approach facilitates the simultaneous assessment of the quality of the imaging data, the filtering of outliers and artefacts, and the mechanistic modelling of the biological process. As this integrated framework circumvents preprocessing and the extraction of summary statistics, it avoids a potential information loss and provides a tailored, unbiased filtering. By avoiding the tuning of parameters in the preprocessing, the approach furthermore simplifies the workflow and promises an improved reproducibility of analysis results.

We implemented the integrated modelling approach and assessed it by studying artificial and experimental data for the formation of gradients of the chemokine CCL21, a process relevant in the immune response. Using this process, we demonstrate the loss of information associated with the use of summary statistics as well as the influence of structured noise on estimation results. Subsequently, we demonstrate how the integrated modelling framework facilitates the direct use of noise-corrupted whole imaging data. We exploit the integrated framework to generate novel hypotheses regarding the underlying biochemistry, which are partially validated using literature data.

## Methods

In this manuscript, we present an integrated modelling approach for tissue-scale imaging data. In the following, we outline the considered modelling approaches, data types, and inference methods.

### Mechanistic model of spatio-temporal biological processes

We consider spatio-temporal biological processes described by reaction-diffusion equations – a class of PDE models. Reaction-diffusion equations are widely used in systems and computational biology, among other to capture the dynamics of intra- and extracellular substances (Efendiev, 2013).

The state variable *u*(*x, t*) *∈* R*n* of the PDE model is the abundance of *n* chemical substances at time *t* and spatial location *x ∈* Ω. The state is defined on the modelled spatial domain Ω and changes due to diffusion and biochemical reactions. The Laplace operator is denoted by *6*, the matrix of diffusion coefficients by *D*(*θ*) and the reaction term by *f* (*u*(*x, t*), *x, θ*). This yields the PDE model

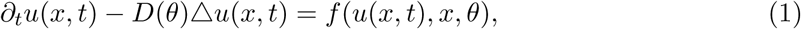

with initial condition *u*(*x*, 0) = *u*_0_(*x, θ*) and boundary conditions defined on the boundary *∂*Ω of the spatial domain Ω, e.g., Dirichlet or Neumann boundary conditions. Diffusion coefficients, reaction term, initial condition and boundary conditions can depend on the unknown parameters *θ*, e.g., binding affinities and degradation rates.

### Statistical modelling of imaging data

We consider standard image acquisition technologies which provide intensity averages over pixels (or voxels). The spatial domain of the *j*-th pixel is denoted by Ω_*j*_, *j* = 1, *…*, *n*_*j*_. This yields the observation model for the *i*-th observable,

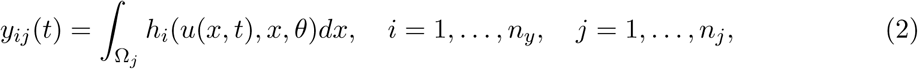

in which *h*_*i*_ describes the dependence of the *i*-th observable on the state variables, *i* = 1, *…*, *n*_*y*_. For biological systems which equilibrate fast, only the stationary distribution might be observed (*t→ ∞*). A typical observation in imaging is the measurement of the relative abundance of a biochemical species, yielding *h*_*i*_(*u*(*x, t*), *x, θ*) = *s*(*u*_*l*_(*x, t*) + *b*) with scaling factor*s*, background *b* and concentration *u*_*l*_(*x, t*) of the *l*-th biochemical species. Saturation effects, unequal elimination, cross-reactivity of antibodies and many other effects can be modelled using the function *h*.

The intensity values of individual pixels, *y*_*ij*_(*t*_*k*_), are corrupted by experimental noise, providing the measured pixel intensities 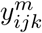. In many applications, the measurement noise is assumed to be independent and identically distributed, e.g., multiplicative log-normally distributed measurement noise,

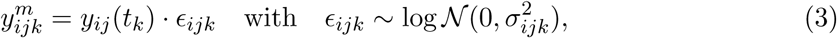

with time points *t*_*k*_, *k* = 1, *…*, *n*_*k*_. In many imaging applications, the measurement noise is however not independent and identically distributed but possesses additional structure. Labelling artefacts or other biological processes which alter the measured intensities result in spatially structured noise. Adjacent pixels exhibit often similar noise levels and regions of high noise might also possess particular shapes. While this is known, a noise model capturing these effects is currently not available. In the following sections, we propose methods to address such structured noise.

The collection of all imaging data is in the remainder denoted by *𝒟*. Furthermore, the unknown observation parameters, i.e., scaling and background, and noise levels are included in the parameter vector *θ*.

### Reconstruction of biological processes from imaging data

To achieve a mechanistic understanding of spatio-temporal biological processes, we want (i) to infer the parameters of model (1) and (ii) to perform model selection to compare competing hypotheses. To address these problems, we consider three alternative statistical approaches:

- *Direct approach:* The presence of outliers is disregarded and the model is fitted to the data using standard noise models (3).
- *Filtering approach:* The measurement data are preprocessed to detect and remove outliers. The model is fitted to the remaining data using standard noise models (3).
- *Integrated modelling approach:* A statistical model for the outlier distribution is formulated. From outlier and noise distribution a likelihood function is derived and used to simultaneously fit the model and quantify the noise level.

In the following, these approaches are described in further detail.

#### Direct approach

The likelihood of observing the imaging data *D* given the parameter vector *θ* is

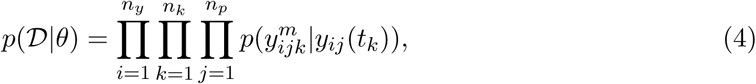

in which 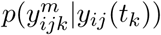denotes the noise model for an individual pixel and *y*_*ij*_(*t*_*k*_) denotes the parameter-dependent solution of the model (1) &(2) (Hock *et al.*, 2013a). For multiplicative log-normally distributed measurement noise (3), we obtain

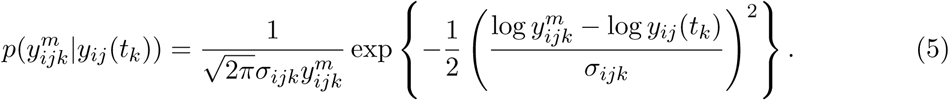

The likelihood function (4) is formulated using the measured intensity values of individual pixels, 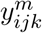, as data points. Alternatively, summary statistics of the pixel intensities can be considered. In the application of gradient formation discussed later, the average intensity as function of the distance from the nearest vessel is used (Weber *et al.*, 2013).

#### Filtering approach

To reduce the impact of outliers and structured noise on the estimation results, image data are preprocessed. We consider filtering methods which provide an index set of filtered pixels, *ℱ*_*ik*_ *⊂ {*1, *…*, *n*_*p*_*}*, for the individual observables *y*_*i*_ and time points *t*_*k*_. These index sets are masks for regions to be excluded from the objective function. Accordingly, the likelihood is only evaluated for the unfiltered pixels, meaning that for the index *j* in (4) only the set *j ∈ {*1, *…*, *n*_*p*_*} \ ℱ*_*ik*_ is considered. Appropriate filtering should render parameter estimation more robust against outliers and structured noise.

Filtering can be performed using a variety of algorithms, most of which possess several tuning parameters which have to be chosen manually or in a semi-automated fashion. The choice of algorithm and tuning parameters depends on the type of structured noise. To remove bright spots from the image, maximally stable extremal region (MSER) filtering (Matas *et al.*, 2004) can be employed. MSER filtering is based on a water shedding mechanism and has been used successfully in a series of studies (see, e.g., (Buggenthin *et al.*, 2013)).

#### Integrated modelling approach

We propose to circumvent the selection of filtering algorithms and the manual tuning of filtering parameters by integrating filtering and parameter estimation. Our integrated modelling approach requires a sufficiently flexible statistical model, ideally accounting for standard measurement noise, structured noise and outliers as well as spatial correlation structure (see Section *Statistical modelling of imaging data*). In this study, we follow ideas from robust regression, i.e., *E*-contamination models (Berger & Berliner, 1986), to address these needs. We assume that the intensity measurement for each pixel is with probability *w*_*o*_ an outlier/artefact generated by structured noise and with probability 1 *- w*_*o*_ no outlier. The outliers are assumed to be distributed according to the density function *p*_*o*_ while the remaining points are distributed according to the standard noise model (3). This yields the likelihood function

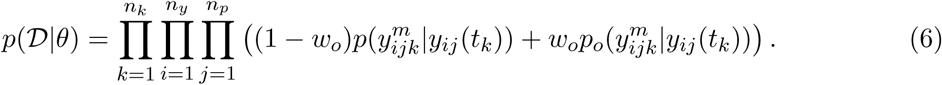

Different outlier distributions can be used given the biological application and the imaging technique. Here, we consider the outliers to be log-normally distributed with location parameter 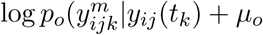 and scale parameter *σ*_*o*_,

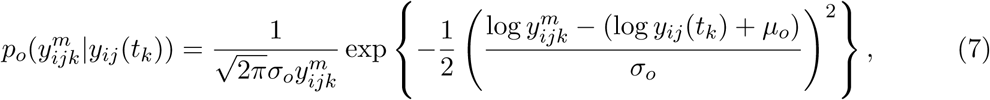

The parameters (*w*_*o*_, *µ*_*o*_ and *σ*_*o*_) ensure the flexibility of the statistical model. This can reduce the bias introduced by measurement artefacts compared to using only the standard noise models (*𝒲*_*o*_ = 0). The inclusion of these additional parameters in the parameter vector *θ* allows for the simultaneous calibration of the models for the biological and the measurement process.

Conceptually, integrated statistical modelling weights the impact of a data-point on the model fit, while the standard filtering approach employs a hard cut-off. The weighting depends on the model-data agreement in different regions of the image, providing a context-dependent filter.

### Parameter estimation and model selection

The analysis of measurement data *D* using the different statistical approaches requires the estimation of the parameters *θ ∈* Θ *⊆* R*^nθ^*. For this, we use maximum likelihood (ML) estimation. The ML estimate of the parameter vector, 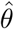, is the solution of the PDE-constrained optimisation problem

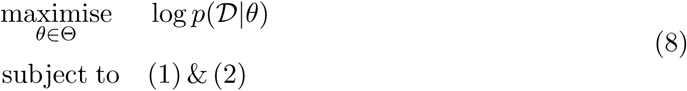

with log-likelihood function log *p*(*𝒟|θ*). The log-likelihood function varies between approaches while the models of the biological process (1) and the measured intensities (2) remain the same.

Optimisation problem (8) is usually nonlinear and can possess multiple local optima. To determine the global optimum, we employ a multi-start local optimisation method. The starting points are sampled from Θ using latin hypercube sampling. For local optimisation an interior point algorithm is used, which is supplied with gradients computed using forward sensitivity equations. This multi-start approach is computationally efficient and reliable for a broad range of applications (see (Raue *et al.*, 2013; Hross & Hasenauer, 2016) and references therein). Instead of multi-start local optimisation, also evolutionary and genetic algorithms (Bäck, 1996), particle swarm optimisers (Yang, 2010) or hybrid optimisers (Vaz & Vicente, 2007) could be employed. For a comprehensive survey and evaluations we refer to the work of Moles *et al.* (2003) and Raue *et al. m* (2013).

The parameter estimates are usually subject to uncertainty due to limited and noise-corrupted data. We determine the uncertainty of the estimated parameters using structural and practical identifiability analysis. For practical identifiability, profile likelihoods are computed (Murphy & van der Vaart, 2000; Raue *et al.*, 2009), which provide parameter confidence intervals to particular confidence levels. For profile likelihood calculation we use the methods recently described for parameter estimation problems with PDE constraints (Boiger *et al.*, 2016).

Biological processes are still poorly understood and there are usually competing hypotheses giving rise to different model structures. To assess the plausibility of hypotheses, we use the Bayesian Information Criterion (BIC) (Schwarz, 1978). The BIC accounts for model-data mismatch and the complexity of the model, measured by the number of parameters. Models with lower BIC values are preferable and a difference of *≥* 10 is considered as substantial (Burnham & Anderson, 2004). Model comparison using BIC and other statistical approaches assumes that all models consider the same dataset. As the filtering approach excludes data points, a comparison between approaches using model selection is not possible. We use model selection merely to compare model alternatives fitted using the same statistical approach.

### Implementation

All methods are implemented in MATLAB and available as Supporting Code S1. The simulation of the PDE model is implemented using the Partial Differential Equation Toolbox of MATLAB. The multi-start local optimisation exploits the MATLAB routine fmincon.m. Parameter estimation and uncertainty analysis are performed using the Parameter EStimation TOolbox (PESTO) available on GitHub (https://github.com/ICB-DCM/PESTO) (Stapor *et al.*, 2017).

## Results

In the following, we will illustrate the reliability achieved using whole imaging data and spatial summary statistic, and compare direct, filtering and integrated modelling approach for statistical inference from imaging data. For this purpose, we studied artificial imaging data, for which the ground truth is known, as well as experimental imaging data, from which new biological insights are gained.

### Biological process

We studied the distribution of the chemokine (C-C motif) ligand 21 (CCL21) in dermal interstitium (Figure 2A). CCL21 gradients facilitate the delivery of antigens to the lymph nodes by guiding mature dendritic cells (Figure 2B) (Schumann *et al.*, 2010). Inside the lymph nodes, mature dendritic cells present the antigens to T-cells, initiating the adaptive immune response.

**Figure 2:**
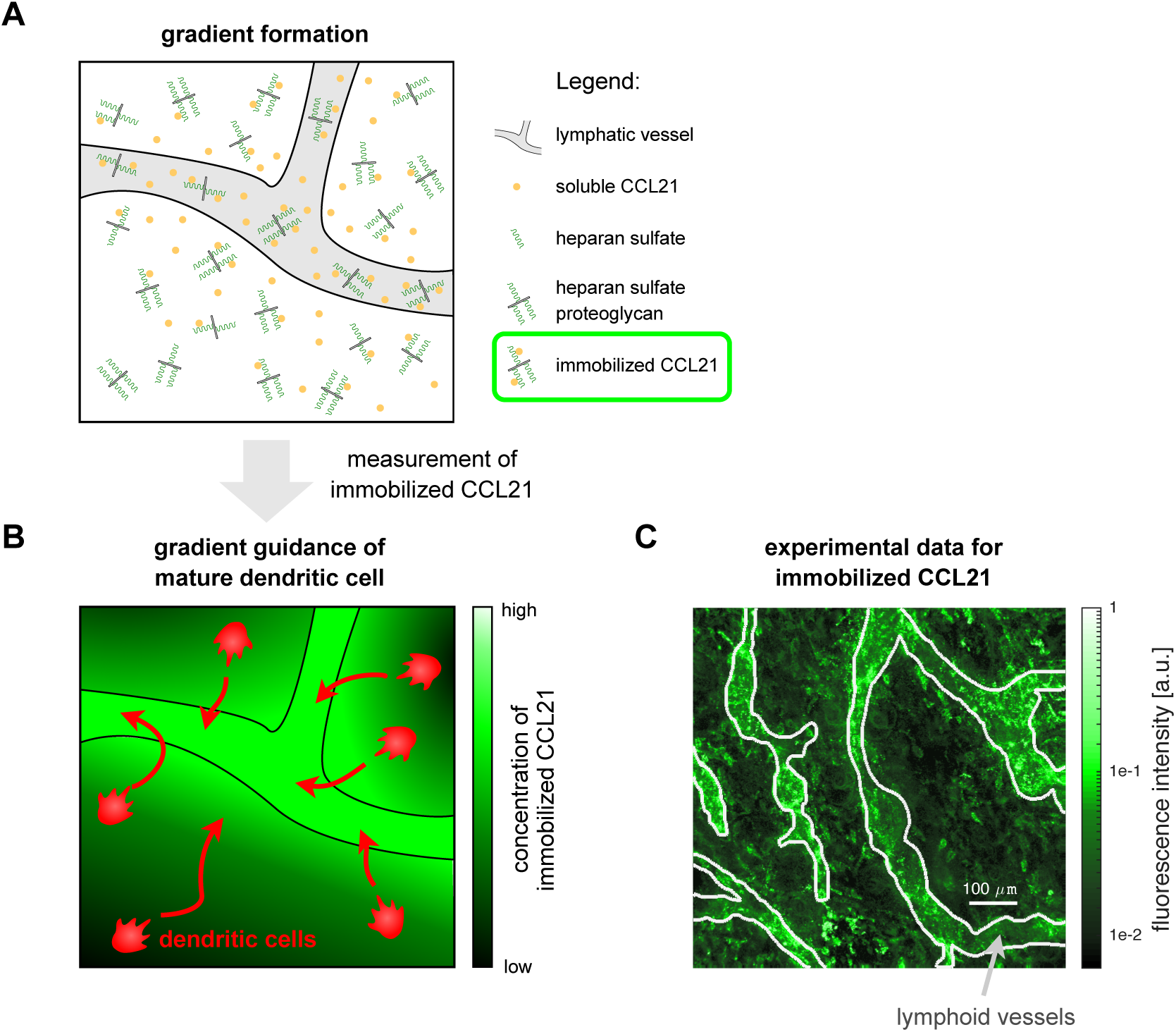
Model for CCL21 gradient formation and experimental data. (A) Schematic of model for CCL21 gradient formation, including diffusive and immobilised CCL21. (B) Illustration of mature dendritic cells guided by a gradient of immobilised CCL21. Experimental data for immobilised CCL21 and lymphatic vessels. CCL21 immunostaining is colour-coded and the outlines of the lymphatic vessels (light grey lines), which were determined using an additional staining, are indicated. These data were collected and provided by Weber *et al.* (2013) and we refer to the original publication for details on materials and methods.

The formation of the CCL21 gradients and their biological functions are relatively well understood and experimentally verified (Weber *et al.*, 2013). It is known that soluble chemokine CCL21 is secreted at the lymphatic vessels, and it is assumed that from there it diffuses into the dermal interstitium. Furthermore, it has been established that CCL21 binds to heparan sulfate proteoglycan, resulting in immobilised CCL21 which guides the migratory dendritic cells. However, the quantitative properties of the individual processes and the detailed mechanisms remain to be analysed. In addition, the available imaging data (Figure 2C) are corrupted by structured noise (see discussion below), rendering the analysis challenging and the process well-suited for the evaluation of the proposed approaches.

### Mathematical model and experimental data

We modelled the dynamics of the concentrations of soluble CCL21 (*u*_1_(*x, t*)), of heparan sulfate (*u*_2_(*x, t*)) and of heparan sulfate-CCL21 dimers (*u*_3_(*x, t*)) by a system of PDEs (Hock *et al.*, 2013a). The PDE model accounted for

- the secretion of soluble CCL21 with rate *α* from *L* lymphatic vessels which cover the spatial location *{x ∈* Ω*|q*_*l*_(*x*) = 1*}, l* = 1, *…*, *L*,
- the diffusion of soluble CCL21 with diffusion coefficient *D*,
- the degradation of soluble CCL21 with rate constant *γ*,
- the binding of soluble CCL21 to heparan sulfate with rate constant *k*_1_, and
- the unbinding of CCL21 from heparan sulfate with rate constant *k*_*-*1_,

and we assumed no flux conditions at the boundaries. Mathematically, we obtained the evolution equation

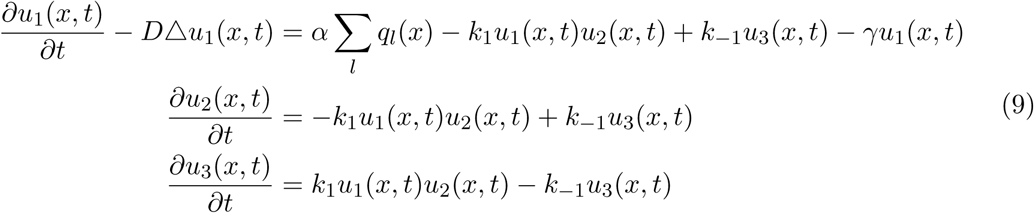

on *x ∈* Ω with initial conditions

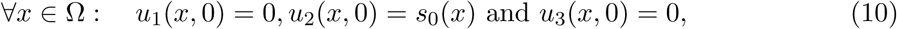

and boundary conditions

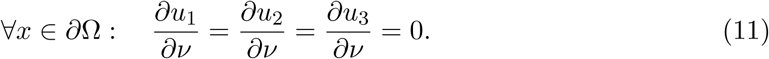

*∂*Ω denotes the boundary of Ω and *ν* denotes its normal vector. The heparan sulfate concentration was assumed to be homogenous, *s*_0_(*x*) = *S*_0_, unless mentioned otherwise.

Weber *et al.* (2013) succeeded to measure the *in vivo* gradients of immobilised CCL21, yielding the 2D images depicted in Figure 2C. As the experiments were performed in unperturbed tissue, the images provide the equilibrium distributions. Accordingly, the experimental readout is

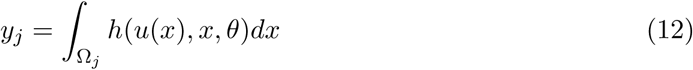

with pixel index *j*, observation function *h*(*u*(*x*), *x, θ*) = *s*(*u*_3_(*x*)+ *b*), and *u*(*x*) solving (9) with *∂u*_*i*_*/∂t* = 0, *i* = 1, *…*, 3. The measured pixel intensities are semi-quantitative, requiring the introduction of the scaling constant *s* and the background intensity *b*. These experimental parameters were estimated along with the kinetic parameters. In addition to immobilised CCL21, Weber *et al.* (2013) assessed the lymphatic vessel masks *q*_*l*_(*x*), *l* = 1, *…*, *L*, in the same mouse ear sheets, providing the basis for the simulation of realistic tissue structures. We assumed that the scaling *s* and the lymphatic vessel masks *q*_*l*_(*x*), *l* = 1, *…*, *L*, differ between images whereas background *b* and mechanistic parameters remain identical.

As the absolute concentration of CCL21 was unknown and merely the equilibrium distribution of immobilised CCL21 was measured, the parameters (*D, γ, k*_1_, *k*_*-*1_, *S*_0_, *s, b*)^*T*^ were structurally non-identifiable (see (Chis *et al.*, 2011) for definition). To circumvent this, we reformulated the model in terms of the parameters (*D/γ*, (*αk*_1_)*/*(*γk*_*-*1_), *s S*_0_, *b*)^*T*^. For details on the reparameterisation, we refer to Supporting Text S2.

The visual inspection of the imaging data revealed a high level of immobilised CCL21 associated with the lymphatic vessel, which was in agreement with the model. However, there were also high intensity spots outside the lymphatic vessels (Figure 2C), which were not explained by the aforementioned processes. As in fixed tissues, the immunostaining performed by Weber *et al.* (2013) labels intracellular and extracellular CCL21 (Kilarski *et al.*, 2013), these spots are most likely caused by previously reported CCL21 expressing cells (Tal *et al.*, 2011). As intracellular CCL21 does not contribute to the extracellular distribution of CCL21 described by model (9), the spots should were considered as structured noise and disregarded in the parameter estimation. This rendered the modelling problem appropriate for the evaluation of the integrated modelling approach.

### Integrative modelling approach outperforms conventional methods on artificial experimental data

In this section, we assess the properties of different image-based modelling using artificial imaging data. This allows us to evaluate the accuracy with which the true parameter vector is recovered using (i) whole imaging data vs. a summary statistic and (ii) direct approach vs. filtering approach vs. integrated modelling approach.

### Generation of artificial data

We derived artificial imaging data closely resembling the experimentally observed images to ensure a realistic test scenario. The spatial structure of the images was conserved by using the measured lymphatic vessel masks and selecting model parameters which roughly reproduced the experimentally observed CCL21 distributions. The employed parameters are provided in Supporting Text S2, Table 1. The structured measurement noise was captured by extracting relevant features of the high intensity spots. Firstly, the high-intensity spots were detected using MSER filtering (Matas *et al.*, 2004) using an implementation by Nistér & Stewenius (2008). Secondly, the identified spots were analysed to obtain the distributions of spot shape parameters and sizes (Figure 3A). Given these distributions, the artificial data were obtained by simulating the model for the selected parameters and adding a varying number of spots with properties sampled from the measured distribution. For simplicity, the spots were assumed to be ellipsoidal. A representative artificial image is depicted in Figure 3C. While the artificial data do not capture the full complexity of experimental data, they facilitate the evaluation of the approaches.

**Figure 3:**
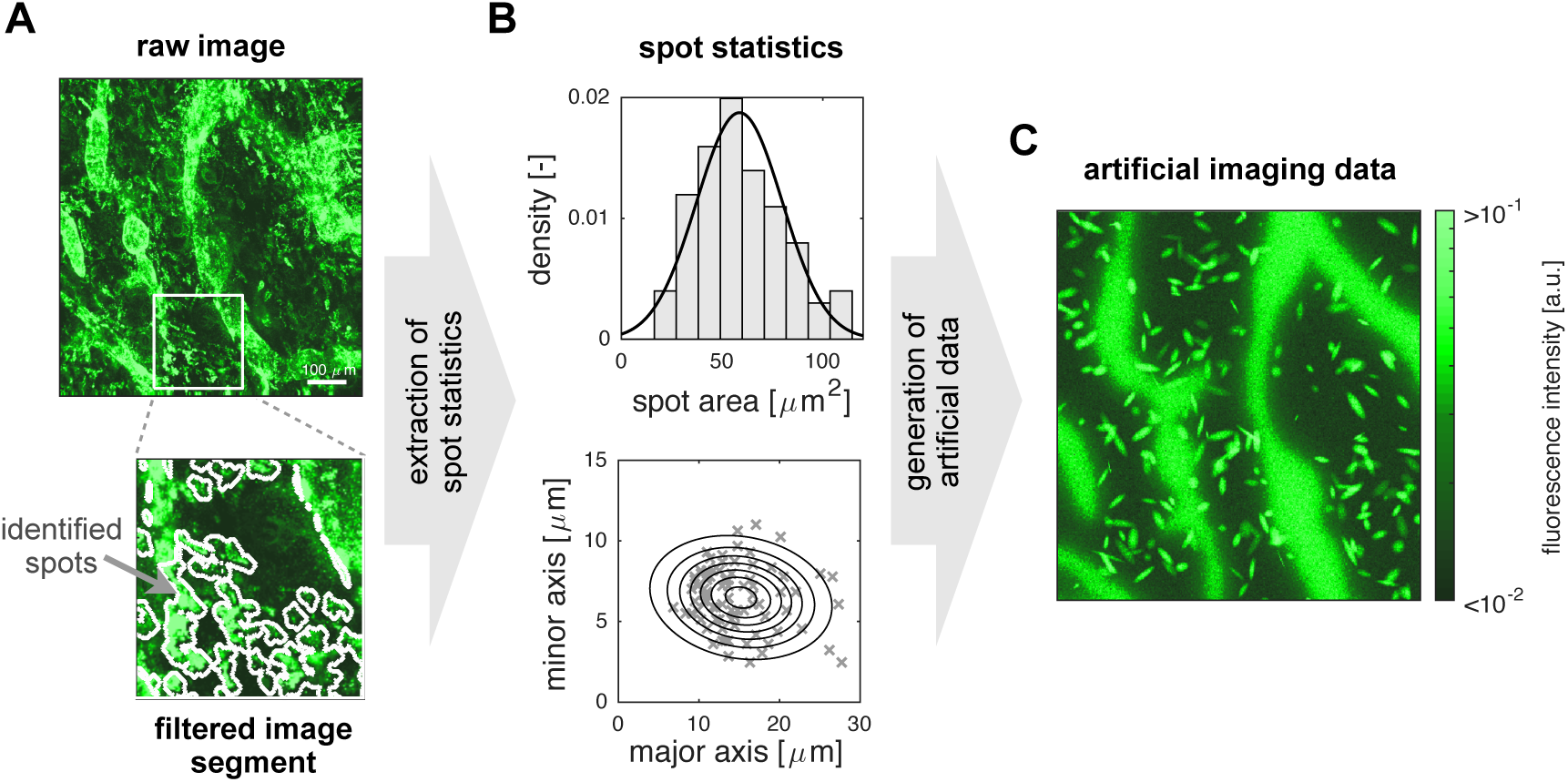
Pipeline for the generation of artificial imaging data. (A) Raw and filtered image. Outlines of spots identified using maximal stable extremal region (MSER) filtering are indicated. (B) Distribution of size and shape parameters of spots. Histogram and points in scatter plot indicate the information extracted using filtering. The lines indicate the densities used for the generation of the artificial data. (C) Artificial data obtained by simulation of the model (9) and subsequent addition of spots and measurement noise.

In addition to the artificial imaging data, we generated the artificial summary statistic. We considered the distance-dependent average intensity as evaluated by Weber *et al.* (2013) for the process of CCL21 gradient formation. To calculate this summary statistic for the artificial data, the minimal distance to the next lymphatic vessel is computed for each pixel. Subsequently, the intensity values of all pixels with the same distance are averaged.

### Detailed spatial information improves estimation accuracy

Given the artificial datasets, we first asked how much information the raw imaging data contain in comparison to the summary statistic computed from them. To address this, we considered three setups:

i. Fitting of the distance-dependent average CCL21 intensity using the 1D model.
ii. Fitting of the distance-dependent average CCL21 using the 2D model accounting for the measured vessel topology.
iii. Fitting of the CCL21 imaging data using the 2D model accounting for the measured vessel topology.

Setup (i) employs a simplified version of model (9) with *x ∈* [0, *L*] denoting the distance from the lymphatic vessel (see Supporting Text S2). The secretion at the lymphatic vessel (at *x* = 0) is modelled via the boundary condition *∂u*_1_*/∂x*|_*x*=0_ = *α*. The diffusion and reaction dynamics stay the same. Setup (ii) and (iii) employed model (9) with the measured lymphatic vessel mask. To study the relevance of detailed spatial information, we considered artificial data without outliers and structured noise but with independent and identically distributed measurement noise, i.e., multiplicative log-normally distributed measurement noise. The signal-to-noise ratio, which is the mean signal intensity divided by the standard deviation of the noise, was approximately 6.

As the considered artificial data do neither contain outliers nor structured noise, we employed the direct approach for statistical modelling. Parameter optimisation and uncertainty analysis for setups (i)-(iii) was performed using multi-start local optimisation and profile likelihood methods, respectively. All parameters were constrained to a regime spanning at least 4-orders of magnitude (see Supporting Text S2, Table 1).

The analysis of a representative artificial dataset revealed that for setups (i) and (ii) a good agreement with the summary statistic was achieved (Figure 4A, B), while for setup (iii) a good agreement with the imaging data was obtained (Figure 4C). The estimated parameters in setup (i) were however far from the true parameters. This was among others due to practical non-identifiabilities of the parameters (*αk*_1_)*/*(*γk*_*-*1_) and *S*_0_ (Figure 4D). While the individual parameters are non-identifiable, their product is practically identifiable. The same phenomenon was observed for setup (ii) (Figure 4D, E), implying that modelling the underlying spatial structure did not improve the information extraction substantially. In contrast, for setup (iii), all parameter estimates were close to the true parameter and practically identifiable (Figure 4D). Thus, not the summary statistic but the whole imaging data should be used as they allow for more accurate parameter estimation.

**Figure 4:**
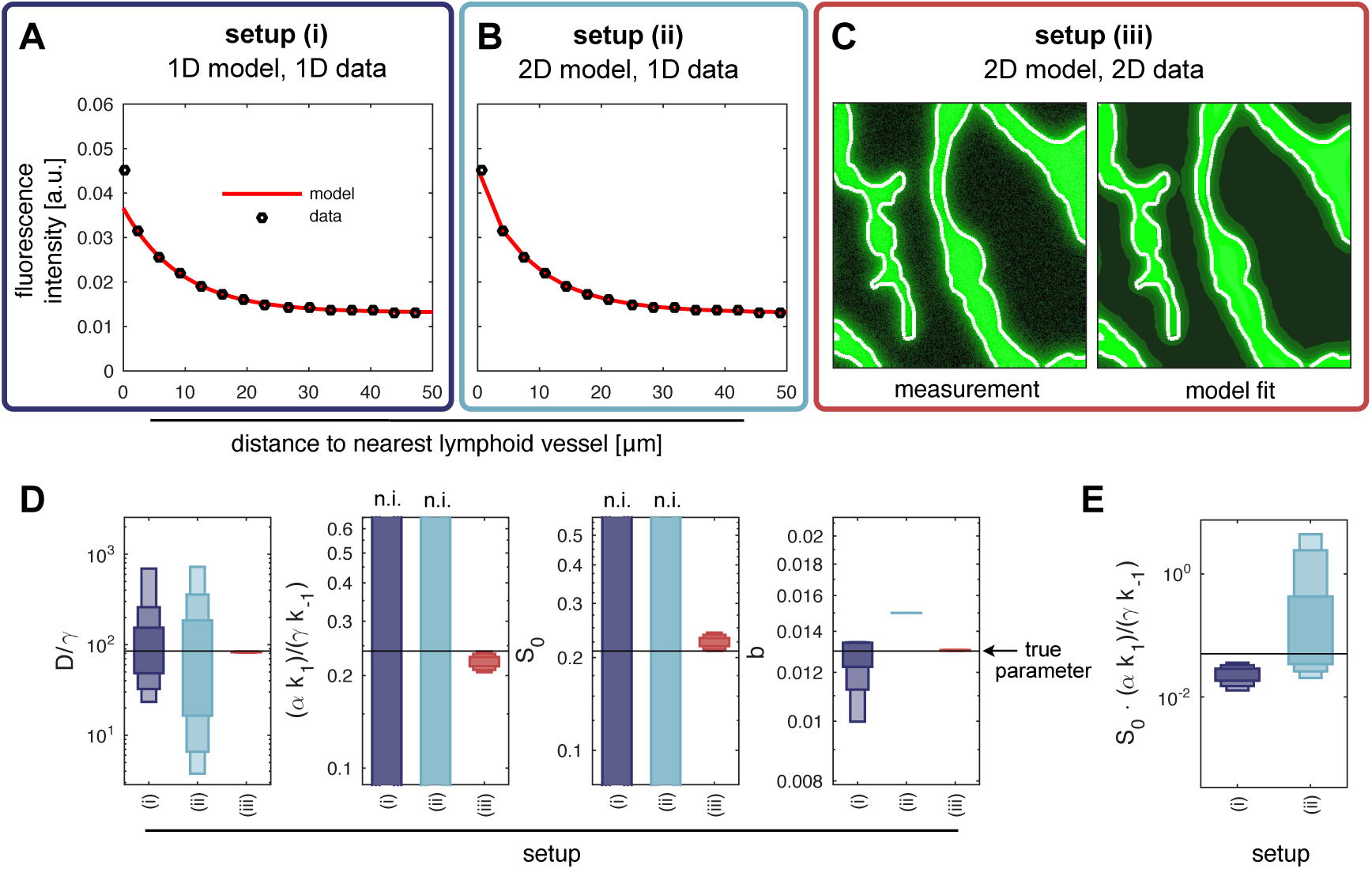
Parameter estimation results for the 1D and 2D models using extracted features and whole imaging data. (A) Fitting result for the 1D model using the distance-dependent average intensity of immobilised CCL21 (1D data). (B) Fitting result for the 2D model using the distance-dependent average intensity of immobilised CCL21 (1D data). (C) Fitting result for the 2D model using the measured intensity distribution of immobilised CCL21 (2D data). (D,E) Profile likelihood derived confidence intervals for parameter estimates for setups (i)-(iii). The horizontal line marks the true parameter and the vertical bars represent the confidence intervals corresponding to different confidence levels (75%, 90% and 99%), with non-identifiable parameters indicated by ’n.i.’. (E) Profile likelihood derived confidence intervals for the product of the parameters which are non-identifiable for setup (i) and (ii).

### Integrated modelling approach yields more accurate and robust results than conventional methods

As whole imaging data are strongly influenced by outliers and structured noise, we compared the accuracy of parameter estimates obtained using the direct approach, the filtering approach and the integrated statistical approach. We considered artificial imaging data with 0 to 620 bright spots and evaluated 30 datasets to obtain robust statistics.

Our analysis revealed that for artificial datasets with a large number of spots, fits obtained using the direct approach overestimated the concentration of immobilised CCL21 outside the spots while filtering and the integrated approach provided consistent results (Figure 5A). Apparently, the direct approach could not explain the structured noise and the bimodal distribution of the residual 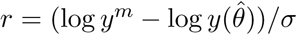 (Figure 5B). The filtering of points and the integrated modelling resulted in a more consistent statistical description.

**Figure 5:**
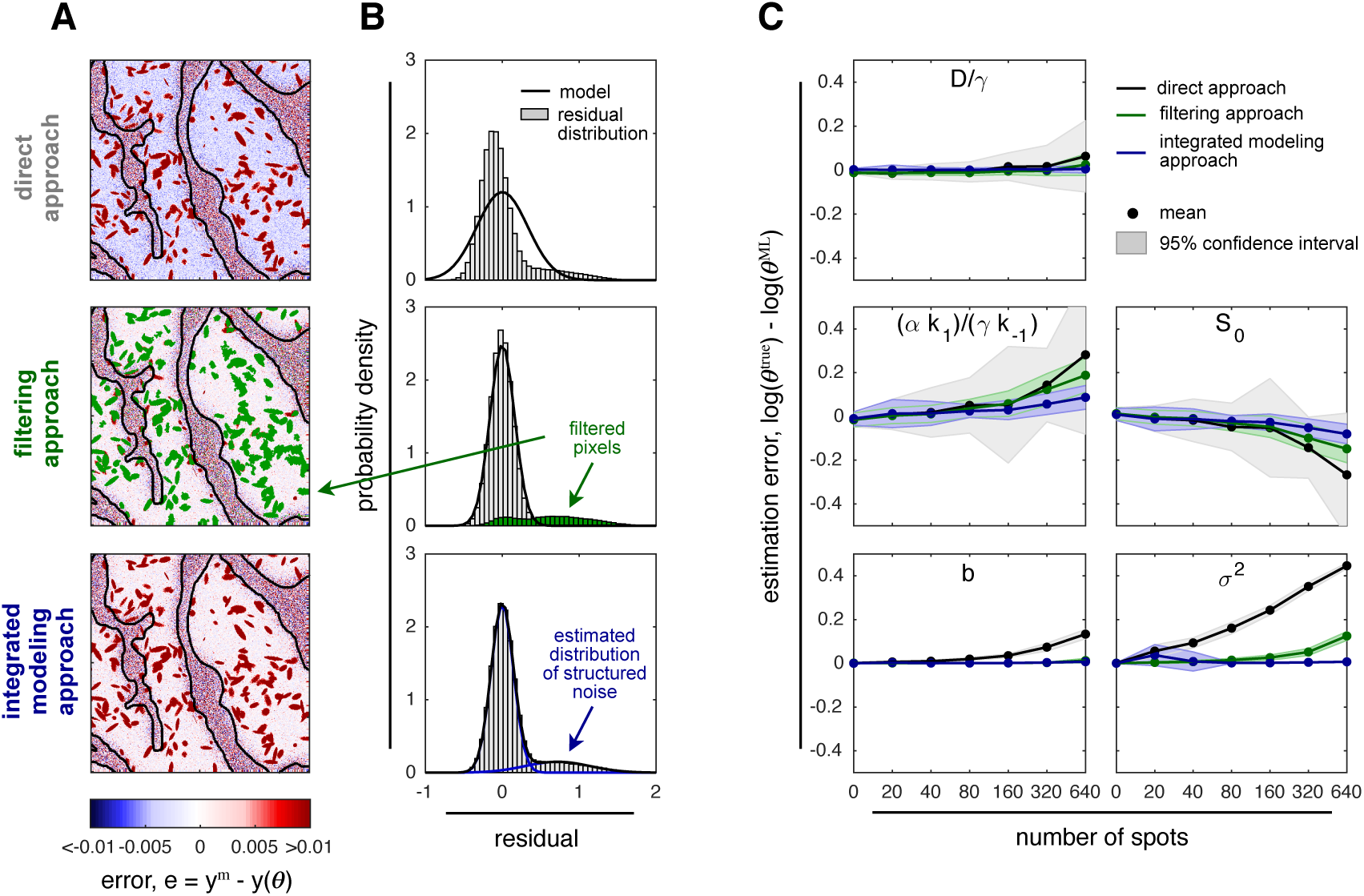
Parameter estimation results for images with structured noise. (A) Spatial structure of residuals and (B) observed (histogram) and modelled (line) residual distribution for representative dataset with 320 spots. The results for the simulation of the 2D model with parameter estimates obtained using (top) the direct approach, (middle) the filtering approach, and (bottom) the integrated modelling approach are depicted. Pixels disregarded by the filtering approach are coded green (histogram). The estimated outlier distribution in the integrated modelling approach is depicted in blue (line). (C) Mean parameter estimation error (black line) 95% percentile interval (area) of the direct approach, the filtering approach and the integrated modelling approach for different spot numbers computed using 30 artificial datasets.

The analysis of the estimation indicated that for low numbers of spots, the filtering approach and the integrated modelling approach yielded almost the same results as without spots while the direct approach already possessed a bias and a large variance (Figure 5C). For medium and high numbers of spots, the integrated modelling yielded the smallest estimation error. This was the case although a) the filter approach employed the same MSER filter settings used to obtain the spot statistics – this parameter setting appeared to be ideal – and b) the integrated modelling approach did not account for the spatial structure of measurement noise. Indeed, the integrated modelling approach yielded almost unbiased results. Thus, integrated noise modelling provided robust parameter estimates from imaging data corrupted with the considered type of structured noise.

In conclusion, our analysis of artificial data suggests that mechanistic modelling of spatial processes should be based on detailed imaging data rather than some spatial summary statistic. Additionally, filtering but even more so the proposed integrated modelling approach can provide robust estimates in the presence of structured noise and outliers.

### Integrative modelling approach predicts lymphatic vessel dependent heparan sulfate concentration

Given the positive results for artificial data, we used the integrated modelling approach on whole imaging data to analyse experimental data for CCL21 gradient formation. Among others, we asked whether the current assumption of uniform heparan sulfate concentration is appropriate or alternative mechanisms need to be considered.

### Model-based image analysis reveals limitation of a literature-based model

We employed model (9) with uniform heparan sulfate concentration, *s*_0_(*x*) = *S*_0_, to describe the imaging data collected by Weber *et al.* (2013). This model was suggested by the literature and experts in the field. As the no-flux boundary conditions (11) are presumable not precisely met in the biological system, we disregarded pixels which are within 40 *µm* of the boundary for the calculation of the objective function (6). This depth was chosen based on preliminary estimates for the diffusion length from the summary statistic (Supporting Text S2, Figure 1) and retrospectively validated given the fitting results.

The fitting results for the model with uniform heparan sulfate concentration for a representative image with multiple lymphatic vessels are depicted in Figure 6. For this image, the comparison of experimental data and the fitting results (Figure 6B,C) revealed that – as expected – bright spots outside lymphatic vessels are not captured. However, there were also larger regions in the image with substantial disagreement. In particular in lymphatic vessels 1 and 2, the concentration of immobilised CCL21 was overestimated. Accordingly, the residuals were not uncorrelated but show a clear spatial structure (Figure 6D), resulting in a pronounced tail in the residual distribution (Figure 6E). This indicated that the model with uniform CCL21 concentration might be too simple.

**Figure 6:**
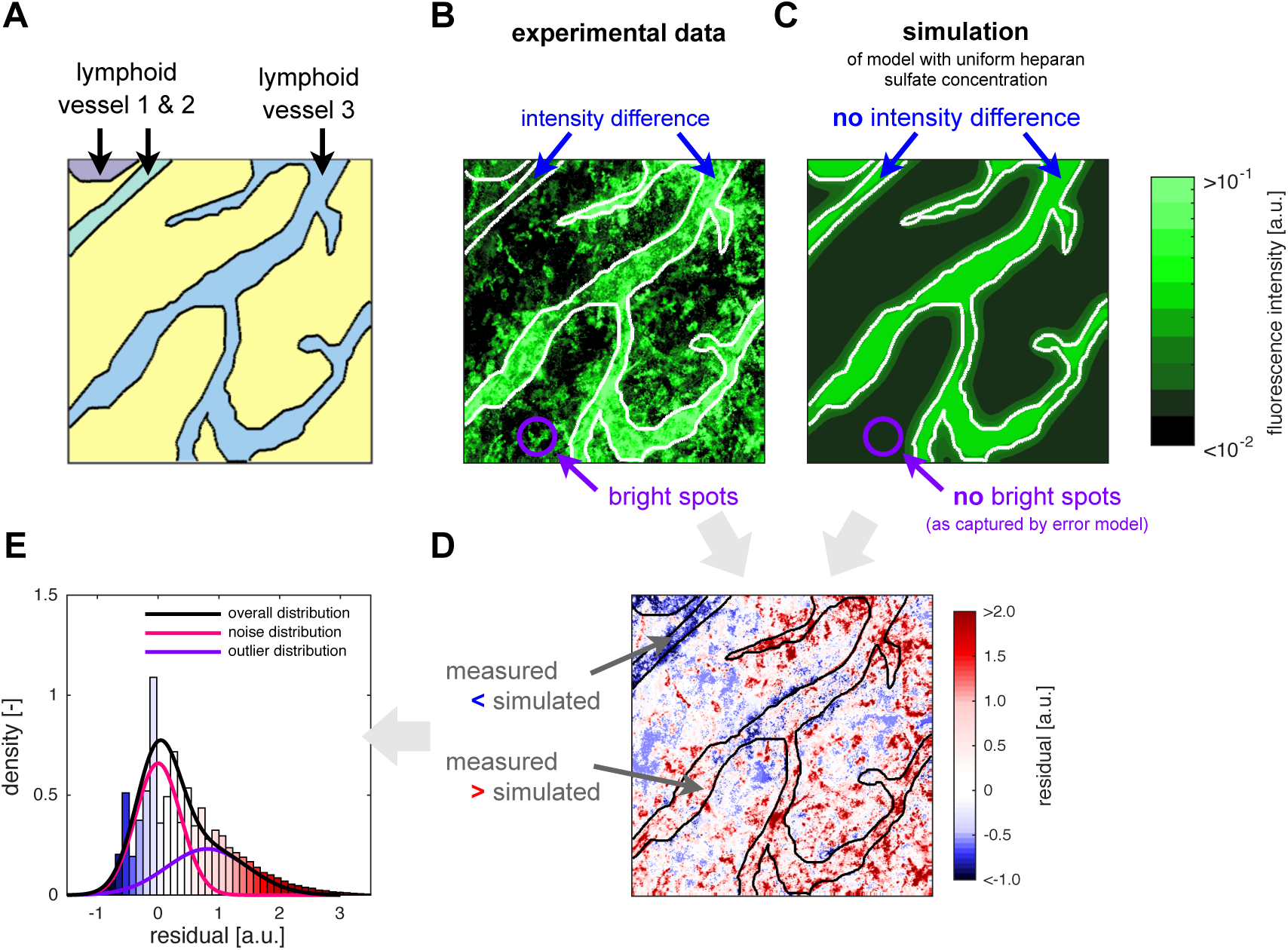
Analysis of model with uniform heparan sulfate concentration. (A) Spatial location of lymphatic vessels. The individual vessels are colour-coded. (B) Experimental data for immobilised CCL21. The difference in the concentration of immobilised CCL21 between lymphatic vessels is indicated along with the presence of spots. (C) Simulation results for immobilised CCL21. The maximum likelihood estimate for the model with uniform heparan sulfate concentration is depicted. Differences between lymphatic vessels are not captured. Spots are filtered using the integrated statistical model. (D) Residuals of experimental data and simulation results. The low measured intensity in lymphatic vessels 1 and 2 are not captured by the model. (E) Observed residual distribution and distributions of unstructured and structured noise indicated by the integrative modelling approach.

### Mathematical modelling supports hypothesis of vessel dependent heparan sulfate concentration

As the detailed analysis of the whole imaging data revealed limitations of the literature-based model, we evaluated possible model refinements. In addition to the hypothesis underlying the model presented in the previous section:

1. Uniform heparan sulfate concentration, *s*_0_(*x*) = *S*_0_. we considered two alternative hypotheses:
2. Different heparan sulfate concentrations in lymphatic vessels and the tissue, 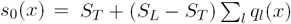.
3. Different hepa n sulfate concentrations in individual lymphatic vessels and the tissue, 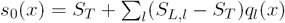.

These hypotheses yield models 1-3 which are illustrated in Figure 7A. We employed the integrated modelling approach to train the models 1-3 on all 9 images recorded by Weber *et al.* (2013), namely image 1 to image 4 and image 12 to image 16. The optimisation converged robustly (Figure 7B) and the fitting results for different images are provided in Supporting Text S2, Table 2-2.

**Figure 7:**
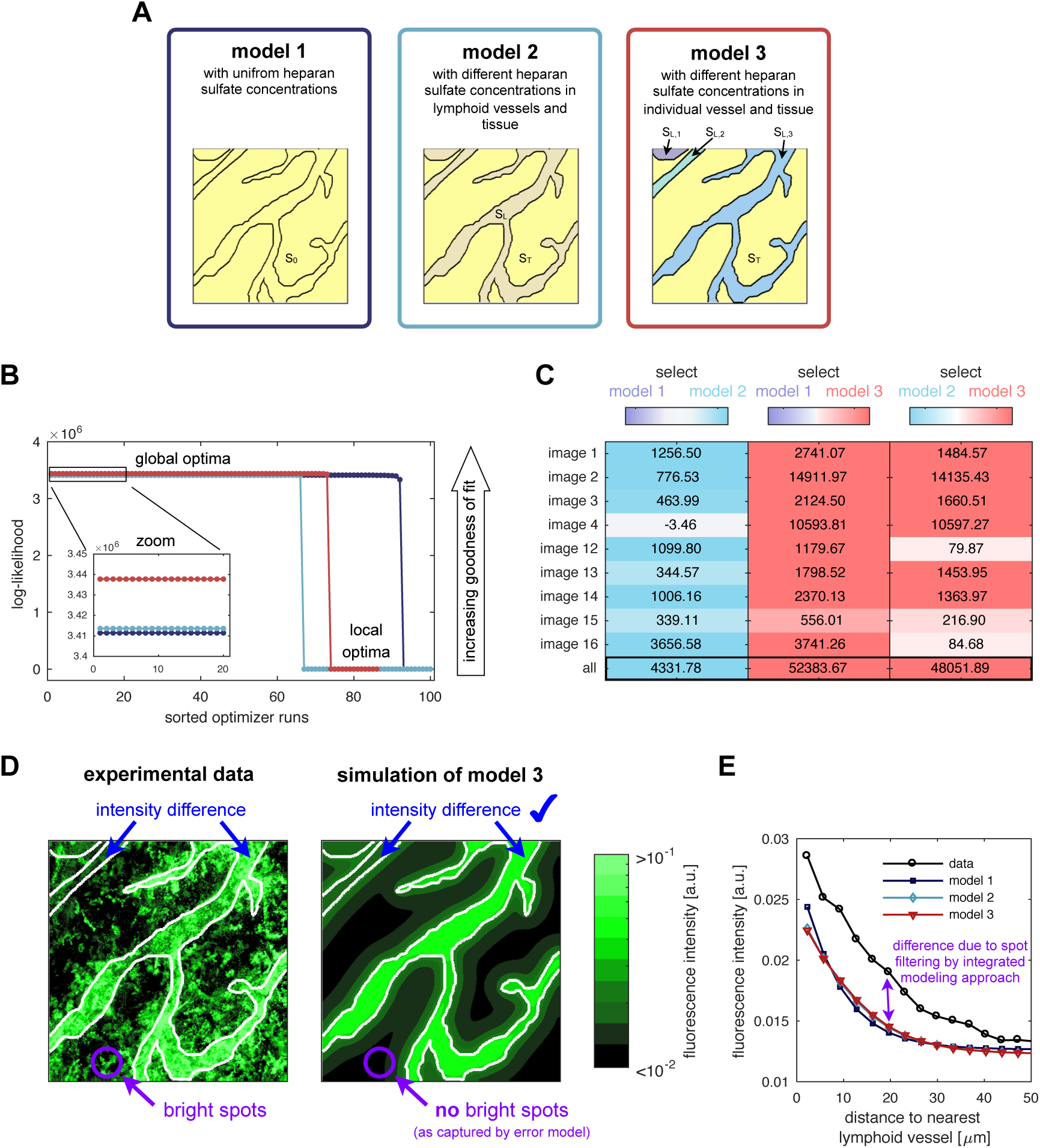
Comparison of three different hypotheses for the CCL21 distributions. (A) Schematics of models 1-3. (B) Multi-start optimisation results for the model alternatives, indicating a unique global optimum for each models. (C) Results of model comparison using BIC for each image and the overall fit. The differences of BIC values for different models are colour-coded, with each model being assigned one color (model 1: purple; model 2: cyan; and model 3: red). (D) Experimental data for the spatial distribution of immobilised CCL21 and simulation results for model 3. (E) Experimental data for the distance-dependent intensity of immobilised CCL21 and simulation results for models 1-3.

For the individual images as well as for the overall dataset, model 3 was substantially better than models 1 and 2 (Figure 7C). Model 3 provided a good agreement with the imaging data (Figure 7D). Furthermore, the prediction of differences in the heparan sulfate concentrations between individual lymphatic vessels is consistent with experimental data indicating different levels of extracellular CCL21 in collecting and initial lymphatics (Kilarski *et al.*, 2013). Thus, our model-based analysis provided a mechanistic hypothesis which we were able to partially validated using published results.

To conclude, in this section we verified the applicability of the integrated modelling approach to experimental imaging data including structured noise. We employed the statistical approach for model-based data analysis and hypothesis testing, thereby providing new insights into the CCL21 gradient formation and dendritic cell guidance. Notably, all models achieved an equally good fit for the summary statistic (Figure 7E), implying that the information content of the summary statistic is too limited for model selection, confirming that models should be based on the whole imaging data.

## Discussion

Imaging data are widely used to assess biological processes. In many studies, the richness of imaging data is, however, disregarded and they are merely used to derive and evaluate simple summary statistics. We illustrated that this can results in a considerable loss of information. While summary statistics are often sufficient to draw qualitative conclusions, quantitative mechanistic models should be trained using whole imaging data – wherever possible – to exploit their richness.

The model-based analysis of imaging data facilitates the unraveling of novel mechanisms and the comparison of competing hypotheses (Uzkudun *et al.*, 2015; Iber *et al.*, 2015; Jagiella *et al.*, 2017), this is often too demanding and error-prone if structured noise and outliers are present. To address this problem, we introduced an integrated statistical approach for the statistical and mechanistic modelling of imaging data. For the considered problems, the integrated modelling approach yields similar or better results than conventional sequential methods. Even without knowledge of the precise structure of the noise, the method was able to reduce estimation bias and variance, providing more reliable parameter estimates.

To evaluate the properties of the integrated modelling approach, we studied CCL21 gradient formation. We established the first quantitative mathematical model of CCL21 gradients measured in tissue. Using experimental data, we quantified the estimation error of different models and performed model selection. Among other, we found indications that the heparan sulfate concentration is vessel dependent. This can influence the gradient formation and cell guidance and might be relevant in some disease conditions (Dudal *et al.*, 2015). Furthermore, it demonstrates that integrated modelling approaches might reveal novel information from available data and can help to unravel causal factors.

In this study, we considered a simple statistical model for outliers and structured noise. A further improvement of the integrated modelling approach could be achieved by considering more tailored statistical models. The correlation of noise in neighbouring pixels could be considered and even sophisticated segmentation methods, e.g., graph-based segmentation approaches (Felzenszwalb & Huttenlocher, 2004), might be incorporated in a likelihood framework. Extension in this direction and towards image regression could improve robustness and applicability further.

In conclusion, mechanistic understanding and rigorous hypothesis testing in biology requires the formulation of mathematical and computational models. For cellular processes, this led to the development of modelling and estimation toolboxes, e.g., Data2Dynamics (Raue *et al.*, 2015), which support the simultaneous inference of kinetic parameters and measurement noise. We illustrated that such a simultaneous inference is also feasible for the case of spatial models, which are usually more challenging. We illustrated parameter optimisation, uncertainty analysis and model selection for PDE model of noise-corrupted imaging data. We expect that the proposed concept and algorithms are well suited for a broad range of applications, including scenarios with time-resolved measurements and time-dependent domains (Uzkudun *et al.*, 2015). This is also facilitated by the availability of the MATLAB code, simplifying reuse and extensions of the methods. Accordingly, this study will contribute to the mechanistic description of spatial processes.

## Supporting Information

### Supporting Code S1

**MATLAB code.** This file includes the code used to generate and analyse the artificial data and to study the experimental data. An implementation of the models, the parameter optimisation and the uncertainty analysis is provided.

### Supporting Text S2

**Supplemental notes.** This document includes a description of the considered models and some additional analysis of the experimental data.

