## Supplementary material for "Mechanistic description of spatial processes using integrative modelling of noise-corrupted imaging data": computeMixtureProbability.m

```
function varargout = computeMixtureProbability(varargin)
% Robust calculation of a mixture distribution likelihood.
%
% USAGE:
%   [logp,dlogpdxi] = computeMixtureProbability(w,q_i,H_i) \n
%   [logp]          = computeMixtureProbability(w,q_i) 
%
% Parameters:
% varargin:
% w: (1 x n_s) vector with weights \f$w_s\f$
% q_i:  (n x n_s) matrix with log(p_i) for every column
% H_i:  (n x n_xi x n_s) s.th. d(w_i*p_i)/dxi = p_i*H_i
%
% Return values:
% logp: n x 1 scalar of loglikelihood
% dlogpdxi: n x n_xi vector of gradient

%% Input assignment and initialization
w = varargin{1};
q_i = varargin{2};
n_s = length(w); % number of subpopulations
n = size(q_i,1); % number of data points

if abs((sum(w)-1)>1e-13) || ~all(w>=0)
    error('Weights need to be positive and sum up to 1!')
end

if nargout == 2
    H_i =  varargin{3};
    n_xi = size(H_i,2); % number of parameters
    Hmax = zeros(n,n_xi);    
end

%% Handling of subpopulations of size zero
ind_w = find(w > 0);
if length(ind_w) < n_s
    w = w(ind_w);
    q_i = q_i(:,ind_w);
    n_s = length(w); % number of non-zero subpopulations
    if nargout == 2
        H_i =  H_i(:,:,ind_w);
    end
end

%% Calculate q = log(p) and gradient dlog(p)/dxi

[qmax,imax] = max(q_i,[],2);

if n_s == 2 % two subpopulations
    imax1 = (imax==1);
    imax2 = (imax==2);
    wmax = w(1).*imax1+w(2).*imax2;
    wmin = w(1).*(~imax1)+w(2).*(~imax2);
    qmin = q_i(:,1).*(~imax1) + q_i(:,2).*(~imax2);
    logp = log(1+(wmin./wmax).*exp(qmin-qmax))+log(wmax)+qmax;
    
    if nargout == 2
        Hmax = bsxfun(@times,H_i(:,:,1),imax1)+ bsxfun(@times,H_i(:,:,2),imax2);
        Hmin = bsxfun(@times,H_i(:,:,1),~imax1)+ bsxfun(@times,H_i(:,:,2),~imax2);
        dlogpdxi = bsxfun(@rdivide,Hmax,(wmax+wmin.*(exp(qmin-qmax))))+...
            bsxfun(@times,Hmin,exp(qmin-qmax)./(wmax+wmin.*(exp(qmin-qmax))));
    end
    
else
    wmax = zeros(n,1);
    for k = 1:n_s-1
        wmin{k} = zeros(n,1);
        qmin{k} = zeros(n,1);
        imin{k} = mod(imax-1+k,n_s)+1;
        if nargout == 2
            Hmin{k} = zeros(n,n_xi);
        end
    end
    
    for s = 1:n_s
        wmax = wmax + w(s).*(imax==s);
        if nargout == 2
            Hmax = Hmax + bsxfun(@times,(imax==s),H_i(:,:,s));
        end
        for k = 1:n_s-1
            ind = (imin{k}==s);
            wmin{k} = wmin{k} + w(s).*ind;
            qmin{k} = qmin{k} + q_i(:,s).*ind;
            if nargout == 2
                Hmin{k} = Hmin{k} + bsxfun(@times,ind,H_i(:,:,s));
            end
        end
    end
    
    temp = 0;
    if nargout == 2
        numer = Hmax;
        denom = wmax;
    end
    for k = 1:n_s-1
        temp = temp + (wmin{k}./wmax).*exp(qmin{k}-qmax);
        if nargout == 2
            numer = numer + bsxfun(@times,exp(qmin{k}-qmax),Hmin{k});
            denom = denom + wmin{k}.*exp(qmin{k}-qmax);
        end
    end
    logp = log(1+temp)+log(wmax)+qmax;
    if nargout == 2
        dlogpdxi = bsxfun(@rdivide,numer,denom);
    end
end

%% Assign output
varargout{1} = logp;
if ~isreal(logp)
    disp('Warning: likelihood is not real!')
end
if nargout == 2
    varargout{2} = dlogpdxi;
end
```
