## Supplementary material for "Mechanistic description of spatial processes using integrative modelling of noise-corrupted imaging data": performFiltering.m

```
function [varargout] = performFiltering(img,mask)
% performFiltering calculates region of interest by performing a maximum 
% stable extremal region ('mser') or a median ('medfilt') filtering.

% Assignment of filter properties
options.type = 'mser';
options.delta = 1; %1-255
options.minSize = 50; % spot area
options.maxSize = 600;
options.dilate = 'true';
options.dilate_size = 1;
options.LV = 'false';

options.plot ='false';
options.cmin = 0.01;
options.cmax = 0.1;

%
Y = adapthisteq(img);
Y = im2uint8(Y);
mask = logical(mask);

% Initialize binary mask
H = zeros(size(Y));
switch options.type
    case 'mser'
        msers = linearMser(Y,options.delta,options.minSize,options.maxSize,1,1);

        % Fill with msers
        for m = 1:numel(msers)
            H(msers{m}) = 1;
        end
    case 'medfilt2'
        B = medfilt2(Y, [options.minSize options.minSize]);
        C = medfilt2(Y, [options.maxSize options.maxSize]);
        mask2 = imdilate(mask,strel('disk',options.safe));
        H(B-C > options.delta & mask2 < 1) = 1;
end

% Optional dilation
if strcmp(options.dilate,'true')
    H = imdilate(H,strel('disk',options.dilate_size));
end

if strcmp(options.LV,'false')
    H(mask) = 0; % ignore points in LV
end
    
H = 1-H;

% Convert mask to logical
H = logical(H);
rm = sum(sum(1-H))/(size(H,1)*size(H,2))*100;

if strcmp(options.plot,'true')
    figure;
    Y_f = zeros(size(options.img));
    Y_f(H) =  options.im(H);

    subplot(1,3,1)
    imagesc(options.im,[options.cmin,options.cmax]); hold on;
    colormap('gray'); %colorbar('Location','East'); axis off
    contour(mask,[0 0],'Color',[0.8 0 0],'linewidth',2)
    title('raw data')
    subplot(1,3,2)
    imagesc(options.im,[options.cmin,options.cmax]); axis off; hold on
    contour(mask,[0 0],'Color',[0.8 0 0],'linewidth',2)
    contour(H,[0,0],'Color',[0 0.8 1]); 
    title('filtering mask')
    subplot(1,3,3)
    imagesc(Y_f,[options.cmin,options.cmax]); axis off; hold on;
    contour(mask,[0 0],'Color',[0.8 0 0],'linewidth',2)
    title(['final image: ',num2str(round(rm)),'% removed'])
end

switch nargout
    case 1
        varargout{1} = H;
    case 2
        varargout{1} = H;
        varargout{2} = msers;
end

end
```
