## Supplementary material for "Mechanistic description of spatial processes using integrative modelling of noise-corrupted imaging data": getDistanceDependentMean.m

```
% getDistanceDependentMean() computes the distance dependent mean intensity
% for an image. 
%
% USAGE:
% * [x,m,s] = getDistanceDependentMean(distance_map,intensity,distance,index_set)
%
% Parameters:
%   distance_map: distance for each pixel.
%   intensity: intensity for each pixel.
%   distance: distance for which the mean intensity is computed.
%   index_set: struct of index sets of pixel with a certain distance. The
%      entry index_set{k} corresponds to the distance(k).
%
% Return values:
%   x: vector of mean distances
%   m: vector of mean intensities
%   s: vector of standard deviation of mean intensity
%
% History:
% * 2017/11/06 Jan Hasenauer

function [x,m,s] = getDistanceDependentMean(distance_map,intensity,distance,index_set)

% Loop: Distances
for k = 1:length(distance)-1
    % Raw data
    x(k,1) = nanmean(distance_map(index_set{k})); % distance
    m(k,1) = nanmean(intensity(index_set{k}));    % mean
    s(k,1) = nanstd(intensity(index_set{k}));     % standard deviation
end
```
