## Supplementary material for "Mechanistic description of spatial processes using integrative modelling of noise-corrupted imaging data": performFilteringAndFeatureExtraction.m

```
% performFilteringAndFeatureExtraction() filters the image and computes 
% features. Among others, it computes the index sets of pixes at a certain
% distance from the lymphatic vessel.
%
% USAGE:
% * [ccl21_data,filtered_ccl21_data,dist_dep_mean,filtered_dist_dep_mean,I] = ...
%       performFilteringAndFeatureExtraction(ccl21_data,lv_mask,dist_from_lv,ind1,ind2,d)
%
% Parameters:
%   ccl21_data: CCL21 data.
%   lv_mask: mask for lymphatic vessel. (Pixels with intensity equal
%       to one are part of a lymphatic vessel.)
%   dist_from_lv: Distance of individual pixel from closest lymphatic vessel.
%   ind1: index set of pixels in x-direction which is considered.
%   ind2: index set of pixels in y-direction which is considered.
%   distance: distance for which the mean intensity is computed.
%
% Return values:
%   ccl21_data: CCL21 data on considered domain.
%   filtered_ccl21_data: filtered CCL21 data on considered domain.
%   dist_dep_mean: distance-dependent mean of CCL21 data.
%   filtered_dist_dep_mean: distance-dependent mean of filtered CCL21 data.
%   dist_from_lv: distance of different pixels from closest lymphoid vessel
%      on the considered domain.
%   I: struct of index sets of pixel with a certain distance. The entry
%      index_set{k} corresponds to the distance(k).
%
% History:
% * 2017/11/06 Jan Hasenauer

function [ccl21_data,filtered_ccl21_data,dist_dep_mean,filtered_dist_dep_mean,lv_mask,dist_from_lv,I] = performFilteringAndFeatureExtraction(ccl21_data,lv_mask,dist_from_lv,ind1,ind2,d)

    % Reduction to region of interest
    ccl21_data = ccl21_data(ind1,ind2)./255;
    dist_from_lv = dist_from_lv(ind1,ind2);
    lv_mask = lv_mask(ind1,ind2);
    
    % Index set I{k} of pixels with distance in the interval [d(k),d(k+1))
    % from the closest lymphatic vessel
    I = [];
    I{1} = find(dist_from_lv(:) == 0);
    for k = 1:length(d)-1
        I{k+1} = find(  dist_from_lv(:) >= d(k) ...
                      & dist_from_lv(:) <  d(k+1));
        if isempty(I{k+1})
            continue
        end
    end

    % Computation of filter and filtered data
    [index_filtered_pixels,~] = performFiltering(ccl21_data,lv_mask);
    
    filtered_ccl21_data = ccl21_data;
    filtered_ccl21_data(~index_filtered_pixels) = nan;
    
    % Calculation of distance-dependent mean and standard deviation
    [dist_dep_mean.dist,dist_dep_mean.mean,dist_dep_mean.std] = ...
        getDistanceDependentMean(dist_from_lv,ccl21_data,d,I);
    [filtered_dist_dep_mean.dist,filtered_dist_dep_mean.mean,filtered_dist_dep_mean.std] = ...
        getDistanceDependentMean(dist_from_lv,filtered_ccl21_data,d,I);
end
```
