## Supplementary material for "Mechanistic description of spatial processes using integrative modelling of noise-corrupted imaging data": propertyFunction_s_S0_a__div__Kd.m

```
function [f, grad_f] = propertyFunction_s_S0_a__div__Kd(theta,index__s,index__S0,index__a__div__Kd)
% propertyFunction
%
% returns the difference of the reaction rates as defined in 
% logLikelihood.m with derivatives
%
% Parameters: 
%  theta: Model parameters [theta_1, theta_2]'
%
% Return values:
%  f: double, value of property function
%  grad_f: double vector, gradient of property function


%% Property Function Evaluation
if isempty(index__s)
    f = theta(index__S0) + theta(index__a__div__Kd);
    grad_f = zeros(length(theta),1);
    grad_f([index__S0,index__a__div__Kd]) = 1;
else
    f = theta(index__s) + theta(index__S0) + theta(index__a__div__Kd);
    grad_f = zeros(length(theta),1);
    grad_f([index__s,index__S0,index__a__div__Kd]) = 1;
end

end
```
