## Supplementary material for "Mechanistic description of spatial processes using integrative modelling of noise-corrupted imaging data": setdefault.m

```
% setdefault sets the not defined fields of the 
%    structure object to the default values specified
%    in the structure defaultobject.
%
% [object] = setdefault(object,defaultobject)

function object = setdefault(object,defaultobject)

% Checks whether input object is empty
if isempty(object)
    % If no input is provided return default
    object = defaultobject;
elseif isstruct(object)
    % Loop over all elements of object and defaultobject
    for j = 1:max(length(defaultobject),length(object))
        % Number of elements in object smaller than number of
        % elements in defaultobject?
        if j <= length(object)
            % Generate list of field names
            fieldlist = fieldnames(defaultobject);
            for i = 1:length(fieldlist)
                % Checks whether field exists
                if any(strcmp(fieldnames(object),fieldlist{i}))
                    % Field exists -> call setdefault.m and set all subfields to the default values
                    object = setfield(object,{j},fieldlist{i},...
                               setdefault(getfield(object,{j},fieldlist{i}),getfield(defaultobject,fieldlist{i})));
                else    
                    % Field does not exist -> set to default value
                    object = setfield(object,{j},fieldlist{i},getfield(defaultobject,fieldlist{i}));
                end
            end
        else
          	object(j) = defaultobject(j);
        end
    end
end
```
