## Supplementary material for "Mechanistic description of spatial processes using integrative modelling of noise-corrupted imaging data": GEOMETRY_CCL21_1.m

```
clear all; close all;
addpath('../LineSegments/')

% Read the sample image in
im = imread('LVMASK-CCL21-1.tif');
edgeim = edge(im,'canny', [0.1 0.2], 1);
[edgelist, labelededgeim] = edgelink(edgeim, 10);

% Fit line segments to the edgelists 
tol = 2;
seglist = lineseg(edgelist, tol);
drawedgelist(seglist,size(im),2, 'rand', 1)

%% Create PDE geometry
%%% create square region 

pdepoly([1 512 512 1],[1 1 512 512],'Out1')

% extend to borders
for i = 1:length(seglist)
    seglist{i}(seglist{i}==2) = 1;
    seglist{i}(seglist{i}==511) = 512;
end
    
%%% create LV Polygones
% Polygone 1   
P1x = seglist{1}(:,1)';
P1y = seglist{1}(:,2)';

% Polygone 2   
P2x = [seglist{2}(:,1);...
       seglist{3}(end:-1:1,1);...
       seglist{5}(end:-1:1,1);...
       seglist{4}(:,1)]'; 
P2y = [seglist{2}(:,2);...
       seglist{3}(end:-1:1,2);...
       seglist{5}(end:-1:1,2);...
       seglist{4}(:,2)]';

% Polygone 3   
P3x = [seglist{8}(:,1);...
       seglist{7}(end:-1:1,1);...
       seglist{9}(:,1)]';
P3y = [seglist{8}(:,2);...
       seglist{7}(end:-1:1,2);...
       seglist{9}(:,2)]';  

% Polygone 4  
P4x = seglist{6}(:,1)';
P4y = seglist{6}(:,2)';

pdepoly(P1x,P1y,'In1'); 
pdepoly(P2x,P2y,'In2'); 
pdepoly(P3x,P3y,'In3');
pdepoly(P4x,P4y,'In4');

%% export geometry by hand and create gd sf ns
%%% create geometry file
lv = [1,1,1,0,0,0,0,0,0,0,1]; % 1 if LV 0 oif not

%%% add boundary conditions by hand and export g, b
save('GEOMETRY-CCL21-1.mat','g','b','lv')

%% test
load('GEOMETRY-CCL21-1.mat')

parameter=struct('S0', 1, 'D',1e-4,'a',1e-3,'KD', 1, 'g', 0.1,'off',1e-4);

% initialize grid for PDE calculation beforehand
[p,e,t] = initmesh(g);

%%% Run model
% coefficients
c = parameter.D;
a = parameter.g;

f = num2str(lv(1)*parameter.a);
for i = 2:length(lv)
    f = [f,'!',num2str(lv(i)*parameter.a)];
end

%%% solve PDE
tic
v = assempde(b, p, e, t, c, a, f);
toc

tic
[v2,p2,e2,t2] = adaptmesh(g, b, c, a, f,'mesh',p,e,t,'Maxt',10000);
toc

comp = parameter.S0 - parameter.S0./(((1/parameter.KD)*v)+1);

figure;
subplot(2,3,1)
    pdemesh(p,e,t);
subplot(2,3,4)
    pdesurf(p,t,v);
subplot(2,3,2)
    pdemesh(p2,e2,t2);
subplot(2,3,5)
    pdesurf(p2,t2,v2);
subplot(2,3,6)
    mx.Nx = 512;
    mx.Ny = 512;
    mx.hx = 1/(mx.Nx-1);
    mx.hy = 1/(mx.Ny-1);
    mx.x = (0:mx.hx:1); 
    mx.y = (0:mx.hy:1);

    g = abs(tri2grid(p,t,v,mx.x,mx.y) - tri2grid(p2,t2,v2,mx.x,mx.y));
    imagesc(g); colorbar;
```
