## Supplementary material for "Mechanistic description of spatial processes using integrative modelling of noise-corrupted imaging data": GEOMETRY_CCL21_4.m

```
clear all; close all;
addpath('../LineSegments/')

% Read the sample image in
im = imread('LVMASK-CCL21-4.tif');
edgeim = edge(im,'canny', [0.1 0.2], 1);
[edgelist, labelededgeim] = edgelink(edgeim, 10);

% Fit line segments to the edgelists 
tol = 2;
seglist = lineseg(edgelist, tol);
drawedgelist(seglist,size(im),2, 'rand', 1)

%% Create PDE geometry
%%% create square region 

pdepoly([1 512 512 1],[1 1 512 512],'Out1')

% extend to borders
for i = 1:length(seglist)
    seglist{i}(seglist{i}==2) = 1;
    seglist{i}(seglist{i}==511) = 512;
end

%%% create LV Polygones
% Polygone 1   
P1x = seglist{3}(:,1)';
P1y = seglist{3}(:,2)';

% Polygone 2   
P2x = [seglist{1}(:,1);...
       seglist{2}(end:-1:1,1)]'; 
P2y = [seglist{1}(:,2);...
       seglist{2}(end:-1:1,2)]';

% Polygone 3   
P3x = [seglist{4}(:,1);...
       seglist{5}(end:-1:1,1)]'; 
P3y = [seglist{4}(:,2);...
       seglist{5}(end:-1:1,2)]';   

% Polygone 4   
P4x = seglist{6}(:,1)'; 
P4y = seglist{6}(:,2)'; 

% Polygone 5   
P5x = seglist{7}(:,1)'; 
P5y = seglist{7}(:,2)';  

pdepoly(P1x,P1y,'In1'); 
pdepoly(P2x,P2y,'In2'); 
pdepoly(P3x,P3y,'In3'); 
pdepoly(P4x,P4y,'In4');
pdepoly(P5x,P5y,'In5'); 

%% export geometry by hand and create gd sf ns
%%% create geometry file
lv = [1,1,1,0,0,0,1,1]; % 1 if LV 0 oif not

%%% add boundary conditions by hand and export g, b
save('GEOMETRY-CCL21-4.mat','g','b','lv')

%% test
load('GEOMETRY-CCL21-4.mat')

parameter=struct('S0', 1, 'D',1e-4,'a',1e-3,'KD', 1, 'g', 0.1,'off',1e-4);

% initialize grid for PDE calculation beforehand
[p,e,t] = initmesh(g);

%%% Run model
% coefficients
c = parameter.D;
a = parameter.g;

f = num2str(lv(1)*parameter.a);
for i = 2:length(lv)
    f = [f,'!',num2str(lv(i)*parameter.a)];
end

%%% solve PDE
v = assempde(b, p, e, t, c, a, f);
comp = parameter.S0 - parameter.S0./(((1/parameter.KD)*v)+1);

figure;
subplot(1,3,1)
    pdemesh(p,e,t);
subplot(1,3,2)
    pdesurf(p,t,v);
subplot(1,3,3)
    pdesurf(p,t,comp);
```
