## Supplementary material for "Mechanistic description of spatial processes using integrative modelling of noise-corrupted imaging data": Contents.m

```
% Functions enabling use of multiple colormaps per figure.
%
% version 2.3, 3/2007
%
%   freezeColors    - Lock colors of plot, enabling multiple colormaps per figure.
%   unfreezeColors  - Restore colors of a plot to original indexed color.
%   
% demo/html
%   freezeColors_pub.html   - Published demonstration.
%
% test
%   test_main       - Test function.

% AUTHOR
% John Iversen, 2005-10
%
% 
% Free for any use, so long as AUTHOR information remains with code.
%
% HISTORY
% 
% JRI 6/2005  (v1)
% JRI 9/2006  (v2.1) now operates on all objects with CData (not just images as before)
% JRI 11/2006 handle NaNs in images/surfaces (was not posted on file exchange, included in v2.3)
% JRI 3/2007  (v2.3) Fix bug in handling of caxis that made colorbar scales on frozen plots incorrect. 
% JRI 4/2007  Add 'published' html documentation.
% JRI 9/2010  Changed documentation for colorbars
```
