## Supplementary material for "Mechanistic description of spatial processes using integrative modelling of noise-corrupted imaging data": freezeColors_pub.html

Demonstration of freezeColors / unfreezeColors


 


### Demonstration of freezeColors / unfreezeColors

**freezeColors** enables you to use multiple colormaps in a single figure.

John Iversen

#### Contents

- Problem: In MATLAB there is only one colormap per figure.
- Solution: freezeColors
- Usage
- Demonstration: Plot a variety of objects using different colormaps in one figure.
- unfreezeColors undoes the effects of freezeColors.
- More information
- Credits

#### Problem: In MATLAB there is only one colormap per figure.

#### Solution: freezeColors

**freezeColors** provides an easy solution to create plots using different colormaps in the same figure.

**freezeColors** will freeze the colors of graphics objects in the current axis so that later changes to the colormap (or caxis) will not
change the colors of these objects. Then, a new colormap can be applied to the next plot without changing the appearance of
the first axis. The original, indexed, color data is saved, and can be restored using **unfreezeColors**, making objects one again subject to change with the colormap.

**freezeColors** / **unfreezeColors** work on images, surfaces, scattergroups, bargroups, patches, etc. (Any object with CData in indexed-color mode).

#### Usage

The basic way to do this is to follow each plot with a call to **freezeColors**, e.g.

subplot(2,1,1); imagesc(peaks); colormap hot; freezeColors

subplot(2,1,2); imagesc(peaks); colormap jet; freezeColors, etc...

colorbars may be frozen using colorbar; cbfreeze

**freezeColors** freezes colors of all indexed-color objects in current axis. **freezeColors(axh)** does the same, but for objects in axis axh.

**unfreezeColors** works similarly, but unfreezes colors, restoring objects to their original state, once again subject to the colormap and
caxis.

**cbfreeze** is by Carlos Adrian Vargas Aguilera and must be downloaded separately from the fileexchange here.

#### Demonstration: Plot a variety of objects using different colormaps in one figure.

Below, you will see plots using different colormaps on the same figure. Hooray! Note how after each plot, **freezeColors** is called, making the plot immune to subsequent changes in colormap used to affect the appearance of the next plot.

The figure demonstrates the range of plots that can be used with **freezeColors**: images (imagesc, pcolor), surfaces (surf and surfl), scatter plots, bar plots, indeed any plot object that has CData.

```
figure; set(gcf,'color',[1 1 1])

% image, colormap JET
subplot(3,2,1); imagesc(peaks); axis xy; colormap jet; title('imagesc, jet');
    freezeColors            %freeze colors of current plot
    colorbar; cbfreeze      %how to freeze a colorbar

% same image, using colormap HOT
subplot(3,2,2); imagesc(peaks); axis xy; title('imagesc, hot');
    colormap hot            %now, changing the colormap affects ONLY the current axis!
    freezeColors
    colorbar; cbfreeze

% surface
subplot(3,2,3); surf(peaks); shading interp; colormap hsv; title('surf, hsv');
    freezeColors; colorbar; cbfreeze

% lighted surface, with hole showing nan transparency is preserved after freezing
pnan = peaks; pnan(4:8,end-7:end-3) = nan; % make a small transparent patch
subplot(3,2,4); surfl(pnan); shading interp; colormap hot; title('surfl with NaNs, hot');
    freezeColors; colorbar; cbfreeze

% scatter plot and bar plot
subplot(3,2,5); scatter(randn(100,1),randn(100,1),rand(100,1)*100,rand(100,1),'filled');
    title('scatter, cool'); colormap cool; axis(3*[-1 1 -1 1]);
    freezeColors; colorbar; cbfreeze

subplot(3,2,6); bar(randn(4,3));xlim([0 5]);title('bar, copper'); colormap copper;
    freezeColors;
```

#### unfreezeColors undoes the effects of freezeColors.

While it is used less often, with **unfreezeColors** it is possible to restore a plot to its original state, meaning that it will now be subject to the current colormap. (The
original color data was stored when freezeColors was first called.

Demo: If we change the colormap, then unfreeze the entire figure, all the plots will use the same colormap. This is Matlab's
standard, dreary behavior of one colormap per figure.

```
colormap gray
unfreezeColors(gcf) %unfreeze entire figure
cbfreeze('off') %unfreeze all colorbars
```

#### More information

```
help freezeColors
help unfreezeColors
```

#### Credits

Free for all uses, but please retain the following:

```
Original Author:
John Iversen, 2005-10

```

```
3/23/05
```

Published with MATLAB® 7.4
