## Supplementary material for "Mechanistic description of spatial processes using integrative modelling of noise-corrupted imaging data": freezeColors.m

```
function freezeColors(varargin)
% freezeColors  Lock colors of plot, enabling multiple colormaps per figure. (v2.3)
%
%   Problem: There is only one colormap per figure. This function provides
%       an easy solution when plots using different colomaps are desired 
%       in the same figure.
%
%   freezeColors freezes the colors of graphics objects in the current axis so 
%       that subsequent changes to the colormap (or caxis) will not change the
%       colors of these objects. freezeColors works on any graphics object 
%       with CData in indexed-color mode: surfaces, images, scattergroups, 
%       bargroups, patches, etc. It works by converting CData to true-color rgb
%       based on the colormap active at the time freezeColors is called.
%
%   The original indexed color data is saved, and can be restored using
%       unfreezeColors, making the plot once again subject to the colormap and
%       caxis.
%
%
%   Usage:
%       freezeColors        applies to all objects in current axis (gca),
%       freezeColors(axh)   same, but works on axis axh.
%
%   Example:
%       subplot(2,1,1); imagesc(X); colormap hot; freezeColors
%       subplot(2,1,2); imagesc(Y); colormap hsv; freezeColors etc...
%
%       Note: colorbars must also be frozen. Due to Matlab 'improvements' this can
%				no longer be done with freezeColors. Instead, please
%				use the function CBFREEZE by Carlos Adrian Vargas Aguilera
%				that can be downloaded from the MATLAB File Exchange
%				(http://www.mathworks.com/matlabcentral/fileexchange/24371)
%
%       h=colorbar; cbfreeze(h), or simply cbfreeze(colorbar)
%
%       For additional examples, see test/test_main.m
%
%   Side effect on render mode: freezeColors does not work with the painters
%       renderer, because Matlab doesn't support rgb color data in
%       painters mode. If the current renderer is painters, freezeColors
%       changes it to zbuffer. This may have unexpected effects on other aspects
%	      of your plots.
%
%       See also unfreezeColors, freezeColors_pub.html, cbfreeze.
%
%
%   John Iversen 3/23/05
%

%   Changes:
%   JRI 4/19/06   Correctly handles scaled integer cdata
%   JRI 9/1/06   should now handle all objects with cdata: images, surfaces, 
%                scatterplots. (v 2.1)
%   JRI 11/11/06 Preserves NaN colors. Hidden option (v 2.2, not uploaded)
%   JRI 3/17/07  Preserve caxis after freezing--maintains colorbar scale (v 2.3)
%   JRI 4/12/07  Check for painters mode as Matlab doesn't support rgb in it.
%   JRI 4/9/08   Fix preserving caxis for objects within hggroups (e.g. contourf)
%   JRI 4/7/10   Change documentation for colorbars

% Hidden option for NaN colors:
%   Missing data are often represented by NaN in the indexed color
%   data, which renders transparently. This transparency will be preserved
%   when freezing colors. If instead you wish such gaps to be filled with 
%   a real color, add 'nancolor',[r g b] to the end of the arguments. E.g. 
%   freezeColors('nancolor',[r g b]) or freezeColors(axh,'nancolor',[r g b]),
%   where [r g b] is a color vector. This works on images & pcolor, but not on
%   surfaces.
%   Thanks to Fabiano Busdraghi and Jody Klymak for the suggestions. Bugfixes 
%   attributed in the code.

% Free for all uses, but please retain the following:
%   Original Author:
%   John Iversen, 2005-10
%  

appdatacode = 'JRI__freezeColorsData';

[h, nancolor] = checkArgs(varargin);

%gather all children with scaled or indexed CData
cdatah = getCDataHandles(h);

%current colormap
cmap = colormap;
nColors = size(cmap,1);
cax = caxis;

% convert object color indexes into colormap to true-color data using 
%  current colormap
for hh = cdatah',
    g = get(hh);
    
    %preserve parent axis clim
    parentAx = getParentAxes(hh);
    originalClim = get(parentAx, 'clim');    
   
    %   Note: Special handling of patches: For some reason, setting
    %   cdata on patches created by bar() yields an error,
    %   so instead we'll set facevertexcdata instead for patches.
    if ~strcmp(g.Type,'patch'),
        cdata = g.CData;
    else
        cdata = g.FaceVertexCData; 
    end
    
    %get cdata mapping (most objects (except scattergroup) have it)
    if isfield(g,'CDataMapping'),
        scalemode = g.CDataMapping;
    else
        scalemode = 'scaled';
    end
    
    %save original indexed data for use with unfreezeColors
    siz = size(cdata);
    setappdata(hh, appdatacode, {cdata scalemode});

    %convert cdata to indexes into colormap
    if strcmp(scalemode,'scaled'),
        %4/19/06 JRI, Accommodate scaled display of integer cdata:
        %       in MATLAB, uint * double = uint, so must coerce cdata to double
        %       Thanks to O Yamashita for pointing this need out
        idx = ceil( (double(cdata) - cax(1)) / (cax(2)-cax(1)) * nColors);
    else %direct mapping
        idx = cdata;
        %10/8/09 in case direct data is non-int (e.g. image;freezeColors)
        % (Floor mimics how matlab converts data into colormap index.)
        % Thanks to D Armyr for the catch
        idx = floor(idx);
    end
    
    %clamp to [1, nColors]
    idx(idx<1) = 1;
    idx(idx>nColors) = nColors;

    %handle nans in idx
    nanmask = isnan(idx);
    idx(nanmask)=1; %temporarily replace w/ a valid colormap index

    %make true-color data--using current colormap
    realcolor = zeros(siz);
    for i = 1:3,
        c = cmap(idx,i);
        c = reshape(c,siz);
        c(nanmask) = nancolor(i); %restore Nan (or nancolor if specified)
        realcolor(:,:,i) = c;
    end
    
    %apply new true-color color data
    
    %true-color is not supported in painters renderer, so switch out of that
    if strcmp(get(gcf,'renderer'), 'painters'),
        set(gcf,'renderer','zbuffer');
    end
    
    %replace original CData with true-color data
    if ~strcmp(g.Type,'patch'),
        set(hh,'CData',realcolor);
    else
        set(hh,'faceVertexCData',permute(realcolor,[1 3 2]))
    end
    
    %restore clim (so colorbar will show correct limits)
    if ~isempty(parentAx),
        set(parentAx,'clim',originalClim)
    end
    
end %loop on indexed-color objects


% ============================================================================ %
% Local functions

%%%%%%%%%%%%%%%%%%%%%%%%%%%%%%%%%%%%%%%%%%%%%%%%%%%%%%%%%%%%%%%%%%%%%%%%%%%%%%%%
%% getCDataHandles -- get handles of all descendents with indexed CData
%%%%%%%%%%%%%%%%%%%%%%%%%%%%%%%%%%%%%%%%%%%%%%%%%%%%%%%%%%%%%%%%%%%%%%%%%%%%%%%%

function hout = getCDataHandles(h)
% getCDataHandles  Find all objects with indexed CData

%recursively descend object tree, finding objects with indexed CData
% An exception: don't include children of objects that themselves have CData:
%   for example, scattergroups are non-standard hggroups, with CData. Changing
%   such a group's CData automatically changes the CData of its children, 
%   (as well as the children's handles), so there's no need to act on them.

error(nargchk(1,1,nargin,'struct'))

hout = [];
if isempty(h),return;end

ch = get(h,'children');
for hh = ch'
    g = get(hh);
    if isfield(g,'CData'),     %does object have CData?
        %is it indexed/scaled?
        if ~isempty(g.CData) && isnumeric(g.CData) && size(g.CData,3)==1, 
            hout = [hout; hh]; %#ok<AGROW> %yes, add to list
        end
    else %no CData, see if object has any interesting children
            hout = [hout; getCDataHandles(hh)]; %#ok<AGROW>
    end
end

%%%%%%%%%%%%%%%%%%%%%%%%%%%%%%%%%%%%%%%%%%%%%%%%%%%%%%%%%%%%%%%%%%%%%%%%%%%%%%%%
%% getParentAxes -- return handle of axes object to which a given object belongs
%%%%%%%%%%%%%%%%%%%%%%%%%%%%%%%%%%%%%%%%%%%%%%%%%%%%%%%%%%%%%%%%%%%%%%%%%%%%%%%%
function hAx = getParentAxes(h)
% getParentAxes  Return enclosing axes of a given object (could be self)

error(nargchk(1,1,nargin,'struct'))
%object itself may be an axis
if strcmp(get(h,'type'),'axes'),
    hAx = h;
    return
end

parent = get(h,'parent');
if (strcmp(get(parent,'type'), 'axes')),
    hAx = parent;
else
    hAx = getParentAxes(parent);
end


%%%%%%%%%%%%%%%%%%%%%%%%%%%%%%%%%%%%%%%%%%%%%%%%%%%%%%%%%%%%%%%%%%%%%%%%%%%%%%%%
%% checkArgs -- Validate input arguments
%%%%%%%%%%%%%%%%%%%%%%%%%%%%%%%%%%%%%%%%%%%%%%%%%%%%%%%%%%%%%%%%%%%%%%%%%%%%%%%%

function [h, nancolor] = checkArgs(args)
% checkArgs  Validate input arguments to freezeColors

nargs = length(args);
error(nargchk(0,3,nargs,'struct'))

%grab handle from first argument if we have an odd number of arguments
if mod(nargs,2),
    h = args{1};
    if ~ishandle(h),
        error('JRI:freezeColors:checkArgs:invalidHandle',...
            'The first argument must be a valid graphics handle (to an axis)')
    end
    % 4/2010 check if object to be frozen is a colorbar
    if strcmp(get(h,'Tag'),'Colorbar'),
      if ~exist('cbfreeze.m'),
        warning('JRI:freezeColors:checkArgs:cannotFreezeColorbar',...
            ['You seem to be attempting to freeze a colorbar. This no longer'...
            'works. Please read the help for freezeColors for the solution.'])
      else
        cbfreeze(h);
        return
      end
    end
    args{1} = [];
    nargs = nargs-1;
else
    h = gca;
end

%set nancolor if that option was specified
nancolor = [nan nan nan];
if nargs == 2,
    if strcmpi(args{end-1},'nancolor'),
        nancolor = args{end};
        if ~all(size(nancolor)==[1 3]),
            error('JRI:freezeColors:checkArgs:badColorArgument',...
                'nancolor must be [r g b] vector');
        end
        nancolor(nancolor>1) = 1; nancolor(nancolor<0) = 0;
    else
        error('JRI:freezeColors:checkArgs:unrecognizedOption',...
            'Unrecognized option (%s). Only ''nancolor'' is valid.',args{end-1})
    end
end
```
