## Supplementary material for "Mechanistic description of spatial processes using integrative modelling of noise-corrupted imaging data": test_main.m

```
% test_main Test freezeColors / unfreezeColors
%
%   JRI 4/2007 

%   Author:
%   John Iversen
%  

disp(' ')
disp('==================================================')
disp('==freezeColors / unfreezeColors interactive test==')
disp('==================================================')
disp(' ')
disp('freezeColors allows multiple colormaps to be used per figure.')
disp(' ')

% ============================================================================ %

disp('As this is a graphics function, you will have to use your eyes to verify')
disp('  correct behavior. In addition, no errors should be reported.')
disp(' ')
disp('First, An image is drawn, using the hot colormap.')
disp(' ')

figure
set(gcf,'color',[1 1 1],'renderer','zbuffer')
% later, freezeColors will change from painters to zbuffer when it
%   is called, which will change the appearance of the unfrozen plot slightly.
%   This is not desired, so start out with zbuffer mode.

subplot(3,2,1)
imagesc(peaks); axis xy; colormap hot; title('colors not frozen'); h1=colorbar;

disp('>> subplot(3,2,1)')
disp('>> imagesc(peaks); axis xy; colormap hot; title(''colors not frozen''); colorbar')

% ============================================================================ %

disp(' ')
disp('Next, if we change the colormap to jet, the plot will change appearance.')
disp('  This is normal Matlab behavior.')
disp(' =Hit a key='), pause, disp(' ')

disp('>> colormap jet')
colormap jet

% ============================================================================ %

disp(' ')
disp('Now, repeat the plot, but using freezeColors. We also freeze the colorbar,')
disp('  using the function cbfreeze by Carlos Adrian Vargas Aguilera. Note: this')
disp('  function must be downloaded separately from the MATLAB Central File Exchange!')

disp(' =Hit a key='), pause, disp(' ')

subplot(3,2,2)
imagesc(peaks); axis xy; colormap jet; title('jet colormap, frozen')
freezeColors
h=colorbar;
try
  cbfreeze(h1)
	cbfreeze(h)
catch
	disp('cbfreeze is not available, so colorbar colors will not be frozen until you')
	disp(' go and download it from the file exchange. Sorry for the extra step, but')
  disp(' it is necessary because of matlab internal changes. Please visit')
  disp('  http://www.mathworks.com/matlabcentral/fileexchange/24371 to download it.')
	freezeColors(h) %tests passing an axis handle 
end

disp('>> subplot(3,2,2)')
disp('>> imagesc(p); axis xy; colormap jet; title(''jet colormap, frozen'')')
disp(' ')
disp('%% The test of freezeColors %%')
disp('>> freezeColors')
disp('>> colorbar')
disp('>> cbfreeze %NOTE: this is separately available from File Exchange')


% ============================================================================ %

disp(' ')
disp('The two plots should appear identical.')
disp(' ')
disp('However, if we now change the colormap again, this time to cool, the left')
disp('  plot should change but the frozen one on the right will remain the same.')
disp('  This shows that freezeColor is working as intended because the frozen')
disp('  plot is no longer affected by the colormap.')
disp(' =Hit a key='), pause, disp(' ')

colormap cool

disp('>> colormap cool')

% ============================================================================ %

disp(' ')
disp('Next, test some other plot types (image, scatter, bar, surf), as well as')
disp('  correct handling of NaNs in images. You will see a variety of plots,')
disp('  using a variety of different colormaps--all on the same figure.')
disp(' =Hit a key='), pause, disp(' ')

clf

subplot(3,2,1); imagesc(peaks); axis xy; colormap hsv; title('imagesc, hsv');
freezeColors; h=colorbar;
try, cbfreeze(h), catch,freezeColors(h), end

subplot(3,2,2); surf(peaks); shading interp; colormap jet; title('surf, jet');
freezeColors; h=colorbar;
try, cbfreeze(h), catch,freezeColors(h), end

subplot(3,2,3); scatter(randn(100,1),randn(100,1),rand(100,1)*100,rand(100,1),'filled');
title('scatter, cool'),colormap cool; axis(3*[-1 1 -1 1]);
freezeColors; h=colorbar;
try, cbfreeze(h), catch,freezeColors(h), end
    
subplot(3,2,4); bar(randn(4,4));xlim([0 5]);title('bar, copper')
colormap copper; freezeColors;
    
%demonstrate handling of NaNs -- ordinarily these are preserved, but there's a hidden
%   option to subsitute another color for NaN

c = magic(50); c(20:30,20:30) = nan;
subplot(3,2,5); pcolor(c); shading interp; colormap hot; title('NaNs remain transparent')
view(-67,56); %so grid shows through
set(gca,'color',[0 1 0])
grid on, box off
freezeColors

subplot(3,2,6); pcolor(c); shading interp; colormap hot; title('NaN (using ''nancolor'' option)')
view(-67,56); %so grid shows through
set(gca,'color',[0 1 0])
grid on, box off
freezeColors('nancolor',[0 0 1]); %test two argument form

% ============================================================================ %

disp(' ')
disp('Finally, test unfreezeColors by unfreezing the entire figure, after')
disp('  changing colormap to gray.')
disp(' ')
disp('All plots are restored to the unfrozen state and are once again')
disp('  influenced by the colormap.')
disp(' =Hit a key='), pause, disp(' ')

colormap gray
unfreezeColors(gcf)
try
	cbfreeze('off')
end

disp('>> colormap gray')
disp('>> unfreezeColors(gcf)')
disp('>> cbfreeze(''off'')')

disp(' ')
disp('Verify that all data is now displayed with the gray colormap. Now that')
disp('  the effects of freezeColors are undone, we see Matlab''s default')
disp('  single colormap per figure.')
disp('  ')
disp('==End of Test==')
```
