## Supplementary material for "Mechanistic description of spatial processes using integrative modelling of noise-corrupted imaging data": unfreezeColors.m

```
function unfreezeColors(h)
% unfreezeColors  Restore colors of a plot to original indexed color. (v2.3)
%
%   Useful if you want to apply a new colormap to plots whose
%       colors were previously frozen with freezeColors.
%
%   Usage:
%       unfreezeColors          unfreezes all objects in current axis, 
%       unfreezeColors(axh)     same, but works on axis axh. axh can be vector.
%       unfreezeColors(figh)    same, but for all objects in figure figh.
%
%       Has no effect on objects on which freezeColors was not already called.
%				(Note: if colorbars were frozen using cbfreeze, use cbfreeze('off') to 
%       unfreeze them. See freezeColors for information on cbfreeze.)
%
%
%   See also freezeColors, freezeColors_pub.html, cbfreeze.
%
%
%   John Iversen 3/23/05
%

%   Changes:
%   JRI 9/1/06 now restores any object with frozen CData;
%              can unfreeze an entire figure at once.
%   JRI 4/7/10 Change documentation for colorbars

% Free for all uses, but please retain the following:
%
%   Original Author:
%   John Iversen, 2005-10
%  

error(nargchk(0,1,nargin,'struct'))

appdatacode = 'JRI__freezeColorsData';

%default: operate on gca
if nargin < 1,
    h = gca;
end

if ~ishandle(h),
     error('JRI:unfreezeColors:invalidHandle',...
            'The argument must be a valid graphics handle to a figure or axis')
end

%if h is a figure, loop on its axes
if strcmp(get(h,'type'),'figure'),
    h = get(h,'children');
end

for h1 = h', %loop on axes

    %process all children, acting only on those with saved CData
    %   ( in appdata JRI__freezeColorsData)
    ch = findobj(h1);
    
    for hh = ch',
        
        %some object handles may be invalidated when their parent changes 
        %   (e.g. restoring colors of a scattergroup unfortunately changes
        %   the handles of all its children). So, first check to make sure
        %   it's a valid handle
        if ishandle(hh)
            if isappdata(hh,appdatacode),
                ad = getappdata(hh,appdatacode);
                %get oroginal cdata
                %patches have to be handled separately (see note in freezeColors)
                if ~strcmp(get(hh,'type'),'patch'),
                    cdata = get(hh,'CData');
                else
                    cdata = get(hh,'faceVertexCData');
                    cdata = permute(cdata,[1 3 2]);
                end
                indexed = ad{1};
                scalemode = ad{2};
                
                %size consistency check
                if all(size(indexed) == size(cdata(:,:,1))),
                    %ok, restore indexed cdata
                    if ~strcmp(get(hh,'type'),'patch'),
                        set(hh,'CData',indexed);
                    else
                        set(hh,'faceVertexCData',indexed);
                    end
                    %restore cdatamapping, if needed
                    g = get(hh);
                    if isfield(g,'CDataMapping'),
                        set(hh,'CDataMapping',scalemode);
                    end
                    %clear appdata
                    rmappdata(hh,appdatacode)
                else
                    warning('JRI:unfreezeColors:internalCdataInconsistency',...
                        ['Could not restore indexed data: it is the wrong size. ' ...
                        'Were the axis contents changed since the call to freezeColors?'])
                end
                
            end %test if has our appdata
        end %test ishandle

    end %loop on children

end %loop on axes
```
