## Supplementary material for "Mechanistic description of spatial processes using integrative modelling of noise-corrupted imaging data": generateArtificialData.m

```
% generateArtificialData.m generates the outlier-corrupted artificial
% experimental data:
%
% Data types:
%   1D Feature ... saved to the folder Feat1D. Contains the distancemeasure
%                  extracted from the 2D images generated. Named
%                  FeatData-NR.mat with NR = 1..30 containing the 
%                  simulation Y and data for YNs with Ns = 0,20,40,80,160,
%                  320 and 640 spots                 
%   2D Images  ... saved to the folder Full2D. Contains the 2D images 
%                  generated. Named FeatData-NR.mat with NR = 1..30 
%                  containing the simulation Y and data for YNs with Ns = 
%                  0,20,40,80,160,20 and 640 spots            
% The data is used to perform the parameter estimation for the 
%
% 2014/04/08 - Sabrina Hross
% 2017/11/11 - Jan Hasenauer

clear all;
close all;
clc;

%% OPTIONS
exp_nr = 1;    % Index of image which is used as basis for data generation
               % (Remark: This has to be a scalar.)
num_sets = 30; % Number of artifical datasets
spots_nr = [0,20,40,80,160,320,640];    % Number of spots

% Parameters used to generate artifical data
true_par = [8.5e+1;   % D
            2.4e-1;   % a
            2.1e-1;   % S0
            1.3e-2;   % off
            1.44e-1]; % sig 

%% ADD PATH AND CHECK AVAILABILITY OF TOOLBOXES
% Add path
addpath(genpath('./Auxillary'));

% Check that necessary MATLAB toolboxes are installed
if ~license('test','PDE_Toolbox')
	error('MATLAB PDE toolbox is required to run this code.');
end
if ~license('test','Statistics_Toolbox')
	error('MATLAB statistics toolbox is required to run this code.');
end

%% LOADING OF MEASURED DATA AND GEOMETRY, AND DEFINITION OF OPTIONS
% Load data and geometry
load('./Auxillary/mycmaps.mat')
data = load(['./ExperimentalData/CCL21-'         ,num2str(exp_nr),'.mat']);
geom = load(['./ExperimentalData/GEOMETRY-CCL21-',num2str(exp_nr),'.mat']);

% Normalization of image and extraction of mask
options.region.im = data.img./255;
options.region.mask = data.mask;

% Filtering for the subsequent calculation of outlier statistics
[options.region.index,options.region.msers] = performFiltering(options.region.im,options.region.mask);
options.region.removed = (size(data.mask,1)*size(data.mask,2)-sum(sum(options.region.index)))/(size(data.mask,1)*size(data.mask,2));
options.region.im_filter = nan(size(options.region.im));
options.region.im_filter(options.region.index) = options.region.im(options.region.index);

% Definition of grid for distance measure
options.dist.muedges = linspace(1,100,20); % in \mum
options.dist.edges = options.dist.muedges*400/604;

options.dist.xtick = linspace(0,66,5); % 0 bis 100 \mum
options.dist.xticklabel = {'0','25','50','75','100'};
options.dist.xlim = [0 66];

% Consideration of inner part of image to avoid boundary effects
options.dist.ind1 = [56:456];
options.dist.ind2 = [56:456];

% Assignment of index set of pixels for source, sink and pixel at a defined
% distance from the lymphoid vessel
options.dist.distmap = data.Distmap(options.dist.ind1,options.dist.ind2);
options.dist.I_sink = find(options.region.mask(options.dist.ind1,options.dist.ind2)<1); 
options.dist.I_source = find(options.region.mask(options.dist.ind1,options.dist.ind2)>=1);

I = [];
for k = 1:length(options.dist.edges)-1
    I{k} = find(options.dist.distmap(:) >= options.dist.edges(k) & options.dist.distmap(:) < options.dist.edges(k+1));
    if isempty(I{k})
        continue
    end
end
options.dist.I = I;

% Definition of filter properties
BW = logical(1-options.region.index);
stats = regionprops(BW,'all');

% - Size of ellipsoid
ell_size = cat(1,stats.Area);
ind = find(50<ell_size & ell_size<250);
[options.spot.size.mu, options.spot.size.sigma] = normfit(ell_size(ind));

% - Axis length of ellipsoid
ell_a = cat(1,stats(ind).MajorAxisLength);
ell_b = cat(1,stats(ind).MinorAxisLength);

% Fitting of spot statistics
options_gm = statset('Display','final','TolFun',1e-8,'MaxIter',300);
options.spot.Ax = gmdistribution.fit([ell_a,ell_b],1,'Options',options_gm);

%% VISUALIZATION OF EXTRACTED PLOT CHARACTERISTICS
figure('name','Distribution of spot properties');

% Distribution of the lengths of major and minor axis of spots 
subplot(1,2,1);
hcont = ezcontour(@(a,b) pdf(options.spot.Ax,[a b]),[6 40],[2 17]); hold on;
set(hcont, 'Color',[0 0 0]);
plot(ell_a,ell_b,'o','Color',[1,0,0]);
axis(1.05*[0,max(ell_a),0,max(ell_a)]);
xlabel('length of major axis');
ylabel('length of minor axis');
title('Lengths of ellipsoid axes')

% Distribution of the area of the detected spots 
subplot(1,2,2)
x_bin = linspace(0,300,20);
histogram(ell_size(ind),x_bin,'Normalization','pdf','FaceColor',[1,0,0]); hold on;
plot(linspace(x_bin(1),x_bin(end),150),pdf('normal',linspace(x_bin(1),x_bin(end),150),options.spot.size.mu, options.spot.size.sigma),'-k','linewidth',2);
xlabel('convex hull area in px');
ylabel('probability density function');
title('area size');
        
%% CONSTRUCTION OF PDE AND SIMULATION OF NOISE-FREE DATA
% Grid based on images
mx.Nx = size(data.img,1);
mx.Ny = size(data.img,2);
mx.x = 1:512; 
mx.y = 1:512;
[mx.xG,mx.yG] = meshgrid(mx.x,mx.y);

% Assignment of parameter values
parameter_t = zeros(9,1);
parameter_t(1:5) = log10(true_par); 
name = {'D','a','S0','off','sig'};

for j = 1:5
    parameter.(name{j}) = 10^parameter_t(j);
end

% Initialization of matrices for unit parameters
c = parameter.D;
a = 1;
f = num2str(geom.lv(1)*parameter.a);
for j = 2:length(geom.lv)
    f = [f,'!',num2str(geom.lv(j)*parameter.a)];
end

% Initialization of mesh
[p_in,e_in,t_in] = initmesh(geom.g);

% Adaptive calculation of mesh
% For the meshing we consider the model -grad(c grad(u)) + au = f with:
[v,p,e,t] = adaptmesh(geom.g, geom.b, c, a, f,'mesh',p_in,e_in,t_in,'Ngen',8);
[K,M,F,Q,~,~,~] = assempde(geom.b, p, e, t, c, a, f);

% Initialization of matrix for interpolation
[~,tn,a2,a3] = tri2grid(p,t,zeros(size(p,2),1),mx.x,mx.y);

% Assignment of PDE properties to option struct
options.disc = struct('K',K,'M',M,'F',F,'Q',Q,'p',p,'t',t,'e',e,'tn',tn,'a2',a2,'a3',a3,'mx',mx);

% Calculation of noise-free observable
y = tri2grid(p,t,v,tn,a2,a3)';
Y = parameter.off + parameter.S0./(1+1./y);

% Visualization of simulation results
figure('name','Model simulation');
imagesc(log10(Y),[-2.2,0]); axis square; axis off;
colormap(cmap_green_data_NaN);
colorbar('Location','WestOutside','YTick',[-2.2,-2,-1,0],'YTickLabel',{'NaN','1e-2','1e-1','1'});
title('immobilized CCL21');

%% GENERATION OF ARTIFICIAL DATA

% Calculation of subplot layout
ns_1 = ceil(sqrt(length(spots_nr)));
ns_2 = ceil(length(spots_nr)/ns_1);

% Definition of random seed to ensure reproducibility
rng(280186); % (= one of our birthdays)

% Generation of artificial data
for n = 1:num_sets
    % Progress
    disp(['Artificical dataset ' num2str(n) ' is generated.']);

    % Loop: Spots
    for j = 1:length(spots_nr)
        % Initalization of intensities and spot mask
        Y_n = Y;
        Spot_Mask_n = zeros(size(Y_n));

        % Addition of spots
        if spots_nr(j) >= 1
            % Sampling the half-axis lengths, intensities, rotation angles
            % and centers of the spot 
            r = 0.45.*random(options.spot.Ax,spots_nr(j));
            intensity = min(min(Y)) + (max(max(Y))-min(min(Y)))*rand(spots_nr(j),1);
            phi = 360*rand(spots_nr(j),1); % rotation of ellipsoids
            ind = randi(length(data.I_sink),spots_nr(j),1);
            [Mx,My] = ind2sub(size(data.mask),data.I_sink(ind));
            [y,x] = meshgrid(1:size(data.mask,1),1:size(data.mask,2));

            % Addition of spots to simulation results
            % Loop: Spots
            for i = 1:spots_nr(j)
                % Definition of spot position
                pos = find(((cosd(phi(i))*(x-Mx(i))-sind(phi(i))*(y-My(i))).^2/r(i,1)^2 + (sind(phi(i))*(x-Mx(i))+cosd(phi(i))*(y-My(i))).^2/r(i,2)^2)<1);
                % Assignment of spot intensity and spot mask
                Y_n(pos) = Y_n(pos) + intensity(i)*(1-((cosd(phi(i))*(x(pos)-Mx(i))-sind(phi(i))*(y(pos)-My(i))).^2/r(i,1)^2 + (sind(phi(i))*(x(pos)-Mx(i))+cosd(phi(i))*(y(pos)-My(i))).^2/r(i,2)^2));
                Spot_Mask_n(pos) = 1;
            end
        end

        % Addition of multiplicative measurement noise
        Y_n = Y_n.*lognrnd(0,parameter.sig,size(Y,1),size(Y,2));
        
        % Struct of artificial data
        img = Y_n*255;
        mask = data.mask;
        I_source = data.I_source;
        I_sink = data.I_sink;
        Distmap = data.Distmap;
        Distfun = data.Distfun;

        % Save artificial data
        save(['./ArtificialData/CCL21-',num2str(exp_nr),'-',num2str(n),'-',num2str(spots_nr(j)),'.mat'],...
            'img','mask','I_source','I_sink','Distmap','Distfun','parameter');
        
        % Visualize data
        if n == 1
            % Open figure
            if j == 1
                fh = figure('name','Artifical measurement data - sample 1');
            else
                figure(fh);
            end
            
            % Open subplot
            subplot(ns_1,ns_2,j);
            
            % Draw figure
            imagesc(log10(Y_n),[-2.2,0]); colormap(cmap_green_data_NaN);
            axis square; axis off;
            title([num2str(spots_nr(j)) ' spots']);
            
            drawnow;
        end
    end
end
```
