## Supplementary material for "Mechanistic description of spatial processes using integrative modelling of noise-corrupted imaging data": getResultsForFigure4.m

```
% Analysis for Figure 4:
% Parameter estimation results for the 1D and 2D models using extracted
% features and whole imaging data.

clear all;
close all;
clc;

%% DATASET
% Type of experimental data
flags.data_type = 'artificial';

% Index set of images
exp_nr = 1;

% For artifical data:
% - number of spots
spot_nr = 0;
% - replicate number
replicate_nr = 1;

%% OPTIMIZATION AND PROFILE CALCULATION OPTIONS
% Number of local optimization runs
n_starts = 10;

% Parameter profile calculation
flags.parameter_profile_likelihood_calculation = true;
par_ind = 1:4;

% Property profile calculation
flags.property_profile_likelihood_calculation = true;

%% PARAMETER ESTIMATION
[parameters_1D_DDM,properties_1D_DDM,info_1D_DDM,options_1D_DDM] = runParameterEstimation('M1_1D_DDM_SNM',flags,n_starts,exp_nr,spot_nr,replicate_nr,par_ind);
[parameters_2D_DDM,properties_2D_DDM,info_2D_DDM,options_2D_DDM] = runParameterEstimation('M1_2D_DDM_SNM',flags,n_starts,exp_nr,spot_nr,replicate_nr,par_ind);
flags.property_profile_likelihood_calculation = false;
[parameters_2D_WID,properties_2D_WID,info_2D_WID,options_2D_WID] = runParameterEstimation('M1_2D_WID_SNM',flags,n_starts,exp_nr,spot_nr,replicate_nr,par_ind);

% Close figures
close all;

% Confidence interval calculation
alpha = [0.75,0.95,0.99];
parameters_1D_DDM = getParameterConfidenceIntervals(parameters_1D_DDM,alpha,options_1D_DDM);
parameters_2D_DDM = getParameterConfidenceIntervals(parameters_2D_DDM,alpha,options_2D_DDM);
parameters_2D_WID = getParameterConfidenceIntervals(parameters_2D_WID,alpha,options_2D_WID);

%% SIMULATION OF MODEL
[~,~,g_1D_DDM,x_1D_DDM,m_1D_DDM] = nll_M1_1D_DDM_SNM(parameters_1D_DDM.MS.par(:,1),info_1D_DDM);
[~,~,g_2D_DDM,x_2D_DDM,m_2D_DDM] = nll_M1_2D_DDM_SNM(parameters_2D_DDM.MS.par(:,1),info_2D_DDM);
[~,~,g_2D_WID,x_2D_WID,m_2D_WID] = nll_M1_2D_WID_SNM(parameters_2D_WID.MS.par(:,1),setfield(info_2D_WID,'plot',true));

%% VISUALIZATION
% Colors
load('./Auxillary/mycmaps.mat')
Col = [0.20 0.20 0.44;
       0.44 0.68 0.76;
       0.76 0.28 0.28];

% Figure 4A: Fitting result for the 1D model using the distance-dependent
% average intensity of immobilised CCL21 (1D data).
figure('name','Figure 4A: setup (i) - 1D model, 1D data');
plot(x_1D_DDM{1},m_1D_DDM{1},'-r'); hold on;
plot(info_1D_DDM.dist_dep_mean{1}.dist,info_1D_DDM.dist_dep_mean{1}.mean,'ok');
set(gca,'xlim',[0,50],'ylim',[0,0.06]);
xlabel('distance to nearest lymphoid vessel [\mum]');
ylabel('fluorescence intensity [a.u.]');
legend({'model','data'},'location','NorthEast','box','off');

% Figure 4B: Fitting result for the 2D model using the distance-dependent
% average intensity of immobilised CCL21 (2D data).
figure('name','Figure 4B: setup (ii) - 2D model, 1D data');
plot(x_2D_DDM{1},m_2D_DDM{1},'-r'); hold on;
plot(info_2D_DDM.dist_dep_mean{1}.dist,info_2D_DDM.dist_dep_mean{1}.mean,'ok');
set(gca,'xlim',[0,50],'ylim',[0,0.06]);
xlabel('distance to nearest lymphoid vessel [\mum]');
ylabel('fluorescence intensity [a.u.]');
legend({'model','data'},'location','NorthEast','box','off');

% Figure 4C: Fitting result for the 2D model using the measured intensity
% distribution of immobilised CCL21 (2D data).
figure('name','Figure 4C: setup (iii) - 2D model, 2D data');
cmap_green_data_NaN_reduced = cmap_green_data_NaN([1,10,22,26,30,33,36,38,42,46],:);
c_min = log10(0.01); c_max = log10(0.1);

% - measurement
subplot(1,2,1);
imagesc(log10(info_2D_DDM.data{1}),[c_min,c_max]); hold on;
colormap(cmap_green_data_NaN_reduced); 
contour(info_2D_DDM.mask{1},[1 1],'Color',[1,1,1],'Linewidth',1.5)  
set(gca,'xlim',[-0.5,402],'ylim',[-0.5,402.5],'box','on','xtick',[],'ytick',[]);
axis square;
title('measurement');
hcol = colorbar('Location','EastOutside','YTick',log10([0.01,0.02,0.03,0.04,0.05,0.06,0.07,0.08,0.09,0.1]),'YTickLabel',{'<0.01','  0.02','','  0.04','','  0.06','','  0.08','','>0.10'});
hcol.Label.String = 'fluorescence intensity [a.u.]';

% - model fit
subplot(1,2,2);
imagesc(log10(g_2D_WID{1}),[c_min,c_max]); hold on;
colormap(cmap_green_data_NaN_reduced); 
contour(info_2D_DDM.mask{1},[1 1],'Color',[1,1,1],'Linewidth',1.5)  
set(gca,'xlim',[-0.5,402],'ylim',[-0.5,402.5],'box','on','xtick',[],'ytick',[]);
axis square;
title('model fit');
hcol = colorbar('Location','EastOutside','YTick',log10([0.01,0.02,0.03,0.04,0.05,0.06,0.07,0.08,0.09,0.1]),'YTickLabel',{'<0.01','  0.02','','  0.04','','  0.06','','  0.08','','>0.10'});
hcol.Label.String = 'fluorescence intensity [a.u.]';

% Figure 4D: Profile likelihood derived confidence intervals for parameter 
% estimates for setups (i)-(iii).
figure('name','Figure 4D: confidence intervals');
theta_true = [8.5e+1;  % D
              2.4e-1;  % a
              2.1e-1;  % S0
              1.3e-2]; % off
xi_true = log10(theta_true);

w = [0.3,0.225,0.15];

% Loop: Parameters
for i = 1:4
    switch i
        case 1 % D
            p_lim = xi_true(i) + 1.50*[-1,1];
            ylab = 'D/\gamma';
        case 2 % a
            p_lim = xi_true(i) + 0.45*[-1,1];
            ylab = '(\alpha k_1)/(\gamma k_{-1})';
        case 3 % S0
            p_lim = xi_true(i) + 0.45*[-1,1];
            ylab = 's S_0';
        case 4 % b
            p_lim = xi_true(i) + 0.22*[-1,1];
            ylab = 's b';
    end
    subplot(1,4,i);
    % Loop: Confidence levels
    for j = length(alpha):-1:1
        r = j/(length(alpha)+2);
        fill([-1,1,1,-1]*w(j)  ,10.^parameters_1D_DDM.CI.PL(i,[1,1,2,2],j),'k','facecolor',(1-r)*Col(1,:)+r,'edgecolor',Col(1,:)); hold on;
        fill([-1,1,1,-1]*w(j)+1,10.^parameters_2D_DDM.CI.PL(i,[1,1,2,2],j),'k','facecolor',(1-r)*Col(2,:)+r,'edgecolor',Col(2,:)); hold on;
        fill([-1,1,1,-1]*w(j)+2,10.^parameters_2D_WID.CI.PL(i,[1,1,2,2],j),'k','facecolor',(1-r)*Col(3,:)+r,'edgecolor',Col(3,:)); hold on;
        plot([-1,3],theta_true(i)*[1,1],'k-');
    end
    xlim([-0.5,2.5]);
    ylim(10.^p_lim);
    set(gca,'yscale','log','xtick',[0,1,2],'xticklabel',{'(i)','(ii)','(iii)'},'xticklabelrotation',90);
    ylabel(ylab);
end
```
