## Supplementary material for "Mechanistic description of spatial processes using integrative modelling of noise-corrupted imaging data": getResultsForFigure5.m

```
% Analysis for Figure 5:
% Parameter estimation results for the 1D and 2D models using extracted
% features and whole imaging data.
%
% Note: The calculations take several hours.

clear all;
close all;
clc;

%% DATASET
% Type of experimental data
flags.data_type = 'artificial';

% Index set of images
exp_nr = 1;

% For artifical data:
% - number of spots
spot_nr = [0,20,40,80,160,320,640];
% - replicate number
replicate_nr = 1:5; % Note: In the manuscript, we used 30 replicates.

%% OPTIMIZATION AND PROFILE CALCULATION OPTIONS
% Number of local optimization runs
n_starts = 3; % Note: In the manuscript, we used 10 local optimizations.

%% PARAMETER ESTIMATION
for r = 1:length(replicate_nr)
    for s = 1:length(spot_nr)
        disp(['Replicate: ' num2str(r) '/' num2str(length(replicate_nr)) '; Spot number: ' num2str(r) '/' num2str(length(spot_nr))]);
        [parameters_2D_WID_SNM{r,s},properties_2D_WID_SNM{r,s},info_2D_WID_SNM{r,s},options_2D_WID_SNM{r,s}] = runParameterEstimation('M1_2D_WID_SNM',flags,n_starts,exp_nr,spot_nr(s),replicate_nr(r));
        [parameters_2D_FID_SNM{r,s},properties_2D_FID_SNM{r,s},info_2D_FID_SNM{r,s},options_2D_FID_SNM{r,s}] = runParameterEstimation('M1_2D_FID_SNM',flags,n_starts,exp_nr,spot_nr(s),replicate_nr(r));
        [parameters_2D_WID_INM{r,s},properties_2D_WID_INM{r,s},info_2D_WID_INM{r,s},options_2D_WID_INM{r,s}] = runParameterEstimation('M1_2D_WID_INM',flags,n_starts,exp_nr,spot_nr(s),replicate_nr(r));
        close all;
    end
end

%% SIMULATION OF MODEL
s_320 = find(spot_nr==320); % index of the spot number 320 in spot_nr
                            % (this indey has to be non-empty for the
                            % following code to work)

% Update options                            
info_2D_WID_SNM{1,s_320}.plot = true; info_2D_WID_SNM{1,s_320}.fh = 1;
info_2D_FID_SNM{1,s_320}.plot = true; info_2D_FID_SNM{1,s_320}.fh = 2;
info_2D_WID_INM{1,s_320}.plot = true; info_2D_WID_INM{1,s_320}.fh = 3;

% Simulation
[~,~,g_2D_WID_SNM_320,x_2D_WID_SNM_320,m_2D_WID_SNM_320] = nll_M1_2D_WID_SNM(parameters_2D_WID_SNM{1,s_320}.MS.par(:,1),info_2D_WID_SNM{1,s_320});
[~,~,g_2D_FID_SNM_320,x_2D_FID_SNM_320,m_2D_FID_SNM_320] = nll_M1_2D_FID_SNM(parameters_2D_FID_SNM{1,s_320}.MS.par(:,1),info_2D_FID_SNM{1,s_320});
[~,~,g_2D_WID_INM_320,x_2D_WID_INM_320,m_2D_WID_INM_320] = nll_M1_2D_WID_INM(parameters_2D_WID_INM{1,s_320}.MS.par(:,1),info_2D_WID_INM{1,s_320});

%% VISUALIZATION OF SIMULATION AND ERROR (FIGURE 5A & B)
% Colors
load('./Auxillary/mycmaps.mat')
Col = [0.0 0.0 0.0;
       0.0 0.6 0.0;
       0.0 0.0 0.8];

% Open figure
figure('name','Figure 5A&B: spatial error and residual distribution');

% Spatial error pattern for "direct approach"
axes('Position',[0.05,0.73,0.38,0.24]);
imagesc(info_2D_WID_SNM{1,s_320}.data{1}-g_2D_WID_SNM_320{1},1e-2*[-1,1]); hold on;
contour(info_2D_WID_SNM{1,s_320}.mask{1},[1 1],'Color','k','linewidth',1);
colormap(cmap_redblue_error_NaN); freezeColors;
set(gca,'XTickLabel','','YTickLabel',''); axis square;

% Spatial error pattern and filter for "filtering approach"
axes('Position',[0.05,0.46,0.38,0.24]);
err_2D_FID_SNM = info_2D_FID_SNM{1,s_320}.filtered_data{1}-g_2D_FID_SNM_320{1};
err_2D_FID_SNM(isnan(err_2D_FID_SNM)) = -1e-2;
imagesc(err_2D_FID_SNM,1e-2*[-1,1]); hold on;
contour(info_2D_FID_SNM{1,s_320}.mask{1},[1 1],'Color','k','linewidth',1);
colormap([Col(:,2)';cmap_redblue_error_NaN(2:end,:)]);  freezeColors;
set(gca,'XTickLabel','','YTickLabel',''); axis square;

% Spatial error pattern for "integrated modelling approach"
axes('Position',[0.05,0.19,0.38,0.24]);
imagesc(info_2D_WID_INM{1,s_320}.data{1}-g_2D_WID_INM_320{1},1e-2*[-1,1]); hold on;
contour(info_2D_WID_INM{1,s_320}.mask{1},[1 1],'Color','k','linewidth',1);
colormap(cmap_redblue_error_NaN); freezeColors;
set(gca,'XTickLabel','','YTickLabel',''); axis square;

% Colorbar for spatial error pattern
hcol = colorbar('Location','SouthOutside',...
                'YTick',linspace(-1e-2,1e-2,5),...
                'YTickLabel',{'<-0.01','-0.005','0','0.005','>0.01'});
set(hcol,'Position', [0.06 0.13 0.36 0.03]); % shift colorbar
set(get(hcol,'YLabel'),'String','error, e=y^m-y(\theta)')

% Histograms of rediuals for "direct approach"
x_bin = linspace(-1,2,50);
axes('Position',[0.61,0.73,0.38,0.24]);
histogram(log(info_2D_WID_SNM{1,s_320}.data{1}(:))-log(g_2D_WID_SNM_320{1}(:)),x_bin,...
    'Normalization','pdf','FaceColor',0.9*[1,1,1]); hold on;
plot(x_bin,pdf('normal',x_bin,0,10.^(parameters_2D_WID_SNM{1,s_320}.MS.par(6,1))),'-k');
set(gca,'xlim',x_bin([1,end]),'xticklabel','','ylim',[0,3],'ytick',[0:3]);

% Histograms of rediuals for "filtering approach"
axes('Position',[0.61,0.46,0.38,0.24]);
res_2D_FID_SNM = log(info_2D_FID_SNM{1,s_320}.data{1})-log(g_2D_FID_SNM_320{1});
c1 = hist(res_2D_FID_SNM(~isnan(info_2D_FID_SNM{1,s_320}.filtered_data{1})),x_bin);
c2 = hist(res_2D_FID_SNM( isnan(info_2D_FID_SNM{1,s_320}.filtered_data{1})),x_bin);
bar(x_bin,c1./trapz(x_bin,c1+c2),1,'FaceColor',0.9*[1,1,1]); hold on;
bar(x_bin,c2./trapz(x_bin,c1+c2),1,'FaceColor',Col(:,2)); 
plot(x_bin,pdf('normal',x_bin,0,10.^(parameters_2D_FID_SNM{1,s_320}.MS.par(6,1))),'-k');
set(gca,'xlim',x_bin([1,end]),'xticklabel','','ylim',[0,3],'ytick',[0:3]);

% Histograms of rediuals for "integrated modelling approach"
axes('Position',[0.61,0.19,0.38,0.24]);
histogram(log(info_2D_WID_INM{1,s_320}.data{1}(:))-log(g_2D_WID_INM_320{1}(:)),x_bin,...
    'Normalization','pdf','FaceColor',0.9*[1,1,1]); hold on;
sig   = 10.^(parameters_2D_WID_INM{1,s_320}.MS.par(6,1));
sig_o = 10.^(parameters_2D_WID_INM{1,s_320}.MS.par(7,1));
mu_o  = 10.^(parameters_2D_WID_INM{1,s_320}.MS.par(8,1));
w     = 10.^(parameters_2D_WID_INM{1,s_320}.MS.par(9,1));
plot(x_bin,(1-w)*pdf('normal',x_bin,mu_o,sig_o) + w*pdf('normal',x_bin,0,sig),'-k');
plot(x_bin,(1-w)*pdf('normal',x_bin,mu_o,sig_o),'-','Color',Col(:,3));
set(gca,'xlim',x_bin([1,end]),'xtick',[-1:2],'ylim',[0,3],'ytick',[0:3]);
xlabel('residual');

%% EVALUATION OF ESTIMATION ERRORS
% True parameters
true_par = [8.5e+1;   % D
            2.4e-1;   % a
            2.1e-1;   % S0
            1.3e-2;   % off
            1.44e-1]; % sig 

% Estimated log-parameters
for r = 1:length(replicate_nr)
    for s = 1:length(spot_nr)
        MLE_2D_WID_SNM(:,r,s) = parameters_2D_WID_SNM{r,s}.MS.par(:,1); 
        MLE_2D_FID_SNM(:,r,s) = parameters_2D_FID_SNM{r,s}.MS.par(:,1); 
        MLE_2D_WID_INM(:,r,s) = parameters_2D_WID_INM{r,s}.MS.par(:,1); 
    end
end

% Estimation error
ERR_2D_WID_SNM = MLE_2D_WID_SNM([1,2,3,4,6],:,:) - log10(true_par);
ERR_2D_FID_SNM = MLE_2D_FID_SNM([1,2,3,4,6],:,:) - log10(true_par);
ERR_2D_WID_INM = MLE_2D_WID_INM([1,2,3,4,6],:,:) - log10(true_par);

%% VISUALIZATION OF ESTIMATION ERROR (FIGURE 5C)
% Open figure
figure('name','Figure 5C: estimation error');

% Axis position
AxesPos = [0.12, 0.73, 0.38, 0.24;  % D/g
           0.12, 0.46, 0.38, 0.24;  % a/gK_D
           0.55, 0.46, 0.38, 0.24;  % S_0
           0.12, 0.19, 0.38, 0.24;  % b
           0.55, 0.19, 0.38, 0.24]; % sigma
       
% Color
ColMean = [0.0 0.0 0.0;
           0.0 0.6 0.0;
           0.0 0.0 0.8];
Col = [0.7 0.7 0.7;
       0.0 0.6 0.0;
       0.0 0.0 0.8];
edgeCol = Col+(1-Col).*0.55;
facealpha = 0.25;

% Labels
tit = {'D/\gamma','(\alpha k_{1})/(\gamma k_{-1})','S_0','b','\sigma^2'};

% Loop: Parameters
for i = 1:5
    % Subplot
    axes('Position',AxesPos(i,:));

    % Mean and standard deviation of estimation errors
    m_diff   = [squeeze(mean(ERR_2D_WID_SNM(i,:,:),2))';
                squeeze(mean(ERR_2D_FID_SNM(i,:,:),2))';
                squeeze(mean(ERR_2D_WID_INM(i,:,:),2))'];
    std_diff = [squeeze(std(ERR_2D_WID_SNM(i,:,:),[],2))';
                squeeze(std(ERR_2D_FID_SNM(i,:,:),[],2))';
                squeeze(std(ERR_2D_WID_INM(i,:,:),[],2))'];

    % Position vector
    X = [1:length(spot_nr),fliplr(1:length(spot_nr))];
    Y = [m_diff-2*std_diff,fliplr(m_diff+2*std_diff)];

    % Loop: Methods
    for j = 1:3
        hplo(  j) = plot(1:length(spot_nr),m_diff(j,:),'-','color',0.7*ColMean(j,:)); hold on;
        hplo(3+j) = plot(1:length(spot_nr),m_diff(j,:),'.','color',0.7*ColMean(j,:),'Markersize',10);
        hplo(6+j) = patch(X,Y(j,:),Col(j,:),'facecolor',Col(j,:),'edgecolor',edgeCol(j,:),'facealpha',facealpha);
    end
    
    % Axis and box
    box on;
    axis([1 length(spot_nr) -0.5 0.5])

    % Ticks
    switch i
        case {1,2,3}
            set(gca,'xtick',1:length(spot_nr),'xticklabel',{})
        case {4,5}
            set(gca,'xtick',1:length(spot_nr),'xticklabel',spot_nr) 
    end
    switch i
        case {1,2,4}
            set(gca,'ylim',[-0.5 0.5],'ytick',[-0.4:0.2:0.4]);
        case {3,5}
            set(gca,'ylim',[-0.5 0.5],'ytick',[-0.4:0.2:0.4],'yticklabel',[]);
    end
    
    % Label
    if i == 2
        ylabel('estimation error, log(\theta^{true}) - log(\theta^{ML})');
    end
    if (i == 4) || (i == 5)
        xlabel('number of spots');
    end
    
    % Title
    text((length(spot_nr)+1)/2,0.4,tit{i},'HorizontalAlignment','center');
end

% Legend
hleg = legend(hplo([1:4,7]),{'direct approach','filtering approach',['integrated modelling' char(10) 'approach'],'mean','95% percentile interval'});
set(hleg,'Position', [0.78,0.85,0,0],'box','off');
```
