## Supplementary material for "Mechanistic description of spatial processes using integrative modelling of noise-corrupted imaging data": getResultsForFigure6.m

```
% Analysis for Figure 6:
% Parameter estimation results and residual distributions for 2D models 
% with constant heparan sulfate concentration using whole imaging data and 
% integrated noise model.
%
% Note: The calculations take roughly one hours.

clear all;
close all;
clc;

%% DATASET
% Type of experimental data
flags.data_type = 'real';

% Index set of images
exp_nr = 2;

%% OPTIMIZATION
% Number of local optimization runs
n_starts = 10;

% Multi-start optimization
[parameters,~,info,options] = runParameterEstimation('M1_2D_WID_INM',flags,n_starts,exp_nr,[],[],[]);

% Maximual likelihood estimates of parameter values
sig_2   = 10.^parameters.MS.par(6,1);
sig_o_2 = 10.^parameters.MS.par(7,1);
mu_2    = 10.^parameters.MS.par(8,1);
w_2     = 10.^parameters.MS.par(9,1);

%% SIMULATION OF MODEL
[~,~,g,x,m] = nll_M1_2D_WID_INM(parameters.MS.par(:,1),info);

%% VISUALIZATION
% Colors
load('./Auxillary/mycmaps.mat')
Col = [0.20 0.20 0.44;
       0.44 0.68 0.76;
       0.76 0.28 0.28];
   
% Figure 6A: Lymphoid vessels
figure('name','Figure 6A: Lymphoid vessels');
Col_map_LV = [0.9908 0.9936 0.6322;
              0.8018 0.9197 0.7392;
              0.6616 0.8361 0.9687;
              0.6969 0.6602 0.8641];
S = + 1.00.*info.disc{1}.Sj(:,:,1) ...
    + 0.66.*info.disc{1}.Sj(:,:,2) ...
    + 0.33.*info.disc{1}.Sj(:,:,3);
surf(S(info.dist{1}.ind1(end:-1:1),info.dist{1}.ind2)); hold on; shading interp; view(0,90);
contour(S(info.dist{1}.ind1(end:-1:1),info.dist{1}.ind2),0.01*[1 1],'Color',[0 0 0],'Linewidth',1);
colormap(Col_map_LV);
set(gca,'xtick',[],'ytick',[]);
set(gca,'xlim',[-0.5,402],'ylim',[-0.5,402.5],'box','on','xtick',[],'ytick',[]);
axis on; box on; axis square

% Figure 6B: Experimental data
figure('name','Figure 6B: Experimental data');
cmap_green_data_NaN_reduced = cmap_green_data_NaN([1,10,22,26,30,33,36,38,42,46],:);
c_min = log10(0.01); c_max = log10(0.1);
imagesc(log10(info.data{1}),[c_min,c_max]); hold on;
colormap(cmap_green_data_NaN_reduced); 
contour(info.mask{1},[1 1],'Color',[1,1,1],'Linewidth',1.5)  
set(gca,'xlim',[-0.5,402],'ylim',[-0.5,402.5],'box','on','xtick',[],'ytick',[]);
axis square;

% Figure 6C: Simulation
figure('name','Figure 6C: Simulation of model with constant heparan sulfate concentration');
cmap_green_data_NaN_reduced = cmap_green_data_NaN([1,10,22,26,30,33,36,38,42,46],:);
c_min = log10(0.01); c_max = log10(0.1);
imagesc(log10(g{1}),[c_min,c_max]); hold on;
colormap(cmap_green_data_NaN_reduced); 
contour(info.mask{1},[1 1],'Color',[1,1,1],'Linewidth',1.5)  
set(gca,'xlim',[-0.5,402],'ylim',[-0.5,402.5],'box','on','xtick',[],'ytick',[]);
axis square;
hcol = colorbar('Location','EastOutside','YTick',log10([0.01,0.02,0.03,0.04,0.05,0.06,0.07,0.08,0.09,0.1]),'YTickLabel',{'<0.01','  0.02','','  0.04','','  0.06','','  0.08','','>0.10'});
hcol.Label.String = 'fluorescence intensity [a.u.]';

% Figure 6D: Residual
figure('name','Figure 6D: Residual for model with constant heparan sulfate concentration');
c_min = -1; c_max = 2;
imagesc((log10(info.data{1})-log10(g{1}))/sig_2,[c_min c_max]); hold on;
colormap(cmap_redblue_error([1:2:32,32,32,32,32,32:end],:))
contour(info.mask{1},[1 1],'Color','k','linewidth',2);
set(gca,'xlim',[-0.5,402],'ylim',[-0.5,402.5],'box','on','xtick',[],'ytick',[]);
axis square;
hcol = colorbar('Location','EastOutside','YTick',[-1.0,-0.5,0,0.5,1.0,1.5,2.0],'YTickLabel',{'<-1.0','-0.5','0','0.5','1.0','1.5','>2.0'});
hcol.Label.String = 'residual [a.u.]';

% Figure 6E: Residual distribution
figure('name','Figure 6E: Residual distribution');
cmap_redblue_error_1 = cmap_redblue_error([1:2:32,32,32,32,32,32:end],:);
cmap_redblue_error_reduced = [repmat(cmap_redblue_error(1,:),6,1);
                              cmap_redblue_error_1(1:2:end,:);
                              repmat(cmap_redblue_error(end,:),12,1)];    
r = (log10(info.data{1}(:))-log10(g{1}(:)))/sig_2;
r_grid = linspace(-1.5,3.5,size(cmap_redblue_error_reduced,1));
dr = r_grid(2) - r_grid(1);
for k = 1:(length(r_grid)-1)
    n_grid(k) = sum((r_grid(k) <= r).*(r < r_grid(k+1)))/length(r)/dr;
    fill(r_grid([k,k+1,k+1,k]),n_grid(k)*[0,0,1,1],'k','facecolor',cmap_redblue_error_reduced(k,:)); hold on;
end
r_grid2 = linspace(-3,3,200);
lh(2) = plot(r_grid2,   w_2 *pdf('norm',r_grid2,0   ,sig_2  ),'-','linewidth',1.5,'color',[1.0,0.0,0.5]);
lh(3) = plot(r_grid2,(1-w_2)*pdf('norm',r_grid2,mu_2,sig_o_2),'-','linewidth',1.5,'color',[0.5,0.0,1.0]);
lh(1) = plot(r_grid2,   w_2 *pdf('norm',r_grid2,0   ,sig_2  ) ...
                   + (1-w_2)*pdf('norm',r_grid2,mu_2,sig_o_2),'-','linewidth',1.5,'color',[0.0,0.0,0.0]);
set(gca,'xlim',[-1.5,3.5],'ylim',[0,1.5]);
xlabel('residual [a.u.]');
ylabel('density [-]');
l_h = legend(lh,{'overall distribution','noise distribution','outlier distribution'},'location','NorthEast','box','off');
set(l_h,'position',[0.6,0.8,0.01,0.05]);
```
