## Supplementary material for "Mechanistic description of spatial processes using integrative modelling of noise-corrupted imaging data": getResultsForFigure7.m

```
% Analysis for Figure 7:
% Model selection of 2D models with
% - unifrom heparan sulfate concentrations (model 1),
% - different heparan sulfate concentrations in lymphoid vessels and
%   tissue (model 2), and
% - different heparan sulfate concentrations in individual vessel and
%   tissue (model 3).
% For the statistical comparison the integrated noise model is employed.
%
% Note: The calculations take several days.

clear all;
close all;
clc;

%% DATASET
% Type of experimental data
flags.data_type = 'real';

% Index set of images
exp_nr = [1,2,3,4,12,13,14,15,16];

%% OPTIMIZATION AND PROFILE CALCULATION OPTIONS
% Number of local optimization runs
n_starts_all = 10;       % Note: In the manuscript, we used 100 local optimizations.
n_starts_individual = 3; % Note: In the manuscript, we used 10 local optimizations.

%% (SIMULTANEOUS) PARAMETER ESTIMATION FOR ALL IMAGES
[parameters_M1_all,~,info_M1_all,options_M1_all] = runParameterEstimation('M1_2D_WID_INM',flags,n_starts_all,exp_nr,[],[]);
[parameters_M2_all,~,info_M2_all,options_M2_all] = runParameterEstimation('M2_2D_WID_INM',flags,n_starts_all,exp_nr,[],[]);
[parameters_M3_all,~,info_M3_all,options_M3_all] = runParameterEstimation('M3_2D_WID_INM',flags,n_starts_all,exp_nr,[],[]);

%% PARAMETER ESTIMATION FOR INDIVIDUAL IMAGES
for i = 1:length(exp_nr)
    disp(['Image: ' num2str(exp_nr(i)) ' (' num2str(i) ' of ' num2str(length(exp_nr)) ')']);
    [parameters_M1_individual{i},~,info_M1_individual{i},options_M1_individual{i}] = runParameterEstimation('M1_2D_WID_INM',flags,n_starts_individual,exp_nr(i),[],[]);
    [parameters_M2_individual{i},~,info_M2_individual{i},options_M2_individual{i}] = runParameterEstimation('M2_2D_WID_INM',flags,n_starts_individual,exp_nr(i),[],[]);
    [parameters_M3_individual{i},~,info_M3_individual{i},options_M3_individual{i}] = runParameterEstimation('M3_2D_WID_INM',flags,n_starts_individual,exp_nr(i),[],[]);
    close all;
end

%% SIMULATION OF MODEL FOR EXPERIMENTAL IMAGE 2
i_exp2 = find(exp_nr == 2);

fh = figure;
[~,~,g1,x1,m1] = nll_M1_2D_WID_INM(parameters_M1_individual{i_exp2}.MS.par(:,1),setfield(info_M1_individual{i_exp2},'fh',fh));
[~,~,g2,x2,m2] = nll_M2_2D_WID_INM(parameters_M2_individual{i_exp2}.MS.par(:,1),setfield(info_M2_individual{i_exp2},'fh',fh));
[~,~,g3,x3,m3] = nll_M3_2D_WID_INM(parameters_M3_individual{i_exp2}.MS.par(:,1),setfield(info_M3_individual{i_exp2},'fh',fh));

%% CALCULATION OF BIC VALUES
% For all images
parameters_M1_all.BIC = -2*parameters_M1_all.MS.logPost(1) + parameters_M1_all.number*log(length(exp_nr)*prod(size(info_M1_all.data{1})));
parameters_M2_all.BIC = -2*parameters_M2_all.MS.logPost(1) + parameters_M2_all.number*log(length(exp_nr)*prod(size(info_M2_all.data{1})));
parameters_M3_all.BIC = -2*parameters_M3_all.MS.logPost(1) + parameters_M3_all.number*log(length(exp_nr)*prod(size(info_M3_all.data{1})));

% For individual images
for i = 1:length(exp_nr)
    parameters_M1_individual{i}.BIC = -2*parameters_M1_individual{i}.MS.logPost(1) + parameters_M1_individual{i}.number*log(prod(size(info_M1_individual{i}.data{1})));
    parameters_M2_individual{i}.BIC = -2*parameters_M2_individual{i}.MS.logPost(1) + parameters_M2_individual{i}.number*log(prod(size(info_M2_individual{i}.data{1})));
    parameters_M3_individual{i}.BIC = -2*parameters_M3_individual{i}.MS.logPost(1) + parameters_M3_individual{i}.number*log(prod(size(info_M3_individual{i}.data{1})));
end

% Collection of all BIC values in a single matrix
BIC = nan(3,length(exp_nr)+1);
for i = 1:length(exp_nr)
    BIC(1,i) = parameters_M1_individual{i}.BIC;
    BIC(2,i) = parameters_M2_individual{i}.BIC;
    BIC(3,i) = parameters_M3_individual{i}.BIC;
end
BIC(1,end) = parameters_M1_all.BIC;
BIC(2,end) = parameters_M2_all.BIC;
BIC(3,end) = parameters_M3_all.BIC;

%% VISUALIZATION
% Colors
load('./Auxillary/mycmaps.mat')
% Colors for models
Col = [0.20 0.20 0.44;
       0.44 0.68 0.76;
       0.76 0.28 0.28];

% Colors for sugar concentrations
Col_Sugar = [0.9908 0.9936 0.6322;
             0.8018 0.9197 0.7392;
             0.6616 0.8361 0.9687;
             0.6969 0.6602 0.8641;
             0.8977 0.8598 0.7197];

% Figure 7A (left): Sugar concentration for model 1:
%   unifrom heparan sulfate concentrations
figure('name','Figure 7A (left): Sugar concentration for model 1');
S = + 1.00.*info_M1_all.disc{2}.Sj(:,:,1) ...
    + 0.66.*info_M1_all.disc{2}.Sj(:,:,2) ...
    + 0.33.*info_M1_all.disc{2}.Sj(:,:,3);
surf(S(info_M1_all.dist{2}.ind1(end:-1:1),info_M1_all.dist{2}.ind2)); hold on; shading interp; view(0,90);
contour(S(info_M1_all.dist{2}.ind1(end:-1:1),info_M1_all.dist{2}.ind2),0.01*[1 1],'Color',[0 0 0],'Linewidth',1);
colormap(Col_Sugar(1,:));
set(gca,'xtick',[],'ytick',[]);
set(gca,'xlim',[-0.5,402],'ylim',[-0.5,402.5],'box','on','xtick',[],'ytick',[]);
axis on; box on; axis square

% Figure 7A (middle): Sugar concentration for model 2:
%   different heparan sulfate concentrations in lymphoid vessels and tissue
figure('name','Figure 7A (middle): Sugar concentration for model 2');
surf(S(info_M2_all.dist{2}.ind1(end:-1:1),info_M2_all.dist{2}.ind2)); hold on; shading interp; view(0,90);
contour(S(info_M2_all.dist{2}.ind1(end:-1:1),info_M2_all.dist{2}.ind2),0.01*[1 1],'Color',[0 0 0],'Linewidth',1);
colormap(Col_Sugar([1,5,5,5],:));
set(gca,'xtick',[],'ytick',[]);
set(gca,'xlim',[-0.5,402],'ylim',[-0.5,402.5],'box','on','xtick',[],'ytick',[]);
axis on; box on; axis square

% Figure 7A (right): Sugar concentration for model 3:
%   different heparan sulfate concentrations in individual vessel and tissue
figure('name','Figure 7A (right): Sugar concentration for model 3');
surf(S(info_M3_all.dist{2}.ind1(end:-1:1),info_M3_all.dist{2}.ind2)); hold on; shading interp; view(0,90);
contour(S(info_M3_all.dist{2}.ind1(end:-1:1),info_M3_all.dist{2}.ind2),0.01*[1 1],'Color',[0 0 0],'Linewidth',1);
colormap(Col_Sugar(1:4,:));
set(gca,'xtick',[],'ytick',[]);
set(gca,'xlim',[-0.5,402],'ylim',[-0.5,402.5],'box','on','xtick',[],'ytick',[]);
axis on; box on; axis square

% Figure 7B: Multi-start optimization results for simulateneous
%   estimation of all images
figure('name','Figure 7B: Multi-start optimization results');
plot(1:n_starts_all,parameters_M1_all.MS.logPost,'.','color',Col(1,:),'markersize',15); hold on;
plot(1:n_starts_all,parameters_M2_all.MS.logPost,'.','color',Col(2,:),'markersize',15);
plot(1:n_starts_all,parameters_M3_all.MS.logPost,'.','color',Col(3,:),'markersize',15);
plot(1:n_starts_all,parameters_M1_all.MS.logPost,'-','color',Col(1,:));
plot(1:n_starts_all,parameters_M2_all.MS.logPost,'-','color',Col(2,:));
plot(1:n_starts_all,parameters_M3_all.MS.logPost,'-','color',Col(3,:));
ylabel('log-likelihood');
xlabel('sorted optimizer runs');
set(gca,'xlim',[0,n_starts_all+1]);
legend({'model 1','model 2','model 3'},'box','off','Location','southwest');

% Figure 7C: Model selection results
figure('name','Figure 7C: Model selection results');
%    Model 1 vs. Model 2
ha(1) = axes('Position',[0.20,0.05,0.25,0.70]);
imagesc((BIC(1,:)-BIC(2,:))',[-1e3,1e3]); colormap(ha(1),cmap_bluecyan_BIC);
hcol1 = colorbar('Location','North','YTick',[-0.7e3 0.7e3],'YTickLabel',{'\color[rgb]{0.57,0.57,0.87} model 1','\color[rgb]{0.53,0.83,0.93} model 2'});
set(hcol1,'Position',[0.2+0.25/2-0.18/2,0.8,0.18,0.05]); hcol1.Label.String = 'select';
text(ones(10,1),1:10,cellstr(num2str((BIC(1,:)-BIC(2,:))','%1.2f')),'color','k','HorizontalAlignment','center','VerticalAlignment','middle');
%    Model 1 vs. Model 3
ha(2) = axes('Position',[0.45,0.05,0.25,0.70]);
imagesc((BIC(1,:)-BIC(3,:))',[-1e3,1e3]); colormap(ha(2),cmap_bluered_BIC);
hcol2 = colorbar('Location','North','YTick',[-0.7e3 0.7e3],'YTickLabel',{'\color[rgb]{0.57,0.57,0.87} model 1','\color[rgb]{1,0.47,0.47} model 3'});
set(hcol2,'Position',[0.45+0.25/2-0.18/2,0.8,0.18,0.05]); hcol2.Label.String = 'select';
text(ones(10,1),1:10,cellstr(num2str((BIC(1,:)-BIC(3,:))','%1.2f')),'color','k','HorizontalAlignment','center','VerticalAlignment','middle');
%    Model 2 vs. Model 3
ha(3) = axes('Position',[0.70,0.05,0.25,0.70]);
imagesc((BIC(2,:)-BIC(3,:))',[-1e3,1e3]); colormap(ha(3),cmap_cyanred_BIC);
hcol3 = colorbar('Location','North','YTick',[-0.7e3 0.7e3],'YTickLabel',{'\color[rgb]{0.53,0.83,0.93} model 2','\color[rgb]{1,0.47,0.47} model 3'});
set(hcol3,'Position',[0.7+0.25/2-0.18/2,0.8,0.18,0.05]); hcol3.Label.String = 'select';
text(ones(10,1),1:10,cellstr(num2str((BIC(2,:)-BIC(3,:))','%1.2f')),'color','k','HorizontalAlignment','center','VerticalAlignment','middle');
%    Labeling
set(ha(1),'yTick',1:10,'YTicklabel',{' image 1',' image 2',' image 3',' image 4',' image 12','image 13','image 14','image 15','image 16','all'})
set(ha([2,3]),'YTickLabel','')
set(ha,'xTick',[],'xTickLabel','');

% Figure 7D (left): Experimental data
figure('name','Figure 7D (left): Experimental data');
cmap_green_data_NaN_reduced = cmap_green_data_NaN([1,10,22,26,30,33,36,38,42,46],:);
c_min = log10(0.01); c_max = log10(0.1);
imagesc(log10(info_M3_individual{i_exp2}.data{1}),[c_min,c_max]); hold on;
colormap(cmap_green_data_NaN_reduced); 
contour(info_M3_individual{i_exp2}.mask{1},[1 1],'Color',[1,1,1],'Linewidth',1.5)  
set(gca,'xlim',[-0.5,402],'ylim',[-0.5,402.5],'box','on','xtick',[],'ytick',[]);
axis square;

% Figure 7D (right): Simulation
figure('name','Figure 7D (right): Simulation of model 3');
cmap_green_data_NaN_reduced = cmap_green_data_NaN([1,10,22,26,30,33,36,38,42,46],:);
c_min = log10(0.01); c_max = log10(0.1);
imagesc(log10(g3{1}),[c_min,c_max]); hold on;
colormap(cmap_green_data_NaN_reduced); 
contour(info_M3_individual{i_exp2}.mask{1},[1 1],'Color',[1,1,1],'Linewidth',1.5)  
set(gca,'xlim',[-0.5,402],'ylim',[-0.5,402.5],'box','on','xtick',[],'ytick',[]);
axis square;
hcol = colorbar('Location','EastOutside','YTick',log10([0.01,0.02,0.03,0.04,0.05,0.06,0.07,0.08,0.09,0.1]),'YTickLabel',{'<0.01','  0.02','','  0.04','','  0.06','','  0.08','','>0.10'});
hcol.Label.String = 'fluorescence intensity [a.u.]';

% Figure 7E: Distance-dependent fluorescence intensity
figure('name','Figure 7E: Distance-dependent fluorescence intensity for all models');
plot(info_M1_individual{i_exp2}.filtered_dist_dep_mean{1}.dist,...
     info_M1_individual{i_exp2}.filtered_dist_dep_mean{1}.mean,'o-k','markersize',3); hold on;
plot(x1{1},m1{1},'-s','Color',[0.0,0.0,0.3],'markersize',3);
plot(x2{1},m2{1},'-d','Color',[0.3,0.6,0.7],'markersize',3);
plot(x3{1},m3{1},'-v','Color',[0.7,0.1,0.1],'markersize',3);

set(gca,'xlim',[0,64],'ylim',[0.01,0.03]);
xlabel({'distance to nearest','lymphoid vessel [\mum]'});
ylabel('fluorescence intensity [a.u.]');

l_h = legend({'data', 'model 1','model 2','model 3'},'location','NorthEast','box','off');
```
