## Supplementary material for "Mechanistic description of spatial processes using integrative modelling of noise-corrupted imaging data": getStarted.m

```
% Illustration of analysis pipeline

clear all;
close all;
clc;

%% MODEL
% (i)  measurement data
%      - whole imaging data (WID)
%      - filtered imaging data (FID)
%      - distance-dependent mean (DDM)
% (ii) noise model
%      - standard noise model (SNM)
%      - integrated noise model (INM)

% model_name = 'M1_1D_DDM_SNM'; % remark: opt. checked!
% model_name = 'M1_2D_DDM_SNM'; % remark: opt. checked!
% model_name = 'M1_2D_WID_SNM'; % remark: opt. checked!
% model_name = 'M1_2D_FID_SNM'; % remark: opt. checked!
model_name = 'M1_2D_WID_INM'; % remark: opt. checked!
% model_name = 'M2_2D_WID_INM'; % remark: opt. checked!
% model_name = 'M3_2D_WID_INM'; % remark: opt. checked!

%% DATASET
% Type of experimental data
flags.data_type = 'real';       % measured data
% flags.data_type = 'artificial'; % simulated data

% Index set of images:
% (a subset of {1,2,3,4,12,13,14,15,16}.)
exp_nr = [1];
%exp_nr = [1:3];
%exp_nr = [1:4,12:16];

% For artifical data:
% - number of spots (a subset of {0,20,40,80,160,320,640})
spot_nr = 320;
% - replicate number (a number between 1 and 30)
replicate_nr = 1;

%% MULTI-START OPTIMIZATION OPTIONS
% Number of local optimization runs
n_starts = 1;

%% PARAMETER ESTIMATION
[parameters,info,options,fh] = runParameterEstimation(model_name,flags,n_starts,exp_nr,spot_nr,replicate_nr,[]);
```
