## Supplementary material for "Mechanistic description of spatial processes using integrative modelling of noise-corrupted imaging data": nll_M1_2D_DDM_SNM.m

```
% nll_M1_2D_DDM_SNM() computes the negative log-likelihood and its 
% gradient for: 
% * Model M1 (2D PDE model with uniform sugar concentrations)
% * Distance-Dependent Mean data (DDM)
% * Standard Noise Model (SNM)
%
% USAGE:
% * [J] = nll_M1_2D_DDM_SNM(p,info)
% * [J,G] = nll_M1_2D_DDM_SNM(p,info)
%
% Parameters:
%   p: log-parameter values in the order
%     (D,a,S0,b,s_1,..,s_N)
%   info: structure needed to evaluate the Likelihood
%     .exp_nr: index of considered experimental data
%     .name: parameter names
%     .data: experimental data for sugar-bound CCL21
%     .disc: struct containing the PDE solver information
%     .dist_dep_mean: distance-dependent mean intensity of raw images
%     .filtered_dist_dep_mean: distance-dependent mean intensity of filtered images
%     .plot: flag indicating if results are visualized (default: false)
%
% Return values:
%   J: negative log-Likelihood
%   G: gradient
%
% History:
% * 2014/05/09 Sabrina Hross
% * 2017/10/30 Jan Hasenauer

function [varargout] = nll_M1_2D_DDM_SNM(varargin)

%% CHECK AND ASSIGN INPUTS
% Parameter vector
if nargin <= 2
    p = varargin{1};
else
    error('nll_M1_FID_SNM requires two input.');
end

% Options
info.plot = false;
if nargin == 2
    info = setdefault(varargin{2},info);
end

%% ASSIGNMENT OF DATA AND PARAMETER PROPERTIES
% Filtered experimental data
Y = info.filtered_data;
n_e = length(info.exp_nr);

% Parameters
n_p = length(p);
for i = 1:n_p
    parameter.(info.name{i}) = 10^p(i);
end

%% SIMULATION OF MODEL
% Loop: Experimental conditions
for i = 1:n_e
    % Free CCL21 and sensitivities
    v = (info.disc{i}.K.*parameter.D+info.disc{i}.M)\info.disc{i}.F.*parameter.a;
    if nargout >= 2
        vS = (info.disc{i}.K.*parameter.D+info.disc{i}.M)\[-info.disc{i}.K*v,info.disc{i}.F];
    end
    
    % Sugar-bound CCL21 and sensitivities
    y = tri2grid(info.disc{i}.p,info.disc{i}.t,v,info.disc{i}.tn,info.disc{i}.a2,info.disc{i}.a3)';
    if nargout >= 2
        yS_i(:,:,1) = tri2grid(info.disc{i}.p,info.disc{i}.t,vS(:,1),info.disc{i}.tn,info.disc{i}.a2,info.disc{i}.a3)';
        yS_i(:,:,2) = tri2grid(info.disc{i}.p,info.disc{i}.t,vS(:,2),info.disc{i}.tn,info.disc{i}.a2,info.disc{i}.a3)';
    end
    
    % Observable and sensitivities
    g{i} = parameter.(['s_',num2str(info.exp_nr(i))]).*(parameter.b + (parameter.S0.*y(info.dist{i}.ind1,info.dist{i}.ind2))./(1+y(info.dist{i}.ind1,info.dist{i}.ind2)));
    if min(g{i}(:)) <= 0
        error('Evaluation of objective function failed as solution of PDE is negative.')
    end
    if nargout >= 2
        dgdk{i} = zeros(size(g{i},1),size(g{i},2),n_p);
        for j = [1,2,3,4,4+i]
            switch j
                case 1 % D
                    dgdk{i}(:,:,1) = parameter.(['s_',num2str(info.exp_nr(i))]).*(parameter.S0./(1+y(info.dist{i}.ind1,info.dist{i}.ind2)).^2).*yS_i(info.dist{i}.ind1,info.dist{i}.ind2,1);
                case 2 % a
                    dgdk{i}(:,:,2) = parameter.(['s_',num2str(info.exp_nr(i))]).*(parameter.S0./(1+y(info.dist{i}.ind1,info.dist{i}.ind2)).^2).*yS_i(info.dist{i}.ind1,info.dist{i}.ind2,2);
                case 3 % S0
                    dgdk{i}(:,:,3) = (parameter.(['s_',num2str(info.exp_nr(i))]).*y(info.dist{i}.ind1,info.dist{i}.ind2))./(1+y(info.dist{i}.ind1,info.dist{i}.ind2));
                case 4 % b
                    dgdk{i}(:,:,4) = parameter.(['s_',num2str(info.exp_nr(i))]).*ones(size(y(info.dist{i}.ind1,info.dist{i}.ind2)));
                case 4+i % s_i
                    dgdk{i}(:,:,4+i) = parameter.b + (parameter.S0.*y(info.dist{i}.ind1,info.dist{i}.ind2))./(1+y(info.dist{i}.ind1,info.dist{i}.ind2));
            end
            % Chain rule for log-transformed parameters
            dgdk{i}(:,:,j) = dgdk{i}(:,:,j)*parameter.(info.name{j})*log(10);
        end
    end
    
    % Evaluation of distance-dependent mean intensity
    [x{i},m{i},~] = getDistanceDependentMean(info.dist{i}.dist_from_lv,g{i},info.dist{i}.edges,info.dist{i}.I);
    if nargout >= 2
        dmdk{i} = zeros(length(x{i}),n_p);
        for j = [1,2,3,4,4+i]
            [~,dmdk{i}(:,j),~] = getDistanceDependentMean(info.dist{i}.dist_from_lv,dgdk{i}(:,:,j),info.dist{i}.edges,info.dist{i}.I);
        end
    end
end

%% PLOT RESULTS
if info.plot
    % Open figure
    figure(info.fh);

    % Parameters
    subplot(1+n_e,6,1:6); hold off;
    plot(1:n_p,p,'-ro','linewidth',2); hold on;
    plot(1:n_p,info.parameters.min,'--k');
    plot(1:n_p,info.parameters.max,'--k');
    legend('parameter values','lower bound','upper bound');
    set(gca,'XTick',1:n_p,'XLim',[1,n_p],'XTickLabel',info.name,'XTickLabelRotation',90);

    % Loop: experimental conditions
    for i = 1:n_e
        % Filter
        subplot(1+n_e,6,6*i+1);
        imagesc(~isnan(Y{i}));
        set(gca,'XTick',[],'YTick',[]); title('data');

        % Filtered data
        subplot(1+n_e,6,6*i+2);
        imagesc(log(Y{i}),[min(log(Y{i}(:))),max(log(Y{i}(:)))]);
        set(gca,'XTick',[],'YTick',[]); title('filtered data');

        % Model
        subplot(1+n_e,6,6*i+3);
        imagesc(log(g{i}),[min(log(Y{i}(:))),max(log(Y{i}(:)))]);
        set(gca,'XTick',[],'YTick',[]); title('simulation');

        % Residual
        subplot(1+n_e,6,6*i+4);
        imagesc(log(Y{i})-log(g{i}),[-2 2]);
        set(gca,'XTick',[],'YTick',[]); title('residual');
        colormap('jet');

        % Residual distributions
        subplot(1+n_e,6,6*i+5); hold off;
        histogram(log(Y{i}(:))-log(g{i}(:)),50,'Normalization','pdf','facecolor',0.5*[1,1,1]); hold on;
        set(gca,'XLim',max(abs(log(Y{i}(:))-log(g{i}(:))))*[-1,1],'YTick',[]); title('error distribution');
        
        % Distance-dependent mean intensity
        subplot(1+n_e,6,6*i+6); hold off;
        plot(info.dist_dep_mean{i}.dist,info.dist_dep_mean{i}.mean,'g-','linewidth',2); hold on
        plot(info.filtered_dist_dep_mean{i}.dist,info.filtered_dist_dep_mean{i}.mean,'b-','linewidth',2); hold on
        plot(x{i},m{i},'r-','linewidth',2);
        if i == 1
            legend('data','filtered data','simulation');
        end
    end
    drawnow
end

%% EVALUATION OF NEGATIVE LOG-LIKELIHOOD AND GRADIENT
% Initialization
J = 0;
G = zeros(n_p,1);

% Noise model
for i = 1:n_e
    % Robust calculation of mixture probability
    w_i = zeros(1,2);
    q_i = zeros(length(info.dist_dep_mean{i}.mean),2);
    H_i = zeros(length(info.dist_dep_mean{i}.mean),n_p,2);
    
    % - Mixture component 1: standard noise
    w_i(1) = 1;
    q_i(:,1) = -log(sqrt(2*pi)*info.dist_dep_mean{i}.std)-(info.dist_dep_mean{i}.mean-m{i}).^2./(2*info.dist_dep_mean{i}.std.^2);
    if nargout >= 2
        H_i(:,:,1) = bsxfun(@times,(info.dist_dep_mean{i}.mean-m{i})./(info.dist_dep_mean{i}.std.^2),dmdk{i}); % derivative with respect to kinetic parameters
    end
    
    % - Mixture component 2: outlier
    % ---- is not included for standard noise model  ----
    
    % Index set of pixels which are not filtered
    ind = ~any(isnan(q_i),2);
    
    switch nargout
        case 1
            [logp] = computeMixtureProbability(w_i,q_i(ind,:));
        otherwise
            [logp,dlogpdxi] = computeMixtureProbability(w_i,q_i(ind,:),H_i(ind,:,:));
    end

    % Summation
    J = J + sum(logp);
    if nargout >= 2
        G = G + sum(dlogpdxi,1)';
    end
end

%% DEFINITION OF OUTPUTS
varargout{1} = J;
if nargout >= 2
    varargout{2} = G;
    if nargout >= 3
        varargout{3} = g;
        varargout{4} = x;
        varargout{5} = m;
    end
end
```
