## Supplementary material for "Mechanistic description of spatial processes using integrative modelling of noise-corrupted imaging data": cleanedgelist.m

```
% CLEANEDGELIST - remove short edges from a set of edgelists
%
% Function to clean up a set of edge lists generated by EDGELINK so that
% isolated edges and spurs that are shorter that a minimum length are removed.
% This code can also be use with a set of line segments generated by LINESEG.
%
% Usage: nedgelist = cleanedgelist(edgelist, minlength)
%
% Arguments:
%                edgelist - a cell array of edge lists in row,column coords in
%                           the form
%                           { [r1 c1   [r1 c1   etc }
%                              r2 c2    ...
%                                       ...
%                              rN cN]   ....]   
%                minlength - minimum edge length of interest
%
% Returns:
%                nedgelist - the new, cleaned up set of edgelists
%
% See also:  EDGELINK, DRAWEDGELIST, LINESEG

% Copyright (c) 2006-2007 Peter Kovesi
% School of Computer Science & Software Engineering
% The University of Western Australia
% pk at csse uwa edu au
% http://www.csse.uwa.edu.au/
% 
% Permission is hereby granted, free of charge, to any person obtaining a copy
% of this software and associated documentation files (the "Software"), to deal
% in the Software without restriction, subject to the following conditions:
% 
% The above copyright notice and this permission notice shall be included in 
% all copies or substantial portions of the Software.
%
% The Software is provided "as is", without warranty of any kind.
%
% December 2006  Original version
% February 2007  A major rework to fix several problems, hope they really are
%                fixed!


function nedgelist = cleanedgelist(edgelist, minlength)
    
    Nedges = length(edgelist);
    Nnodes = 2*Nedges;

    % Each edgelist has two end nodes - the starting point and the ending point.
    % We build up an adjacency/connection matrix for each node so that we can
    % determine which, if any, edgelists are connected to a node. We also
    % maintain an adjacency matrix for the edges themselves.
    % 
    % It is tricky maintaining all this information but it does allow the
    % code to run much faster.

    % First extract the end nodes from each edgelist.  The nodes are numbered
    % so that the start node has number 2*edgenumber-1 and the end node has
    % number 2*edgenumber
    node = zeros(Nnodes, 2);
    for n = 1:Nedges
        node(2*n-1,:) = edgelist{n}(1,:);
        node(2*n  ,:) = edgelist{n}(end,:);     
    end
    
    % Now build the adjacency/connection matrices. 
    A = zeros(Nnodes);   % Adjacency matrix for nodes
    B = zeros(Nedges);   % Adjacency matrix for edges
    
    for n = 1:Nnodes-1
        for m = n+1:Nnodes
            % If nodes m & n are connected
            A(n,m) =  node(n,1)==node(m,1) && node(n,2)==node(m,2);
            A(m,n) = A(n,m);
            
            if A(n,m)
                edgen = fix((n+1)/2);
                edgem = fix((m+1)/2);           
                B(edgen, edgem) = 1;
                B(edgem, edgen) = 1;            
            end
        end
    end

    % If we sum the columns of the adjacency matrix we get the number of
    % other edgelists that are connected to an edgelist
    Nconnections = sum(A);   % Connection count array for nodes
    Econnections = sum(B);   % Connection count array for edges
    

    % Check every edge to see if any of its ends are connected to just one edge.
    % This should not happen, but occasionally does due to a problem in
    % EDGELINK.  Here we simply merge it with the edge it is connected to.
    % Ultimately I want to be able to remove this block of code.
    % I think there are also some cases that are (still) not properly handled
    % by CLEANEDGELIST and there may be a case for repeating this block of
    % code at the end for another final cleanup pass
    for n = 1:Nedges
        if ~B(n,n) && ~isempty(edgelist{n}) % if edge is not connected to itself
            [spurdegree, spurnode, startnode, sconns, endnode, econns] = connectioninfo(n);
            if sconns == 1
                node2merge = find(A(startnode,:));
                mergenodes(node2merge,startnode);
            end
            
            if ~isempty(edgelist{n})   % If we have not removed this edge in
                                       % the code above check the other end.
                if econns == 1
                    node2merge = find(A(endnode,:));
                    mergenodes(node2merge,endnode);
                end         
            end
        end
    end
    
    
    % Now check every edgelist, if the edgelength is below the minimum length
    % check if we should remove it.

    if minlength > 0
        
    for n = 1:Nedges
        
        [spurdegree, spurnode] = connectioninfo(n);
        
        if ~isempty(edgelist{n}) && edgelistlength(edgelist{n}) < minlength  

            % Remove unconnected lists, or lists that are only connected to
            % themselves. 
            if ~Econnections(n) || (Econnections(n)==1 && B(n,n) == 1)
                removeedge(n);
            
            % Process edges that are spurs coming from a 3-way junction.
            elseif spurdegree == 2
                %fprintf('%d is a spur\n',n)  %%debug

                linkingedges = find(B(n,:));
                
                if length(linkingedges) == 1 % We have a loop with a spur
                                             % coming from the join in the
                                             % loop
                   % Just remove the spur, leaving the loop intact.
                   removeedge(n);  
                   
                else   % Check the other edges coming from this point. If any
                       % are also spurs make sure we remove the shortest one
                   spurs = n;
                   len = edgelistlength(edgelist{n});
                   for i = 1:length(linkingedges)
                       spurdegree = connectioninfo(linkingedges(i));
                       if spurdegree
                           spurs = [spurs linkingedges(i)];  
                           len = [len edgelistlength(edgelist{linkingedges(i)})];  
                       end
                   end

                   linkingedges = [linkingedges n];
                   
                   [mn,i] = min(len);
                   edge2delete = spurs(i);
                   [spurdegree, spurnode] = connectioninfo(edge2delete);

                   nodes2merge = find(A(spurnode,:));
                   
                   if length(nodes2merge) ~= 2
                       error('attempt to merge other than 2 nodes');
                   end
                   
                   removeedge(edge2delete);             
                   mergenodes(nodes2merge(1),nodes2merge(2))               
                   
                end 

            % Look for spurs coming from 4-way junctions that are below the minimum length
            elseif spurdegree == 3
                removeedge(n);    % Just remove it, no subsequent merging needed.
            end
        end
    end
    
    % Final cleanup of any new isolated edges that might have been created by
    % removing spurs.  An edge is isolated if it has no connections to other
    % edges, or is only connected to itself (in a loop).
    
    for n = 1:Nedges
        if ~isempty(edgelist{n}) && edgelistlength(edgelist{n}) < minlength  
            if ~Econnections(n) || (Econnections(n)==1 && B(n,n) == 1)
                removeedge(n);          
            end
        end
    end
    
    end % if minlength > 0
    
    % Run through the edgelist and extract out the non-empty lists
    m = 0;
    for n = 1:Nedges
       if ~isempty(edgelist{n})
           m = m+1;
           nedgelist{m} = edgelist{n};
       end
    end
        
    
%---------------------------------------------------------------------- 
% Internal function to merge 2 edgelists together at the specified nodes and
% perform the necessary updates to the edge adjacency and node adjacency
% matrices and the connection count arrays

function mergenodes(n1,n2)
    
    edge1 = fix((n1+1)/2);   % Indices of the edges associated with the nodes
    edge2 = fix((n2+1)/2);    

    % Get indices of nodes at each end of the two edges
    s1 = 2*edge1-1; e1 = 2*edge1;
    s2 = 2*edge2-1; e2 = 2*edge2;    
    
    if edge1==edge2
        % We should not get here, but somehow we occasionally do
        % fprintf('Nodes %d %d\n',n1,n2)    %% debug
        % warning('Attempt to merge an edge with itself')
        return
    end
    
    if ~A(n1,n2)
        error('Attempt to merge nodes that are not connected');
    end
    
    if mod(n1,2)  % node n1 is the start of edge1
        flipedge1 = 1;  % edge1 will need to be reversed in order to join edge2
    else 
        flipedge1 = 0; 
    end
    
    if mod(n2,2)  % node n2 is the start of edge2       
        flipedge2 = 0;
    else
        flipedge2 = 1;
    end
    
    % Join edgelists together - with appropriate reordering depending on which
    % end is connected to which.  The result is stored in edge1
    
    if  ~flipedge1 && ~flipedge2 
        edgelist{edge1} = [edgelist{edge1}; edgelist{edge2}];

        A(e1,:) = A(e2,:);      A(:,e1) = A(:,e2);
        Nconnections(e1) = Nconnections(e2);
        
    elseif  ~flipedge1 && flipedge2
        edgelist{edge1} = [edgelist{edge1}; flipud(edgelist{edge2})];   
        
        A(e1,:) = A(s2,:);      A(:,e1) = A(:,s2);
        Nconnections(e1) = Nconnections(s2);

    elseif  flipedge1 && ~flipedge2
        edgelist{edge1} = [flipud(edgelist{edge1}); edgelist{edge2}]; 
        
        A(s1,:) = A(e1,:);      A(:,s1) = A(:,e1);
        A(e1,:) = A(e2,:);      A(:,e1) = A(:,e2);      
        Nconnections(s1) = Nconnections(e1);
        Nconnections(e1) = Nconnections(e2);    

    elseif  flipedge1 && flipedge2
        edgelist{edge1} = [flipud(edgelist{edge1}); flipud(edgelist{edge2})];

        A(s1,:) = A(e1,:);      A(:,s1) = A(:,e1);          
        A(e1,:) = A(s2,:);      A(:,e1) = A(:,s2);
        Nconnections(s1) = Nconnections(e1);
        Nconnections(e1) = Nconnections(s2);
        
    else
        fprintf('merging edges %d and %d\n',edge1, edge2); %%debug      
        error('We should not have got here - edgelists cannot be merged');
    end
    
    % Now correct the edge adjacency matrix to reflect the new arrangement
    % The edges that the new edge1 is connected to is all the edges that
    % edge1 and edge2 were connected to
    B(edge1,:) = B(edge1,:) | B(edge2,:);
    B(:,edge1) = B(:,edge1) | B(:,edge2);    
    B(edge1, edge1) = 0;

    % Recompute connection counts because we have shuffled the adjacency matrices
    Econnections = sum(B);
    Nconnections = sum(A);
  
    removeedge(edge2);  % Finally discard edge2
    
end  % end of mergenodes

%--------------------------------------------------------------------    

% Function to provide information about the connections at each end of an
% edgelist 
%
%   [spurdegree, spurnode, startnode, sconns, endnode, econns] = connectioninfo(n)
%
%  spurdegree - If this is non-zero it indicates this edgelist is a spur, the
%               value is the number of edges this spur is connected to.
%  spurnode   - If this is a spur spurnode is the index of the node that is
%               connected to other edges, 0 otherwise.
%  startnode  - index of starting node of edgelist.
%  endnode    - index of end node of edgelist.
%  sconns     - number of connections to start node.
%  econns     - number of connections to end node.
    
function [spurdegree, spurnode, startnode, sconns, endnode, econns] = connectioninfo(n)

    if isempty(edgelist{n})
        spurdegree = 0; spurnode = 0;
        startnode = 0; sconns = 0; endnode = 0; econns = 0;
        return
    end
    
    startnode = 2*n-1;
    endnode   = 2*n;
    sconns = Nconnections(startnode);  % No of connections to start node
    econns = Nconnections(endnode);    % No of connections to end node    
    
    if sconns == 0 && econns >= 1
        spurdegree = econns;
        spurnode = endnode;
    elseif sconns >= 1 && econns == 0
        spurdegree = sconns;
        spurnode = startnode;   
    else
        spurdegree = 0;
        spurnode = 0;
    end
    
end

%--------------------------------------------------------------------
% Function to remove an edgelist and perform the necessary updates to the edge
% adjacency and node adjacency matrices and the connection count arrays

function removeedge(n)
    
    edgelist{n} = [];
    Econnections = Econnections - B(n,:); 
    Econnections(n) = 0; 
    B(n,:) = 0;
    B(:,n) = 0;
    
    nodes2delete = [2*n-1, 2*n];
    
    Nconnections = Nconnections - A(nodes2delete(1),:);    
    Nconnections = Nconnections - A(nodes2delete(2),:);        
    
    A(nodes2delete, :) = 0;
    A(:, nodes2delete) = 0;                 
    
end

%--------------------------------------------------------------------
% Function to compute the path length of an edgelist

function l = edgelistlength(edgelist)
    l = sum(sqrt(sum((edgelist(1:end-1,:)-edgelist(2:end,:)).^2, 2)));
end

%--------------------------------------------------------------------
end % End of cleanedgelists
    
```
