## Supplementary material for "Mechanistic description of spatial processes using integrative modelling of noise-corrupted imaging data": drawedgelist.m

```
% DRAWEDGELIST - plots pixels in edgelists
%
% Usage:    h =  drawedgelist(edgelist, rowscols, lw, col, figno, mid)
%
% Arguments:
%    edgelist   - Cell array of edgelists in the form
%                     { [r1 c1   [r1 c1   etc }
%                        ...
%                        rN cN]   ....]
%    rowscols -   Optional 2 element vector [rows cols] specifying the size
%                 of the image from which edges were detected (used to set
%                 size of plotted image).  If omitted or specified as [] this
%                 defaults to the bounds of the linesegment points
%    lw         - Optional line width specification. If omitted or specified
%                 as [] it defaults to a value of 1;
%    col        - Optional colour specification. Eg [0 0 1] for blue.  This
%                 can also be specified as the string 'rand' to generate a
%                 random color coding for each edgelist so that it is easier
%                 to see how the edges have been broken up into separate
%                 lists. If omitted or specified as [] it defaults to blue.
%                 col can also be a N x 3 array of colours where N is the
%                 length of edgelist, thus specifying a particulr colour for
%                 each edge.
%    figno      - Optional figure number in which to display image.
%    mid        - Optional flag 0/1.  If set each edge is drawn by joining
%                 the mid points betwen points along the edge lists.  This is
%                 intended only to be used for edgelists at pixel resolution.
%                 It reduces staircasing on diagonal lines giving nicer output.
%
% Returns:
%       h       - Array of handles to each plotted edgelist
%
% See also: EDGELINK, LINESEG

% Copyright (c) 2003-2013 Peter Kovesi
% Centre for Exploration Targeting
% The University of Western Australia
% peter.kovesi at uwa edu au
% 
% Permission is hereby granted, free of charge, to any person obtaining a copy
% of this software and associated documentation files (the "Software"), to deal
% in the Software without restriction, subject to the following conditions:
% 
% The above copyright notice and this permission notice shall be included in 
% all copies or substantial portions of the Software.
%
% The Software is provided "as is", without warranty of any kind.

% February  2003 - Original version
% September 2004 - Revised and updated
% December  2006 - Colour and linewidth specification updated
% January   2011 - Axis setting corrected (thanks to Stzpz)
% July      2013 - Colours can be specified for each edge
% Feb       2014 - Code cleanup + mid segment drawing option

function h = drawedgelist(edgelist, rowscols, lw, col, figno, mid)
    
    if ~exist('rowscols', 'var') | isempty(rowscols), rowscols = [1 1]; end
    if ~exist('lw',       'var') | isempty(lw),       lw = 1;           end
    if ~exist('col',      'var') | isempty(col),      col = [0 0 1];    end
    if  exist('figno',    'var'), figure(figno);    end
    if ~exist('mid',      'var'), mid = 0;          end
    
    debug = 0;
    Nedge = length(edgelist);
    h = zeros(length(edgelist),1);
    
%    clf;
    subplot('position',[0 0 1 1]); % Use the whole window
    
    % Set up edge colours
    if strcmp(col,'rand')
        col = hsv(Nedge);              % HSV colour map with Nedge entries
	col = col(randperm(Nedge),:);  % Form random permutation of colours
    
    elseif all(size(col) == [1 3])     % Single colour for all edges
        col = repmat(col,[Nedge 1]);
        
    elseif all(size(col) == [Nedge 3]) % Colour for each edge specified
        ;                              % we do not need to do anything
    
    else
        error('Colour not specified properly');        
    end
    
    if mid
        for I = 1:Nedge
            melist = midedge(edgelist{I});
            h(I) = line(melist(:,2), melist(:,1),...
                        'LineWidth', lw, 'Color', col(I,:));
        end	
        
    else
        for I = 1:Nedge
            h(I) = line(edgelist{I}(:,2), edgelist{I}(:,1),...
                        'LineWidth', lw, 'Color', col(I,:));
        end	
    end
    
    if debug
	for I = 1:Nedge
	    mid = fix(length(edgelist{I})/2);
	    text(edgelist{I}(mid,2), edgelist{I}(mid,1),sprintf('%d',I))
	end
    end
    
    % Check whether we need to expand bounds
    minx = 1; miny = 1;
    maxx = rowscols(2); maxy = rowscols(1);

    for I = 1:Nedge
	minx = min(min(edgelist{I}(:,2)),minx);
	miny = min(min(edgelist{I}(:,1)),miny);
	maxx = max(max(edgelist{I}(:,2)),maxx);
	maxy = max(max(edgelist{I}(:,1)),maxy);	
    end	    

    axis('equal'); axis('ij');
    axis off
    axis([minx maxx miny maxy]);

    % Set width and height of window to match aspect ratio specified by
    % rowscols scaled to a maximum size of 1000 pixels
    scale = 1000/max(rowscols);
    set(gcf, 'Position', [100 100 rowscols(2)*scale rowscols(1)*scale])
    
    if nargout == 0
        clear h
    end
    
%------------------------------------------------------------------------

% Function to construct a new edge list formed by stepping through the mid
% points of each segment of the edgelist.  The idea is to form a smoother path
% along the edgelist.  This is intended for edgelists formed at pixel
% resolution.

function melist = midedge(elist)
    
    melist =  [elist(1,:); (elist(1:end-1, :) + elist(2:end, :))/2; elist(end,:)];
    
    
```
