## Supplementary material for "Mechanistic description of spatial processes using integrative modelling of noise-corrupted imaging data": edgelink.m

```
% EDGELINK - Link edge points in an image into lists
%
% Usage: [edgelist edgeim, etype] = edgelink(im, minlength, location)
%
%    **Warning** 'minlength' is ignored at the moment because 'cleanedgelist'
%                 has some bugs and can be memory hungry
%
% Arguments:  im         - Binary edge image, it is assumed that edges
%                          have been thinned (or are nearly thin).
%             minlength  - Optional minimum edge length of interest, defaults
%                          to 1 if omitted or specified as []. Ignored at the
%                          moment. 
%             location   - Optional complex valued image holding subpixel
%                          locations of edge points. For any pixel the
%                          real part holds the subpixel row coordinate of
%                          that edge point and the imaginary part holds
%                          the column coordinate.  See NONMAXSUP.  If
%                          this argument is supplied the edgelists will
%                          be formed from the subpixel coordinates,
%                          otherwise the the integer pixel coordinates of
%                          points in 'im' are used.
%
% Returns:  edgelist - a cell array of edge lists in row,column coords in
%                      the form
%                     { [r1 c1   [r1 c1   etc }
%                        r2 c2    ...
%                        ...
%                        rN cN]   ....]   
%
%           edgeim   - Image with pixels labeled with edge number. 
%                      Note that junctions in the labeled edge image will be
%                      labeled with the edge number of the last edge that was
%                      tracked through it.  Note that this image also includes
%                      edges that do not meet the minimum length specification.
%                      If you want to see just the edges that meet the
%                      specification you should pass the edgelist to
%                      DRAWEDGELIST.
%
%            etype   - Array of values, one for each edge segment indicating
%                      its type
%                      0  - Start free, end free
%                      1  - Start free, end junction
%                      2  - Start junction, end free (should not happen)
%                      3  - Start junction, end junction
%                      4  - Loop
%
% This function links edge points together into lists of coordinate pairs.
% Where an edge junction is encountered the list is terminated and a separate
% list is generated for each of the branches.
%
% Note I am not sure if using bwmorph's 'thin' or 'skel' is best for
% preprocessing the edge image prior to edgelinking.  The main issue is the
% treatment of junctions.  Skel can result in an image where multiple adjacent
% junctions are produced (maybe this is more a problem with my junction
% detection code).  Thin, on the other hand, can produce different output when
% you rotate an image by 90 degrees. On balance I think using 'thin' is better.
% Note, however, the input image should be 'nearly thin' otherwise the thinning
% operation could shorten the ends of structures. Skeletonisation and thinning
% is surprisingly awkward.
%
% See also:  DRAWEDGELIST, LINESEG, MAXLINEDEV, CLEANEDGELIST,
%            FINDENDSJUNCTIONS, FILLEDGEGAPS

% Copyright (c) 1996-2013 Peter Kovesi
% Centre for Exploration Targeting
% The University of Western Australia
% peter.kovesi at uwa edu au
% 
% Permission is hereby granted, free of charge, to any person obtaining a copy
% of this software and associated documentation files (the "Software"), to deal
% in the Software without restriction, subject to the following conditions:
% 
% The above copyright notice and this permission notice shall be included in 
% all copies or substantial portions of the Software.
%
% The Software is provided "as is", without warranty of any kind.

% February  2001 - Original version
% September 2004 - Revised to allow subpixel edge data to be used
% November  2006 - Changed so that edgelists start and stop at every junction 
% January   2007 - Trackedge modified to discard isolated pixels and the
%                  problems they cause (thanks to Jeff Copeland)
% January   2007 - Fixed so that closed loops are closed!
% May       2013 - Completely redesigned with a new linking strategy that
%                  hopefully handles adjacent junctions correctly. It runs
%                  about twice as fast too.

function [edgelist, edgeim, etype] = edgelink(im, minlength, location)
    
    % Set up some global variables to avoid passing (and copying) of arguments,
    % this improves speed.
    global EDGEIM;
    global ROWS;
    global COLS;
    global JUNCT;
    
    if ~exist('minlength','var') || isempty(minlength), minlength = 0; end
    
    EDGEIM = im ~= 0;                     % Make sure image is binary.
    EDGEIM = bwmorph(EDGEIM,'clean');     % Remove isolated pixels

    % Make sure edges are thinned.  Use 'thin' rather than 'skel', see
    % comments in header.
    EDGEIM = bwmorph(EDGEIM,'thin',Inf); 
    [ROWS, COLS] = size(EDGEIM);
    
    % Find endings and junctions in edge data
    [RJ, CJ, re, ce] = findendsjunctions(EDGEIM);
    Njunct = length(RJ);
    Nends = length(re);
    
    % Create a sparse matrix to mark junction locations. This makes junction
    % testing much faster.  A value of 1 indicates a junction, a value of 2
    % indicates we have visited the junction.
    JUNCT = spalloc(ROWS,COLS, Njunct);
    for n = 1:Njunct
        JUNCT(RJ(n),CJ(n)) = 1;    
    end

    % ? Think about using labels >= 2 so that EDGEIM can be uint16, say. %
    EDGEIM = double(EDGEIM);   % Cast to double to allow the use of -ve labelings
    edgeNo = 0;
    
    % Summary of strategy:
    % 1) From every end point track until we encounter an end point or
    % junction.  As we track points along an edge image pixels are labeled with
    % the -ve of their edge No.
    % 2) From every junction track out on any edges that have not been
    % labeled yet.
    % 3) Scan through the image looking for any unlabeled pixels.  These
    % correspond to isolated loops that have no junctions.


    %% 1) Form tracks from each unlabeled endpoint until we encounter another
    % endpoint or junction.
    for n = 1:Nends
        if EDGEIM(re(n),ce(n)) == 1  % Endpoint is unlabeled
            edgeNo = edgeNo + 1;
            [edgelist{edgeNo} endType] = trackedge(re(n), ce(n), edgeNo);
            etype(edgeNo) = endType;
        end
    end
    
    %% 2) Handle junctions.
    % Junctions are awkward when they are adjacent to other junctions.  We
    % start by looking at all the neighbours of a junction.  
    % If there is an adjacent junction we first create a 2-element edgetrack
    % that links the two junctions together.  We then look to see if there are
    % any non-junction edge pixels that are adjacent to both junctions. We then
    % test to see which of the two junctions is closest to this common pixel and
    % initiate an edge track from the closest of the two junctions through this
    % pixel.  When we do this we set the 'avoidJunction' flag in the call to
    % trackedge so that the edge track does not immediately loop back and
    % terminate on the other adjacent junction.
    % Having checked all the common neighbours of both junctions we then
    % track out on any remaining untracked neighbours of the junction 
    
    for j = 1:Njunct
        if JUNCT(RJ(j),CJ(j)) ~= 2;  % We have not visited this junction
            JUNCT(RJ(j),CJ(j)) = 2;

            % Call availablepixels with edgeNo = 0 so that we get a list of
            % available neighbouring pixels that can be linked to and a list of
            % all neighbouring pixels that are also junctions.
            [ra, ca, rj, cj] =  availablepixels(RJ(j), CJ(j), 0);
        
            for k = 1:length(rj)        % For all adjacent junctions...
                % Create a 2-element edgetrack to each adjacent junction
                edgeNo = edgeNo + 1;
                edgelist{edgeNo} = [RJ(j) CJ(j); rj(k) cj(k)];
                etype(edgeNo) = 3;  % Edge segment is junction-junction
                EDGEIM(RJ(j), CJ(j)) = -edgeNo;
                EDGEIM(rj(k), cj(k)) = -edgeNo;
            
                % Check if the adjacent junction has some untracked pixels that
                % are also adjacent to the initial junction.  Thus we need to
                % get available pixels adjacent to junction (rj(k) cj(k))
                [rak, cak] = availablepixels(rj(k), cj(k));
                
                % If both junctions have untracked neighbours that need checking...
                if ~isempty(ra) && ~isempty(rak)
                    
                    % Find untracked neighbours common to both junctions. 
                    commonrc =  intersect([ra ca], [rak cak], 'rows');
                    
                    for n = 1:size(commonrc, 1);
                        % If one of the junctions j or k is closer to this common
                        % neighbour use that as the start of the edge track and the
                        % common neighbour as the 2nd element. When we call
                        % trackedge we set the avoidJunction flag to prevent the
                        % track immediately connecting back to the other junction.
                        distj = norm(commonrc(n,:) - [RJ(j) CJ(j)]);
                        distk = norm(commonrc(n,:) - [rj(k) cj(k)]);
                        edgeNo = edgeNo + 1;
                        if distj < distk
                            edgelist{edgeNo} = trackedge(RJ(j), CJ(j), edgeNo, ...
                                                   commonrc(n,1), commonrc(n,2), 1);
                        else                                                
                            edgelist{edgeNo} = trackedge(rj(k), cj(k), edgeNo, ...
                                                   commonrc(n,1), commonrc(n,2), 1);
                        end
                        etype(edgeNo) = 3;  % Edge segment is junction-junction
                    end
                end
            
                % Track any remaining unlabeled pixels adjacent to this junction k
                for m = 1:length(rak)
                    if EDGEIM(rak(m), cak(m)) == 1
                        edgeNo = edgeNo + 1;
                        edgelist{edgeNo} = trackedge(rj(k), cj(k), edgeNo, rak(m), cak(m));
                        etype(edgeNo) = 3;  % Edge segment is junction-junction
                    end    
                end
                
                % Mark that we have visited junction (rj(k) cj(k))
                JUNCT(rj(k), cj(k)) = 2;
                
            end % for all adjacent junctions

            % Finally track any remaining unlabeled pixels adjacent to original junction j
            for m = 1:length(ra)
                if EDGEIM(ra(m), ca(m)) == 1
                    edgeNo = edgeNo + 1;
                    edgelist{edgeNo} = trackedge(RJ(j), CJ(j), edgeNo, ra(m), ca(m));
                    etype(edgeNo) = 3;  % Edge segment is junction-junction                
                end    
            end
        
        end  % If we have not visited this junction
    end   % For each junction
    
    %% 3) Scan through the image looking for any unlabeled pixels.  These
    % should correspond to isolated loops that have no junctions or endpoints.
    for ru = 1:ROWS
        for cu = 1:COLS
            if EDGEIM(ru,cu) == 1  % We have an unlabeled edge
                edgeNo = edgeNo + 1; 
                [edgelist{edgeNo} endType] = trackedge(ru, cu, edgeNo);
                etype(edgeNo) = endType; 
            end
        end
    end
    
    edgeim = -EDGEIM;  % Finally negate image to make edge encodings +ve.

    % Eliminate isolated edges and spurs that are below the minimum length
    % ** DISABLED for the time being **
%    if nargin >= 2 && ~isempty(minlength)
%	edgelist = cleanedgelist2(edgelist, minlength);
%    end
   
    % If subpixel edge locations are supplied upgrade the integer precision
    % edgelists that were constructed with data from 'location'.
    if nargin == 3
	for I = 1:length(edgelist)
	    ind = sub2ind(size(im),edgelist{I}(:,1),edgelist{I}(:,2));
	    edgelist{I}(:,1) = real(location(ind))';
	    edgelist{I}(:,2) = imag(location(ind))';    
	end
    end

    clear global EDGEIM;
    clear global ROWS;
    clear global COLS;
    clear global JUNCT;
    
%----------------------------------------------------------------------    
% TRACKEDGE
%
% Function to track all the edge points starting from an end point or junction.
% As it tracks it stores the coords of the edge points in an array and labels the
% pixels in the edge image with the -ve of their edge number. This continues
% until no more connected points are found, or a junction point is encountered.
%
% Usage:   edgepoints = trackedge(rstart, cstart, edgeNo, r2, c2, avoidJunction)
% 
% Arguments:   rstart, cstart   - Row and column No of starting point.
%              edgeNo           - The current edge number.
%              r2, c2           - Optional row and column coords of 2nd point.
%              avoidJunction    - Optional flag indicating that (r2,c2)
%                                 should not be immediately connected to a
%                                 junction (if possible).
%
% Returns:     edgepoints       - Nx2 array of row and col values for
%                                 each edge point.
%              endType          - 0 for a free end
%                                 1 for a junction
%                                 5 for a loop

function [edgepoints endType] = trackedge(rstart, cstart, edgeNo, r2, c2, avoidJunction)
    
    global EDGEIM;
    global JUNCT;
    
    if ~exist('avoidJunction', 'var'), avoidJunction = 0; end
    
    edgepoints = [rstart cstart];      % Start a new list for this edge.
    EDGEIM(rstart,cstart) = -edgeNo;   % Edge points in the image are 
			               % encoded by -ve of their edgeNo.

    preferredDirection = 0;            % Flag indicating we have/not a
                                       % preferred direction.
    
    % If the second point has been supplied add it to the track and set the
    % path direction
    if exist('r2', 'var') && exist('c2', 'var')
        edgepoints = [edgepoints
                      r2    c2 ];
        EDGEIM(r2, c2) = -edgeNo;
        % Initialise direction vector of path and set the current point on
        % the path
        dirn = unitvector([r2-rstart c2-cstart]);
        r = r2;
        c = c2;
        preferredDirection = 1;
    else
        dirn = [0 0];  
        r = rstart;
        c = cstart;
    end
    
    % Find all the pixels we could link to
    [ra, ca, rj, cj] = availablepixels(r, c, edgeNo);
    
    while ~isempty(ra) || ~isempty(rj)
        
        % First see if we can link to a junction. Choose the junction that
        % results in a move that is as close as possible to dirn. If we have no
        % preferred direction, and there is a choice, link to the closest
        % junction
        % We enter this block:
        % IF there are junction points and we are not trying to avoid a junction
        % OR there are junction points and no non-junction points, ie we have
        % to enter it even if we are trying to avoid a junction
        if (~isempty(rj) && ~avoidJunction)  || (~isempty(rj) && isempty(ra))

            % If we have a prefered direction choose the junction that results
            % in a move that is as close as possible to dirn.
            if preferredDirection  
                dotp = -inf;
                for n = 1:length(rj)
                    dirna = unitvector([rj(n)-r  cj(n)-c]); 
                    dp = dirn*dirna';
                    if dp > dotp
                        dotp = dp;
                        rbest = rj(n); cbest = cj(n);
                        dirnbest = dirna;
                    end
                end            
            
            % Otherwise if we have no established direction, we should pick a
            % 4-connected junction if possible as it will be closest.  This only
            % affects tracks of length 1 (Why do I worry about this...?!).
            else
                distbest = inf;
                for n = 1:length(rj)
                    dist = sum([rj(n)-r;  cj(n)-c]); 
                    if dist < distbest
                        rbest = rj(n); cbest = cj(n);
                        distbest = dist;
                        dirnbest = unitvector([rj(n)-r  cj(n)-c]); 
                    end
                end
                preferredDirection = 1;
            end
            
        % If there were no junctions to link to choose the available
        % non-junction pixel that results in a move that is as close as possible
        % to dirn
        else    
            dotp = -inf;
            for n = 1:length(ra)
                dirna = unitvector([ra(n)-r  ca(n)-c]); 
                dp = dirn*dirna';
                if dp > dotp
                    dotp = dp;
                    rbest = ra(n); cbest = ca(n);
                    dirnbest = dirna;
                end
            end

            avoidJunction = 0; % Clear the avoidJunction flag if it had been set
        end
        
        % Append the best pixel to the edgelist and update the direction and EDGEIM
        r = rbest; c = cbest;
        edgepoints = [edgepoints
                        r    c  ];
        dirn = dirnbest;
        EDGEIM(r, c) = -edgeNo;

        % If this point is a junction exit here
        if JUNCT(r, c);
            endType = 1;  % Mark end as being a junction
            return;
        else
            % Get the next set of available pixels to link.
            [ra, ca, rj, cj] = availablepixels(r, c, edgeNo);  
        end
    end
    
    % If we get here we are at an endpoint or our sequence of pixels form a
    % loop.  If it is a loop the edgelist should have start and end points
    % matched to form a loop.  If the number of points in the list is four or
    % more (the minimum number that could form a loop), and the endpoints are
    % within a pixel of each other, append a copy of the first point to the end
    % to complete the loop
     
    endType = 0;  % Mark end as being free, unless it is reset below
    
    if length(edgepoints) >= 4
	if abs(edgepoints(1,1) - edgepoints(end,1)) <= 1  &&  ...
           abs(edgepoints(1,2) - edgepoints(end,2)) <= 1 
	    edgepoints = [edgepoints
			  edgepoints(1,:)];
            endType = 5; % Mark end as being a loop
        end
    end

%----------------------------------------------------------------------    
% AVAILABLEPIXELS
%
% Find all the pixels that could be linked to point r, c
%
% Arguments:  rp, cp - Row, col coordinates of pixel of interest.
%             edgeNo - The edge number of the edge we are seeking to
%                      track. If not supplied its value defaults to 0
%                      resulting in all adjacent junctions being returned,
%                      (see note below)
%
% Returns:    ra, ca - Row and column coordinates of available non-junction
%                      pixels.
%             rj, cj - Row and column coordinates of available junction
%                      pixels.
%
% A pixel is avalable for linking if it is:
% 1) Adjacent, that is it is 8-connected.
% 2) Its value is 1 indicating it has not already been assigned to an edge
% 3) or it is a junction that has not been labeled -edgeNo indicating we have
%    not already assigned it to the current edge being tracked.  If edgeNo is
%    0 all adjacent junctions will be returned
    
function  [ra, ca, rj, cj] = availablepixels(rp, cp, edgeNo)

    global EDGEIM;
    global JUNCT;
    global ROWS;
    global COLS;
    
    % If edgeNo not supplied set to 0 to allow all adjacent junctions to be returned
    if ~exist('edgeNo', 'var'), edgeNo = 0; end
    
    ra = []; ca = [];
    rj = []; cj = [];
    
    % row and column offsets for the eight neighbours of a point
    roff = [-1  0  1  1  1  0 -1 -1];
    coff = [-1 -1 -1  0  1  1  1  0];
    
    r = rp+roff;
    c = cp+coff;
    
    % Find indices of arrays of r and c that are within the image bounds
    ind = find((r>=1 & r<=ROWS) & (c>=1 & c<=COLS));

    % A pixel is avalable for linking if its value is 1 or it is a junction
    % that has not been labeled -edgeNo
    for i = ind
        if EDGEIM(r(i),c(i)) == 1 && ~JUNCT(r(i), c(i));
            ra = [ra; r(i)];
            ca = [ca; c(i)];
        elseif (EDGEIM(r(i),c(i)) ~= -edgeNo) && JUNCT(r(i), c(i));
            rj = [rj; r(i)];
            cj = [cj; c(i)];
        end
    end
    
    
%---------------------------------------------------------------------    
% UNITVECTOR Normalises a vector to unit magnitude
%

function nv = unitvector(v)
    
    nv = v./sqrt(v(:)'*v(:));    
```
