## Supplementary material for "Mechanistic description of spatial processes using integrative modelling of noise-corrupted imaging data": edgelist2image.m

```
% EDGELIST2IMAGE - transfers edgelist data back into a 2D image array
%
% Usage:      im = edgelist2image(edgelist, rowscols)
%
%    edgelist   - Cell array of edgelists in the form
%                     { [r1 c1   [r1 c1   etc }
%                        ...
%                        rN cN]   ....]
%    rowscols -   Optional 2 element vector [rows cols] specifying the size
%                 of the image from which edges were detected (used to set
%                 size of plotted image).  If omitted or specified as [] this
%                 defaults to the bounds of the linesegment points
%
% Note this function will only work effectively on 'dense' edgelist data
% obtained, say, directly from edgelink.  If you have subsequently fitted
% line segments to the edgelist data this function will only mark the endpoints
% of the segments,  use DRAWEDGELIST instead.
%
% See also: EDGELINK, DRAWEDGELIST

% Copyright (c) 2007 Peter Kovesi
% 
% Permission is hereby granted, free of charge, to any person obtaining a copy
% of this software and associated documentation files (the "Software"), to deal
% in the Software without restriction, subject to the following conditions:
% 
% The above copyright notice and this permission notice shall be included in 
% all copies or substantial portions of the Software.
%
% The Software is provided "as is", without warranty of any kind.

% September 2007


function im = edgelist2image(edgelist, rowscols)

    if nargin < 2, rowscols = [1 1]; end

    Nedge = length(edgelist);
    
    % Establish bounds of image
    minx = 1; miny = 1;
    maxx = rowscols(2); maxy = rowscols(1);

    for I = 1:Nedge
	minx = min(min(edgelist{I}(:,2)),minx);
	miny = min(min(edgelist{I}(:,1)),miny);
	maxx = max(max(edgelist{I}(:,2)),maxx);
	maxy = max(max(edgelist{I}(:,1)),maxy);	
    end	    
    
    % Draw the edgelist data into an image array
    im = zeros(maxy,maxx);
    
    for I = 1:Nedge
	ind = sub2ind([maxy maxx], edgelist{I}(:,1), edgelist{I}(:,2));
	im(ind) = 1;
    end	
    
```
