## Supplementary material for "Mechanistic description of spatial processes using integrative modelling of noise-corrupted imaging data": filledgegaps.m

```
% FILLEDGEGAPS  Fills small gaps in a binary edge map image
%
% Usage: bw2 = filledgegaps(bw, gapsize)
%
% Arguments:    bw - Binary edge image
%          gapsize - The edge gap size that you wish to be able to fill.
%                    Use the smallest value you can. (Odd values work best). 
%
% Returns:     bw2 - The binary edge image with gaps filled.
%
%
% Strategy: A binary circular blob of radius = gapsize/2 is placed at the end of
% every edge segment.  If the ends of two edge segments are in close proximity
% the circular blobs will overlap.  The image is then thinned.  Where circular
% blobs at end points overlap the thinning process will leave behind a line of
% pixels linking the two end points.  Where an end point is isolated the
% thinning process will erode the circular blob away so that the original edge
% segment is restored.
%
% Use the smallest gapsize value you can.  With large values all sorts of
% unwelcome linking can occur.
%
% The circular blobs are generated using the function CIRCULARSTRUCT which,
% unlike MATLAB's STREL, will accept real valued radius values.  Note that I
% suggest that you use an odd value for 'gapsize'.  This results in a radius
% value of the form x.5 being passed to CIRCULARSTRUCT which results in discrete
% approximation to a circle that seems to respond to thinning in a 'good'
% way. With integer radius values CIRCULARSTRUCT can produce circles that
% result in minor artifacts being generated by the thinning process.
%
% See also: FINDENDSJUNCTIONS, FINDISOLATEDPIXELS, CIRCULARSTRUCT, EDGELINK

% Copyright (c) 2013 Peter Kovesi
% Centre for Exploration Targeting
% The University of Western Australia
% peter.kovesi at uwa edu au
% 
% Permission is hereby granted, free of charge, to any person obtaining a copy
% of this software and associated documentation files (the "Software"), to deal
% in the Software without restriction, subject to the following conditions:
% 
% The above copyright notice and this permission notice shall be included in 
% all copies or substantial portions of the Software.
%
% The Software is provided "as is", without warranty of any kind.

% PK May 2013

function bw = filledgegaps(bw, gapsize)
    
    [rows, cols] = size(bw);
    
    % Generate a binary circle with radius gapsize/2 (but not less than 1)
    blob = circularstruct(max(gapsize/2, 1));
    
    rad = (size(blob,1)-1)/2;  % Radius of resulting blob matrix. Note
                               % circularstruct returns an odd sized matrix
    
    % Get coordinates of end points and of isolated pixels
    [~, ~, re, ce] = findendsjunctions(bw);
    [ri, ci] = findisolatedpixels(bw);
    
    re = [re;ri];
    ce = [ce;ci];

    % Place a circular blob at every endpoint and isolated pixel
    for n = 1:length(re)
        
        if (re(n) > rad) && (re(n) < rows-rad) && ...
                (ce(n) > rad) && (ce(n) < cols-rad)

            bw(re(n)-rad:re(n)+rad, ce(n)-rad:ce(n)+rad) = ...
                bw(re(n)-rad:re(n)+rad, ce(n)-rad:ce(n)+rad) | blob;
        end
    end
    
    bw = bwmorph(bw, 'thin', inf);  % Finally thin
    
    % At this point, while we may have joined endpoints that were close together
    % we typically have also generated a number of small loops where there were
    % more than one endpoint close to an edge.  To address this we identfy the
    % loops by finding 4-connected blobs in the inverted image.  Blobs that are
    % less than or equal to the size of the blobs we used to link edges are
    % filled in, the image reinverted and then rethinned.
    
    L = bwlabel(~bw,4); 
    stats = regionprops(L, 'Area');
    
    % Get blobs with areas <= pi* (gapsize/2)^2
    ar = cat(1,stats.Area);
    ind = find(ar <= pi*(gapsize/2)^2);
    
    % Fill these blobs in image bw
    for n = ind'
        bw(L==n) = 1;
    end
    
    bw = bwmorph(bw, 'thin', inf);  % thin again
    
```
