## Supplementary material for "Mechanistic description of spatial processes using integrative modelling of noise-corrupted imaging data": findendsjunctions.m

```
% FINDENDSJUNCTIONS - find junctions and endings in a line/edge image
%
% Usage: [rj, cj, re, ce] = findendsjunctions(edgeim, disp)
% 
% Arguments:  edgeim - A binary image marking lines/edges in an image.  It is
%                      assumed that this is a thinned or skeleton image 
%             disp   - An optional flag 0/1 to indicate whether the edge
%                      image should be plotted with the junctions and endings
%                      marked.  This defaults to 0.
%
% Returns:    rj, cj - Row and column coordinates of junction points in the
%                      image. 
%             re, ce - Row and column coordinates of end points in the
%                      image.
%
% See also: EDGELINK
%
% Note I am not sure if using bwmorph's 'thin' or 'skel' is best for finding
% junctions.  Skel can result in an image where multiple adjacent junctions are
% produced (maybe this is more a problem with this junction detection code).
% Thin, on the other hand, can produce different output when you rotate an image
% by 90 degrees.  On balance I think using 'thin' is better. Skeletonisation and
% thinning is surprisingly awkward.

% Copyright (c) 2006-2013 Peter Kovesi
% Centre for Exploration Targeting
% The University of Western Australia
% peter.kovesi at uwa edu au
% 
% Permission is hereby granted, free of charge, to any person obtaining a copy
% of this software and associated documentation files (the "Software"), to deal
% in the Software without restriction, subject to the following conditions:
% 
% The above copyright notice and this permission notice shall be included in 
% all copies or substantial portions of the Software.
%
% The Software is provided "as is", without warranty of any kind.

% November 2006  - Original version
% May      2013  - Call to bwmorph to ensure image is thinned was removed as
%                  this might cause problems if the image used to find
%                  junctions is different from the image used for, say,
%                  edgelinking 

function [rj, cj, re, ce] = findendsjunctions(b, disp)

    if nargin == 1
	disp = 0;
    end
    
    % Set up look up table to find junctions.  To do this we use the function
    % defined at the end of this file to test that the centre pixel within a 3x3
    % neighbourhood is a junction.
    lut = makelut(@junction, 3);
    junctions = applylut(b, lut);
    [rj,cj] = find(junctions);
    
    % Set up a look up table to find endings.  
    lut = makelut(@ending, 3);
    ends = applylut(b, lut);
    [re,ce] = find(ends);    

    if disp    
	show(edgeim,1), hold on
	plot(cj,rj,'r+')
	plot(ce,re,'g+')    
    end

%----------------------------------------------------------------------
% Function to test whether the centre pixel within a 3x3 neighbourhood is a
% junction. The centre pixel must be set and the number of transitions/crossings
% between 0 and 1 as one traverses the perimeter of the 3x3 region must be 6 or
% 8.
%
% Pixels in the 3x3 region are numbered as follows
%
%       1 4 7
%       2 5 8
%       3 6 9

function b = junction(x)
    
    a = [x(1) x(2) x(3) x(6) x(9) x(8) x(7) x(4)]';
    b = [x(2) x(3) x(6) x(9) x(8) x(7) x(4) x(1)]';    
    crossings = sum(abs(a-b));
    
    b = x(5) && crossings >= 6;
    
%----------------------------------------------------------------------
% Function to test whether the centre pixel within a 3x3 neighbourhood is an
% ending. The centre pixel must be set and the number of transitions/crossings
% between 0 and 1 as one traverses the perimeter of the 3x3 region must be 2.
%
% Pixels in the 3x3 region are numbered as follows
%
%       1 4 7
%       2 5 8
%       3 6 9

function b = ending(x)
    a = [x(1) x(2) x(3) x(6) x(9) x(8) x(7) x(4)]';
    b = [x(2) x(3) x(6) x(9) x(8) x(7) x(4) x(1)]';    
    crossings = sum(abs(a-b));
    
    b = x(5) && crossings == 2;
```
