## Supplementary material for "Mechanistic description of spatial processes using integrative modelling of noise-corrupted imaging data": findisolatedpixels.m

```
% FINDENDSJUNCTIONS - find isolated pixels in a binary image
%
% Usage: [r, c] = findisolatedpixels(b)
% 
% Argument:      b - A binary image 
%
% Returns:    r, c - Row and column coordinates of isolated pixels in the
%                    image. 
%
% See also: FINDENDSJUNCTIONS
%

% Copyright (c) 2013 Peter Kovesi
% Centre for Exploration Targeting
% The University of Western Australia
% peter.kovesi at uwa edu au
% 
% Permission is hereby granted, free of charge, to any person obtaining a copy
% of this software and associated documentation files (the "Software"), to deal
% in the Software without restriction, subject to the following conditions:
% 
% The above copyright notice and this permission notice shall be included in 
% all copies or substantial portions of the Software.
%
% The Software is provided "as is", without warranty of any kind.

% May      2013 

function [r, c] = findisolatedpixels(b)

    lut = makelut(@isolated, 3);
    isolated = applylut(b, lut);
    [r, c] = find(isolated);

%----------------------------------------------------------------------
% Function to test whether the centre pixel within a 3x3 neighbourhood is
% isolated.
%
% Pixels in the 3x3 region are numbered as follows
%
%       1 4 7
%       2 5 8
%       3 6 9

function b = isolated(x)
    
    b = x(5) && sum(x(:)) == 1;
```
