## Supplementary material for "Mechanistic description of spatial processes using integrative modelling of noise-corrupted imaging data": lineseg.m

```
% LINESEG - Form straight line segements from an edge list.
%
% Usage: seglist = lineseg(edgelist, tol)
%
% Arguments:  edgelist - Cell array of edgelists where each edgelist is an
%                        Nx2 array of (row col) coords.
%             tol      - Maximum deviation from straight line before a
%                        segment is broken in two (measured in pixels).
% Returns:
%             seglist  - A cell array of in the same format of the input
%                        edgelist but each seglist is a subsampling of its
%                        corresponding edgelist such that straight line
%                        segments between these subsampled points do not
%                        deviate from the original points by more than tol.
%
% This function takes each array of edgepoints in edgelist, finds the
% size and position of the maximum deviation from the line that joins the
% endpoints, if the maximum deviation exceeds the allowable tolerance the
% edge is shortened to the point of maximum deviation and the test is
% repeated.  In this manner each edge is broken down to line segments,
% each of which adhere to the original data with the specified tolerance.
%
% See also:  EDGELINK, MAXLINEDEV, DRAWEDGELIST
%

% Copyright (c) 2000-2006 Peter Kovesi
% School of Computer Science & Software Engineering
% The University of Western Australia
% http://www.csse.uwa.edu.au/
% 
% Permission is hereby granted, free of charge, to any person obtaining a copy
% of this software and associated documentation files (the "Software"), to deal
% in the Software without restriction, subject to the following conditions:
% 
% The above copyright notice and this permission notice shall be included in 
% all copies or substantial portions of the Software.
%
% The Software is provided "as is", without warranty of any kind.

% December 2000 - Original version
% February 2003 - Added the returning of nedgelist data.
% December 2006 - Changed so that separate cell arrays of line segments are
%                 formed, in the same format used for edgelists


function seglist = lineseg(edgelist, tol)
    
    Nedge = length(edgelist);
    seglist = cell(1,Nedge);
    
    for e = 1:Nedge
        y = edgelist{e}(:,1);   % Note that (col, row) corresponds to (x,y)
	x = edgelist{e}(:,2);

	fst = 1;                % Indices of first and last points in edge
	lst = length(x);        % segment being considered.

	Npts = 1;	
	seglist{e}(Npts,:) = [y(fst) x(fst)];
	
	while  fst<lst
	    [m,i] = maxlinedev(x(fst:lst),y(fst:lst));  % Find size & posn of
                                                        % maximum deviation.
	    
	    while m > tol       % While deviation is > tol  
		lst = i+fst-1;  % Shorten line to point of max deviation by adjusting lst
		[m,i] = maxlinedev(x(fst:lst),y(fst:lst));
	    end
	
	    Npts = Npts+1;
	    seglist{e}(Npts,:) = [y(lst) x(lst)];
	    
	    fst = lst;        % reset fst and lst for next iteration
	    lst = length(x);
	end
    end

    
```
