## Supplementary material for "Mechanistic description of spatial processes using integrative modelling of noise-corrupted imaging data": maxlinedev.m

```
% MAXLINEDEV - Find max deviation from a line in an edge contour.
%
% Function finds the point of maximum deviation from a line joining the
% endpoints of an edge contour.
%
% Usage:   [maxdev, index, D, totaldev] = maxlinedev(x,y)
%
% Arguments:
%          x, y   - arrays of x,y  (col,row) indicies of connected pixels 
%                   on the contour.
% Returns:
%          maxdev   - Maximum deviation of contour point from the line
%                     joining the end points of the contour (pixels).
%          index    - Index of the point having maxdev.
%          D        - Distance between end points of the contour so that
%                     one can calculate maxdev/D - the normalised error.
%          totaldev - Sum of the distances of all the pixels from the
%                     line joining the endpoints.
%
% See also:  EDGELINK, LINESEG
%

% Peter Kovesi  
% School of Computer Science & Software Engineering
% The University of Western Australia
% pk at csse uwa edu au
% http://www.csse.uwa.edu.au/~pk
%
% December 2000 - Original version
% February 2003 - Added calculation of total deviation
% August   2006 - Avoid degeneracy when endpoints are coincident
% February 2007 - Corrected distance calculation when endpoints are
%                 coincident.

function [maxdev, index, D, totaldev] = maxlinedev(x,y)

    Npts = length(x);
    
    if Npts == 1
	warning('Contour of length 1');
	maxdev = 0; index = 1;
	D = 1; totaldev = 0;
	return;
    elseif Npts == 0
	error('Contour of length 0');
    end

    %    D = norm([x(1) y(1)] - [x(Npts) y(Npts)]);  % Distance between end points
    D = sqrt((x(1)-x(Npts))^2 + (y(1)-y(Npts))^2); % This runs much faster

    if D > eps    
	
	% Eqn of line joining end pts (x1 y1) and (x2 y2) can be parameterised by
	%    
	%    x*(y1-y2) + y*(x2-x1) + y2*x1 - y1*x2 = 0
	%
	% (See Jain, Rangachar and Schunck, "Machine Vision", McGraw-Hill
	% 1996. pp 194-196)
	
	y1my2 = y(1)-y(Npts);                       % Pre-compute parameters
	x2mx1 = x(Npts)-x(1);
	C = y(Npts)*x(1) - y(1)*x(Npts);
	
	% Calculate distance from line segment for each contour point
	d = abs(x*y1my2 + y*x2mx1 + C)/D;          
	
    else    % End points are coincident, calculate distances from 1st point
    
        d = sqrt((x - x(1)).^2 + (y - y(1)).^2);
	D = 1;  % Now set D to 1 so that normalised error can be used
	
    end						

    [maxdev, index] = max(d);

    if nargout == 4
	totaldev = sum(d.^2);
    end
```
