## Supplementary material for "Mechanistic description of spatial processes using integrative modelling of noise-corrupted imaging data": DRAMOptions.m

```
classdef DRAMOptions < matlab.mixin.CustomDisplay
   % DRAMOptions provides an option container to pass options 
   % into the PestoSamplingOptions class for Delayed Rejection Adaption Metropolis (DRAM).
   % The DRAM algorithm uses a delayed rejection scheme for better mixing.
   % 
   % This file is based on AMICI amioptions.m (http://icb-dcm.github.io/AMICI/)
   
   properties      
      % This factor is used for regularization in cases where the single-chain proposal
      % covariance matrices are ill conditioned. Larger values equal
      % stronger regularization.
      regFactor = 1e-6;
      
      % The number of tries in the delayed rejection scheme
      nTry = 1;
      
      % Defines the level of verbosity 'silent', 'visual', 'debug' or 'text'
      verbosityMode = 'text';
      
      %  Update the proposal density only every adaptionInterval time
      adaptionInterval = 1;
      
   end
   
   methods
      function obj = DRAMOptions(varargin)
         % DRAMOptions Construct a new DRAMOptions object
         %
         %   OPTS = DRAMOptions() creates a set of options with
         %   each option set to itsdefault value.
         %
         %   OPTS = DRAMOptions(PARAM, VAL, ...) creates a set
         %   of options with the named parameters altered with the
         %   specified values.
         %
         %   OPTS = DRAMOptions(OLDOPTS, PARAM, VAL, ...)
         %   creates a copy of OLDOPTS with the named parameters altered
         %   with the specified value
         %
         %   Note to see the parameters, check the
         %   documentation page for DRAMOptions
         %
         % Parameters:
         % varargin:
         
         % adapted from SolverOptions
         
         if nargin > 0
            
            % Deal with the case where the first input to the
            % constructor is a amioptions/struct object.
            if isa(varargin{1},'DRAMOptions')
               if strcmp(class(varargin{1}),class(obj))
                  obj = varargin{1};
               else
                  % Get the properties from options object passed
                  % into the constructor.
                  thisProps = properties(obj);
                  % Set the common properties. Note that we
                  % investigated first finding the properties that
                  % are common to both objects and just looping over
                  % those. We found that in most cases this was no
                  % quicker than just looping over the properties of
                  % the object passed in.
                  for i = 1:length(thisProps)
                     try %#ok
                        % Try to set one of the properties of the
                        % old object in the new one.
                        obj.(thisProps{i}) = varargin{1}.(thisProps{i});
                     end
                  end
               end
               firstInputObj = true;
            elseif isstruct(varargin{1})
               fieldlist = fieldnames(varargin{1});
               for ifield = 1:length(fieldlist)
                  obj.(fieldlist{ifield}) = varargin{1}.(fieldlist{ifield});
               end
               firstInputObj = true;
            elseif isempty(varargin{1})
               firstInputObj = true;
            else
               firstInputObj = false;
            end
            
            % Extract the options that the caller of the constructor
            % wants to set.
            if firstInputObj
               pvPairs = varargin(2:end);
            else
               pvPairs = varargin;
            end
            
            % Loop through each param-value pair and just try to set
            % the option. When the option has been fully specified with
            % the correct case, this is fast. The catch clause deals
            % with partial matches or errors.
            haveCreatedInputParser = false;
            for i = 1:2:length(pvPairs)
               try
                  obj.(pvPairs{i}) = pvPairs{i+1};
               catch ME %#ok
                  
                  % Create the input parser if we haven't already. We
                  % do it here to avoid creating it if possible, as
                  % it is slow to set up.
                  if ~haveCreatedInputParser
                     ip = inputParser;
                     % Structures are currently not supported as
                     % an input to optimoptions. Setting the
                     % StructExpand property of the input parser to
                     % false, forces the parser to treat the
                     % structure as a single input and not a set of
                     % param-value pairs.
                     ip.StructExpand =  false;
                     % Get list of option names
                     allOptionNames = properties(obj);
                     for j = 1:length(allOptionNames)
                        % Just specify an empty default as we already have the
                        % defaults in the options object.
                        ip.addParameter(allOptionNames{j}, []);
                     end
                     haveCreatedInputParser = true;
                  end
                  
                  % Get the p-v pair to parse.
                  thisPair = pvPairs(i:min(i+1, length(pvPairs)));
                  ip.parse(thisPair{:});
                  
                  % Determine the option that was specified in p-v pairs.
                  % These options will now be matched even if only partially
                  % specified (by 13a). Now set the specified value in the
                  % options object.
                  optionSet = setdiff(allOptionNames, ip.UsingDefaults);
                  obj.(optionSet{1}) = ip.Results.(optionSet{1});
               end
            end
         end
      end
      
      function new = copy(this)
         % Creates a copy of the passed DRAMOptions instance
         new = feval(class(this));
         
         p = properties(this);
         for i = 1:length(p)
            new.(p{i}) = this.(p{i});
         end
      end
      
      %% Part for checking the correct setting of options
      function this = set.regFactor(this, value)
         if(isnumeric(value) && value > 0)
            this.regFactor = lower(value);
         else
            error(['Please specify a positive regularization factor for ill conditioned covariance'...
                ' matrices of the adapted proposal density, e.g. ' ...
                'PestoSamplingOptions.DRAM.regFactor = 1e-5']);
         end
      end
      
      function this = set.nTry(this, value)
         if(isnumeric(value) && value > 0)
            this.nTry = lower(value);
         else
            error(['Please specify a positive number of maximum tries, e.g. ' ...
                'PestoSamplingOptions.DRAM.nTry = 3']);
         end
      end     
      
      function this = set.verbosityMode(this, value)
         if (strcmp(value, 'text') || strcmp(value, 'silent') || strcmp(value, 'debug'))
            this.verbosityMode = lower(value);
         else
            error(['Please specify a verbosity mode, e.g. ' ...
                'PestoSamplingOptions.DRAM.verbosityMode = "text"']);
         end
      end           
 
      function this = set.adaptionInterval(this, value)
         if(isnumeric(value) && value > 0)
            this.adaptionInterval = lower(value);
         else
            error(['Please specify the adaption inteval, e.g. ' ...
                'PestoSamplingOptions.DRAM.adaptionInterval = 1']);
         end
      end        
      
   end
end
```
