## Supplementary material for "Mechanistic description of spatial processes using integrative modelling of noise-corrupted imaging data": PestoOptions.m

```
classdef PestoOptions < matlab.mixin.CustomDisplay
    % PestoOptions provides an option container to pass options to various
    % PESTO functions. Not all options are used by all functions, consult the respective function documentation for details.
    %
    % This file is based on AMICI amioptions.m (http://icb-dcm.github.io/AMICI/)
    
    properties        
        % <!-- General options -->
        
        % Perform calculations sequentially (''sequential'', default), or
        % in parallel (''parallel''). Parallel mode will speed-up the
        % calculations on multi-core systems, but requires the 
        % [MATLAB Parallel Computing Toolbox](https://mathworks.com/products/parallel-computing/) 
        % to be installed.
        comp_type = 'sequential';
                
        % Determine whether results are saved or not.
        % * false: results are not saved
        % * true: results are stored to an extra folder
        save = false;
        
        % Name of the folder in which results are stored. If no folder is 
        % provided, a random foldername is generated.
        foldername = strrep(datestr(now,31),' ','__'); 
        
        
        
        % <!-- Options for the objective function -->
        
        % Type of objective function provided: 'log-posterior' or 'negative log-posterior'
        % Tells the algorithm that log-posterior or log-likelihood are provided so it performs
        % a maximization of the objective function or that the negative
        % log-posterior or negative log-likelihood are provided so that
        % a minimization of the objective function is performed.
        obj_type = 'log-posterior';
        
        % Maximum number of outputs, the objective function can provide:
        % * 1 ... only objective value
        % * 2 ... objective value with gradient
        % * 3 ... objective value, gradient and Hessian
        %
        % (Missing values will be approximated by finite differences.)
        % Don't confuse this with the number of outputs from the objective
        % function that Pesto really calls! objOutNumber just tells Pesto,
        % with how many outputs it can call the objective function. Also,
        % optimization is still possible to be done without gradient 
        % information and without the Pesto FD routine.
        objOutNumber = 3;
        
        % Parameter inidices to be fixed, if any
        fixedParameters = [];
        
        % Values of fixed parameters, same sorting as indices, if any
        fixedParameterValues = [];
        
        % Number of data points used for parameter estimation, which should 
        % be used to compute the BIC (optional)
        nDatapoints = [];
        
        
        % <!-- Options concerning the output -->
        
        % Output mode of algorithm: 
        % * 'visual': plots showing the progress are generated
        % * 'text': optimization results for multi-start are printed on screen
        % * 'silent': no output during the multi-start local optimization
        % * 'debug': print extra debug information (only available in
        % certain functions
        mode = 'visual';
        
        % Figure handle in which results are printed. If no handle is 
        % provided, a new figure is used. TODO: move to plot options
        fh = [];
        
        % Plotting options of class PestoPlottingOptions.m
        plot_options = PestoPlottingOptions();
        
        
        
        % <!-- Options for getMultiStarts -->

        % Number of local optimizations.
        n_starts = 20;
        
        % vector of indices which starts should be performed.
        % default is 1:n_starts
        start_index = [];

        % log-likelihood / log-posterior threshold for initialization of 
        % optimization.
        init_threshold = -inf;
        
        % Which optimizer to use?
        % Current options: ['fmincon', 'meigo-ess', 'meigo-vns', 'pswarm']
        %
        % For 'meigo-ess' or 'meigo-vns', MEIGO
        % (http://gingproc.iim.csic.es/meigom.html) has to be installed 
        % separately.
        %
        % For 'pswarm' PSwarm (http://www.norg.uminho.pt/aivaz/pswarm/) has
        % to be installed separately
        localOptimizer = 'fmincon';
               
        % Options for the chosen local optimizer. Setting fmincon options as default local optimizer. See *help('fmincon')*
        %
        % MaxIter: fmincon default, necessary to be set for tracing
        %
        % Options for 'meigo-ess' are described in ess_kernel.m in the
        % MEIGO folder.
        localOptimizerOptions = optimset( ...
            'algorithm', 'interior-point',...
            'display', 'off', ...
            'GradObj', 'on', ...
            'MaxIter', 2000, ...
            'PrecondBandWidth', inf);
        
        % Method used to propose starting points for fmincon. Can be
        % * 'latin hypercube': latin hypercube sampling
        % * 'uniform': uniform random sampling
        % * 'user-supplied': user supplied function PestoOptions.init_fun
        proposal = 'latin hypercube'; 
        
        % determine whether objective function, parameter values and
        % computation time are stored over iterations
        % * false:  not saved
        % * true: stored in fields par_trace, fval_trace and time_trace
        trace = false;
        
        % determine whether intermediate results are stored every
        % 10 iterations
        % * false: not saved
        % * true: results are stored to an extra folder
        tempsave = false;
                         
        % clears the objective function before every multi-start.
        % * false: persistent variables are preserved.
        % * true: remove all temporary/persistent variables.
        % 
        % WHEN TRUE THIS OPTION REMOVES ALL OBJECTIVE FUNCTION BREAK POINTS
        resetobjective = false;
        
        
        
        % <!-- Options for getParameterProfiles -->
        
        % flag for profile calculation
        % * true: profiles are calculated
        % * false: profiles are not calculated
        calc_profiles = true;
        
        % parameter bounds (can be removed in a future release, since
        % parameters.min/max can be used instead)
        % * .P.min ... lower bound for profiling parameters, having same
        %   dimension as the parameter vector (default = parameters.min).
        % * .P.max ... lower bound for profiling parameters, having same
        %   dimension as the parameter vector (default = parameters.max).
        P = {};
                
        % index MAP - parameter vector starting from which the
        %       profile is calculated. This option is helpful if local
        %       optima are present.
        MAP_index = 1;
        
        % Minimal ratio down to which the profile is calculated
        R_min = 0.03;
        
        % Indices of the parameters for which the profile is calculated. 
        %
        % Default: profile_optim_index will be set to 1:parameters.number
        % if left empty
        parameter_index = [];
        
        % Indices of the parameters for which the profile is calculated by 
        % reoptimization. 
        profile_optim_index = [];
        
        % Indices of the parameters for which the profile is calculated by 
        % integration. 
        profile_integ_index = [];
        
        % How should profiles be computed (if no more precise options are 
        % set like profile_optim_index or profile_integ_index)? 
        % Possibilities: {'optimization' (default), 'integration', 'mixed'}
        profile_method = 'default';
        
        
        
        % <!-- Detailed options for profile optimization -->
        
        % Optimizer options for profile likelihood
        % *     .algorithm ... choice of algorithm
        % *         = 'interior-point' (default) 
        % *         = 'trust-region-reflective'
        % *         = 'active-set'
        % *     .display ... output of optimization process
        % *         = 'off' (default) ... no output
        % *         = 'iter' ... output for every optimization step
        % *     .MaxIter ... maximum of optimization steps
        % *     .GradObj ... are gradients provided?
        % *     .GradConstr ... do we have constraints?
        % *     .TolCon ... Tolerance for constraints
        profileReoptimizationOptions = optimset( ...
            'algorithm', 'interior-point', ...
            'display', 'off', ...
            'MaxIter', 400, ...
            'GradObj', 'on', ...
            'GradConstr', 'on', ...
            'TolCon', 1e-6 ...
            );
        
        % Maximal relative decrease of ratio allowed for two adjacent 
        % points in the profile (default = 0.10) if options.dJ = 0;
        dR_max = 0.10;
        
        % influences step size at small likelihood ratio values
        dJ = 0.5;
        
        % options for the generation fo the next profile point
        % *     .mode ... choice of proposal direction
        % *         = 'multi-dimensional' (default) ... all parameters are updated.
        % *             The direct is the same as between the last two points.
        % *         = 'one-dimensional' ... only parameter for which profile is
        % *             currently calculated is updated.
        % *     .guess = 1e-2 ... guess for initial update stepsize
        % *     .min = 1e-6 ... lower bound for update stepsize
        % *     .min = 1e2 ... upper bound for update stepsize
        % *     .update = 1.25 ... incremental change if stepsize is too large or
        % *         too small, must be > 1.
        options_getNextPoint = struct('mode', 'multi-dimensional', ...
            'guess', 1e-2, ...
            'min', 1e-6, ...
            'max', 1, ...
            'update', 1.25);
        
        
        
        % <!-- Detailed options for profile integration -->
        
        % Options for profile integration 
        % *     .type ... choice of ODE integrator
        % *         = 'ode113' (default) ... Adams-Bashf.-Solver by Matlab
        % *         = 'ode15s' ... BDF-Solver by Matlab
        % *         = 'ode45' ... Runge-Kutta-Solver by Matlab
        % *         = 'CVODE' ... BDF and AB-Solver by sundials (to be implemented!)
        % *     .algorithm ... choice of algorithm (CVODE only)
        % *         = 'Adams' (default) ... Adams-Bashford solver
        % *         = 'BDF' ... BDF solver
        % *     .nonlinSolver ... choice of nonlinear solver (CVODE only)
        % *         = 'Newton' (default)
        % *         = 'Functional'
        % *     .linSolver ... choice of linear solver (CVODE only)
        % *         = 'Dense' (default) ... solver for small problems
        % *         = 'Band' ... solver for bigger problems
        % *     .gamma ... Retraction factor
        % *     .eps ... regularization for poorly conditioned Hessian
        % *     .minCond ... Minimum condition number, when regularization is to be used 
        % *     .hessian ... how is the Hessian Matrix provided?
        % *         = 'user-supplied' (default) ... Hessian is 3rd output of ObjFun 
        % *         = 'bfgs' ... BFGS approximation to Hessian
        % *         = 'sr1' ... symmetric-rank 1 approximation to Hessian
        % *     .gradient ... is a gradient provided?        
        % *     .MinStep ... minimum step size of the solver
        % *     .MaxStep ... maximum step size of the solver
        % *     .MaxNumSteps ... maximum steps to be taken
        % *     .GradTol ... maximum remaining gradient to be tolerated
        % *     .RelTol ... maximum relative integration error
        % *     .AbsTol ... maximum absolute integration error
        solver = struct('type', 'ode113', ...
            'algorithm', 'Adams', ...
            'nonlinSolver', 'Newton', ...
            'linSolver', 'Dense', ...
            'gamma', 0, ...
            'eps', 1e-8, ...
            'minCond', 1e-10, ...
            'hessian', 'user-supplied', ... 
            'gradient', true, ...
            'MaxStep', 0.1, ...
            'MinStep', 1e-5, ...
            'MaxNumSteps', 1e5, ...
            'GradTol', 1, ...
            'RelTol', 1e-4, ...
            'AbsTol', 1e-6 ...
            );
        

        
        % <!-- Options for getPropertyProfiles -->
        
        % Tolance for the maximal distance of the list point 
        % the lower and upper bounds for the properties.
        boundary_tol = 1e-5;
        
        % Indices of the properties for which the profile is to be  
        % calculated (default = 1:properties.number, reoptimization only). 
        property_index = [];         

        % Set MCMC options by calling an PestoSamplingOptions Class object
        MCMC;

    end
    
    properties (Hidden)
    end
    
    methods
        function obj = PestoOptions(varargin)
            %PestoOptions Construct a new PestoOptions object
            %
            %   OPTS = PestoOptions() creates a set of options with each option set to its
            %   default value.
            %
            %   OPTS = PestoOptions(PARAM, VAL, ...) creates a set of options with the named
            %   parameters altered with the specified values.
            %
            %   OPTS = PestoOptions(OLDOPTS, PARAM, VAL, ...) creates a copy of OLDOPTS with
            %   the named parameters altered with the specified value
            %
            %   Note to see the parameters, check the
            %   documentation page for PestoOptions
            %
            % Parameters:
            %   varargin:
            
            % adapted from SolverOptions
                        
            if nargin > 0 
                
                % Deal with the case where the first input to the
                % constructor is a PestoOptions/struct object.
                if isa(varargin{1},'PestoOptions')
                    if strcmp(class(varargin{1}),class(obj))
                        obj = varargin{1};
                    else
                        % Get the properties from options object passed
                        % into the constructor.
                        thisProps = properties(obj);
                        % Set the common properties. Note that we
                        % investigated first finding the properties that
                        % are common to both objects and just looping over
                        % those. We found that in most cases this was no
                        % quicker than just looping over the properties of
                        % the object passed in.
                        for i = 1:length(thisProps)
                            try %#ok
                                % Try to set one of the properties of the
                                % old object in the new one.
                                obj.(thisProps{i}) = varargin{1}.(thisProps{i});
                            end
                        end
                    end
                    firstInputObj = true;
                elseif isstruct(varargin{1})
                    fieldlist = fieldnames(varargin{1});
                    for ifield = 1:length(fieldlist)
                        obj.(fieldlist{ifield}) = varargin{1}.(fieldlist{ifield});
                    end
                    firstInputObj = true;
                elseif isempty(varargin{1})
                    firstInputObj = true;
                else
                    firstInputObj = false;
                end
                
                % Extract the options that the caller of the constructor
                % wants to set.
                if firstInputObj
                    pvPairs = varargin(2:end);
                else
                    pvPairs = varargin;
                end
                
                % Loop through each param-value pair and just try to set
                % the option. When the option has been fully specified with
                % the correct case, this is fast. The catch clause deals
                % with partial matches or errors.
                haveCreatedInputParser = false;
                for i = 1:2:length(pvPairs)
                    try
                        obj.(pvPairs{i}) = pvPairs{i+1};
                    catch ME %#ok
                        
                        % Create the input parser if we haven't already. We
                        % do it here to avoid creating it if possible, as
                        % it is slow to set up.
                        if ~haveCreatedInputParser
                            ip = inputParser;
                            % Structures are currently not supported as
                            % an input to optimoptions. Setting the
                            % StructExpand property of the input parser to
                            % false, forces the parser to treat the
                            % structure as a single input and not a set of
                            % param-value pairs.
                            ip.StructExpand =  false;
                            % Get list of option names
                            allOptionNames = properties(obj);
                            for j = 1:length(allOptionNames)
                                % Just specify an empty default as we already have the
                                % defaults in the options object.
                                ip.addParameter(allOptionNames{j}, []);
                            end
                            haveCreatedInputParser = true;
                        end
                        
                        % Get the p-v pair to parse.
                        thisPair = pvPairs(i:min(i+1, length(pvPairs)));
                        ip.parse(thisPair{:});
                        
                        % Determine the option that was specified in p-v pairs.
                        % These options will now be matched even if only partially
                        % specified (by 13a). Now set the specified value in the
                        % options object.
                        optionSet = setdiff(allOptionNames, ip.UsingDefaults);
                        obj.(optionSet{1}) = ip.Results.(optionSet{1});
                    end
                end
            end      
            
            % Add required subclasses
            obj.MCMC = PestoSamplingOptions();
            
        end
        
        function new = copy(this)
            % Creates a copy of the passed PestoOptions instance
            new = feval(class(this));
            
            p = properties(this);
            for iProp = 1:length(p)
                new.(p{iProp}) = this.(p{iProp});
            end
        end

        % Part for checking the correct setting of options
        
        function this = set.obj_type(this, value)
            if(strcmpi(value, 'log-posterior') || strcmpi(value, 'negative log-posterior'))
                this.obj_type = lower(value);
            else
                error('PestoOptions.obj_type must be ''log-posterior'' or ''negative log-posterior''.');
            end
        end
        
        function this = set.comp_type(this, value)
            if(strcmpi(value, 'sequential') || strcmpi(value, 'parallel'))
                this.comp_type = lower(value);
            else
                error('PestoOptions.comp_type must be ''sequential'' or ''parallel''.');
            end
        end

        function this = set.n_starts(this, value)
            if(isnumeric(value) && floor(value) == value && value > 0)
                this.n_starts = value;
                this.start_index = [];
            else
                error('PestoOptions.n_starts must be a positive integer value.');
            end
        end
        
        function this = set.mode(this, value)
            this.mode = value;
            if (strcmp(value, 'visual') || strcmp(value, 'text') || strcmp(value, 'silent') || strcmp(value, 'debug'))
                this.mode = value;
            else
                error('PestoOptions.mode must be set to either "visual", "text", "silent" or "debug".');
            end
        end
        
        function this = set.proposal(this, value)
            if (strcmp(value, 'latin hypercube') || strcmp(value, 'uniform') || strcmp(value, 'user-supplied'))
                this.proposal = value;
            else
                error('PestoOptions.proposal must be set to either "latin hypercube", "uniform" or "user-supplied".');
            end
        end
        
        function this = set.save(this, value)
            if islogical(value)
                this.save = value;
            else
                error('PestoOptions.save must ba a logical value.');
            end
        end
        
        function this = set.tempsave(this, value)
            if islogical(value)
                this.tempsave = value;
            else
                error('PestoOptions.tempsave must ba a logical value.');
            end
        end
        
        function this = set.trace(this, value)
            if islogical(value)
                this.trace = value;
            else
                error('PestoOptions.trace must ba a logical value.');
            end
        end

        function this = set.calc_profiles(this, value)
            if islogical(value)
                this.calc_profiles = value;
            else
                error('PestoOptions.calc_profiles must ba a logical value.');
            end
        end
        
        function this = set.resetobjective(this, value)
            if islogical(value)
                this.resetobjective = value;
            else
                error('PestoOptions.resetobjective must ba a logical value.');
            end
        end
        
        function this = set.start_index(this, value)
            if isvector(value) || isempty(value)
                this.start_index = value;
            else
                error(['PestoOptions.start_index must ba a numeric vector.']);
            end
        end
        
        function this = set.parameter_index(this, value)
            if isvector(value) || isempty(value)
                this.parameter_index = value;
            else
                error(['PestoOptions.parameter_index must ba a numeric vector.']);
            end
        end
        
        function this = set.property_index(this, value)
            if isvector(value) || isempty(value)
                this.property_index = value;
            else
                error(['PestoOptions.property_index must ba a numeric vector.']);
            end
        end
        
        function this = set.MAP_index(this, value)
            if(isempty(value) || isnumeric(value) && floor(value) == value && value > 0)
                this.MAP_index = value;
            else
                error('PestoOptions.MAP_index must be a positive integer value.');
            end
        end
        
        function this = set.dR_max(this, value)
            if(isnumeric(value) && value >= 0 && value <= 1)
                this.dR_max = value;
            else
                error('PestoOptions.dR_max positive numeric value between 0 and 1.');
            end
        end

        function this = set.R_min(this, value)
            if(isnumeric(value) && value >= 0 && value <= 1)
                this.R_min = value;
            else
                error('PestoOptions.R_min positive numeric value between 0 and 1.');
            end
        end
        
        function this = set.localOptimizer(this, value)
            switch(value)
                case {'fmincon', 'meigo-ess', 'meigo-vns', 'pswarm', 'lsqnonlin', 'cs', 'dhc', 'bobyqa'}
                    this.localOptimizer = value;
                    if strcmp(value, 'pswarm')
                        this.localOptimizerOptions = PSwarm('defaults');
                    end
                otherwise
                    error('PestoOptions.localOptimizer only supports the following choices: fmincon, meigo-ess, meigo-vns, pswarm, hctt, cs, dhc, bobyqa.');
            end
        end

    end
end
```
