## Supplementary material for "Mechanistic description of spatial processes using integrative modelling of noise-corrupted imaging data": PestoPlottingOptions.m

```
% @file PestoPlottingOptions
% @brief A class for checking and holding information on optimization
% parameters

classdef PestoPlottingOptions < matlab.mixin.CustomDisplay
    % PestoPlottingOptions is a class for checking and holding information on optimization
    % parameters
    %
    % This file is based on AMICI amioptions.m (http://icb-dcm.github.io/AMICI/)
    
    properties
        % Title of PESTO-generated plots
        % * true: show
        % * false: don't show
        title = true;
        
        % Additional points to include in the plots, e.g. true
        % parameter in the case of test examples
        %
        % Struct with the following fields
        % * .par: n x m matrix of m additional points
        % * .col: color used for additional points (default = [0,0,0]).
        %                  This can also be a m x 3 matrix of colors.
        % * .ls: line style (default = '-')
        % * .lw: line width (default = 2)
        % * .m: marker style (default = 's')
        % * .ms: line width (default = 8)
        % * .name: name of legend entry (default = 'add. point')
        % * .property_MS: line width (default = 8).
        % * .logPost
        
        add_points = struct('par', [], ...
            'logPost', [], ...
            'col', [0,0.8,0], ...
            'ls', '-', ...
            'lw', 1, ...
            'm', 'd', ...
            'ms', 8, ...
            'name', 'add. point');
        
        % TODO: from plotmultistarts
        mark_constraint = false;
        
        % TODO from plotparameteruncertainty
        subplot_size_1D = [];
        
        % TODO
        subplot_indexing_1D = [];
        
        % TODO 
        labels = struct('y_always', true, ...
            'y_name', []);
        
        % Indicates whether plots are redrawn or whether something
        %  is added to the plot
        % * true: extension of plot
        % * false: new plot
        hold_on = false;
        
        % Way of choosing x limits for plotting
        % * 'dynamic': x limits depending on analysis results
        % * 'static': x limits depending on parameters.min and .max or on
        %          user-defined bound options.bounds.min and .max. The later are
        %          used if provided.
        interval = 'dynamic';
        
        % Draw bounds
        % * true: yes
        % * false: no
        draw_bounds = true;
        
        % Bounds used for visualization if options.interval = 'static'
        %
        % struct with 
        %  * .min: lower bound
        %  * .max: upper bound
        bounds = {};
        
        % Options for profile plots
        %
        % Struct with 
        % * .plot_type: plot type
        %   * = 0 (default if no profiles are provided) ... no plot
        %   * = 1 (default if profiles are provided) ... likelihood ratio
        %   * = 2 ... negative log-likelihood
        % * .col: color of profile lines (default: [1,0,0])
        % * .lw: line width of profile lines (default: 1.5)
        P = struct('plot_type', 1, ...
            'col', [1,0,0], ...
            'lw', 2, ...
            'name', 'P');
        
        % Options for sample plots
        % * .plot_type: plot type
        %   * = 0 (default if no samples are provided) ... no plot
        %   * = 1 (default if samples are provided) ... histogram
        %   * = 2 ... kernel-density estimates
        % * .col ... color of profile lines (default: [0.7,0.7,0.7])
        % * .hist_col ... color of histogram (default = [0.7,0.7,0.7])
        % * .bins ... number of histogram bins (default: 30)
        %   * = 'optimal' ... selection using Scott's rule
        %   * = 'conservative' ... selection using Scott's rule / 2
        %   * = N (with N being an integer) ... N bins
        % * .sp_col: color of scatter plot (default = [0.7,0.7,0.7])
        % * .sp_m: marker for scatter plot (default = '.')
        % * .sp_ms: marker size for scatter plot (default = 5)
        % * .name: name of legend entry (default = 'S')
        S = struct('plot_type', 0, ...
            'bins', 'conservative', ...
            'scaling', [], ...
            'hist_col',  [0.7,0.7,0.7], ...
            'sp_col', [0.7,0.7,0.7], ...
            'lin_col', [1,0,0], ...
            'lin_lw', 2, ...
            'sp_m', '.', ...
            'sp_ms', 5, ...
            'col', [1,0,0], 'lw', 2, ...
            'PT', struct('sp_m', '.', ...
                'sp_ms', 5, ...
                'lw', 1.5, ...
                'ind', [], ...
                'col', [], ...
                'plot_type', 0), ...
            'name', 'S');
        
        % Options for multi-start optimization plots
        %
        % Struct with:
        % * .plot_type: plot type
        %   * = 0 (default if no MS are provided) ... no plot
        %   * = 1 (default if MS are provided) ... likelihood ratio and
        %               position of optima above threshold
        %   * = 2 ... negative log-likelihood and position of optima
        %               above threshold
        % * .col: color of local optima (default: [1,0,0])
        % * .lw: line width of local optima (default: 1.5)
        % * .name_conv: name of legend entry (default = 'MS - conv.')
        % * .name_nconv: name of legend entry (default = 'MS - not conv.')
        % * .only_optimum: only optimum is plotted
        
        MS = struct('plot_type', 1, ...
            'col', [1,0,0], ...
            'lw' , 2, ...
            'name_conv', 'MS - conv.', ...
            'name_nconv', 'MS - not conv.', ...
            'only_optimum', false);
        
        % Options for distribution approximation plots
        % 
        % Struct with:
        % * .plot_type: plot type
        %   * = 0 (default if no MS are provided) ... no plot
        %   * = 1 (default if MS are provided) ... likelihood ratio
        %   * = 2 ... negative log-likelihood
        % * .col: color of approximation lines (default: [0,0,1])
        % * .lw: line width of approximation lines (default: 1.5)
        % * .sigma_level: sigma-level which is visualized (default = 2)
        % * .name: name of legend entry (default = 'P_{app}')
        
        A = struct('plot_type', 1, ...
            'col', [0,0,1], ...
            'lw', 2, ...
            'sigma_level', 2, ...
            'name', 'P_{app}');
        
        % Option if a user provided sampling initialization should be used
        % for plotting an approximation of the distribution
        %
        % * 'user-provided'
        % * 'multistart' (default)
        
        MCMC = 'multistart';
        
        % Options for boundary visualization
        % 
        % Struct with 
        % * .mark: marking of profile points which are on the boundary
        %   * = 0 ... no visualization
        %   * = 1 (default) ... indicates points which ar close to the
        %               boundaries in one or more dimensions.
        % * .eps: minimal distance from boundary for which points are
        %           consider to e close do the boundary (default = 1e-4). Note
        %               that a one-norm is used.
        boundary = struct('mark', true, ...
            'eps', 1e-4);
        
        % Options for confidence level plots
        % 
        % Struct with
        % * .plot_type: plot type
        %   * = 0 (default) ... no plot
        %   * = 1 ... likelihood ratio
        %   * = 2 ... negative log-likelihood
        % * .alpha: visualized confidence level (default = 0.95)
        % * .type: type of confidence interval
        %   * = 'point-wise' (default) ... point-wise confidence interval
        %   * = 'simultanous' ... point-wise confidence interval
        %   * = {'point-wise','simultanous'} ... both
        % * .col: color of profile lines (default: [0,0,0])
        % * .lw: line width of profile lines (default: 1.5)
        % * .name: name of legend entry (default = 'cut-off')
        
        CL = struct('plot_type', 0, ...
            'alpha', 0.95, ...
            'type', 'point-wise', ...
            'col', [0,0,0], ...
            'lw', 2,  ...
            'name', 'cut-off');
        
        % Options for the way to plot confidence intervals
        %
        % Either all confidence intervals of one method are plotted to one
        % window 'params', or the confidence intervals for one parameter 
        % from all methods are plotted to one window 'methods', or 
        % everthing is grouped together 'all'.
        
        group_CI_by = 'parprop';
        
        % Settings for 2D plot to position subplot axes.
        % 
        % Struct with:
        % * .b1 ... offset from left and bottom border (default = 0.15)
        % * .b2 ... offset from left and bottom border (default = 0.02)
        % * .r ... relative width of subplots (default = 0.95)
        op2D = struct('b1', 0.15, 'b2', 0.02, 'r', 0.95);
        
        % Legend options
        % * .color: background color (default = 'none').
        % * .box: legend outine (default = 'on').
        % * .orientation: orientation of list (default = 'vertical')
        
        legend = struct('color', 'none', ...
            'box', 'on', ...
            'orientation', 'vertical', ...
            'position', []);
        
        % Fontsize for labels
        % * .tick: fontsize for ticklabels (default = 12)
        fontsize = struct ('tick', 12);
        
        % figure handle for log-posterior trace plot
        fh_logPost_trace = [];
        
        % figure handle for parameter trace plots.
        fh_par_trace = [];
        
        % figure handle for the 1D parameter distribution plot.
        fh_par_dis_1D = [];

        % figure handle for the 2D parameter distribution plot.
        fh_par_dis_2D = [];
        
        % plot type
        plot_type = {'parameter','posterior'};
        
        % max
        n_max = 1e4;
    end
    
    properties (Hidden)
    end
    
    methods
    function obj = PestoPlottingOptions(varargin)
            %PestoPlottingOptions Construct a new PestoPlottingOptions object
            %
            %   OPTS = PestoPlottingOptions() creates a set of options with each option set to its
            %   default value.
            %
            %   OPTS = PestoPlottingOptions(PARAM, VAL, ...) creates a set of options with the named
            %   parameters altered with the specified values.
            %
            %   OPTS = PestoPlottingOptions(OLDOPTS, PARAM, VAL, ...) creates a copy of OLDOPTS with
            %   the named parameters altered with the specified value
            %
            %   Note to see the parameters, check the
            %   documentation page for PestoPlottingOptions
            %
            % Parameters:
            %   varargin:

            
            % adapted from SolverOptions
            
            if nargin > 0
                
                % Deal with the case where the first input to the
                % constructor is a struct object.
                if isa(varargin{1},'PestoPlottingOptions')
                    if strcmp(class(varargin{1}),class(obj))
                        obj = varargin{1};
                    else
                        % Get the properties from options object passed
                        % into the constructor.
                        thisProps = properties(obj);
                        % Set the common properties. Note that we
                        % investigated first finding the properties that
                        % are common to both objects and just looping over
                        % those. We found that in most cases this was no
                        % quicker than just looping over the properties of
                        % the object passed in.
                        for i = 1:length(thisProps)
                            try %#ok
                                % Try to set one of the properties of the
                                % old object in the new one.
                                obj.(thisProps{i}) = varargin{1}.(thisProps{i});
                            end
                        end
                    end
                    firstInputObj = true;
                elseif isstruct(varargin{1})
                    fieldlist = fieldnames(varargin{1});
                    for ifield = 1:length(fieldlist)
                        obj.(fieldlist{ifield}) = varargin{1}.(fieldlist{ifield});
                    end
                    firstInputObj = true;
                elseif isempty(varargin{1})
                    firstInputObj = true;
                else
                    firstInputObj = false;
                end
                
                % Extract the options that the caller of the constructor
                % wants to set.
                if firstInputObj
                    pvPairs = varargin(2:end);
                else
                    pvPairs = varargin;
                end
                
                % Loop through each param-value pair and just try to set
                % the option. When the option has been fully specified with
                % the correct case, this is fast. The catch clause deals
                % with partial matches or errors.
                haveCreatedInputParser = false;
                for i = 1:2:length(pvPairs)
                    try
                        obj.(pvPairs{i}) = pvPairs{i+1};
                    catch ME %#ok
                        
                        % Create the input parser if we haven't already. We
                        % do it here to avoid creating it if possible, as
                        % it is slow to set up.
                        if ~haveCreatedInputParser
                            ip = inputParser;
                            % Structures are currently not supported as
                            % an input to optimoptions. Setting the
                            % StructExpand property of the input parser to
                            % false, forces the parser to treat the
                            % structure as a single input and not a set of
                            % param-value pairs.
                            ip.StructExpand =  false;
                            % Get list of option names
                            allOptionNames = properties(obj);
                            for j = 1:length(allOptionNames)
                                % Just specify an empty default as we already have the
                                % defaults in the options object.
                                ip.addParameter(allOptionNames{j}, []);
                            end
                            haveCreatedInputParser = true;
                        end
                        
                        % Get the p-v pair to parse.
                        thisPair = pvPairs(i:min(i+1, length(pvPairs)));
                        ip.parse(thisPair{:});
                        
                        % Determine the option that was specified in p-v pairs.
                        % These options will now be matched even if only partially
                        % specified (by 13a). Now set the specified value in the
                        % options object.
                        optionSet = setdiff(allOptionNames, ip.UsingDefaults);
                        obj.(optionSet{1}) = ip.Results.(optionSet{1});
                    end
                end
            end
    end
    
    function new = copy(this)
        % Creates a copy of the passed PestoPlottingOptions instance

        new = feval(class(this));
        
        p = properties(this);
        for i = 1:length(p)
            new.(p{i}) = this.(p{i});
        end
    end

        %% Part for checking the correct setting of options
        
        function this = set.MCMC(this, value)
            if (strcmp(value, 'multistart') || strcmp(value, 'user-provided'))
                this.MCMC = value;
            else
                error('PestoOptions.MCMC must be set to either "multistart" or "user-provided".');
            end
        end
        
        function this = set.interval(this, value)
            if (strcmp(value, 'dynamic') || strcmp(value, 'static'))
                this.interval = value;
            else
                error('PestoOptions.interval must be set to either "dynamic" or "static".');
            end
        end

        function this = set.group_CI_by(this, value)
            if (strcmp(value, 'parprop') || strcmp(value, 'methods') || strcmp(value, 'all'))
                this.group_CI_by = value;
            else
                error('PestoOptions.group_CI_by must be set to either "parprop", "methods" or "all".');
            end
        end
        
        function this = set.n_max(this, value)
            if(isnumeric(value) && value > 0)
                this.n_max = value;
            else
                error('PestoOptions.n_max must be a positive number.');
            end
        end
        
        function this = set.title(this, value)
            if islogical(value)
                this.title = value;
            else
                error('PestoOptions.title must ba a logical value.');
            end
        end
        
        function this = set.draw_bounds(this, value)
            if islogical(value)
                this.draw_bounds = value;
            else
                error('PestoOptions.draw_bounds must ba a logical value.');
            end
        end
        
        function this = set.mark_constraint(this, value)
            if islogical(value)
                this.mark_constraint = value;
            else
                error('PestoOptions.mark_constraint must ba a logical value.');
            end
        end
        
        function this = set.hold_on(this, value)
            if islogical(value)
                this.hold_on = value;
            else
                error('PestoOptions.hold_on must ba a logical value.');
            end
        end
        
    end
end
```
