## Supplementary material for "Mechanistic description of spatial processes using integrative modelling of noise-corrupted imaging data": PestoSamplingOptions.m

```
classdef PestoSamplingOptions < matlab.mixin.CustomDisplay
   % PestoSamplingOptions provides an option container to pass options to
   % various PESTO functions. Not all options are used by all functions,
   % consult the respective function documentation for details.
   %
   % This file is based on AMICI amioptions.m (http://icb-dcm.github.io/AMICI/)
   
   properties
      % <!-- General options -->
      
      % Type of objective function provided:
      % 'log-posterior' (default) or 'negative log-posterior'
      %
      % Tells the algorithm that log-posterior or log-likelihood are
      % provided so it takes into account the corect sign for perfoming
      % all algorithms correctly.
      obj_type = 'log-posterior';
            
      % Sampling algorithm, can be 'PT' (parallel tempering), 'PHS',
      % (parallel hierarchical sampling), 'MALA' (Metropolis adjusted
      % Langevin algorithm), or 'DRAM' (delayed rejection adapted
      % Metropolis algorithm, only if the DRAM toolbox is installed).
      % Default value is 'PT'
      samplingAlgorithm = 'PT'
      
      % Number of iterations, integer (1e5 is not too high, e.g. 1e6 is a
      % morereasonable value for reliable results, but computationally
      % more intensive.
      nIterations = 1e5;
      
      % Initialization points for all chains. If the algorithm uses
      % multiple chains (as 'PT' and 'PHS'), one can specify multiple
      % theta0,
      % e.g.: opt.theta0 = repmat([0.1,1.5,-2.5,-0.5,0.4],opt.nTemps,1)';
      % If there is just one chain, please specify as
      % opt.theta0 = [1;2;3;4];
      % It is recommendet to set theta0 by taking into account the
      % results from a preceeding optimization (see Pesto examples).
      theta0 = [];
      
      % Initial covariance matrices for all chains.
      % Example for single-chain algorithms:
      % opt.sigma0 = 1e5 * diag(ones(1,5));
      % Example for multi-chain algorithms:
      % opt.sigma0 = repmat(1e5*diag(ones(1,5)),opt.nTemps,1);
      % It is recommendet to set sigma0 by taking into account the
      % results from a preceeding optimization.
      sigma0 = [];
      
      % Output mode for sampling algorithms (except DRAM, which has its
      % own format), can be chosen as 'visual', 'text', 'silent', or
      % 'debug'. Default: visual
      mode = 'visual';
      
      % Maximum number of outputs, the objective function can provide:
      % * 1 ... only objective value
      % * 2 ... objective value with gradient
      % * 3 ... objective value, gradient and Hessian (Default)
      %
      % Missing values will be approximated by finite differences.
      objOutNumber = 1;
      
      % Parallel Tempering Options, an instance of PTOptions
      PT;
      
      % Parallel Hierarchical Sampling options, an instance of PHSOptions
      PHS;
            
      % Metropolis Adaptive Langevin Algorithm options, an instance of
      % MALAOptions
      MALA;
      
      % Delayed Rejection Adaptive Metropolis options, an instance of
      % DRAMOptions
      DRAM;      
   end
      
   methods
      function obj = PestoSamplingOptions(varargin)
         % PestoSamplingOptions Construct a new PestoSamplingOptions object
         %
         %   OPTS = PestoSamplingOptions() creates a set of options with
         %   each option set to itsdefault value.
         %
         %   OPTS = PestoSamplingOptions(PARAM, VAL, ...) creates a set
         %   of options with the named parameters altered with the
         %   specified values.
         %
         %   OPTS = PestoSamplingOptions(OLDOPTS, PARAM, VAL, ...)
         %   creates a copy of OLDOPTS with the named parameters altered
         %   with the specified value
         %
         %   Note to see the parameters, check the
         %   documentation page for PestoSamplingOptions
         % Parameters:
         % varargin:

         % adapted from SolverOptions
         
         if nargin > 0
            
            % Deal with the case where the first input to the
            % constructor is a amioptions/struct object.
            if isa(varargin{1},'PestoSamplingOptions')
               if strcmp(class(varargin{1}),class(obj))
                  obj = varargin{1};
               else
                  % Get the properties from options object passed
                  % into the constructor.
                  thisProps = properties(obj);
                  % Set the common properties. Note that we
                  % investigated first finding the properties that
                  % are common to both objects and just looping over
                  % those. We found that in most cases this was no
                  % quicker than just looping over the properties of
                  % the object passed in.
                  for i = 1:length(thisProps)
                     try %#ok
                        % Try to set one of the properties of the
                        % old object in the new one.
                        obj.(thisProps{i}) = varargin{1}.(thisProps{i});
                     end
                  end
               end
               firstInputObj = true;
            elseif isstruct(varargin{1})
               fieldlist = fieldnames(varargin{1});
               for ifield = 1:length(fieldlist)
                  obj.(fieldlist{ifield}) = varargin{1}.(fieldlist{ifield});
               end
               firstInputObj = true;
            elseif isempty(varargin{1})
               firstInputObj = true;
            else
               firstInputObj = false;
            end
            
            % Extract the options that the caller of the constructor
            % wants to set.
            if firstInputObj
               pvPairs = varargin(2:end);
            else
               pvPairs = varargin;
            end
            
            % Loop through each param-value pair and just try to set
            % the option. When the option has been fully specified with
            % the correct case, this is fast. The catch clause deals
            % with partial matches or errors.
            haveCreatedInputParser = false;
            for i = 1:2:length(pvPairs)
               try
                  obj.(pvPairs{i}) = pvPairs{i+1};
               catch ME %#ok
                  
                  % Create the input parser if we haven't already. We
                  % do it here to avoid creating it if possible, as
                  % it is slow to set up.
                  if ~haveCreatedInputParser
                     ip = inputParser;
                     % Structures are currently not supported as
                     % an input to optimoptions. Setting the
                     % StructExpand property of the input parser to
                     % false, forces the parser to treat the
                     % structure as a single input and not a set of
                     % param-value pairs.
                     ip.StructExpand =  false;
                     % Get list of option names
                     allOptionNames = properties(obj);
                     for j = 1:length(allOptionNames)
                        % Just specify an empty default as we already have the
                        % defaults in the options object.
                        ip.addParameter(allOptionNames{j}, []);
                     end
                     haveCreatedInputParser = true;
                  end
                  
                  % Get the p-v pair to parse.
                  thisPair = pvPairs(i:min(i+1, length(pvPairs)));
                  ip.parse(thisPair{:});
                  
                  % Determine the option that was specified in p-v pairs.
                  % These options will now be matched even if only partially
                  % specified (by 13a). Now set the specified value in the
                  % options object.
                  optionSet = setdiff(allOptionNames, ip.UsingDefaults);
                  obj.(optionSet{1}) = ip.Results.(optionSet{1});
               end
            end
            
            % Add required subclasses
            obj.PT = PTOptions;
            
         end
      end
      
      function new = copy(this)
        % Creates a copy of the passed PestoSamplingOptions instance
         new = feval(class(this));
         
         p = properties(this);
         for i = 1:length(p)
            new.(p{i}) = this.(p{i});
         end
      end
      
      %% Part for checking the correct setting of options
      function this = set.obj_type(this, value)
         if(strcmpi(value, 'log-posterior') || strcmpi(value, 'negative log-posterior'))
            this.obj_type = lower(value);
         else
            error('PestoSamplingOptions.obj_type must be ''log-posterior'' or ''negative log-posterior''.');
         end
      end
      
      function this = set.mode(this, value)
         if (strcmp(value, 'visual') || strcmp(value, 'text') || strcmp(value, 'silent') || strcmp(value, 'debug'))
            this.mode = value;
         else
            error('PestoSamplingOptions.mode must be set to either "visual", "text", "silent" or "debug".');
         end
      end
            
      function this = set.nIterations(this, value)
         if (value == floor(value) && value > 0)
            this.nIterations = value;
         else
            error('Please enter the number of desired iterations as integer, e.g. opt.nIterations = 1e6.');
         end
      end
      
      function this = set.samplingAlgorithm(this, value)
         if ~isstr(value) || isempty(value)
            error('Please specify the algorithm which should be used, e.g. opt.samplingAlgorithm = ''PT''');
         end
         if (strcmp(value, 'MALA') || strcmp(value, 'DRAM') || strcmp(value, 'PT') || strcmp(value, 'PHS'))
            this.samplingAlgorithm = value;
            switch value
               case 'MALA'
                  this.MALA = MALAOptions();
                  this.DRAM = struct;
                  this.PT   = struct;
                  this.PHS  = struct;
               case 'DRAM'
                  this.MALA = struct;
                  this.DRAM = DRAMOptions();
                  this.PT   = struct;
                  this.PHS  = struct;
               case 'PT'
                  this.MALA = struct;
                  this.DRAM = struct;
                  this.PT   = PTOptions();
                  this.PHS  = struct;
               case 'PHS'
                  this.MALA = struct;
                  this.DRAM = struct;
                  this.PT   = struct;
                  this.PHS  = PHSOptions();
            end
         else
            error('You have entered an sampling algorithm which does not exist.')
         end
      end
      
      function this = set.objOutNumber(this, value)
         if value == floor(value) && ( value == 1 || value == 2 || value == 3 )
            this.objOutNumber = lower(value);
         else
            error(['Please enter wheter finite differences and Hessians opt.objOutNumber = 1' ...
               'or sensitivity based gradients and Hessians opt.objOutNumber = 3 should be used.']);
         end
      end
      
      function this = checkDependentDefaults(this, par)
          % checkDependentDefaults sets default values for sampling options which are problem-specific
          % and will be adapted to the problem properties in par.
          % Should be called providing the parameter struct for dependent
          % defaults as theta0. Does both, checking already set dependent
          % options and defaulting based on par if not set yet.
          %
          % Parameters:
          % par: parameters as passed getParameterSamples() to use for choosing default options
          %
          % Return values:
          % this: the updated PestoSamplingOptions instance
          
         if ~isempty(this.theta0)
            switch this.samplingAlgorithm
               case {'DRAM','MALA'}
                  if size(this.theta0,1) ~= par.number
                     error('Please make sure opt.theta0, the par.number are consistent.')
                  end
               case 'PT'
                  if size(this.theta0,1) ~= par.number || ...
                        (size(this.theta0,2) ~= this.PT.nTemps && size(this.theta0,2) ~= 1)
                     error('Please make sure opt.theta0, the par.number and opt.PT.nTemps are consistent.')
                  end
               case 'PHS'
                  if size(this.theta0,1) ~= par.number || ...
                        (size(this.theta0,2) ~= this.PHS.nChains && size(this.theta0,2) ~= 1)
                     error('Please make sure opt.theta0, the par.number and opt.PHS.nChains are consistent.')
                  end
            end
            
         else
            warning('No user-provided initial point found. Setting Initial points randomly.')
            switch this.samplingAlgorithm
               case {'DRAM','MALA'}
                  this.theta0 = bsxfun(@plus, par.min', ...
                     bsxfun(@times, par.max' - par.min', rand(par.number,1)))';
               case 'PT'
                  this.theta0 = bsxfun(@plus, par.min', ...
                     bsxfun(@times, par.max' - par.min', rand(par.number,this.PT.nTemps)))';
               case 'PHS'
                  this.theta0 = bsxfun(@plus, par.min', ...
                     bsxfun(@times, par.max' - par.min', rand(par.number,this.PHS.nChains)))';
            end
         end
         if ~isempty(this.sigma0)
            switch this.samplingAlgorithm
               case {'DRAM','MALA'}
                  if size(this.sigma0,1) ~= par.number || ...
                        size(this.sigma0,2) ~= par.number
                     error('Please make sure opt.sigma0, the par.number are consistent.')
                  end
               case 'PT'
                  if size(this.sigma0,1) ~= par.number || ...
                        size(this.sigma0,2) ~= par.number || ...
                        (size(this.sigma0,3) ~= this.PT.nTemps && size(this.sigma0,3) ~= 1)
                     error('Please make sure opt.sigma0, the par.number and opt.PT.nTemps are consistent.')
                  end
               case 'PHS'
                  if size(this.sigma0,1) ~= par.number || ...
                        size(this.sigma0,2) ~= par.number || ...
                        (size(this.sigma0,3) ~= this.PHS.nChains && size(this.sigma0,3) ~= 1)
                     error('Please make sure opt.sigma0, the par.number and opt.PHS.nChains are consistent.')
                  end
            end
         else
            warning('No user-provided initial covariance sigma0 found. Setting to diagonal matrix with small entries.')
            switch this.samplingAlgorithm
               case {'DRAM','MALA'}
                  this.sigma0 = 1e4 * diag(ones(1,5));
               case 'PT'
                  this.sigma0 = 1e4 * diag(ones(1,5));
               case 'PHS'
                  this.sigma0 = 1e4 * diag(ones(1,5));
            end
         end
      end
   end
end
```
