## Supplementary material for "Mechanistic description of spatial processes using integrative modelling of noise-corrupted imaging data": PHSOptions.m

```
classdef PHSOptions < matlab.mixin.CustomDisplay
   % PHSOptions provides an option container to set options for Parallel
   % Hierarchical Sampling (PHS) in PestoSamplingOptions.PHS .
   %
   % This file is based on AMICI amioptions.m (http://icb-dcm.github.io/AMICI/)
   
   properties      
      % Initial number of temperatures
      nChains = 10;
      
      % Parameter which controlls the adaption degeneration
      % velocity of the single-chain proposals.
      % Value between 0 and 1.
      % No adaption (classical Metropolis-Hastings) for 0.
      alpha = 0.51;

      % The higher the value the more it lowers the impact of early adaption steps.
      memoryLength = 1;
      
      % Regularization factor for ill conditioned covariance matrices of the adapted proposal density.
      % Regularization might happen if the eigenvalues of the covariance matrix strongly differ in order of
      % magnitude. In this case, the algorithm adds a small diag-matrix to the covariance matrix with elements
      % regFactor.
      regFactor = 1e-6;
      
      % The number of iterations before the chains start to swap. 
      % Might get used to ensure a proper local adaption of the single chains before exchanging them.
      trainingTime = 1;      
   end
   
   methods
      function obj = PHSOptions(varargin)
         % PHSOptions Construct a new PHSOptions object
         %
         %   OPHSS = PHSOptions() creates a set of options with
         %   each option set to itsdefault value.
         %
         %   OPHSS = PHSOptions(PARAM, VAL, ...) creates a set
         %   of options with the named parameters altered with the
         %   specified values.
         %
         %   OPHSS = PHSOptions(OLDOPHSS, PARAM, VAL, ...)
         %   creates a copy of OLDOPHSS with the named parameters altered
         %   with the specified value
         %
         %   Note to see the parameters, check the
         %   documentation page for PHSOptions
         % 
         % Parameters:
         %  varargin:
         % 
         % adapted from SolverOptions
         
         if nargin > 0
            
            % Deal with the case where the first input to the
            % constructor is a amioptions/struct object.
            if isa(varargin{1},'PHSOptions')
               if strcmp(class(varargin{1}),class(obj))
                  obj = varargin{1};
               else
                  % Get the properties from options object passed
                  % into the constructor.
                  thisProps = properties(obj);
                  % Set the common properties. Note that we
                  % investigated first finding the properties that
                  % are common to both objects and just looping over
                  % those. We found that in most cases this was no
                  % quicker than just looping over the properties of
                  % the object passed in.
                  for i = 1:length(thisProps)
                     try %#ok
                        % Try to set one of the properties of the
                        % old object in the new one.
                        obj.(thisProps{i}) = varargin{1}.(thisProps{i});
                     end
                  end
               end
               firstInputObj = true;
            elseif isstruct(varargin{1})
               fieldlist = fieldnames(varargin{1});
               for ifield = 1:length(fieldlist)
                  obj.(fieldlist{ifield}) = varargin{1}.(fieldlist{ifield});
               end
               firstInputObj = true;
            elseif isempty(varargin{1})
               firstInputObj = true;
            else
               firstInputObj = false;
            end
            
            % Extract the options that the caller of the constructor
            % wants to set.
            if firstInputObj
               pvPairs = varargin(2:end);
            else
               pvPairs = varargin;
            end
            
            % Loop through each param-value pair and just try to set
            % the option. When the option has been fully specified with
            % the correct case, this is fast. The catch clause deals
            % with partial matches or errors.
            haveCreatedInputParser = false;
            for i = 1:2:length(pvPairs)
               try
                  obj.(pvPairs{i}) = pvPairs{i+1};
               catch ME %#ok
                  
                  % Create the input parser if we haven't already. We
                  % do it here to avoid creating it if possible, as
                  % it is slow to set up.
                  if ~haveCreatedInputParser
                     ip = inputParser;
                     % Structures are currently not supported as
                     % an input to optimoptions. Setting the
                     % StructExpand property of the input parser to
                     % false, forces the parser to treat the
                     % structure as a single input and not a set of
                     % param-value pairs.
                     ip.StructExpand =  false;
                     % Get list of option names
                     allOptionNames = properties(obj);
                     for j = 1:length(allOptionNames)
                        % Just specify an empty default as we already have the
                        % defaults in the options object.
                        ip.addParameter(allOptionNames{j}, []);
                     end
                     haveCreatedInputParser = true;
                  end
                  
                  % Get the p-v pair to parse.
                  thisPair = pvPairs(i:min(i+1, length(pvPairs)));
                  ip.parse(thisPair{:});
                  
                  % Determine the option that was specified in p-v pairs.
                  % These options will now be matched even if only partially
                  % specified (by 13a). Now set the specified value in the
                  % options object.
                  optionSet = setdiff(allOptionNames, ip.UsingDefaults);
                  obj.(optionSet{1}) = ip.Results.(optionSet{1});
               end
            end
         end
      end
      
      function new = copy(this)
        % Creates a copy of the passed PHSOptions instance
         new = feval(class(this));
         
         p = properties(this);
         for i = 1:length(p)
            new.(p{i}) = this.(p{i});
         end
      end
      
      %% Part for checking the correct setting of options
      function this = set.regFactor(this, value)
         if(isnumeric(value) && value > 0)
            this.regFactor = lower(value);
         else
            error(['Please specify a positive regularization factor for ill conditioned covariance'...
                ' matrices of the adapted proposal density, e.g. ' ...
                'PestoSamplingOptions.PHS.regFactor = 1e-5']);
         end
      end
      
      function this = set.nChains(this, value)
         if(value == floor(value) && value > 0)
            this.nChains = lower(value);
         else
            error(['Please enter a positive integer for the number auxiliary chains,' ...
               'e.g. PestoSamplingOptions.PHS.nChains = 10.']);
         end
      end            

      function this = set.alpha(this, value)
         if(isnumeric(value) && value > 0.5 && value < 1)
            this.alpha = lower(value);
         else
            error(['Please an adaption decay constant between 0.5 and 1.0, e.g. PestoSamplingOptions.PHS.alpha = 0.51']);
         end
      end  
      
      function this = set.memoryLength(this, value)
         if(value == floor(value) && value > 0)
            this.memoryLength = lower(value);
         else
            error(['Please enter a positive interger memoryLength constant, '...
               'e.g. PestoSamplingOptions.PHS.memoryLength = 1']);
         end
      end   
      
      function this = set.trainingTime(this, value)
         if(value == floor(value) && value > 0)
            this.trainingTime = lower(value);
         else
            error(['Please enter a positive interger training time constant, '...
               'e.g. PestoSamplingOptions.PHS.trainingTime = 1']);
         end
      end         
      
   
            
   end
end
```
