## Supplementary material for "Mechanistic description of spatial processes using integrative modelling of noise-corrupted imaging data": collectResults.m

```
function obj = collectResults(foldername)
% collectResults() collects and plots the results stored in a common folder
%
% USAGE:
% [parameters] = collectResults(foldername)
%
% Parameters:
% foldername: Name of folder from which results are collected.
%
% Return values:
% parameters: parameter struct.
%
% History:
% 2014/06/12 Jan Hasenauer
%% Initialization
warning off;
try
    init = load(fullfile(foldername,'init'),'parameters');
    obj = init.parameters;
    type = 'parameters';
catch
    init = load(fullfile(foldername,'init'),'properties');
    obj = init.properties;
    type = 'properties';
end
warning on;

%% Collection of data
files = dir(fullfile(foldername,'*.csv'));

ntheta = size(init.parameters.MS.par0,1);
nstarts = init.parameters.MS.n_starts;

obj.MS.logPost = NaN(nstarts,1);
obj.MS.logPost0 = NaN(nstarts,1);
obj.MS.par = NaN(ntheta,nstarts);
obj.MS.par0 = NaN(ntheta,nstarts);
obj.MS.gradient = NaN(ntheta,nstarts);
obj.MS.hessian = NaN(ntheta,ntheta,nstarts);
obj.MS.t_cpu = NaN(nstarts,1);
obj.MS.n_objfun = NaN(nstarts,1);
obj.MS.n_iter = NaN(nstarts,1);
obj.MS.exitflag = NaN(nstarts,1);
% should do this for prop and prop_Sigma aswell, but what are the dimensions?

% Loop: files
for j = 1:length(files)
    % Read file
    v = csvread(fullfile(foldername,files(j).name));
    
    % Determine index and fieldname
    fn1 = files(j).name(1);
    fn2 = files(j).name((strfind(files(j).name,'__')+2):(length(files(j).name)-4));
    
    % Assignment
    switch fn1
        case 'M' % -> Multi-start optimization results
            i = str2num(files(j).name(3:(strfind(files(j).name,'__')-1)));
            switch fn2
                case 'logPost', obj.MS.logPost(i,1) = v;
                case 'logPost0', obj.MS.logPost0(i,1) = v;
                case 'par', obj.MS.par(:,i) = v;
                case 'par0', obj.MS.par0(:,i) = v;
                case 'gradient', obj.MS.gradient(:,i) = v;
                case 'hessian', obj.MS.hessian(:,:,i) = v;
                case 't_cpu', obj.MS.t_cpu(i,1) = v;
                case 'n_objfun', obj.MS.n_objfun(i,1) = v;
                case 'n_iter', obj.MS.n_iter(i,1) = v;
                case 'exitflag', obj.MS.exitflag(i,1) = v;
                case 'prop', obj.MS.prop(:,i) = v;
                case 'prop_Sigma', obj.MS.prop_Sigma(:,:,i) = v;
                case 'fval_trace'
                    if(~isfield(obj.MS,'fval_trace'))
                        obj.MS.fval_trace = NaN(size(v,1),nstarts);
                    end
                    obj.MS.fval_trace(:,i) = v;
                case 'time_trace'
                    if(~isfield(obj.MS,'time_trace'))
                        obj.MS.time_trace = NaN(size(v,1),nstarts);
                    end
                    obj.MS.time_trace(:,i) = v;
                case 'par_trace'
                    if(~isfield(obj.MS,'par_trace'))
                        obj.MS.par_trace = NaN(ntheta,size(v,2),nstarts);
                    end
                    obj.MS.par_trace(:,:,i) = v;
            end
        case 'P' % -> Profile calculation results
            i = str2num(files(j).name(2:(strfind(files(j).name,'__')-1)));
            switch fn2
                case 'par', obj.P(i).par = v;
                case 'logPost', obj.P(i).logPost = v;
                case 'R', obj.P(i).R = v;
                case 'exitflag', obj.P(i).exitflag = v;
                case 'prop', obj.P(i).prop = v;
            end
    end
end

%% Sort and save results
switch type
    case 'parameters'
        parameters = sortMultiStarts(obj);
        save(fullfile(foldername,['parameters_' foldername]),'parameters');
        obj = parameters;
end

%% Visualization
switch type
    case 'parameters'
        if isfield(obj,'MS')
            disp(['progress of MS = ' num2str(100*sum(~isnan(obj.MS.par(1,:)))/obj.MS.n_starts) '%']);
            plotMultiStarts(obj);
        end
        if isfield(obj,'P')
            plotParameterProfiles(obj);
        end
    case 'properties'
        if isfield(obj,'MS')
            plotPropertyMultiStarts(obj);
        end
        if isfield(obj,'P')
            plotPropertyProfiles(obj);
        end
end
```
