## Supplementary material for "Mechanistic description of spatial processes using integrative modelling of noise-corrupted imaging data": class_substitutes.c

```
/**
 * @class rowvec
 * @brief A matlab row vector.
 *
 * This class is an artificially created class in doxygen to allow more precise type declarations
 *
 * @class colvec
 * @brief A matlab column vector.
 *
 * This class is an artificially created class in doxygen to allow more precise type declarations
 *
 * @class integer
 * @brief An integer value
 *
 * This class is an artificially created class in doxygen to allow more precise type declarations.
 * Matlab types associated with this class are all int-types (int8, uint8 etc).
 *
 * @class double
 * @brief A double value
 *
 * This class is an artificially created class in doxygen to allow more precise type declarations.
 * The MatLab type associated with this class is double.
 * 
 * @class logical
 * @brief A boolean value
 *
 * This class can be seen as synonym for boolean values/flags used inside classes. In order to stick with
 * matlab conventions/datatypes, this class was named logical instead of bool or boolean.
 *
 * This class is an artificially created class in doxygen to allow more precise type declarations
 *
 * @class function_handle
 * @brief A MatLab function handle
 *
 * This class is an artificially created class in doxygen to allow more precise type declarations
 *
 * @class char
 * @brief A MatLab character array
 *
 * This class is an artificially created class in doxygen to allow more precise type declarations
 * and represents string-like types.
 *
 * @class cell
 * @brief A MatLab cell array or matrix
 *
 * This class is an artificially created class in doxygen to allow more precise type declarations
 *
 * @class struct
 * @brief A MatLab struct
 *
 * This class is an artificially created class in doxygen to allow more precise type declarations
 *
 * @class varargin
 * @brief A variable number of input arguments
 *
 * This class is an artificially created class in doxygen to allow more precise type declarations.
 *
 * For more information about the varargin argument see the
 * <a href="http://www.mathworks.de/help/techdoc/ref/varargin.html" target="_blank">
 * MatLab documentation on varargin</a>.
 *
 *
 * @class varargout
 * @brief A variable number of output arguments
 *
 * This class is an artificially created class in doxygen to allow more precise type declarations.
 *
 * For more information about the varargout argument see the
 * <a href="http://www.mathworks.de/help/techdoc/ref/varargout.html" target="_blank">
 * MatLab documentation on varargout</a>.
 */

class matrix {
    /**
     * @class matrix
     * @brief A matlab matrix
     *
     * This class is an artificially created class in doxygen to allow more precise type declarations
     */
};
class sparsematrix
    :public ::matrix {
    /**
     * @class sparsematrix
     * @brief A matlab sparse matrix
     *
     * This class is an artificially created class in doxygen to allow more precise type declarations
     */
};

class handle {
	/**
		@class handle
		@brief Matlab's base handle class (documentation generation substitute)

		As doxygen does not know the class "handle" from itself, many classes do not get rendered within the documentation and the correct root class is not even displayed.
		This workaround guarantees a correct (also graphical) representation of the class hierarchy.
     *  
     *  Note here that by having the type handle it could also mean to have a vector or matrix of handles.
     *
	*/
public:
/** @brief  Creates a listener for the specified event and assigns a callback function to execute when the event occurs.
  * @sa notify
 */
matlabtypesubstitute addlistener;


/**
	@brief Broadcast a notice that a specific event is occurring on a specified handle object or array of handle objects.
*/
matlabtypesubstitute notify;

/**
	@brief Handle object destructor method that is called when the object's lifecycle ends.
*/
matlabtypesubstitute delete;

/**
	@brief Handle object disp method which is called by the display method. See the MATLAB disp function.
*/
matlabtypesubstitute disp;

/**
	@brief Handle object display method called when MATLAB software interprets an expression returning a handle object that is not terminated by a semicolon. See the MATLAB display function.
*/
matlabtypesubstitute display;

/**
	@brief Finds objects matching the specified conditions from the input array of handle objects.
*/
matlabtypesubstitute findobj;

/**
	@brief Returns a meta.property objects associated with the specified property name.
*/
matlabtypesubstitute findprop;

/**
	@brief Returns a cell array of string containing the names of public properties.
*/
matlabtypesubstitute fields;

/**
	@brief Returns a cell array of string containing the names of public properties. See the MATLAB fieldnames function.
*/
matlabtypesubstitute fieldnames;

/**
	@brief Returns a logical array in which elements are true if the corresponding elements in the input array are valid handles.
	This method is Sealed so you cannot override it in a handle subclass.
*/
matlabtypesubstitute isvalid;

/**
	@brief Relational functions example. See details for more information.
	
	@par Other possible relational operators:
		-ne
		-lt
		-le
		-gt
		-ge

	Relational functions return a logical array of the same size as the pair of input handle object arrays. Comparisons use a number associated with each handle. You can assume that the same two handles will compare as equal and the repeated comparison of any two handles will yield the same result in the same MATLAB session. Different handles are always not-equal. The order of handles is purely arbitrary, but consistent.
*/
matlabtypesubstitute eq;

/**
	@brief Transposes the elements of the handle object array.
*/
matlabtypesubstitute transpose;

/**
	@brief Rearranges the dimensions of the handle object array. See the MATLAB permute function.
*/
matlabtypesubstitute permute;

/**
	@brief hanges the dimensions of the handle object array to the specified dimensions. See the MATLAB reshape function.
*/
matlabtypesubstitute reshape;

/**
	@brief ort the handle objects in any array in ascending or descending order. 

	The order of handles is purely arbitrary, but reproducible in a given MATLAB session. See the MATLAB sort function.
*/
matlabtypesubstitute sort;
}
```
