## Supplementary material for "Mechanistic description of spatial processes using integrative modelling of noise-corrupted imaging data": developers.c

```
/*
 * This file is included in order to give correct links to the developers of mtoc++ for the MatLabDocMaker,
 * if included in the documentation.
 *
 * If you are also making use of the @change/@new tags you will also have a similar project-specific development page.
 * However, then you might want to add an ingroup-doxygen tag here in order to neatly include this generated page at an appropriate place.
 * Unfortunately, the versions used in the MatLabDocMaker.m file naturally correspond to the
 * internal mtoc++-versions and will of course differ from those used in your project. If this bothers you all too much, feel free to
 * remove the corresponding tags from the MatlabDocMaker file, as probably wont come up with a hack for this too soon :-)
 */
/**
 * @page developers PESTO Developers
 * @section jh Jan Hasenauer
 * https://www.helmholtz-muenchen.de/icb/dcm
 *
 *- Lead Programmer
 */
```
