## Supplementary material for "Mechanistic description of spatial processes using integrative modelling of noise-corrupted imaging data": MatlabDocMaker.m

```
classdef MatlabDocMaker
% MatlabDocMaker: Automated documentation creation using doxygen and mtoc++ from within MatLab
%
% Currently documentation creation for unix and windows environment is
% supported.
%
% Prerequisites:
% The following tools must be installed and present on the PATH or
% reside inside the folder returned by
% MatlabDocMaker.getConfigDirectory.
% - \c mtocpp, \c mtocpp_post (the main tool)
% - \c doxygen (mtoc++ is a filter for doxygen)
%
% Strongly recommended:
% - \c latex Doxygen supports built-in latex formulas and
% MatlabDocMaker/mtoc++ allows for easy extra latex inclusions and
% notation in code
% - \c dot Doxygen creates really nice inheritance graphs and
% collaboration diagrams with dot.
%
% @author Daniel Wirtz @date 2011-10-13
%
% @change{1,5,dw,2013-12-03} Fixed default value selection for properties,
% now not having set a description or logo does not cause an error to be
% thrown.
% 
% @change{1,5,dw,2013-02-21} Fixed the callback for suggested direct documentation creation
% after MatlabDocMaker.setup (Thanks to Aurelien Queffurust)
%
% @change{1,5,dw,2013-02-12} Also added the escaping for the Logo file. Thanks to Chris Olien
% for the hint.
%
% @change{1,5,dw,2013-01-07} Included some backslash escaping for paths on windows platforms.
% Thanks to MathWorks Pilot Engineer '''Arvind Jayaraman''' for providing the feedback and code!
%
% @change{1,4,dw,2012-10-18} Removed \c m4 dependency and included constant properties for
% configuration file names.
%
% @new{1,4,dw,2012-10-16}
% - Added two more configuration variables "ProjectDescription" and "ProjectLogo" for easier
% configuration of the MatlabDocMaker in many cases. Thanks to Wolfgang
% Mennerich <http://www.mathworks.com/matlabcentral/fileexchange/authors/272859> for the suggestion.
% - Restructured the configuration, now only the project name function has to be implemented
% (the preferences tag depends on it, there might be more than one project within the same
% Matlab installation whos documentation is created using this tool). The rest can be provided
% either at setup time or later via suitable setters for the version, description and logo.
% - Automatically setting HaveDot in the doxygen config whenever its found on the environment
% path.
% - Added basic support for LaTeX documentation creation. Using the parameter 'latex'=true for
% the create method creates the LaTeX version of the documentation in a folder "latex" in the
% OutputDirectory (default behaviour)
%
% @change{1,4,dw,2012-09-27} Added automatic dot Graphviz tool detection on
% call to create.
%
% @change{1,3,dw,2012-02-16}
% - Now also collecting error messages from mtocpp_post and adding them to
% the warnings.log file.
% - Added the directive "LD_LIBRARY_PATH= " for unix systems, as MatLab
% sets it inside its executing environment. This can lead to errors if
% doxygen and/or mtoc++ have been built using never GLIBC (libstd) versions
% than the one shipped with MatLab.
%
% @change{1,3,dw,2012-01-16}
% - Properly using the correct file separators everywhere now
% - Hyperlinked the log file so it can be opened directly
%
% @change{1,3,dw,2012-01-14} Not displaying the "generated warnings"-text if there have not
% been any during documentation creation.
%
% @change{1,2,dw,2011-11-27}
% - Included documentation creation for the Windows platform and
% combined the old methods into one (small effective differences)
% - No longer storing the doxygen binary file in the prefs as a lot of
% tools must be present on the path anyways. The new paradigm is to
% expect all required 3rd-party programmes to be available on PATH. As
% backup the configuration files directory is
% added to the Matlab PATH environment \b nonpermanently and any
% executables found there will thus also be usable.
% - Included checks for \c dot and \c latex at the setup stage to
% recommend installation of those tools if not present (the default
% doxygen settings in Doxyfile.m4 are to use both)
%
% @change{1,2,dw,2011-11-08} Improved the createUnix method by displaying the warnings and writing the output to
% a log file afterwards. Not using cprintf anymore as this is 3rd party software.
%
% @change{1,2,dw,2011-11-07} Fixed a recursion bug caused by copy and paste. Now the preferences
% are stored on an per-application basis.
%
% @change{1,2,dw,2011-11-04} Changed the name to MatlabDocMaker in
% order to export it into the mtoc++ distribution later.
%
% @new{1,2,dw,2011-10-13} Added this class and moved documentation related
% stuff here from the KerMor class.
%
% This class is part of the mtoc++ tool
% - \c Homepage http://www.morepas.org/software/mtocpp/
% - \c License http://www.morepas.org/software/mtocpp/docs/licensing.html
%
% Copyright (c) 2012, Daniel Wirtz
% All rights reserved.
%
% Redistribution and use in source and binary forms, with or without
% modification, are permitted only in compliance with the BSD license, see
% http://www.opensource.org/licenses/bsd-license.php

    properties(Constant)
        % File name for the doxygen configuration file processed by the MatlabDocMaker.
        %
        % Assumed to reside in the MatlabDocMaker.getConfigDirectory
        %
        % @type char @default 'Doxyfile.template'
        DOXYFILE_TEMPLATE = 'Doxyfile.template';
        
        % File name for the latex extras style file processed by the MatlabDocMaker.
        %
        % Assumed to reside in the MatlabDocMaker.getConfigDirectory.
        % If not found, no latex extras are used.
        %
        % @type char @default 'latexextras.template'
        LATEXEXTRAS_TEMPLATE = 'latexextras.template';
        
        % File name the mtoc++ configuration file.
        %
        % Assumed to reside in the MatlabDocMaker.getConfigDirectory.
        % If not found, no special configuration is used.
        %
        % @type char @default 'mtocpp.conf'
        MTOCPP_CONFIGFILE = 'mtocpp.conf';
    end

    methods(Static)
        function name = getProjectName
            % Returns the project name.
            %
            % @note Changing the return value of this method will require
            % another execution of MatlabDocMaker.setup as the preferences
            % storage key also depends on it.
            %
            % Return values:
            % name: The project name @type char
            
            name = 'PESTO';
            % Example:
            % name = 'My mtoc++-powered project';
        end
    end
    
    %%%%%%%%%%%%%%%%%%%%%%%%%%%%%%%%%%%%%%%%%%%%%%%%%%%%%%%%%%%%%%%%%%%%%%
    %% End of user defined part.
    %%%%%%%%%%%%%%%%%%%%%%%%%%%%%%%%%%%%%%%%%%%%%%%%%%%%%%%%%%%%%%%%%%%%%%
    
    methods(Static, Sealed)
        function dir = getOutputDirectory
            % Returns the directory where the applications source files
            % reside
            %
            % Return values:
            % dir: The output directory @type char
            dir = MatlabDocMaker.getPref('outdir');
        end
        
        function dir = getSourceDirectory
            % Returns the directory where the applications source files
            % reside
            %
            % Return values:
            % dir: The project source directory @type char
            dir = MatlabDocMaker.getPref('srcdir');
        end
        
        function dir = getConfigDirectory
            % Returns the directory where the applications documentation
            % configuration files reside
            %
            % This folder must contain at least the files "mtoc.conf" and
            % "Doxyfile.template"
            %
            % Return values:
            % dir: The documentation configuration directory @type char
            dir = MatlabDocMaker.getPref('confdir');
        end
        
        function desc = getProjectDescription
            % Returns the short project description.
            %
            % Return values:
            % desc: The short project description @type char @default []
            %
            % See also: setProjectDescription
            desc =  MatlabDocMaker.getPref('proj_desc', '');
        end
        
        function setProjectDescription(value)
            % Sets the project description.
            %
            % Parameters:
            % value: The description @type char
            %
            % See also: getProjectDescription
            if ~ischar(value)
                error('The description must be a char array.');
            end
            MatlabDocMaker.setPref('proj_desc', value);
        end
        
        function version = getProjectVersion
            % Returns the current version of the project.
            %
            % @note The built-in @@new and @@change tags from the
            % Doxyfile.template support two-level versioning a la X.X.
            %
            % Return values:
            % version: The project version @type char @default []
            %
            % See also: setProjectVersion
            version = MatlabDocMaker.getPref('proj_ver', '0');
        end
        
        function setProjectVersion(value)
            % Sets the project version.
            %
            % Parameters:
            % value: The version string @type char
            %
            % See also: getProjectVersion
            if ~ischar(value)
                error('The project version must be a char array.');
            end
            MatlabDocMaker.setPref('proj_ver', value);
        end
        
        function fullPath = getProjectLogo
            % Returns the logo image file for the project. Either an absolute path or a plain
            % filename. For the latter case the image file is assumed to reside inside the
            % directory returned by MatlabDocMaker.getConfigDirectory.
            %
            % Return values:
            % logoFile: The projects logo image file. @type char @default []
            %
            % See also: setProjectLogo
            logoFile = MatlabDocMaker.getPref('proj_logo','');
            fullPath = '';
            if ~isempty(logoFile)
                if isempty(fileparts(logoFile))
                    fullPath = fullfile(MatlabDocMaker.getConfigDirectory,logoFile);
                else
                    fullPath = logoFile;
                end
                if exist(fullPath,'file') ~= 2
                    warning('MatlabDocMaker:getLogo',['Could not find logo file "%s".\n'...
                        'No logo will be shown in documentation output.'],logoFile);
                end
            end
        end
        
        function setProjectLogo(value)
            % Sets the project logo. Set to '' to unset.
            %
            % See the doxygen documentation for valid logo file types (wont be checked here).
            %
            % Parameters:
            % value: The logo file to use. Must be either an absolute path or a plain filename,
            % in which case the image is assumed to reside inside the
            % MatlabDocMaker.getConfigDirectory directory. @type char
            %
            % See also: getProjectLogo
            if nargin < 1
                [f, p] = uigetfile('*.*', 'Select project logo', pwd);
                if f ~= 0
                    value = fullfile(p,f);
                else
                    fprintf(2,'No file selected. Aborting.\n');
                    return;
                end
            end
            if ~ischar(value)
                error('The project logo file must be a char array.');
            end
            if ~isempty(value)
                fullPath = value;
                if isempty(fileparts(value))
                    fullPath = fullfile(MatlabDocMaker.getConfigDirectory,value);
                end
                if ~exist(fullPath,'file')
                    error('Invalid logo file: Could not find "%s"',fullPath);
                end
            end
            MatlabDocMaker.setPref('proj_logo', value);
        end
    end
        
    methods(Static)
        
        function open
            % Opens the generated documentation.
            %
            % Depending on the system's type the generated index.html is opened in the system's
            % default browser.
            index = fullfile(MatlabDocMaker.getOutputDirectory, 'index.html');
            if ispc
                winopen(index);
            else
                [s, m] = system(sprintf('xdg-open "%s"',index));
                if s ~= 0
                    fprintf(2,'Could not find/execute xdg-open: %s',m);
                    web(index);
                end
            end
        end
        
        function create(varargin)
            % Creates the Doxygen documentation
            %
            % Parameters:
            % varargin: Optional parameters for creation.
            % open: Set to true if the documentation should be opened after
            % successful compilation @type logical @default false
            % latex: Set to true if `\text{\LaTeX}` output should be generated, too. @type logical
            % @default false
           
            %% Preparations
            ip = inputParser;
            ip.addParamValue('open',false,@islogical);
            ip.addParamValue('latex',true,@islogical);
            ip.parse(varargin{:});
            genlatex = ip.Results.latex;
            
            % Check for correct setup
            cdir = MatlabDocMaker.getConfigDirectory;
            srcdir = MatlabDocMaker.getSourceDirectory;
            outdir = MatlabDocMaker.getOutputDirectory;
            % Check if doxygen config file template exists
            doxyfile_in = fullfile(cdir,MatlabDocMaker.DOXYFILE_TEMPLATE);
            if exist(doxyfile_in,'file') ~= 2
                error('No doxygen configuration file template found at "%s"',doxyfile_in);
            end
            
            lstr = '';
            if genlatex
                lstr = '(+Latex)';
            end
            fprintf(['Starting creation of doxygen/mtoc++ powered HTML%s documentation for "%s" (%s)\n'...
                'Sources: %s\nOutput to: <a href="matlab:MatlabDocMaker.open">%s</a>\nCreating config files...'],lstr,...
                MatlabDocMaker.getProjectName,MatlabDocMaker.getProjectVersion,...
                srcdir,outdir);
            
            % Operation-system dependent strings
            strs = struct;
            if isunix
                strs.null = '/dev/null';
                strs.silencer = '';
            elseif ispc
                strs.null = 'NUL';
                strs.silencer = '@'; % argh that took a while to remember..
            else
                error('Current platform not supported.');
            end
            
            % Save current working dir and change into the KerMor home
            % directory; only from there all classes and packages are
            % detected properly.
            curdir = pwd;
            cd(srcdir);
            
            % Append the configuration file directory to the current PATH
            pathadd = [pathsep cdir];
            setenv('PATH',[getenv('PATH') pathadd]);
            
            mtoc_conf = fullfile(cdir,MatlabDocMaker.MTOCPP_CONFIGFILE);
            filter = sprintf('%smtocpp',strs.silencer);
            if exist(mtoc_conf,'file')
                if isunix
                    strs.filter = 'mtocpp_filter.sh';
                    strs.farg = '$1';
                elseif ispc
                    strs.filter = 'mtocpp_filter.bat';
                    strs.farg = '%1';
                else
                    error('Current platform not supported.');
                end
                %% Creation part
                cdir = MatlabDocMaker.getConfigDirectory;
                % Create "configured" filter script for inclusion in doxygen 
                filter = fullfile(cdir,strs.filter);
                f = fopen(filter,'w');
                fprintf(f,'%smtocpp %s %s',strs.silencer,strs.farg,mtoc_conf);
                fclose(f);
                if isunix
                    unix(['chmod +x ' filter]);
                end
            end
            
            %% Prepare placeholders in the Doxyfile template
            m = {'_OutputDir_' strrep(outdir,'\','\\'); ...
                 '_SourceDir_' strrep(MatlabDocMaker.getSourceDirectory,'\','\\');...
                 '_ConfDir_' strrep(cdir,'\','\\');...
                 '_ProjectName_' MatlabDocMaker.getProjectName; ...
                 '_ProjectDescription_' MatlabDocMaker.getProjectDescription; ...
                 '_ProjectLogo_' strrep(MatlabDocMaker.getProjectLogo,'\','\\'); ...
                 '_ProjectVersion_' MatlabDocMaker.getProjectVersion; ...
                 '_MTOCFILTER_' strrep(filter,'\','\\'); ...
                 };
             
            % Check for latex extra stuff
            texin = fullfile(cdir,MatlabDocMaker.LATEXEXTRAS_TEMPLATE);
            latexextras = '';
            if exist(texin,'file') == 2  
                latexstr = strrep(fileread(texin),'_ConfDir_',strrep(cdir,'\','/'));
                latexextras = fullfile(cdir,'latexextras.sty');
                fid = fopen(latexextras,'w+'); fprintf(fid,'%s',latexstr); fclose(fid);
            end
            % Always use "/" for latex usepackage commands, so replace "\" (effectively windows
            % only) by "/", and dont pass the extension ".sty" as latex automatically adds it.
            m(end+1,:) = {'_LatexExtras_' strrep(latexextras(1:end-4),'\','/')};
            L = 'NO';
            if genlatex
                L = 'YES';
            end
            m(end+1,:) = {'_GenLatex_',L};
            
            % Check how to set the HAVE_DOT flag
            [s, ~] = system('dot -V');
            if s == 0
                HD = 'YES';
            else
                HD = 'NO';
                fprintf('no "dot" found...');
            end
            m(end+1,:) = {'_HaveDot_',HD};
            
            % Read, replace & write doxygen config file
            doxyfile = fullfile(cdir,'Doxyfile');
            doxyconfstr = regexprep(fileread(doxyfile_in),m(:,1),m(:,2));
            fid = fopen(doxyfile,'w'); fprintf(fid,'%s',doxyconfstr); fclose(fid);
            
            % Fix for unix systems where the MatLab installation uses older
            % GLIBSTD libraries than doxygen/mtoc++
            ldpath = '';
            if isunix
                setenv('PATH', [getenv('PATH'), ':/usr/local/mtocpp/bin/'])
                ldpath = 'LD_LIBRARY_PATH= ';
            end
            % Call doxygen
            fprintf('running doxygen with mtoc++ filter...');
            [~,warn] = system(sprintf('%sdoxygen "%s" 1>%s',ldpath, doxyfile, strs.null));
             
            % Postprocess
            fprintf('running mtoc++ postprocessor...');
            [~,postwarn] = system(sprintf('%smtocpp_post "%s" 1>%s',ldpath,...
                outdir, strs.null));
            if ~isempty(postwarn)
                warn = [warn sprintf('mtoc++ postprocessor messages:\n') postwarn];
            end
            
            % Create latex document if desired
            if genlatex
                oldd = pwd;
                latexerr = false;
                latexdir = fullfile(outdir,'latex');
                if exist(latexdir,'dir') == 7
                    if exist(fullfile(latexdir,'refman.tex'),'file') == 2
                        fprintf('compiling LaTeX output...');
                        cd(latexdir);
                        [s, latexmsg] = system('make');
                        if s ~= 0
                            warn = [warn sprintf('LaTeX compiler output:\n') latexmsg];
                            latexerr = true;
                        else
                            copyfile(fullfile(latexdir,'refman.pdf'), fullfile(fileparts(mfilename('fullpath')),'PESTO-doc.pdf'))
                        end
                    else
                        fprintf('cannot compile LaTeX output: no refman.tex found...');
                    end
                end
                cd(oldd);
            end
            
            % Tidy up
            fprintf('cleaning up...');
            if isfield(strs,'filter')
                % delete(filter);
            end
            if ~isempty(latexextras)
                % delete(latexextras);
            end
            % delete(doxyfile);
            
            %% Post generation phase 
            cd(curdir);
            % Restore PATH to previous value
            curpath = getenv('PATH');
            setenv('PATH',curpath(1:end-length(pathadd)));
            fprintf('done!\n');
            
            % Process warnings
            showchars = 800;
            warn = strtrim(warn);
            if ~isempty(warn)
                dispwarn = [warn(1:min(showchars,length(warn))) ' [...]'];
                fprintf('First %d characters of warnings generated during documentation creation:\n%s\n',showchars,dispwarn);
                % Write to log file
                log = fullfile(outdir,'warnings.log');
                f = fopen(log,'w'); fprintf(f,'%s',warn); fclose(f);
                fprintf(2,'MatlabDocMaker finished with warnings!\n');
                fprintf('Complete log file at <a href="matlab:edit(''%s'')">%s</a>.\n',log,log);
                % Check for latex log file
                log = fullfile(outdir,'_formulas.log');
                if exist(log,'file')
                    fprintf('Found LaTeX formula log file. Check <a href="matlab:edit(''%s'')">%s</a> for any errors.\n',log,log);
                end
                % Check for errors on latex generation
                if genlatex && latexerr
                    log = fullfile(latexdir,'refman.log');
                    fprintf('There have been errors with LaTeX compilation. See log file at <a href="matlab:edit(''%s'')">%s</a>.\n',log,log);
                end
            else
                fprintf('MatlabDocMaker finished with no warnings!\n');
            end
            
            % Open index.html if wanted
            if ip.Results.open
                MatlabDocMaker.open;
            end
        end
        
        function setup
            % Runs the setup script for MatlabDocMaker and collects all
            % necessary paths in order for the documentation creation to
            % work properly.
            
            %% Validity checks
            fprintf('<<<< Welcome to the MatlabDocMaker setup for your project "%s"! >>>>\n',MatlabDocMaker.getProjectName);
            
            %% Setup directories
            % Source directory
            srcdir = MatlabDocMaker.getPref('srcdir','');
            word = 'keep';
            if isempty(srcdir) || exist(srcdir,'dir') ~= 7
                srcdir = pwd;
                word = 'set';
            end
            str = sprintf('Do you want to %s %s as your projects source directory?\n(Y)es/(N)o?: ',word,strrep(srcdir,'\','\\'));
            ds = lower(input(str,'s'));
            if isequal(ds,'n')
                d = uigetdir(srcdir,'Please select the projects source directory');
                if d == 0
                    error('No source directory specified. Aborting setup.');
                end
                srcdir = d;
            end
            MatlabDocMaker.setPref('srcdir',srcdir);
            
            % Config directory
            confdir = MatlabDocMaker.getPref('confdir','');
            word = 'keep';
            if isempty(confdir) || exist(confdir,'dir') ~= 7
                confdir = fullfile(srcdir,'documentation');
                word = 'set';
            end
            str = sprintf('Do you want to %s %s as your documentation configuration files directory?\n(Y)es/(N)o?: ',word,strrep(confdir,'\','\\'));
            ds = lower(input(str,'s'));
            if isequal(ds,'n')
                d = uigetdir(confdir,'Please select the documentation configuration directory');
                if d == 0
                    error('No documentation configuration directory specified. Aborting setup.');
                end
                confdir = d;
            end
            MatlabDocMaker.setPref('confdir',confdir);
            
            % Output directory
            outdir = MatlabDocMaker.getPref('outdir','');
            word = 'keep';
            if isempty(outdir) || exist(outdir,'dir') ~= 7
                outdir = confdir;
                word = 'set';
            end
            str = sprintf('Do you want to %s %s as your documentation output directory?\n(Y)es/(N)o?: ',word,strrep(outdir,'\','\\'));
            ds = lower(input(str,'s'));
            if isequal(ds,'n')
                d = uigetdir(outdir,'Please select the documentation output directory');
                if d == 0
                    error('No documentation output directory specified. Aborting setup.');
                end
                outdir = d;
            end
            MatlabDocMaker.setPref('outdir',outdir);
            
            %% Additional Project properties
            if isequal(lower(input(['Do you want to specify further project details?\n'...
                    'You can set them later using provided set methods. (Y)es/(N)o?: '],'s')),'y')
                MatlabDocMaker.setPref('proj_ver',input('Please specify the project version, e.g. "0.1": ','s'));
                MatlabDocMaker.setPref('proj_desc',input('Please specify a short project description: ','s'));
                MatlabDocMaker.setProjectLogo;
            end
            
            %% Check for necessary and recommended tools
            hasall = true;
            setenv('PATH',[getenv('PATH') pathsep confdir]);
            fprintf('[Required] Checking for doxygen... ');
            [st, vers] = system('doxygen --version');
            if st == 0
                fprintf(' found %s\n',vers(1:end-1));
            else
                fprintf(2,' not found!\n');
                hasall = false;                
            end
            fprintf('[Required] Checking for mtoc++... ');
            ldpath = '';
            if isunix
                ldpath = 'LD_LIBRARY_PATH= ';
            end
            [st, vers] = system(sprintf('%smtocpp --version',ldpath));
            if st == 0
                fprintf(' found %s\n',vers(1:end-1));
            else
                fprintf(2,' not found!\n');
                hasall = false;
            end
            fprintf('[Recommended] Checking for dot... ');
            [st, vers] = system('dot -V');
            if st == 0
                fprintf(' found %s\n',vers(1:end-1));
            else
                fprintf(2,'dot Graphviz tool not found!\nInstall dot for nicer class diagrams.\n');
            end
            fprintf('[Recommended] Checking for latex... ');
            [st, vers] = system('latex --version');
            if st == 0
                fprintf(' found %s\n',vers(1:strfind(vers,sprintf('\n'))-1));
            else
                fprintf(2,'latex not found!\nA LaTeX installation available on the system path is strongly recommended with mtoc++.\n');
            end
            % Ghostscript
            fprintf('[Recommended] Checking for ghostscript... ');
            if ispc
                [st, vers] = system('gswin32c --version');
                if st ~= 0
                    [st, vers] = system('gswin64c --version');
                end
            else
                [st, vers] = system('gs --version');
            end
            if st == 0
                fprintf(' found %s\n',vers(1:strfind(vers,sprintf('\n'))-1));
            else
                fprintf(2,'ghostscript not found!\nCreating LaTeX formulas might not work properly.\n');
            end

            if ~hasall
                fprintf(2,'Please make sure all prerequisites can be found on PATH or copy the executables into %s.\n',confdir);
                fprintf('<<<< MatlabDocMaker setup finished with warnings. >>>>\n');
            else
                fprintf('<<<< MatlabDocMaker setup successful. >>>>\nYou can now create your projects documentation by running <a href="matlab:MatlabDocMaker.create(''open'',true)">MatlabDocMaker.create</a>!\n');
            end
        end
    end
    
    methods(Static, Access=private)
        function value = getProjPrefTag
            % Gets the tag for the MatLab preferences struct.
            % 
            % @change{0,7,dw,2013-04-02} Now also removing "~" and "-" characters from ProjectName tags for preferences.
            str = regexprep(strrep(strtrim(MatlabDocMaker.getProjectName),' ','_'),'[^\d\w]','');
            value = sprintf('MatlabDocMaker_on_%s',str);
        end
        
        function value = getPref(name, default)
            if nargin < 2
                def = [];
            else 
                def = default;
            end
            value = getpref(MatlabDocMaker.getProjPrefTag,name,def);
            if nargin < 2 && isempty(value)
                error('MatlabDocMaker preferences not found/set correctly. (Re-)Run the MatlabDocMaker.setup method.');
            end
        end
        
        function value = setPref(name, value)
            setpref(MatlabDocMaker.getProjPrefTag,name,value);
        end
    end
end
```
