## Supplementary material for "Mechanistic description of spatial processes using integrative modelling of noise-corrupted imaging data": PESTO-doc.pdf

1.0

Generated by Doxygen 1.8.11

Index 97

### 1 PESTO Documentation

#### 1.1 Introduction

Computational models are commonly used in diverse disciplines such as computational biology, engineering, or meteorology. The parameterization of the these models is usually based on measurements or observations. The process of inferring model parameters from such data is called model calibration or parameter estimation. This parameter estimation is often not straightforward due to non-linearities in the model equations or due to the mere size and the resulting computational challenges. Therefore, efficient algorithms are required to provide robust results within acceptable time.

PESTO is a freely available Parameter EStimation TOolbox for MATLAB (MathWorks) implementing a number of state-of-the-art algorithms for parameter estimation. It provides the following features, which are explained in more detail below:

- Parameter estimation from measurement data by global optimization based on multi-start local optimization (some algorithms require the `MATLAB Optimization Toolbox`)
- Parameter sampling using Markov Chain Monte Carlo (MCMC) algorithms
- Uncertainty analysis based on local approximations, parameter samples and profile-likelihood analysis (some algorithms require the `MATLAB Optimization Toolbox`)
- Visualization routines for all analyses
- Parallel processing (requires `MATLAB Parallel Computing Toolbox`)
- ...

PESTO functions can be applied to any user-provided formulation of an optimization problem with an objective function that can be evaluated in MATLAB. Besides the objective function, upper and lower bounds for the function parameters need to be specified.

#### 1.2 Availability

PESTO can be freely obtained from `https://github.com/ICB-DCM/PESTO/` by downloading the zip archive at `https://github.com/ICB-DCM/PESTO/archive/master.zip` or cloning the `git` repository via

```
git clone git@github.com:ICB-DCM/PESTO.git
```

#### 1.3 Installation

If the zip archive was downloaded, it needs to be unzipped and the main folder has to be added to the MATLAB search path (non-recursively).

If the repository was cloned, the main folder needs to be added to the MATLAB search path (non-recursively).

*Note:* Detailed instructions on how to modify your MATLAB search path are provided on the `web`.

###### Third-party packages

PESTO provides an interfaces to several other toolboxes which are not included in the PESTO archive:

- PSwarm (optimizer): http://www.norg.uminho.pt/aivaz/pswarm/
- MEIGO (optimizer): http://gingproc.iim.csic.es/meigo.html
- DRAM (MCMC): http://helios.fmi.fi/~lainema/dram/

To use their functionality, these toolboxes have to be installed separately. Please consult the respective user manuals for details.

#### 1.4 Licensing

See `LICENSE` file in the PESTO source directory.

#### 1.5 How to cite

This section will be updated upon publication of PESTO.

#### 1.6 Code organization

The end-user interface is provided by the MATLAB functions and classes in the top-level directory. PESTO example applications are provided in `/examples/`. All other folders only contain files used internally in PESTO.

#### 1.7 Features

PESTO implements a number of state-of-the-art algorithms related to parameter estimation. The main features are described below. Various examples demonstrate their application.

###### Notations and Terminology

Since most of the examples use analytical approaches for computing the gradient of the respective objective function, which quantifies the deviation of the fit for the current model parameters from the actual measurement data, the usage of the term ‚sensitivity analysis‘ may be misleading. In our context, ‚sensitivity analysis‘ is used in the context of ODE or PDE models and describes the sensitivity of the ODE/PDE state with respect to the model parameters. Those state sensitivities can be implemented in the ODE/PDE system and then used for an analytical calculation of the sensitivity of the objective function. This objective functions sensitivity will always be called the objective function gradient in our context. Finally, the behavior of the objective function by the variation of single parameters in order to find possible (non-)identifiabilities will always be referred to as ‚uncertainty analysis‘.

##### 1.7.1 Global optimization

Non-linear optimization problems like those in parameter estimation problems tend to have multiple optima. Usually, nothing is known beforehand about their number or their location, but the user is interested in finding the global optimum. There are different techniques for this kind of problem. PESTO provides a multi-start local optimization framework and provides an interface to two global optimizers.

###### Multi-start local optimization

Multi-start local optimization has turned out to be a very efficient method for "global optimization": Here, random points from across the parameter space are chosen as starting points for local optimization. If an adequate number of starting points spanning the domain of interest of the parameter space is selected, the lowest/highest minimum/maximum is accepted to be the global minimum/maximum. By default, `fmincon` from the MATLAB optimization toolbox is used as a local solver.

Multi-start local optimization functionality is provided by getMultiStarts.m, getPropertyMultiStarts.m and the respective plotting routines plotMultiStarts.m and plotPropertyMultiStarts.m. See examples/conversion_reaction/main↵ConversionReaction.m for an example.

###### Global optimizers

PESTO provides an interface to `PSwarm` and `MEIGO`. Once these toolboxes have been installed - they are not included in the PESTO archive - they can be used for parameter estimation. These optimizers are also accessed via getMultiStarts.m by setting PestoOptions::localOptimizer and PestoOptions::localOptimizerOptions accordingly. In principle, a single optimizer run (PestoOptions::n_starts = 1) should be enough for these global optimizers.

An example of both global optimizers is included in mainConversionReaction.m, another example for PSwarm in mainTransfection.m and an additional example for MEIGO can be found in mainEnzymaticCatalysis.m.

##### 1.7.2 Uncertainty analysis

When parameters are inferred from measurement data, the deviation of the data from the fit for the best parameter guess is usually assumed to be of stochastic nature. This means that the estimated parameters themselves are stochastic and underly an uncertainty. This can be quantified by performing uncertainty analysis and computing confidence intervals.

The easiest way to do this is using local approximations (based on the Hessian matrix of the objective function) at the best parameter guess. From those approximations, either threshold-based or mass-based methods can be used to compute confidence intervals for the inferred parameters. Another more reliable approach uses sampling based methods in combination with local approximations. It is described in more detail in the next section.

The probably most reliable way to compute confidence intervals is a third approach, based on profile likelihoods. There are two different ways to compute them: In the classical optimization based approach, each model parameter is varied separately while the others are constantly reoptimized. In this way one finds profiles for every parameter. A more recent approach uses an ODE, which follows the path in parameter space which describes the profile. After the profile computation, one fixes a confidence level using the inverse chi-squared-distribution to get a threshold. This threshold, together with the profile likelihood, gives reliable confidence intervals for each parameter. In this way, non-identifiable parameters can be detected.

Those functionalities are provided in getParameterProfiles.m, getPropertyProfiles.m (for the profile likelihoods), getParameterConfidenceIntervals.m and getPropertyConfidenceIntervals.m (for the confidence intervals). In order to get confidence intervals based on local approximations or sampling methods, one needs to run the routines get↵MultiStarts.m / getPropertyMultiStarts.m or getParameterSamples.m / getPropertySamples.m first. The respective visualization routines are plotParameterProfiles.m and plotPropertyProfiles.m. Examples for profile likelihoods can be found in mainConversionReaction.m and mainExampleRing.m (optimization based), mainEnzymaticCatalysis.m (integration based) and mainTransfection.m (a mixture of both).

##### 1.7.3 Parameter sampling

PESTO provides Markov Chain Monte Carlo (MCMC) algorithms for sampling the posterior distribution. Currently, two multi-chain methods (which can also be used in single-chain mode) and two additional single-chain methods can be chosen. The two multi-chain methods are `parallel tempering (PT)`, which is based on an adaptive version of the `Metropolis-Hastings algorithm (MH)`, and `parallel hierarchical sampling (PHS)`. For further single-chain sampling, also `Metropolis-adjusted Langevin algorithm (MALA)` can be chosen. Additionally, PESTO provides an interface to the `Delayed Rejection Adaptive Metropolis (DRAM)` toolbox.

Details on the recommended settings can be found in PestoSamplingOptions.m and the corresponding option files for each algorithm. Examples for the algorithms can be found in mainConversionReaction.m (PT as single-chain algorithm), mainEnzymaticCatalysis.m (PT as multi-chain algorithm), mainTransfection.m (PHS as multi-chain algorithm) and mainExampleRing.m and mainExampleGauss.m (all algorithms, respectively).

##### 1.7.4 Plotting

An integral part of PESTO are its highly customizable plotting functions for each type of analysis.

Details are provided in the documentation of the specific plotting functions:

- plotMultiStarts.m
- plotParameterProfiles.m
- plotParameterSamples.m
- plotParameterUncertainty.m
- plotPropertyMultiStarts.m
- plotPropertyProfiles.m
- plotPropertySamples.m
- plotPropertyUncertainty.m

Here some examples:

Plot of model fit using plotMultiStarts.m:

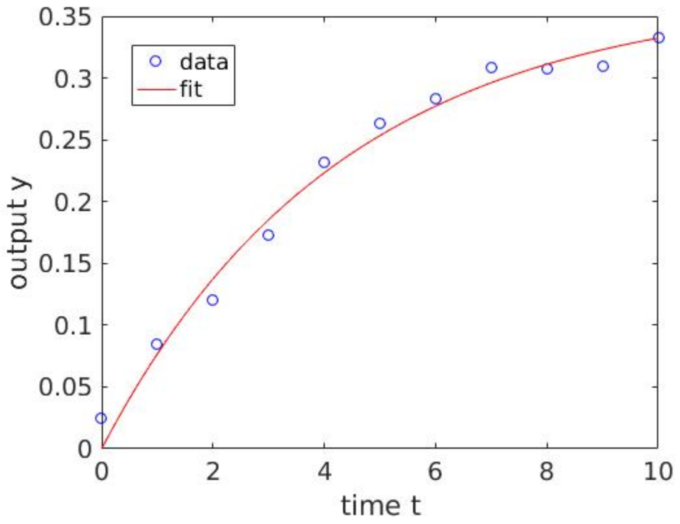

**Figure 1 Plot of model fit using plotMultiStarts.m**

Plot of different variants of parameter confidence intervals using plotParameterUncertainty.m:

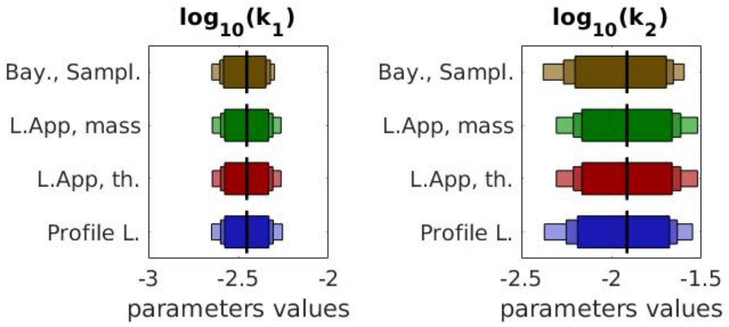

**Figure 2 Plot of different variants of parameter confidence intervals using plotParameterUncertainty.m**

2D plot of parameter samples using plotParameterSamples.m:

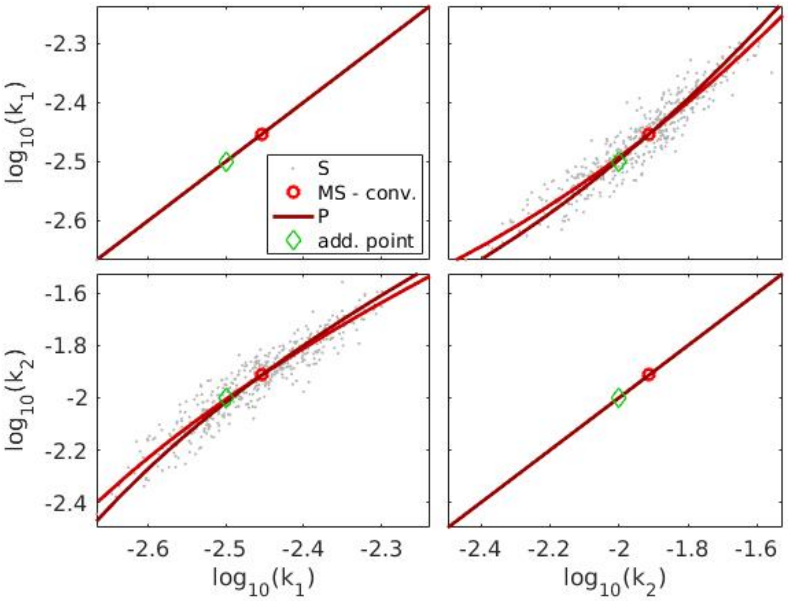

**Figure 3 2D plot of parameter samples using plotParameterSamples.m**

Plot of parameter samples using plotParameterSamples.m:

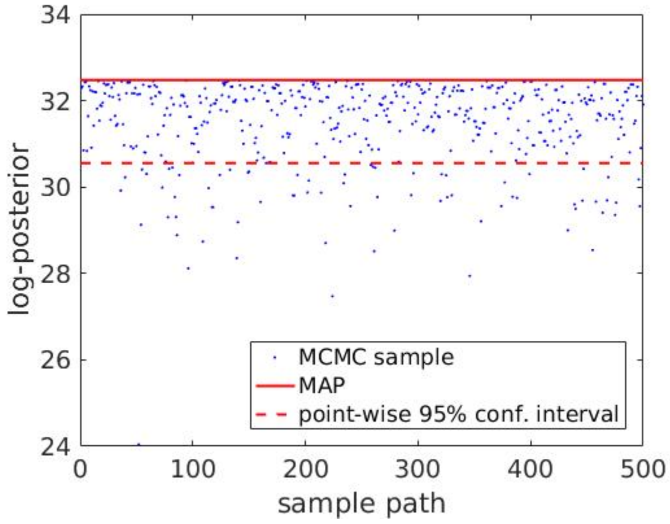

**Figure 4 Plot of parameter samples using plotParameterSamples.m**

Plot of property samples using plotPropertySamples.m:

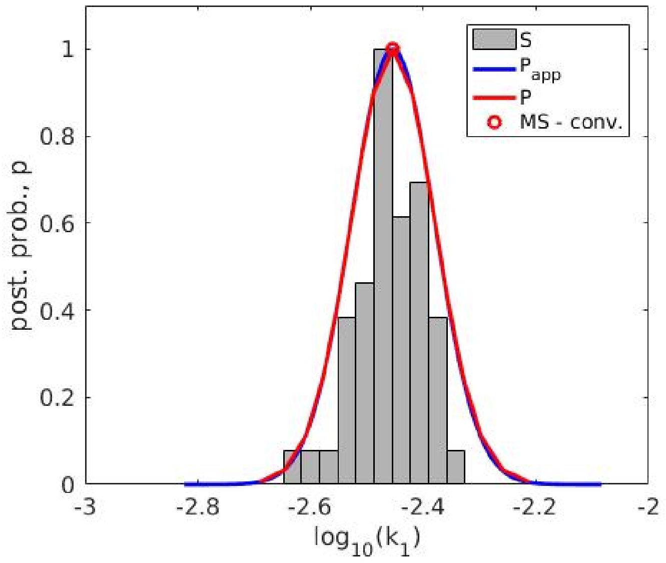

**Figure 5 Plot of property samples using plotPropertySamples.m**

See mainConversionReaction.m for live examples.

##### 1.7.5 Properties

The above-mentioned methods for parameter estimation, confidence intervals, parameter profiles and parameter samples (getMultiStarts.m, getParameterConfidenceIntervals.m, getParameterProfiles.m, getParameterSamples.↩ m) all operate on the objective function parameters directly. However, sometimes not the parameters themselves, but some function thereof is of interest. To this end, PESTO provides a simple interface to achieve this without having to change the objective function. Arbitrary user-provided functions which take the objective function parameter vector as an argument are referred to as 'properties'. After having used any of the getParameter∗.m functions, the respective getProperty∗.m function can be called, to evaluate a user-provided property function with the parameters values/samples/confidences obtained from the getParameter∗.m functions.

The following functions are available to analyze and plot properties:

- getPropertyConfidenceIntervals.m
- getPropertyMultiStarts.m
- getPropertyProfiles.m
- getPropertySamples.m
- plotPropertyMultiStarts.m
- plotPropertyProfiles.m
- plotPropertySamples.m
- plotPropertyUncertainty.m

See mainConversionReaction.m for examples.

### 2 Examples

#### 2.1 Examples

##### 2.1.1 Overview

PESTO comes with a number of ready-to-run examples to demonstrate its usage and functionality. The examples mostly stem from computational biology and comprise ordinary and partial differential equation (ODE / PDE), and Gaussian mixture models. More background information is provided in the respective example folder in the `main*.m` script. The three first example files toghether demonstrate basically all features and tools of PESTO.

The following examples are included:

- *Conversion reaction*, the simplest example and demonstrating most of the PESTO features (`examples/conversion_reaction/mainConversionReaction.m`, ODE)
- *mRNA transfection*, another very simple example, also demonstrating many of the PESTO features (`examples/mRNA_transfection/mainTransfection.m`, ODE)
- *Enzymatic catalysis*, the third small example case, also demonstrating a variety of the PESTO features (`examples/enzymatic_catalysis/mainEnzymaticCatalysis.m`, ODE)
- *Gaussian mixture*, a benchmark model for the MCMC sampling routines in PESTO (`examples/GaussExample/mainExampleGauss.m`)
- *Hyperring*, a second benchmark model (with non-identifiabilities) for the MCMC sampling routines in PESTO, (`examples/RingExample/mainExampleRing.m`)
- *Pom1p gradient formation*, a PDE problem which is discretized in space and reformulated as ODE § (`examples/Pom1p_gradient_formation/mainPom1.m`, PDE)
- *Jak-Stat-signaling*, a challenging example for the optimization routines in PESTO, § (`examples/jakstat_signaling/mainJakstatSignaling.m`, ODE)
- *ERBB signaling*, largest model among the examples § (`examples/erbb_signaling/mainErbBSignaling.m`, ODE)

§ These models require the freely available AMICI toolbox to run (http://icb-dcm.github.io/AMICI/).

The following table provides an overview of which of the PESTO functions are demonstrated in the different examples:

| | conversion reaction | enzymatic catalysis | Gaussian mixture | Hyperring | transfectior | Pom1p | JakStat | ErbB |
|---|---|---|---|---|---|---|---|---|
| getMultiStarts() | X | X | | X | X | X | X | X |
| plotMultiStarts() | X | X | | X | X | X | X | X |
| plotMultiStartDiagnosis() | | X | | | | | | |
| getParameterProfiles() | X | X | | X | X | | | |
| plotParameterProfiles() | X | X | | X | X | | | |
| getParameterSamples() | X | X | X | X | X | | | |
| plotParameterSamples() | X | X | X | X | X | | | |
| plotMCMCdiagnosis() | | X | X | | | | | |
| getParameterConfidenceIntervals() | X | X | | X | X | | | |
| plotConfidenceIntervals() | X | X | | X | X | | | |
| getPropertyMultiStarts() | X | | | X | X | | | |
| plotPropertyMultiStarts() | X | | | X | X | | | |
| getPropertyProfiles() | X | | | | X | | | |
| plotPropertyProfiles() | X | | | | X | | | |
| getPropertySamples() | X | | | X | X | | | |
| plotPropertySamples() | X | | | X | X | | | |
| getPropertyConfidenceIntervals() | X | | | | X | | | |

| | conversion reaction | enzymatic catalysis | Gaussian mixture | Hyperring | transfection | Pom1p | JakStat | ErbB |
|---|---|---|---|---|---|---|---|---|
| test←Gradient() | | | | | | | | |
| collect←Results() | | | | | | | | |

In addition to the principal routines, the examples also highlight how analysis results may look like in certain situations (especially in the case of modelling related problems), or how the results from the PESTO routines can be manipulated (the following list is of course non-exhaustive):

- *multi-modal posterior distributions* (transfection and Gaussian mixture)
- *structurally non-identifiable parameters* (Hyperring and transfection)
- *practically non-identifiable parameters* (enzymatic catalysis)
- *analysis of sampling results and Markov chain properties* (enzymatic catalysis, Gaussian mixture, and Hyperring)
- *cutting off burn-in phase of Markov chain after sampling* (transfection)
- *advanced usage of PESTO plotting routines* (Gaussian mixture)
- *integration of profile likelihoods* (enzymatic catalysis and transfection)
- *usage of non-default sampling algorithms* (transfection, Gaussian mixture, Hyperring)

### 3 Hierarchical Index

#### 3.1 Class Hierarchy

This inheritance list is sorted roughly, but not completely, alphabetically:

SetGet

**DRAMOptions** 13

**MALAOptions** 16

**PestoOptions** 18

**PestoPlottingOptions** 32

**PestoSamplingOptions** 45

**PHSOptions** 50

**PTOptions** 53

### 4 Class Index

#### 4.1 Class List

Here are the classes, structs, unions and interfaces with brief descriptions:

**DRAMOptions**
DRAMOptions provides an option container to pass options into the PestoSamplingOptions class for Delayed Rejection Adaption Metropolis (DRAM). The DRAM algorithm uses a delayed rejection scheme for better mixing 13

**MALAOptions**
MALAOptions provides an option container to pass options as subclass into the Pesto←SamplingOptions class for the Metropolis Adaptive Langevin Algorithm (MALA) 16

**PestoOptions**
PestoOptions provides an option container to pass options to various PESTO functions. Not all options are used by all functions, consult the respective function documentation for details 18

**PestoPlottingOptions**
PestoPlottingOptions is a class for checking and holding information on optimization parameters 32

**PestoSamplingOptions**
PestoSamplingOptions provides an option container to pass options to various PESTO functions. Not all options are used by all functions, consult the respective function documentation for details 45

**PHSOptions**
PHSOptions provides an option container to set options for Parallel Hierarchical Sampling (P←HS) in PestoSamplingOptions.PHS 50

**PTOptions**
PTOptions provides an option container to specify parallel tempering (PT) options in Pesto←SamplingOptions.PT 53

### 5 File Index

#### 5.1 File List

Here is a list of all documented files with brief descriptions:

**collectResults.m**
CollectResults() collects and plots the results stored in a common folder 57

**getMultiStarts.m**
GetMultiStarts() computes the maximum a posterior estimate of the parameters of a user-supplied posterior function. Therefore, a multi-start local optimization is used. The parameters from the best value of the posterior function arethen used as the global optimum. To ensure that the found maximum is a global one, a sufficiently high number of multistarts must be done. Those starts can be initialized with either randomly sampled parameter values, following either a uniform distribution or a latin hypercube, or they can be sampled by a user provided initial function (provided as option.init_fun) 58

**getParameterConfidenceIntervals.m**
GetParameterConfidenceIntervals() calculates the confidence intervals for the model parameters. This is done by four approaches: The values of CI.local_PL and CI.PL are determined by the point on which a threshold according to the confidence level alpha (calculated by a chi2-distribution) is reached. local_PL computes this point by a local approximation around the MAP estimate using the Hessian matrix, PL uses the profile likelihoods instead. The value of CI.←local_B is computed by using the cummulative distribution function of a local approximation of the profile based on the Hessian matrix at the MAP estimate. The value of CI.S is calculated using samples for the model parameters and the according percentiles based on the confidence levels alpha 61

getParameterProfiles.m
GetParameterProfiles.m calculates the profiles likelihoods for the model parameters, starting from the maximum a posteriori estimate. This calculation is done by fixing the i-th parameter and repeatedly reoptimizing the likelihood/posterior estimate (for all i). The initial guess for the next reoptimization point is computed by extrapolation from the previous points to ensure a quick optimization 63

getParameterSamples.m
GetParameterSamples.m performs MCMC sampling of the posterior distribution 65

getPropertyConfidenceIntervals.m
GetPropertyConfidenceIntervals.m calculates the confidence intervals for the model properties. This is done by three approaches: The values of CI.local_PL and CI.PL are determined by the point on which a threshold according to the confidence level alpha (calculated by a chi2-distribution) is reached. local_PL computes this point by a local approximation around the MAP estimate using the Hessian matrix, PL uses the profile likelihoods instead. The value of CI.local_B is computed by using the cummulative distribution function of a local approximation of the profile based on the Hessian matrix at the MAP estimate 67

getPropertyMultiStarts.m
GetPropertyMultiStarts.m evaluates the properties for the different mutli-start results 69

getPropertyProfiles.m
GetPropertyProfiles.m calculates the profiles of user-supplied property functions, starting from the maximum a posteriori estimate. This calculation is done by varying the value of each property function respectively, starting from the value of this function at the global optimum and by reoptimizing the likelihood/posterior estimate in each variational step of the property. The initial guess for the next reoptimization point is computed by extrapolation from the previous points to ensure a quick optimization 71

getPropertySamples.m
GetPropertySamples.m evaluates the properties for the sampled parameters 74

meigoDummy.m
Objective function wrapper for MEIGO / PSwarm / ... which need objective function $file*name$ and cannot use function handles directly 76

plotConfidenceIntervals.m
PlotConfidenceIntervals.m visualizes confidence itervals stored in either the parameters or properties struct .CI 77

plotMCMCdiagnosis.m
PlotMCMCdiagnosis.m visualizes the Markov chains generated by getSamples.m 79

plotMultiStartDiagnosis.m ??

plotMultiStarts.m
PlotMultiStarts plots the result of the multi-start optimization stored in parameters 80

plotParameterProfiles.m
PlotParameterProfiles.m visualizes profile likelihood. Note: This routine provides an interface for plotUncertainty.m 82

plotParameterSamples.m
PlotParameterSamples.m visualizes MCMC samples. Note: This routine provides an interface for plotUncertainty.m 84

plotParameterUncertainty.m
PlotParameterUncertainty.m visualizes profile likelihood and MCMC samples stored in parameters 85

**plotPropertyMultiStarts.m**
PlotPropertyMultiStarts plots the result of the multi-start optimization stored in properties 87

**plotPropertyProfiles.m**
PlotPropertyProfiles.m visualizes profile likelihood of model properties. Note: This routine provides an interface for plotPropertyUncertainty.m 88

**plotPropertySamples.m**
PlotPropertySamples.m visualizes samples of model properties. Note: This routine provides an interface for plotPropertyUncertainty.m 90

**plotPropertyUncertainty.m**
PlotPropertyUncertainty.m visualizes profile likelihood and MCMC samples stored in properties 92

**runPestoTests.m**
RunPestoTests Run a set of PESTO unit tests 93

**testGradient.m**
TestGradient.m calculates finite difference approximations to the gradient to check an analytical version 94

**@DRAMOptions/DRAMOptions.m** ??

**@MALAOptions/MALAOptions.m** ??

**@PestoOptions/PestoOptions.m** ??

**@PestoPlottingOptions/PestoPlottingOptions.m** ??

**@PestoSamplingOptions/PestoSamplingOptions.m** ??

**@PHSOptions/PHSOptions.m** ??

**@PTOptions/PTOptions.m** ??

### 6 Class Documentation

#### 6.1 DRAMOptions Class Reference

DRAMOptions provides an option container to pass options into the PestoSamplingOptions class for Delayed Rejection Adaption Metropolis (DRAM). The DRAM algorithm uses a delayed rejection scheme for better mixing.

Inheritance diagram for DRAMOptions:

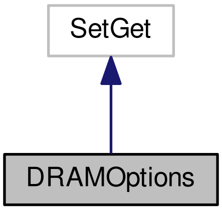

Collaboration diagram for DRAMOptions:

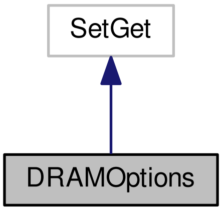

##### Public Member Functions

- DRAMOptions (matlabtypesubstitute varargin)

  *DRAMOptions Construct a new DRAMOptions object.*
- mlhsInnerSubst< matlabtypesubstitute, new > copy ()

  *Creates a copy of the passed DRAMOptions instance.*

##### Public Attributes

- matlabtypesubstitute regFactor = 1e-6

  *This factor is used for regularization in cases where the single-chain proposal covariance matrices are ill conditioned. Larger values equal stronger regularization.*
- matlabtypesubstitute nTry = 1

  *The number of tries in the delayed rejection scheme.*
- matlabtypesubstitute verbosityMode = "text"

  *Defines the level of verbosity `silent`, `visual`, `debug` or `text`*
- matlabtypesubstitute adaptionInterval = 1

  *Update the proposal density only every adaptionInterval time.*

##### 6.1.1 Detailed Description

DRAMOptions provides an option container to pass options into the PestoSamplingOptions class for Delayed Rejection Adaption Metropolis (DRAM). The DRAM algorithm uses a delayed rejection scheme for better mixing.

This file is based on AMICI amioptions.m (`http://icb-dcm.github.io/AMICI/`)

Definition at line 17 of file DRAMOptions.m.

##### 6.1.2 Constructor & Destructor Documentation

###### 6.1.2.1 DRAMOptions::DRAMOptions ( matlabtypesubstitute *varargin* )

DRAMOptions Construct a new DRAMOptions object.

OPTS = DRAMOptions() creates a set of options with each option set to itsdefault value.

OPTS = DRAMOptions(PARAM, VAL, ...) creates a set of options with the named parameters altered with the specified values.

OPTS = DRAMOptions(OLDOPTS, PARAM, VAL, ...) creates a copy of OLDOPTS with the named parameters altered with the specified value

Note to see the parameters, check the documentation page for DRAMOptions

**Parameters**

| |
|---|
| *varargin* |

Definition at line 72 of file DRAMOptions.m.

##### 6.1.3 Member Data Documentation

###### 6.1.3.1 DRAMOptions::adaptionInterval = 1

Update the proposal density only every adaptionInterval time.

**Default:** 1

**Note**

This property has custom functionality when its value is changed.

Definition at line 60 of file DRAMOptions.m.

Referenced by copy().

###### 6.1.3.2 DRAMOptions::nTry = 1

The number of tries in the delayed rejection scheme.

**Default:** 1

**Note**

This property has custom functionality when its value is changed.

Definition at line 42 of file DRAMOptions.m.

Referenced by copy().

###### 6.1.3.3 DRAMOptions::regFactor = 1e-6

This factor is used for regularization in cases where the single-chain proposal covariance matrices are ill conditioned. Larger values equal stronger regularization.

% Part for checking the correct setting of options

**Default:** 1e-6

**Note**

This property has custom functionality when its value is changed.

Definition at line 31 of file DRAMOptions.m.

Referenced by copy().

###### 6.1.3.4 DRAMOptions::verbosityMode = "text"

Defines the level of verbosity `silent`, `visual`, `debug` or `text`

**Default:** "text"

**Note**

This property has custom functionality when its value is changed.

Definition at line 51 of file DRAMOptions.m.

Referenced by copy().

The documentation for this class was generated from the following file:

- @DRAMOptions/DRAMOptions.m

#### 6.2 MALAOptions Class Reference

MALAOptions provides an option container to pass options as subclass into the PestoSamplingOptions class for the Metropolis Adaptive Langevin Algorithm (MALA).

Inheritance diagram for MALAOptions:

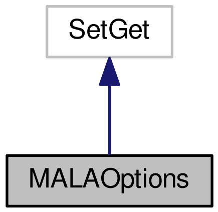

Collaboration diagram for MALAOptions:

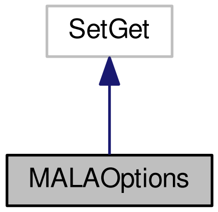

##### Public Member Functions

- MALAOptions (matlabtypesubstitute varargin)

  *MALAOptions Construct a new MALAOptions object.*
- mlhsInnerSubst< matlabtypesubstitute, new > copy ()

  *Creates a copy of the passed MALAOptions instance.*

##### Public Attributes

- matlabtypesubstitute regFactor = 1e-6

  *This factor is used for regularization in cases where the proposal covariance matrices are ill conditioned. Larger values equal stronger regularization.*

##### 6.2.1 Detailed Description

MALAOptions provides an option container to pass options as subclass into the PestoSamplingOptions class for the Metropolis Adaptive Langevin Algorithm (MALA).

Note: This algorithm uses gradients & Hessians either provided by the user or computed by finite differences.

This file is based on AMICI amioptions.m (`http://icb-dcm.github.io/AMICI/`)

Definition at line 17 of file MALAOptions.m.

##### 6.2.2 Constructor & Destructor Documentation

###### 6.2.2.1 MALAOptions::MALAOptions ( matlabtypesubstitute *varargin* )

MALAOptions Construct a new MALAOptions object.

OPTS = MALAOptions() creates a set of options with each option set to itsdefault value.

OPTS = MALAOptions(PARAM, VAL, ...) creates a set of options with the named parameters altered with the specified values.

OPTS = MALAOptions(OLDOPTS, PARAM, VAL, ...) creates a copy of OLDOPTS with the named parameters altered with the specified value

Note to see the parameters, check the documentation page for MALAOptions

**Parameters**

| |
|---|
| *varargin* |

Definition at line 47 of file MALAOptions.m.

##### 6.2.3 Member Data Documentation

###### 6.2.3.1 MALAOptions::regFactor = 1e-6

This factor is used for regularization in cases where the proposal covariance matrices are ill conditioned. Larger values equal stronger regularization.

% Part for checking the correct setting of options

**Default:** 1e-6

**Note**

This property has custom functionality when its value is changed.

Definition at line 33 of file MALAOptions.m.

Referenced by copy().

The documentation for this class was generated from the following file:

- @MALAOptions/MALAOptions.m

#### 6.3 PestoOptions Class Reference

PestoOptions provides an option container to pass options to various PESTO functions. Not all options are used by all functions, consult the respective function documentation for details.

Inheritance diagram for PestoOptions:

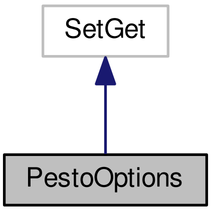

Collaboration diagram for PestoOptions:

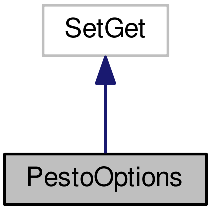

##### Public Member Functions

- PestoOptions (matlabtypesubstitute varargin)

  *PestoOptions Construct a new PestoOptions object.*
- mlhsInnerSubst< matlabtypesubstitute, new > copy ()

  *Creates a copy of the passed PestoOptions instance.*

##### Public Attributes

- matlabtypesubstitute comp_type = "sequential"

  *Perform calculations sequentially ('sequential', default), or in parallel ('parallel'). Parallel mode will speed-up the calculations on multi-core systems, but requires the* `MATLAB Parallel Computing Toolbox` *to be installed.*
- matlabtypesubstitute save = false

  *Determine whether results are saved or not.*
- matlabtypesubstitute foldername = strrep("datestr(now,31),' ','__'")

  *Name of the folder in which results are stored. If no folder is provided, a random foldername is generated.*
- matlabtypesubstitute obj_type = "log-posterior"

  *Type of objective function provided:* `log-posterior` *or* `negative log-posterior` *Tells the algorithm that log-posterior or log-likelihood are provided so it performs a maximization of the objective function or that the negative log-posterior or negative log-likelihood are provided so that a minimization of the objective function is performed.*
- matlabtypesubstitute objOutNumber = 3

  *Maximum number of outputs, the objective function can provide:*
- matlabtypesubstitute mode = "visual"

  *Output mode of algorithm:*
- matlabtypesubstitute fh = "[ ]"

  *Figure handle in which results are printed. If no handle is provided, a new figure is used. TODO: move to plot options.*
- matlabtypesubstitute plot_options = PestoPlottingOptions("")

  *Plotting options of class PestoPlottingOptions.m.*
- matlabtypesubstitute n_starts = 20

  *Number of local optimizations.*
- matlabtypesubstitute start_index = "[ ]"

  *vector of indices which starts should be performed. default is 1:n_starts*
- matlabtypesubstitute init_threshold = -inf

  *log-likelihood / log-posterior threshold for initialization of optimization.*
- matlabtypesubstitute localOptimizer = "fmincon"

  *Which optimizer to use? Current options: [*`fmincon`*,* `meigo-ess`*,* `meigo-vns`*,* `pswarm`*].*
- matlabtypesubstitute localOptimizerOptions

  *Options for the chosen local optimizer. Setting fmincon options as default local optimizer. See help(*`fmincon`*)*
- matlabtypesubstitute proposal = "latin hypercube"

  *Method used to propose starting points for fmincon. Can be.*
- matlabtypesubstitute trace = false

  *determine whether objective function, parameter values and computation time are stored over iterations*
- matlabtypesubstitute tempsave = false

  *determine whether intermediate results are stored every 10 iterations*
- matlabtypesubstitute resetobjective = false

  *clears the objective function before every multi-start.*
- matlabtypesubstitute calc_profiles = true

  *flag for profile calculation*
- matlabtypesubstitute P = {""}

  *parameter bounds (can be removed in a future release, since parameters.min/max can be used instead)*
- matlabtypesubstitute MAP_index = 1

*index MAP - parameter vector starting from which the profile is calculated. This option is helpful if local optima are present.*

- matlabtypesubstitute R_min = 0.03

  *Minimal ratio down to which the profile is calculated.*
- matlabtypesubstitute parameter_index = "[ ]"

  *Indices of the parameters for which the profile is calculated.*
- matlabtypesubstitute profile_optim_index = "[ ]"

  *Indices of the parameters for which the profile is calculated by reoptimization.*
- matlabtypesubstitute profile_integ_index = "[ ]"

  *Indices of the parameters for which the profile is calculated by integration.*
- matlabtypesubstitute profile_method = "default"

  *How should profiles be computed (if no more precise options are set like profile_optim_index or profile_integ_index)? Possibilities: {`optimization` (default), `integration`, `mixed`}.*
- matlabtypesubstitute profileReoptimizationOptions

  *Optimizer options for profile likelihood.*
- matlabtypesubstitute dR_max = 0.10

  *Maximal relative decrease of ratio allowed for two adjacent points in the profile (default = 0.10) if options.dJ = 0;.*
- matlabtypesubstitute dJ = 0.5

  *influences step size at small likelihood ratio values*
- matlabtypesubstitute options_getNextPoint

  *options for the generation fo the next profile point*
- matlabtypesubstitute solver

  *Options for profile integration.*
- matlabtypesubstitute boundary_tol = 1e-5

  *Tolance for the maximal distance of the list point the lower and upper bounds for the properties.*
- matlabtypesubstitute property_index = "[ ]"

  *Indices of the properties for which the profile is to be calculated (default = 1:properties.number, reoptimization only).*
- matlabtypesubstitute MCMC

  *Set MCMC options by calling an PestoSamplingOptions Class object.*

##### 6.3.1 Detailed Description

PestoOptions provides an option container to pass options to various PESTO functions. Not all options are used by all functions, consult the respective function documentation for details.

This file is based on AMICI amioptions.m (`http://icb-dcm.github.io/AMICI/`)

Definition at line 17 of file PestoOptions.m.

##### 6.3.2 Constructor & Destructor Documentation

###### 6.3.2.1 PestoOptions::PestoOptions ( matlabtypesubstitute *varargin* )

PestoOptions Construct a new PestoOptions object.

OPTS = PestoOptions() creates a set of options with each option set to its default value.

OPTS = PestoOptions(PARAM, VAL, ...) creates a set of options with the named parameters altered with the specified values.

OPTS = PestoOptions(OLDOPTS, PARAM, VAL, ...) creates a copy of OLDOPTS with the named parameters altered with the specified value

Note to see the parameters, check the documentation page for PestoOptions

**Parameters**

| |
|---|
| *varargin* |

Definition at line 544 of file PestoOptions.m.

##### 6.3.3 Member Data Documentation

###### 6.3.3.1 PestoOptions::boundary_tol = 1e-5

Tolance for the maximal distance of the list point the lower and upper bounds for the properties.

**Default:** 1e-5

Definition at line 509 of file PestoOptions.m.

###### 6.3.3.2 PestoOptions::calc_profiles = true

flag for profile calculation

- true: profiles are calculated
- false: profiles are not calculated

**Default:** true

**Note**

This property has custom functionality when its value is changed.

Definition at line 265 of file PestoOptions.m.

Referenced by copy().

###### 6.3.3.3 PestoOptions::comp_type = "sequential"

Perform calculations sequentially ('sequential', default), or in parallel ('parallel'). Parallel mode will speed-up the calculations on multi-core systems, but requires the `MATLAB Parallel Computing Toolbox` to be installed.

**Default:** "sequential"

**Note**

This property has custom functionality when its value is changed.

Definition at line 31 of file PestoOptions.m.

Referenced by copy().

###### 6.3.3.4 PestoOptions::dJ = 0.5

influences step size at small likelihood ratio values

**Default:** 0.5

Definition at line 402 of file PestoOptions.m.

###### 6.3.3.5 PestoOptions::dR_max = 0.10

Maximal relative decrease of ratio allowed for two adjacent points in the profile (default = 0.10) if options.dJ = 0;.

**Default:** 0.10

**Note**

This property has custom functionality when its value is changed.

Definition at line 392 of file PestoOptions.m.

Referenced by copy().

###### 6.3.3.6 PestoOptions::fh = "[ ]"

Figure handle in which results are printed. If no handle is provided, a new figure is used. TODO: move to plot options.

**Default:** "[ ]"

Definition at line 120 of file PestoOptions.m.

###### 6.3.3.7 PestoOptions::foldername = strrep("datestr(now,31),' ','__'")

Name of the folder in which results are stored. If no folder is provided, a random foldername is generated.

**Default:** strrep("datestr(now,31),' ','__'")

Definition at line 56 of file PestoOptions.m.

###### 6.3.3.8 PestoOptions::init_threshold = -inf

log-likelihood / log-posterior threshold for initialization of optimization.

**Default:** -inf

Definition at line 162 of file PestoOptions.m.

###### 6.3.3.9 PestoOptions::localOptimizer = "fmincon"

Which optimizer to use? Current options: [`fmincon`, `meigo-ess`, `meigo-vns`, `pswarm`].

For `meigo-ess` or `meigo-vns`, MEIGO (`http://gingproc.iim.csic.es/meigom.html`) has to be installed separately.

For `pswarm` PSwarm (`http://www.norg.uminho.pt/aivaz/pswarm/`) has to be installed separately

**Default:** "fmincon"

**Note**

This property has custom functionality when its value is changed.

Definition at line 172 of file PestoOptions.m.

Referenced by copy().

###### 6.3.3.10 PestoOptions::localOptimizerOptions

**Initial value:**

```
= optimset("    \
           'algorithm', 'interior-point',   \
           'display', 'off',    \
           'GradObj', 'on',    \
           'MaxIter', 2000,    \
           'PrecondBandWidth', inf")
```

Options for the chosen local optimizer. Setting fmincon options as default local optimizer. See *help(`fmincon`)*

MaxIter: fmincon default, necessary to be set for tracing

Options for `meigo-ess` are described in ess_kernel.m in the MEIGO folder.

**Default:** optimset(" \ 'algorithm', 'interior-point', \ 'display', 'off', \ 'GradObj', 'on', \ 'MaxIter', 2000, \ 'PrecondBand↩ Width', inf")

Definition at line 189 of file PestoOptions.m.

###### 6.3.3.11 PestoOptions::MAP_index = 1

index MAP - parameter vector starting from which the profile is calculated. This option is helpful if local optima are present.

**Default:** 1

**Note**

This property has custom functionality when its value is changed.

Definition at line 291 of file PestoOptions.m.

Referenced by copy().

###### 6.3.3.12 PestoOptions::mode = "visual"

Output mode of algorithm:

- `visual`: plots showing the progress are generated
- `text`: optimization results for multi-start are printed on screen
- `silent`: no output during the multi-start local optimization
- `debug`: print extra debug information (only available in certain functions

**Default:** "visual"

**Note**

This property has custom functionality when its value is changed.

Definition at line 105 of file PestoOptions.m.

Referenced by copy().

###### 6.3.3.13 PestoOptions::n_starts = 20

Number of local optimizations.

**Default:** 20

**Note**

This property has custom functionality when its value is changed.

Definition at line 142 of file PestoOptions.m.

Referenced by copy().

###### 6.3.3.14 PestoOptions::obj_type = "log-posterior"

Type of objective function provided: `log-posterior` or `negative log-posterior` Tells the algorithm that log-posterior or log-likelihood are provided so it performs a maximization of the objective function or that the negative log-posterior or negative log-likelihood are provided so that a minimization of the objective function is performed.

**Default:** "log-posterior"

**Note**

This property has custom functionality when its value is changed.

Definition at line 69 of file PestoOptions.m.

Referenced by copy().

###### 6.3.3.15 PestoOptions::objOutNumber = 3

Maximum number of outputs, the objective function can provide:

- 1 ... only objective value
- 2 ... objective value with gradient
- 3 ... objective value, gradient and Hessian

(Missing values will be approximated by finite differences.) Don't confuse this with the number of outputs from the objective function that Pesto really calls! objOutNumber just tells Pesto, with how many outputs it can call the objective function. Also, optimization is still possible to be done without gradient information and without the Pesto FD routine.

**Default:** 3

Definition at line 83 of file PestoOptions.m.

###### 6.3.3.16 PestoOptions::options_getNextPoint

**Initial value:**

```
= struct("'mode', 'multi-dimensional',    \
           'guess', 1e-2,    \
           'min', 1e-6,    \
           'max', 1,    \
           'update', 1.25")
```

options for the generation fo the next profile point

- .mode ... choice of proposal direction
- = `multi-dimensional` (default) ... all parameters are updated.
- The direct is the same as between the last two points.
- = `one-dimensional` ... only parameter for which profile is
- currently calculated is updated.
- .guess = 1e-2 ... guess for initial update stepsize
- .min = 1e-6 ... lower bound for update stepsize
- .min = 1e2 ... upper bound for update stepsize
- .update = 1.25 ... incremental change if stepsize is too large or
- too small, must be $>$ 1.

**Default:** struct("'mode', 'multi-dimensional', \ 'guess', 1e-2, \ 'min', 1e-6, \ 'max', 1, \ 'update', 1.25")

Definition at line 411 of file PestoOptions.m.

###### 6.3.3.17 PestoOptions::P = {""}

parameter bounds (can be removed in a future release, since parameters.min/max can be used instead)

- .P.min ... lower bound for profiling parameters, having same dimension as the parameter vector (default = parameters.min).
- .P.max ... lower bound for profiling parameters, having same dimension as the parameter vector (default = parameters.max).

**Default:** {""}

Definition at line 277 of file PestoOptions.m.

###### 6.3.3.18 PestoOptions::parameter_index = "[ ]"

Indices of the parameters for which the profile is calculated.

Default: profile_optim_index will be set to 1:parameters.number if left empty

**Default:** "[ ]"

**Note**

This property has custom functionality when its value is changed.

Definition at line 311 of file PestoOptions.m.

Referenced by copy().

###### 6.3.3.19 PestoOptions::plot_options = PestoPlottingOptions("")

Plotting options of class PestoPlottingOptions.m.

**Default:** PestoPlottingOptions("")

Definition at line 130 of file PestoOptions.m.

###### 6.3.3.20 PestoOptions::profile_integ_index = "[ ]"

Indices of the parameters for which the profile is calculated by integration.

**Default:** "[ ]"

Definition at line 333 of file PestoOptions.m.

###### 6.3.3.21 PestoOptions::profile_method = "default"

How should profiles be computed (if no more precise options are set like profile_optim_index or profile_integ_↩ index)? Possibilities: {`optimization` (default), `integration`, `mixed`}.

**Default:** "default"

Definition at line 343 of file PestoOptions.m.

###### 6.3.3.22 PestoOptions::profile_optim_index = "[ ]"

Indices of the parameters for which the profile is calculated by reoptimization.

**Default:** "[ ]"

Definition at line 323 of file PestoOptions.m.

###### 6.3.3.23 PestoOptions::profileReoptimizationOptions

**Initial value:**

```
= optimset("    \
            'algorithm', 'interior-point',    \
            'display', 'off',    \
            'MaxIter', 400,    \
            'GradObj', 'on',    \
            'GradConstr', 'on',    \
            'TolCon', 1e-6    \
            ")
```

Optimizer options for profile likelihood.

- .algorithm ... choice of algorithm
- = `interior-point` (default)
- = `trust-region-reflective`
- = `active-set`
- .display ... output of optimization process
- = `off` (default) ... no output
- = `iter` ... output for every optimization step
- .MaxIter ... maximum of optimization steps
- .GradObj ... are gradients provided?
- .GradConstr ... do we have constraints?
- .TolCon ... Tolerance for constraints

**Default:** optimset(" \ 'algorithm', 'interior-point', \ 'display', 'off', \ 'MaxIter', 400, \ 'GradObj', 'on', \ 'GradConstr', 'on', \ 'TolCon', 1e-6 \ ")

Definition at line 357 of file PestoOptions.m.

###### 6.3.3.24 PestoOptions::property_index = "[ ]"

Indices of the properties for which the profile is to be calculated (default = 1:properties.number, reoptimization only).

**Default:** "[ ]"

**Note**

This property has custom functionality when its value is changed.

Definition at line 520 of file PestoOptions.m.

Referenced by copy().

###### 6.3.3.25 PestoOptions::proposal = "latin hypercube"

Method used to propose starting points for fmincon. Can be.

- `latin hypercube`: latin hypercube sampling
- `uniform`: uniform random sampling
- `user-supplied`: user supplied function PestoOptions.init_fun

**Default:** "latin hypercube"

**Note**

This property has custom functionality when its value is changed.

Definition at line 213 of file PestoOptions.m.

Referenced by copy().

###### 6.3.3.26 PestoOptions::R_min = 0.03

Minimal ratio down to which the profile is calculated.

**Default:** 0.03

**Note**

This property has custom functionality when its value is changed.

Definition at line 302 of file PestoOptions.m.

Referenced by copy().

###### 6.3.3.27 PestoOptions::resetobjective = false

clears the objective function before every multi-start.

- false: persistent variables are preserved.
- true: remove all temporary/persistent variables.

WHEN TRUE THIS OPTION REMOVES ALL OBJECTIVE FUNCTION BREAK POINTS

**Default:** false

**Note**

This property has custom functionality when its value is changed.

Definition at line 249 of file PestoOptions.m.

Referenced by copy().

###### 6.3.3.28 PestoOptions::save = false

Determine whether results are saved or not.

- false: results are not saved
- true: results are stored to an extra folder

**Default:** false

**Note**

This property has custom functionality when its value is changed.

Definition at line 45 of file PestoOptions.m.

Referenced by copy().

###### 6.3.3.29 PestoOptions::solver

**Initial value:**

```
= struct("'type', 'ode113',     \
           'algorithm', 'Adams',     \
           'nonlinSolver', 'Newton',     \
           'linSolver', 'Dense',     \
           'gamma', 0,     \
           'eps', 1e-8,     \
           'minCond', 1e-10,     \
           'hessian', 'user-supplied',     \
           'gradient', true,     \
           'MaxStep', 0.1,     \
           'MinStep', 1e-4,     \
           'MaxNumSteps', 1e5,     \
           'RelTol', 1e-5,     \
           'AbsTol', 1e-8
           ")
```

Options for profile integration.

- .type ... choice of ODE integrator
- = `ode113` (default) ... Adams-Bashf.-Solver by Matlab
- = `ode15s` ... BDF-Solver by Matlab
- = `ode45` ... Runge-Kutta-Solver by Matlab
- = `CVODE` ... BDF and AB-Solver by sundials (to be implemented!)
- .algorithm ... choice of algorithm (CVODE only)
- = `Adams` (default) ... Adams-Bashford solver
- = `BDF` ... BDF solver
- .nonlinSolver ... choice of nonlinear solver (CVODE only)
- = `Newton` (default)
- = `Functional`
- .linSolver ... choice of linear solver (CVODE only)
- = `Dense` (default) ... solver for small problems
- = `Band` ... solver for bigger problems
- .gamma ... Retraction factor
- .eps ... regularization for poorly conditioned Hessian
- .minCond ... Minimum condition number, when regularization is to be used
- .hessian ... how is the Hessian Matrix provided?
- = `user-supplied` (default) ... Hessian is 3rd output of ObjFun
- = `bfgs` ... BFGS approximation to Hessian
- = `sr1` ... symmetric-rank 1 approximation to Hessian
- .gradient ... is a gradient provided?
- .MaxStep ... minimum step size of the solver
- .MaxStep ... maximum step size of the solver

- .MaxNumSteps ... maximum steps to be taken
- .RelTol ... maximum relative integration error
- .AbsTol ... maximum absolute integration error

**Default:** struct("'type', 'ode113', \ 'algorithm', 'Adams', \ 'nonlinSolver', 'Newton', \ 'linSolver', 'Dense', \ 'gamma', 0, \ 'eps', 1e-8, \ 'minCond', 1e-10, \ 'hessian', 'user-supplied', \ 'gradient', true, \ 'MaxStep', 0.1, \ 'MinStep', 1e-4, \ 'MaxNumSteps', 1e5, \ 'RelTol', 1e-5, \ 'AbsTol', 1e-8 ")

Definition at line 441 of file PestoOptions.m.

###### 6.3.3.30 PestoOptions::start_index = "[ ]"

vector of indices which starts should be performed. default is 1:n_starts

**Default:** "[ ]"

**Note**

This property has custom functionality when its value is changed.

Definition at line 152 of file PestoOptions.m.

Referenced by copy().

###### 6.3.3.31 PestoOptions::tempsave = false

determine whether intermediate results are stored every 10 iterations

- false: not saved
- true: results are stored to an extra folder

**Default:** false

**Note**

This property has custom functionality when its value is changed.

Definition at line 237 of file PestoOptions.m.

Referenced by copy().

###### 6.3.3.32 PestoOptions::trace = false

determine whether objective function, parameter values and computation time are stored over iterations

- false: not saved
- true: stored in fields par_trace, fval_trace and time_trace

**Default:** false

**Note**

This property has custom functionality when its value is changed.

Definition at line 225 of file PestoOptions.m.

Referenced by copy().

The documentation for this class was generated from the following file:

- @PestoOptions/PestoOptions.m

#### 6.4 PestoPlottingOptions Class Reference

PestoPlottingOptions is a class for checking and holding information on optimization parameters.

Inheritance diagram for PestoPlottingOptions:

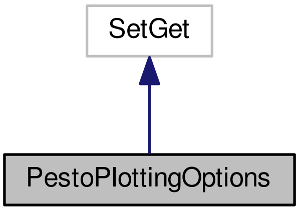

Collaboration diagram for PestoPlottingOptions:

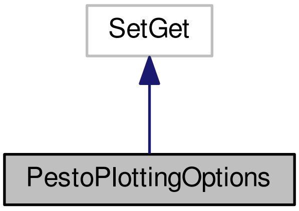

##### Public Member Functions

- PestoPlottingOptions (matlabtypesubstitute varargin)

  *PestoPlottingOptions Construct a new PestoPlottingOptions object.*
- mlhsInnerSubst< matlabtypesubstitute, new > copy ()

  *Creates a copy of the passed PestoPlottingOptions instance.*

##### Public Attributes

- matlabtypesubstitute title = true

  *Title of PESTO-generated plots.*
- matlabtypesubstitute add_points

  *Additional points to include in the plots, e.g. true parameter in the case of test examples.*
- matlabtypesubstitute mark_constraint = false

  *TODO: from plotmultistarts.*
- matlabtypesubstitute subplot_size_1D = "[ ]"

  *TODO from plotparameteruncertainty.*
- matlabtypesubstitute subplot_indexing_1D = "[ ]"

  *TODO.*
- matlabtypesubstitute labels

  *TODO.*
- matlabtypesubstitute hold_on = false

  *Indicates whether plots are redrawn or whether something is added to the plot.*
- matlabtypesubstitute interval = "dynamic"

  *Way of choosing x limits for plotting.*
- matlabtypesubstitute draw_bounds = true

  *Draw bounds.*
- matlabtypesubstitute bounds = {""}

  *Bounds used for visualization if options.interval =* `static`
- matlabtypesubstitute P

  *Options for profile plots.*
- matlabtypesubstitute S

  *Options for sample plots.*
- matlabtypesubstitute MS

  *Options for multi-start optimization plots.*
- matlabtypesubstitute A

  *Options for distribution approximation plots.*
- matlabtypesubstitute MCMC = "multistart"

  *Option if a user provided sampling initialization should be used for plotting an approximation of the distribution.*
- matlabtypesubstitute boundary

  *Options for boundary visualization.*
- matlabtypesubstitute CL

  *Options for confidence level plots.*
- matlabtypesubstitute group_CI_by = "parprop"

  *Options for the way to plot confidence intervals.*
- matlabtypesubstitute op2D = struct("'b1', 0.15, 'b2', 0.02, 'r', 0.95")

  *Settings for 2D plot to position subplot axes.*
- matlabtypesubstitute legend

  *Legend options.*
- matlabtypesubstitute fontsize = struct ("'tick', 12")

*Fontsize for labels.*

- matlabtypesubstitute fh_logPost_trace = "[ ]"

  *figure handle for log-posterior trace plot*
- matlabtypesubstitute fh_par_trace = "[ ]"

  *figure handle for parameter trace plots.*
- matlabtypesubstitute fh_par_dis_1D = "[ ]"

  *figure handle for the 1D parameter distribution plot.*
- matlabtypesubstitute fh_par_dis_2D = "[ ]"

  *figure handle for the 2D parameter distribution plot.*
- matlabtypesubstitute plot_type = {"'parameter','posterior'"}

  *plot type*
- matlabtypesubstitute n_max = 1e4

  *max*

##### 6.4.1 Detailed Description

PestoPlottingOptions is a class for checking and holding information on optimization parameters.

This file is based on AMICI amioptions.m (`http://icb-dcm.github.io/AMICI/`)

Definition at line 17 of file PestoPlottingOptions.m.

##### 6.4.2 Constructor & Destructor Documentation

###### 6.4.2.1 PestoPlottingOptions::PestoPlottingOptions ( matlabtypesubstitute *varargin* )

PestoPlottingOptions Construct a new PestoPlottingOptions object.

OPTS = PestoPlottingOptions() creates a set of options with each option set to its default value.

OPTS = PestoPlottingOptions(PARAM, VAL, ...) creates a set of options with the named parameters altered with the specified values.

OPTS = PestoPlottingOptions(OLDOPTS, PARAM, VAL, ...) creates a copy of OLDOPTS with the named parameters altered with the specified value

Note to see the parameters, check the documentation page for PestoPlottingOptions

**Parameters**

| |
|---|
| *varargin* |

Definition at line 489 of file PestoPlottingOptions.m.

##### 6.4.3 Member Data Documentation

###### 6.4.3.1 PestoPlottingOptions::A

**Initial value:**

```
= struct("'plot_type', 1,    \
           'col', [0,0,1],    \
           'lw', 2,    \
           'sigma_level', 2,    \
           'name', 'P_{app}'")
```

Options for distribution approximation plots.

###### Struct with

- .plot_type: plot type
  - = 0 (default if no MS are provided) ... no plot
  - = 1 (default if MS are provided) ... likelihood ratio
  - = 2 ... negative log-likelihood
- .col: color of approximation lines (default: [0,0,1])
- .lw: line width of approximation lines (default: 1.5)
- .sigma_level: sigma-level which is visualized (default = 2)
- .name: name of legend entry (default = `P_{app}`)

**Default:** struct("'plot_type', 1, \ 'col', [0,0,1], \ 'lw', 2, \ 'sigma_level', 2, \ 'name', 'P_{app}'")

Definition at line 277 of file PestoPlottingOptions.m.

###### 6.4.3.2 PestoPlottingOptions::add_points

**Initial value:**

```
= struct("'par', [],    \
           'logPost', [],    \
           'col', [0,0.8,0],    \
           'ls', '-',    \
           'lw', 1,    \
           'm', 'd',    \
           'ms', 8,    \
           'name', 'add. point'")
```

Additional points to include in the plots, e.g. true parameter in the case of test examples.

Struct with the following fields

- .par: n x m matrix of m additional points
- .col: color used for additional points (default = [0,0,0]). This can also be a m x 3 matrix of colors.
- .ls: line style (default = `-`)
- .lw: line width (default = 2)
- .m: marker style (default = `s`)
- .ms: line width (default = 8)
- .name: name of legend entry (default = `add.  point`)
- .property_MS: line width (default = 8).
- .logPost

**Default:** struct("'par', [], \ 'logPost', [], \ 'col', [0,0.8,0], \ 'ls', '-', \ 'lw', 1, \ 'm', 'd', \ 'ms', 8, \ 'name', 'add. point'")

Definition at line 42 of file PestoPlottingOptions.m.

###### 6.4.3.3 PestoPlottingOptions::boundary

**Initial value:**

```
= struct("'mark', true,    \
          'eps', 1e-4")
```

Options for boundary visualization.

Struct with

- .mark: marking of profile points which are on the boundary
  - = 0 ... no visualization
  - = 1 (default) ... indicates points which ar close to the boundaries in one or more dimensions.
- .eps: minimal distance from boundary for which points are consider to e close do the boundary (default = 1e-4). Note that a one-norm is used.

**Default:** struct("'mark', true, \ 'eps', 1e-4")

Definition at line 318 of file PestoPlottingOptions.m.

###### 6.4.3.4 PestoPlottingOptions::bounds = {""}

Bounds used for visualization if options.interval = `static`

struct with

- .min: lower bound
- .max: upper bound

**Default:** {""}

Definition at line 152 of file PestoPlottingOptions.m.

###### 6.4.3.5 PestoPlottingOptions::CL

**Initial value:**

```
= struct("'plot_type', 0,    \
          'alpha', 0.95,    \
          'type', 'point-wise',    \
          'col', [0,0,0],    \
          'lw', 2,     \
          'name', 'cut-off'")
```

Options for confidence level plots.

Struct with

- .plot_type: plot type
  - = 0 (default) ... no plot
  - = 1 ... likelihood ratio
  - = 2 ... negative log-likelihood
- .alpha: visualized confidence level (default = 0.95)
- .type: type of confidence interval
  - = `point-wise` (default) ... point-wise confidence interval
  - = `simultanous` ... point-wise confidence interval
  - = {`point-wise`,`simultanous`} ... both
- .col: color of profile lines (default: [0,0,0])
- .lw: line width of profile lines (default: 1.5)
- .name: name of legend entry (default = `cut-off`)

**Default:** struct("'plot_type', 0, \ 'alpha', 0.95, \ 'type', 'point-wise', \ 'col', [0,0,0], \ 'lw', 2, \ 'name', 'cut-off'")

Definition at line 339 of file PestoPlottingOptions.m.

###### 6.4.3.6 PestoPlottingOptions::draw_bounds = true

Draw bounds.

- true: yes
- false: no

**Default:** true

**Note**

This property has custom functionality when its value is changed.

Definition at line 141 of file PestoPlottingOptions.m.

Referenced by copy().

###### 6.4.3.7 PestoPlottingOptions::fh_logPost_trace = "[ ]"

figure handle for log-posterior trace plot

**Default:** "[ ]"

Definition at line 430 of file PestoPlottingOptions.m.

###### 6.4.3.8 PestoPlottingOptions::fh_par_dis_1D = "[ ]"

figure handle for the 1D parameter distribution plot.

**Default:** "[ ]"

Definition at line 448 of file PestoPlottingOptions.m.

###### 6.4.3.9 PestoPlottingOptions::fh_par_dis_2D = "[ ]"

figure handle for the 2D parameter distribution plot.

**Default:** "[ ]"

Definition at line 457 of file PestoPlottingOptions.m.

###### 6.4.3.10 PestoPlottingOptions::fh_par_trace = "[ ]"

figure handle for parameter trace plots.

**Default:** "[ ]"

Definition at line 439 of file PestoPlottingOptions.m.

###### 6.4.3.11 PestoPlottingOptions::fontsize = struct ("'tick', 12")

Fontsize for labels.

- .tick: fontsize for ticklabels (default = 12)

**Default:** struct ("'tick', 12")

Definition at line 420 of file PestoPlottingOptions.m.

###### 6.4.3.12 PestoPlottingOptions::group_CI_by = "parprop"

Options for the way to plot confidence intervals.

Either all confidence intervals of one method are plotted to one window `params`, or the confidence intervals for one parameter from all methods are plotted to one window `methods`, or everthing is grouped together `all`.

**Default:** "parprop"

**Note**

This property has custom functionality when its value is changed.

Definition at line 373 of file PestoPlottingOptions.m.

Referenced by copy().

###### 6.4.3.13 PestoPlottingOptions::hold_on = false

Indicates whether plots are redrawn or whether something is added to the plot.

- true: extension of plot
- false: new plot

**Default:** false

**Note**

This property has custom functionality when its value is changed.

Definition at line 116 of file PestoPlottingOptions.m.

Referenced by copy().

###### 6.4.3.14 PestoPlottingOptions::interval = "dynamic"

Way of choosing x limits for plotting.

- `dynamic`: x limits depending on analysis results
- `static`: x limits depending on parameters.min and .max or on user-defined bound options.bounds.min and .max. The later are used if provided.

**Default:** "dynamic"

**Note**

This property has custom functionality when its value is changed.

Definition at line 128 of file PestoPlottingOptions.m.

Referenced by copy().

###### 6.4.3.15 PestoPlottingOptions::labels

**Initial value:**

```
= struct("'y_always', true,    \
           'y_name', []")
```

TODO.

**Default:** struct("'y_always', true, \ 'y_name', [ ]")

Definition at line 105 of file PestoPlottingOptions.m.

###### 6.4.3.16 PestoPlottingOptions::legend

**Initial value:**

```
= struct("'color', 'none',    \
           'box', 'on',    \
           'orientation', 'vertical',    \
           'position', []")
```

Legend options.

- .color: background color (default = `none`).
- .box: legend outine (default = `on`).
- .orientation: orientation of list (default = `vertical`)

**Default:** struct("'color', 'none', \ 'box', 'on', \ 'orientation', 'vertical', \ 'position', []")

Definition at line 402 of file PestoPlottingOptions.m.

###### 6.4.3.17 PestoPlottingOptions::mark_constraint = false

TODO: from plotmultistarts.

**Default:** false

**Note**

This property has custom functionality when its value is changed.

Definition at line 78 of file PestoPlottingOptions.m.

Referenced by copy().

###### 6.4.3.18 PestoPlottingOptions::MCMC = "multistart"

Option if a user provided sampling initialization should be used for plotting an approximation of the distribution.

- `user-provided`
- `multistart` (default)

**Default:** "multistart"

**Note**

This property has custom functionality when its value is changed.

Definition at line 305 of file PestoPlottingOptions.m.

Referenced by copy().

###### 6.4.3.19 PestoPlottingOptions::MS

**Initial value:**

```
= struct("'plot_type', 1,    \
            'col', [1,0,0],    \
            'lw' , 2,    \
            'name_conv', 'MS - conv.',    \
            'name_nconv', 'MS - not conv.',    \
            'only_optimum', false")
```

Options for multi-start optimization plots.

**Struct with**

- .plot_type: plot type
  - = 0 (default if no MS are provided) ... no plot
  - = 1 (default if MS are provided) ... likelihood ratio and position of optima above threshold
  - = 2 ... negative log-likelihood and position of optima above threshold
- .col: color of local optima (default: [1,0,0])
- .lw: line width of local optima (default: 1.5)
- .name_conv: name of legend entry (default = `MS - conv.`)
- .name_nconv: name of legend entry (default = `MS - not conv.`)
- .only_optimum: only optimum is plotted

**Default:** struct("'plot_type', 1, \ 'col', [1,0,0], \ 'lw' , 2, \ 'name_conv', 'MS - conv.', \ 'name_nconv', 'MS - not conv.', \ 'only_optimum', false")

Definition at line 244 of file PestoPlottingOptions.m.

###### 6.4.3.20 PestoPlottingOptions::n_max = 1e4

max

**Default:** 1e4

**Note**

This property has custom functionality when its value is changed.

Definition at line 475 of file PestoPlottingOptions.m.

Referenced by copy().

###### 6.4.3.21 PestoPlottingOptions::op2D = struct("'b1', 0.15, 'b2', 0.02, 'r', 0.95")

Settings for 2D plot to position subplot axes.

**Struct with**

- .b1 ... offset from left and bottom border (default = 0.15)
- .b2 ... offset from left and bottom border (default = 0.02)
- .r ... relative width of subplots (default = 0.95)

**Default:** struct("'b1', 0.15, 'b2', 0.02, 'r', 0.95")

Definition at line 387 of file PestoPlottingOptions.m.

###### 6.4.3.22 PestoPlottingOptions::P

**Initial value:**

```
= struct("'plot_type', 1,    \
            'col', [1,0,0],    \
            'lw', 2,    \
            'name', 'P'")
```

Options for profile plots.

Struct with

- .plot_type: plot type
  - = 0 (default if no profiles are provided) ... no plot
  - = 1 (default if profiles are provided) ... likelihood ratio
  - = 2 ... negative log-likelihood
- .col: color of profile lines (default: [1,0,0])
- .lw: line width of profile lines (default: 1.5)

**Default:** struct("'plot_type', 1, \ 'col', [1,0,0], \ 'lw', 2, \ 'name', 'P'")

Definition at line 165 of file PestoPlottingOptions.m.

###### 6.4.3.23 PestoPlottingOptions::plot_type = {"'parameter','posterior'"}

plot type

**Default:** {"'parameter','posterior'"}

Definition at line 466 of file PestoPlottingOptions.m.

###### 6.4.3.24 PestoPlottingOptions::S

**Initial value:**

```
= struct("'plot_type', 0,    \
           'bins', 'conservative',    \
           'scaling', [],    \
           'hist_col',  [0.7,0.7,0.7],    \
           'sp_col', [0.7,0.7,0.7],    \
           'lin_col', [1,0,0],    \
           'lin_lw', 2,    \
           'sp_m', '.',    \
           'sp_ms', 5,    \
           'col', [1,0,0], 'lw', 2,    \
           'PT', struct('sp_m', '.',    \
               'sp_ms', 5,    \
               'lw', 1.5,    \
               'ind', [],    \
               'col', [],    \
               'plot_type', 0),    \
           'name', 'S'")
```

Options for sample plots.

- .plot_type: plot type
  - = 0 (default if no samples are provided) ... no plot
  - = 1 (default if samples are provided) ... histogram
  - = 2 ... kernel-density estimates
- .col ... color of profile lines (default: [0.7,0.7,0.7])
- .hist_col ... color of histogram (default = [0.7,0.7,0.7])
- .bins ... number of histogram bins (default: 30)
  - = `optimal` ... selection using Scott's rule
  - = `conservative` ... selection using Scott's rule / 2
  - = N (with N being an integer) ... N bins
- .sp_col: color of scatter plot (default = [0.7,0.7,0.7])
- .sp_m: marker for scatter plot (default = `.`)
- .sp_ms: marker size for scatter plot (default = 5)
- .name: name of legend entry (default = `S`)

**Default:** struct("'plot_type', 0, \ 'bins', 'conservative', \ 'scaling', [], \ 'hist_col', [0.7,0.7,0.7], \ 'sp_col', [0.7,0.7,0.7], \ 'lin_col', [1,0,0], \ 'lin_lw', 2, \ 'sp_m', '.', \ 'sp_ms', 5, \ 'col', [1,0,0], 'lw', 2, \ 'PT', struct('sp_m', '.', \ 'sp_ms', 5, \ 'lw', 1.5, \ 'ind', [], \ 'col', [], \ 'plot_type', 0), \ 'name', 'S'")

Definition at line 188 of file PestoPlottingOptions.m.

###### 6.4.3.25 PestoPlottingOptions::subplot_indexing_1D = "[ ]"

TODO.

**Default:** "[ ]"

Definition at line 96 of file PestoPlottingOptions.m.

###### 6.4.3.26 PestoPlottingOptions::subplot_size_1D = "[ ]"

TODO from plotparameteruncertainty.

**Default:** "[ ]"

Definition at line 87 of file PestoPlottingOptions.m.

###### 6.4.3.27 PestoPlottingOptions::title = true

Title of PESTO-generated plots.

- true: show
- false: don't show

**Default:** true

**Note**

This property has custom functionality when its value is changed.

Definition at line 30 of file PestoPlottingOptions.m.

Referenced by copy().

The documentation for this class was generated from the following file:

- @PestoPlottingOptions/PestoPlottingOptions.m

#### 6.5 PestoSamplingOptions Class Reference

PestoSamplingOptions provides an option container to pass options to various PESTO functions. Not all options are used by all functions, consult the respective function documentation for details.

Inheritance diagram for PestoSamplingOptions:

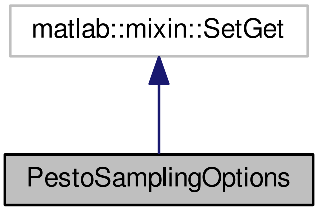

Collaboration diagram for PestoSamplingOptions:

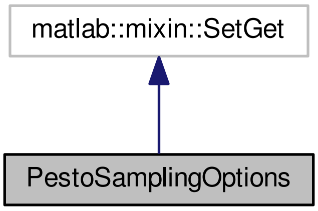

##### Public Member Functions

- PestoSamplingOptions (matlabtypesubstitute varargin)

  *PestoSamplingOptions Construct a new PestoSamplingOptions object.*
- mlhsInnerSubst< matlabtypesubstitute, new > copy ()

  *Creates a copy of the passed PestoSamplingOptions instance.*
- mlhsInnerSubst< matlabtypesubstitute, this > checkDependentDefaults (matlabtypesubstitute par)

  *checkDependentDefaults sets default values for sampling options which are problem-specific and will be adapted to the problem properties in par. Should be called providing the parameter struct for dependent defaults as theta0. Does both, checking already set dependent options and defaulting based on par if not set yet.*

**Public Attributes**

- matlabtypesubstitute obj_type = "log-posterior"

  *Type of objective function provided:* `log-posterior` *(default) or* `negative log-posterior`
- matlabtypesubstitute samplingAlgorithm = "PT"

  *Sampling algorithm, can be* `PT` *(parallel tempering),* `PHS`*, (parallel hierarchical sampling),* `MALA` *(Metropolis adjusted Langevin algorithm), or* `DRAM` *(delayed rejection adapted Metropolis algorithm, only if the DRAM toolbox is installed). Default value is* `PT`
- matlabtypesubstitute nIterations = 1e5

  *Number of iterations, integer (1e5 is not too high, e.g. 1e6 is a morereasonable value for reliable results, but computationally more intensive.*
- matlabtypesubstitute theta0 = "[ ]"

  *Initialization points for all chains. If the algorithm uses multiple chains (as* `PT` *and* `PHS`*), one can specify multiple theta0, e.g.: opt.theta0 = repmat([0.1,1.5,-2.5,-0.5,0.4],opt.nTemps,1)'; If there is just one chain, please specify as opt.theta0 = [1;2;3;4]; It is recommendet to set theta0 by taking into account the results from a preceeding optimization (see Pesto examples).*
- matlabtypesubstitute sigma0 = "[ ]"

  *Initial covariance matrices for all chains. Example for single-chain algorithms: opt.sigma0 = 1e5 ∗ diag(ones(1,5)); Example for multi-chain algorithms: opt.sigma0 = repmat(1e5∗diag(ones(1,5)),opt.nTemps,1); It is recommendet to set sigma0 by taking into account the results from a preceeding optimization.*
- matlabtypesubstitute mode = "visual"

  *Output mode for sampling algorithms (except DRAM, which has its own format), can be chosen as* `visual`*,* `text`*,* `silent`*, or* `debug`*. Default: visual.*
- matlabtypesubstitute objOutNumber = 1

  *Maximum number of outputs, the objective function can provide:*
- matlabtypesubstitute PT

  *Parallel Tempering Options, an instance of PTOptions.*
- matlabtypesubstitute PHS

  *Parallel Hierarchical Sampling options, an instance of PHSOptions.*
- matlabtypesubstitute MALA

  *Metropolis Adaptive Langevin Algorithm options, an instance of MALAOptions.*
- matlabtypesubstitute DRAM

  *Delayed Rejection Adaptive Metropolis options, an instance of DRAMOptions.*

##### 6.5.1 Detailed Description

PestoSamplingOptions provides an option container to pass options to various PESTO functions. Not all options are used by all functions, consult the respective function documentation for details.

This file is based on AMICI amioptions.m (`http://icb-dcm.github.io/AMICI/`)

Definition at line 17 of file PestoSamplingOptions.m.

##### 6.5.2 Constructor & Destructor Documentation

###### 6.5.2.1 PestoSamplingOptions::PestoSamplingOptions ( matlabtypesubstitute *varargin* )

PestoSamplingOptions Construct a new PestoSamplingOptions object.

OPTS = PestoSamplingOptions() creates a set of options with each option set to itsdefault value.

OPTS = PestoSamplingOptions(PARAM, VAL, ...) creates a set of options with the named parameters altered with the specified values.

OPTS = PestoSamplingOptions(OLDOPTS, PARAM, VAL, ...) creates a copy of OLDOPTS with the named parameters altered with the specified value

Note to see the parameters, check the documentation page for PestoSamplingOptions

**Parameters**

| |
|---|
| *varargin* |

Definition at line 162 of file PestoSamplingOptions.m.

##### 6.5.3 Member Function Documentation

###### 6.5.3.1 mlhsInnerSubst< matlabtypesubstitute, this > PestoSamplingOptions::checkDependentDefaults ( matlabtypesubstitute *par* )

checkDependentDefaults sets default values for sampling options which are problem-specific and will be adapted to the problem properties in par. Should be called providing the parameter struct for dependent defaults as theta0. Does both, checking already set dependent options and defaulting based on par if not set yet.

**Parameters**

| | |
|---|---|
| *par* | parameters as passed getParameterSamples() to use for choosing default options |

**Return values**

| | |
|---|---|
| *this* | the updated PestoSamplingOptions instance |

**Required fields of par:**

**Generated fields of this:**

Definition at line 397 of file PestoSamplingOptions.m.

##### 6.5.4 Member Data Documentation

###### 6.5.4.1 PestoSamplingOptions::mode = "visual"

Output mode for sampling algorithms (except DRAM, which has its own format), can be chosen as `visual`, `text`, `silent`, or `debug`. Default: visual.

**Default:** "visual"

**Note**

This property has custom functionality when its value is changed.

Definition at line 101 of file PestoSamplingOptions.m.

Referenced by copy().

###### 6.5.4.2 PestoSamplingOptions::nIterations = 1e5

Number of iterations, integer (1e5 is not too high, e.g. 1e6 is a morereasonable value for reliable results, but computationally more intensive.

**Default:** 1e5

**Note**

This property has custom functionality when its value is changed.

Definition at line 59 of file PestoSamplingOptions.m.

Referenced by copy().

###### 6.5.4.3 PestoSamplingOptions::obj_type = "log-posterior"

Type of objective function provided: `log-posterior` (default) or `negative log-posterior`

% Part for checking the correct setting of options

Tells the algorithm that log-posterior or log-likelihood are provided so it takes into account the corect sign for perfoming all algorithms correctly.

**Default:** "log-posterior"

**Note**

This property has custom functionality when its value is changed.

Definition at line 32 of file PestoSamplingOptions.m.

Referenced by copy().

###### 6.5.4.4 PestoSamplingOptions::objOutNumber = 1

Maximum number of outputs, the objective function can provide:

- 1 ... only objective value
- 2 ... objective value with gradient
- 3 ... objective value, gradient and Hessian (Default)

Missing values will be approximated by finite differences.

**Default:** 1

**Note**

This property has custom functionality when its value is changed.

Definition at line 112 of file PestoSamplingOptions.m.

Referenced by copy().

###### 6.5.4.5 PestoSamplingOptions::samplingAlgorithm = "PT"

Sampling algorithm, can be `PT` (parallel tempering), `PHS`, (parallel hierarchical sampling), `MALA` (Metropolis adjusted Langevin algorithm), or `DRAM` (delayed rejection adapted Metropolis algorithm, only if the DRAM toolbox is installed). Default value is `PT`

**Default:** "PT"

**Note**

This property has custom functionality when its value is changed.

Definition at line 47 of file PestoSamplingOptions.m.

Referenced by copy().

###### 6.5.4.6 PestoSamplingOptions::sigma0 = "[ ]"

Initial covariance matrices for all chains. Example for single-chain algorithms: opt.sigma0 = 1e5 ∗ diag(ones(1,5)); Example for multi-chain algorithms: opt.sigma0 = repmat(1e5∗diag(ones(1,5)),opt.nTemps,1); It is recommendet to set sigma0 by taking into account the results from a preceeding optimization.

**Default:** "[ ]"

Definition at line 86 of file PestoSamplingOptions.m.

###### 6.5.4.7 PestoSamplingOptions::theta0 = "[ ]"

Initialization points for all chains. If the algorithm uses multiple chains (as `PT` and `PHS`), one can specify multiple theta0, e.g.: opt.theta0 = repmat([0.1,1.5,-2.5,-0.5,0.4],opt.nTemps,1)'; If there is just one chain, please specify as opt.theta0 = [1;2;3;4]; It is recommendet to set theta0 by taking into account the results from a preceeding optimization (see Pesto examples).

**Default:** "[ ]"

Definition at line 70 of file PestoSamplingOptions.m.

The documentation for this class was generated from the following file:

- @PestoSamplingOptions/PestoSamplingOptions.m

#### 6.6 PHSOptions Class Reference

PHSOptions provides an option container to set options for Parallel Hierarchical Sampling (PHS) in Pesto←SamplingOptions.PHS .

Inheritance diagram for PHSOptions:

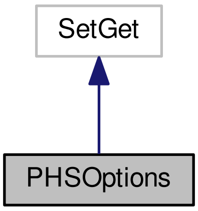

Collaboration diagram for PHSOptions:

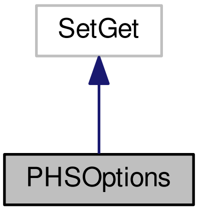

##### Public Member Functions

- PHSOptions (matlabtypesubstitute varargin)

  *PHSOptions Construct a new PHSOptions object.*
- mlhsInnerSubst< matlabtypesubstitute, new > copy ()

  *Creates a copy of the passed PHSOptions instance.*

##### Public Attributes

- matlabtypesubstitute nChains = 10

  *Initial number of temperatures.*
- matlabtypesubstitute alpha = 0.51

  *Parameter which controlls the adaption degeneration velocity of the single-chain proposals. Value between 0 and 1. No adaption (classical Metropolis-Hastings) for 0.*

- matlabtypesubstitute memoryLength = 1

  *The higher the value the more it lowers the impact of early adaption steps.*
- matlabtypesubstitute regFactor = 1e-6

  *Regularization factor for ill conditioned covariance matrices of the adapted proposal density. Regularization might happen if the eigenvalues of the covariance matrix strongly differ in order of magnitude. In this case, the algorithm adds a small diag-matrix to the covariance matrix with elements regFactor.*
- matlabtypesubstitute trainingTime = 1

  *The number of iterations before the chains start to swap. Might get used to ensure a proper local adaption of the single chains before exchanging them.*

##### 6.6.1 Detailed Description

PHSOptions provides an option container to set options for Parallel Hierarchical Sampling (PHS) in Pesto←SamplingOptions.PHS .

This file is based on AMICI amioptions.m (`http://icb-dcm.github.io/AMICI/`)

Definition at line 17 of file PHSOptions.m.

##### 6.6.2 Constructor & Destructor Documentation

###### 6.6.2.1 PHSOptions::PHSOptions ( matlabtypesubstitute *varargin* )

PHSOptions Construct a new PHSOptions object.

OPHSS = PHSOptions() creates a set of options with each option set to itsdefault value.

OPHSS = PHSOptions(PARAM, VAL, ...) creates a set of options with the named parameters altered with the specified values.

OPHSS = PHSOptions(OLDOPHSS, PARAM, VAL, ...) creates a copy of OLDOPHSS with the named parameters altered with the specified value

Note to see the parameters, check the documentation page for PHSOptions

adapted from SolverOptions

**Parameters**

| |
|---|
| *varargin* |

Definition at line 84 of file PHSOptions.m.

##### 6.6.3 Member Data Documentation

###### 6.6.3.1 PHSOptions::alpha = 0.51

Parameter which controlls the adaption degeneration velocity of the single-chain proposals. Value between 0 and 1. No adaption (classical Metropolis-Hastings) for 0.

**Default:** 0.51

**Note**

This property has custom functionality when its value is changed.

Definition at line 39 of file PHSOptions.m.

Referenced by copy().

###### 6.6.3.2 PHSOptions::memoryLength = 1

The higher the value the more it lowers the impact of early adaption steps.

**Default:** 1

**Note**

This property has custom functionality when its value is changed.

Definition at line 51 of file PHSOptions.m.

Referenced by copy().

###### 6.6.3.3 PHSOptions::nChains = 10

Initial number of temperatures.

**Default:** 10

**Note**

This property has custom functionality when its value is changed.

Definition at line 30 of file PHSOptions.m.

Referenced by copy().

###### 6.6.3.4 PHSOptions::regFactor = 1e-6

Regularization factor for ill conditioned covariance matrices of the adapted proposal density. Regularization might happen if the eigenvalues of the covariance matrix strongly differ in order of magnitude. In this case, the algorithm adds a small diag-matrix to the covariance matrix with elements regFactor.

% Part for checking the correct setting of options

**Default:** 1e-6

**Note**

This property has custom functionality when its value is changed.

Definition at line 60 of file PHSOptions.m.

Referenced by copy().

###### 6.6.3.5 PHSOptions::trainingTime = 1

The number of iterations before the chains start to swap. Might get used to ensure a proper local adaption of the single chains before exchanging them.

**Default:** 1

**Note**

This property has custom functionality when its value is changed.

Definition at line 72 of file PHSOptions.m.

Referenced by copy().

The documentation for this class was generated from the following file:

- @PHSOptions/PHSOptions.m

#### 6.7 PTOptions Class Reference

PTOptions provides an option container to specify parallel tempering (PT) options in PestoSamplingOptions.PT.

Inheritance diagram for PTOptions:

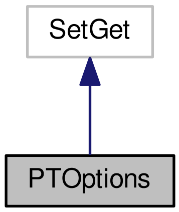

Collaboration diagram for PTOptions:

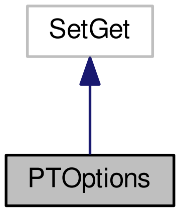

##### Public Member Functions

- PTOptions (matlabtypesubstitute varargin)

  *PTOptions Construct a new PTOptions object.*
- mlhsInnerSubst< matlabtypesubstitute, new > copy ()

  *Creates a copy of the passed PTOptions instance.*

##### Public Attributes

- matlabtypesubstitute nTemps = 10

  *Initial number of temperatures.*
- matlabtypesubstitute exponentT = 4

  *The initial temperatures are set by a power law to ^opt.exponentT.*
- matlabtypesubstitute alpha = 0.51

  *Parameter which controlls the adaption degeneration velocity of the single-chain proposals. Value between 0 and 1. No adaption (classical Metropolis-Hastings) for 0.*
- matlabtypesubstitute temperatureAlpha = 0.51

  *Parameter which controlls the adaption degeneration velocity of the temperature adaption. Value between 0 and 1. No effect for value = 0.*
- matlabtypesubstitute memoryLength = 1

  *The higher the value the more it lowers the impact of early adaption steps.*
- matlabtypesubstitute regFactor = 1e-6

  *Regularization factor for ill conditioned covariance matrices of the adapted proposal density. Regularization might happen if the eigenvalues of the covariance matrix strongly differ in order of magnitude. In this case, the algorithm adds a small diag-matrix to the covariance matrix with elements regFactor.*
- matlabtypesubstitute temperatureAdaptionScheme = "Vousden16"

  *Follows the temperature adaption scheme from `Vousden16` or `Lacki15`. Can be set to `none` for no temperature adaption.*

##### 6.7.1 Detailed Description

PTOptions provides an option container to specify parallel tempering (PT) options in PestoSamplingOptions.PT.

This file is based on AMICI amioptions.m (`http://icb-dcm.github.io/AMICI/`)

Definition at line 17 of file PTOptions.m.

##### 6.7.2 Constructor & Destructor Documentation

###### 6.7.2.1 PTOptions::PTOptions ( matlabtypesubstitute *varargin* )

PTOptions Construct a new PTOptions object.

OPTS = PTOptions() creates a set of options with each option set to itsdefault value.

OPTS = PTOptions(PARAM, VAL, ...) creates a set of options with the named parameters altered with the specified values.

OPTS = PTOptions(OLDOPTS, PARAM, VAL, ...) creates a copy of OLDOPTS with the named parameters altered with the specified value

Note to see the parameters, check the documentation page for PTOptions

**Parameters**

| |
|---|
| *varargin* |

Definition at line 106 of file PTOptions.m.

##### 6.7.3 Member Data Documentation

###### 6.7.3.1 PTOptions::alpha = 0.51

Parameter which controlls the adaption degeneration velocity of the single-chain proposals. Value between 0 and 1. No adaption (classical Metropolis-Hastings) for 0.

**Default:** 0.51

**Note**

This property has custom functionality when its value is changed.

Definition at line 49 of file PTOptions.m.

Referenced by copy().

###### 6.7.3.2 PTOptions::exponentT = 4

The initial temperatures are set by a power law to ^opt.exponentT.

**Default:** 4

**Note**

This property has custom functionality when its value is changed.

Definition at line 39 of file PTOptions.m.

Referenced by copy().

###### 6.7.3.3 PTOptions::memoryLength = 1

The higher the value the more it lowers the impact of early adaption steps.

**Default:** 1

**Note**

This property has custom functionality when its value is changed.

Definition at line 71 of file PTOptions.m.

Referenced by copy().

###### 6.7.3.4 PTOptions::nTemps = 10

Initial number of temperatures.

**Default:** 10

**Note**

This property has custom functionality when its value is changed.

Definition at line 30 of file PTOptions.m.

Referenced by copy().

###### 6.7.3.5 PTOptions::regFactor = 1e-6

Regularization factor for ill conditioned covariance matrices of the adapted proposal density. Regularization might happen if the eigenvalues of the covariance matrix strongly differ in order of magnitude. In this case, the algorithm adds a small diag-matrix to the covariance matrix with elements regFactor.

% Part for checking the correct setting of options

**Default:** 1e-6

**Note**

This property has custom functionality when its value is changed.

Definition at line 80 of file PTOptions.m.

Referenced by copy().

###### 6.7.3.6 PTOptions::temperatureAdaptionScheme = "Vousden16"

Follows the temperature adaption scheme from `Vousden16` or `Lacki15`. Can be set to `none` for no temperature adaption.

**Default:** "Vousden16"

**Note**

This property has custom functionality when its value is changed.

Definition at line 93 of file PTOptions.m.

Referenced by copy().

###### 6.7.3.7 PTOptions::temperatureAlpha = 0.51

Parameter which controlls the adaption degeneration velocity of the temperature adaption. Value between 0 and 1. No effect for value = 0.

**Default:** 0.51

**Note**

This property has custom functionality when its value is changed.

Definition at line 61 of file PTOptions.m.

Referenced by copy().

The documentation for this class was generated from the following file:

- @PTOptions/PTOptions.m

### 7 File Documentation

#### 7.1 collectResults.m File Reference

collectResults() collects and plots the results stored in a common folder

**Functions**

- mlhsInnerSubst< matlabtypesubstitute, obj > collectResults (matlabtypesubstitute foldername)

  *collectResults() collects and plots the results stored in a common folder*

##### 7.1.1 Detailed Description

collectResults() collects and plots the results stored in a common folder

##### 7.1.2 Function Documentation

###### 7.1.2.1 mlhsInnerSubst< matlabtypesubstitute, obj > collectResults ( matlabtypesubstitute *foldername* )

collectResults() collects and plots the results stored in a common folder

**USAGE**

[parameters] = collectResults(foldername)

**History**

2014/06/12 Jan Hasenauer % Initialization

**Parameters**

| | |
|---|---|
| *foldername* | Name of folder from which results are collected. |

**Return values**

| | |
|---|---|
| *parameters* | parameter struct. |

**Generated fields of obj:**

Definition at line 17 of file collectResults.m.

References plotMultiStarts(), plotParameterProfiles(), plotPropertyMultiStarts(), and plotPropertyProfiles().

Here is the call graph for this function:

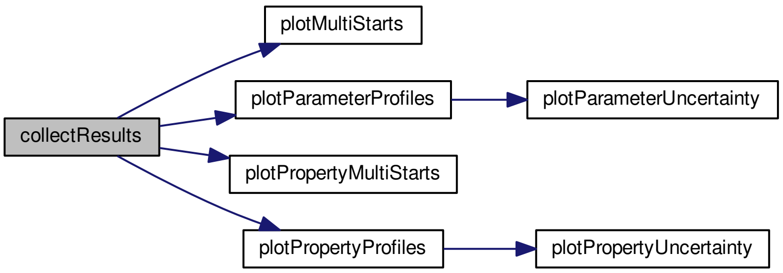

#### 7.2 getMultiStarts.m File Reference

getMultiStarts() computes the maximum a posterior estimate of the parameters of a user-supplied posterior function. Therefore, a multi-start local optimization is used. The parameters from the best value of the posterior function arethen used as the global optimum. To ensure that the found maximum is a global one, a sufficiently high number of multistarts must be done. Those starts can be initialized with either randomly sampled parameter values, following either a uniform distribution or a latin hypercube, or they can be sampled by a user provided initial function (provided as option.init_fun).

**Functions**

- mlhsSubst< mlhsInnerSubst< matlabtypesubstitute, parameters >,mlhsInnerSubst< matlabtypesubstitute, fh > > getMultiStarts (matlabtypesubstitute parameters, matlabtypesubstitute objective_function, matlabtypesubstitute varargin)

  *getMultiStarts() computes the maximum a posterior estimate of the parameters of a user-supplied posterior function. Therefore, a multi-start local optimization is used. The parameters from the best value of the posterior function arethen used as the global optimum. To ensure that the found maximum is a global one, a sufficiently high number of multistarts must be done. Those starts can be initialized with either randomly sampled parameter values, following either a uniform distribution or a latin hypercube, or they can be sampled by a user provided initial function (provided as option.init_fun).*
- mlhsInnerSubst< matlabtypesubstitute, stringTimePrediction > **mtoc_subst_getMultiStarts_m_tsbus_cotm_updateWaitBar** (matlabtypesubstitute timePredicted)
- noret::substitute **mtoc_subst_getMultiStarts_m_tsbus_cotm_saveResults** (matlabtypesubstitute parameters, matlabtypesubstitute options, matlabtypesubstitute i)

##### 7.2.1 Detailed Description

getMultiStarts() computes the maximum a posterior estimate of the parameters of a user-supplied posterior function. Therefore, a multi-start local optimization is used. The parameters from the best value of the posterior function arethen used as the global optimum. To ensure that the found maximum is a global one, a sufficiently high number of multistarts must be done. Those starts can be initialized with either randomly sampled parameter values, following either a uniform distribution or a latin hypercube, or they can be sampled by a user provided initial function (provided as option.init_fun).

##### 7.2.2 Function Documentation

###### 7.2.2.1 mlhsSubst< mlhsInnerSubst< matlabtypesubstitute, parameters >,mlhsInnerSubst< matlabtypesubstitute, fh > > getMultiStarts ( matlabtypesubstitute *parameters,* matlabtypesubstitute *objective_function,* matlabtypesubstitute *varargin* )

getMultiStarts() computes the maximum a posterior estimate of the parameters of a user-supplied posterior function. Therefore, a multi-start local optimization is used. The parameters from the best value of the posterior function arethen used as the global optimum. To ensure that the found maximum is a global one, a sufficiently high number of multistarts must be done. Those starts can be initialized with either randomly sampled parameter values, following either a uniform distribution or a latin hypercube, or they can be sampled by a user provided initial function (provided as option.init_fun).

Note: This function can exploit up to (n_start + 1) workers when running in `parallel` mode.

**USAGE**

- [...] = getMultiStarts(parameters,objective_function)
- [...] = getMultiStarts(parameters,objective_function,options)
- [parameters,fh] = getMultiStarts(...)

**getMultiStarts() uses the following PestoOptions members**

- PestoOptions::start_index
- PestoOptions::n_starts
- PestoOptions::mode
- PestoOptions::fh
- PestoOptions::fmincon
- PestoOptions::proposal
- PestoOptions::save
- PestoOptions::foldername
- PestoOptions::trace
- PestoOptions::comp_type
- PestoOptions::tempsave
- PestoOptions::resetobjective
- PestoOptions::obj_type
- PestoOptions::init_threshold
- PestoOptions::plot_options

**History**

- 2012/05/31 Jan Hasenauer

- 2012/07/11 Jan Hasenauer
- 2014/06/11 Jan Hasenauer
- 2015/07/28 Fabian Froehlich
- 2015/11/10 Fabian Froehlich
- 2016/06/07 Paul Stapor
- 2016/10/04 Daniel Weindl
- 2016/12/04 Paul Stapor

**Parameters**

| | |
|---|---|
| *parameters* | parameter struct |
| *objective_function* | objective function to be optimized. This function should accept one input, the parameter vector. |
| *varargin* | `1 getMultiStarts ( ..., options )` *Required Parameters for varargin:* • options A PestoOptions object holding various options for the algorithm. |

**Return values**

| | |
|---|---|
| *parameters* | updated parameter object |
| *fh* | figure handle |

**Required fields of parameters:**

- `number --` Number of parameters
- `min --` Lower bound for each parameter
- `max --` upper bound for each parameter name = {`name1`, ...}: names of the parameters
- `guess --` initial guess for the parameters (Optional, will be initialized empty if not provided)
- `init_fun --` function to draw starting points for local optimization, must have the structure init_fun(theta_0, theta_min, theta_max). (Only required if proposal == `user-supplied`)

**Generated fields of parameters:**

- `MS --` information about multi-start optimization
  - par0(:,i): starting point yielding ith MAP
  - par(:,i): ith MAP
  - logPost(i): log-posterior for ith MAP
  - logPost0(i): log-posterior for starting point yielding ith MAP
  - gradient(_,i): gradient of log-posterior at ith MAP
  - hessian(:,:,i): hessian of log-posterior at ith MAP
  - n_objfun(i): # objective evaluations used to calculate ith MAP
  - n_iter(i): # iterations used to calculate ith MAP
  - t_cpu(i): CPU time for calculation of ith MAP
  - exitflag(i): exitflag the optimizer returned for ith MAP
  - par_trace(:,:,i): parameter trace for ith MAP (if options.trace == true)
  - fval_trace(:,i): objective function value trace for ith MAP (if options.trace == true)

- time_trace(:,i): computation time trace for ith MAP (if options.trace == true)

Definition at line 17 of file getMultiStarts.m.

References plotMultiStarts().

Here is the call graph for this function:

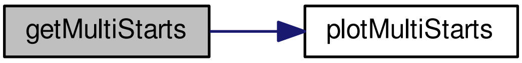

#### 7.3 getParameterConfidenceIntervals.m File Reference

getParameterConfidenceIntervals() calculates the confidence intervals for the model parameters. This is done by four approaches: The values of CI.local_PL and CI.PL are determined by the point on which a threshold according to the confidence level alpha (calculated by a chi2-distribution) is reached. local_PL computes this point by a local approximation around the MAP estimate using the Hessian matrix, PL uses the profile likelihoods instead. The value of CI.local_B is computed by using the cummulative distribution function of a local approximation of the profile based on the Hessian matrix at the MAP estimate. The value of CI.S is calculated using samples for the model parameters and the according percentiles based on the confidence levels alpha.

###### Functions

- mlhsInnerSubst< matlabtypesubstitute, parameters > getParameterConfidenceIntervals (matlabtypesubstitute parameters, matlabtypesubstitute alpha, matlabtypesubstitute varargin)

  *getParameterConfidenceIntervals() calculates the confidence intervals for the model parameters. This is done by four approaches: The values of CI.local_PL and CI.PL are determined by the point on which a threshold according to the confidence level alpha (calculated by a chi2-distribution) is reached. local_PL computes this point by a local approximation around the MAP estimate using the Hessian matrix, PL uses the profile likelihoods instead. The value of CI.local_B is computed by using the cummulative distribution function of a local approximation of the profile based on the Hessian matrix at the MAP estimate. The value of CI.S is calculated using samples for the model parameters and the according percentiles based on the confidence levels alpha.*

##### 7.3.1 Detailed Description

getParameterConfidenceIntervals() calculates the confidence intervals for the model parameters. This is done by four approaches: The values of CI.local_PL and CI.PL are determined by the point on which a threshold according to the confidence level alpha (calculated by a chi2-distribution) is reached. local_PL computes this point by a local approximation around the MAP estimate using the Hessian matrix, PL uses the profile likelihoods instead. The value of CI.local_B is computed by using the cummulative distribution function of a local approximation of the profile based on the Hessian matrix at the MAP estimate. The value of CI.S is calculated using samples for the model parameters and the according percentiles based on the confidence levels alpha.

###### 7.3.2 Function Documentation

###### 7.3.2.1 mlhsInnerSubst< matlabtypesubstitute, parameters > getParameterConfidenceIntervals ( matlabtypesubstitute *parameters*, matlabtypesubstitute *alpha*, matlabtypesubstitute *varargin* )

getParameterConfidenceIntervals() calculates the confidence intervals for the model parameters. This is done by four approaches: The values of CI.local_PL and CI.PL are determined by the point on which a threshold according to the confidence level alpha (calculated by a chi2-distribution) is reached. local_PL computes this point by a local approximation around the MAP estimate using the Hessian matrix, PL uses the profile likelihoods instead. The value of CI.local_B is computed by using the cummulative distribution function of a local approximation of the profile based on the Hessian matrix at the MAP estimate. The value of CI.S is calculated using samples for the model parameters and the according percentiles based on the confidence levels alpha.

**USAGE**

- parameters = getParameterConfidenceIntervals(parameters, alpha)

**History**

- 2013/11/29 Jan Hasenauer
- 2016/12/01 Paul Stapor

**Parameters**

| | |
|---|---|
| *parameters* | parameter struct |
| *alpha* | vector with desired confidence levels for the intervals |
| *varargin* | `1 getParameterConfidenceIntervals ( ..., options )` *Required Parameters for varargin:* • options A PestoOptions instance |

**Return values**

| | |
|---|---|
| *parameters* | updated parameter struct |

**Required fields of parameters:**

**Generated fields of parameters:**

- `CI --` Information about confidence levels
  - local_PL: Threshold based approach, uses a local approximation by the Hessian matrix at the MAP estimate (requires parameters.MS, e.g. from getMultiStarts)
  - PL: Threshold based approach, uses profile likelihoods (requires parameters.P, e.g. from get↩ParameterProfiles)
  - local_B: Mass based approach, uses a local approximation by the Hessian matrix at the MAP estimate (requires parameters.MS, e.g. from getMultiStarts)
  - S: Bayesian approach, uses percentiles based on samples (requires parameters.S, e.g. from get↩ParameterSamples)

Definition at line 17 of file getParameterConfidenceIntervals.m.

References plotConfidenceIntervals().

Here is the call graph for this function:

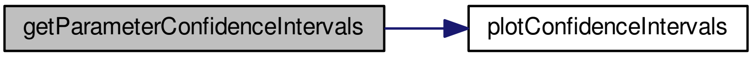

#### 7.4 getParameterProfiles.m File Reference

getParameterProfiles.m calculates the profiles likelihoods for the model parameters, starting from the maximum a posteriori estimate. This calculation is done by fixing the i-th parameter and repeatedly reoptimizing the likelihood/posterior estimate (for all i). The initial guess for the next reoptimization point is computed by extrapolation from the previous points to ensure a quick optimization.

##### Functions

- mlhsSubst$<$ mlhsInnerSubst$<$ matlabtypesubstitute, parameters $>$,mlhsInnerSubst$<$ matlabtypesubstitute, fh $>$ $>$ getParameterProfiles (matlabtypesubstitute parameters, matlabtypesubstitute objective_function, matlabtypesubstitute varargin)

  *getParameterProfiles.m calculates the profiles likelihoods for the model parameters, starting from the maximum a posteriori estimate. This calculation is done by fixing the i-th parameter and repeatedly reoptimizing the likelihood/posterior estimate (for all i). The initial guess for the next reoptimization point is computed by extrapolation from the previous points to ensure a quick optimization.*

##### 7.4.1 Detailed Description

getParameterProfiles.m calculates the profiles likelihoods for the model parameters, starting from the maximum a posteriori estimate. This calculation is done by fixing the i-th parameter and repeatedly reoptimizing the likelihood/posterior estimate (for all i). The initial guess for the next reoptimization point is computed by extrapolation from the previous points to ensure a quick optimization.

##### 7.4.2 Function Documentation

###### 7.4.2.1 mlhsSubst$<$ mlhsInnerSubst$<$ matlabtypesubstitute, parameters $>$,mlhsInnerSubst$<$ matlabtypesubstitute, fh $>$ $>$ getParameterProfiles ( matlabtypesubstitute *parameters*, matlabtypesubstitute *objective_function*, matlabtypesubstitute *varargin* )

getParameterProfiles.m calculates the profiles likelihoods for the model parameters, starting from the maximum a posteriori estimate. This calculation is done by fixing the i-th parameter and repeatedly reoptimizing the likelihood/posterior estimate (for all i). The initial guess for the next reoptimization point is computed by extrapolation from the previous points to ensure a quick optimization.

Note: This function can exploit up to (n_theta + 1) workers when running in `parallel` mode.

###### USAGE

[...] = getParameterProfiles(parameters, objective_function) [...] = getParameterProfiles(parameters, objective_function, options) [parameters, fh] = getParameterProfiles(...)

###### getParameterProfiles() uses the following PestoOptions members

- PestoOptions::calc_profiles
- PestoOptions::comp_type
- PestoOptions::dJ
- PestoOptions::dR_max
- PestoOptions::fh
- PestoOptions::MAP_index
- PestoOptions::mode
- PestoOptions::obj_type
- PestoOptions::options_getNextPoint .guess .min .max .update .mode
- PestoOptions::parameter_index
- PestoOptions::parameter_method_index
- PestoOptions::profile_method
- PestoOptions::profileReoptimizationOptions
- PestoOptions::plot_options
- PestoOptions::R_min
- PestoOptions::save

###### History

- 2012/05/16 Jan Hasenauer
- 2014/06/12 Jan Hasenauer
- 2016/10/04 Daniel Weindl
- 2016/10/12 Paul Stapor

###### Parameters

| | |
|---|---|
| *parameters* | parameter struct |
| *objective_function* | objective function to be optimized. This function should accept one input, the parameter vector. |
| *varargin* | `1 getParameterProfiles ( ..., options )`<br>*Required Parameters for varargin:*<br>• options A PestoOptions object holding various options for the algorithm. |

###### Return values

| | |
|---|---|
| *parameters* | updated parameter struct |
| *fh* | figure handle |

**Required fields of parameters:**

- `number --` Number of parameters
- `min --` Lower bound for each parameter
- `max --` upper bound for each parameter name = {`name1`, ...}: names of the parameters
- `MS --` results of global optimization, obtained using for instance the routine `getMultiStarts.m`. MS has to contain at least
  - par: sorted list n_theta x n_starts of parameter estimates. The first entry is assumed to be the best one.
  - logPost: sorted list n_starts x 1 of of log-posterior values corresponding to the parameters listed in .par.
  - hessian: Hessian matrix (or approximation) at the optimal point

**Generated fields of parameters:**

- `P(i) --` profile for i-th parameter
  - par: MAPs along profile
  - logPost: maximum log-posterior along profile
  - R: ratio

Definition at line 17 of file getParameterProfiles.m.

References plotParameterProfiles().

Here is the call graph for this function:

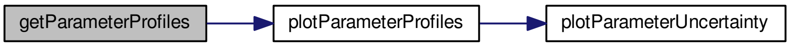

#### 7.5 getParameterSamples.m File Reference

getParameterSamples.m performs MCMC sampling of the posterior distribution.

##### Functions

- mlhsInnerSubst< matlabtypesubstitute, parameters > getParameterSamples (matlabtypesubstitute parameters, matlabtypesubstitute objFkt, matlabtypesubstitute opt)

  *getParameterSamples.m performs MCMC sampling of the posterior distribution.*

##### 7.5.1 Detailed Description

getParameterSamples.m performs MCMC sampling of the posterior distribution.

##### 7.5.2 Function Documentation

###### 7.5.2.1 mlhsInnerSubst< matlabtypesubstitute, parameters > getParameterSamples ( matlabtypesubstitute *parameters,* matlabtypesubstitute *objFkt,* matlabtypesubstitute *opt* )

getParameterSamples.m performs MCMC sampling of the posterior distribution.

Note, the DRAM library routine tooparameters.minox is used internally. This function is capable of sampling with MH, AM, DRAM, MALA, PT and PHS. The sampling plotting routines should no longer be contained in here but as standalone scripts capable of using the resulting par.S.

**History**

- 2012/07/11 Jan Hasenauer
- 2015/04/29 Jan Hasenauer
- 2016/10/17 Benjamin Ballnus
- 2016/10/19 Daniel Weindl
- 2016/11/04 Paul Stapor
- 2017/02/01 Benjamin Ballnus

**Parameters**

| | |
|---|---|
| *parameters* | parameter struct covering model options and results obtained by optimization, profiles and sampling. Optimization results can be used for initialization. |
| *objFkt* | Objective function which measures the difference of model output and data |
| *opt* | An options object holding various options for the sampling. Depending on the algorithm and particular flavor, different options must be set: For details, please visit PestoSamplingOptions.m |

**Return values**

| | |
|---|---|
| *parameters* | The provided parameters struct |

**Required fields of parameters:**

- `min` -- Lower parameter bounds
- `max` -- Upper parameter bounds
- `number` -- Number of parameters

**Required fields of opt:**

**Generated fields of parameters:**

- `S` -- The obtained sampling results

Definition at line 17 of file getParameterSamples.m.

References plotParameterSamples().

Here is the call graph for this function:

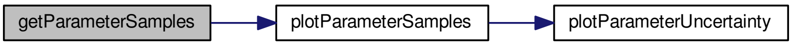

#### 7.6 getPropertyConfidenceIntervals.m File Reference

getPropertyConfidenceIntervals.m calculates the confidence intervals for the model properties. This is done by three approaches: The values of CI.local_PL and CI.PL are determined by the point on which a threshold according to the confidence level alpha (calculated by a chi2-distribution) is reached. local_PL computes this point by a local approximation around the MAP estimate using the Hessian matrix, PL uses the profile likelihoods instead. The value of CI.local_B is computed by using the cummulative distribution function of a local approximation of the profile based on the Hessian matrix at the MAP estimate.

##### Functions

- mlhsInnerSubst$<$ matlabtypesubstitute, properties $>$ getPropertyConfidenceIntervals (matlabtypesubstitute properties, matlabtypesubstitute alpha, matlabtypesubstitute varargin)

  *getPropertyConfidenceIntervals.m calculates the confidence intervals for the model properties. This is done by three approaches: The values of CI.local_PL and CI.PL are determined by the point on which a threshold according to the confidence level alpha (calculated by a chi2-distribution) is reached. local_PL computes this point by a local approximation around the MAP estimate using the Hessian matrix, PL uses the profile likelihoods instead. The value of CI.local_B is computed by using the cummulative distribution function of a local approximation of the profile based on the Hessian matrix at the MAP estimate.*

##### 7.6.1 Detailed Description

getPropertyConfidenceIntervals.m calculates the confidence intervals for the model properties. This is done by three approaches: The values of CI.local_PL and CI.PL are determined by the point on which a threshold according to the confidence level alpha (calculated by a chi2-distribution) is reached. local_PL computes this point by a local approximation around the MAP estimate using the Hessian matrix, PL uses the profile likelihoods instead. The value of CI.local_B is computed by using the cummulative distribution function of a local approximation of the profile based on the Hessian matrix at the MAP estimate.

##### 7.6.2 Function Documentation

###### 7.6.2.1 mlhsInnerSubst$<$ matlabtypesubstitute, properties $>$ getPropertyConfidenceIntervals ( matlabtypesubstitute *properties*, matlabtypesubstitute *alpha*, matlabtypesubstitute *varargin* )

getPropertyConfidenceIntervals.m calculates the confidence intervals for the model properties. This is done by three approaches: The values of CI.local_PL and CI.PL are determined by the point on which a threshold according to the confidence level alpha (calculated by a chi2-distribution) is reached. local_PL computes this point by a local approximation around the MAP estimate using the Hessian matrix, PL uses the profile likelihoods instead. The value of CI.local_B is computed by using the cummulative distribution function of a local approximation of the profile based on the Hessian matrix at the MAP estimate.

**USAGE**

- properties = getPropertyConfidenceIntervals(properties, alpha)

**History**

- 2013/11/29 Jan Hasenauer
- 2016/12/01 Paul Stapor

**Parameters**

| | |
|---|---|
| *properties* | property struct |
| *alpha* | vector with desired confidence levels for the intervals |
| *varargin* | `1 getPropertyConfidenceIntervals ( ..., options )`<br>*Required Parameters for varargin:*<br>• options A PestoOptions instance |

**Return values**

| | |
|---|---|
| *properties* | updated properties struct |

**Required fields of properties:**

**Generated fields of properties:**

- `CI --` Information about confidence levels
  - local_PL: Threshold based approach, uses a local approximation by the Hessian matrix at the MAP estimate (requires parameters.MS, e.g. from getMultiStarts)
  - PL: Threshold based approach, uses profile likelihoods (requires parameters.P, e.g. from get←ParameterProfiles)
  - local_B: Mass based approach, uses a local approximation by the Hessian matrix at the MAP estimate (requires parameters.MS, e.g. from getMultiStarts)

Definition at line 17 of file getPropertyConfidenceIntervals.m.

References plotConfidenceIntervals().

Here is the call graph for this function:

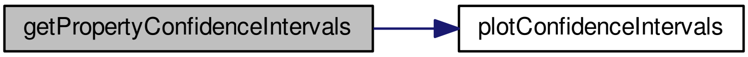

#### 7.7 getPropertyMultiStarts.m File Reference

getPropertyMultiStarts.m evaluates the properties for the different mutli-start results.

**Functions**

- mlhsSubst< mlhsInnerSubst< matlabtypesubstitute, properties >,mlhsInnerSubst< matlabtypesubstitute, fh > > getPropertyMultiStarts (matlabtypesubstitute properties, matlabtypesubstitute parameters, matlabtypesubstitute varargin)

  *getPropertyMultiStarts.m evaluates the properties for the different mutli-start results.*

##### 7.7.1 Detailed Description

getPropertyMultiStarts.m evaluates the properties for the different mutli-start results.

##### 7.7.2 Function Documentation

###### 7.7.2.1 mlhsSubst< mlhsInnerSubst< matlabtypesubstitute, properties >,mlhsInnerSubst< matlabtypesubstitute, fh > > getPropertyMultiStarts ( matlabtypesubstitute *properties*, matlabtypesubstitute *parameters*, matlabtypesubstitute *varargin* )

getPropertyMultiStarts.m evaluates the properties for the different mutli-start results.

**USAGE**

[...] = getPropertyMultiStarts(properties,parameters) [...] = getPropertyMultiStarts(properties,parameters,options) [parameters,fh] = getPropertyMultiStarts(...)

**getPropertyMultiStarts() uses the following PestoOptions members**

- PestoOptions::mode
- PestoOptions::fh
- PestoOptions::save
- PestoOptions::foldername
- PestoOptions::comp_type

**History**

- 2015/03/03 Jan Hasenauer
- 2016/04/10 Daniel Weindl

**Parameters**

| | |
|---|---|
| *properties* | property struct containing at least: |
| *parameters* | parameter struct containing at least: |
| *varargin* | `1 getPropertyMultiStarts ( ..., MS, number, min, max, options )`<br>*Required Parameters for varargin:*<br>• MS information about multi-start optimization<br>• number Number of properties<br>• min lower bound for property values<br>• max upper bound for property values name = {`name1`,...}: names of the properties function = {`function1`,...}: functions to evaluate property values. These functions provide the values of the respective properties and the corresponding 1st and 2nd order derivatives.<br>• options A PestoOptions object holding the options for the algorithm. |

**Return values**

| | |
|---|---|
| *properties* | updated parameter object containing: |
| *fh* | figure handle |
| *MS* | properties for multi-start optimization results<br>• par(:,i): ith MAP<br>• logPost(i): log-posterior for ith MAP<br>• exitflag(i): exit flag of ith MAP<br>• prop(j,i): values of jth property for ith MAP<br>• prop_Sigma(:,:,i): covariance of properties for ith MAP |

**Required fields of parameters:**

**Required fields of properties:**

**Generated fields of properties:**

Definition at line 17 of file getPropertyMultiStarts.m.

References plotPropertyMultiStarts().

Here is the call graph for this function:

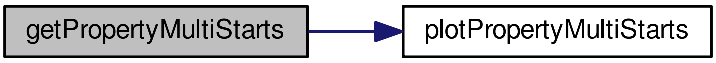

#### 7.8 getPropertyProfiles.m File Reference

getPropertyProfiles.m calculates the profiles of user-supplied property functions, starting from the maximum a posteriori estimate. This calculation is done by varying the value of each property function respectively, starting from the value of this function at the global optimum and by reoptimizing the likelihood/posterior estimate in each variational step of the property. The initial guess for the next reoptimization point is computed by extrapolation from the previous points to ensure a quick optimization.

###### Functions

- mlhsSubst< mlhsInnerSubst< matlabtypesubstitute, properties >,mlhsInnerSubst< matlabtypesubstitute, fh > > getPropertyProfiles (matlabtypesubstitute properties, matlabtypesubstitute parameters, matlabtypesubstitute objective_function, matlabtypesubstitute varargin)

  *getPropertyProfiles.m calculates the profiles of user-supplied property functions, starting from the maximum a posteriori estimate. This calculation is done by varying the value of each property function respectively, starting from the value of this function at the global optimum and by reoptimizing the likelihood/posterior estimate in each variational step of the property. The initial guess for the next reoptimization point is computed by extrapolation from the previous points to ensure a quick optimization.*
- mlhsInnerSubst< matlabtypesubstitute, varargout > **mtoc_subst_getPropertyProfiles_m_tsbus_cotm↩_obj** (matlabtypesubstitute theta, matlabtypesubstitute fun, matlabtypesubstitute type)
- mlhsInnerSubst< matlabtypesubstitute, varargout > **mtoc_subst_getPropertyProfiles_m_tsbus_cotm↩_obj_con** (matlabtypesubstitute theta, matlabtypesubstitute fun, matlabtypesubstitute fun_min, matlabtypesubstitute type)
- mlhsInnerSubst< matlabtypesubstitute, varargout > **mtoc_subst_getPropertyProfiles_m_tsbus_cotm_↩prop_fun** (matlabtypesubstitute theta, matlabtypesubstitute fun, matlabtypesubstitute prop_min, matlabtypesubstitute prop_max, matlabtypesubstitute s)
- mlhsInnerSubst< matlabtypesubstitute, varargout > **mtoc_subst_getPropertyProfiles_m_tsbus_cotm↩_prop_con_fun** (matlabtypesubstitute theta, matlabtypesubstitute fun, matlabtypesubstitute prop_min, matlabtypesubstitute prop_max, matlabtypesubstitute s)

##### 7.8.1 Detailed Description

getPropertyProfiles.m calculates the profiles of user-supplied property functions, starting from the maximum a posteriori estimate. This calculation is done by varying the value of each property function respectively, starting from the value of this function at the global optimum and by reoptimizing the likelihood/posterior estimate in each variational step of the property. The initial guess for the next reoptimization point is computed by extrapolation from the previous points to ensure a quick optimization.

##### 7.8.2 Function Documentation

###### 7.8.2.1 mlhsSubst< mlhsInnerSubst< matlabtypesubstitute, properties >,mlhsInnerSubst< matlabtypesubstitute, fh > > getPropertyProfiles ( matlabtypesubstitute *properties,* matlabtypesubstitute *parameters,* matlabtypesubstitute *objective_function,* matlabtypesubstitute *varargin* )

getPropertyProfiles.m calculates the profiles of user-supplied property functions, starting from the maximum a posteriori estimate. This calculation is done by varying the value of each property function respectively, starting from the value of this function at the global optimum and by reoptimizing the likelihood/posterior estimate in each variational step of the property. The initial guess for the next reoptimization point is computed by extrapolation from the previous points to ensure a quick optimization.

Note: This function can exploit up to (n_theta + 1) workers when running in `parallel` mode.

###### USAGE

[...] = getPropertyProfiles(properties, parameters, objective_function) [...] = getPropertyProfiles(properties, parameters, objective_function, options) [parameters, fh] = getPropertyProfiles(...)

%

###### getPropertyProfiles() uses the following PestoOptions members

- PestoOptions::boundary

- PestoOptions::calc_profiles
- PestoOptions::comp_type
- PestoOptions::dJ
- PestoOptions::dR_max
- PestoOptions::fh
- PestoOptions::fmincon
- PestoOptions::foldername
- PestoOptions::MAP_index
- PestoOptions::mode
- PestoOptions::obj_type
- PestoOptions::options_getNextPoint .guess .min .max .update .mode
- PestoOptions::plot_options
- PestoOptions::property_index
- PestoOptions::R_min
- PestoOptions::save

**History**

- 2012/03/02 Jan Hasenauer
- 2016/04/10 Daniel Weindl
- 2016/10/12 Paul Stapor

**Parameters**

| | |
|---|---|
| *properties* | property struct |
| *parameters* | parameter struct |
| *objective_function* | objective function to be optimized. This function should accept one input, the parameter vector. |
| *varargin* | `getPropertyProfiles ( ..., options )`<br>*Required Parameters for varargin:*<br>• options A PestoOptions object holding various options for the algorithm. |

**Return values**

| | |
|---|---|
| *properties* | updated property struct |
| *fh* | figure handle |

**Required fields of properties:**

- `number --` Number of properties
- `min --` Lower bound for each properties
- `max --` upper bound for each properties name = {`name1`, ...}: names of the properties function = {`function1`, ...}: functions to evaluate property values. These functions provide the values of the respective properties and the corresponding 1st and 2nd order derivatives.

**Required fields of parameters:**

- `number --` Number of parameters
- `min --` Lower bound for each parameter
- `max --` upper bound for each parameter name = {`name1`, ...}: names of the parameters
- `MS --` results of global optimization, obtained using for instance the routine `getMultiStarts.m`. MS has to contain at least
  - par: sorted list n_theta x n_starts of parameter estimates. The first entry is assumed to be the best one.
  - logPost: sorted list n_starts x 1 of of log-posterior values corresponding to the parameters listed in .par.
  - hessian: Hessian matrix (or approximation) at the optimal point

**Generated fields of properties:**

- `P(i) --` profile for i-th parameter
  - prop: MAPs along profile
  - par: MAPs along profile
  - logPost: maximum log-posterior along profile
  - R: ratio

Definition at line 17 of file getPropertyProfiles.m.

References plotPropertyProfiles().

Here is the call graph for this function:

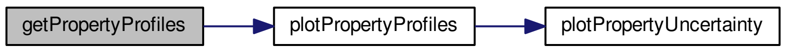

#### 7.9 getPropertySamples.m File Reference

getPropertySamples.m evaluates the properties for the sampled parameters.

##### Functions

- mlhsSubst< mlhsInnerSubst< matlabtypesubstitute, properties >,mlhsInnerSubst< matlabtypesubstitute, fh > > getPropertySamples (matlabtypesubstitute properties, matlabtypesubstitute parameters, matlabtypesubstitute varargin)

  *getPropertySamples.m evaluates the properties for the sampled parameters.*

##### 7.9.1 Detailed Description

getPropertySamples.m evaluates the properties for the sampled parameters.

##### 7.9.2 Function Documentation

###### 7.9.2.1 mlhsSubst< mlhsInnerSubst< matlabtypesubstitute, properties >,mlhsInnerSubst< matlabtypesubstitute, fh > > getPropertySamples ( matlabtypesubstitute *properties*, matlabtypesubstitute *parameters*, matlabtypesubstitute *varargin* )

getPropertySamples.m evaluates the properties for the sampled parameters.

**USAGE**

[...] = getPropertySamples(properties,parameters) [...] = getPropertySamples(properties,parameters,options) [parameters,fh] = getPropertySamples(...)

**getPropertySamples() uses the following PestoOptions members**

- PestoOptions::property_index
- PestoOptions::mode
- PestoOptions::fh
- PestoOptions::save
- PestoOptions::foldername
- PestoOptions::comp_type
- PestoOptions::plot_options
- PestoOptions::MCMC.thinning

**History**

- 2015/04/01 Jan Hasenauer
- 2016/10/04 Daniel Weindl

**Parameters**

| | |
|---|---|
| *properties* | property struct |
| *parameters* | parameter struct |
| *varargin* | `1 getPropertySamples ( ..., options )` *Required Parameters for varargin:* • options A PestoOptions object holding various options for the algorithm. |

**Return values**

| | |
|---|---|
| *properties* | updated parameter object |
| *fh* | figure handle |

**Required fields of properties:**

- `number --` number of parameter
- `min --` lower bound for property values

- `max --` upper bound for property values
- `name --` = {`name1`,...} ... names of the parameters
- `function --` = {`function1`,...} ... functions to evaluate property values. These functions provide the values of the respective properties and the corresponding 1st and 2nd order derivatives.

**Required fields of parameters:**

- `S --` parameter and posterior sample. logPost ... log-posterior function along chain par ... parameters along chain *Note* This struct is obtained using getSamples.m.

**Generated fields of properties:**

- `S --` properties for sampling results
  - par(∗,i): ith samples parameter vector
  - logPost(i): log-posterior for ith samples parameter vector
  - prop(j,i): values of jth property for ith samples parameter vector
  - prop_Sigma(∗,∗,i): covariance of properties for ith samples parameter vector

Definition at line 17 of file getPropertySamples.m.

References plotPropertySamples().

Here is the call graph for this function:

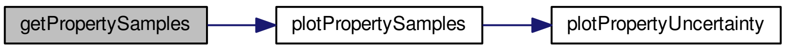

#### 7.10 meigoDummy.m File Reference

Objective function wrapper for MEIGO / PSwarm / ... which need objective function *file∗name* and cannot use function handles directly.

##### Functions

- mlhsInnerSubst< matlabtypesubstitute, f > meigoDummy (matlabtypesubstitute theta, matlabtypesubstitute fun, matlabtypesubstitute varargin)

  *Objective function wrapper for MEIGO / PSwarm / ... which need objective function file∗name and cannot use function handles directly.*

##### 7.10.1 Detailed Description

Objective function wrapper for MEIGO / PSwarm / ... which need objective function *file∗name* and cannot use function handles directly.

##### 7.10.2 Function Documentation

###### 7.10.2.1 mlhsInnerSubst< matlabtypesubstitute, f > meigoDummy ( matlabtypesubstitute *theta,* matlabtypesubstitute *fun,* matlabtypesubstitute *varargin* )

Objective function wrapper for MEIGO / PSwarm / ... which need objective function *file∗name* and cannot use function handles directly.

**Parameters**

| | |
|---|---|
| *theta* | parameter vector |
| *fun* | objective function handle |
| *varargin* | |

**Return values**

| | |
|---|---|
| *f* | Objective function value |

Definition at line 17 of file meigoDummy.m.

#### 7.11 plotConfidenceIntervals.m File Reference

plotConfidenceIntervals.m visualizes confidence itervals stored in either the parameters or properties struct .CI

**Functions**

- mlhsInnerSubst< matlabtypesubstitute, fh > plotConfidenceIntervals (matlabtypesubstitute pStruct, matlabtypesubstitute alpha, matlabtypesubstitute varargin)

  *plotConfidenceIntervals.m visualizes confidence itervals stored in either the parameters or properties struct .CI*
- mlhsInnerSubst< matlabtypesubstitute, methodsOut > **mtoc_subst_plotConfidenceIntervals_m_tsbus_cotm_checkMeth** (matlabtypesubstitute methodsIn, matlabtypesubstitute pStruct, matlabtypesubstitute boolWarning)

##### 7.11.1 Detailed Description

plotConfidenceIntervals.m visualizes confidence itervals stored in either the parameters or properties struct .CI

##### 7.11.2 Function Documentation

###### 7.11.2.1 mlhsInnerSubst< matlabtypesubstitute, fh > plotConfidenceIntervals ( matlabtypesubstitute *pStruct,* matlabtypesubstitute *alpha,* matlabtypesubstitute *varargin* )

plotConfidenceIntervals.m visualizes confidence itervals stored in either the parameters or properties struct .CI

**USAGE**

fh = plotParameterUncertainty(pStruct) fh = plotParameterUncertainty(pStruct, methods) fh = plotParameterUncertainty(pStruct, methods, options)

**plotMultiStarts() uses the following PestoPlottingOptions members**

- PestoPlottingOptions::P
- PestoPlottingOptions::S
- PestoPlottingOptions::MS
- PestoPlottingOptions::boundary

- PestoPlottingOptions::subplot_size_1D
- PestoPlottingOptions::subplot_indexing_1D
- PestoPlottingOptions::CL
- PestoPlottingOptions::hold_on
- PestoPlottingOptions::interval
- PestoPlottingOptions::bounds
- PestoPlottingOptions::A
- PestoPlottingOptions::add_points
- PestoPlottingOptions::labels
- PestoPlottingOptions::legend
- PestoPlottingOptions::op2D
- PestoPlottingOptions::fontsize

**History**

- 2016/11/14 Paul Stapor

**Parameters**

| | |
|---|---|
| *pStruct* | either the parameter or the property struct containing information about parameters and results of optimization (.MS) and uncertainty analysis (.P and .S). This structures is the output of plotMultiStarts.m, getProfiles.m or plotSamples.m. |
| *alpha* | significance levels |
| *varargin* | `1 plotConfidenceIntervals ( ..., method, options )`<br>*Required Parameters for varargin:*<br>• method integer array, from which method confidence intervals should be plotted:<br>• options options of plotting as instance of PestoPlottingOptions |

**Return values**

| | |
|---|---|
| *fh* | figure handle |

**Required fields of pStruct:**

Definition at line 17 of file plotConfidenceIntervals.m.

Referenced by getParameterConfidenceIntervals(), and getPropertyConfidenceIntervals().

Here is the caller graph for this function:

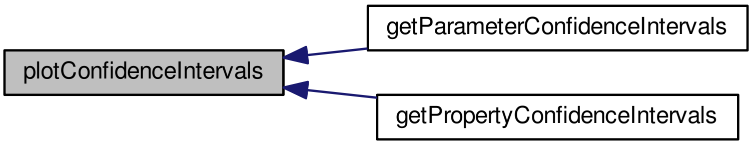

#### 7.12 plotMCMCdiagnosis.m File Reference

plotMCMCdiagnosis.m visualizes the Markov chains generated by getSamples.m.

##### Functions

- mlhsInnerSubst< matlabtypesubstitute, fh > plotMCMCdiagnosis (matlabtypesubstitute parameters, matlabtypesubstitute varargin)

    *plotMCMCdiagnosis.m visualizes the Markov chains generated by getSamples.m.*

##### 7.12.1 Detailed Description

plotMCMCdiagnosis.m visualizes the Markov chains generated by getSamples.m.

##### 7.12.2 Function Documentation

###### 7.12.2.1 mlhsInnerSubst< matlabtypesubstitute, fh > plotMCMCdiagnosis ( matlabtypesubstitute *parameters,* matlabtypesubstitute *varargin* )

plotMCMCdiagnosis.m visualizes the Markov chains generated by getSamples.m.

**USAGE**

fh = plotMCMCdiagnosis(parameters) fh = plotMCMCdiagnosis(parameters,type) fh = plotMCM↩ Cdiagnosis(parameters,type,fh) fh = plotMCMCdiagnosis(parameters,type,fh,I) fh = plotMCMCdiagnosis(parameters,type,fh,↩ I,options)

**History**

- 2014/06/20 Jan Hasenauer
- 2016/10/10 Daniel Weindl

###### Parameters

| | |
|---|---|
| *parameters* | parameter struct containing information about parameters and results of optimization (.MS) and uncertainty analysis (.S). This structures is the output of plotMultiStarts.m, getProfiles.m or plotSamples.m. |
| *varargin* | `1 plotMCMCdiagnosis ( ..., type, fh, I, options )` <br> *Required Parameters for varargin:* <br> • type string indicating the type of visualization: `parameters` (default) and `log-posterior` <br> • fh handle of figure. If no figure handle is provided, a new figure is opened. <br> • I index of parameters which are updated. If no index is provided all parameters are updated. <br> • options options of plotting as instance of PestoPlottingOptions |

###### Return values

| | |
|---|---|
| *fh* | figure handle |

###### Required fields of parameters:

Definition at line 17 of file plotMCMCdiagnosis.m.

#### 7.13 plotMultiStarts.m File Reference

plotMultiStarts plots the result of the multi-start optimization stored in parameters.

###### Functions

- mlhsInnerSubst< matlabtypesubstitute, fh > plotMultiStarts (matlabtypesubstitute parameters, matlabtype-substitute varargin)

  *plotMultiStarts plots the result of the multi-start optimization stored in parameters.*

##### 7.13.1 Detailed Description

plotMultiStarts plots the result of the multi-start optimization stored in parameters.

##### 7.13.2 Function Documentation

###### 7.13.2.1 mlhsInnerSubst< matlabtypesubstitute, fh > plotMultiStarts ( matlabtypesubstitute *parameters,* matlabtypesubstitute *varargin* )

plotMultiStarts plots the result of the multi-start optimization stored in parameters.

**USAGE**

fh = plotMultiStarts(parameters) fh = plotMultiStarts(parameters,fh) fh = plotMultiStarts(parameters,fh,options)

**plotMultiStarts() uses the following PestoPlottingOptions members**

- PestoPlottingOptions::add_points
- PestoPlottingOptions::title
- PestoPlottingOptions::draw_bounds

History:

- 2012/05/31 Jan Hasenauer
- 2016/10/07 Daniel Weindl

**Parameters**

<table>
<tr><td>parameters</td><td>parameter struct containing information about parameters and log-posterior.</td></tr>
<tr><td>varargin</td><td><pre>1 plotMultiStarts ( ..., fh, options )</pre>Required Parameters for varargin:<ul><li>fh handle of figure in which profile likelihood is plotted. If no figure handle is provided, a new figure is opened.</li><li>options options of plotting as instance of PestoPlottingOptions</li></ul></td></tr>
</table>

**Return values**

| | |
|---|---|
| *fh* | figure handle |

**Required fields of parameters:**

Definition at line 17 of file plotMultiStarts.m.

Referenced by collectResults(), and getMultiStarts().

Here is the caller graph for this function:

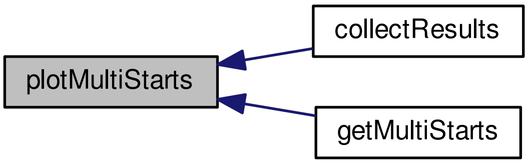

#### 7.14 plotParameterProfiles.m File Reference

plotParameterProfiles.m visualizes profile likelihood. Note: This routine provides an interface for plotUncertainty.m.

##### Functions

- mlhsInnerSubst< matlabtypesubstitute, fh > plotParameterProfiles (matlabtypesubstitute parameters, matlabtypesubstitute varargin)

  *plotParameterProfiles.m visualizes profile likelihood. Note: This routine provides an interface for plotUncertainty.m.*

##### 7.14.1 Detailed Description

plotParameterProfiles.m visualizes profile likelihood. Note: This routine provides an interface for plotUncertainty.m.

##### 7.14.2 Function Documentation

###### 7.14.2.1 mlhsInnerSubst< matlabtypesubstitute, fh > plotParameterProfiles ( matlabtypesubstitute *parameters,* matlabtypesubstitute *varargin* )

plotParameterProfiles.m visualizes profile likelihood. Note: This routine provides an interface for plotUncertainty.m.

**USAGE**

fh = plotParameterProfiles(parameters) fh = plotParameterProfiles(parameters,type) fh = plotParameter↩
Profiles(parameters,type,fh) fh = plotParameterProfiles(parameters,type,fh,I) fh = plotParameterProfiles(parameters,type,fh,↩
I,options)

**History**

- 2012/05/31 Jan Hasenauer
- 2014/06/20 Jan Hasenauer
- 2016/10/10 Daniel Weindl

**Parameters**

| | |
|---|---|
| *parameters* | parameter struct containing information about parameters and results of optimization (.MS) and uncertainty analysis (.P and .S). This structures is the output of plotMultiStarts.m, getProfiles.m or plotSamples.m. |
| *varargin* | `1 plotParameterProfiles ( ..., type, fh, I, options )`<br>*Required Parameters for varargin:*<br>• type string indicating the type of visualization: `1D` or `2D`<br>• fh handle of figure. If no figure handle is provided, a new figure is opened.<br>• I index of parameters which are updated. If no index is provided all parameters are updated.<br>• options options of plotting as instance of PestoPlottingOptions |

**Return values**

| | |
|---|---|
| *fh* | figure handle |

Definition at line 17 of file plotParameterProfiles.m.

References plotParameterUncertainty().

Referenced by collectResults(), and getParameterProfiles().

Here is the call graph for this function:

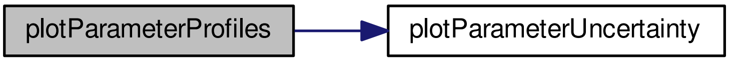

Here is the caller graph for this function:

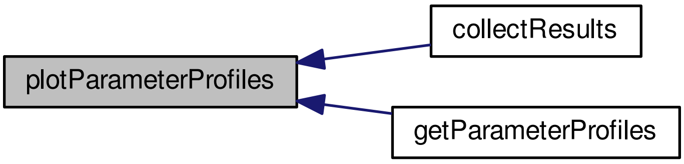

#### 7.15 plotParameterSamples.m File Reference

plotParameterSamples.m visualizes MCMC samples. Note: This routine provides an interface for plotUncertainty.m.

##### Functions

- mlhsInnerSubst< matlabtypesubstitute, fh > plotParameterSamples (matlabtypesubstitute parameters, matlabtypesubstitute varargin)

  *plotParameterSamples.m visualizes MCMC samples. Note: This routine provides an interface for plotUncertainty.m.*

##### 7.15.1 Detailed Description

plotParameterSamples.m visualizes MCMC samples. Note: This routine provides an interface for plotUncertainty.m.

##### 7.15.2 Function Documentation

###### 7.15.2.1 mlhsInnerSubst< matlabtypesubstitute, fh > plotParameterSamples ( matlabtypesubstitute *parameters,* matlabtypesubstitute *varargin* )

plotParameterSamples.m visualizes MCMC samples. Note: This routine provides an interface for plotUncertainty.m.

**USAGE**

fh = plotParameterSamples(parameters) fh = plotParameterSamples(parameters,type) fh = plotParameter←Samples(parameters,type,fh) fh = plotParameterSamples(parameters,type,fh,I) fh = plotParameter←Samples(parameters,type,fh,I,options)

**History**

- 2012/05/31 Jan Hasenauer
- 2014/06/20 Jan Hasenauer
- 2016/10/10 Daniel Weindl

**Parameters**

| | |
|---|---|
| *parameters* | parameter struct containing information about parameters and results of optimization (.MS) and uncertainty analysis (.P and .S). This structures is the output of plotMultiStarts.m, getProfiles.m or plotSamples.m. |
| *varargin* | `1 plotParameterSamples ( ..., type, fh, I, options )`<br>*Required Parameters for varargin:*<br>• type string indicating the type of visualization: `1D` or `2D`<br>• fh handle of figure. If no figure handle is provided, a new figure is opened.<br>• I index of parameters which are updated. If no index is provided all parameters are updated.<br>• options options of plotting as instance of PestoPlottingOptions |

**Return values**

| | |
|---|---|
| *fh* | figure handle |

Definition at line 17 of file plotParameterSamples.m.

References plotParameterUncertainty().

Referenced by getParameterSamples().

Here is the call graph for this function:

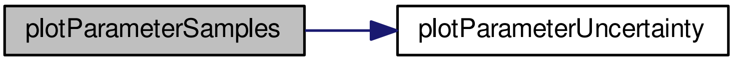

Here is the caller graph for this function:

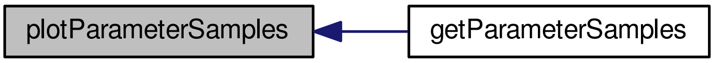

#### 7.16 plotParameterUncertainty.m File Reference

plotParameterUncertainty.m visualizes profile likelihood and MCMC samples stored in parameters.

##### Functions

- mlhsInnerSubst< matlabtypesubstitute, fh > plotParameterUncertainty (matlabtypesubstitute parameters, matlabtypesubstitute varargin)

  *plotParameterUncertainty.m visualizes profile likelihood and MCMC samples stored in parameters.*

##### 7.16.1 Detailed Description

plotParameterUncertainty.m visualizes profile likelihood and MCMC samples stored in parameters.

##### 7.16.2 Function Documentation

###### 7.16.2.1 mlhsInnerSubst< matlabtypesubstitute, fh > plotParameterUncertainty ( matlabtypesubstitute *parameters,* matlabtypesubstitute *varargin* )

plotParameterUncertainty.m visualizes profile likelihood and MCMC samples stored in parameters.

**USAGE**

fh = plotParameterUncertainty(parameters) fh = plotParameterUncertainty(parameters,type) fh = plot←ParameterUncertainty(parameters,type,fh) fh = plotParameterUncertainty(parameters,type,fh,I) fh = plot←ParameterUncertainty(parameters,type,fh,I,options)

**plotMultiStarts() uses the following PestoPlottingOptions members**

- PestoPlottingOptions::P
- PestoPlottingOptions::S
- PestoPlottingOptions::MS
- PestoPlottingOptions::boundary
- PestoPlottingOptions::subplot_size_1D
- PestoPlottingOptions::subplot_indexing_1D
- PestoPlottingOptions::CL
- PestoPlottingOptions::hold_on
- PestoPlottingOptions::interval
- PestoPlottingOptions::bounds
- PestoPlottingOptions::A
- PestoPlottingOptions::add_points
- PestoPlottingOptions::labels
- PestoPlottingOptions::legend
- PestoPlottingOptions::op2D
- PestoPlottingOptions::fontsize

**History**

- 2012/05/31 Jan Hasenauer
- 2014/06/20 Jan Hasenauer
- 2016/10/10 Daniel Weindl

**Parameters**

| | |
|---|---|
| *parameters* | parameter struct containing information about parameters and results of optimization (.MS) and uncertainty analysis (.P and .S). This structures is the output of plotMultiStarts.m, getProfiles.m or plotSamples.m. |
| *varargin* | `1 plotParameterUncertainty ( ..., type, fh, I, options )` <br> *Required Parameters for varargin:* <br> • type string indicating the type of visualization: `1D` or `2D` <br> • fh handle of figure. If no figure handle is provided, a new figure is opened. <br> • I index of parameters which are updated. If no index is provided all parameters are updated. <br> • options options of plotting as instance of PestoPlottingOptions |

**Return values**

| | |
|---|---|
| *fh* | figure handle |

**Required fields of parameters:**

Definition at line 17 of file plotParameterUncertainty.m.

Referenced by plotParameterProfiles(), and plotParameterSamples().

Here is the caller graph for this function:

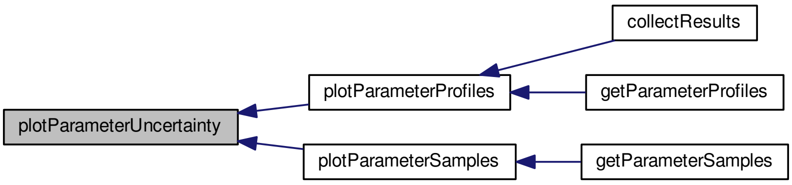

#### 7.17 plotPropertyMultiStarts.m File Reference

plotPropertyMultiStarts plots the result of the multi-start optimization stored in properties.

##### Functions

- mlhsInnerSubst< matlabtypesubstitute, fh > plotPropertyMultiStarts (matlabtypesubstitute properties, matlabtypesubstitute varargin)

    *plotPropertyMultiStarts plots the result of the multi-start optimization stored in properties.*

##### 7.17.1 Detailed Description

plotPropertyMultiStarts plots the result of the multi-start optimization stored in properties.

##### 7.17.2 Function Documentation

###### 7.17.2.1 mlhsInnerSubst< matlabtypesubstitute, fh > plotPropertyMultiStarts ( matlabtypesubstitute *properties*, matlabtypesubstitute *varargin* )

plotPropertyMultiStarts plots the result of the multi-start optimization stored in properties.

**USAGE**

fh = plotPropertyMultiStarts(properties) fh = plotPropertyMultiStarts(properties,fh) fh = plotPropertyMulti↵
Starts(properties,fh,options)

**History**

- 2015/03/03 Jan Hasenauer

**Parameters**

| | |
|---|---|
| *properties* | property struct containing information about properties and log-posterior. |
| *varargin* | `1 plotPropertyMultiStarts ( ..., fh, options )` <br> *Required Parameters for varargin:* <br> • fh handle of figure in which profile likelihood is plotted. If no figure handle is provided, a new figure is opened. <br> • options options of plotting as instance of PestoPlottingOptions |

**Return values**

| | |
|---|---|
| *fh* | figure handle |

**Required fields of properties:**

Definition at line 17 of file plotPropertyMultiStarts.m.

Referenced by collectResults(), and getPropertyMultiStarts().

Here is the caller graph for this function:

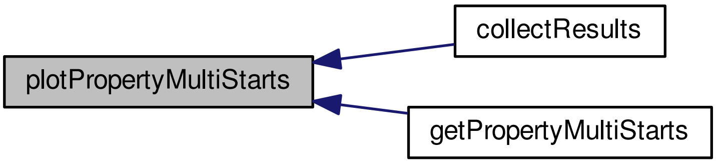

#### 7.18 plotPropertyProfiles.m File Reference

plotPropertyProfiles.m visualizes profile likelihood of model properties. Note: This routine provides an interface for plotPropertyUncertainty.m.

**Functions**

- mlhsInnerSubst< matlabtypesubstitute, fh > plotPropertyProfiles (matlabtypesubstitute properties, matlabtypesubstitute varargin)

  *plotPropertyProfiles.m visualizes profile likelihood of model properties. Note: This routine provides an interface for plotPropertyUncertainty.m.*

##### 7.18.1 Detailed Description

plotPropertyProfiles.m visualizes profile likelihood of model properties. Note: This routine provides an interface for plotPropertyUncertainty.m.

##### 7.18.2 Function Documentation

###### 7.18.2.1 mlhsInnerSubst< matlabtypesubstitute, fh > plotPropertyProfiles ( matlabtypesubstitute *properties*, matlabtypesubstitute *varargin* )

plotPropertyProfiles.m visualizes profile likelihood of model properties. Note: This routine provides an interface for plotPropertyUncertainty.m.

**USAGE**

fh = plotPropertyProfiles(properties) fh = plotPropertyProfiles(properties,type) fh = plotProperty↩ Profiles(properties,type,fh) fh = plotPropertyProfiles(properties,type,fh,I) fh = plotPropertyProfiles(properties,type,fh,↩ I,options)

**History**

- 2015/03/02 Jan Hasenauer
- 2016/10/10 Daniel Weindl

**Parameters**

| | |
|---|---|
| *properties* | property struct containing information about properties and results of optimization (.MS) and uncertainty analysis (.P and .S). |
| *varargin* | `1 plotPropertyProfiles ( ..., type, fh, I, options )` <br> *Required Parameters for varargin:* <br> • type string indicating the type of visualization: `1D` or `2D` <br> • fh handle of figure. If no figure handle is provided, a new figure is opened. <br> • I index of properties which are updated. If no index is provided all properties are updated. <br> • options options of plotting as instance of PestoPlottingOptions |

**Return values**

| | |
|---|---|
| *fh* | figure handle |

Definition at line 17 of file plotPropertyProfiles.m.

References plotPropertyUncertainty().

Referenced by collectResults(), and getPropertyProfiles().

Here is the call graph for this function:

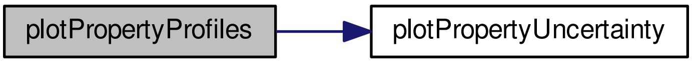

Here is the caller graph for this function:

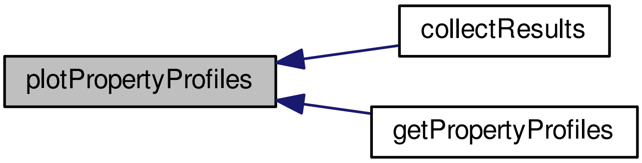

#### 7.19 plotPropertySamples.m File Reference

plotPropertySamples.m visualizes samples of model properties. Note: This routine provides an interface for plot←PropertyUncertainty.m.

##### Functions

- mlhsInnerSubst< matlabtypesubstitute, fh > plotPropertySamples (matlabtypesubstitute properties, matlab-typesubstitute varargin)

    *plotPropertySamples.m visualizes samples of model properties. Note: This routine provides an interface for plot←PropertyUncertainty.m.*

##### 7.19.1 Detailed Description

plotPropertySamples.m visualizes samples of model properties. Note: This routine provides an interface for plot←PropertyUncertainty.m.

##### 7.19.2 Function Documentation

###### 7.19.2.1 mlhsInnerSubst< matlabtypesubstitute, fh > plotPropertySamples ( matlabtypesubstitute *properties*, matlabtypesubstitute *varargin* )

plotPropertySamples.m visualizes samples of model properties. Note: This routine provides an interface for plot←PropertyUncertainty.m.

**USAGE**

fh = plotPropertySamples(properties) fh = plotPropertySamples(properties,type) fh = plotProperty←Samples(properties,type,fh) fh = plotPropertySamples(properties,type,fh,I) fh = plotPropertySamples(properties,type,fh,←I,options)

**History**

- 2015/04/01 Jan Hasenauer
- 2016/10/10 Daniel Weindl

**Parameters**

| | |
|---|---|
| *properties* | property struct containing information about properties and results of optimization (.MS) and uncertainty analysis (.P and .S). |
| *varargin* | `1 plotPropertySamples ( ..., type, fh, I, options )` *Required Parameters for varargin:* • type string indicating the type of visualization: `1D` or `2D` • fh handle of figure. If no figure handle is provided, a new figure is opened. • I index of properties which are updated. If no index is provided all properties are updated. • options options of plotting as instance of PestoPlottingOptions |

**Return values**

| | |
|---|---|
| *fh* | figure handle |

Definition at line 17 of file plotPropertySamples.m.

References plotPropertyUncertainty().

Referenced by getPropertySamples().

Here is the call graph for this function:

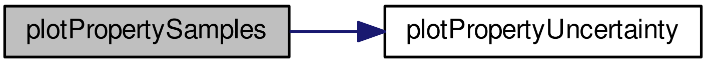

Here is the caller graph for this function:

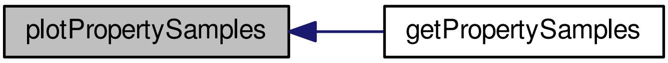

#### 7.20 plotPropertyUncertainty.m File Reference

plotPropertyUncertainty.m visualizes profile likelihood and MCMC samples stored in properties.

###### Functions

- mlhsInnerSubst< matlabtypesubstitute, fh > plotPropertyUncertainty (matlabtypesubstitute properties, matlabtypesubstitute varargin)

    *plotPropertyUncertainty.m visualizes profile likelihood and MCMC samples stored in properties.*

##### 7.20.1 Detailed Description

plotPropertyUncertainty.m visualizes profile likelihood and MCMC samples stored in properties.

##### 7.20.2 Function Documentation

###### 7.20.2.1 mlhsInnerSubst< matlabtypesubstitute, fh > plotPropertyUncertainty ( matlabtypesubstitute *properties,* matlabtypesubstitute *varargin* )

plotPropertyUncertainty.m visualizes profile likelihood and MCMC samples stored in properties.

**USAGE**

fh = plotPropertyUncertainty(properties,type) fh = plotPropertyUncertainty(properties,type,fh) fh = plotPropertyUncertainty(properties,type,fh,I) fh = plotPropertyUncertainty(properties,type,fh,I,options)

**History**

- 2012/05/31 Jan Hasenauer
- 2014/06/20 Jan Hasenauer
- 2016/10/10 Daniel Weindl

**Parameters**

| | |
|---|---|
| *properties* | properties struct. |
| *varargin* | `1 plotPropertyUncertainty ( ..., type, fh, I, options )`<br>*Required Parameters for varargin:*<br>• type string indicating the type of visualization: `1D`<br>• fh handle of figure. If no figure handle is provided, a new figure is opened.<br>• I index of properties which are updated. If no index is provided all parameters are updated.<br>• options options of plotting as instance of PestoPlottingOptions |

**Return values**

| | |
|---|---|
| *fh* | figure handle |

**Required fields of properties:**

Definition at line 17 of file plotPropertyUncertainty.m.

Referenced by plotPropertyProfiles(), and plotPropertySamples().

Here is the caller graph for this function:

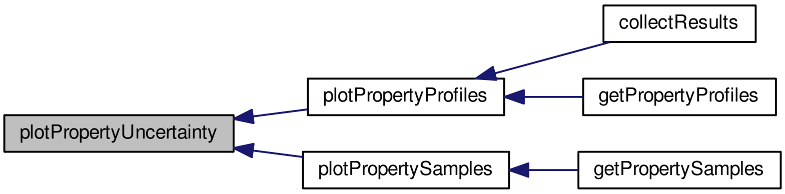

#### 7.21 runPestoTests.m File Reference

runPestoTests Run a set of PESTO unit tests

**Functions**

- noret::substitute runPestoTests ()
  *runPestoTests Run a set of PESTO unit tests*

###### 7.21.1 Detailed Description

runPestoTests Run a set of PESTO unit tests

#### 7.22 testGradient.m File Reference

testGradient.m calculates finite difference approximations to the gradient to check an analytical version.

##### Functions

- mlhsSubst< mlhsInnerSubst< matlabtypesubstitute, g >,mlhsInnerSubst< matlabtypesubstitute, g_fd_↩ f >,mlhsInnerSubst< matlabtypesubstitute, g_fd_b >,mlhsInnerSubst< matlabtypesubstitute, g_fd_c > > testGradient (matlabtypesubstitute varargin)

  *testGradient.m calculates finite difference approximations to the gradient to check an analytical version.*
- noret::substitute **mtoc_subst_testGradient_m_tsbus_cotm_error_plot** (matlabtypesubstitute g1, matlabtypesubstitute g2, matlabtypesubstitute ee)
- noret::substitute **mtoc_subst_testGradient_m_tsbus_cotm_ratio_plot** (matlabtypesubstitute g1, matlabtypesubstitute g2, matlabtypesubstitute rr, matlabtypesubstitute ee)

###### 7.22.1 Detailed Description

testGradient.m calculates finite difference approximations to the gradient to check an analytical version.

###### 7.22.2 Function Documentation

###### 7.22.2.1 mlhsSubst< mlhsInnerSubst< matlabtypesubstitute, g >,mlhsInnerSubst< matlabtypesubstitute, g_fd_f >,mlhsInnerSubst< matlabtypesubstitute, g_fd_b >,mlhsInnerSubst< matlabtypesubstitute, g_fd_c > > testGradient ( matlabtypesubstitute *varargin* )

testGradient.m calculates finite difference approximations to the gradient to check an analytical version.

backward differences: g_fd_f = (f(theta+eps∗e_i) - f(theta))/eps
forward differences: g_fd_b = (f(theta) - f(theta-eps∗e_i))/eps
central differences: g_fd_c = (f(theta+eps∗e_i) - f(theta-eps∗e_i))/(2∗eps)
in order to work with tensors of order n the gradient must be returned as tensor of order n+1 where the n+1th tensor dimension indexes the parameters with respect to which the differentiation was carried out

##### USAGE

[...] = testGradient(theta,fun,eps,il,ig) [g,g_fd_f,g_fd_b,g_fd_c] = testGradient(...)

##### History

- 2014/06/11 Jan Hasenauer
- 2015/01/16 Fabian Froehlich
- 2015/04/03 Jan Hasenauer
- 2015/07/28 Fabian Froehlich

**Parameters**

| | |
|---|---|
| *varargin* | `1 testGradient ( theta, fun, eps, il, ig )`<br>*Required Parameters for varargin:*<br>• theta parameter vector at which gradient is evaluated.<br>• fun function of theta for which gradients are checked.<br>• eps epsilon used for finite difference approximation of gradient (eps = 1e-4).<br>• il argout index/fieldname at which function values are returned (default = 1).<br>• ig argout index/fieldname at which gradient values are returned (default = 2). |

**Return values**

| | |
|---|---|
| *g* | gradient computed by f |
| *g_fd_f* | backward differences |
| *g_fd_b* | forward differences |
| *g_fd_c* | central differences |

Definition at line 17 of file testGradient.m.

### Index

A
PestoPlottingOptions, 34
adaptionInterval
DRAMOptions, 15
add_points
PestoPlottingOptions, 35
alpha
PHSOptions, 51
PTOptions, 55

boundary
PestoPlottingOptions, 35
boundary_tol
PestoOptions, 21
bounds
PestoPlottingOptions, 36

calc_profiles
PestoOptions, 21
checkDependentDefaults
PestoSamplingOptions, 47
CL
PestoPlottingOptions, 36
collectResults
collectResults.m, 57
collectResults.m, 57
collectResults, 57
comp_type
PestoOptions, 21

dR_max
PestoOptions, 22
DRAMOptions, 13
adaptionInterval, 15
DRAMOptions, 14
nTry, 15
regFactor, 15
verbosityMode, 15
dJ
PestoOptions, 21
draw_bounds
PestoPlottingOptions, 37

exponentT
PTOptions, 55

fh
PestoOptions, 22
fh_logPost_trace
PestoPlottingOptions, 37
fh_par_dis_1D
PestoPlottingOptions, 38
fh_par_dis_2D
PestoPlottingOptions, 38
fh_par_trace
PestoPlottingOptions, 38
foldername
PestoOptions, 22
fontsize
PestoPlottingOptions, 38

getMultiStarts
getMultiStarts.m, 59
getMultiStarts.m, 58
getMultiStarts, 59
getParameterConfidenceIntervals
getParameterConfidenceIntervals.m, 62
getParameterConfidenceIntervals.m, 61
getParameterConfidenceIntervals, 62
getParameterProfiles
getParameterProfiles.m, 63
getParameterProfiles.m, 63
getParameterProfiles, 63
getParameterSamples
getParameterSamples.m, 66
getParameterSamples.m, 65
getParameterSamples, 66
getPropertyConfidenceIntervals
getPropertyConfidenceIntervals.m, 67
getPropertyConfidenceIntervals.m, 67
getPropertyConfidenceIntervals, 67
getPropertyMultiStarts
getPropertyMultiStarts.m, 70
getPropertyMultiStarts.m, 69
getPropertyMultiStarts, 70
getPropertyProfiles
getPropertyProfiles.m, 72
getPropertyProfiles.m, 71
getPropertyProfiles, 72
getPropertySamples
getPropertySamples.m, 75
getPropertySamples.m, 74
getPropertySamples, 75
group_CI_by
PestoPlottingOptions, 38

hold_on
PestoPlottingOptions, 39

init_threshold
PestoOptions, 22
interval
PestoPlottingOptions, 39

labels
PestoPlottingOptions, 39
legend
PestoPlottingOptions, 40
localOptimizer
PestoOptions, 22
localOptimizerOptions
PestoOptions, 23

MALAOptions, 16
MALAOptions, 17
regFactor, 17
MAP_index
PestoOptions, 23
MCMC
PestoPlottingOptions, 40
mark_constraint
PestoPlottingOptions, 40
meigoDummy
meigoDummy.m, 76
meigoDummy.m, 76
meigoDummy, 76
memoryLength
PHSOptions, 52
PTOptions, 55
mode
PestoOptions, 23
PestoSamplingOptions, 47
MS
PestoPlottingOptions, 41

n_max
PestoPlottingOptions, 41
n_starts
PestoOptions, 24
nChains
PHSOptions, 52
nIterations
PestoSamplingOptions, 47
nTemps
PTOptions, 55
nTry
DRAMOptions, 15

obj_type
PestoOptions, 24
PestoSamplingOptions, 48
objOutNumber
PestoOptions, 24
PestoSamplingOptions, 48
op2D
PestoPlottingOptions, 42
options_getNextPoint
PestoOptions, 25

P
PestoOptions, 25
PestoPlottingOptions, 42
PHSOptions, 50
alpha, 51
memoryLength, 52
nChains, 52
PHSOptions, 51
regFactor, 52
trainingTime, 52
PTOptions, 53
alpha, 55
exponentT, 55
memoryLength, 55
nTemps, 55
PTOptions, 54
regFactor, 56
temperatureAdaptionScheme, 56
temperatureAlpha, 56
parameter_index
PestoOptions, 26
PestoOptions, 18
boundary_tol, 21
calc_profiles, 21
comp_type, 21
dR_max, 22
dJ, 21
fh, 22
foldername, 22
init_threshold, 22
localOptimizer, 22
localOptimizerOptions, 23
MAP_index, 23
mode, 23
n_starts, 24
obj_type, 24
objOutNumber, 24
options_getNextPoint, 25
P, 25
parameter_index, 26
PestoOptions, 20
plot_options, 26
profile_integ_index, 26
profile_method, 26
profile_optim_index, 27
profileReoptimizationOptions, 27
property_index, 27
proposal, 28
R_min, 28
resetobjective, 28
save, 29
solver, 29
start_index, 31
tempsave, 31
trace, 31
PestoPlottingOptions, 32
A, 34
add_points, 35
boundary, 35
bounds, 36
CL, 36
draw_bounds, 37
fh_logPost_trace, 37
fh_par_dis_1D, 38
fh_par_dis_2D, 38
fh_par_trace, 38
fontsize, 38
group_CI_by, 38
hold_on, 39
interval, 39
labels, 39

legend, 40
MCMC, 40
mark_constraint, 40
MS, 41
n_max, 41
op2D, 42
P, 42
PestoPlottingOptions, 34
plot_type, 42
S, 43
subplot_indexing_1D, 43
subplot_size_1D, 44
title, 44
PestoSamplingOptions, 45
checkDependentDefaults, 47
mode, 47
nIterations, 47
obj_type, 48
objOutNumber, 48
PestoSamplingOptions, 46
samplingAlgorithm, 48
sigma0, 49
theta0, 49
plot_options
PestoOptions, 26
plot_type
PestoPlottingOptions, 42
plotConfidenceIntervals
plotConfidenceIntervals.m, 77
plotConfidenceIntervals.m, 77
plotConfidenceIntervals, 77
plotMCMCdiagnosis
plotMCMCdiagnosis.m, 79
plotMCMCdiagnosis.m, 79
plotMCMCdiagnosis, 79
plotMultiStarts
plotMultiStarts.m, 81
plotMultiStarts.m, 80
plotMultiStarts, 81
plotParameterProfiles
plotParameterProfiles.m, 82
plotParameterProfiles.m, 82
plotParameterProfiles, 82
plotParameterSamples
plotParameterSamples.m, 84
plotParameterSamples.m, 84
plotParameterSamples, 84
plotParameterUncertainty
plotParameterUncertainty.m, 86
plotParameterUncertainty.m, 85
plotParameterUncertainty, 86
plotPropertyMultiStarts
plotPropertyMultiStarts.m, 87
plotPropertyMultiStarts.m, 87
plotPropertyMultiStarts, 87
plotPropertyProfiles
plotPropertyProfiles.m, 89
plotPropertyProfiles.m, 88
plotPropertyProfiles, 89
plotPropertySamples
plotPropertySamples.m, 91
plotPropertySamples.m, 90
plotPropertySamples, 91
plotPropertyUncertainty
plotPropertyUncertainty.m, 92
plotPropertyUncertainty.m, 92
plotPropertyUncertainty, 92
profile_integ_index
PestoOptions, 26
profile_method
PestoOptions, 26
profile_optim_index
PestoOptions, 27
profileReoptimizationOptions
PestoOptions, 27
property_index
PestoOptions, 27
proposal
PestoOptions, 28

R_min
PestoOptions, 28
regFactor
DRAMOptions, 15
MALAOptions, 17
PHSOptions, 52
PTOptions, 56
resetobjective
PestoOptions, 28
runPestoTests.m, 93

S
PestoPlottingOptions, 43
samplingAlgorithm
PestoSamplingOptions, 48
save
PestoOptions, 29
sigma0
PestoSamplingOptions, 49
solver
PestoOptions, 29
start_index
PestoOptions, 31
subplot_indexing_1D
PestoPlottingOptions, 43
subplot_size_1D
PestoPlottingOptions, 44

temperatureAdaptionScheme
PTOptions, 56
temperatureAlpha
PTOptions, 56
tempsave
PestoOptions, 31
testGradient
testGradient.m, 94
testGradient.m, 94

testGradient, 94
theta0
PestoSamplingOptions, 49
title
PestoPlottingOptions, 44
trace
PestoOptions, 31
trainingTime
PHSOptions, 52

verbosityMode
DRAMOptions, 15
