## Supplementary material for "Mechanistic description of spatial processes using integrative modelling of noise-corrupted imaging data": logLikelihoodCR.m

```
function [logL, dlogLdtheta, FIM] = logLikelihoodCR(theta, t, Y, sigma2, scale)
% Objective function for examples/conversion_reaction
%
% logLikelihood.m provides the log-likelihood, its gradient and an 
% approximation of the Hessian matrix based on Fisher information matrix
% (FIM) for the conversion reaction process.
% 
% Parameters:
%  theta: Model parameters [theta_1, theta_2]'
%  t: vector of time points
%  Y: measurement vector
%  sigma2: variance of the measurements (noise)
%  scale: 'lin' or 'log'
%
% Return values:
%  logL: double, value of log-likelihood
%  dlogLdtheta: double vector, gradient of log-likelihood
%  FIM: double array, negative Fisher information matrix, 
%   an approximation of Hessian of log-likelihood


%% Model Definition
% Definition of the biological process:
% Interconversion of two species (X_1 and X_2) following first-order mass 
% action kinetics with the parameters theta_1 and theta_2 respectively
%
% Reaction:
% X_1 -> X_2, rate = theta_1 * [X_1]
% X_2 -> X_1, rate = theta_2 * [X_2]
%
% State of the system:
% x   = [x_1, x_2] with
% x_1 = [X_1]
% x_2 = [X_2]
%
% Augmented state:
% (In order to get gradient information, the state of the system is
% augmented by forward sensitivities. Here, s_{theta_i}^j defines the 
% sensitivity of x_j = [X_j] with respect to parameter theta_i)
% X = [x_1; x_2; s_{theta_1}^1; s_{theta_1}^2; s_{theta_2}^1; s_{theta_2}^2]
%
% Measurement:
% Y = x_2
%
% Observables of the system:
% y(t) = h(theta, x(t))
%
% Measurement noise:
% We assume additive normally distributed noise with mean 0 and standard
% deviation 0.015 ( = sqrt(sigma2))
%
% Right hand side of the ODE of the system:
% f = @(t,x,theta) [- theta(1) * x(1) + theta(2) * x(2);...
%                   + theta(1) * x(1) - theta(2) * x(2)];

% Number of states, parameters, observables
n_x = 2;
n_theta = 2;
n_y = 1;

% Initial values
x0 = @(theta) [1; 0; 0; 0; 0; 0];

% Right hand side of the ODE (augmented system)
f = @(t,x,theta) [- theta(1) * x(1) + theta(2) * x(2);...
                  + theta(1) * x(1) - theta(2) * x(2);...
                  - theta(1) * x(3) + theta(2) * x(4) - x(1);...
                  + theta(1) * x(3) - theta(2) * x(4) + x(1);...
                  - theta(1) * x(5) + theta(2) * x(6) + x(2);...
                  + theta(1) * x(5) - theta(2) * x(6) - x(2)];

% Measurement function              
h = @(x,theta) x(:,2);
dhdx = @(x,theta) [0, 1];

%% Simulation and ODE Integration
% Initialization of observable sensitivities

dydtheta = zeros(length(t), n_theta * n_y);
switch scale
    case 'lin'
        odeOptions = odeset('RelTol', 1e-5, 'AbsTol', 1e-8);
        [~,X] = ode15s(@(t,x) f(t,x,theta), t, x0(theta), odeOptions);
        y = h(X(:,1:n_x), theta);
        for i = 1:n_theta
            dxdxi_i = X(:, i*n_x+(1:n_x));
            dydtheta(1:length(t), (i-1)*n_y+(1:n_y)) = dxdxi_i*dhdx(X, theta)';
        end
    case 'log'
        odeOptions = odeset('RelTol', 1e-5, 'AbsTol', 1e-8);
        [~,X] = ode15s(@(t,x) f(t, x, exp(theta)), t, x0(exp(theta)), odeOptions);
        y = h(X(:,1:n_x), exp(theta));
        for i = 1:n_theta
            dxdxi_i = X(:, i*n_x+(1:n_x))*exp(theta(i));
            dydtheta(1:length(t), (i-1)*n_y+(1:n_y)) = dxdxi_i*dhdx(X, exp(theta))';
        end
    otherwise
        error('Scale argument must be either "lin" or "log".')
end

%% Objective Function Evaluation
% Initialization of output values
logL = 0;
dlogLdtheta = zeros(n_theta, 1);
FIM = zeros(n_theta, n_theta);

% Assignment
for i = 1:n_y
    logL = logL - 0.5*sum(log(2*pi*sigma2) + (Y(:,i)-y(:,i)).^2/sigma2);
    dlogLdtheta = dlogLdtheta + dydtheta(:,i+(0:n_y:n_theta*n_y-i))' * ((Y(:,i)-y(:,i))/sigma2);
    FIM = FIM - dydtheta(:,i+(0:n_y:n_theta*n_y-i))' * dydtheta(:,i+(0:n_y:n_theta*n_y-i))/sigma2;
end

end
```
