## Supplementary material for "Mechanistic description of spatial processes using integrative modelling of noise-corrupted imaging data": mainConversionReaction.m

```
% Main file of the conversion reaction example
%
% Demonstrates the use of:
% * getMultiStarts()
% * getParameterProfiles()
% * getParameterSamples()
% * getParameterConfidenceIntervals()
% * getPropertyMultiStarts()
% * getPropertyProfiles()
% * getPropertySamples()
% * getPropertyConfidenceIntervals()
%
% Demonstrates furthermore:
% * how to use the MEIGO and the PSwarm toolbox for optimization (commented code version)
%
% This example provides a model for the interconversion of two species 
% (X_1 and X_2) following first-order mass action kinetics with the 
% parameters theta_1 and theta_2 respectively:
%
% * X_1 -> X_2, rate = theta_1 * [X_1]
% * X_2 -> X_1, rate = theta_2 * [X_2]
%
% Measurement of [X_2] are provided as: Y = [X_2]
%
% This file provides time-series measurement data Y and 
% performs a multistart maximum likelihood parameter estimation based on
% these measurements, demonstrating the use of getMultiStarts(). The model 
% fit is then visualized.
% 
% Profile likelihood calculation is done using getParameterProfiles() 
% applying a classical optimization algorithm and plotted.
%
% Single-chain Monte-Carlo sampling is performed by getParameterSamples() 
% and plotted in 1D and 2D.


%% Preliminary
clear all;
close all;
clc;

TextSizes.DefaultAxesFontSize = 14;
TextSizes.DefaultTextFontSize = 18;
set(0,TextSizes);

% Seed the random number generator. Seeding the random number generator
% ensures that everytime this example is run, the same sequence of random
% numbers is generated, and thus, the same starting points for multi-start 
% optimization will be used. This is helpful for debugging or comparing
% results across different machines. 
% Results might vary though if PestoOptions.comp_type is set to 'parallel'
rng(0);

%% Model Definition
% See logLikelihoodCR.m for a detailed description

%% Data
% We fix an artificial data set. It consists of a vector of time points t
% and a measurement vector y. This data was created using the parameter 
% values which are assigned to theta_true and by adding normally distributed 
% measurement noise with variance sigma2. 

% True parameters
theta_true = [-2.5;-2];

% Time points, measurement noise and measurement data
t = (0:10)';
sigma2 = 0.015^2;
y = [0.0244; 0.0842; 0.1208; 0.1724; 0.2315; 0.2634; ... 
    0.2831; 0.3084; 0.3079; 0.3097; 0.3324];


%% Definition of the Parameter Estimation Problem
% In order to run any PESTO routine, at least the parameters struct with 
% the fields shown here and the objective function need to be defined, 
% since they are mandatory for getMultiStarts, which is usually the first 
% routine needed for any parameter estimation problem

% parameters
parameters.name = {'log_{10}(k_1)','log_{10}(k_2)'};
parameters.min = [-7,-7];
parameters.max = [ 3, 3];
parameters.number = length(parameters.name);

% Log-likelihood function
objectiveFunction = @(theta) logLikelihoodCR(theta, t, y, sigma2, 'log');

% properties (parameter function to be analyzed)
properties.name = {'log_{10}(k_1)','log_{10}(k_2)',...
                   'log_{10}(k_1)-log_{10}(k_2)','log_{10}(k_1)^2',...
                   'x_2(t=3)','x_2(t=10)'};
properties.function = {@propertyFunction_theta1,...
                       @propertyFunction_theta2,...
                       @propertyFunction_theta1_minus_theta2,...
                       @propertyFunction_theta1_square,...
                       @(theta) propertyFunction_x2(theta,3,'log'),...
                       @(theta) propertyFunction_x2(theta,10,'log')};
properties.min = [-2.6;-2.2;-5;-10; 0; 0];
properties.max = [-2.4;-1.7; 5; 10; 1; 1];
properties.number = length(properties.name);


%% Multi-start local optimization
% A multi-start local optimization is performed within the bounds defined in
% parameters.min and .max in order to infer the unknown parameters from 
% measurement data. Therefore, a PestoOptions object is created and
% some of its properties are set accordingly.

% Options
optionsPesto = PestoOptions();
optionsPesto.obj_type = 'log-posterior';
optionsPesto.n_starts = 20;
optionsPesto.comp_type = 'sequential';
optionsPesto.mode = 'visual';
optionsPesto.plot_options.add_points.par = theta_true;
optionsPesto.plot_options.add_points.logPost = objectiveFunction(theta_true);
optionsPesto.plot_options.add_points.prop = nan(properties.number,1);
for j = 1 : properties.number
    optionsPesto.plot_options.add_points.prop(j) = properties.function{j}(optionsPesto.plot_options.add_points.par);
end

% The example can also be run in parallel mode: Uncomment this, if wanted
optionsMultistart.comp_type = 'parallel'; 
optionsMultistart.mode = 'text';
optionsMultistart.save = true; 
optionsMultistart.foldername = 'results';
n_workers = 4;

% Open parpool
if strcmp(optionsPesto.comp_type, 'parallel') && (n_workers >= 2)
    parpool(n_workers); 
else
    optionsPesto.comp_type = 'sequential';
end

% Optimization
parameters = getMultiStarts(parameters, objectiveFunction, optionsPesto);


%% Choosing different optimizers

% Besides the default fmincon local optimizer, alternative optimizers can be chosen. 
% Currently, PESTO provides an interface to MEIGO and PSwarm, which have to be installed separately.
% These algorithms aim at finding the global optimum, and therefore, a
% low number or a single optimizer run should be enough.

% The following uses the MEIGO toolbox with default settings:
% (Install MEIGO from http://gingproc.iim.csic.es/meigom.html and
% uncomment:

% MeigoOptions = struct(...
%     'maxeval', 1e4, ...
%     'local', struct('solver', 'fmincon', ...
%     'finish', 'fmincon', ...
%     'iterprint', 1) ...
%     );
% 
% optionsMultistartMeigo = optionsMultistart.copy();
% optionsMultistartMeigo.localOptimizer = 'meigo-ess';
% optionsMultistartMeigo.localOptimizerOptions = MeigoOptions;
% optionsMultistartMeigo.n_starts = 2;
% parameters = getMultiStarts(parameters, objectiveFunction, optionsMultistartMeigo);

% This section uses PSwarm, a particle swarm optimizer
% (Install from http://www.norg.uminho.pt/aivaz/pswarm/ and uncomment)

% optionsMultistartPSwarm = optionsMultistart.copy();
% optionsMultistartPSwarm.localOptimizer = 'pswarm';
% optionsMultistartPSwarm.n_starts = 10;
% parameters = getMultiStarts(parameters, objectiveFunction, optionsMultistartPSwarm);


%% Visualization of fit
% The measured data is visualized in plot, together with fit for the best
% parameter value found during getMultiStarts

if strcmp(optionsPesto.mode,'visual')
    % Simulation
    tsim = linspace(t(1),t(end),100);
    ysim = simulateConversionReaction(exp(parameters.MS.par(:,1)),tsim);

    % Plot: Fit
    figure('Name','Conversion reaction: Visualization of fit');
    plot(t,y,'bo'); hold on;
    plot(tsim,ysim,'r-'); 
    xlabel('time t');
    ylabel('output y');
    legend('data','fit');
end


%% Profile likelihood calculation -- Parameters
% The uncertainty of the estimated parameters is visualized by computing
% and plotting profile likelihoods. In getParameterProfiles, this is done
% by using repeated reoptimization, if standard setings are used.
parameters = getParameterProfiles(parameters, objectiveFunction, optionsPesto);


%% Markov Chain Monte Carlo sampling -- Parameters
% Values for the parameters are sampled by using an Parallel Tempering (PT)
% algorithm. This way, the underlying probability density of the parameter 
% distribution can be captured. Since only one temperature is used, this is
% effectively an adapted Metropolis algorithm single-chain algorithm.

% Building a struct covering all sampling options:
optionsPesto.MCMC.nIterations = 1e4;
optionsPesto.MCMC.mode = optionsPesto.mode;

% PT specific options:
optionsPesto.MCMC.samplingAlgorithm   = 'PT';
optionsPesto.MCMC.PT.nTemps           = 1;
optionsPesto.MCMC.PT.temperatureAdaptionScheme = 'Lacki15'; %'Vousden16'; 

% Initialize the chains by making use of the preceeding multi-start local
% optimization, all of them starting from the same point
optionsPesto.MCMC.theta0 = parameters.MS.par(:,1); 
optionsPesto.MCMC.sigma0 = 0.5 * inv(squeeze(parameters.MS.hessian(:,:,1)));

% Run the sampling
parameters = getParameterSamples(parameters, objectiveFunction, optionsPesto);


%% Confidence interval evaluation -- Parameters
% Confidence intervals to the confidence levels fixed in the array
% alpha
% are computed based on local approximations from the Hessian matrix at the
% optimum, based on the profile likelihoods and on the parameter sampling.

alpha = [0.9,0.95,0.99];
parameters = getParameterConfidenceIntervals(parameters, alpha, optionsPesto);


%% Evaluation of properties for multi-start local optimization results -- Properties
% The values of the properties are evaluated at the end points of the
% multi-start optimization runs by getPropertyMultiStarts.

optionsProperties = optionsPesto.copy();
optionsProperties.fh = [];
properties = getPropertyMultiStarts(properties,parameters,optionsProperties);


%% Profile likelihood calculation -- Properties
% Profile likelihoods are computed for the properties in the same fashion,
% as they were computed for the parameters.

properties = getPropertyProfiles(properties, parameters, objectiveFunction, optionsProperties);


%% Evaluation of properties for sampling results -- Properties
% From the samples of the parameters, the properties are calculated and
% hence a probability distribution for the properties can be reconstructed
% from that.

properties = getPropertySamples(properties, parameters, optionsProperties);


%% Confidence interval evaluation -- Properties
% As for the parameters, confidence intervals are computed for the
% properties in different fashion, based on local approximations, profile
% likelihoods and samples.

properties = getPropertyConfidenceIntervals(properties, alpha, optionsProperties);


%% Comparison of calculated parameter profiles

if strcmp(optionsPesto.mode, 'visual')
    % Open figure
    figure('Name','Conversion reaction: Comparison of parameter profiles');
    
    % Loop: parameters
    for i = 1:min(parameters.number, properties.number)
        subplot(ceil(parameters.number/ceil(sqrt(parameters.number))),ceil(sqrt(parameters.number)),i);
        plot(parameters.P(i).par(i,:),parameters.P(i).R,'bx-'); hold on;
        plot(properties.P(i).prop,properties.P(i).R,'r-o');
        xlabel(properties.name{i});
        ylabel('likelihood ratio');
        if i == 1
            legend({'unconst. opt. (= standard)','unconst. op. (= new)'},'color','none');
        end
    end
end


%% Close the pools of parallel working threads

if strcmp(optionsPesto.comp_type, 'parallel') && (n_workers >= 2)
    delete(gcp('nocreate'))
end
```
