## Supplementary material for "Mechanistic description of spatial processes using integrative modelling of noise-corrupted imaging data": propertyFunction_theta2.m

```
function [f, grad_f, hess_f] = propertyFunction_theta2(theta)
% propertyFunction_theta2 for examples/conversion_reaction
%
% returns the the second reaction rate as defined in logLikelihood.m with 
% derivatives
%
% Parameters: 
%  theta: Model parameters [theta_1, theta_2]'
%
% Return values:
%  f: double, value of property function
%  grad_f: double vector, gradient of property function
%  hess_f: double array, hessian of property function


%% Property Function Evaluation
f = theta(2);
grad_f = [0; 1];
hess_f = [0, 0; 0, 0];

end
```
