## Supplementary material for "Mechanistic description of spatial processes using integrative modelling of noise-corrupted imaging data": propertyFunction_x2.m

```
function [f, grad_f] = propertyFunction_x2(theta, T, scale)
% propertyFunction_x2 for examples/conversion_reaction
%
% returns the value of x_2 at time T as defined in logLikelihood.m with 
% derivatives
%
% Parameters: 
%  theta: Model parameters [theta_1, theta_2]'
%  T: stopping time for simulation
%  scale: 'lin' or 'log'
%
% Return values:
%  f: double, value of property function
%  grad_f: double vector, gradient of property function


%% Model Definition
% For more details, see logLikelihood.m

% Initial values
x0 = @(theta) [1; 0; 0; 0; 0; 0];

% Right hand side of the ODE
f = @(t,x,theta) [- theta(1) * x(1) + theta(2) * x(2);...
                  + theta(1) * x(1) - theta(2) * x(2);...
                  - theta(1) * x(3) + theta(2) * x(4) - x(1);...
                  + theta(1) * x(3) - theta(2) * x(4) + x(1);...
                  - theta(1) * x(5) + theta(2) * x(6) + x(2);...
                  + theta(1) * x(5) - theta(2) * x(6) - x(2)];

%% Simulation and ODE Integration
switch scale
    case 'lin'
        [~,X] = ode15s(@(t,x) f(t,x,theta),[0,T],x0(theta));
    case 'log'
        [~,X] = ode15s(@(t,x) f(t,x,exp(theta)),[0,T],x0(exp(theta)));
        X(:,3:4) = exp(theta(1))*X(:,3:4);
        X(:,5:6) = exp(theta(2))*X(:,5:6);
end

%% Property Function Evaluation
% Assignment
f = X(end,2);
grad_f = [X(end,4); X(end,6)];

end
```
