## Supplementary material for "Mechanistic description of spatial processes using integrative modelling of noise-corrupted imaging data": simulateConversionReaction.m

```
function [y] = simulateConversionReaction(theta, t)
% simulateConversionReaction for examples/conversion_reaction
%
% simulateConversionReaction performs a simulation of the conversion 
% reaction model for the given timepoints t and parameters theta
%
% Parameters:
%  theta: Model parameters [theta_1, theta_2]'
%  t: vector of time points
%
% Return values:
%  y: double vector, values of the observable Y = [x_2] at time points t


%% Model Definition
% Initial values
x0 = @(theta) [1;0];

% Right hand side of the ODE
f = @(t,x,theta) [- theta(1) * x(1) + theta(2) * x(2);...
                  + theta(1) * x(1) - theta(2) * x(2)];

% Measurement function 
h = @(x,theta) x(:,2);

%% Simulation and Assignment
[~,X] = ode15s(@(t,x) f(t,x,theta),t,x0(theta));
y = h(X, theta);

end
```
