## Supplementary material for "Mechanistic description of spatial processes using integrative modelling of noise-corrupted imaging data": getMeasuredData.m

```
function yMeasure = getMeasuredData()

yMeasure = nan(1, 50 , 2);
yMeasure(1, 1, :) = [1.3244      1.5386];
yMeasure(1, 2, :) = [0.58495      1.6576];
yMeasure(1, 3, :) = [0.34931      1.6084];
yMeasure(1, 4, :) = [0.2424      1.7396];
yMeasure(1, 5, :) = [0.32102      1.7862];
yMeasure(1, 6, :) = [0.23102       1.929];
yMeasure(1, 7, :) = [-0.0050333       1.892];
yMeasure(1, 8, :) = [0.19572      2.0183];
yMeasure(1, 9, :) = [0.068784       2.111];
yMeasure(1, 10, :) = [0.022116      2.2016];
yMeasure(1, 11, :) = [0.056302      2.2833];
yMeasure(1, 12, :) = [0.0071388      2.2664];
yMeasure(1, 13, :) = [0.0089219      2.2403];
yMeasure(1, 14, :) = [0.087921      2.3273];
yMeasure(1, 15, :) = [0.082555      2.3578];
yMeasure(1, 16, :) = [0.081864      2.5721];
yMeasure(1, 17, :) = [0.043644      2.4648];
yMeasure(1, 18, :) = [-0.051116      2.5714];
yMeasure(1, 19, :) = [0.044404      2.5605];
yMeasure(1, 20, :) = [0.089414      2.6484];
yMeasure(1, 21, :) = [0.031771      2.5976];
yMeasure(1, 22, :) = [0.058539      2.5955];
yMeasure(1, 23, :) = [0.042675      2.6225];
yMeasure(1, 24, :) = [-0.0092716      2.7443];
yMeasure(1, 25, :) = [0.020202      2.7356];
yMeasure(1, 26, :) = [-0.034216      2.7577];
yMeasure(1, 27, :) = [0.049238      2.8602];
yMeasure(1, 28, :) = [-0.052839      2.8241];
yMeasure(1, 29, :) = [-0.049209      2.8384];
yMeasure(1, 30, :) = [-0.036498      2.9258];
yMeasure(1, 31, :) = [-0.14347      2.8229];
yMeasure(1, 32, :) = [0.075439      2.9137];
yMeasure(1, 33, :) = [0.019577      2.9354];
yMeasure(1, 34, :) = [-0.034617      2.8952];
yMeasure(1, 35, :) = [0.071471      2.9312];
yMeasure(1, 36, :) = [-0.08278      2.8742];
yMeasure(1, 37, :) = [-0.0024658      2.8865];
yMeasure(1, 38, :) = [-0.009565      2.9598];
yMeasure(1, 39, :) = [0.018339      3.0007];
yMeasure(1, 40, :) = [0.017902      3.1033];
yMeasure(1, 41, :) = [-0.041097      2.9494];
yMeasure(1, 42, :) = [0.00054081      3.0004];
yMeasure(1, 43, :) = [-0.0062971      2.9947];
yMeasure(1, 44, :) = [0.033242      2.9094];
yMeasure(1, 45, :) = [0.056436      2.9909];
yMeasure(1, 46, :) = [0.057158      2.9295];
yMeasure(1, 47, :) = [-0.041561      3.0672];
yMeasure(1, 48, :) = [0.0054217      2.9864];
yMeasure(1, 49, :) = [-0.059216      3.0411];
yMeasure(1, 50, :) = [-0.054244      3.0138];

 end
```
