## Supplementary material for "Mechanistic description of spatial processes using integrative modelling of noise-corrupted imaging data": logLikelihoodEC.m

```
function varargout = logLikelihoodEC(theta, yMeasured, sigma2, con0, nTimepoints, nMeasure)
% Objective function for examples/enzymatic_catalysis
%
% logLikelihood.m provides the log-likelihood, its gradient and an 
% approximation of the Hessian matrix based on Fisher information matrix
% (FIM) for the enzymatic catalysis example.
% 
% Parameters:
%  theta: Model parameters [theta_1, theta_2, theta_3, theta_4]'
%  yMeasured: measurement array returned from getMeasuredData()
%  sigma2: variance of the measurements (noise)
%  con0: inititial concentrations of the experiments returnd 
%    from getInitialConcentrations()
%  nTimepoints: number of Time points (equidistad between 0 and 5)
%  nMeasure: number of experiments
%
% Return values:
%  J: double, value of log-likelihood
%  gradJ: double vector, gradient of log-likelihood
%  FIM: double array, negative Fisher information matrix, 
%    an approximation of Hessian of log-likelihood


%% Model Definition
% Definition of the biological process:
% A species X_1 is bound to an enzyme X_2. They form a complex X_3 which 
% falls apart to the product X_4 and the enzyme X_2 again. The reaction 
% follows first-order mass action kinetics with the parameters theta_1 to
% theta_4
%
% Reaction:
% * X_1 + X_2 -> X_3, rate = theta_1 * [X_1] * [X_2]
% * X_3 -> X_1 + X_2, rate = theta_2 * [X_3]
% * X_3 -> X_4 + X_2, rate = theta_3 * [X_3]
% * X_4 + X_2 -> X_3, rate = theta_4 * [X_4] * [X_2]
%
% State of the system:
% x = [x_1; x_2; x_3; x_4]
% x_1 = [X_1]
% x_2 = [X_2]
% x_3 = [X_3]
% x_4 = [X_4]
%
% Augmented state:
% (In order to get gradient information, the state of the system is
% augmented by forward sensitivities. Here, s_{theta_i}^j defines the 
% sensitivity of x_j = [X_j] with respect to parameter theta_i)
% X = [x_1; x_2; s_{theta_1}^1; s_{theta_1}^2; ...; s_{theta_2}^1; s_{theta_2}^2; ...]
%
% Measurements:
% Y = [x_1; x_4]
%
% Observables of the system:
% y(t) = h(theta, x(t))
%
% Measurement noise:
% We assume additive normally ditributed noise with mean 0 and standard
% deviation sqrt(sigma2)
%
% Right hand side of the original ODE of the system:
% f = @(theta, x) [...
%     - theta(1)*x(1)*x(2) + theta(2)*x(3);...
%     - theta(1)*x(1)*x(2) + (theta(2)+theta(3))*x(3) - theta(4)*x(2)*x(4);...
%       theta(1)*x(1)*x(2) - (theta(2)+theta(3))*x(3) + theta(4)*x(2)*x(4);...
%       theta(3)*x(3) - theta(4)*x(2)*x(4)]

% Number of states, parameters, obeservables
nStates = 4;
nParams = 4;
nObserv = 2;

% Creation of the time vector and initialization for return values
t     = linspace(0, 5, nTimepoints)';
Y     = nan(nTimepoints, nObserv);
J     = 0;
gradJ = zeros(nParams, 1);
FIM   = zeros(nParams, nParams);

if (nargout > 1)
% Right hand side of the ODE (augmented system)
f = @(theta, x) [...
    - theta(1)*x(1)*x(2) + theta(2)*x(3);...
    - theta(1)*x(1)*x(2) + (theta(2)+theta(3))*x(3) - theta(4)*x(2)*x(4);...
      theta(1)*x(1)*x(2) - (theta(2)+theta(3))*x(3) + theta(4)*x(2)*x(4);...
      theta(3)*x(3) - theta(4)*x(2)*x(4);...
    - x(1)*x(2) - theta(1)*(x(5)*x(2)+x(1)*x(6)) + theta(2)*x(7);...
    - x(1)*x(2) - theta(1)*(x(5)*x(2)+x(1)*x(6)) + (theta(2)+theta(3))*x(7) - theta(4)*(x(6)*x(4)+x(2)*x(8));...
      x(1)*x(2) + theta(1)*(x(5)*x(2)+x(1)*x(6)) - (theta(2)+theta(3))*x(7) + theta(4)*(x(6)*x(4)+x(2)*x(8));...
      theta(3)*x(7) - theta(4)*(x(6)*x(4)+x(2)*x(8));...
      x(3) - theta(1)*(x(9)*x(2)+x(1)*x(10)) + theta(2)*x(11);...
      x(3) - theta(1)*(x(9)*x(2)+x(1)*x(10)) + (theta(2)+theta(3))*x(11) - theta(4)*(x(10)*x(4)+x(2)*x(12));...
    - x(3) + theta(1)*(x(9)*x(2)+x(1)*x(10)) - (theta(2)+theta(3))*x(11) + theta(4)*(x(10)*x(4)+x(2)*x(12));...
      theta(3)*x(11) - theta(4)*(x(10)*x(4)+x(2)*x(12));...
    - theta(1)*(x(13)*x(2)+x(1)*x(14)) + theta(2)*x(15);...
      x(3) - theta(1)*(x(13)*x(2)+x(1)*x(14)) + (theta(2)+theta(3))*x(15) - theta(4)*(x(14)*x(4)+x(2)*x(16));...
    - x(3) + theta(1)*(x(13)*x(2)+x(1)*x(14)) - (theta(2)+theta(3))*x(15) + theta(4)*(x(14)*x(4)+x(2)*x(16));...
      x(3) + theta(3)*x(15) - theta(4)*(x(14)*x(4)+x(2)*x(16));...
    - theta(1)*(x(17)*x(2)+x(1)*x(18)) + theta(2)*x(19);...
    - x(2)*x(4) - theta(1)*(x(17)*x(2)+x(1)*x(18)) + (theta(2)+theta(3))*x(19) - theta(4)*(x(18)*x(4)+x(2)*x(20));...
      x(2)*x(4) + theta(1)*(x(17)*x(2)+x(1)*x(18)) - (theta(2)+theta(3))*x(19) + theta(4)*(x(18)*x(4)+x(2)*x(20));...
    - x(2)*x(4) + theta(3)*x(19) - theta(4)*(x(18)*x(4)+x(2)*x(20))];
    fillX = zeros(16,1);
else
    f = @(theta, x) [...
        - theta(1)*x(1)*x(2) + theta(2)*x(3);...
        - theta(1)*x(1)*x(2) + (theta(2)+theta(3))*x(3) - theta(4)*x(2)*x(4);...
          theta(1)*x(1)*x(2) - (theta(2)+theta(3))*x(3) + theta(4)*x(2)*x(4);...
          theta(3)*x(3) - theta(4)*x(2)*x(4)];
    fillX = [];
end

% Measurement functions
h = @(x,theta) [x(:,1), x(:,4)];
dhdx = @(x,theta) [1, 0, 0, 0; 0, 0, 0, 1];
    

%% Simulation and ODE Integration
% Initialization of observable sensitivities
dydtheta = zeros(length(t), nParams * nObserv);

% Loop over the experiments and simulation for each experiment
for iMeasure = 1 : nMeasure
    % Simulation
    odeOptions = odeset('RelTol', 1e-6, 'AbsTol', 1e-9);
    [~,X] = ode15s(@(t,x) f(exp(theta),x), t, [con0(:,iMeasure); fillX], odeOptions);
    y = h(X(:,1:nStates), exp(theta));
    Y(:, :) = yMeasured(iMeasure, :, :);
    if (nargout > 1)
        for iTheta = 1 : nParams
            dxdxi_i = X(:, iTheta * nStates + (1:nStates)) * exp(theta(iTheta));
            dydtheta(1:length(t), (iTheta-1) * nObserv + (1 : nObserv)) = dxdxi_i * dhdx(X, exp(theta))';
        end
    end

    % Computation and assignment of the objective function and its
    % derivatives
    for iObserv = 1 : nObserv
        J = J - 0.5 * sum(log(2*pi*sigma2) + (Y(:,iObserv) - y(:,iObserv)).^2 / sigma2);
        if (nargout > 1)
            gradJ = gradJ + dydtheta(:,iObserv + (0 : nObserv : nParams*nObserv - iObserv))' * ((Y(:,iObserv) - y(:,iObserv)) / sigma2);
            FIM = FIM + dydtheta(:,iObserv + (0 : nObserv : nParams*nObserv - iObserv))' * dydtheta(:,iObserv + (0 : nObserv : nParams*nObserv - iObserv))/sigma2;
        end
    end
end

% Normalization by the number of experiments
varargout{1} = J / nMeasure;
if (nargout > 1)
    varargout{2} = gradJ / nMeasure;
    varargout{3} = FIM / nMeasure;
end

end
```
