## Supplementary material for "Mechanistic description of spatial processes using integrative modelling of noise-corrupted imaging data": mainEnzymaticCatalysis.m

```
% Main file of the enzymatic catalysis example
%
% Demonstrates the use of:
% * getParameterSamples()
% * getMultiStarts()
% * getParameterProfiles()
% * getParameterConfidenceIntervals()
%
% Demonstrates furthermore:
% * how to do sampling without multi-start local optimization beforehand
% * how to use the MEIGO toolbox for optimization (commented)
% * how to compute profile likelihoods via the integration method
% * how to use plotting functions outside the get... routines
% * the reliability of sampling and profiling in the case of
%   non-identifiabilites
% * how to use diagnosis tool (e.g. plotMCMCDiagnosis and
%   plotMultiStartDiagnosis)
%
% This example provides a model for the reaction of a species X_1 to a
% species X_4, which is catalyzed by an enzyme X_2.
%
% * X_1 + X_2 -> X_3, rate = theta_1 * [X_1] * [X_2]
% * X_3 -> X_1 + X_2, rate = theta_2 * [X_3]
% * X_3 -> X_4 + X_2, rate = theta_3 * [X_3]
% * X_4 + X_2 -> X_3, rate = theta_4 * [X_4] * [X_2]
%
% Measurements of [X_1] and [X_4] are provided as: Y = [[X_1]; [X_4]]
%
% This file sets a parameter vector, creates and saves artificial
% measurement data as a time series and performs a multi-start local
% optimization based on these measurements, demonstrating the use of
% getMultiStarts().
%
% Parameter sampling is done first without prior information from
% optimization, then with information from optimization.
%
% Parameter optimization is done using multi-start local optimization.
%
% The Profile likelihoods are calculated by integrating an ODE following
% the profile path using getParameterProfiles with the option
% optionsPesto.profile_method = 'integration'.


%% Preliminary
clear;
close all;
clc;

TextSizes.DefaultAxesFontSize = 14;
TextSizes.DefaultTextFontSize = 18;
set(0,TextSizes);

% Seed random number generator
rng(0);

%% Model Definition
% See logLikelihood.m for a detailed description

%% Create Artificial Data for Parameter Estimation
% The necessary variables are set (Parameter bounds, variance, ...)
nTimepoints = 50;      % Time points of Measurement
nMeasure    = 1;        % Number of experiments
sigma2      = 0.05^2;   % Variance of Measurement noise
lowerBound  = -10;      % Lower bound for parameters
upperBound  = 5;        % Upper bound for parameters
theta       = [1.1770; -2.3714; -0.4827; -5.5387]; % True parameter values

% Creation of data
% Once the two files getMeasuredData.m and getInitialConcentrations.m are
% written, the two following lines can be commented
% display(' Write new measurement data...');
% performNewMeasurement(theta, nMeasure, nTimepoints, sigma2);

% The measurement data is read out from the files where it is saved
yMeasured = getMeasuredData();
con0 = getInitialConcentrations();

%% Generation of the structs and options for PESTO
% The structs and the PestoOptions object, which are necessary for the 
% PESTO routines to work are created and set to convenient values

% parameters
display(' Prepare structs and options...')
parameters.name   = {'log(theta_1)', 'log(theta_2)', 'log(theta_3)', 'log(theta_4)'};
parameters.min    = lowerBound * ones(4, 1);
parameters.max    = upperBound * ones(4, 1);
parameters.number = length(parameters.name);

% objective function
objectiveFunction = @(theta) logLikelihoodEC(theta, yMeasured, sigma2, con0, nTimepoints, nMeasure);

% PestoOptions
optionsPesto           = PestoOptions();
optionsPesto.obj_type  = 'log-posterior';
optionsPesto.comp_type = 'sequential'; 
optionsPesto.mode      = 'visual';
optionsPesto.plot_options.add_points.par = theta;
optionsPesto.plot_options.add_points.logPost = objectiveFunction(theta);

%% Parameter Sampling
% Covering all sampling options in one struct
display(' Sampling without prior information...');
optionsPesto.MCMC.nIterations  = 1e4;

% PT (with only 1 chain -> AM) specific options:
optionsPesto.MCMC.samplingAlgorithm = 'PT';
optionsPesto.MCMC.PT.nTemps         = 6;
optionsPesto.MCMC.PT.exponentT      = 6;    
optionsPesto.MCMC.PT.regFactor      = 1e-8;
optionsPesto.MCMC.PT.temperatureAdaptionScheme = 'Lacki15'; %'Vousden16';

% Initialize the chains by choosing a random initial point and a 'large'
% covariance matrix
optionsPesto.MCMC.theta0 = lowerBound * ones(4, 1) + ...
    (upperBound * ones(4, 1) - lowerBound * ones(4, 1)) .* rand(4,1); 
optionsPesto.MCMC.sigma0 = 1e5 * eye(4);

% Run the sampling
parameters = getParameterSamples(parameters, objectiveFunction, optionsPesto);

% Use a diagnosis tool to see, how plotting worked (see burn-in etc.)
plotMCMCdiagnosis(parameters, 'parameters');

%% Calculate Confidence Intervals
% Confidence Intervals for the Parameters are inferred from the local 
% optimization and the sampling information.

% Set alpha levels
alpha = [0.8, 0.9, 0.95, 0.99];

display(' Computing confidence intervals...');
parameters = getParameterConfidenceIntervals(parameters, alpha, optionsPesto);


%% Perform Multistart optimization
% A multi-start local optimization is performed within the bound defined in
% parameters.min and .max in order to infer the unknown parameters from 
% measurement data.

% The following section uses the MEIGO toolbox with following settings:
% (Install MEIGO from http://gingproc.iim.csic.es/meigom.html and
% uncomment:

MeigoOptions = struct(...
    'maxeval', 1000, ...
    'local', struct('solver', 'fmincon', ...
    'finish', 'fmincon', ...
    'iterprint', 0) ...
    );

optionsPesto.localOptimizer = 'meigo-ess';
optionsPesto.localOptimizerOptions = MeigoOptions;
optionsPesto.n_starts = 3;
parameters = getMultiStarts(parameters, objectiveFunction, optionsPesto);

% % Options for an alternative multi-start local optimization
% display(' Optimizing parameters...');
% optionsPesto.n_starts = 5;
% parameters = getMultiStarts(parameters, objectiveFunction, optionsPesto);

% Use a diagnosis tool to see, how optimization worked
plotMultiStartDiagnosis(parameters);


%% Calculate Confidence Intervals
% Confidence Intervals for the Parameters are inferred from the local 
% optimization and the sampling information.
display(' Computing confidence intervals...');
parameters = getParameterConfidenceIntervals(parameters, alpha, optionsPesto);


%% Calculate Profile Likelihoods
% The result of the sampling is compared with profile likelihoods.
optionsPesto.profile_method = 'integration';
optionsPesto.solver.gamma = 1;
optionsPesto.objOutNumber = 2;
optionsPesto.solver.hessian = 'user-supplied';

display(' Computing parameter profiles...');
parameters = getParameterProfiles(parameters, objectiveFunction, optionsPesto);


%% Do additional plots
% In order to check how well sampling and profiling agree with each other,
% we do two additional plots.
PlottingOptionsSampling = PestoPlottingOptions();
PlottingOptionsSampling.S.plot_type = 1;
PlottingOptionsSampling.S.ind = 1;

fh = figure('Name','plotParameterSamples - 1D');
plotParameterSamples(parameters,'1D',fh,[],PlottingOptionsSampling);

fh = figure('Name','plotParameterSamples - 2D');
plotParameterSamples(parameters,'2D',fh,[],PlottingOptionsSampling);

%% Perform a second Sampling, now based on Multistart Optimization
% To compare the effect of previous multi-start optimization, we perform a
% second sampling.
optionsPesto.MCMC.theta0 = parameters.MS.par(:,1); 
optionsPesto.MCMC.sigma0 = 0.5*inv(squeeze(parameters.MS.hessian(:,:,1)));

% Run the sampling
display(' Sampling with information from optimization...');
optionsPesto.MCMC.nIterations  = 2e3;
parametersNew = parameters;
parametersNew = getParameterSamples(parametersNew, objectiveFunction, optionsPesto);


%% Calculate Confidence Intervals
% Confidence Intervals for the Parameters are inferred from the local 
% optimization, the sampling and the profile information.
parameters = getParameterConfidenceIntervals(parametersNew, alpha, optionsPesto);
```
