## Supplementary material for "Mechanistic description of spatial processes using integrative modelling of noise-corrupted imaging data": performNewMeasurement.m

```
function performNewMeasurement(theta, nMeasure, nTimepoints, sigma2)
% performNewMeasurement.m for examples/enzymatic_catalysis
%
% creates artificial data for the parameter estimation of the enzymatic 
% catalysis example for a given number of time points and a given number of
% measurements with different initial conditions of the chemical species.
% 
% Parameters:
%  theta: Model parameters [theta_1, theta_2, theta_3, theta_4]'
%  nMeasure: number of experiments
%  nTimepoints: number of Time points (equidistad between 0 and 5)
%  sigma2: variance of the measurements (noise)
%
% Return values:
%  No return values, but the files getInitialConcentrations.m and
%   getMeasuredData.m are written to the folder of this example
    


%% Model Definition
% For more details, see logLikelihood.m

%% Creation of initial concentrations
% Writing the initial concentrations using a log-normal distribution
con0 = exp(normrnd(0, 0.5, 4, nMeasure));

% Create file with initial concentrations
[exdir, ~, ~] = fileparts(which('performNewMeasurement.m'));
fid = fopen([exdir '/getInitialConcentrations.m'], 'w');
fprintf(fid, 'function con0 = getInitialConcentrations()\n\n');
fprintf(fid, ['    con0 = nan(4, ' num2str(nMeasure) ');\n']);

% Write initial concentrations
for iMeasure = 1 : nMeasure
    fprintf(fid, ['    con0(:, ' num2str(iMeasure) ') = [']);
    fprintf(fid, '%11.7f; %11.7f; %11.7f; %11.7f];\n', ...
        con0(1, iMeasure), con0(2, iMeasure), con0(3, iMeasure), con0(4, iMeasure));
end

% Close file
fprintf(fid, '\n end');
fclose(fid);

%% Creation of measurement data
% Right hand side of the ODE
f = @(theta, x) [...
    - theta(1)*x(1)*x(2) + theta(2)*x(3);...
    - theta(1)*x(1)*x(2) + (theta(2)+theta(3))*x(3) - theta(4)*x(2)*x(4);...
      theta(1)*x(1)*x(2) - (theta(2)+theta(3))*x(3) + theta(4)*x(2)*x(4);...
      theta(3)*x(3) - theta(4)*x(2)*x(4)];

% Creation of the time vector and the observable function
t = linspace(0, 5, nTimepoints);
h = @(x,theta) [x(:,1), x(:,4)];

% Create file with measurement data
fid = fopen([exdir '/getMeasuredData.m'], 'w');
fprintf(fid, 'function yMeasure = getMeasuredData()\n\n');
fprintf(fid, ['yMeasure = nan(' num2str(nMeasure) ', ' num2str(nTimepoints) ' , 2);\n']);

% Write the measurement data
for iMeasure = 1 : nMeasure                
    [~,X] = ode15s(@(t,x) f(exp(theta),x), t, con0(:,iMeasure));
    y = h(X(:,1:4), exp(theta));
    y = y + normrnd(0, sqrt(sigma2), nTimepoints, 2);
    for iTime = 1 : nTimepoints
        fprintf(fid, ['yMeasure(' num2str(iMeasure) ', ' num2str(iTime) ', :) = [']);
        fprintf(fid, num2str(y(iTime, :)));
        fprintf(fid, '];\n');
    end
end

% Close file
fprintf(fid, '\n end');
fclose(fid);

end
```
