## Supplementary material for "Mechanistic description of spatial processes using integrative modelling of noise-corrupted imaging data": erbb_signaling_pesto_syms.m

```
function model = erbb_signaling_pesto_syms()
% erbb_signaling_pesto_syms for examples/erbb_signaling
%
% creates an amimodel-object for the AMICI solver
%
% Parameters: 
% 
% Return values:
%  model: amimodel object


%% STATES
syms c2 c3 c288 c335 c117 c336 c286 c499 c500 c501 c4 c10 c11 c141 c145 c140 c146 c143 c147 c155 c159 c154 c160 c156 c161 c492 c493 c494 c502 c504 c503 c505 c506 c507 c105 c123 c124 c125 c126 c116 c122 c127 c128 c284 c129 c427 c130 c428 c131 c429 c132 c436 c133 c439 c134 c442 c135 c483 c136 c516 c137 c517 c138 c345 c139 c355 c168 c421 c169 c422 c170 c23 c12 c7 c25 c88 c27 c89 c29 c90 c34 c91 c35 c92 c36 c93 c37 c94 c189 c195 c190 c196 c191 c197 c198 c204 c199 c205 c200 c206 c207 c213 c208 c214 c209 c215 c216 c222 c217 c223 c218 c224 c225 c231 c226 c232 c227 c233 c243 c249 c244 c250 c245 c251 c252 c258 c253 c259 c254 c260 c234 c240 c235 c241 c236 c242 c300 c301 c303 c304 c306 c307 c309 c310 c312 c313 c315 c316 c318 c319 c321 c322 c357 c358 c360 c361 c366 c367 c369 c370 c372 c373 c375 c376 c378 c379 c381 c382 c363 c364 c384 c385 c387 c388 c390 c391 c393 c394 c396 c397 c399 c400 c402 c403 c9 c404 c401 c398 c395 c392 c389 c386 c365 c383 c380 c377 c374 c371 c368 c362 c359 c323 c320 c317 c314 c311 c308 c305 c302 c239 c238 c237 c257 c256 c255 c248 c247 c246 c230 c229 c228 c221 c220 c219 c212 c211 c210 c203 c202 c201 c194 c193 c192 c68 c67 c66 c65 c21 c20 c18 c19 c6 c5 c8 c15 c17 c32 c63 c33 c64 c347 c349 c348 c350 c351 c353 c508 c512 c354 c356 c148 c162 c149 c163 c150 c164 c289 c290 c337 c338 c291 c293 c294 c296 c297 c299 c14 c165 c166 c167 c151 c152 c153 c341 c344 c343 c346 c16 c157 c22 c40 c39 c180 c181 c182 c183 c184 c185 c24 c26 c28 c69 c43 c71 c31 c171 c172 c173 c174 c175 c176 c41 c42 c70 c72 c45 c38 c30 c44 c73 c46 c75 c76 c47 c74 c48 c49 c50 c51 c77 c53 c78 c52 c54 c79 c55 c56 c57 c58 c80 c81 c82 c59 c83 c60 c61 c84 c62 c85 c86 c425 c13 c518 c519 c339 c340 c95 c96 c97 c98 c101 c102 c99 c419 c103 c100 c420 c287 c486 c104 c447 c263 c445 c261 c446 c262 c454 c324 c457 c405 c460 c408 c106 c453 c452 c448 c449 c450 c451 c467 c468 c469 c470 c471 c107 c108 c112 c495 c109 c496 c110 c111 c497 c113 c498 c114 c280 c281 c282 c415 c283 c417 c418 c416 c87 c531 c330 c331 c332 c509 c510 c511 c513 c461 c444 c462 c279 c482 c426 c455 c463 c464 c465 c466 c473 c476 c479 c489 c431 c432 c433 c434 c435 c437 c438 c440 c474 c475 c477 c478 c480 c481 c491 c490 c410 c409 c430 c488 c487 c264 c265 c266 c267 c268 c269 c325 c472 c484 c485 c520 c521 c522 c523 c411 c412 c456 c424 c407 c142 c144 c158 c532 c525 c526 c527 c528 c524 c529 c530 c115 c121 c550 c551 c552 c553 c554 c555 c556 c557 c558
model.sym.x = [c2,c3,c288,c335,c117,c336,c286,c499,c500,c501,c4,c10,c11,c141,c145,c140,c146,c143,c147,c155,c159,c154,c160,c156,c161,c492,c493,c494,c502,c504,c503,c505,c506,c507,c105,c123,c124,c125,c126,c116,c122,c127,c128,c284,c129,c427,c130,c428,c131,c429,c132,c436,c133,c439,c134,c442,c135,c483,c136,c516,c137,c517,c138,c345,c139,c355,c168,c421,c169,c422,c170,c23,c12,c7,c25,c88,c27,c89,c29,c90,c34,c91,c35,c92,c36,c93,c37,c94,c189,c195,c190,c196,c191,c197,c198,c204,c199,c205,c200,c206,c207,c213,c208,c214,c209,c215,c216,c222,c217,c223,c218,c224,c225,c231,c226,c232,c227,c233,c243,c249,c244,c250,c245,c251,c252,c258,c253,c259,c254,c260,c234,c240,c235,c241,c236,c242,c300,c301,c303,c304,c306,c307,c309,c310,c312,c313,c315,c316,c318,c319,c321,c322,c357,c358,c360,c361,c366,c367,c369,c370,c372,c373,c375,c376,c378,c379,c381,c382,c363,c364,c384,c385,c387,c388,c390,c391,c393,c394,c396,c397,c399,c400,c402,c403,c9,c404,c401,c398,c395,c392,c389,c386,c365,c383,c380,c377,c374,c371,c368,c362,c359,c323,c320,c317,c314,c311,c308,c305,c302,c239,c238,c237,c257,c256,c255,c248,c247,c246,c230,c229,c228,c221,c220,c219,c212,c211,c210,c203,c202,c201,c194,c193,c192,c68,c67,c66,c65,c21,c20,c18,c19,c6,c5,c8,c15,c17,c32,c63,c33,c64,c347,c349,c348,c350,c351,c353,c508,c512,c354,c356,c148,c162,c149,c163,c150,c164,c289,c290,c337,c338,c291,c293,c294,c296,c297,c299,c14,c165,c166,c167,c151,c152,c153,c341,c344,c343,c346,c16,c157,c22,c40,c39,c180,c181,c182,c183,c184,c185,c24,c26,c28,c69,c43,c71,c31,c171,c172,c173,c174,c175,c176,c41,c42,c70,c72,c45,c38,c30,c44,c73,c46,c75,c76,c47,c74,c48,c49,c50,c51,c77,c53,c78,c52,c54,c79,c55,c56,c57,c58,c80,c81,c82,c59,c83,c60,c61,c84,c62,c85,c86,c425,c13,c518,c519,c339,c340,c95,c96,c97,c98,c101,c102,c99,c419,c103,c100,c420,c287,c486,c104,c447,c263,c445,c261,c446,c262,c454,c324,c457,c405,c460,c408,c106,c453,c452,c448,c449,c450,c451,c467,c468,c469,c470,c471,c107,c108,c112,c495,c109,c496,c110,c111,c497,c113,c498,c114,c280,c281,c282,c415,c283,c417,c418,c416,c87,c531,c330,c331,c332,c509,c510,c511,c513,c461,c444,c462,c279,c482,c426,c455,c463,c464,c465,c466,c473,c476,c479,c489,c431,c432,c433,c434,c435,c437,c438,c440,c474,c475,c477,c478,c480,c481,c491,c490,c410,c409,c430,c488,c487,c264,c265,c266,c267,c268,c269,c325,c472,c484,c485,c520,c521,c522,c523,c411,c412,c456,c424,c407,c142,c144,c158,c532,c525,c526,c527,c528,c524,c529,c530,c115,c121,c550,c551,c552,c553,c554,c555,c556,c557,c558];

%% PARAMETERS
model.param = 'log10';
syms EGFR_t ERK_t AKT_t k1d k2 kd2 k2b kd2b k3 kd3 k4 kd4 k4b kd4b k5 kd5 k5b kd5b kd5c k6 k8 kd8 kd8b k8b k10 k10b kd10 k13 kd13 k14 kd14 k15 kd15 k16 k16b k17 kd17 k18 kd18 k19 kd19 k20 kd20 k21 kd21 k22 kd22 k23 kd23 kd24 k25 kd25 k28 kd28 k28b kd28b k29 kd29 kd32 k32 kd33 k33 kd34 k34 kd35 k35 k36 kd36 kd37 k37 k40 kd40 k41 kd41 k42 kd42 kd43 k43 kd44 kd45 k45 kd47 k47 k48 kd48 kd49 k49 k50 kd50 kd52 kd53 k53 kd55 k55 kd56 k56 kd57 k57 k58 kd58 k52 k44 k60 kd60 k61 kd61 kd63 k64 kd64 kd65 k65 k66 kd66 k67 kd67 kd68 k68 kd68b k69 kd69 k70 kd70 k71 kd71 k72 kd72 k73 kd73 k74 kd74 kd75 k75 k76 kd76 kd60d k22b kd22b kd34b k34b k94b k94 kd94 k95 kd95 k96 kd96 kd6b k7 kd7 k62b kd60b k60c k60b KD_iressa_wt kd97 KD_iressa_mt kd97c Kinh2 kd98 k98 Kinh4 kd99 k99 Kinh3 kd100 k100 k101 kd101 k102 kd102 k103 kd103 k104 kd104 k105 kd105 k106 kd106 k106b kd106b k107 kd107 k108 kd108 k109 kd109 k110 kd110 kd111 k111 k112 kd112 k113 kd113 k114 kd114 k115 kd115 kd116 k117 kd117 k118 kd118 kd119 k120 kd120 k120b kd120b Ks k121 kd121 k123 kd123 k6b KD_ATP k123h kd123h KD kd_v853
model.sym.p = [EGFR_t,ERK_t,AKT_t,k1d,k2,kd2,k2b,kd2b,k3,kd3,k4,kd4,k4b,kd4b,k5,kd5,k5b,kd5b,kd5c,k6,k8,kd8,kd8b,k8b,k10,k10b,kd10,k13,kd13,k14,kd14,k15,kd15,k16,k16b,k17,kd17,k18,kd18,k19,kd19,k20,kd20,k21,kd21,k22,kd22,k23,kd23,kd24,k25,kd25,k28,kd28,k28b,kd28b,k29,kd29,kd32,k32,kd33,k33,kd34,k34,kd35,k35,k36,kd36,kd37,k37,k40,kd40,k41,kd41,k42,kd42,kd43,k43,kd44,kd45,k45,kd47,k47,k48,kd48,kd49,k49,k50,kd50,kd52,kd53,k53,kd55,k55,kd56,k56,kd57,k57,k58,kd58,k52,k44,k60,kd60,k61,kd61,kd63,k64,kd64,kd65,k65,k66,kd66,k67,kd67,kd68,k68,kd68b,k69,kd69,k70,kd70,k71,kd71,k72,kd72,k73,kd73,k74,kd74,kd75,k75,k76,kd76,kd60d,k22b,kd22b,kd34b,k34b,k94b,k94,kd94,k95,kd95,k96,kd96,kd6b,k7,kd7,k62b,kd60b,k60c,k60b,KD_iressa_wt,kd97,KD_iressa_mt,kd97c,Kinh2,kd98,k98,Kinh4,kd99,k99,Kinh3,kd100,k100,k101,kd101,k102,kd102,k103,kd103,k104,kd104,k105,kd105,k106,kd106,k106b,kd106b,k107,kd107,k108,kd108,k109,kd109,k110,kd110,kd111,k111,k112,kd112,k113,kd113,k114,kd114,k115,kd115,kd116,k117,kd117,k118,kd118,kd119,k120,kd120,k120b,kd120b,Ks,k121,kd121,k123,kd123,k6b,KD_ATP,k123h,kd123h,KD,kd_v853];

%% CONDITIONS
syms c1 c515 c285 c514
model.sym.k = [c1,c515,c285,c514];

%%
% DYNAMICS
time = sym('t');

model.sym.xdot = [c516*kd120 - c2*k6 + c517*kd120 - c2*stepfunc(time, 1799.99, 0, 1800, 1) + c6*stepfunc(time, 1799.99, 0, 1800, 0.00005) + c3*stepfunc(time, 1799.99, 0, 1800, 0.0033) + c105*c531*stepfunc(time, 1799.99, 0, 1800, 0.000000018704) - c2*c142*k120b - c2*c144*k120b - c1*c2*stepfunc(time, 1799.99, 0, 1800, 10000000), ...
c4*kd2 + c145*kd2b + c146*kd2b + c147*kd2b + c500*kd2 + c527*kd2 + c550*kd2 + c504*kd2b + c505*kd2b + c507*kd2b - c3^2*k2 - c3*stepfunc(time, 1799.99, 0, 1800, 0.0033) - c3*c499*k2 - c3*c526*k2 - c3*c529*k2 - c3*c140*k2b - c3*c141*k2b - c3*c143*k2b - c3*c502*k2b - c3*c503*k2b - c3*c506*k2b + c1*c2*stepfunc(time, 1799.99, 0, 1800, 10000000), ...
c335*stepfunc(time, 1799.99, 0, 1800, 0) - c288*kd103 + c140*c141*k103 - c1*c288*stepfunc(time, 1799.99, 0, 1800, 0), ...
c168*kd123 - c335*k7 + c337*kd7 + c341*kd8 - c335*kd103 - c335*stepfunc(time, 1799.99, 0, 1800, 0) - c14*c335*k8 + c87*c331*k103 - c105*c335*k123 + c1*c288*stepfunc(time, 1799.99, 0, 1800, 0), ...
c336*stepfunc(time, 1799.99, 0, 1800, 0) - c117*kd103 + c141*c143*k103 - c1*c117*k1d, ...
c139*kd123 - c336*k7 + c338*kd7 + c344*kd8 - c336*kd103 - c336*stepfunc(time, 1799.99, 0, 1800, 0) - c14*c336*k8 + c87*c332*k103 - c105*c336*k123 + c1*c117*k1d, ...
c499*stepfunc(time, 1799.99, 0, 1800, 0.0033) - c286*kd97 - c1*c286*stepfunc(time, 1799.99, 0, 1800, 10000000) + (c285*c531*kd97)/KD_iressa_wt, ...
c500*kd2 + c501*kd2 + c551*kd2 + c554*kd2 + c492*kd2b + c493*kd2b + c494*kd2b - c499^2*k2 - c499*stepfunc(time, 1799.99, 0, 1800, 0.0033) - c3*c499*k2 - c499*c526*k2 - c499*c529*k2 - c140*c499*k2b - c141*c499*k2b - c143*c499*k2b + c1*c286*stepfunc(time, 1799.99, 0, 1800, 10000000), ...
c115*stepfunc(time, 1799.99, 0, 1800, 1) - c500*kd2 - c105*c500*stepfunc(time, 1799.99, 0, 1800, 0.000000018704) + c3*c499*k2, ...
c499^2*k2 - c501*kd2, ...
c3^2*k2 - c4*kd2 + c116*stepfunc(time, 1799.99, 0, 1800, 1) - c4*c105*stepfunc(time, 1799.99, 0, 1800, 0.000000018704), ...
c11*kd2 - 2*c10*kd10 + c159*kd2b + c160*kd2b + c161*kd2b - c10^2*k2 + c6*c16*k10b + c16*c530*k10b - c10*c154*k2b - c10*c155*k2b - c10*c156*k2b, ...
c86*kd60 - c11*kd2 - c11*k60 + c415*kd95 + c10^2*k2 + c126*stepfunc(time, 1799.99, 0, 1800, 1) - c11*c105*stepfunc(time, 1799.99, 0, 1800, 0.000000018704) - c11*c280*k95, ...
c117*kd103 - c141*k6b + c145*kd2b + c155*kd6b + c284*kd103 + c288*kd103 + c345*kd120 + c355*kd120 + c492*kd2b + c502*kd98 + c513*kd103 - c87*c141*k103 - c140*c141*k103 - c141*c143*k103 - c141*c142*k120 - c141*c144*k120 - c141*c285*k98 - c141*c503*k103 - c3*c141*k2b - c141*c499*k2b, ...
c122*stepfunc(time, 1799.99, 0, 1800, 1) - c145*kd2b - c105*c145*stepfunc(time, 1799.99, 0, 1800, 0.000000018704) + c3*c141*k2b, ...
c146*kd2b - c140*k6b + c154*kd6b + c142*kd119 + c288*kd103 + c493*kd2b + c506*kd100 + c510*kd103 - c140*c141*k103 - c140*c285*k100 - c140*c502*k103 - c3*c140*k2b - c140*c499*k2b - c140*c514*stepfunc(time, 1799.99, 0, 1800, 10000000), ...
c127*stepfunc(time, 1799.99, 0, 1800, 1) - c146*kd2b - c105*c146*stepfunc(time, 1799.99, 0, 1800, 0.000000018704) + c3*c140*k2b, ...
c117*kd103 - c143*k6b + c147*kd2b + c156*kd6b + c144*kd119 + c494*kd2b + c503*kd99 + c511*kd103 - c141*c143*k103 - c143*c285*k99 - c143*c502*k103 - c3*c143*k2b - c143*c499*k2b - c143*c514*stepfunc(time, 1799.99, 0, 1800, 10000000), ...
c128*stepfunc(time, 1799.99, 0, 1800, 1) - c147*kd2b - c105*c147*stepfunc(time, 1799.99, 0, 1800, 0.000000018704) + c3*c143*k2b, ...
c141*k6b - c155*k60b + c86*kd60b - c155*kd6b + c159*kd2b + c421*kd120 + c422*kd120 - c155*c157*k120 - c155*c158*k120 - c10*c155*k2b, ...
c86*kd60b - c159*k62b - c159*kd2b + c416*kd95 + c123*stepfunc(time, 1799.99, 0, 1800, 1) - c105*c159*stepfunc(time, 1799.99, 0, 1800, 0.000000018704) - c159*c280*k95 + c10*c155*k2b, ...
c140*k6b - c154*k60b + c157*kd10 + c86*kd60b - c154*kd6b + c160*kd2b - c154*c515*k10b - c10*c154*k2b, ...
c86*kd60b - c160*k62b - c160*kd2b + c281*kd95 + c124*stepfunc(time, 1799.99, 0, 1800, 1) - c105*c160*stepfunc(time, 1799.99, 0, 1800, 0.000000018704) - c160*c280*k95 + c10*c154*k2b, ...
c143*k6b - c156*k60b + c86*kd60b - c156*kd6b + c161*kd2b - c10*c156*k2b, ...
c86*kd60b - c161*k62b - c161*kd2b + c282*kd95 + c125*stepfunc(time, 1799.99, 0, 1800, 1) - c105*c161*stepfunc(time, 1799.99, 0, 1800, 0.000000018704) - c161*c280*k95 + c10*c156*k2b, ...
c141*c499*k2b - c492*kd2b, ...
c140*c499*k2b - c493*kd2b, ...
c143*c499*k2b - c494*kd2b, ...
c504*kd2b - c502*kd98 - c502*k6b + c508*kd6b + c509*kd103 + c510*kd103 + c511*kd103 + c141*c285*k98 - c87*c502*k103 - c140*c502*k103 - c143*c502*k103 - c3*c502*k2b, ...
c3*c502*k2b - c504*kd2b, ...
c505*kd2b - c503*kd99 - c503*k6b + c512*kd6b + c513*kd103 + c143*c285*k99 - c141*c503*k103 - c3*c503*k2b, ...
c3*c503*k2b - c505*kd2b, ...
c507*kd2b - c506*kd100 + c140*c285*k100 - c3*c506*k2b, ...
c3*c506*k2b - c507*kd2b, ...
c116*kd123 + c122*kd123 + c123*kd123 + c124*kd123 + c125*kd123 + c126*kd123 + c127*kd123 + c128*kd123 + c129*kd123 + c130*kd123 + c131*kd123 + c132*kd123 + c133*kd123 + c134*kd123 + c135*kd123 + c136*kd123 + c137*kd123 + c138*kd123 + c139*kd123 + c168*kd123 + c169*kd123 + c170*kd123 + c115*kd123h + c121*kd123h + c555*kd123 + c557*kd123 + c556*kd123h + c558*kd123h + c2*stepfunc(time, 1799.99, 0, 1800, 1) + c115*stepfunc(time, 1799.99, 0, 1800, 1) + c116*stepfunc(time, 1799.99, 0, 1800, 1) + c121*stepfunc(time, 1799.99, 0, 1800, 1) + c122*stepfunc(time, 1799.99, 0, 1800, 1) + c123*stepfunc(time, 1799.99, 0, 1800, 1) + c124*stepfunc(time, 1799.99, 0, 1800, 1) + c125*stepfunc(time, 1799.99, 0, 1800, 1) + c126*stepfunc(time, 1799.99, 0, 1800, 1) + c127*stepfunc(time, 1799.99, 0, 1800, 1) + c128*stepfunc(time, 1799.99, 0, 1800, 1) + c129*stepfunc(time, 1799.99, 0, 1800, 1) + c130*stepfunc(time, 1799.99, 0, 1800, 1) + c131*stepfunc(time, 1799.99, 0, 1800, 1) + c132*stepfunc(time, 1799.99, 0, 1800, 1) + c133*stepfunc(time, 1799.99, 0, 1800, 1) + c134*stepfunc(time, 1799.99, 0, 1800, 1) + c135*stepfunc(time, 1799.99, 0, 1800, 1) + c136*stepfunc(time, 1799.99, 0, 1800, 1) + c137*stepfunc(time, 1799.99, 0, 1800, 1) + c138*stepfunc(time, 1799.99, 0, 1800, 1) + c139*stepfunc(time, 1799.99, 0, 1800, 1) + c168*stepfunc(time, 1799.99, 0, 1800, 1) + c169*stepfunc(time, 1799.99, 0, 1800, 1) + c170*stepfunc(time, 1799.99, 0, 1800, 1) + c524*stepfunc(time, 1799.99, 0, 1800, 1) + c555*stepfunc(time, 1799.99, 0, 1800, 1) + c556*stepfunc(time, 1799.99, 0, 1800, 1) + c557*stepfunc(time, 1799.99, 0, 1800, 1) + c558*stepfunc(time, 1799.99, 0, 1800, 1) - c4*c105*stepfunc(time, 1799.99, 0, 1800, 0.000000018704) - c11*c105*stepfunc(time, 1799.99, 0, 1800, 0.000000018704) - c105*c145*stepfunc(time, 1799.99, 0, 1800, 0.000000018704) - c105*c146*stepfunc(time, 1799.99, 0, 1800, 0.000000018704) - c105*c147*stepfunc(time, 1799.99, 0, 1800, 0.000000018704) - c105*c159*stepfunc(time, 1799.99, 0, 1800, 0.000000018704) - c105*c160*stepfunc(time, 1799.99, 0, 1800, 0.000000018704) - c105*c161*stepfunc(time, 1799.99, 0, 1800, 0.000000018704) - c105*c284*stepfunc(time, 1799.99, 0, 1800, 0.000000018704) - c105*c345*stepfunc(time, 1799.99, 0, 1800, 0.000000018704) - c105*c355*stepfunc(time, 1799.99, 0, 1800, 0.000000018704) - c105*c421*stepfunc(time, 1799.99, 0, 1800, 0.000000018704) - c105*c422*stepfunc(time, 1799.99, 0, 1800, 0.000000018704) - c105*c427*stepfunc(time, 1799.99, 0, 1800, 0.000000018704) - c105*c428*stepfunc(time, 1799.99, 0, 1800, 0.000000018704) - c105*c429*stepfunc(time, 1799.99, 0, 1800, 0.000000018704) - c105*c436*stepfunc(time, 1799.99, 0, 1800, 0.000000018704) - c105*c439*stepfunc(time, 1799.99, 0, 1800, 0.000000018704) - c105*c442*stepfunc(time, 1799.99, 0, 1800, 0.000000018704) - c105*c483*stepfunc(time, 1799.99, 0, 1800, 0.000000018704) - c105*c500*stepfunc(time, 1799.99, 0, 1800, 0.000000018704) - c105*c516*stepfunc(time, 1799.99, 0, 1800, 0.000000018704) - c105*c517*stepfunc(time, 1799.99, 0, 1800, 0.000000018704) - c105*c527*stepfunc(time, 1799.99, 0, 1800, 0.000000018704) - c105*c531*stepfunc(time, 1799.99, 0, 1800, 0.000000018704) - c105*c532*stepfunc(time, 1799.99, 0, 1800, 0.000000018704) - c105*c550*stepfunc(time, 1799.99, 0, 1800, 0.000000018704) - c105*c551*stepfunc(time, 1799.99, 0, 1800, 0.000000018704) - c105*c552*stepfunc(time, 1799.99, 0, 1800, 0.000000018704) - c105*c553*stepfunc(time, 1799.99, 0, 1800, 0.000000018704) - 3*c5*c105*k123 - c8*c105*k123 - 4*c5*c105*k123h - c105*c148*k123 - 2*c105*c149*k123 - 2*c105*c150*k123 - c105*c162*k123 - c105*c163*k123 - c105*c164*k123 - c105*c289*k123 - c105*c335*k123 - c105*c336*k123 - c105*c337*k123 - c105*c338*k123 - c105*c445*k123 - c105*c446*k123 - c105*c447*k123 - c105*c454*k123 - c105*c457*k123 - c105*c460*k123 - c105*c486*k123, ...
c105*c159*stepfunc(time, 1799.99, 0, 1800, 0.000000018704) - c123*stepfunc(time, 1799.99, 0, 1800, 1) - c123*kd123 + c105*c162*k123, ...
c105*c160*stepfunc(time, 1799.99, 0, 1800, 0.000000018704) - c124*stepfunc(time, 1799.99, 0, 1800, 1) - c124*kd123 + c105*c163*k123, ...
c105*c161*stepfunc(time, 1799.99, 0, 1800, 0.000000018704) - c125*stepfunc(time, 1799.99, 0, 1800, 1) - c125*kd123 + c105*c164*k123, ...
c11*c105*stepfunc(time, 1799.99, 0, 1800, 0.000000018704) - c126*stepfunc(time, 1799.99, 0, 1800, 1) - c126*kd123 + c8*c105*k123, ...
c4*c105*stepfunc(time, 1799.99, 0, 1800, 0.000000018704) - c116*stepfunc(time, 1799.99, 0, 1800, 1) - c116*kd123 + c5*c105*k123, ...
c105*c145*stepfunc(time, 1799.99, 0, 1800, 0.000000018704) - c122*stepfunc(time, 1799.99, 0, 1800, 1) - c122*kd123 + c105*c148*k123, ...
c105*c146*stepfunc(time, 1799.99, 0, 1800, 0.000000018704) - c127*stepfunc(time, 1799.99, 0, 1800, 1) - c127*kd123 + c105*c149*k123, ...
c105*c147*stepfunc(time, 1799.99, 0, 1800, 0.000000018704) - c128*stepfunc(time, 1799.99, 0, 1800, 1) - c128*kd123 + c105*c150*k123, ...
c129*stepfunc(time, 1799.99, 0, 1800, 1) - c284*kd103 - c105*c284*stepfunc(time, 1799.99, 0, 1800, 0.000000018704) + c87*c141*k103, ...
c105*c284*stepfunc(time, 1799.99, 0, 1800, 0.000000018704) - c129*stepfunc(time, 1799.99, 0, 1800, 1) - c129*kd123 + c105*c289*k123, ...
c464*kd108 - c427*kd105 + c130*stepfunc(time, 1799.99, 0, 1800, 1) - c105*c427*stepfunc(time, 1799.99, 0, 1800, 0.000000018704) + c225*c426*k105 - c427*c463*k108, ...
c105*c427*stepfunc(time, 1799.99, 0, 1800, 0.000000018704) - c130*stepfunc(time, 1799.99, 0, 1800, 1) - c130*kd123 + c105*c445*k123, ...
c465*kd108 - c428*kd105 + c131*stepfunc(time, 1799.99, 0, 1800, 1) - c105*c428*stepfunc(time, 1799.99, 0, 1800, 0.000000018704) + c226*c426*k105 - c428*c463*k108, ...
c105*c428*stepfunc(time, 1799.99, 0, 1800, 0.000000018704) - c131*stepfunc(time, 1799.99, 0, 1800, 1) - c131*kd123 + c105*c446*k123, ...
c466*kd108 - c429*kd105 + c132*stepfunc(time, 1799.99, 0, 1800, 1) - c105*c429*stepfunc(time, 1799.99, 0, 1800, 0.000000018704) + c227*c426*k105 - c429*c463*k108, ...
c105*c429*stepfunc(time, 1799.99, 0, 1800, 0.000000018704) - c132*stepfunc(time, 1799.99, 0, 1800, 1) - c132*kd123 + c105*c447*k123, ...
c473*kd108 - c436*kd105 + c133*stepfunc(time, 1799.99, 0, 1800, 1) - c105*c436*stepfunc(time, 1799.99, 0, 1800, 0.000000018704) + c312*c426*k105 - c436*c463*k108, ...
c105*c436*stepfunc(time, 1799.99, 0, 1800, 0.000000018704) - c133*stepfunc(time, 1799.99, 0, 1800, 1) - c133*kd123 + c105*c454*k123, ...
c476*kd108 - c439*kd105 + c134*stepfunc(time, 1799.99, 0, 1800, 1) - c105*c439*stepfunc(time, 1799.99, 0, 1800, 0.000000018704) + c381*c426*k105 - c439*c463*k108, ...
c105*c439*stepfunc(time, 1799.99, 0, 1800, 0.000000018704) - c134*stepfunc(time, 1799.99, 0, 1800, 1) - c134*kd123 + c105*c457*k123, ...
c479*kd108 - c442*kd105 + c135*stepfunc(time, 1799.99, 0, 1800, 1) - c105*c442*stepfunc(time, 1799.99, 0, 1800, 0.000000018704) + c384*c426*k105 - c442*c463*k108, ...
c105*c442*stepfunc(time, 1799.99, 0, 1800, 0.000000018704) - c135*stepfunc(time, 1799.99, 0, 1800, 1) - c135*kd123 + c105*c460*k123, ...
c489*kd108 - c483*kd105 + c136*stepfunc(time, 1799.99, 0, 1800, 1) - c105*c483*stepfunc(time, 1799.99, 0, 1800, 0.000000018704) + c23*c426*k105 - c463*c483*k108, ...
c105*c483*stepfunc(time, 1799.99, 0, 1800, 0.000000018704) - c136*stepfunc(time, 1799.99, 0, 1800, 1) - c136*kd123 + c105*c486*k123, ...
c137*stepfunc(time, 1799.99, 0, 1800, 1) - c516*kd120 - c105*c516*stepfunc(time, 1799.99, 0, 1800, 0.000000018704) + c2*c142*k120b, ...
c105*c516*stepfunc(time, 1799.99, 0, 1800, 0.000000018704) - c137*stepfunc(time, 1799.99, 0, 1800, 1) - c137*kd123 + c105*c149*k123, ...
c138*stepfunc(time, 1799.99, 0, 1800, 1) - c517*kd120 - c105*c517*stepfunc(time, 1799.99, 0, 1800, 0.000000018704) + c2*c144*k120b, ...
c105*c517*stepfunc(time, 1799.99, 0, 1800, 0.000000018704) - c138*stepfunc(time, 1799.99, 0, 1800, 1) - c138*kd123 + c105*c150*k123, ...
c139*stepfunc(time, 1799.99, 0, 1800, 1) - c345*kd120 - c105*c345*stepfunc(time, 1799.99, 0, 1800, 0.000000018704) + c141*c144*k120, ...
c105*c345*stepfunc(time, 1799.99, 0, 1800, 0.000000018704) - c139*stepfunc(time, 1799.99, 0, 1800, 1) - c139*kd123 + c105*c336*k123, ...
c168*stepfunc(time, 1799.99, 0, 1800, 1) - c355*kd120 - c105*c355*stepfunc(time, 1799.99, 0, 1800, 0.000000018704) + c141*c142*k120, ...
c105*c355*stepfunc(time, 1799.99, 0, 1800, 0.000000018704) - c168*stepfunc(time, 1799.99, 0, 1800, 1) - c168*kd123 + c105*c335*k123, ...
c86*kd60b - c421*k62b - c421*kd120 + c169*stepfunc(time, 1799.99, 0, 1800, 1) - c105*c421*stepfunc(time, 1799.99, 0, 1800, 0.000000018704) + c155*c157*k120, ...
c105*c421*stepfunc(time, 1799.99, 0, 1800, 0.000000018704) - c169*stepfunc(time, 1799.99, 0, 1800, 1) - c169*kd123 + c105*c337*k123, ...
c86*kd60b - c422*k62b - c422*kd120 + c170*stepfunc(time, 1799.99, 0, 1800, 1) - c105*c422*stepfunc(time, 1799.99, 0, 1800, 0.000000018704) + c155*c158*k120, ...
c105*c422*stepfunc(time, 1799.99, 0, 1800, 0.000000018704) - c170*stepfunc(time, 1799.99, 0, 1800, 1) - c170*kd123 + c105*c338*k123, ...
c7*kd4 - c23*k6 + c25*kd17 - c23*kd63 + c99*kd101 + c483*kd105 + c18*stepfunc(time, 1799.99, 0, 1800, 0.00005) - c12*c23*k4 + c15*c22*k16 - c23*c24*k17 - c23*c103*k101 - c23*c426*k105, ...
c9*k15 + c7*kd4 - c12*kd15 + c88*kd4 + c89*kd4 + c90*kd4 + c91*kd4 + c92*kd4 + c93*kd4 + c94*kd4 + c195*kd4 + c196*kd4 + c197*kd4 + c204*kd4 + c205*kd4 + c206*kd4 + c213*kd4 + c214*kd4 + c215*kd4 + c222*kd4 + c223*kd4 + c224*kd4 + c231*kd4 + c232*kd4 + c233*kd4 + c240*kd4 + c241*kd4 + c242*kd4 + c249*kd4 + c250*kd4 + c251*kd4 + c258*kd4 + c259*kd4 + c260*kd4 + c301*kd4 + c304*kd4 + c307*kd4 + c310*kd4 + c313*kd4 + c316*kd4 + c319*kd4 + c322*kd4 + c358*kd4 + c361*kd4 + c364*kd4 + c367*kd4 + c370*kd4 + c373*kd4 + c376*kd4 + c379*kd4 + c382*kd4 + c385*kd4 + c388*kd4 + c391*kd4 + c394*kd4 + c397*kd4 + c400*kd4 + c403*kd4 - c12*c23*k4 - c12*c25*k4 - c12*c27*k4 - c12*c29*k4 - c12*c34*k4 - c12*c35*k4 - c12*c36*k4 - c12*c37*k4 - c12*c252*k4 - c12*c253*k4 - c12*c254*k4 - c12*c189*k4b - c12*c190*k4b - c12*c191*k4b - c12*c198*k4b - c12*c199*k4b - c12*c200*k4b - c12*c207*k4b - c12*c208*k4b - c12*c209*k4b - c12*c216*k4b - c12*c217*k4b - c12*c218*k4b - c12*c225*k4b - c12*c226*k4b - c12*c227*k4b - c12*c234*k4b - c12*c235*k4b - c12*c236*k4b - c12*c243*k4b - c12*c244*k4b - c12*c245*k4b - c12*c300*k4b - c12*c303*k4b - c12*c306*k4b - c12*c309*k4b - c12*c312*k4b - c12*c315*k4b - c12*c318*k4b - c12*c321*k4b - c12*c357*k4b - c12*c360*k4b - c12*c363*k4b - c12*c366*k4b - c12*c369*k4b - c12*c372*k4b - c12*c375*k4b - c12*c378*k4b - c12*c381*k4b - c12*c384*k4b - c12*c387*k4b - c12*c390*k4b - c12*c393*k4b - c12*c396*k4b - c12*c399*k4b - c12*c402*k4b, ...
c9*c18*k5 - c7*kd5 - c7*kd4 + c12*c23*k4, ...
c27*kd18 - c25*kd17 - c25*k6 + c27*kd19 + c29*kd20 + c29*kd21 - c25*kd34 + c88*kd4 + c95*kd64 + c19*stepfunc(time, 1799.99, 0, 1800, 0.00005) - c12*c25*k4 + c23*c24*k17 - c25*c26*k18 - c25*c26*k21 - c25*c28*k19 + c15*c30*k34 - c25*c43*k20 - c25*c59*k64, ...
c9*c19*k5 - c88*kd5 - c88*kd4 + c12*c25*k4, ...
c89*kd4 - c27*kd18 - c27*kd19 - c27*k6 + c20*stepfunc(time, 1799.99, 0, 1800, 0.00005) - c12*c27*k4 + c25*c26*k18 + c25*c28*k19, ...
c9*c20*k5 - c89*kd5 - c89*kd4 + c12*c27*k4, ...
c90*kd4 - c29*kd20 - c29*kd21 - c29*k6 + c21*stepfunc(time, 1799.99, 0, 1800, 0.00005) - c12*c29*k4 + c25*c26*k21 + c25*c43*k20, ...
c9*c21*k5 - c90*kd5 - c90*kd4 + c12*c29*k4, ...
c35*kd25 - c34*kd24 - c34*k6 - c34*kd37 + c91*kd4 + c419*kd101 + c65*stepfunc(time, 1799.99, 0, 1800, 0.00005) - c12*c34*k4 + c22*c33*k16 - c24*c34*k25 + c15*c39*k37 - c34*c103*k101, ...
c12*c34*k4 - c91*kd5 - c91*kd4 + c9*c65*k5, ...
c36*kd18 - c35*k6 + c36*kd19 + c37*kd20 + c37*kd21 - c35*kd25 - c35*kd32 - c35*kd41 + c92*kd4 + c97*kd64 + c66*stepfunc(time, 1799.99, 0, 1800, 0.00005) - c12*c35*k4 - c26*c35*k18 - c26*c35*k21 - c28*c35*k19 + c24*c34*k25 + c15*c38*k32 - c35*c43*k20 + c30*c33*k41 - c35*c59*k64, ...
c12*c35*k4 - c92*kd5 - c92*kd4 + c9*c66*k5, ...
c93*kd4 - c36*kd18 - c36*kd19 - c36*k6 + c67*stepfunc(time, 1799.99, 0, 1800, 0.00005) - c12*c36*k4 + c26*c35*k18 + c28*c35*k19, ...
c12*c36*k4 - c93*kd5 - c93*kd4 + c9*c67*k5, ...
c94*kd4 - c37*kd20 - c37*kd21 - c37*k6 + c68*stepfunc(time, 1799.99, 0, 1800, 0.00005) - c12*c37*k4 + c26*c35*k21 + c35*c43*k20, ...
c12*c37*k4 - c94*kd5 - c94*kd4 + c9*c68*k5, ...
c195*kd4 - c189*kd24 + c198*kd25 - c189*kd37 + c22*c180*k16 + c39*c151*k37 - c24*c189*k25 - c12*c189*k4b, ...
c9*c192*k5 - c195*kd5b - c195*kd4 + c12*c189*k4b, ...
c196*kd4 - c190*kd24 + c199*kd25 - c190*kd37 + c22*c181*k16 + c39*c152*k37 - c24*c190*k25 - c12*c190*k4b, ...
c9*c193*k5 - c196*kd5b - c196*kd4 + c12*c190*k4b, ...
c197*kd4 - c191*kd24 + c200*kd25 - c191*kd37 + c22*c182*k16 + c39*c153*k37 - c24*c191*k25 - c12*c191*k4b, ...
c9*c194*k5 - c197*kd5b - c197*kd4 + c12*c191*k4b, ...
c204*kd4 - c198*kd25 + c207*kd18 + c207*kd19 - c198*kd32 + c216*kd20 + c216*kd21 - c198*kd41 + c38*c151*k32 + c24*c189*k25 - c26*c198*k18 - c26*c198*k21 - c28*c198*k19 + c30*c180*k41 - c43*c198*k20 - c12*c198*k4b, ...
c9*c201*k5 - c204*kd5b - c204*kd4 + c12*c198*k4b, ...
c205*kd4 - c199*kd25 + c208*kd18 + c208*kd19 - c199*kd32 + c217*kd20 + c217*kd21 - c199*kd41 + c38*c152*k32 + c24*c190*k25 - c26*c199*k18 - c26*c199*k21 - c28*c199*k19 + c30*c181*k41 - c43*c199*k20 - c12*c199*k4b, ...
c9*c202*k5 - c205*kd5b - c205*kd4 + c12*c199*k4b, ...
c206*kd4 - c200*kd25 + c209*kd18 + c209*kd19 - c200*kd32 + c218*kd20 + c218*kd21 - c200*kd41 + c38*c153*k32 + c24*c191*k25 - c26*c200*k18 - c26*c200*k21 - c28*c200*k19 + c30*c182*k41 - c43*c200*k20 - c12*c200*k4b, ...
c9*c203*k5 - c206*kd5b - c206*kd4 + c12*c200*k4b, ...
c213*kd4 - c207*kd18 - c207*kd19 + c26*c198*k18 + c28*c198*k19 - c12*c207*k4b, ...
c9*c210*k5 - c213*kd5b - c213*kd4 + c12*c207*k4b, ...
c214*kd4 - c208*kd18 - c208*kd19 + c26*c199*k18 + c28*c199*k19 - c12*c208*k4b, ...
c9*c211*k5 - c214*kd5b - c214*kd4 + c12*c208*k4b, ...
c215*kd4 - c209*kd18 - c209*kd19 + c26*c200*k18 + c28*c200*k19 - c12*c209*k4b, ...
c9*c212*k5 - c215*kd5b - c215*kd4 + c12*c209*k4b, ...
c222*kd4 - c216*kd20 - c216*kd21 + c26*c198*k21 + c43*c198*k20 - c12*c216*k4b, ...
c9*c219*k5 - c222*kd5b - c222*kd4 + c12*c216*k4b, ...
c223*kd4 - c217*kd20 - c217*kd21 + c26*c199*k21 + c43*c199*k20 - c12*c217*k4b, ...
c9*c220*k5 - c223*kd5b - c223*kd4 + c12*c217*k4b, ...
c224*kd4 - c218*kd20 - c218*kd21 + c26*c200*k21 + c43*c200*k20 - c12*c218*k4b, ...
c9*c221*k5 - c224*kd5b - c224*kd4 + c12*c218*k4b, ...
c231*kd4 - c225*kd24 + c234*kd17 + c427*kd105 + c22*c151*k16 - c24*c225*k17 - c225*c426*k105 - c12*c225*k4b, ...
c9*c228*k5 - c231*kd5b - c231*kd4 + c12*c225*k4b, ...
c232*kd4 - c226*kd24 + c235*kd17 + c428*kd105 + c22*c152*k16 - c24*c226*k17 - c226*c426*k105 - c12*c226*k4b, ...
c9*c229*k5 - c232*kd5b - c232*kd4 + c12*c226*k4b, ...
c233*kd4 - c227*kd24 + c236*kd17 + c429*kd105 + c22*c153*k16 - c24*c227*k17 - c227*c426*k105 - c12*c227*k4b, ...
c9*c230*k5 - c233*kd5b - c233*kd4 + c12*c227*k4b, ...
c249*kd4 - c243*kd18 - c243*kd19 + c26*c234*k18 + c28*c234*k19 - c12*c243*k4b, ...
c9*c246*k5 - c249*kd5b - c249*kd4 + c12*c243*k4b, ...
c250*kd4 - c244*kd18 - c244*kd19 + c26*c235*k18 + c28*c235*k19 - c12*c244*k4b, ...
c9*c247*k5 - c250*kd5b - c250*kd4 + c12*c244*k4b, ...
c251*kd4 - c245*kd18 - c245*kd19 + c26*c236*k18 + c28*c236*k19 - c12*c245*k4b, ...
c9*c248*k5 - c251*kd5b - c251*kd4 + c12*c245*k4b, ...
c258*kd4 - c252*kd20 - c252*kd21 - c12*c252*k4 + c26*c234*k21 + c43*c234*k20, ...
c12*c252*k4 - c258*kd5b - c258*kd4 + c9*c255*k5, ...
c259*kd4 - c253*kd20 - c253*kd21 - c12*c253*k4 + c26*c235*k21 + c43*c235*k20, ...
c12*c253*k4 - c259*kd5b - c259*kd4 + c9*c256*k5, ...
c260*kd4 - c254*kd20 - c254*kd21 - c12*c254*k4 + c26*c236*k21 + c43*c236*k20, ...
c12*c254*k4 - c260*kd5b - c260*kd4 + c9*c257*k5, ...
c240*kd4 - c234*kd17 + c243*kd18 + c243*kd19 - c234*kd34 + c252*kd20 + c252*kd21 + c30*c151*k34 + c24*c225*k17 - c26*c234*k18 - c26*c234*k21 - c28*c234*k19 - c43*c234*k20 - c12*c234*k4b, ...
c9*c237*k5 - c240*kd5b - c240*kd4 + c12*c234*k4b, ...
c241*kd4 - c235*kd17 + c244*kd18 + c244*kd19 - c235*kd34 + c253*kd20 + c253*kd21 + c30*c152*k34 + c24*c226*k17 - c26*c235*k18 - c26*c235*k21 - c28*c235*k19 - c43*c235*k20 - c12*c235*k4b, ...
c9*c238*k5 - c241*kd5b - c241*kd4 + c12*c235*k4b, ...
c242*kd4 - c236*kd17 + c245*kd18 + c245*kd19 - c236*kd34 + c254*kd20 + c254*kd21 + c30*c153*k34 + c24*c227*k17 - c26*c236*k18 - c26*c236*k21 - c28*c236*k19 - c43*c236*k20 - c12*c236*k4b, ...
c9*c239*k5 - c242*kd5b - c242*kd4 + c12*c236*k4b, ...
c301*kd4 - c300*kd24 + c303*kd25 - c300*kd37 + c22*c297*k16 - c24*c300*k25 + c39*c291*k37 - c12*c300*k4b, ...
c9*c302*k5 - c301*kd5b - c301*kd4 + c12*c300*k4b, ...
c304*kd4 + c306*kd18 + c306*kd19 - c303*kd25 + c309*kd20 + c309*kd21 - c303*kd32 - c303*kd41 - c26*c303*k18 + c24*c300*k25 - c26*c303*k21 - c28*c303*k19 + c38*c291*k32 - c43*c303*k20 + c30*c297*k41 - c12*c303*k4b, ...
c9*c305*k5 - c304*kd5b - c304*kd4 + c12*c303*k4b, ...
c307*kd4 - c306*kd18 - c306*kd19 + c26*c303*k18 + c28*c303*k19 - c12*c306*k4b, ...
c9*c308*k5 - c307*kd5b - c307*kd4 + c12*c306*k4b, ...
c310*kd4 - c309*kd20 - c309*kd21 + c26*c303*k21 + c43*c303*k20 - c12*c309*k4b, ...
c9*c311*k5 - c310*kd5b - c310*kd4 + c12*c309*k4b, ...
c313*kd4 + c315*kd17 - c312*kd63 + c436*kd105 + c22*c291*k16 - c24*c312*k17 - c312*c426*k105 - c12*c312*k4b, ...
c9*c314*k5 - c313*kd5b - c313*kd4 + c12*c312*k4b, ...
c316*kd4 - c315*kd17 + c318*kd18 + c318*kd19 + c321*kd20 + c321*kd21 - c315*kd34 + c24*c312*k17 + c30*c291*k34 - c26*c315*k18 - c26*c315*k21 - c28*c315*k19 - c43*c315*k20 - c12*c315*k4b, ...
c9*c317*k5 - c316*kd5b - c316*kd4 + c12*c315*k4b, ...
c319*kd4 - c318*kd18 - c318*kd19 + c26*c315*k18 + c28*c315*k19 - c12*c318*k4b, ...
c9*c320*k5 - c319*kd5b - c319*kd4 + c12*c318*k4b, ...
c322*kd4 - c321*kd20 - c321*kd21 + c26*c315*k21 + c43*c315*k20 - c12*c321*k4b, ...
c9*c323*k5 - c322*kd5b - c322*kd4 + c12*c321*k4b, ...
c358*kd4 - c357*kd24 + c363*kd25 - c357*kd37 + c22*c351*k16 - c24*c357*k25 + c39*c341*k37 - c12*c357*k4b, ...
c9*c359*k5b - c358*kd5b - c358*kd4 + c12*c357*k4b, ...
c361*kd4 - c360*kd24 + c366*kd25 - c360*kd37 + c22*c354*k16 - c24*c360*k25 + c39*c344*k37 - c12*c360*k4b, ...
c9*c362*k5b - c361*kd5b - c361*kd4 + c12*c360*k4b, ...
c367*kd4 + c372*kd18 - c366*kd25 + c372*kd19 - c366*kd32 + c378*kd20 + c378*kd21 - c366*kd41 + c24*c360*k25 - c26*c366*k18 - c26*c366*k21 - c28*c366*k19 + c38*c344*k32 + c30*c354*k41 - c43*c366*k20 - c12*c366*k4b, ...
c9*c368*k5b - c367*kd5b - c367*kd4 + c12*c366*k4b, ...
c370*kd4 - c369*kd18 - c369*kd19 + c26*c363*k18 + c28*c363*k19 - c12*c369*k4b, ...
c9*c371*k5b - c370*kd5b - c370*kd4 + c12*c369*k4b, ...
c373*kd4 - c372*kd18 - c372*kd19 + c26*c366*k18 + c28*c366*k19 - c12*c372*k4b, ...
c9*c374*k5b - c373*kd5b - c373*kd4 + c12*c372*k4b, ...
c376*kd4 - c375*kd20 - c375*kd21 + c26*c363*k21 + c43*c363*k20 - c12*c375*k4b, ...
c9*c377*k5b - c376*kd5b - c376*kd4 + c12*c375*k4b, ...
c379*kd4 - c378*kd20 - c378*kd21 + c26*c366*k21 + c43*c366*k20 - c12*c378*k4b, ...
c9*c380*k5b - c379*kd5b - c379*kd4 + c12*c378*k4b, ...
c382*kd4 + c387*kd17 - c381*kd63 + c439*kd105 + c22*c341*k16 - c24*c381*k17 - c381*c426*k105 - c12*c381*k4b, ...
c9*c383*k5b - c382*kd5b - c382*kd4 + c12*c381*k4b, ...
c364*kd4 + c369*kd18 - c363*kd25 + c369*kd19 - c363*kd32 + c375*kd20 + c375*kd21 - c363*kd41 + c24*c357*k25 - c26*c363*k18 - c26*c363*k21 - c28*c363*k19 + c38*c341*k32 + c30*c351*k41 - c43*c363*k20 - c12*c363*k4b, ...
c9*c365*k5b - c364*kd5b - c364*kd4 + c12*c363*k4b, ...
c385*kd4 + c390*kd17 - c384*kd24 + c442*kd105 + c22*c344*k16 - c24*c384*k17 - c384*c426*k105 - c12*c384*k4b, ...
c9*c386*k5b - c385*kd5b - c385*kd4 + c12*c384*k4b, ...
c388*kd4 - c387*kd17 + c393*kd18 + c393*kd19 + c399*kd20 + c399*kd21 - c387*kd34 + c30*c341*k34 + c24*c381*k17 - c26*c387*k18 - c26*c387*k21 - c28*c387*k19 - c43*c387*k20 - c12*c387*k4b, ...
c9*c389*k5b - c388*kd5b - c388*kd4 + c12*c387*k4b, ...
c391*kd4 - c390*kd17 + c396*kd18 + c396*kd19 + c402*kd20 + c402*kd21 - c390*kd34 + c30*c344*k34 + c24*c384*k17 - c26*c390*k18 - c26*c390*k21 - c28*c390*k19 - c43*c390*k20 - c12*c390*k4b, ...
c9*c392*k5b - c391*kd5b - c391*kd4 + c12*c390*k4b, ...
c394*kd4 - c393*kd18 - c393*kd19 + c26*c387*k18 + c28*c387*k19 - c12*c393*k4b, ...
c9*c395*k5b - c394*kd5b - c394*kd4 + c12*c393*k4b, ...
c397*kd4 - c396*kd18 - c396*kd19 + c26*c390*k18 + c28*c390*k19 - c12*c396*k4b, ...
c9*c398*k5b - c397*kd5b - c397*kd4 + c12*c396*k4b, ...
c400*kd4 - c399*kd20 - c399*kd21 + c26*c387*k21 + c43*c387*k20 - c12*c399*k4b, ...
c9*c401*k5b - c400*kd5b - c400*kd4 + c12*c399*k4b, ...
c403*kd4 - c402*kd20 - c402*kd21 + c26*c390*k21 + c43*c390*k20 - c12*c402*k4b, ...
c9*c404*k5b - c403*kd5b - c403*kd4 + c12*c402*k4b, ...
c7*kd5 - c9*k15 + c12*kd15 + c88*kd5 + c89*kd5 + c90*kd5 + c91*kd5 + c92*kd5 + c93*kd5 + c94*kd5 + c195*kd5b + c196*kd5b + c197*kd5b + c204*kd5b + c205*kd5b + c206*kd5b + c213*kd5b + c214*kd5b + c215*kd5b + c222*kd5b + c223*kd5b + c224*kd5b + c231*kd5b + c232*kd5b + c233*kd5b + c240*kd5b + c241*kd5b + c242*kd5b + c249*kd5b + c250*kd5b + c251*kd5b + c258*kd5b + c259*kd5b + c260*kd5b + c301*kd5b + c304*kd5b + c307*kd5b + c310*kd5b + c313*kd5b + c316*kd5b + c319*kd5b + c322*kd5b + c358*kd5b + c361*kd5b + c364*kd5b + c367*kd5b + c370*kd5b + c373*kd5b + c376*kd5b + c379*kd5b + c382*kd5b + c385*kd5b + c388*kd5b + c391*kd5b + c394*kd5b + c397*kd5b + c400*kd5b + c403*kd5b - c9*c18*k5 - c9*c19*k5 - c9*c20*k5 - c9*c21*k5 - c9*c65*k5 - c9*c66*k5 - c9*c67*k5 - c9*c68*k5 - c9*c192*k5 - c9*c193*k5 - c9*c194*k5 - c9*c201*k5 - c9*c202*k5 - c9*c203*k5 - c9*c210*k5 - c9*c211*k5 - c9*c212*k5 - c9*c219*k5 - c9*c220*k5 - c9*c221*k5 - c9*c228*k5 - c9*c229*k5 - c9*c230*k5 - c9*c237*k5 - c9*c238*k5 - c9*c239*k5 - c9*c246*k5 - c9*c247*k5 - c9*c248*k5 - c9*c255*k5 - c9*c256*k5 - c9*c257*k5 - c9*c302*k5 - c9*c305*k5 - c9*c308*k5 - c9*c311*k5 - c9*c314*k5 - c9*c317*k5 - c9*c320*k5 - c9*c323*k5 - c9*c359*k5b - c9*c362*k5b - c9*c365*k5b - c9*c368*k5b - c9*c371*k5b - c9*c374*k5b - c9*c377*k5b - c9*c380*k5b - c9*c383*k5b - c9*c386*k5b - c9*c389*k5b - c9*c392*k5b - c9*c395*k5b - c9*c398*k5b - c9*c401*k5b - c9*c404*k5b, ...
c86*kd60 - c404*k60c - c404*kd20 - c404*kd21 + c403*kd5b + c26*c392*k21 + c71*c392*k20 - c9*c404*k5b, ...
c86*kd60 - c401*k60c - c401*kd20 - c401*kd21 + c400*kd5b + c26*c389*k21 + c71*c389*k20 - c9*c401*k5b, ...
c86*kd60 - c398*k60c - c398*kd18 - c398*kd19 + c397*kd5b + c26*c392*k18 + c69*c392*k19 - c9*c398*k5b, ...
c86*kd60 - c395*k60c - c395*kd18 - c395*kd19 + c394*kd5b + c26*c389*k18 + c69*c389*k19 - c9*c395*k5b, ...
c86*kd60 - c392*k60c - c392*kd17 + c398*kd18 + c398*kd19 + c404*kd20 + c404*kd21 - c392*kd34 + c391*kd5b + c30*c346*k34 + c24*c386*k17 - c26*c392*k18 - c26*c392*k21 - c69*c392*k19 - c71*c392*k20 - c9*c392*k5b, ...
c86*kd60 - c389*k60c - c389*kd17 + c395*kd18 + c395*kd19 + c401*kd20 + c401*kd21 - c389*kd34 + c388*kd5b + c30*c343*k34 + c24*c383*k17 - c26*c389*k18 - c26*c389*k21 - c69*c389*k19 - c71*c389*k20 - c9*c389*k5b, ...
c86*kd60 - c386*k60c + c392*kd17 - c386*kd63 + c385*kd5b + c22*c346*k16 - c24*c386*k17 - c9*c386*k5b, ...
c86*kd60 - c365*k60c + c371*kd18 - c365*kd25 + c371*kd19 - c365*kd32 + c377*kd20 + c377*kd21 - c365*kd41 + c364*kd5b + c24*c359*k25 - c26*c365*k18 - c26*c365*k21 + c38*c343*k32 + c30*c353*k41 - c69*c365*k19 - c71*c365*k20 - c9*c365*k5b, ...
c86*kd60 - c383*k60c + c389*kd17 - c383*kd24 + c382*kd5b + c22*c343*k16 - c24*c383*k17 - c9*c383*k5b, ...
c86*kd60 - c380*k60c - c380*kd20 - c380*kd21 + c379*kd5b + c26*c368*k21 + c71*c368*k20 - c9*c380*k5b, ...
c86*kd60 - c377*k60c - c377*kd20 - c377*kd21 + c376*kd5b + c26*c365*k21 + c71*c365*k20 - c9*c377*k5b, ...
c86*kd60 - c374*k60c - c374*kd18 - c374*kd19 + c373*kd5b + c26*c368*k18 + c69*c368*k19 - c9*c374*k5b, ...
c86*kd60 - c371*k60c - c371*kd18 - c371*kd19 + c370*kd5b + c26*c365*k18 + c69*c365*k19 - c9*c371*k5b, ...
c86*kd60 - c368*k60c + c374*kd18 - c368*kd25 + c374*kd19 - c368*kd32 + c380*kd20 + c380*kd21 - c368*kd41 + c367*kd5b + c24*c362*k25 - c26*c368*k18 - c26*c368*k21 + c38*c346*k32 + c30*c356*k41 - c69*c368*k19 - c71*c368*k20 - c9*c368*k5b, ...
c86*kd60 - c362*k60c - c362*kd24 + c368*kd25 - c362*kd37 + c361*kd5b + c22*c356*k16 - c24*c362*k25 + c39*c346*k37 - c9*c362*k5b, ...
c86*kd60 - c359*k60c - c359*kd24 + c365*kd25 - c359*kd37 + c358*kd5b + c22*c353*k16 - c24*c359*k25 + c39*c343*k37 - c9*c359*k5b, ...
c86*kd60 - c323*k60b - c323*kd20 - c323*kd21 + c322*kd5b - c9*c323*k5 + c26*c317*k21 + c71*c317*k20, ...
c86*kd60 - c320*k60b - c320*kd18 - c320*kd19 + c319*kd5b - c9*c320*k5 + c26*c317*k18 + c69*c317*k19, ...
c86*kd60 - c317*k60b - c317*kd17 + c320*kd18 + c320*kd19 + c323*kd20 + c323*kd21 - c317*kd34 + c316*kd5b - c9*c317*k5 + c24*c314*k17 + c30*c293*k34 - c26*c317*k18 - c26*c317*k21 - c69*c317*k19 - c71*c317*k20, ...
c86*kd60 - c314*k60b + c317*kd17 - c314*kd63 + c313*kd5b - c9*c314*k5 + c22*c293*k16 - c24*c314*k17, ...
c86*kd60 - c311*k60b - c311*kd20 - c311*kd21 + c310*kd5b - c9*c311*k5 + c26*c305*k21 + c71*c305*k20, ...
c86*kd60 - c308*k60b - c308*kd18 - c308*kd19 + c307*kd5b - c9*c308*k5 + c26*c305*k18 + c69*c305*k19, ...
c308*kd18 + c308*kd19 - c305*kd25 + c311*kd20 + c311*kd21 - c305*kd32 - c305*kd41 + c304*kd5b - c9*c305*k5 - c26*c305*k18 + c24*c302*k25 - c26*c305*k21 + c38*c293*k32 + c30*c299*k41 - c69*c305*k19 - c71*c305*k20, ...
c86*kd60 - c302*k60b - c302*kd24 + c305*kd25 - c302*kd37 + c301*kd5b - c9*c302*k5 + c22*c299*k16 - c24*c302*k25 + c39*c293*k37, ...
c86*kd60 - c239*k60b - c239*kd17 + c248*kd18 + c248*kd19 - c239*kd34 + c257*kd20 + c257*kd21 + c242*kd5b + c30*c167*k34 - c9*c239*k5 + c24*c230*k17 - c26*c239*k18 - c26*c239*k21 - c69*c239*k19 - c71*c239*k20, ...
c86*kd60 - c238*k60b - c238*kd17 + c247*kd18 + c247*kd19 - c238*kd34 + c256*kd20 + c256*kd21 + c241*kd5b + c30*c166*k34 - c9*c238*k5 + c24*c229*k17 - c26*c238*k18 - c26*c238*k21 - c69*c238*k19 - c71*c238*k20, ...
c86*kd60 - c237*k60b - c237*kd17 + c246*kd18 + c246*kd19 - c237*kd34 + c255*kd20 + c255*kd21 + c240*kd5b + c30*c165*k34 - c9*c237*k5 + c24*c228*k17 - c26*c237*k18 - c26*c237*k21 - c69*c237*k19 - c71*c237*k20, ...
c86*kd60 - c257*k60b - c257*kd20 - c257*kd21 + c260*kd5b - c9*c257*k5 + c26*c239*k21 + c71*c239*k20, ...
c86*kd60 - c256*k60b - c256*kd20 - c256*kd21 + c259*kd5b - c9*c256*k5 + c26*c238*k21 + c71*c238*k20, ...
c86*kd60 - c255*k60b - c255*kd20 - c255*kd21 + c258*kd5b - c9*c255*k5 + c26*c237*k21 + c71*c237*k20, ...
c86*kd60 - c248*k60b - c248*kd18 - c248*kd19 + c251*kd5b - c9*c248*k5 + c26*c239*k18 + c69*c239*k19, ...
c86*kd60 - c247*k60b - c247*kd18 - c247*kd19 + c250*kd5b - c9*c247*k5 + c26*c238*k18 + c69*c238*k19, ...
c86*kd60 - c246*k60b - c246*kd18 - c246*kd19 + c249*kd5b - c9*c246*k5 + c26*c237*k18 + c69*c237*k19, ...
c86*kd60 - c230*k60b - c230*kd24 + c239*kd17 + c233*kd5b + c22*c167*k16 - c9*c230*k5 - c24*c230*k17, ...
c86*kd60 - c229*k60b - c229*kd24 + c238*kd17 + c232*kd5b + c22*c166*k16 - c9*c229*k5 - c24*c229*k17, ...
c86*kd60 - c228*k60b - c228*kd24 + c237*kd17 + c231*kd5b + c22*c165*k16 - c9*c228*k5 - c24*c228*k17, ...
c86*kd60 - c221*k60b - c221*kd20 - c221*kd21 + c224*kd5b - c9*c221*k5 + c26*c203*k21 + c71*c203*k20, ...
c86*kd60 - c220*k60b - c220*kd20 - c220*kd21 + c223*kd5b - c9*c220*k5 + c26*c202*k21 + c71*c202*k20, ...
c86*kd60 - c219*k60b - c219*kd20 - c219*kd21 + c222*kd5b - c9*c219*k5 + c26*c201*k21 + c71*c201*k20, ...
c86*kd60 - c212*k60b - c212*kd18 - c212*kd19 + c215*kd5b - c9*c212*k5 + c26*c203*k18 + c69*c203*k19, ...
c86*kd60 - c211*k60b - c211*kd18 - c211*kd19 + c214*kd5b - c9*c211*k5 + c26*c202*k18 + c69*c202*k19, ...
c86*kd60 - c210*k60b - c210*kd18 - c210*kd19 + c213*kd5b - c9*c210*k5 + c26*c201*k18 + c69*c201*k19, ...
c86*kd60 - c203*k60b - c203*kd25 + c212*kd18 + c212*kd19 - c203*kd32 + c221*kd20 + c221*kd21 - c203*kd41 + c206*kd5b - c9*c203*k5 + c38*c167*k32 + c24*c194*k25 - c26*c203*k18 - c26*c203*k21 + c30*c185*k41 - c69*c203*k19 - c71*c203*k20, ...
c86*kd60 - c202*k60b - c202*kd25 + c211*kd18 + c211*kd19 - c202*kd32 + c220*kd20 + c220*kd21 - c202*kd41 + c205*kd5b - c9*c202*k5 + c38*c166*k32 + c24*c193*k25 - c26*c202*k18 - c26*c202*k21 + c30*c184*k41 - c69*c202*k19 - c71*c202*k20, ...
c86*kd60 - c201*k60b - c201*kd25 + c210*kd18 + c210*kd19 - c201*kd32 + c219*kd20 + c219*kd21 - c201*kd41 + c204*kd5b - c9*c201*k5 + c38*c165*k32 + c24*c192*k25 - c26*c201*k18 - c26*c201*k21 + c30*c183*k41 - c69*c201*k19 - c71*c201*k20, ...
c86*kd60 - c194*k60b - c194*kd24 + c203*kd25 - c194*kd37 + c197*kd5b - c9*c194*k5 + c22*c185*k16 - c24*c194*k25 + c39*c167*k37, ...
c86*kd60 - c193*k60b - c193*kd24 + c202*kd25 - c193*kd37 + c196*kd5b - c9*c193*k5 + c22*c184*k16 - c24*c193*k25 + c39*c166*k37, ...
c86*kd60 - c192*k60b - c192*kd24 + c201*kd25 - c192*kd37 + c195*kd5b - c9*c192*k5 + c22*c183*k16 - c24*c192*k25 + c39*c165*k37, ...
c37*k6 - c68*k60 - c68*kd20 - c68*kd21 + c94*kd5 + c86*kd60 - c68*stepfunc(time, 1799.99, 0, 1800, 0.00005) - c9*c68*k5 + c26*c66*k21 + c66*c71*k20, ...
c36*k6 - c67*k60 - c67*kd18 - c67*kd19 + c93*kd5 + c86*kd60 - c67*stepfunc(time, 1799.99, 0, 1800, 0.00005) - c9*c67*k5 + c26*c66*k18 + c66*c69*k19, ...
c35*k6 - c66*k60 + c67*kd18 + c67*kd19 + c68*kd20 + c68*kd21 - c66*kd25 + c92*kd5 - c66*kd32 - c66*kd41 + c86*kd60 + c98*kd64 - c66*stepfunc(time, 1799.99, 0, 1800, 0.00005) - c9*c66*k5 + c17*c38*k32 - c26*c66*k18 - c26*c66*k21 + c24*c65*k25 + c30*c64*k41 - c66*c69*k19 - c66*c71*k20 - c66*c83*k64, ...
c34*k6 - c65*k60 - c65*kd24 + c66*kd25 + c91*kd5 - c65*kd37 + c86*kd60 + c420*kd101 - c65*stepfunc(time, 1799.99, 0, 1800, 0.00005) - c9*c65*k5 + c17*c39*k37 + c22*c64*k16 - c24*c65*k25 - c65*c103*k101, ...
c29*k6 - c21*k60 - c21*kd20 - c21*kd21 + c90*kd5 + c86*kd60 - c21*stepfunc(time, 1799.99, 0, 1800, 0.00005) - c9*c21*k5 + c19*c26*k21 + c19*c71*k20, ...
c27*k6 - c20*k60 - c20*kd18 - c20*kd19 + c89*kd5 + c86*kd60 - c20*stepfunc(time, 1799.99, 0, 1800, 0.00005) - c9*c20*k5 + c19*c26*k18 + c19*c69*k19, ...
c23*k6 - c18*k60 + c7*kd5 + c19*kd17 - c18*kd63 + c86*kd60 + c100*kd101 - c18*stepfunc(time, 1799.99, 0, 1800, 0.00005) - c9*c18*k5 + c17*c22*k16 - c18*c24*k17 - c18*c103*k101, ...
c25*k6 - c19*k60 - c19*kd17 + c20*kd18 + c20*kd19 + c21*kd20 + c21*kd21 - c19*kd34 + c88*kd5 + c86*kd60 + c96*kd64 - c19*stepfunc(time, 1799.99, 0, 1800, 0.00005) - c9*c19*k5 + c18*c24*k17 - c19*c26*k18 - c19*c26*k21 + c17*c30*k34 - c19*c69*k19 - c19*c71*k20 - c19*c83*k64, ...
c2*k6 - c6*k60 + c10*kd10 + c86*kd60 + c518*kd120 + c519*kd120 - c6*stepfunc(time, 1799.99, 0, 1800, 0.00005) - c6*c16*k10b - c6*c157*k120b - c6*c158*k120b, ...
c15*kd8 - c5*k6 - c5*kd102 + c116*kd123 + c115*kd123h + c121*kd123h + c555*kd123 + c557*kd123 + c556*kd123h + c558*kd123h + c330^2*k102 + c8*stepfunc(time, 1799.99, 0, 1800, 0.00005) - c5*c14*k8 - 3*c5*c105*k123 - 4*c5*c105*k123h, ...
c5*k6 + c17*kd8 + c126*kd123 + c415*kd94 - c8*stepfunc(time, 1799.99, 0, 1800, 0.00005) - c8*c14*k8 - c8*c105*k123 - c8*c280*k94, ...
c32*kd22 - c15*kd8 - c15*k6 + c25*kd34 + c35*kd32 + c33*kd37 + c34*kd37 + c23*kd63 + c17*stepfunc(time, 1799.99, 0, 1800, 0.00005) + c5*c14*k8 - c15*c22*k16 - c15*c31*k22 - c15*c30*k34 - c15*c38*k32 - c15*c39*k37 - c15*c40*k37, ...
c15*k6 - c17*k60 - c17*kd8 + c19*kd34 + c18*kd63 + c63*kd22 + c66*kd32 + c64*kd37 + c65*kd37 + c86*kd60 - c17*stepfunc(time, 1799.99, 0, 1800, 0.00005) + c8*c14*k8 - c17*c22*k16 - c17*c31*k22 - c17*c30*k34 - c17*c38*k32 - c17*c39*k37 - c17*c40*k37, ...
c33*kd23 - c32*k23 - c32*kd22 - c32*k6 + c63*stepfunc(time, 1799.99, 0, 1800, 0.00005) + c15*c31*k22, ...
c32*k6 - c63*k23 - c63*k60 - c63*kd22 + c64*kd23 + c86*kd60 - c63*stepfunc(time, 1799.99, 0, 1800, 0.00005) + c17*c31*k22, ...
c32*k23 - c33*k6 - c33*kd23 + c34*kd24 - c33*kd37 + c35*kd41 + c64*stepfunc(time, 1799.99, 0, 1800, 0.00005) - c22*c33*k16 + c15*c40*k37 - c30*c33*k41, ...
c33*k6 + c63*k23 - c64*k60 - c64*kd23 + c65*kd24 - c64*kd37 + c66*kd41 + c86*kd60 - c64*stepfunc(time, 1799.99, 0, 1800, 0.00005) + c17*c40*k37 - c22*c64*k16 - c30*c64*k41, ...
c351*kd23 - c347*k6b - c347*k23 + c349*kd6b - c347*kd22b + c31*c341*k22, ...
c347*k6b - c349*k60c - c349*k23 + c86*kd60 + c353*kd23 - c349*kd6b - c349*kd22b + c31*c343*k22, ...
c354*kd23 - c348*k23 - c348*kd22 - c348*k6 + c350*stepfunc(time, 1799.99, 0, 1800, 0.00005) + c31*c344*k22, ...
c348*k6 - c350*k23 - c350*k60c + c86*kd60 - c350*kd22 + c356*kd23 - c350*stepfunc(time, 1799.99, 0, 1800, 0.00005) + c31*c346*k22, ...
c347*k23 - c351*k6b - c351*kd23 + c357*kd24 - c351*kd37 + c363*kd41 + c353*kd6b - c22*c351*k16 + c40*c341*k37 - c30*c351*k41, ...
c349*k23 - c353*k60c + c351*k6b + c86*kd60 - c353*kd23 + c359*kd24 - c353*kd37 + c365*kd41 - c353*kd6b - c22*c353*k16 + c40*c343*k37 - c30*c353*k41, ...
c502*k6b - c508*kd6b, ...
c503*k6b - c512*kd6b, ...
c348*k23 - c354*k6b - c354*kd23 + c360*kd24 - c354*kd37 + c366*kd41 + c356*kd6b - c22*c354*k16 + c40*c344*k37 - c30*c354*k41, ...
c350*k23 - c356*k60c + c354*k6b + c86*kd60 - c356*kd23 + c362*kd24 - c356*kd37 + c368*kd41 - c356*kd6b - c22*c356*k16 + c40*c346*k37 - c30*c356*k41, ...
c151*kd8 - c148*k7 + c162*kd7 + c122*kd123 - c148*kd102 - c14*c148*k8 - c105*c148*k123 + c87*c330*k102, ...
c148*k7 - c162*kd7 + c123*kd123 + c165*kd8b + c416*kd94 - c105*c162*k123 - c162*c280*k94 - c14*c162*k8b, ...
c163*kd7 - c149*k7 + c127*kd123 - c149*kd102 + c152*kd8b + c137*kd123 - 2*c105*c149*k123 + c330*c331*k102 - c14*c149*k8b, ...
c149*k7 - c163*kd7 + c124*kd123 + c166*kd8b + c281*kd94 - c105*c163*k123 - c163*c280*k94b - c14*c163*k8b, ...
c164*kd7 - c150*k7 + c128*kd123 - c150*kd102 + c153*kd8b + c138*kd123 - 2*c105*c150*k123 + c330*c332*k102 - c14*c150*k8b, ...
c150*k7 - c164*kd7 + c125*kd123 + c167*kd8b + c282*kd94 - c105*c164*k123 - c164*c280*k94b - c14*c164*k8b, ...
c129*kd123 - c289*k7 + c290*kd7 + c291*kd8 - c289*kd96 + c87^2*k96 - c14*c289*k8 - c105*c289*k123, ...
c289*k7 - c290*kd7 + c293*kd8 + c283*kd94 - c14*c290*k8 - c280*c290*k94, ...
c335*k7 + c169*kd123 - c337*kd7 + c343*kd8 + c417*kd94 - c14*c337*k8 - c105*c337*k123 - c280*c337*k94, ...
c336*k7 + c170*kd123 - c338*kd7 + c346*kd8 + c418*kd94 - c14*c338*k8 - c105*c338*k123 - c280*c338*k94, ...
c293*kd7 - c291*kd8 - c291*k7 + c297*kd37 + c303*kd32 + c300*kd37 + c315*kd34 + c312*kd63 + c294*kd22b + c14*c289*k8 - c22*c291*k16 - c31*c291*k22 - c30*c291*k34 - c38*c291*k32 - c39*c291*k37 - c40*c291*k37, ...
c291*k7 - c293*k60b + c86*kd60 - c293*kd7 - c293*kd8 + c299*kd37 + c305*kd32 + c302*kd37 + c317*kd34 + c314*kd63 + c296*kd22b + c14*c290*k8 - c22*c293*k16 - c31*c293*k22 - c30*c293*k34 - c38*c293*k32 - c39*c293*k37 - c40*c293*k37, ...
c296*kd7 - c294*k23 - c294*k7 + c297*kd23 - c294*kd22b + c31*c291*k22, ...
c294*k7 - c296*k23 - c296*k60b + c86*kd60 - c296*kd7 + c299*kd23 - c296*kd22b + c31*c293*k22, ...
c294*k23 - c297*k7 + c299*kd7 - c297*kd23 + c300*kd24 - c297*kd37 + c303*kd41 - c22*c297*k16 - c30*c297*k41 + c40*c291*k37, ...
c297*k7 + c296*k23 - c299*k60b + c86*kd60 - c299*kd7 - c299*kd23 + c302*kd24 - c299*kd37 + c305*kd41 - c22*c299*k16 - c30*c299*k41 + c40*c293*k37, ...
c15*kd8 + c17*kd8 + c151*kd8 + c152*kd8b + c153*kd8b + c165*kd8b + c166*kd8b + c167*kd8b + c291*kd8 + c293*kd8 + c341*kd8 + c343*kd8 + c344*kd8 + c346*kd8 - c5*c14*k8 - c8*c14*k8 - c14*c148*k8 - c14*c289*k8 - c14*c290*k8 - c14*c335*k8 - c14*c336*k8 - c14*c337*k8 - c14*c338*k8 - c14*c149*k8b - c14*c150*k8b - c14*c162*k8b - c14*c163*k8b - c14*c164*k8b, ...
c183*kd37 + c192*kd37 + c201*kd32 + c228*kd24 - c165*kd8b + c237*kd34 + c174*kd22b - c22*c165*k16 - c31*c165*k22 - c30*c165*k34 - c38*c165*k32 - c39*c165*k37 - c40*c165*k37 + c14*c162*k8b, ...
c184*kd37 + c193*kd37 + c202*kd32 + c229*kd24 - c166*kd8b + c238*kd34 + c175*kd22b - c22*c166*k16 - c31*c166*k22 - c30*c166*k34 - c38*c166*k32 - c39*c166*k37 - c40*c166*k37 + c14*c163*k8b, ...
c185*kd37 + c194*kd37 + c203*kd32 + c230*kd24 - c167*kd8b + c239*kd34 + c176*kd22b - c22*c167*k16 - c31*c167*k22 - c30*c167*k34 - c38*c167*k32 - c39*c167*k37 - c40*c167*k37 + c14*c164*k8b, ...
c180*kd37 - c151*kd8 + c189*kd37 + c198*kd32 + c225*kd24 + c234*kd34 + c171*kd22b + c14*c148*k8 - c22*c151*k16 - c31*c151*k22 - c30*c151*k34 - c38*c151*k32 - c39*c151*k37 - c40*c151*k37, ...
c181*kd37 + c190*kd37 + c199*kd32 + c226*kd24 - c152*kd8b + c235*kd34 + c172*kd22b - c22*c152*k16 - c31*c152*k22 - c30*c152*k34 - c38*c152*k32 - c39*c152*k37 - c40*c152*k37 + c14*c149*k8b, ...
c182*kd37 + c191*kd37 + c200*kd32 + c227*kd24 - c153*kd8b + c236*kd34 + c173*kd22b - c22*c153*k16 - c31*c153*k22 - c30*c153*k34 - c38*c153*k32 - c39*c153*k37 - c40*c153*k37 + c14*c150*k8b, ...
c351*kd37 - c341*kd8 + c357*kd37 + c363*kd32 + c387*kd34 + c381*kd63 + c347*kd22b + c14*c335*k8 - c22*c341*k16 - c31*c341*k22 - c30*c341*k34 - c38*c341*k32 - c39*c341*k37 - c40*c341*k37, ...
c348*kd22 - c344*kd8 + c354*kd37 + c360*kd37 + c366*kd32 + c384*kd24 + c390*kd34 + c14*c336*k8 - c22*c344*k16 - c31*c344*k22 - c30*c344*k34 - c38*c344*k32 - c39*c344*k37 - c40*c344*k37, ...
c353*kd37 - c343*kd8 + c359*kd37 + c365*kd32 + c383*kd24 + c389*kd34 + c349*kd22b + c14*c337*k8 - c22*c343*k16 - c31*c343*k22 - c30*c343*k34 - c38*c343*k32 - c39*c343*k37 - c40*c343*k37, ...
c350*kd22 - c346*kd8 + c356*kd37 + c362*kd37 + c368*kd32 + c392*kd34 + c386*kd63 + c14*c338*k8 - c22*c346*k16 - c31*c346*k22 - c30*c346*k34 - c38*c346*k32 - c39*c346*k37 - c40*c346*k37, ...
2*c10*kd10 - c16*k61 + c13*kd61 - c6*c16*k10b - c16*c530*k10b, ...
c421*kd120 - c157*kd10 + c518*kd120 - c6*c157*k120b - c155*c157*k120 + c154*c515*k10b, ...
c34*kd24 + c39*kd24 + c30*kd35 + c18*kd63 + c23*kd63 + c65*kd24 + c189*kd24 + c190*kd24 + c191*kd24 + c192*kd24 + c193*kd24 + c194*kd24 + c225*kd24 + c226*kd24 + c227*kd24 + c228*kd24 + c229*kd24 + c230*kd24 + c300*kd24 + c302*kd24 + c312*kd63 + c314*kd63 + c357*kd24 + c359*kd24 + c360*kd24 + c362*kd24 + c383*kd24 + c384*kd24 + c381*kd63 + c386*kd63 - c15*c22*k16 - c17*c22*k16 - c22*c33*k16 - c22*c40*k16 - c22*c24*k35 - c22*c64*k16 - c22*c151*k16 - c22*c152*k16 - c22*c153*k16 - c22*c165*k16 - c22*c166*k16 - c22*c167*k16 - c22*c180*k16 - c22*c181*k16 - c22*c182*k16 - c22*c183*k16 - c22*c184*k16 - c22*c185*k16 - c22*c291*k16 - c22*c293*k16 - c22*c297*k16 - c22*c299*k16 - c22*c341*k16 - c22*c343*k16 - c22*c344*k16 - c22*c346*k16 - c22*c351*k16 - c22*c353*k16 - c22*c354*k16 - c22*c356*k16, ...
c39*kd24 - c40*k36 + c31*kd36 + c33*kd37 + c38*kd33 + c64*kd37 + c180*kd37 + c181*kd37 + c182*kd37 + c183*kd37 + c184*kd37 + c185*kd37 + c297*kd37 + c299*kd37 + c351*kd37 + c353*kd37 + c354*kd37 + c356*kd37 - c22*c40*k16 - c15*c40*k37 - c17*c40*k37 - c30*c40*k33 - c40*c151*k37 - c40*c152*k37 - c40*c153*k37 - c40*c165*k37 - c40*c166*k37 - c40*c167*k37 - c40*c291*k37 - c40*c293*k37 - c40*c341*k37 - c40*c343*k37 - c40*c344*k37 - c40*c346*k37, ...
c34*kd37 - c39*kd24 + c38*kd40 + c65*kd37 + c189*kd37 + c190*kd37 + c191*kd37 + c192*kd37 + c193*kd37 + c194*kd37 + c300*kd37 + c302*kd37 + c357*kd37 + c359*kd37 + c360*kd37 + c362*kd37 + c22*c40*k16 - c15*c39*k37 - c17*c39*k37 - c24*c39*k40 - c39*c151*k37 - c39*c152*k37 - c39*c153*k37 - c39*c165*k37 - c39*c166*k37 - c39*c167*k37 - c39*c291*k37 - c39*c293*k37 - c39*c341*k37 - c39*c343*k37 - c39*c344*k37 - c39*c346*k37, ...
c171*k23 - c180*kd23 + c189*kd24 - c180*kd37 + c198*kd41 - c22*c180*k16 + c40*c151*k37 - c30*c180*k41, ...
c172*k23 - c181*kd23 + c190*kd24 - c181*kd37 + c199*kd41 - c22*c181*k16 + c40*c152*k37 - c30*c181*k41, ...
c173*k23 - c182*kd23 + c191*kd24 - c182*kd37 + c200*kd41 - c22*c182*k16 + c40*c153*k37 - c30*c182*k41, ...
c174*k23 - c183*kd23 + c192*kd24 - c183*kd37 + c201*kd41 - c22*c183*k16 + c40*c165*k37 - c30*c183*k41, ...
c175*k23 - c184*kd23 + c193*kd24 - c184*kd37 + c202*kd41 - c22*c184*k16 + c40*c166*k37 - c30*c184*k41, ...
c176*k23 - c185*kd23 + c194*kd24 - c185*kd37 + c203*kd41 - c22*c185*k16 + c40*c167*k37 - c30*c185*k41, ...
c19*kd17 + c25*kd17 + c35*kd25 + c30*kd35 + c38*kd40 + c66*kd25 + c101*kd64 + c102*kd64 + c198*kd25 + c199*kd25 + c200*kd25 + c201*kd25 + c202*kd25 + c203*kd25 + c234*kd17 + c235*kd17 + c236*kd17 + c237*kd17 + c238*kd17 + c239*kd17 + c303*kd25 + c305*kd25 + c315*kd17 + c317*kd17 + c363*kd25 + c365*kd25 + c366*kd25 + c368*kd25 + c387*kd17 + c389*kd17 + c390*kd17 + c392*kd17 - c18*c24*k17 - c23*c24*k17 - c22*c24*k35 - c24*c34*k25 - c24*c39*k40 - c24*c65*k25 - c24*c59*k64 - c24*c83*k64 - c24*c189*k25 - c24*c190*k25 - c24*c191*k25 - c24*c192*k25 - c24*c193*k25 - c24*c194*k25 - c24*c225*k17 - c24*c226*k17 - c24*c227*k17 - c24*c228*k17 - c24*c229*k17 - c24*c230*k17 - c24*c300*k25 - c24*c302*k25 - c24*c312*k17 - c24*c314*k17 - c24*c357*k25 - c24*c359*k25 - c24*c360*k25 - c24*c362*k25 - c24*c381*k17 - c24*c383*k17 - c24*c384*k17 - c24*c386*k17, ...
c20*kd18 + c21*kd21 + c27*kd18 + c29*kd21 + c36*kd18 + c37*kd21 + c67*kd18 + c68*kd21 + c207*kd18 + c208*kd18 + c209*kd18 + c210*kd18 + c211*kd18 + c212*kd18 + c216*kd21 + c217*kd21 + c218*kd21 + c219*kd21 + c220*kd21 + c221*kd21 + c243*kd18 + c244*kd18 + c245*kd18 + c246*kd18 + c247*kd18 + c248*kd18 + c252*kd21 + c253*kd21 + c254*kd21 + c255*kd21 + c256*kd21 + c257*kd21 + c306*kd18 + c308*kd18 + c309*kd21 + c311*kd21 + c318*kd18 + c320*kd18 + c321*kd21 + c323*kd21 + c264*kd112 + c265*kd112 + c266*kd112 + c267*kd112 + c268*kd112 + c269*kd112 + c369*kd18 + c371*kd18 + c372*kd18 + c374*kd18 + c375*kd21 + c377*kd21 + c378*kd21 + c380*kd21 + c393*kd18 + c395*kd18 + c396*kd18 + c398*kd18 + c399*kd21 + c401*kd21 + c402*kd21 + c404*kd21 + c325*kd112 - c19*c26*k18 - c19*c26*k21 - c25*c26*k18 - c25*c26*k21 - c26*c35*k18 - c26*c35*k21 - c26*c66*k18 - c26*c66*k21 - c26*c104*k112 - c26*c198*k18 - c26*c199*k18 - c26*c200*k18 - c26*c198*k21 - c26*c201*k18 - c26*c199*k21 - c26*c202*k18 - c26*c200*k21 - c26*c203*k18 - c26*c201*k21 - c26*c202*k21 - c26*c203*k21 - c26*c234*k18 - c26*c235*k18 - c26*c236*k18 - c26*c234*k21 - c26*c237*k18 - c26*c235*k21 - c26*c238*k18 - c26*c236*k21 - c26*c239*k18 - c26*c237*k21 - c26*c238*k21 - c26*c239*k21 - c26*c303*k18 - c26*c305*k18 - c26*c303*k21 - c26*c305*k21 - c26*c315*k18 - c26*c317*k18 - c26*c315*k21 - c26*c317*k21 - c26*c261*k112 - c26*c262*k112 - c26*c263*k112 - c26*c363*k18 - c26*c365*k18 - c26*c363*k21 - c26*c366*k18 - c26*c365*k21 - c26*c368*k18 - c26*c366*k21 - c26*c368*k21 - c26*c387*k18 - c26*c389*k18 - c26*c387*k21 - c26*c390*k18 - c26*c389*k21 - c26*c392*k18 - c26*c390*k21 - c26*c392*k21 - c26*c324*k112 - c26*c405*k112 - c26*c408*k112, ...
c27*kd19 + c36*kd19 + c42*kd28 + c207*kd19 + c208*kd19 + c209*kd19 + c243*kd19 + c244*kd19 + c245*kd19 + c306*kd19 + c318*kd19 + c264*kd113 + c265*kd113 + c266*kd113 + c267*kd113 + c268*kd113 + c269*kd113 + c369*kd19 + c372*kd19 + c393*kd19 + c396*kd19 + c325*kd113 - c25*c28*k19 - c28*c35*k19 - c28*c41*k28 - c28*c104*k113 - c28*c198*k19 - c28*c199*k19 - c28*c200*k19 - c28*c234*k19 - c28*c235*k19 - c28*c236*k19 - c28*c303*k19 - c28*c315*k19 - c28*c261*k113 - c28*c262*k113 - c28*c263*k113 - c28*c363*k19 - c28*c366*k19 - c28*c387*k19 - c28*c390*k19 - c28*c324*k113 - c28*c405*k113 - c28*c479*k113, ...
c20*kd19 + c67*kd19 + c70*kd28 + c210*kd19 + c211*kd19 + c212*kd19 + c246*kd19 + c247*kd19 + c248*kd19 + c308*kd19 + c320*kd19 + c371*kd19 + c374*kd19 + c395*kd19 + c398*kd19 - c19*c69*k19 - c41*c69*k28 - c66*c69*k19 - c69*c201*k19 - c69*c202*k19 - c69*c203*k19 - c69*c237*k19 - c69*c238*k19 - c69*c239*k19 - c69*c305*k19 - c69*c317*k19 - c69*c365*k19 - c69*c368*k19 - c69*c389*k19 - c69*c392*k19, ...
c29*kd20 + c37*kd20 + c42*kd29 + c216*kd20 + c217*kd20 + c218*kd20 + c252*kd20 + c253*kd20 + c254*kd20 + c309*kd20 + c321*kd20 + c375*kd20 + c378*kd20 + c399*kd20 + c402*kd20 - c25*c43*k20 - c35*c43*k20 - c43*c45*k29 - c43*c198*k20 - c43*c199*k20 - c43*c200*k20 - c43*c234*k20 - c43*c235*k20 - c43*c236*k20 - c43*c303*k20 - c43*c315*k20 - c43*c363*k20 - c43*c366*k20 - c43*c387*k20 - c43*c390*k20, ...
c21*kd20 + c68*kd20 + c70*kd29 + c219*kd20 + c220*kd20 + c221*kd20 + c255*kd20 + c256*kd20 + c257*kd20 + c311*kd20 + c323*kd20 + c377*kd20 + c380*kd20 + c401*kd20 + c404*kd20 - c19*c71*k20 - c66*c71*k20 - c71*c72*k29 - c71*c201*k20 - c71*c202*k20 - c71*c203*k20 - c71*c237*k20 - c71*c238*k20 - c71*c239*k20 - c71*c305*k20 - c71*c317*k20 - c71*c365*k20 - c71*c368*k20 - c71*c389*k20 - c71*c392*k20, ...
c40*k36 + c32*kd22 - c31*kd36 + c63*kd22 + c171*kd22b + c172*kd22b + c173*kd22b + c174*kd22b + c175*kd22b + c176*kd22b + c348*kd22 + c350*kd22 + c294*kd22b + c296*kd22b + c347*kd22b + c349*kd22b - c15*c31*k22 - c17*c31*k22 - c31*c151*k22 - c31*c152*k22 - c31*c153*k22 - c31*c165*k22 - c31*c166*k22 - c31*c167*k22 - c31*c291*k22 - c31*c293*k22 - c31*c341*k22 - c31*c343*k22 - c31*c344*k22 - c31*c346*k22, ...
c180*kd23 - c171*k23 - c171*kd22b + c31*c151*k22, ...
c181*kd23 - c172*k23 - c172*kd22b + c31*c152*k22, ...
c182*kd23 - c173*k23 - c173*kd22b + c31*c153*k22, ...
c183*kd23 - c174*k23 - c174*kd22b + c31*c165*k22, ...
c184*kd23 - c175*k23 - c175*kd22b + c31*c166*k22, ...
c185*kd23 - c176*k23 - c176*kd22b + c31*c167*k22, ...
c42*kd28 + c46*kd43 + c70*kd28 + c73*kd43 - c28*c41*k28 - 2*c41*c44*k43 - c41*c69*k28, ...
c28*c41*k28 - c42*kd29 - c42*kd28 + c43*c45*k29, ...
c41*c69*k28 - c70*kd29 - c70*kd28 + c71*c72*k29, ...
c70*kd29 + c73*kd42 + c74*kd45 + c76*kd47 + c74*kd52 + c76*kd52 + c484*kd114 - c44*c72*k42 - c47*c72*k44 - c71*c72*k29 - c72*c75*k44 - c72*c75*k45 - c72*c77*k47 - c72*c497*k114, ...
c42*kd29 + c46*kd42 + c48*kd45 + c50*kd47 + c48*kd52 + c50*kd52 + c472*kd114 - c43*c45*k29 - c44*c45*k42 - c45*c47*k44 - c45*c49*k44 - c45*c49*k45 - c45*c51*k47 - c45*c497*k114, ...
c35*kd32 - c38*kd33 - c38*kd40 + c66*kd32 + c198*kd32 + c199*kd32 + c200*kd32 + c201*kd32 + c202*kd32 + c203*kd32 + c303*kd32 + c305*kd32 + c363*kd32 + c365*kd32 + c366*kd32 + c368*kd32 - c15*c38*k32 - c17*c38*k32 + c24*c39*k40 + c30*c40*k33 - c38*c151*k32 - c38*c152*k32 - c38*c153*k32 - c38*c165*k32 - c38*c166*k32 - c38*c167*k32 - c38*c291*k32 - c38*c293*k32 - c38*c341*k32 - c38*c343*k32 - c38*c344*k32 - c38*c346*k32, ...
c19*kd34 + c25*kd34 - c30*kd35 + c38*kd33 + c35*kd41 + c66*kd41 + c198*kd41 + c199*kd41 + c200*kd41 + c201*kd41 + c202*kd41 + c203*kd41 + c234*kd34 + c235*kd34 + c236*kd34 + c237*kd34 + c238*kd34 + c239*kd34 + c303*kd41 + c305*kd41 + c315*kd34 + c317*kd34 + c363*kd41 + c365*kd41 + c366*kd41 + c368*kd41 + c387*kd34 + c389*kd34 + c390*kd34 + c392*kd34 - c15*c30*k34 - c17*c30*k34 + c22*c24*k35 - c30*c40*k33 - c30*c33*k41 - c30*c64*k41 - c30*c151*k34 - c30*c152*k34 - c30*c153*k34 - c30*c165*k34 - c30*c166*k34 - c30*c167*k34 - c30*c180*k41 - c30*c181*k41 - c30*c182*k41 - c30*c183*k41 - c30*c184*k41 - c30*c185*k41 - c30*c291*k34 - c30*c293*k34 - c30*c297*k41 - c30*c299*k41 - c30*c341*k34 - c30*c343*k34 - c30*c344*k34 - c30*c346*k34 - c30*c351*k41 - c30*c353*k41 - c30*c354*k41 - c30*c356*k41, ...
c46*kd42 + c46*kd43 + c73*kd42 + c73*kd43 - 2*c41*c44*k43 - c44*c45*k42 - c44*c72*k42, ...
c41*c44*k43 - c73*kd43 - c73*kd42 + c44*c72*k42, ...
c41*c44*k43 - c46*kd43 - c46*kd42 + c44*c45*k42, ...
c74*kd45 + c78*kd49 + c76*kd52 + c79*kd50 - c53*c75*k49 - c53*c75*k50 - c72*c75*k44 - c72*c75*k45, ...
c72*c75*k44 - c76*kd52 - c76*kd47 + c72*c77*k47, ...
c48*kd52 + c54*kd49 + c74*kd52 + c79*kd49 - c45*c47*k44 - 2*c47*c53*k49 - c47*c72*k44, ...
c47*c72*k44 - c74*kd52 - c74*kd45 + c72*c75*k45, ...
c45*c47*k44 - c48*kd52 - c48*kd45 + c45*c49*k45, ...
c48*kd45 + c52*kd49 + c50*kd52 + c54*kd50 - c45*c49*k44 - c45*c49*k45 - c49*c53*k49 - c49*c53*k50, ...
c45*c49*k44 - c50*kd52 - c50*kd47 + c45*c51*k47, ...
c50*kd47 + c52*kd48 + c56*kd44 + c58*kd44 + c56*kd53 + c58*kd55 - c45*c51*k47 - c51*c53*k48 - c51*c55*k52 - c51*c57*k52 - c51*c57*k53 - c51*c59*k55, ...
c76*kd47 + c80*kd44 + c78*kd48 + c82*kd44 + c80*kd53 + c82*kd55 - c53*c77*k48 - c55*c77*k52 - c72*c77*k47 - c77*c81*k52 - c77*c81*k53 - c77*c83*k55, ...
c52*kd48 + c52*kd49 + c54*kd49 + c54*kd50 + c78*kd48 + c78*kd49 + c79*kd49 + c79*kd50 - 2*c47*c53*k49 - c49*c53*k49 - c49*c53*k50 - c51*c53*k48 - c53*c75*k49 - c53*c75*k50 - c53*c77*k48, ...
c53*c75*k49 - c78*kd49 - c78*kd48 + c53*c77*k48, ...
c49*c53*k49 - c52*kd49 - c52*kd48 + c51*c53*k48, ...
c47*c53*k49 - c54*kd50 - c54*kd49 + c49*c53*k50, ...
c47*c53*k49 - c79*kd50 - c79*kd49 + c53*c75*k50, ...
c56*kd44 + c62*kd57 + c80*kd44 + c85*kd57 - c51*c55*k52 - 2*c55*c60*k57 - c55*c77*k52, ...
c51*c55*k52 - c56*kd53 - c56*kd44 + c51*c57*k53, ...
c58*kd44 + c56*kd53 + c61*kd57 + c62*kd58 - c51*c57*k52 - c51*c57*k53 - c57*c60*k57 - c57*c60*k58, ...
c51*c57*k52 - c58*kd55 - c58*kd44 + c51*c59*k55, ...
c55*c77*k52 - c80*kd53 - c80*kd44 + c77*c81*k53, ...
c82*kd44 + c80*kd53 + c84*kd57 + c85*kd58 - c60*c81*k57 - c60*c81*k58 - c77*c81*k52 - c77*c81*k53, ...
c77*c81*k52 - c82*kd55 - c82*kd44 + c77*c83*k55, ...
c58*kd55 + c61*kd56 + c95*kd64 + c95*kd65 + c97*kd64 + c97*kd65 + c101*kd64 + c101*kd65 + c431*kd110 + c431*kd111 + c433*kd110 + c433*kd111 + c435*kd110 + c435*kd111 + c438*kd110 + c438*kd111 + c474*kd110 + c474*kd111 + c477*kd110 + c477*kd111 + c480*kd110 + c480*kd111 - c24*c59*k64 - c25*c59*k64 - c35*c59*k64 - c51*c59*k55 - c59*c60*k56 - c59*c99*k65 - c59*c103*k65 - c59*c419*k65 - c59*c409*k111 - c59*c410*k111 - c59*c430*k111 - c59*c445*k110 - c59*c446*k110 - c59*c447*k110 - c59*c454*k110 - c59*c457*k110 - c59*c460*k110 - c59*c486*k110 - c59*c487*k111 - c59*c488*k111 - c59*c490*k111 - c59*c491*k111, ...
c82*kd55 + c84*kd56 + c96*kd64 + c96*kd65 + c98*kd64 + c98*kd65 + c102*kd64 + c102*kd65 + c432*kd110 + c432*kd111 + c434*kd110 + c434*kd111 + c437*kd110 + c437*kd111 + c440*kd110 + c440*kd111 + c475*kd110 + c475*kd111 + c478*kd110 + c478*kd111 + c481*kd110 + c481*kd111 - c19*c83*k64 - c24*c83*k64 - c60*c83*k56 - c66*c83*k64 - c77*c83*k55 - c83*c100*k65 - c83*c103*k65 - c83*c420*k65 - c83*c409*k111 - c83*c410*k111 - c83*c430*k111 - c83*c445*k110 - c83*c446*k110 - c83*c447*k110 - c83*c454*k110 - c83*c457*k110 - c83*c460*k110 - c83*c486*k110 - c83*c487*k111 - c83*c488*k111 - c83*c490*k111 - c83*c491*k111, ...
c61*kd56 + c61*kd57 + c62*kd57 + c62*kd58 + c84*kd56 + c84*kd57 + c85*kd57 + c85*kd58 + c520*kd116 - c60*stepfunc(time, 2659.99, 0, 2660, 0.0150356) - 2*c55*c60*k57 - c57*c60*k57 - c57*c60*k58 - c59*c60*k56 - c60*c81*k57 - c60*c81*k58 - c60*c83*k56, ...
c57*c60*k57 - c61*kd57 - c61*kd56 + c59*c60*k56, ...
c60*c81*k57 - c84*kd57 - c84*kd56 + c60*c83*k56, ...
c55*c60*k57 - c62*kd58 - c62*kd57 + c57*c60*k58, ...
c55*c60*k57 - c85*kd58 - c85*kd57 + c60*c81*k58, ...
c6*k60 + c11*k60 + c17*k60 + c18*k60 + c19*k60 + c20*k60 + c21*k60 + c63*k60 + c64*k60 + c65*k60 + c66*k60 + c67*k60 + c68*k60 + c154*k60b + c155*k60b + c156*k60b + c159*k62b + c160*k62b + c161*k62b + c192*k60b + c193*k60b + c194*k60b + c201*k60b + c202*k60b + c203*k60b + c210*k60b + c211*k60b + c212*k60b + c219*k60b + c220*k60b + c221*k60b + c228*k60b + c229*k60b + c230*k60b + c237*k60b + c238*k60b + c239*k60b + c246*k60b + c247*k60b + c248*k60b + c255*k60b + c256*k60b + c257*k60b + c293*k60b + c296*k60b + c299*k60b + c302*k60b + c308*k60b + c311*k60b + c314*k60b + c317*k60b + c320*k60b + c323*k60b + c339*k62b + c340*k62b + c349*k60c + c350*k60c + c353*k60c + c356*k60c + c359*k60c + c362*k60c + c365*k60c + c368*k60c + c371*k60c + c374*k60c + c377*k60c + c380*k60c + c383*k60c + c386*k60c + c389*k60c + c392*k60c + c395*k60c + c398*k60c + c401*k60c + c404*k60c + c421*k62b + c422*k62b + c425*k60b + c518*k62b + c519*k62b - 68*c86*kd60 - 12*c86*kd60b, ...
c86*kd60 - c425*k60b + c283*kd95 - c280*c425*k95, ...
c16*k61 - c13*kd61, ...
c86*kd60b - c518*k62b - c518*kd120 + c6*c157*k120b, ...
c86*kd60b - c519*k62b - c519*kd120 + c6*c158*k120b, ...
c86*kd60b - c339*k62b + c417*kd95 - c280*c339*k95, ...
c86*kd60b - c340*k62b + c418*kd95 - c280*c340*k95, ...
c25*c59*k64 - c95*kd65 - c95*kd64 + c59*c99*k65, ...
c19*c83*k64 - c96*kd65 - c96*kd64 + c83*c100*k65, ...
c35*c59*k64 - c97*kd65 - c97*kd64 + c59*c419*k65, ...
c66*c83*k64 - c98*kd65 - c98*kd64 + c83*c420*k65, ...
c24*c59*k64 - c101*kd65 - c101*kd64 + c59*c103*k65, ...
c24*c83*k64 - c102*kd65 - c102*kd64 + c83*c103*k65, ...
c95*kd65 - c99*kd101 - c59*c99*k65 + c23*c103*k101, ...
c97*kd65 - c419*kd101 + c34*c103*k101 - c59*c419*k65, ...
c101*kd65 + c102*kd65 + c99*kd101 + c100*kd101 + c419*kd101 + c420*kd101 - c18*c103*k101 - c23*c103*k101 - c59*c103*k65 - c34*c103*k101 - c83*c103*k65 - c65*c103*k101, ...
c96*kd65 - c100*kd101 + c18*c103*k101 - c83*c100*k65, ...
c98*kd65 - c420*kd101 + c65*c103*k101 - c83*c420*k65, ...
c104*kd66 + c261*kd66 + c262*kd67 + c263*kd66 + c324*kd67 + c405*kd67 + c408*kd66 - c287*c445*k66 - c287*c446*k67 - c287*c447*k66 - c287*c454*k67 - c287*c457*k67 - c287*c460*k66 - c287*c486*k66, ...
c104*kd66 + c136*kd123 + c431*kd110 + c432*kd110 + c489*kd107 + c522*kd118 - c59*c486*k110 - c83*c486*k110 - c105*c486*k123 - c287*c486*k66 - c463*c486*k107 - c486*c521*k118, ...
c264*kd112 - c104*kd66 + c264*kd113 + c448*kd68 + c448*kd106b - c26*c104*k112 - c28*c104*k113 - c104*c106*k68 - c104*c444*k106b + c287*c486*k66, ...
c132*kd123 + c263*kd66 + c412*kd118 + c438*kd110 + c440*kd110 + c466*kd107 - c59*c447*k110 - c83*c447*k110 - c105*c447*k123 - c287*c447*k66 - c447*c463*k107 - c447*c521*k118, ...
c267*kd112 - c263*kd66 + c267*kd113 + c451*kd68 + c451*kd106b - c26*c263*k112 - c28*c263*k113 - c106*c263*k68 + c287*c447*k66 - c263*c444*k106b, ...
c130*kd123 + c261*kd66 + c424*kd118 + c433*kd110 + c434*kd110 + c464*kd107 - c59*c445*k110 - c83*c445*k110 - c105*c445*k123 - c287*c445*k66 - c445*c463*k107 - c445*c521*k118, ...
c265*kd112 - c261*kd66 + c265*kd113 + c449*kd68 + c449*kd106b - c26*c261*k112 - c28*c261*k113 - c106*c261*k68 + c287*c445*k66 - c261*c444*k106b, ...
c131*kd123 + c262*kd67 + c411*kd118 + c435*kd110 + c437*kd110 + c465*kd107 - c59*c446*k110 - c83*c446*k110 - c105*c446*k123 - c287*c446*k67 - c446*c463*k107 - c446*c521*k118, ...
c266*kd112 - c262*kd67 + c266*kd113 + c450*kd68 + c450*kd106b - c26*c262*k112 - c28*c262*k113 - c106*c262*k68 + c287*c446*k67 - c262*c444*k106b, ...
c133*kd123 + c324*kd67 + c473*kd107 + c474*kd110 + c475*kd110 + c523*kd118 - c59*c454*k110 - c83*c454*k110 - c105*c454*k123 - c287*c454*k67 - c454*c463*k107 - c454*c521*k118, ...
c268*kd112 + c268*kd113 - c324*kd67 + c452*kd68 + c452*kd106 - c26*c324*k112 - c28*c324*k113 - c106*c324*k68 + c287*c454*k67 - c324*c444*k106, ...
c134*kd123 + c405*kd67 + c456*kd118 + c476*kd107 + c477*kd110 + c478*kd110 - c59*c457*k110 - c83*c457*k110 - c105*c457*k123 - c287*c457*k67 - c457*c463*k107 - c457*c521*k118, ...
c269*kd112 + c269*kd113 - c405*kd67 + c453*kd106 + c453*kd68b - c26*c405*k112 - c28*c405*k113 - c106*c405*k68 + c287*c457*k67 - c405*c444*k106, ...
c135*kd123 + c408*kd66 + c407*kd118 + c479*kd107 + c480*kd110 + c481*kd110 - c59*c460*k110 - c83*c460*k110 - c105*c460*k123 - c287*c460*k66 - c460*c463*k107 - c460*c521*k118, ...
c325*kd112 - c408*kd66 + c455*kd106 - c26*c408*k112 + c287*c460*k66 - c408*c444*k106, ...
c108*kd69 + c111*kd76 + c448*kd68 + c449*kd68 + c450*kd68 + c451*kd68 + c452*kd68 + c495*kd69 + c462*kd109 + c482*kd109 + c453*kd68b + c467*kd68b + c468*kd68b + c469*kd68b + c470*kd68b + c471*kd68b - c104*c106*k68 - c106*c107*k69 - c106*c112*k69 - c106*c109*k76 - c106*c261*k68 - c106*c262*k68 - c106*c263*k68 - c106*c279*k109 - c106*c324*k68 - c106*c405*k68 - c106*c453*k68 - c106*c467*k68 - c106*c468*k68 - c106*c469*k68 - c106*c470*k68 - c106*c461*k109, ...
c467*kd106 - c453*kd106 - c453*kd68b + c467*kd68b + c106*c405*k68 - c106*c453*k68 + c405*c444*k106 - c444*c453*k106, ...
c106*c324*k68 - c452*kd106 - c452*kd68 + c324*c444*k106, ...
c104*c106*k68 - c448*kd106b - c448*kd68 + c104*c444*k106b, ...
c106*c261*k68 - c449*kd106b - c449*kd68 + c261*c444*k106b, ...
c106*c262*k68 - c450*kd106b - c450*kd68 + c262*c444*k106b, ...
c106*c263*k68 - c451*kd106b - c451*kd68 + c263*c444*k106b, ...
c468*kd106 - c467*kd106 - c467*kd68b + c468*kd68b + c106*c453*k68 - c106*c467*k68 + c444*c453*k106 - c444*c467*k106, ...
c469*kd106 - c468*kd106 - c468*kd68b + c469*kd68b + c106*c467*k68 - c106*c468*k68 + c444*c467*k106 - c444*c468*k106, ...
c470*kd106 - c469*kd106 - c469*kd68b + c470*kd68b + c106*c468*k68 - c106*c469*k68 + c444*c468*k106 - c444*c469*k106, ...
c471*kd106 - c470*kd106 - c470*kd68b + c471*kd68b + c106*c469*k68 - c106*c470*k68 + c444*c469*k106 - c444*c470*k106, ...
c106*c470*k68 - c471*kd68b - c471*kd106 + c444*c470*k106, ...
c108*kd69 + c114*kd75 - c106*c107*k69 - c107*c113*k75, ...
c110*kd70 - c108*kd69 + c106*c107*k69 - c108*c109*k70, ...
c110*kd71 + c114*kd73 + c495*kd69 + c498*kd75 - c106*c112*k69 - c111*c112*k71 - c112*c113*k73 - c112*c113*k75, ...
c496*kd70 - c495*kd69 + c106*c112*k69 - c109*c495*k70, ...
c110*kd70 + c111*kd76 + c496*kd70 - c108*c109*k70 - c106*c109*k76 - c109*c495*k70, ...
c109*c495*k70 - c496*kd72 - c496*kd70 + c111*c497*k72, ...
c108*c109*k70 - c110*kd71 - c110*kd70 + c111*c112*k71, ...
c110*kd71 - c111*kd76 + c496*kd72 + c106*c109*k76 - c111*c112*k71 - c111*c497*k72, ...
c496*kd72 + c498*kd74 + c472*kd114 + c472*kd115 + c484*kd114 + c484*kd115 - c45*c497*k114 - c111*c497*k72 - c72*c497*k114 - c113*c497*k74 - 2*c485*c497*k115, ...
c114*kd73 + c114*kd75 + c498*kd74 + c498*kd75 - c107*c113*k75 - c112*c113*k73 - c112*c113*k75 - c113*c497*k74, ...
c112*c113*k75 - c498*kd75 - c498*kd74 + c113*c497*k74, ...
c107*c113*k75 - c114*kd75 - c114*kd73 + c112*c113*k73, ...
c281*kd94 + c281*kd95 + c282*kd94 + c282*kd95 + c283*kd94 + c283*kd95 + c415*kd94 + c415*kd95 + c416*kd94 + c416*kd95 + c417*kd94 + c417*kd95 + c418*kd94 + c418*kd95 - c8*c280*k94 - c11*c280*k95 - c159*c280*k95 - c160*c280*k95 - c161*c280*k95 - c162*c280*k94 - c163*c280*k94b - c164*c280*k94b - c280*c290*k94 - c280*c337*k94 - c280*c338*k94 - c280*c339*k95 - c280*c340*k95 - c280*c425*k95, ...
c160*c280*k95 - c281*kd95 - c281*kd94 + c163*c280*k94b, ...
c161*c280*k95 - c282*kd95 - c282*kd94 + c164*c280*k94b, ...
c8*c280*k94 - c415*kd95 - c415*kd94 + c11*c280*k95, ...
c280*c290*k94 - c283*kd95 - c283*kd94 + c280*c425*k95, ...
c280*c337*k94 - c417*kd95 - c417*kd94 + c280*c339*k95, ...
c280*c338*k94 - c418*kd95 - c418*kd94 + c280*c340*k95, ...
c159*c280*k95 - c416*kd95 - c416*kd94 + c162*c280*k94, ...
c148*kd102 + c289*kd96 + c284*kd103 + c335*kd103 + c336*kd103 + c509*kd103 - c87^2*k96 - c87*c141*k103 - c87*c330*k102 - c87*c331*k103 - c87*c332*k103 - c87*c502*k103, ...
c286*kd97 + c2*stepfunc(time, 1799.99, 0, 1800, 1) - c105*c531*stepfunc(time, 1799.99, 0, 1800, 0.000000018704) - (c285*c531*kd97)/KD_iressa_wt, ...
c5*kd102 + c148*kd102 + c149*kd102 + c150*kd102 - c330^2*k102 - c87*c330*k102 - c330*c331*k102 - c330*c332*k102, ...
c149*kd102 + c335*kd103 - c87*c331*k103 - c330*c331*k102, ...
c150*kd102 + c336*kd103 - c87*c332*k103 - c330*c332*k102, ...
c87*c502*k103 - c509*kd103, ...
c140*c502*k103 - c510*kd103, ...
c143*c502*k103 - c511*kd103, ...
c141*c503*k103 - c513*kd103, ...
c462*kd104 + c462*kd109 - c106*c461*k109 - c444*c461*k104, ...
c452*kd106 + c453*kd106 + c455*kd106 + c462*kd104 + c467*kd106 + c468*kd106 + c469*kd106 + c470*kd106 + c471*kd106 + c482*kd104 + c448*kd106b + c449*kd106b + c450*kd106b + c451*kd106b - c104*c444*k106b - c279*c444*k104 - c324*c444*k106 - c261*c444*k106b - c262*c444*k106b - c263*c444*k106b - c405*c444*k106 - c408*c444*k106 - c444*c453*k106 - c444*c461*k104 - c444*c467*k106 - c444*c468*k106 - c444*c469*k106 - c444*c470*k106, ...
c106*c461*k109 - c462*kd109 - c462*kd104 + c444*c461*k104, ...
c482*kd104 + c482*kd109 - c106*c279*k109 - c279*c444*k104, ...
c106*c279*k109 - c482*kd109 - c482*kd104 + c279*c444*k104, ...
c427*kd105 + c428*kd105 + c429*kd105 + c436*kd105 + c439*kd105 + c442*kd105 + c483*kd105 - c23*c426*k105 - c225*c426*k105 - c226*c426*k105 - c227*c426*k105 - c312*c426*k105 - c381*c426*k105 - c384*c426*k105, ...
c408*c444*k106 - c455*kd106, ...
c464*kd107 + c464*kd108 + c465*kd107 + c465*kd108 + c466*kd107 + c466*kd108 + c473*kd107 + c473*kd108 + c476*kd107 + c476*kd108 + c479*kd107 + c479*kd108 + c489*kd107 + c489*kd108 - c427*c463*k108 - c428*c463*k108 - c429*c463*k108 - c436*c463*k108 - c439*c463*k108 - c442*c463*k108 - c445*c463*k107 - c446*c463*k107 - c447*c463*k107 - c454*c463*k107 - c457*c463*k107 - c460*c463*k107 - c463*c483*k108 - c463*c486*k107, ...
c427*c463*k108 - c464*kd108 - c464*kd107 + c445*c463*k107, ...
c428*c463*k108 - c465*kd108 - c465*kd107 + c446*c463*k107, ...
c429*c463*k108 - c466*kd108 - c466*kd107 + c447*c463*k107, ...
c436*c463*k108 - c473*kd108 - c473*kd107 + c454*c463*k107, ...
c439*c463*k108 - c476*kd108 - c476*kd107 + c457*c463*k107, ...
c325*kd113 - c479*kd107 - c479*kd108 - c28*c479*k113 + c442*c463*k108 + c460*c463*k107, ...
c463*c483*k108 - c489*kd108 - c489*kd107 + c463*c486*k107, ...
c59*c486*k110 - c431*kd111 - c431*kd110 + c59*c488*k111, ...
c83*c486*k110 - c432*kd111 - c432*kd110 + c83*c488*k111, ...
c59*c430*k111 - c433*kd111 - c433*kd110 + c59*c445*k110, ...
c83*c430*k111 - c434*kd111 - c434*kd110 + c83*c445*k110, ...
c59*c409*k111 - c435*kd111 - c435*kd110 + c59*c446*k110, ...
c83*c409*k111 - c437*kd111 - c437*kd110 + c83*c446*k110, ...
c59*c410*k111 - c438*kd111 - c438*kd110 + c59*c447*k110, ...
c83*c410*k111 - c440*kd111 - c440*kd110 + c83*c447*k110, ...
c59*c454*k110 - c474*kd111 - c474*kd110 + c59*c490*k111, ...
c83*c454*k110 - c475*kd111 - c475*kd110 + c83*c490*k111, ...
c59*c457*k110 - c477*kd111 - c477*kd110 + c59*c491*k111, ...
c83*c457*k110 - c478*kd111 - c478*kd110 + c83*c491*k111, ...
c59*c460*k110 - c480*kd111 - c480*kd110 + c59*c487*k111, ...
c83*c460*k110 - c481*kd111 - c481*kd110 + c83*c487*k111, ...
c456*kd117 + c477*kd111 + c478*kd111 - c59*c491*k111 - c83*c491*k111 - c491*c521*k117, ...
c474*kd111 + c475*kd111 + c523*kd117 - c59*c490*k111 - c83*c490*k111 - c490*c521*k117, ...
c412*kd117 + c438*kd111 + c440*kd111 - c59*c410*k111 - c83*c410*k111 - c410*c521*k117, ...
c411*kd117 + c435*kd111 + c437*kd111 - c59*c409*k111 - c83*c409*k111 - c409*c521*k117, ...
c424*kd117 + c433*kd111 + c434*kd111 - c59*c430*k111 - c83*c430*k111 - c430*c521*k117, ...
c431*kd111 + c432*kd111 + c522*kd117 - c59*c488*k111 - c83*c488*k111 - c488*c521*k117, ...
c407*kd117 + c480*kd111 + c481*kd111 - c59*c487*k111 - c83*c487*k111 - c487*c521*k117, ...
c26*c104*k112 - c264*kd113 - c264*kd112 + c28*c104*k113, ...
c26*c261*k112 - c265*kd113 - c265*kd112 + c28*c261*k113, ...
c26*c262*k112 - c266*kd113 - c266*kd112 + c28*c262*k113, ...
c26*c263*k112 - c267*kd113 - c267*kd112 + c28*c263*k113, ...
c26*c324*k112 - c268*kd113 - c268*kd112 + c28*c324*k113, ...
c26*c405*k112 - c269*kd113 - c269*kd112 + c28*c405*k113, ...
c26*c408*k112 - c325*kd113 - c325*kd112 + c28*c479*k113, ...
c45*c497*k114 - c472*kd115 - c472*kd114 + c485*c497*k115, ...
c72*c497*k114 - c484*kd115 - c484*kd114 + c485*c497*k115, ...
c472*kd115 + c484*kd115 - 2*c485*c497*k115, ...
c60*stepfunc(time, 2659.99, 0, 2660, 0.0150356) - c520*kd116, ...
c407*kd117 + c407*kd118 + c411*kd117 + c411*kd118 + c412*kd117 + c412*kd118 + c424*kd117 + c424*kd118 + c456*kd117 + c456*kd118 + c522*kd117 + c522*kd118 + c523*kd117 + c523*kd118 - c409*c521*k117 - c410*c521*k117 - c430*c521*k117 - c445*c521*k118 - c446*c521*k118 - c447*c521*k118 - c454*c521*k118 - c457*c521*k118 - c460*c521*k118 - c486*c521*k118 - c487*c521*k117 - c488*c521*k117 - c490*c521*k117 - c491*c521*k117, ...
c486*c521*k118 - c522*kd118 - c522*kd117 + c488*c521*k117, ...
c454*c521*k118 - c523*kd118 - c523*kd117 + c490*c521*k117, ...
c409*c521*k117 - c411*kd118 - c411*kd117 + c446*c521*k118, ...
c410*c521*k117 - c412*kd118 - c412*kd117 + c447*c521*k118, ...
c457*c521*k118 - c456*kd118 - c456*kd117 + c491*c521*k117, ...
c430*c521*k117 - c424*kd118 - c424*kd117 + c445*c521*k118, ...
c460*c521*k118 - c407*kd118 - c407*kd117 + c487*c521*k117, ...
c355*kd120 - c142*kd119 + c516*kd120 - c2*c142*k120b - c141*c142*k120 + c140*c514*stepfunc(time, 1799.99, 0, 1800, 10000000), ...
c345*kd120 - c144*kd119 + c517*kd120 - c2*c144*k120b - c141*c144*k120 + c143*c514*stepfunc(time, 1799.99, 0, 1800, 10000000), ...
c422*kd120 + c519*kd120 - c6*c158*k120b - c155*c158*k120, ...
c525*kd97c + c524*stepfunc(time, 1799.99, 0, 1800, 1) - c105*c532*stepfunc(time, 1799.99, 0, 1800, 0.000000018704) - (c285*c532*kd97c)/KD_iressa_mt, ...
c526*stepfunc(time, 1799.99, 0, 1800, 0.0033) - c525*kd97c - c1*c525*stepfunc(time, 1799.99, 0, 1800, 10000000) + (c285*c532*kd97c)/KD_iressa_mt, ...
c527*kd2 + c528*kd2 + c553*kd2 + c554*kd2 - c526^2*k2 - c526*stepfunc(time, 1799.99, 0, 1800, 0.0033) - c3*c526*k2 - c499*c526*k2 - c526*c529*k2 + c1*c525*stepfunc(time, 1799.99, 0, 1800, 10000000), ...
c121*stepfunc(time, 1799.99, 0, 1800, 1) - c527*kd2 - c105*c527*stepfunc(time, 1799.99, 0, 1800, 0.000000018704) + c3*c526*k2, ...
c526^2*k2 - c528*kd2, ...
c530*stepfunc(time, 1799.99, 0, 1800, 0.00005) - c524*stepfunc(time, 1799.99, 0, 1800, 1) - c524*k6 + c529*stepfunc(time, 1799.99, 0, 1800, 0.0033) + c105*c532*stepfunc(time, 1799.99, 0, 1800, 0.000000018704) - c1*c524*stepfunc(time, 1799.99, 0, 1800, 10000000), ...
c550*kd2 + c551*kd2 + c552*kd2 + c553*kd2 - c529^2*k2 - c529*stepfunc(time, 1799.99, 0, 1800, 0.0033) - c3*c529*k2 - c499*c529*k2 - c526*c529*k2 + c1*c524*stepfunc(time, 1799.99, 0, 1800, 10000000), ...
c524*k6 + c10*kd10 - c530*stepfunc(time, 1799.99, 0, 1800, 0.00005) - c16*c530*k10b, ...
c105*c500*stepfunc(time, 1799.99, 0, 1800, 0.000000018704) - c115*stepfunc(time, 1799.99, 0, 1800, 1) - c115*kd123h + c5*c105*k123h, ...
c105*c527*stepfunc(time, 1799.99, 0, 1800, 0.000000018704) - c121*stepfunc(time, 1799.99, 0, 1800, 1) - c121*kd123h + c5*c105*k123h, ...
c555*stepfunc(time, 1799.99, 0, 1800, 1) - c550*kd2 - c105*c550*stepfunc(time, 1799.99, 0, 1800, 0.000000018704) + c3*c529*k2, ...
c556*stepfunc(time, 1799.99, 0, 1800, 1) - c551*kd2 - c105*c551*stepfunc(time, 1799.99, 0, 1800, 0.000000018704) + c499*c529*k2, ...
c529^2*k2 - c552*kd2 + c557*stepfunc(time, 1799.99, 0, 1800, 1) - c105*c552*stepfunc(time, 1799.99, 0, 1800, 0.000000018704), ...
c558*stepfunc(time, 1799.99, 0, 1800, 1) - c553*kd2 - c105*c553*stepfunc(time, 1799.99, 0, 1800, 0.000000018704) + c526*c529*k2, ...
c499*c526*k2 - c554*kd2, ...
c105*c550*stepfunc(time, 1799.99, 0, 1800, 0.000000018704) - c555*stepfunc(time, 1799.99, 0, 1800, 1) - c555*kd123 + c5*c105*k123, ...
c105*c551*stepfunc(time, 1799.99, 0, 1800, 0.000000018704) - c556*stepfunc(time, 1799.99, 0, 1800, 1) - c556*kd123h + c5*c105*k123h, ...
c105*c552*stepfunc(time, 1799.99, 0, 1800, 0.000000018704) - c557*stepfunc(time, 1799.99, 0, 1800, 1) - c557*kd123 + c5*c105*k123, ...
c105*c553*stepfunc(time, 1799.99, 0, 1800, 0.000000018704) - c558*stepfunc(time, 1799.99, 0, 1800, 1) - c558*kd123h + c5*c105*k123h];

%%
% INITIALIZATION

model.sym.x0 = [0, ...
0, ...
0, ...
0, ...
0, ...
0, ...
0, ...
0, ...
0, ...
0, ...
0, ...
0, ...
0, ...
462000, ...
0, ...
6230, ...
0, ...
794, ...
0, ...
0, ...
0, ...
0, ...
0, ...
0, ...
0, ...
0, ...
0, ...
0, ...
0, ...
0, ...
0, ...
0, ...
0, ...
0, ...
1200000000, ...
0, ...
0, ...
0, ...
0, ...
0, ...
0, ...
0, ...
0, ...
0, ...
0, ...
0, ...
0, ...
0, ...
0, ...
0, ...
0, ...
0, ...
0, ...
0, ...
0, ...
0, ...
0, ...
0, ...
0, ...
0, ...
0, ...
0, ...
0, ...
0, ...
0, ...
0, ...
0, ...
0, ...
0, ...
0, ...
0, ...
0, ...
1236601486306181/274877906944, ...
0, ...
0, ...
0, ...
0, ...
0, ...
0, ...
0, ...
0, ...
0, ...
0, ...
0, ...
0, ...
0, ...
0, ...
0, ...
0, ...
0, ...
0, ...
0, ...
0, ...
0, ...
0, ...
0, ...
0, ...
0, ...
0, ...
0, ...
0, ...
0, ...
0, ...
0, ...
0, ...
0, ...
0, ...
0, ...
0, ...
0, ...
0, ...
0, ...
0, ...
0, ...
0, ...
0, ...
0, ...
0, ...
0, ...
0, ...
0, ...
0, ...
0, ...
0, ...
0, ...
0, ...
0, ...
0, ...
0, ...
0, ...
0, ...
0, ...
0, ...
0, ...
0, ...
0, ...
0, ...
0, ...
0, ...
0, ...
0, ...
0, ...
0, ...
0, ...
0, ...
0, ...
0, ...
0, ...
0, ...
0, ...
0, ...
0, ...
0, ...
0, ...
0, ...
0, ...
0, ...
0, ...
0, ...
0, ...
0, ...
0, ...
0, ...
0, ...
0, ...
0, ...
0, ...
0, ...
0, ...
0, ...
0, ...
0, ...
0, ...
0, ...
0, ...
0, ...
0, ...
0, ...
0, ...
0, ...
0, ...
0, ...
0, ...
0, ...
0, ...
0, ...
0, ...
0, ...
0, ...
0, ...
0, ...
0, ...
0, ...
0, ...
0, ...
0, ...
0, ...
0, ...
0, ...
0, ...
0, ...
0, ...
0, ...
0, ...
0, ...
0, ...
0, ...
0, ...
0, ...
0, ...
0, ...
0, ...
0, ...
0, ...
0, ...
0, ...
0, ...
0, ...
0, ...
0, ...
0, ...
0, ...
0, ...
0, ...
0, ...
0, ...
0, ...
0, ...
0, ...
0, ...
0, ...
0, ...
0, ...
0, ...
0, ...
0, ...
0, ...
0, ...
0, ...
0, ...
0, ...
0, ...
0, ...
0, ...
0, ...
0, ...
0, ...
0, ...
0, ...
0, ...
0, ...
0, ...
0, ...
0, ...
0, ...
0, ...
0, ...
0, ...
0, ...
0, ...
0, ...
0, ...
0, ...
0, ...
0, ...
0, ...
0, ...
0, ...
0, ...
0, ...
0, ...
0, ...
0, ...
0, ...
0, ...
0, ...
534751, ...
0, ...
0, ...
0, ...
0, ...
0, ...
0, ...
0, ...
0, ...
0, ...
0, ...
0, ...
0, ...
5563133012360561/4398046511104, ...
0, ...
0, ...
0, ...
0, ...
0, ...
0, ...
0, ...
0, ...
0, ...
3992271744873267/68719476736, ...
0, ...
0, ...
0, ...
0, ...
1100000, ...
0, ...
0, ...
0, ...
0, ...
0, ...
0, ...
4888098843603763/68719476736, ...
0, ...
0, ...
0, ...
0, ...
0, ...
88914000, ...
50000, ...
0, ...
0, ...
0, ...
0, ...
3020000, ...
0, ...
0, ...
0, ...
0, ...
0, ...
0, ...
124480, ...
0, ...
0, ...
0, ...
0, ...
695000, ...
0, ...
0, ...
0, ...
0, ...
0, ...
0, ...
0, ...
0, ...
84351/5, ...
0, ...
0, ...
0, ...
0, ...
0, ...
0, ...
0, ...
0, ...
0, ...
0, ...
0, ...
0, ...
0, ...
0, ...
0, ...
0, ...
0, ...
0, ...
0, ...
0, ...
0, ...
0, ...
35565600, ...
0, ...
0, ...
0, ...
0, ...
0, ...
0, ...
0, ...
0, ...
0, ...
0, ...
0, ...
0, ...
0, ...
0, ...
0, ...
0, ...
0, ...
0, ...
0, ...
0, ...
0, ...
0, ...
0, ...
0, ...
0, ...
0, ...
905000, ...
0, ...
0, ...
0, ...
300416000, ...
0, ...
0, ...
0, ...
0, ...
450000, ...
0, ...
0, ...
70000, ...
0, ...
0, ...
0, ...
0, ...
0, ...
0, ...
0, ...
0, ...
1080000, ...
0, ...
0, ...
0, ...
0, ...
0, ...
0, ...
0, ...
608466986032169/274877906944, ...
393639, ...
0, ...
7710448984837325/137438953472, ...
0, ...
6519299934833869/68719476736, ...
0, ...
1000000, ...
0, ...
0, ...
0, ...
0, ...
0, ...
0, ...
0, ...
0, ...
0, ...
0, ...
0, ...
0, ...
0, ...
0, ...
0, ...
0, ...
0, ...
0, ...
0, ...
0, ...
0, ...
0, ...
0, ...
0, ...
0, ...
0, ...
0, ...
0, ...
0, ...
0, ...
0, ...
0, ...
0, ...
0, ...
0, ...
0, ...
0, ...
0, ...
0, ...
0, ...
0, ...
0, ...
0, ...
0, ...
0, ...
0, ...
0, ...
0, ...
0, ...
0, ...
0, ...
0, ...
0, ...
0, ...
0, ...
0, ...
0, ...
0, ...
0, ...
0, ...
0, ...
0, ...
0, ...
0, ...
0, ...
0, ...
0, ...
0, ...
0];

%% Observables

% [ERB_B1_P_tot; ERK_PP; AKT_PP]
model.sym.y = [(2*c5 + 2*c7 + 2*c8 + 2*c15 + 2*c17 + 2*c18 + 2*c19 + 2*c20 + 2*c21 + 2*c23 + 2*c25 + 2*c27 + 2*c29 + 2*c32 + 2*c33 + 2*c34 + 2*c35 + 2*c36 + 2*c37 + 2*c63 + 2*c64 + 2*c65 + 2*c66 + 2*c67 + 2*c68 + 2*c88 + 2*c89 + 2*c90 + 2*c91 + 2*c92 + 2*c93 + 2*c94 + 2*c95 + 2*c96 + 2*c97 + 2*c98 + 2*c99 + 2*c100 + 2*c104 + c130 + c131 + c132 + 2*c136 + c148 + c149 + c150 + c151 + c152 + c153 + c162 + c163 + c164 + c165 + c166 + c167 + c171 + c172 + c173 + c174 + c175 + c176 + c180 + c181 + c182 + c183 + c184 + c185 + c189 + c190 + c191 + c192 + c193 + c194 + c195 + c196 + c197 + c198 + c199 + c200 + c201 + c202 + c203 + c204 + c205 + c206 + c207 + c208 + c209 + c210 + c211 + c212 + c213 + c214 + c215 + c216 + c217 + c218 + c219 + c220 + c221 + c222 + c223 + c224 + c225 + c226 + c227 + c228 + c229 + c230 + c231 + c232 + c233 + c234 + c235 + c236 + c237 + c238 + c239 + c240 + c241 + c242 + c243 + c244 + c245 + c246 + c247 + c248 + c249 + c250 + c251 + c252 + c253 + c254 + c255 + c256 + c257 + c258 + c259 + c260 + c261 + c262 + c263 + 2*c264 + c265 + c266 + c267 + c281 + c282 + c409 + c410 + c411 + c412 + 2*c415 + c416 + 2*c419 + 2*c420 + c427 + c428 + c429 + 2*c431 + 2*c432 + c433 + c434 + c435 + c437 + c438 + c440 + c445 + c446 + c447 + 2*c448 + c449 + c450 + c451 + c464 + c465 + c466 + 2*c483 + 2*c486 + 2*c489)/EGFR_t, ...
(c59 + c61 + c83 + c84 + c95 + c96 + c97 + c98 + c101 + c102 + c431 + c432 + c433 + c434 + c435 + c437 + c438 + c440 + c474 + c475 + c477 + c478 + c480 + c481)/ERK_t, ...
(c472 + c484 + c497 + c498)/AKT_t];

end


function r = pow(x,y)
    r = x^y;
end


function r = power(x,y)
    r = x^y;
end


function r = stepfunc2(t,t_start,v_start,t_end,v_end)
    r = am_piecewise(v_start,am_lt(t,t_start),v_end,am_gt(t,t_end),(v_end-v_start)/(t_end-t_start)*(t-t_start)+v_start);
end


function r = stepfunc(t,t_start,v_start,t_end,v_end)
    r = am_piecewise(v_start,am_lt(t,t_start),am_piecewise(v_end,am_gt(t,t_end),(v_start+v_end)/2+(v_end-v_start)/2*sin(3.14/2*(2*t-t_end-t_start)/(t_end-t_start))));
end
```
