## Supplementary material for "Mechanistic description of spatial processes using integrative modelling of noise-corrupted imaging data": getData_ErbB_signaling.m

```
function [ DD ] = getData_ErbB_signaling()
% getData_erbb_signaling.m for examples/Chen2009
%
% getData_erbb_signaling.m provides measurement data for the parameter 
% estimation of the erbb_signaling example file. 
%
% Parameters:
%  void
%
% Return values:
%  D: struct, containing the time-points of measurement, the measured Data, 
%      the variance and the initial conditions


%% Write measured data into a table
% Cell line: A431

% pAkt
M = [...
    0	0	0;...
    2.5	0	0;...
    5	24	0.26;...
    7.5	34	0.59;...
    10	40	2.2;...
    15	50	2;...
    30	64	1.4;...
    45	52	3.4;...
    60	49	2.2;...
    1.2e+02	57	7.3;...
    ];

D(1).t = M(:,1)*60+1800; % minutes -> sec + preequi
D(1).condition = [5e-9,0,0,0];
D(1).Y(:,3) = M(:,2);
D(1).Sigma_Y(:,3) = M(:,3);

% pErbB1
M= [...
    0	0	0;...
    2.5	65	7.3;...
    5	72	6.7;...
    7.5	75	8.8;...
    10	77	4.6;...
    15	83	4.6;...
    30	87	2.4;...
    45	98	2.3;...
    60	87	13;...
    1.2e+02	86	13;...
    ];

D(1).Y(:,1) = M(:,2);
D(1).Sigma_Y(:,1) = M(:,3);

% pERK
M = [...
    0	0	0;...
    2.5	60	0.36;...
    5	88	1.8;...
    7.5	97	1.4;...
    10	98	1.1;...
    15	86	3;...
    30	65	2.7;...
    45	56	2.8;...
    60	50	3.6;...
    1.2e+02	46	1.4;...
    ];

D(1).Y(:,2) = M(:,2);
D(1).Sigma_Y(:,2) = M(:,3);

% pAkt
M = [...
    0	0	0;...
    2.5	45	4.6;...
    5	79	3.1;...
    7.5	86	4.5;...
    10	97	3.1;...
    15	91	2.8;...
    30	85	2.7;...
    45	72	2.5;...
    60	73	1.7;...
    1.2e+02	68	2.2;...
    ];

D(2).t = M(:,1)*60+1800; % minutes -> sec + preequi
D(2).condition = [0,0,0,5e-9];
D(2).Y(:,3) = M(:,2);
D(2).Sigma_Y(:,3) = M(:,3);

% pErbB1
M= [...
    0	0	0;...
    2.5	0	0;...
    5	0	0;...
    7.5	0	0;...
    10	0	0;...
    15	0	0;...
    30	0	0;...
    45	0	0;...
    60	0	0;...
    1.2e+02	0	0;...
    ];

D(2).Y(:,1) = M(:,2);
D(2).Sigma_Y(:,1) = M(:,3);

% pERK
M = [...
    0	0	0;...
    2.5	9.7	3.3;...
    5	46	6.7;...
    7.5	58	1.8;...
    10	54	3.9;...
    15	26	4.2;...
    30	27	2.6;...
    45	24	1.1;...
    60	23	0.84;...
    1.2e+02	17	2.2;...
    ];

D(2).Y(:,2) = M(:,2);
D(2).Sigma_Y(:,2) = M(:,3);

% pAkt
M = [...
    0	0	0;...
    2.5	1.7	0.88;...
    5	44	11;...
    7.5	43	5.5;...
    10	50	3;...
    15	45	4.4;...
    30	35	5;...
    45	40	14;...
    60	28	9.6;...
    1.2e+02	28	5.5;...
    ];

D(3).t = M(:,1)*60+1800; % minutes -> sec + preequi
D(3).condition = [1e-11,0,0,0];
D(3).Y(:,3) = M(:,2);
D(3).Sigma_Y(:,3) = M(:,3);

% pErbB1
M= [...
    0	0	0;...
    2.5	0	0;...
    5	0	0;...
    7.5	0	0;...
    10	0	0;...
    15	0	0;...
    30	0	0;...
    45	0	0;...
    60	0	0;...
    1.2e+02	0	0;...
    ];

D(3).Y(:,1) = M(:,2);
D(3).Sigma_Y(:,1) = M(:,3);

% pERK
M = [...
    0	0	0;...
    2.5	8.8	2.2;...
    5	43	8.8;...
    7.5	46	5.7;...
    10	40	3.2;...
    15	26	2.4;...
    30	27	0.28;...
    45	18	3.5;...
    60	11	2.4;...
    1.2e+02	2.7	0.65;...
    ];

D(3).Y(:,2) = M(:,2);
D(3).Sigma_Y(:,2) = M(:,3);

% pAkt
M = [...
    0	0	0;...
    2.5	1.7	0.88;...
    5	44	11;...
    7.5	43	5.5;...
    10	50	3;...
    15	45	4.4;...
    30	35	5;...
    45	40	14;...
    60	28	9.6;...
    1.2e+02	28	5.5;...
    ];

D(4).t = M(:,1)*60+1800; % minutes -> sec + preequi
D(4).condition = [0,0,0,1e-10];
D(4).Y(:,3) = M(:,2);
D(4).Sigma_Y(:,3) = M(:,3);

% pErbB1
M= [...
    0	0	0;...
    2.5	0	0;...
    5	0	0;...
    7.5	0	0;...
    10	0	0;...
    15	0	0;...
    30	0	0;...
    45	0	0;...
    60	0	0;...
    1.2e+02	0	0;...
    ];

D(4).Y(:,1) = M(:,2);
D(4).Sigma_Y(:,1) = M(:,3);

% pERK
M = [...
    0	0	0;...
    2.5	8.8	2.2;...
    5	43	8.8;...
    7.5	46	5.7;...
    10	40	3.2;...
    15	26	2.4;...
    30	27	0.28;...
    45	18	3.5;...
    60	11	2.4;...
    1.2e+02	2.7	0.65;...
    ];

D(4).Y(:,2) = M(:,2);
D(4).Sigma_Y(:,2) = M(:,3);

for iD = 1:length(D)
    D(iD).Sigma_Y = D(iD).Sigma_Y + 0.1*ones(size(D(iD).Sigma_Y));
    DD(iD) = amidata(D(iD));
end

end
```
