## Supplementary material for "Mechanistic description of spatial processes using integrative modelling of noise-corrupted imaging data": logLikelihoodErbBSignaling.m

```
function varargout = logLikelihoodErbBSignaling(theta, D)
% Objective function for examples/erbb_signaling
%
% logLikelihoodErbBSignaling.m provides the log-likelihood and its gradient 
% for the model defined in erbb_signaling_pesto_syms.m
% 
% USAGE:
% [llh] = logLikelihoodErbBSignaling(theta, amiData)
% [llh, sllh] = logLikelihoodErbBSignaling(theta, amiData)
%
% Parameters:
%  theta: Model parameters 
%  amiData: an amidata object for the AMICI solver
%
% Return values:
%   varargout:
%     llh: Log-Likelihood, only the LogLikelihood will be returned, no 
%         sensitivity analysis is performed
%     sllh: Gradient of llh, The LogLikelihood and its gradient will be 
%         returned, first order adjoint sensitivity analysis is performed


%% Model Definition
% The ODE model is set up using the AMICI toolbox. To access the AMICI
% model setup, see erbb_signaling_pesto_syms.m
% For a detailed description for the biological model see the referenced
% papers on the ErbB signaling pathways by Chen et al.

%% AMICI
% Setting the options for the AMICI solver
optionsAmici = amioption;
optionsAmici.atol = 1e-8;
optionsAmici.maxsteps = 2e5;
optionsAmici.interpType = 2;

if(nargout > 1)
    optionsAmici.sensi = 1;
    optionsAmici.sensi_meth = 'adjoint';
    sol = simulate_erbb_pesto(D.t, theta, D.condition, D, optionsAmici);
    if(sol.status < 0)
        error('integration error');
    else
        varargout{1} = sol.llh;
        varargout{2} = sol.sllh;
    end
else
    optionsAmici.sensi = 0;
    sol = simulate_erbb_pesto(D.t, theta, D.condition, D, optionsAmici);
    if(sol.status<0)
        error('integration error');
    else
        varargout{1} = sol.llh;
    end
end
    
end
```
