## Supplementary material for "Mechanistic description of spatial processes using integrative modelling of noise-corrupted imaging data": mainErbBSignaling.m

```
% Main file of the erbb_signaling example
%
% Demonstrates the use of:
% * getMultiStarts()
%
% This example is a rather big biological model. It is included here to 
% show the ability of PESTO to handle ODE-based models with some hundred 
% state variables and some hundred parameters. This model is taken from the
% paper "Input-output behavior of ErbB signaling pathways as revealed by a 
% mass action model trained against dynamic data", by Chen et al., in 2009,
% PubMed, vol.5, no.239 (see https://www.ncbi.nlm.nih.gov/pubmed/19156131).
%
% The data used is measurement data provided in the publication.
%
% This file performs a multistart local optimization based on measured data 
% from the referenced papers, demonstrating the use of getMultiStarts().


%% Preliminary
% Clean up

clear all;
close all;
clc;

TextSizes.DefaultAxesFontSize = 14;
TextSizes.DefaultTextFontSize = 18;
set(0,TextSizes);

% Seed random number generator
rng(0);

%% Model Definition
% The ODE model is set up using the AMICI toolbox. To access the AMICI
% model setup, see erbb_signaling_pesto_syms.m
% For a detailed description of the biological model see the referenced
% papers on the ErbB signaling pathways by Chen et al.

[exdir,~,~] = fileparts(which('mainErbBSignaling.m'));
try
    amiwrap('erbb_pesto', 'erbb_signaling_pesto_syms', exdir);
catch ME
    warning('There was a problem with the AMICI toolbox (available at https://github.com/ICB-DCM/AMICI), which is needed to run this example file. The original error message was:');
    rethrow(ME);
end

%% Data
% Experimental data is read out and written to an AMICI-data object which 
% is used for the ODE integration

load('erbb_signaling_pnom.mat');
D = getData_ErbB_signaling();

%% Generation of the structs and options for PESTO
% The structs and the PestoOptions object, which are necessary for the 
% PESTO routines to work are created and set to convenient values

% Set the best value of theta
theta = log10(pnom);
theta(isinf(theta)) = log10(eps);

% Write the parameters struct
parameters.min = theta - 2;
parameters.max = theta + 3;
parameters.number = length(theta);

% Set the PESTO-options
optionsMultistart           = PestoOptions();
optionsMultistart.n_starts  = 5;
optionsMultistart.comp_type = 'sequential';
optionsMultistart.mode      = 'text';
optionsMultistart.localOptimizerOptions = optimset('Algorithm','interior-point',...
    'GradObj', 'on',...
    'Display', 'iter', ...
    'MaxIter', 1000,...
    'TolFun', 0,...
    'TolX', 1e-10,...
    'MaxFunEvals', 2000);

% Set the objective function
objectiveFunction = @(theta) logLikelihoodErbBSignaling(theta, D(1));

%% Perform Multistart optimization
% A multi-start local optimization is performed within the bound defined in
% parameters.min and .max in order to infer the unknown parameters from 
% measurement data.

% REMARK: The problem is rather intermediate to large-scale, each 
% evaluation of the objective function takes a while, parameter space is
% high dimensional. Hence, optimization takes a while (up to some hours)
% for this example.
fprintf('\n Perform optimization...');
parameters_adjoint = getMultiStarts(parameters, objectiveFunction, optionsMultistart);
```
