## Supplementary material for "Mechanistic description of spatial processes using integrative modelling of noise-corrupted imaging data": define_Gauss_LLH.m

```
% Definition script for examples/GaussExample
%
% define_Gauss_LLH.m creates the log-posterior for Gaussian mixture model


%% Definition of the Gaussian mixture

% % Hard version
% rng(15);
% angle = pi/2+2*pi/8;
% dimi = 18;
% scale = blkdiag(1e0*[250,0;0,1],diag(ones(1,dimi)));

% Easy Version
rng(3);
angle = 2*pi/8;
dimi = 1;
scale = blkdiag(1e0*[50,0;0,1],diag(ones(1,dimi)));

% Setting the dimension
n = 2;

% Setting of mean, rotation and covariance
mu = [rand(2,n)*50; repmat(25,dimi,n) ];
rot1 = [cos(angle),-sin(angle);sin(angle),cos(angle)];
rot1 = blkdiag(rot1,diag( ones(length(scale) - length(rot1), 1) ));
cov1 = rot1'*scale*rot1;
sig = cat(3,repmat(cov1,1,1,n));
rng('shuffle');

% Definition of log-posterior function
logP = @(theta) simulateGaussLLH(theta, mu, sig);
```
