## Supplementary material for "Mechanistic description of spatial processes using integrative modelling of noise-corrupted imaging data": mainExampleGauss.m

```
% Main file of the Gaussian mixture example
%
% Demonstrates the use of:
% * getParameterSamples()
%
% Demonstrates furthermore:
% * how to use different sampling methods without optimization
% * how to analyze the results from sampling routines
% * how to use advanced plotting tools from PESTO
%
% The example problem is a mixture of Gaussian modes in the first two
% dimensions and a single mode in any other dimension. The dimensionality,
% angle, scaling and position of modes can be altered in defineGaussLLH.m.
% This example was written by B. Ballnus.
% 
% Single-chain and multi-chain Monte-Carlo sampling is performed by 
% getParameterSamples() and with different settings (commented code 
% version) and plotted in 1D and 2D.


%% Preliminary
clear all
close all
clc


%% Problem initialization

% Seed random number generator
rng(0);

% Settings for this example
define_Gauss_LLH();
gaussDimension = 2 + dimi;


% Set required sampling options for Parallel Tempering
parameters.number = gaussDimension;
parameters.min    = -3*ones(dimi+2,1);
parameters.max    = 50*ones(dimi+2,1);
parameters.name   = {};
for i = 1 : dimi + 2
   parameters.name{end+1} = ['\theta_' num2str(i)];
end

% Sampling Options
options                     = PestoSamplingOptions();
options.objOutNumber        = 1;
options.nIterations         = 1e5;

% Using PT
options.samplingAlgorithm   = 'PT';
options.PT.nTemps           = 3;
options.PT.exponentT        = 4;    
options.PT.alpha            = 0.51;
options.PT.temperatureAlpha = 0.51;
options.PT.memoryLength     = 1;
options.PT.regFactor        = 1e-4;
options.PT.temperatureAdaptionScheme = 'Vousden16'; % 'Lacki15'; 

options.theta0              = repmat([0,20,repmat(25,1,dimi)]',1,options.PT.nTemps); 
options.theta0(:,1:2:end)   = repmat([40,5,repmat(25,1,dimi)]',1,ceil(options.PT.nTemps/2));
options.sigma0              = 1e1*blkdiag([50,0;0,1],diag(ones(1,dimi)));

% Using DRAM
% options.samplingAlgorithm     = 'DRAM';
% options.DRAM.nTry             = 5;
% options.DRAM.verbosityMode    = 'debug';    
% options.DRAM.adaptionInterval = 1;
% options.DRAM.regFactor        = 1e-4;
% options.theta0                = [0,20,repmat(25,1,dimi)]'; 
% options.sigma0                = 1e1*blkdiag([50,0;0,1],diag(ones(1,dimi)));

% Using MALA
% options.samplingAlgorithm     = 'MALA';
% options.MALA.regFactor        = 1e-4;
% options.theta0                = [0,20,repmat(25,1,dimi)]'; 
% options.sigma0                = 1e1*blkdiag([50,0;0,1],diag(ones(1,dimi)));

% Using PHS
% options.samplingAlgorithm     = 'PHS';
% options.PHS.nChains           = 3;
% options.PHS.alpha             = 0.51;
% options.PHS.memoryLength      = 1;
% options.PHS.regFactor         = 1e-4;
% options.PHS.trainingTime      = ceil(options.nIterations / 5);
% options.theta0                = [0,20,repmat(25,1,dimi)]'; 
% options.sigma0                = 1e1*blkdiag([50,0;0,1],diag(ones(1,dimi)));

optionsPesto = PestoOptions();
optionsPesto.MCMC = options;

% Perform the parameter estimation via sampling
parameters = getParameterSamples(parameters, logP, optionsPesto);

% Additional visualization
figure('Name', 'Chain analysis and theoretical vs true sampling distribution');

% Visualize the mixing properties and the exploration of the chain
subplot(2,1,1);
plot(squeeze(parameters.S.par(:,:,1))');

% Visualize the samples...
subplot(2,1,2); 
plot(squeeze(parameters.S.par(1,:,1))',squeeze(parameters.S.par(2,:,1))','.'); 

% ...and their theoretical distribution
hold on;
plot_Gauss_LH();
hold off;

% Use a diagnosis tool to see, how plotting worked (another look at 
% burn-in etc.)
plotMCMCdiagnosis(parameters, 'parameters');
plotMCMCdiagnosis(parameters, 'log-posterior');


% The kernel density estimates for all three chains/temepratures are
% plotted using the accoring PESTO routines
samplingPlottingOpt = PestoPlottingOptions();

% Density estimate
samplingPlottingOpt.S.plot_type = 2; 

% Ind set to 3 to show all temperatures (multi chain only)
samplingPlottingOpt.S.ind = 3;

% Set colors for different chains
samplingPlottingOpt.S.col = [0.4,0.4,0.4; 0.6,0.6,0.6; 0.8,0.8,0.8];

% Plotting
samplingPlottingOpt.S.sp_col = samplingPlottingOpt.S.col;
plotParameterUncertainty(parameters,'1D',[],[],samplingPlottingOpt);

% Density estimate in 2D again
samplingPlottingOpt.S.plot_type = 2; 

% Ind set to 1 for only one temperature this time
samplingPlottingOpt.S.ind = 1;

% Set colors for different chains
samplingPlottingOpt.S.col = [0.4,0.4,0.4];

% Plotting
samplingPlottingOpt.S.sp_col = samplingPlottingOpt.S.col;
plotParameterUncertainty(parameters,'2D',[],[],samplingPlottingOpt);
```
