## Supplementary material for "Mechanistic description of spatial processes using integrative modelling of noise-corrupted imaging data": plot_Gauss_LH.m

```
% Additional plotting script for examples/GaussExample
%
% plot_Gauss_llh.m computes and plots the gound truth for the density of 
% the model output in examples/GaussExample


%% Preliminaries
% rng(5);
% mu = rand(2,20)*10;
% sigma = 0.05*ones(1,20) + (1-0.05) * rand(1,20);
% sigma = 2.5*ones(1,20);
% rng('shuffle');

%% Likelihood
% Seeting up a grid for plotting
x = linspace(-3,50,400);
y = linspace(-3,50,400);
[X,Y] = meshgrid(x,y);

% Log-Likelihood function
target = @(p1,p2) (logP([p1; p2; 25 * ones(dimi,1)]));
Z = arrayfun(target,X,Y);

%% Plotting
% Plotting and labeling
contour(X,Y,exp(Z),'linewidth',1.5);
xlabel('X_1');
ylabel('X_2');
```
