## Supplementary material for "Mechanistic description of spatial processes using integrative modelling of noise-corrupted imaging data": plot_gaussian_ellipsoid.m

```
function h = plot_gaussian_ellipsoid(m, C, sdwidth, npts, axh)
% PLOT_GAUSSIAN_ELLIPSOIDS plots 2-d and 3-d Gaussian distributions
%
% H = PLOT_GAUSSIAN_ELLIPSOIDS(M, C) plots the distribution specified by 
%  mean M and covariance C. The distribution is plotted as an ellipse (in 
%  2-d) or an ellipsoid (in 3-d).  By default, the distributions are 
%  plotted in the current axes. H is the graphics handle to the plotted 
%  ellipse or ellipsoid.
%
% PLOT_GAUSSIAN_ELLIPSOIDS(M, C, SD) uses SD as the standard deviation 
%  along the major and minor axes (larger SD => larger ellipse). By 
%  default, SD = 1. Note: 
%  * For 2-d distributions, SD=1.0 and SD=2.0 cover ~ 39% and 86% 
%     of the total probability mass, respectively. 
%  * For 3-d distributions, SD=1.0 and SD=2.0 cover ~ 19% and 73%
%     of the total probability mass, respectively.
%  
% PLOT_GAUSSIAN_ELLIPSOIDS(M, C, SD, NPTS) plots the ellipse or 
%  ellipsoid with a resolution of NPTS (ellipsoids are generated 
%  on an NPTS x NPTS mesh; see SPHERE for more details). By
%  default, NPTS = 50 for ellipses, and 20 for ellipsoids.
%
% PLOT_GAUSSIAN_ELLIPSOIDS(M, C, SD, NPTS, AX) adds the plot to the
%  axes specified by the axis handle AX.
%
% Examples: 
% -------------------------------------------
%  % Plot three 2-d Gaussians
%  figure; 
%  h1 = plot_gaussian_ellipsoid([1 1], [1 0.5; 0.5 1]);
%  h2 = plot_gaussian_ellipsoid([2 1.5], [1 -0.7; -0.7 1]);
%  h3 = plot_gaussian_ellipsoid([0 0], [1 0; 0 1]);
%  set(h2,'color','r'); 
%  set(h3,'color','g');
% 
%  % "Contour map" of a 2-d Gaussian
%  figure;
%  for sd = [0.3:0.4:4],
%    h = plot_gaussian_ellipsoid([0 0], [1 0.8; 0.8 1], sd);
%  end
%
%  % Plot three 3-d Gaussians
%  figure;
%  h1 = plot_gaussian_ellipsoid([1 1  0], [1 0.5 0.2; 0.5 1 0.4; 0.2 0.4 1]);
%  h2 = plot_gaussian_ellipsoid([1.5 1 .5], [1 -0.7 0.6; -0.7 1 0; 0.6 0 1]);
%  h3 = plot_gaussian_ellipsoid([1 2 2], [0.5 0 0; 0 0.5 0; 0 0 0.5]);
%  set(h2,'facealpha',0.6);
%  view(129,36); set(gca,'proj','perspective'); grid on; 
%  grid on; axis equal; axis tight;
% -------------------------------------------
% 
%  Gautam Vallabha, Sep-23-2007,

%  Revision 1.0, Sep-23-2007
%    - File created
%  Revision 1.1, 26-Sep-2007
%    - NARGOUT==0 check added.
%    - Help added on NPTS for ellipsoids

if ~exist('sdwidth', 'var'), sdwidth = 1; end
if ~exist('npts', 'var'), npts = []; end
if ~exist('axh', 'var'), axh = gca; end

if numel(m) ~= length(m), 
    error('M must be a vector'); 
end
if ~( all(numel(m) == size(C)) )
    error('Dimensionality of M and C must match');
end
if ~(isscalar(axh) && ishandle(axh) && strcmp(get(axh,'type'), 'axes'))
    error('Invalid axes handle');
end

set(axh, 'nextplot', 'add');

switch numel(m)
   case 2, h=show2d(m(:),C,sdwidth,npts,axh);
   case 3, h=show3d(m(:),C,sdwidth,npts,axh);
   otherwise
      error('Unsupported dimensionality');
end

if nargout==0,
    clear h;
end

%-----------------------------
function h = show2d(means, C, sdwidth, npts, axh)
if isempty(npts), npts=50; end
% plot the gaussian fits
tt=linspace(0,2*pi,npts)';
x = cos(tt); y=sin(tt);
ap = [x(:) y(:)]';
[v,d]=eig(C); 
d = sdwidth * sqrt(d); % convert variance to sdwidth*sd
bp = (v*d*ap) + repmat(means, 1, size(ap,2)); 
h = plot(bp(1,:), bp(2,:), '-', 'parent', axh, 'linewidth', 1.5, 'color', 'black');

%-----------------------------
function h = show3d(means, C, sdwidth, npts, axh)
if isempty(npts), npts=20; end
[x,y,z] = sphere(npts);
ap = [x(:) y(:) z(:)]';
[v,d]=eig(C); 
if any(d(:) < 0)
   fprintf('warning: negative eigenvalues\n');
   d = max(d,0);
end
d = sdwidth * sqrt(d); % convert variance to sdwidth*sd
bp = (v*d*ap) + repmat(means, 1, size(ap,2)); 
xp = reshape(bp(1,:), size(x));
yp = reshape(bp(2,:), size(y));
zp = reshape(bp(3,:), size(z));
h = surf(axh, xp,yp,zp);
```
