## Supplementary material for "Mechanistic description of spatial processes using integrative modelling of noise-corrupted imaging data": simulateGaussLLH.m

```
% Simulation function for examples/GaussExample
%
% simulateGauss.m computes a log-likelihood which is constructed from a
% Gaussian mixture model. The code is base on papers from Liang & Wong 
% (2001) and Lacki et al. (2015)
%
% Parameters:
%  par: Model parameters 
%  mu: mean value of the model
%  sigma: standard deviation of the model
%
% Return values:
%  llh: double, value of log-likelihood


function [ llh ] = simulateGaussLLH( par, mu, sigma )

    % Dimension of the model
    n = size(mu,2);
    if size(par,1) == 1
        par = par';
    end

    % Computing the log-likelihood
    llh = 0;
    for j = 1:n
        llh = llh + 1/(sqrt(2*pi)^2*sqrt(det(sigma(:,:,j)))) * ...
                exp(-0.5 * (par-mu(:,j))' / sigma(:,:,j) * (par-mu(:,j)));
    end
    llh = log(llh);
    
end
```
