## Supplementary material for "Mechanistic description of spatial processes using integrative modelling of noise-corrupted imaging data": jakstat_pesto_syms.m

```
function [model] = jakstat_pesto_syms()
% jakstat_pesto_syms for examples/jakstat_signaling
%
% creates an amimodel-object for the AMICI solver
%
% Parameters: 
% 
% Return values:
%  model: amimodel object


%% States
syms STAT pSTAT pSTAT_pSTAT npSTAT_npSTAT nSTAT1 nSTAT2 nSTAT3 nSTAT4 nSTAT5

model.sym.x = [
    STAT, pSTAT, pSTAT_pSTAT, npSTAT_npSTAT, nSTAT1, nSTAT2, nSTAT3, nSTAT4, nSTAT5 ...
    ];

%% Parameters
syms p1 p2 p3 p4 init_STAT Omega_cyt Omega_nuc sp1 sp2 sp3 sp4 sp5 offset_tSTAT offset_pSTAT scale_tSTAT scale_pSTAT sigma_pSTAT sigma_tSTAT sigma_pEpoR

model.sym.p = [p1,p2,p3,p4,init_STAT,sp1,sp2,sp3,sp4,sp5,offset_tSTAT,offset_pSTAT,scale_tSTAT,scale_pSTAT,sigma_pSTAT,sigma_tSTAT,sigma_pEpoR];
model.param = 'log10';
model.sym.k = [Omega_cyt,Omega_nuc];

%% Inputs
syms t

u(1) = am_spline_pos(t, 5, 0.0, sp1, 5.0, sp2, 10.0, sp3, 20.0, sp4, 60.0, sp5, 0, 0.0);

%% Right hand side of the ODE
model.sym.xdot = sym(zeros(size(model.sym.x)));

model.sym.xdot(1) = (Omega_nuc*p4*nSTAT5 - Omega_cyt*STAT*p1*u(1))/Omega_cyt;
model.sym.xdot(2) = STAT*p1*u(1) - 2*p2*pSTAT^2;
model.sym.xdot(3) = p2*pSTAT^2 - p3*pSTAT_pSTAT;
model.sym.xdot(4) = -(Omega_nuc*p4*npSTAT_npSTAT - Omega_cyt*p3*pSTAT_pSTAT)/Omega_nuc;
model.sym.xdot(5) = -p4*(nSTAT1 - 2*npSTAT_npSTAT);
model.sym.xdot(6) = p4*(nSTAT1 - nSTAT2);
model.sym.xdot(7) = p4*(nSTAT2 - nSTAT3);
model.sym.xdot(8) = p4*(nSTAT3 - nSTAT4);
model.sym.xdot(9) = p4*(nSTAT4 - nSTAT5);

%% Initial concentrations
model.sym.x0 = sym(zeros(size(model.sym.x)));

model.sym.x0(1) = init_STAT;

%% Observables
model.sym.y = sym(zeros(3,1));

model.sym.y(1) = offset_pSTAT + scale_pSTAT/init_STAT*(pSTAT + 2*pSTAT_pSTAT);
model.sym.y(2) = offset_tSTAT + scale_tSTAT/init_STAT*(STAT + pSTAT + 2*(pSTAT_pSTAT));
model.sym.y(3) = u(1);

%% Variances
model.sym.sigma_y = sym(size(model.sym.y));

model.sym.sigma_y(1) = sigma_pSTAT;
model.sym.sigma_y(2) = sigma_tSTAT;
model.sym.sigma_y(3) = sigma_pEpoR;
    
end
```
