## Supplementary material for "Mechanistic description of spatial processes using integrative modelling of noise-corrupted imaging data": logLikelihoodJakstat.m

```
function varargout = logLikelihoodJakstat(theta, amiData)
% Objective function for examples/jakstat_signaling
%
% logLikelihoodJakstat.m provides the log-likelihood, its gradient and an 
% the Hessian matrix for the model for the JakStat signaling pathway as
% defined in jakstat_pesto_syms.m
% 
% USAGE:
% [llh] = getParameterProfiles(theta, amiData)
% [llh, sllh] = getParameterProfiles(theta, amiData)
% [llh, sllh, s2llh] = getParameterProfiles(theta, amiData)
%
% Parameters:
%  theta: Model parameters 
%  amiData: an amidata object for the AMICI solver
%
% Return values:
%   varargout:
%     llh: Log-Likelihood, only the LogLikelihood will be returned, no 
%         sensitivity analysis is performed
%     sllh: Gradient of llh, The LogLikelihood and its gradient will be 
%         returned, first order adjoint sensitivity analysis is performed
%     s2llh: Hessian of llh, The LogLikelihood, its gradient and the 
%         Hessian matrix will be returned, second order adjoint sensitivity 
%         analysis is performed


%% Model Definition
% The ODE model is set up using the AMICI toolbox. To access the AMICI
% model setup, see jakstat_pesto_syms.m
% For a detailed description for the biological model see the referenced
% papers on the JakStat signaling pathway by Swameye et al. and Schelker et
% al.

    %% AMICI
    % Setting the options for the AMICI solver

    amiOptions.rtol = 1e-12;
    amiOptions.atol = 1e-14;
    amiOptions.sensi_meth = 'adjoint';

    %% Objective Function Evaluation
    % Assignment
    if (nargout == 1)
        amiOptions.sensi = 0;
        sol = simulate_jakstat_pesto([], theta, amiData.condition, amiData, amiOptions);
        varargout{1} = sol.llh;  
    elseif (nargout == 2)
        amiOptions.sensi = 1;
        sol = simulate_jakstat_pesto([], theta, amiData.condition, amiData, amiOptions);
        varargout{1} = sol.llh;
        varargout{2} = sol.sllh;
    elseif (nargout == 3)
        amiOptions.sensi = 2;
        sol = simulate_jakstat_pesto([], theta, amiData.condition, amiData, amiOptions);
        varargout{1} = sol.llh;
        varargout{2} = sol.sllh;
        varargout{3} = sol.s2llh;
    end
    
end
```
