## Supplementary material for "Mechanistic description of spatial processes using integrative modelling of noise-corrupted imaging data": logLikelihoodJakstatLsqnonlin.m

```
function varargout = logLikelihoodJakstatLsqnonlin(theta, amiData)
% Objective function for examples/jakstat_signaling
%
% logLikelihoodJakstat.m provides the log-likelihood, its gradient and an 
% the Hessian matrix for the model for the JakStat signaling pathway as
% defined in jakstat_pesto_syms.m
% 
% USAGE:
% [llh] = getParameterProfiles(theta, amiData)
% [llh, sllh] = getParameterProfiles(theta, amiData)
% [llh, sllh, s2llh] = getParameterProfiles(theta, amiData)
%
% Parameters:
%  theta: Model parameters 
%  amiData: an amidata object for the AMICI solver
%
% Return values:
%   varargout:
%     llh: Log-Likelihood, only the LogLikelihood will be returned, no 
%         sensitivity analysis is performed
%     sllh: Gradient of llh, The LogLikelihood and its gradient will be 
%         returned, first order adjoint sensitivity analysis is performed
%     s2llh: Hessian of llh, The LogLikelihood, its gradient and the 
%         Hessian matrix will be returned, second order adjoint sensitivity 
%         analysis is performed


%% Model Definition
% The ODE model is set up using the AMICI toolbox. To access the AMICI
% model setup, see jakstat_pesto_syms.m
% For a detailed description for the biological model see the referenced
% papers on the JakStat signaling pathway by Swameye et al. and Schelker et
% al.

    %% AMICI
    % Setting the options for the AMICI solver

    amiOptions = amioption();
    amiOptions.rtol = 1e-9;
    amiOptions.atol = 1e-12;
    amiOptions.sensi_meth = 'forward';
    
    if (nargout == 2)
        amiOptions.sensi = 1;
    else
        amiOptions.sensi = 0;
    end
    sol = simulate_jakstat_pesto([], theta, amiData.condition, amiData, amiOptions);
    
    % Set residuals
    res = (sol.y(:) - amiData.Y(:)) ./ sol.sigmay(:);
    res_sigma = sqrt(log(2*pi*sol.sigmay(:).^2) - log(2*pi*10^(-5)^2));
    res([1 48]) = 0;
    res_sigma([1 48]) = 0;
    varargout{1} = [res; res_sigma];
    
    if (nargout == 2)
        sres1 = ((1 ./ sol.sigmay(:)) * ones(1,17)) .* reshape(sol.sy, 16*3, 17);
        sres2 = (((sol.y(:) - amiData.Y(:)) ./ sol.sigmay(:).^2) * ones(1,17)) .* reshape(sol.ssigmay, 16*3, 17);
        sres3 = ((1 ./ (res_sigma .* sol.sigmay(:))) * ones(1,17)) .* reshape(sol.ssigmay, 16*3, 17);
        sres = [sres1 - sres2; sres3];
        sres([1 48 49 96], :) = 0;
        varargout{2} = sres;
    elseif (nargout == 3)
        varargout{2} = [];
        varargout{3} = sol.llh;
    end
    
end
```
