## Supplementary material for "Mechanistic description of spatial processes using integrative modelling of noise-corrupted imaging data": mainJakstatSignaling.m

```
% Main file of the JakStat signaling example
%
% Demonstrates the use of:
% * getMultiStarts()
% * getParameterProfiles()
%
% Demostrates furhtermore
% * how to implement a user-supplied guess for intial parameters
% * that non-evaluable points in parameter space can occur
% * how different optimization methods perform (multi-start local, hybrid,
%   global) -> Not necessary to perform computations, data-sheet is in this
%   folder and can be loaded (comparison_optimization_methods.mat)
% * how providing Hessians can improve optimization
% * how to use profile calculation in hybrid mode on a complex example
% * how to use parallelization
%
% This example provides a model for the JakStat signaling pathway with an
% time resolved input of the drug EPO. The model has been taken from the
% papers "Identification of nucleocytoplasmic cycling as a remote sensor in 
% cellular signaling by databased modeling" by Swameye et al. in 2003, 
% PNAS, vol.100, no.3 (see http://www.pnas.org/content/100/3/1028.long) and
% "Comprehensive estimation of input signals and dynamics in biochemical 
% reaction networks" by Schelker et al. in 2012, Bioinformatics, vol.28 
% (see http://bioinformatics.oxfordjournals.org/content/28/18/i529.full).
%
% The data used is measurement data provided in the publications.
%
% This file performs a multistart local optimization based on measured data 
% from the referenced papers, demonstrating the use of getMultiStarts().


%% Preliminary

clear;
close all;
clc;

TextSizes.DefaultAxesFontSize = 14;
TextSizes.DefaultTextFontSize = 18;
set(0,TextSizes);

% Seed random number generator
rng(0);

%% Model Definition
% The ODE model is set up using the AMICI toolbox. To access the AMICI
% model setup, see jakstat_pesto_syms.m
% For a detailed description for the biological model see the referenced
% papers on the JakStat signaling pathway by Swameye et al. and Schelker et
% al.

[exdir,~,~]=fileparts(which('mainJakstatSignaling.m'));
try
    amiwrap('jakstat_pesto','jakstat_pesto_syms', exdir);
catch ME
    warning('There was a problem with the AMICI toolbox (available at https://github.com/ICB-DCM/AMICI), which is needed to run this example file. The original error message was:');
    rethrow(ME);
end

%% Data
% Experimental data is read out from an .xls-file and written to an AMICI
% object which is used for the ODE integration
datatable         = xlsread(fullfile(exdir,'pnas_data_original.xls'));
amiData.t         = datatable(:,1);       % time points
amiData.Y         = datatable(:,[2,4,6]); % measurement
amiData.condition = [1.4,0.45];           % initial conditions
amiData.Sigma_Y   = NaN(size(amiData.Y)); % preallocation of variances
amiData           = amidata(amiData);     % calling the AMICI routine

%% Generation of the structs and options for PESTO
% The structs and the PestoOptions object, which are necessary for the 
% PESTO routines to work are created and set to convenient values

% parameters
parameters.min     = -5 * ones(17,1);
parameters.max     =  3 * ones(17,1);
parameters.max(4)  =  6;
parameters.max(2)  =  6;
parameters.min(10) = -6;
parameters.min(4)  = -3;
parameters.min(2)  = -3;
parameters.number  = length(parameters.min);
parameters.name    = {'log_{10}(p1)','log_{10}(p2)','log_{10}(p3)','log_{10}(p4)','log_{10}(init_{STAT})',...
    'log_{10}(sp1)','log_{10}(sp2)','log_{10}(sp3)','log_{10}(sp4)','log_{10}(sp5)',...
    'log_{10}(offset_{tSTAT})','log_{10}(offset_{pSTAT})','log_{10}(scale_{tSTAT})','log_{10}(scale_{pSTAT})',...
    'log_{10}(\sigma_{pSTAT})','log_{10}(\sigma_{tSTAT})','log_{10}(\sigma_{pEpoR})'};

% Initial guess for the parameters
par0 = bsxfun(@plus,parameters.min,bsxfun(@times,parameters.max ...
       - parameters.min, lhsdesign(50,parameters.number,'smooth','off')'));
parameters.guess = par0(:,1:50);

% objective function
objectiveFunction = @(theta) logLikelihoodJakstat(theta, amiData);

% PestoOptions
optionsPesto          = PestoOptions();
optionsPesto.trace    = true;
optionsPesto.proposal = 'user-supplied';
optionsPesto.obj_type = 'log-posterior';
optionsPesto.mode     = 'visual';

%% Perform optimization
% A parameters optimization is performed within the bound defined in
% parameters.min and .max in order to infer the unknown parameters from 
% measurement data.

% REMARK: The optimization in this case is rather challenging and the
% box constraints in the parameter space are set generously. So
% optimization will encounter many points in which the ODE can not be
% evaluated, leading to warnings of the ODE simulator AMICI. This is normal
% and not a bug. It just shows how paramter estimation can look like in
% complicated situations.

% Different parameter optimization methods are compared with each other.
% The uncommented version is a simple multi-start local optimization.
% A version with a hybrid optimization technique (MEIGO-ESS) is also
% implemented and commented, as well as a purely global optimization scheme
% (PSwarm). The two alternative (and global) optimization methods are run
% three times, to ensure that the found optimum is indeed the global one.


% % Multi-start local optimization part (fmincon)
% optionsPesto.n_starts = 25;
% optionsPesto.localOptimizer = 'fmincon';
% optionsPesto.localOptimizerOptions = optimset(...
%     'Algorithm', 'interior-point',...
%     'GradObj', 'on',...
%     'Display', 'iter', ... 'Hessian', 'on', ... uncomment this to use the Hessian for optimization 
%     'MaxIter', 1000,...
%     'TolFun', 1e-10,...
%     'MaxFunEvals', 1000*parameters.number);

% Multi-start local optimization part (lsqnonlin)
optionsPesto.n_starts = 25;
optionsPesto.localOptimizer = 'lsqnonlin';
optionsPesto.obj_type = 'log-posterior';
optionsPesto.localOptimizerOptions  = optimset(...
    'Algorithm', 'trust-region-reflective',...
    'MaxIter', 1000,...
    'TolFun', 1e-10,...
    'GradObj', 'on', ...
    'Display', 'iter', ...
    'Jacobian', 'on', ...
    'TolX', 1e-12,...
    'TolGrad', 1e-6, ...
    'PrecondBandWidth', Inf,...
    'MaxFunEvals', 2000*parameters.number);
objectiveFunction = @(theta) logLikelihoodJakstatLsqnonlin(theta, amiData);
    
% % Hybrid-type optimization part (requires the MEIGO toolbox)
% % (Install MEIGO from http://gingproc.iim.csic.es/meigom.html and
% % uncomment):
% optionsPestoHybrid.localOptimizerOptions = MeigoOptions;

% % Global optimization part (requires the PSwarm toolbox)
% % (Install from http://www.norg.uminho.pt/aivaz/pswarm/ and uncomment)
% optionsPestoGlobal = optionsPesto.copy;
% optionsPestoGlobal.localOptimizer = 'pswarm';
% optionsPestoGlobal.localOptimizerOptions.MaxObj  = 10000;

% The example can also be run in parallel mode: Uncomment the following, if wanted
% optionsPesto.comp_type = 'parallel'; 
% optionsPesto.mode = 'text';
% % optionsPesto.save = true; 
% optionsPesto.foldername = 'results';
% n_workers = 10;

if strcmp(optionsPesto.comp_type, 'parallel') && (n_workers >= 2)
    parpool(n_workers); 
else
    optionsPesto.comp_type = 'sequential';
end
    
% Run getMultiStarts
fprintf('\n Perform optimization...');
parametersMultistart = getMultiStarts(parameters, objectiveFunction, optionsPesto);
% parametersHybrid = getMultiStarts(parameters, objectiveFunction, optionsPestoHybrid);
% parametersGlobal = getMultiStarts(parameters, objectiveFunction, optionsPestoGlobal);


%% Perform uncertainty analysis
% The uncertainty of the estimated parameters is visualized by computing
% and plotting profile likelihoods. Different mathod can be used.

% Use the hybrid approach for profiles: uncomment this, if wanted
% optionsPesto.profile_method = 'integration';

% Profile likelihood calculation
parametersMultistart = getParameterProfiles(parametersMultistart, objectiveFunction, optionsPesto);


%% Cleaning up

% Close parpool
if strcmp(optionsPesto.comp_type, 'parallel') && (n_workers >= 2)
    delete(gcp('nocreate'))
end
```
