## Supplementary material for "Mechanistic description of spatial processes using integrative modelling of noise-corrupted imaging data": simulate_jakstat_pesto.m

```
% simulate_jakstat_pesto.m is the matlab interface to the cvodes mex
%   which simulates the ordinary differential equation and respective
%   sensitivities according to user specifications.
%   this routine was generated using AMICI commit f2d86cf027e2f2bb7415ed72025acc6a13c80406 in branch master in repo https://github.com/ICB-DCM/AMICI.
%
% USAGE:
% ======
% [...] = simulate_jakstat_pesto(tout,theta)
% [...] = simulate_jakstat_pesto(tout,theta,kappa,data,options)
% [status,tout,x,y,sx,sy] = simulate_jakstat_pesto(...)
%
% INPUTS:
% =======
% tout ... 1 dimensional vector of timepoints at which a solution to the ODE is desired
% theta ... 1 dimensional parameter vector of parameters for which sensitivities are desired.
%           this corresponds to the specification in model.sym.p
% kappa ... 1 dimensional parameter vector of parameters for which sensitivities are not desired.
%           this corresponds to the specification in model.sym.k
% data ... struct containing the following fields:
%     Y ... 2 dimensional matrix containing data.
%           columns must correspond to observables and rows to time-points
%     Sigma_Y ... 2 dimensional matrix containing standard deviation of data.
%           columns must correspond to observables and rows to time-points
%     T ... (optional) 2 dimensional matrix containing events.
%           columns must correspond to event-types and rows to possible event-times
%     Sigma_T ... (optional) 2 dimensional matrix containing standard deviation of events.
%           columns must correspond to event-types and rows to possible event-times
% options ... additional options to pass to the cvodes solver. Refer to the cvodes guide for more documentation.
%    .atol ... absolute tolerance for the solver. default is specified in the user-provided syms function.
%    .rtol ... relative tolerance for the solver. default is specified in the user-provided syms function.
%    .maxsteps    ... maximal number of integration steps. default is specified in the user-provided syms function.
%    .tstart    ... start of integration. for all timepoints before this, values will be set to initial value.
%    .sens_ind ... 1 dimensional vector of indexes for which sensitivities must be computed.
%           default value is 1:length(theta).
%    .x0 ... user-provided state initialisation. This should be a vector of dimension [#states, 1].
%        default is state initialisation based on the model definition.
%    .sx0 ... user-provided sensitivity initialisation. this should be a matrix of dimension [#states x #parameters].
%        default is sensitivity initialisation based on the derivative of the state initialisation.
%    .lmm    ... linear multistep method for forward problem.
%        1: Adams-Bashford
%        2: BDF (DEFAULT)
%    .iter    ... iteration method for linear multistep.
%        1: Functional
%        2: Newton (DEFAULT)
%    .linsol   ... linear solver module.
%        direct solvers:
%        1: Dense (DEFAULT)
%        2: Band (not implemented)
%        3: LAPACK Dense (not implemented)
%        4: LAPACK Band  (not implemented)
%        5: Diag (not implemented)
%        implicit krylov solvers:
%        6: SPGMR
%        7: SPBCG
%        8: SPTFQMR
%        sparse solvers:
%        9: KLU
%    .stldet   ... flag for stability limit detection. this should be turned on for stiff problems.
%        0: OFF
%        1: ON (DEFAULT)
%    .qPositiveX   ... vector of 0 or 1 of same dimension as state vector. 1 enforces positivity of states.
%    .sensi_meth   ... method for sensitivity analysis.
%        'forward': forward sensitivity analysis (DEFAULT)
%        'adjoint': adjoint sensitivity analysis 
%    .adjoint   ... flag for adjoint sensitivity analysis.
%        true: on 
%        false: off (DEFAULT)
%    .ism   ... only available for sensi_meth == 1. Method for computation of forward sensitivities.
%        1: Simultaneous (DEFAULT)
%        2: Staggered
%        3: Staggered1
%    .Nd   ... only available for sensi_meth == 2. Number of Interpolation nodes for forward solution. 
%              Default is 1000. 
%    .interpType   ... only available for sensi_meth == 2. Interpolation method for forward solution.
%        1: Hermite (DEFAULT for problems without discontinuities)
%        2: Polynomial (DEFAULT for problems with discontinuities)
%    .ordering   ... online state reordering.
%        0: AMD reordering (default)
%        1: COLAMD reordering
%        2: natural reordering
%
% Outputs:
% ========
% sol.status ... flag for status of integration. generally status<0 for failed integration
% sol.t ... vector at which the solution was computed
% sol.llh ... likelihood value
% sol.chi2 ... chi2 value
% sol.sllh ... gradient of likelihood
% sol.s2llh ... hessian or hessian-vector-product of likelihood
% sol.x ... time-resolved state vector
% sol.y ... time-resolved output vector
% sol.sx ... time-resolved state sensitivity vector
% sol.sy ... time-resolved output sensitivity vector
% sol.z ... event output
% sol.sz ... sensitivity of event output
function varargout = simulate_jakstat_pesto(varargin)

% DO NOT CHANGE ANYTHING IN THIS FILE UNLESS YOU ARE VERY SURE ABOUT WHAT YOU ARE DOING
% MANUAL CHANGES TO THIS FILE CAN RESULT IN WRONG SOLUTIONS AND CRASHING OF MATLAB
if(nargin<2)
    error('Not enough input arguments.');
else
    tout=varargin{1};
    theta=varargin{2};
end
if(nargin>=3)
    kappa=varargin{3};
else
    kappa=[];
end

if(length(theta)<17)
    error('provided parameter vector is too short');
end


xscale = [];
if(nargin>=5)
    if(isa(varargin{5},'amioption'))
        options_ami = varargin{5};
    else
        options_ami = amioption(varargin{5});
    end
else
    options_ami = amioption();
end
if(isempty(options_ami.sens_ind))
    options_ami.sens_ind = 1:17;
end
if(options_ami.sensi>1)
    error('Second order sensitivities were requested but not computed');
end

if(~isempty(options_ami.pbar))
    pbar = options_ami.pbar;
else
    pbar = ones(size(theta));
end

if(isempty(options_ami.pscale))
    options_ami.pscale = 'log10' ;
end
switch (options_ami.pscale)
    case 1
        chainRuleFactor = exp(theta(options_ami.sens_ind));
    case 2
        chainRuleFactor = 10.^theta(options_ami.sens_ind)*log(10);
    otherwise
        chainRuleFactor = ones(size(options_ami.sens_ind));
end

if(nargout>1)
    if(nargout>4)
        options_ami.sensi = 1;
        options_ami.sensi_meth = 'forward';
    else
        options_ami.sensi = 0;
    end
end
nplist = length(options_ami.sens_ind); % MUST NOT CHANGE THIS VALUE
if(nplist == 0)
    options_ami.sensi = 0;
end
nxfull = 9;
if(isempty(options_ami.qpositivex))
    options_ami.qpositivex = zeros(nxfull,1);
else
    if(numel(options_ami.qpositivex)>=nxfull)
        options_ami.qpositivex = options_ami.qpositivex(:);
    else
        error(['Number of elements in options_ami.qpositivex does not match number of states ' num2str(nxfull) ]);
    end
end
plist = options_ami.sens_ind-1;
if(nargin>=4)
    if(isempty(varargin{4}));
        data=[];
    else
        if(isa(varargin{4},'amidata'));
             data=varargin{4};
        else
            data=amidata(varargin{4});
        end
        if(data.ne>0);
            options_ami.nmaxevent = data.ne;
        else
            data.ne = options_ami.nmaxevent;
        end
        if(isempty(kappa))
            kappa = data.condition;
        end
        if(isempty(tout))
            tout = data.t;
        end
    end
else
    data=[];
end
if(~all(tout==sort(tout)))
    error('Provided time vector is not monotonically increasing!');
end
if(not(length(tout)==length(unique(tout))))
    error('Provided time vector has non-unique entries!!');
end
if(max(options_ami.sens_ind)>17)
    error('Sensitivity index exceeds parameter dimension!')
end
if(length(kappa)<2)
    error('provided condition vector is too short');
end
init = struct();
if(~isempty(options_ami.x0))
    if(size(options_ami.x0,2)~=1)
        error('x0 field must be a row vector!');
    end
    if(size(options_ami.x0,1)~=nxfull)
        error('Number of columns in x0 field does not agree with number of states!');
    end
    init.x0 = options_ami.x0;
end
if(~isempty(options_ami.sx0))
    if(size(options_ami.sx0,2)~=nplist)
        error('Number of rows in sx0 field does not agree with number of model parameters!');
    end
    if(size(options_ami.sx0,1)~=nxfull)
        error('Number of columns in sx0 field does not agree with number of states!');
    end
    init.sx0 = bsxfun(@times,options_ami.sx0,1./permute(chainRuleFactor(:),[2,1]));
end
sol = ami_jakstat_pesto(tout,theta(1:17),kappa(1:2),options_ami,plist,pbar(plist+1),xscale,init,data);
if(nargout>1)
    varargout{1} = sol.status;
    varargout{2} = sol.t;
    varargout{3} = sol.x;
    varargout{4} = sol.y;
    if(nargout>4)
        varargout{5} = sol.sx;
        varargout{6} = sol.sy;
    end
else
    varargout{1} = sol;
end
end
```
