## Supplementary material for "Mechanistic description of spatial processes using integrative modelling of noise-corrupted imaging data": logLikelihoodT.m

```
function [logL, dlogLdtheta, FIM] = logLikelihoodT(varargin)
% Objective function for examples/mRNA_transfection
%
% logLikelihoodT.m provides the log-likelihood, its gradient and an 
% approximation of the Hessian matrix based on Fisher information matrix
% (FIM) for the conversion reaction process.
% 
% Parameters:
%   varargin:
%     theta: Model parameters
%     t: vector of time points
%     D: measurement data
%
% Return values:
%   logL: double, value of log-likelihood
%   dlogLdtheta: double vector, gradient of the log-likelihood
%   FIM: double array, approximation of the Hessian matrix based on the
%       Fisher information matrix


%% Model Definition
% Definition of the biological process:
% Some text I still need to write
%
% Reaction:
% m -> m + G, rate: k_TL * [m]
% G -> 0    , rate: beta
% m -> 0    , rate: delta
% 
% State of the system:
% X = [X_1, X_2] with
% X_1 = [G]: eGFP concentration
% X_2 = [m]: mRNA concentration
%
% Dynamics:
% dX_1/dt = k_TL * X_2 - beta * X_1
% dX_2/dt = -delta * X_2
%
% Measurement:
% Y = X_2
%
% Observables of the system:
% y(t) = h(theta, x(t))
%
% Measurement noise:
% sigma, parameter to be estimated from the data

%% Initialization
% Initialization of observable sensitivities

theta = varargin{1}; 
theta = theta(:);
t     = varargin{2};
D     = varargin{3};

% Parameter assignment
t0     = 10.^theta(1);
kTL_m0 = 10.^theta(2);
beta   = 10.^theta(3);
delta  = 10.^theta(4);
sigma  = 10.^theta(5);

%% Evaluation of likelihood function
% Simulation

% State
X = [exp(-delta*(t-t0)) .* (t>t0), ...
     kTL_m0 * (exp(-beta*(t-t0)) - exp(-delta*(t-t0))) / (delta-beta) .* (t>t0)];
 
% State sensitivities
SX(:,:,1) = -X*[-delta,0;kTL_m0,-beta]';
SX(:,:,2) = [zeros(size(t)), ...
             (exp(-beta*(t-t0)) - exp(-delta*(t-t0))) / (delta-beta) .* (t>t0)];
SX(:,:,3) = [zeros(size(t)), ...
             kTL_m0 * ( (-(t-t0).*exp( -beta*(t-t0))) / (delta-beta) + ...
             (exp(-beta*(t-t0)) - exp(-delta*(t-t0))) / (delta-beta)^2 ) .* (t>t0)];
SX(:,:,4) = [-exp(-delta*(t-t0)) .* (t-t0) .* (t>t0), ...
             kTL_m0 * ( ((t-t0).*exp(-delta*(t-t0))) / (delta-beta) - ...
             (exp(-beta*(t-t0)) - exp(-delta*(t-t0))) / (delta-beta)^2 ) .* (t>t0)];
       
% Chain rule and Observables
SX = bsxfun(@times, SX, 10.^permute(theta(1:4), [3,2,1]) * log(10));
Y = X(:,2);
SY = squeeze(SX(:,2,:));

%% Objective Function Evaluation
% Log-likelihood
logL = - 1/2 * sum(log(2*pi*sigma^2) + ((D-Y) ./ sigma).^2);

% Gradient
dlogLdtheta(1:4,1) = 1/(sigma^2) * sum(bsxfun(@times,(D-Y), SY))';
dlogLdtheta(5,1)   = - sum(1/sigma - (D-Y).^2 ./ (sigma^3)) * (10.^theta(5) * log(10));

% Fisher information matrix
FIM = zeros(5,5);
FIM = FIM - 1/sigma^2 * [SY, zeros(length(t),1)]' * [SY, zeros(length(t),1)];
FIM(5,1:4) = FIM(5,1:4) - (2/sigma) * dlogLdtheta(1:4)' * (10.^theta(5) * log(10));
FIM(1:4,5) = FIM(1:4,5) - (2/sigma) * dlogLdtheta(1:4) * (10.^theta(5) * log(10));
FIM(5,5)   = FIM(5,5) - sum(-(1/sigma^2) + 3*(D-Y).^2 ./ sigma^4) * (10.^theta(5) * log(10))^2 ...
                      - sum( (1/sigma)   -   (D-Y).^2 ./ sigma^3) * (10.^theta(5) * log(10)^2);
                  
end
                  
```
