## Supplementary material for "Mechanistic description of spatial processes using integrative modelling of noise-corrupted imaging data": mainTransfection.m

```
% Main file of the mRNA transfection example
%
% Demonstrates the use of:
% * getMultiStarts()
% * getParameterProfiles()
% * getParameterSamples()
% * getParameterConfidenceIntervals()
% * getPropertyMultiStarts()
% * getPropertyProfiles()
% * getPropertySamples()
% * getPropertyConfidenceIntervals()
%
% Demonstrates furthermore:
% * how to carry out uncertainty analysis for local (non-global) optima
% * how to use a multi-chain method for sampling which is initialized at
%   different parameter optima
% * how to use the PHS algorithm for sampling
% * how to deal with burn-in time for sampling algorithms and how to plot
%   only the samples which are important for the posterior
% * how to use the PSwarm toolbox for optimization (commented code version)
%   and what problems may occur when no gradient based approach is used
% * How to use the profile computation mode 'mixed'
% * How to use Hessian matrices computed by the automated finite difference
%   scheme of PESTO (in getParameterProfiles)
%
% This example is a model for mRNA transfection, taken from the paper
% "Single-cell mRNA transfection studies: Delivery, kinetics and statistics
% by numbers", by Leonhardt C et al., Nanomedicine: NBM, 2014, vol.10
% (see http://dx.doi.org/10.1016/j.nano.2013.11.008)
%
% The data used is measurement data provided in the publication.
%
% This file provides time-series measurement data Y and
% performs a multistart maximum likelihood parameter estimation based on
% these measurements, demonstrating the use of getMultiStarts(). The model
% fit is then visualized.
% 
% Profile likelihood calculation is done by optimization and integration
% using getParameterProfiles() with the option 
% optionsMultistart.profile_method = 'mixed'to have comparison of both
% methods.
%
% Multi-chain Monte-Carlo sampling is performed by getParameterSamples()


%% Preliminary
clear all;
close all;
clc;

TextSizes.DefaultAxesFontSize = 14;
TextSizes.DefaultTextFontSize = 18;
set(0,TextSizes);

% Seed random number generator
rng(0);

%% Model Definition
% See logLikelihoodT.m for a detailed description

%% Data
% We fix a data set. It consists of a vector of time points t and a
% measurement vector ym. This data is taken from the referenced publication.

t = (0:0.2:10)';
ym = [   0
   0
   0
   0
   0
   0
   0
   0
   0
   0
   0
   1.8309
   3.3559
   4.6091
   5.4235
   5.9757
   6.6298
   7.0080
   7.8280
   7.5852
   7.9247
   7.8363
   8.0107
   7.7077
   7.5316
   7.4208
   7.5734
   7.3197
   7.1489
   7.1987
   6.8493
   6.6425
   6.6268
   6.1223
   6.1078
   5.9242
   5.6034
   5.4618
   5.1281
   4.9489
   4.8930
   4.7747
   4.7750
   4.3095
   4.2211
   4.0416
   3.7485
   3.7164
   3.4799
   3.5286
   3.2785];

%% Definition of the Parameter Estimation Problem
% In order to run any PESTO routine, at least the parameters struct with
% the fields shown here and the objective function need to be defined,
% since they are mandatory for getMultiStarts, which is usually the first
% routine needed for any parameter estimation problem.
% In this case, also a properties struct is created. For this struct,
% basically the same routines can be called as for the parameters, just the
% naming is different. Therefore, basically the same fields as for the
% parameters struct have to be created.

% Parameters
parameters.min    = [-2; -5; -5; -5; -2];
parameters.max    = [log10(max(t)); 5; 5; 5; 2];
parameters.number = 5;
parameters.name   = {'log_{10}(t_0)';'log_{10}(k_{TL}*m_0)';...
   'log_{10}(\beta)';'log_{10}(\delta)';'log_{10}(\sigma)'};

% Properties
properties.function = {@(theta) propertyFunction_theta(theta,1),...
   @(theta) propertyFunction_theta(theta,2),...
   @(theta) propertyFunction_theta(theta,3),...
   @(theta) propertyFunction_theta(theta,4),...
   @(theta) propertyFunction_theta(theta,5)};
properties.min      = [-2; -5; -5; -5; -2];
properties.max      = [log10(max(t)); 5; 5; 5; 2];
properties.number   = length(properties.min);
properties.name     = {'log_{10}(t_0)';'log_{10}(k_{TL}*m_0)';...
   'log_{10}(\beta)';'log_{10}(\delta)';'log_{10}(\sigma)'};

% Objective function (Log-likelihood)
objectiveFunction = @(theta) logLikelihoodT(theta, t, ym);

%% Multi-start local optimization
% A multi-start local optimization is performed within the bounds defined in
% parameters.min and .max in order to infer the unknown parameters from
% measurement data. Therefore, a PestoOptions object is created and
% some of its properties are set. Since the field obj_type is set to
% 'log-posterior', the objective function is maximized.

% Options
optionsPesto           = PestoOptions();
optionsPesto.obj_type  = 'log-posterior';
optionsPesto.comp_type = 'sequential';
optionsPesto.mode      = 'visual';
optionsPesto.n_starts  = 15;
optionsPesto.plot_options.group_CI_by = 'methods';

% The example can also be run in parallel mode: Uncomment this, if wanted
% optionsMultistart.comp_type = 'parallel';
% optionsMultistart.mode = 'silent';
% optionsMultistart.save = true;
% optionsMultistart.foldername = 'results';
% n_workers = 20;

% Open matlabpool
if (strcmp(optionsPesto.comp_type, 'parallel') && (n_workers >= 2))
   parpool(n_workers);
end

% This section uses PSwarm, a particle swarm optimizer
% (Install from http://www.norg.uminho.pt/aivaz/pswarm/ and uncomment)
% 
% optionsPestoPSwarm = optionsPesto.copy();
% optionsPestoPSwarm.localOptimizer = 'pswarm';
% optionsPestoPSwarm.localOptimizerOptions.MaxObj  = 25000;
% optionsPestoPSwarm.localOptimizerOptions.MaxIter = 1000;
% optionsPestoPSwarm.localOptimizerOptions.Size    = 100;
% optionsPestoPSwarm.localOptimizerOptions.Social  = 0.5;
% optionsPestoPSwarm.localOptimizerOptions.Cognitial = 0.9;
% optionsPestoPSwarm.localOptimizerOptions.IPrint  = -1;
% 
% parameters = getMultiStarts(parameters, objectiveFunction, optionsPestoPSwarm);


% This section uses multi-start local optimization
parameters = getMultiStarts(parameters, objectiveFunction, optionsPesto);

%% Collection of results, check for bimodality

% Check if a second optimum was found, which is good enough
for iMode = 2 : optionsPesto.n_starts
   if (parameters.MS.logPost(iMode) > 39.5)
      if (abs(parameters.MS.par(3,iMode) - parameters.MS.par(3,1)) > 0.1)
         MAP_index2 = iMode;
         break;
      end
   end
end

% Create information for a second parameter struct
parametersAlt = parameters;

%% Visualization of fit
% The measured data is visualized in a plot, together with fit for the best
% parameter value found during getMultiStarts.

if strcmp(optionsPesto.mode,'visual')
   % Simulation
   tsim = linspace(t(1), t(end), 100);
   ysim = simulate_mRNA_Transfection(10.^parameters.MS.par(:,1), tsim);
   
   % Plot: Fit
   figure();
   plot(t,ym,'bo'); hold on;
   plot(tsim,ysim,'r-');
   xlabel('time t');
   ylabel('output y');
   legend('data', 'fit');
end

%% Profile likelihood calculation -- Parameters
% The uncertainty of the estimated parameters is visualized by computing
% and plotting profile likelihoods. In getParameterProfiles, this is done
% by using repeated reoptimization. The information about the profiles is
% then written to the parameters struct.

% Profiles are computed using optimization and integratiob by 
% getParameterProfiles() with profile_method = 'mixed'.
% Profiles for parameters 1, 2, and 5 are integrated, profiles for
% parameters 3 and 4 are optimized.
optionsPesto.profile_method      = 'mixed';
optionsPesto.profile_optim_index = [2 3];
optionsPesto.profile_integ_index = [1 4 5];

% Hessians for profile integration should be calculated by finite
% differences, so PESTO gets told that he objective function can only
% compute 2 outputs (objective and gradient), hence it will compute the 
% necessary Hessians by finite differences.
optionsPesto.objOutNumber = 2;

parameters = getParameterProfiles(parameters, objectiveFunction, optionsPesto);

% Computation for the second mode
optionsPesto.fh = [];
optionsPesto.MAP_index = MAP_index2;

% Now only the profiles 3 and 4 should be recomputed, since all the other
% profiles are identical for the second mode
optionsPesto.parameter_index = [3 4];
optionsPesto.profile_optim_index = 4;
optionsPesto.profile_integ_index = 3;
parametersAlt = getParameterProfiles(parametersAlt, objectiveFunction, optionsPesto);

% Reset the default value
optionsPesto.objOutNumber = 3;

%% Markov Chain Monte Carlo sampling -- Parameters
% Values for the parameters are sampled by using an Parallel Tempering (PT)
% algorithm. This way, the underlying probability density of the parameter
% distribution can be captured.

% Building a struct covering all sampling options:
optionsPesto.MCMC.nIterations = 1e5;
trainingTime = ceil(optionsPesto.MCMC.nIterations / 5);

% PHS specific options:
optionsPesto.MCMC.samplingAlgorithm = 'PHS';
optionsPesto.MCMC.objOutNumber      = 1;
optionsPesto.MCMC.PHS.nChains       = 4;
optionsPesto.MCMC.PHS.alpha         = 0.51;
optionsPesto.MCMC.PHS.memoryLength  = 1;
optionsPesto.MCMC.PHS.regFactor     = 1e-6;
optionsPesto.MCMC.PHS.trainingTime  = trainingTime;


optionsPesto.MCMC.theta0(:,[1 3]) = parameters.MS.par(:, [1 1]);
optionsPesto.MCMC.sigma0(:,:,[1 3]) = repmat(inv(squeeze(parameters.MS.hessian(:, :, 1))), [1 1 2]);
optionsPesto.MCMC.theta0(:,[2 4]) = parameters.MS.par(:, [MAP_index2 MAP_index2]);
optionsPesto.MCMC.sigma0(:,:,[2 4]) = repmat(inv(squeeze(parameters.MS.hessian(:, :, MAP_index2))), [1 1 2]);

% Run the sampling
parameters = getParameterSamples(parameters, objectiveFunction, optionsPesto);

% Now, the plots for the sampling include the burn-in phase, which the
% sampling algorithm needs to explore the posterior. Those burn-in samples
% corrupt the shape of the posterior, if they are not removed from the
% histogram. So we clean the parameters.S struct from them (it may yet be
% good to retain a copy of all samples, to do some more datailed analysis
% and diagnosis with it.

% Cleaning up
parameters.S.par(:,1:trainingTime,:) = [];
parameters.S.logPost(1:trainingTime,:) = [];

% Plotting of the new sampling set
optionsPesto.plot_options.S.plot_type = 1;
optionsPesto.plot_options.S.ind = 1;
optionsPesto.plot_options.S.bins = 'optimal';
fh = figure('Name','plotParameterSamples after CleanUp - 1D');
plotParameterSamples(parameters,'1D',fh,[],optionsPesto.plot_options);
fh = figure('Name','plotParameterSamples after CleanUp - 2D');
plotParameterSamples(parameters,'2D',fh,[],optionsPesto.plot_options);


%% Confidence interval evaluation -- Parameters
% Confidence intervals to the confidence levels fixed in the array alpha
% are computed based on local approximations from the Hessian matrix at the
% optimum, based on the profile likelihoods and on the parameter sampling.

alpha = [0.9,0.95,0.99];

% Computation for the first mode
optionsPesto.MAP_index = 1;
optionsPesto.parameter_index = 1 : parameters.number;
parameters = getParameterConfidenceIntervals(parameters, alpha, optionsPesto);

% Computation for the second mode
optionsPesto.MAP_index = MAP_index2;
optionsPesto.parameter_index = [3, 4];
parametersAlt = getParameterConfidenceIntervals(parametersAlt, alpha, optionsPesto);

%% Evaluation of properties for multi-start local optimization results -- Properties
% The values of the properties are evaluated at the end points of the
% multi-start optimization runs by getPropertyMultiStarts.

optionsPesto.MAP_index = 1;
optionsPesto.parameter_index = 1 : parameters.number;
properties = getPropertyMultiStarts(properties, parameters, optionsPesto);

%% Profile likelihood calculation -- Properties
% Profile likelihoods are computed for the properties in the same fashion,
% as they were computed for the parameters.

properties = getPropertyProfiles(properties, parameters, objectiveFunction, optionsPesto);

%% Evaluation of properties for sampling results -- Properties
% From the samples of the parameters, the properties are calculated and
% hence a probability distribution for the properties can be reconstructed
% from that.

properties = getPropertySamples(properties, parameters, optionsPesto);

%% Confidence interval evaluation -- Properties
% As for the parameters, confidence intervals are computed for the
% properties in different fashion, based on local approximations, profile
% likelihoods and samples.

properties = getPropertyConfidenceIntervals(properties, alpha);

%% Comparison of calculated parameter profiles

if strcmp(optionsPesto.mode, 'visual')
   % Open figure
   figure
   
   % Loop: parameters
   for i = 1:parameters.number
      subplot(ceil(parameters.number/ceil(sqrt(parameters.number))),ceil(sqrt(parameters.number)),i);
      plot(parameters.P(i).par(i,:),parameters.P(i).R,'bx-'); hold on;
      plot(properties.P(i).prop, properties.P(i).R,'r-o');
      xlabel(properties.name{i});
      ylabel('likelihood ratio');
      if i == 1
         legend('unconst. opt. (= standard)','unconst. op. (= new)');
      end
   end
end

%% Close the pools of parallel working threads

if strcmp(optionsPesto.comp_type, 'parallel')
   delete(gcp('nocreate'));
end
```
