## Supplementary material for "Mechanistic description of spatial processes using integrative modelling of noise-corrupted imaging data": propertyFunction_theta.m

```
function varargout = propertyFunction_theta(theta, i)
% propertyFunction_theta for examples/mRNA_transfection
%
% returns the value of the i-th parameter as defined in 
% logLikelihood.m with derivatives
%
% Parameters: 
%  theta: parameter vector
%  i: index for the parameter
%
% Return values:
%  f: double, value of property function
%  grad_f: double vector, gradient of property function
%  hess_f: double array, hessian of property function


%% Property Function Evaluation
n = length(theta);

f         = theta(i);
grad_f    = zeros(n, 1);
grad_f(i) = 1;
hess_f    = zeros(n, n);

varargout{1} = f;
varargout{2} = grad_f;
varargout{3} = hess_f;

end
```
