## Supplementary material for "Mechanistic description of spatial processes using integrative modelling of noise-corrupted imaging data": simulate_mRNA_Transfection.m

```
function [Y, X] = simulate_mRNA_Transfection(theta, t)
% simulate_mRNA_Transfection for examples/mRNA_transfection
%
% simulateConversionReaction performs a simulation of the transfection 
% model for the given timepoints t and parameters theta
%
% Parameters:
%  theta: Model parameters [theta_1, theta_2]'
%  t: vector of time points
%
% Return values:
%  Y: Vector with values of the observables Y = [X_2] at timepoints t
%  X: State vector at timepoints t


%% Parameter assignment
t0 = theta(1);
kTL_m0 = theta(2);
beta = theta(3);
delta = theta(4);

%% Simulation
t = t(:);
X = [exp(-delta*(t-t0)).*(t>t0),...
     kTL_m0 * (exp(-beta*(t-t0)) - exp(-delta*(t-t0))) / (delta-beta) .* (t>t0)];
 
%% Output assignment
Y = X(:, 2);

end
```
