## Supplementary material for "Mechanistic description of spatial processes using integrative modelling of noise-corrupted imaging data": logLikelihoodPom1.m

```
function [varargout] = logLikelihoodPom1(varargin)
%function [ F ] = MLEPom1P_w_grad(p, options)
% Calculates the MLE vector and gradient for fmincon for the kinetic parameters
%
% USAGE:
% ======
% [...] = logLikelihoodPom1(parameter,options)
%
% INPUTS:
% =======
% parameter ... log-parameter values in the order given by options.name
% options ... structure needed to evaluate the Likelihood
%   .options.datafile path to load options.Data file
%   .disc     structure containing the PDE solver information
%   .name     parameter names
%   .sign     default: negative
%   .grad_ind default: 1:length(p)
%   .type     default: 'none'
%
% Outputs:
% ========
% F  ... negative log-Likelihood
% dF ... gradient of negative log-Likelihood in the 
%    order of options.name
%
% 2013/04/20 Sabrina Hock
% 2016/10/17 Anna Fiedler

%% CHECK AND ASSIGN INPUTS
if nargin >= 1
    p = varargin{1};
else
    error('logLikelihoodPom1 requires a parameter object as input.');
end

% Check and assign options
if nargin == 2
    options = varargin{2};
else
    error('logLikelihoodPom1 requires a option object as input.');
end

% Check function evaluation counter
if strcmp(options.counter,'true')
    global fcn_count
    fcn_count = fcn_count + 1;
end


%% asign data
n_data = [196,0,13,14,82];

Data{1}.Y = options.Estdata{1}.Data(:,2);
Data{1}.Sigma_Y = options.Estdata{1}.Data(:,3)./sqrt(n_data(1))';

Data{2}.Y = options.Estdata{2}.Data(:,2);
Data{2}.Sigma_Y = options.Estdata{2}.Data(:,3)./sqrt(n_data(3));

Data{3}.Y = options.Estdata{3}.Data(:,2);
Data{3}.Sigma_Y = options.Estdata{3}.Data(:,3)./sqrt(n_data(4));

Data{4}.Y = options.Estdata{4}.Data(1);
Data{4}.Sigma_Y = options.Estdata{4}.Data(2)./sqrt(n_data(5));

% asign model functions
modelEq = str2func(['simulate_Pom1p_',options.model_type,'_model_hess_eq']);
modelFt = str2func(['simulate_Pom1p_',options.model_type,'_model_hess_ft']);
modelHt = str2func(['simulate_Pom1p_',options.model_type,'_model_hess_ht']);

% asign options
options_simu.sensi = 1;
options_simu.maxsteps = 1e6;

% asign dimensions
n_grid = length(options.disc.p);
switch options.model_type 
    case 'SDD'
        n_comp = 1;
    case 'MSP'
        n_comp = 1;
    case 'AP'
        n_comp = 2;
    case 'NLIC'
        n_comp = 2;
end

%% compute objective function value     
J = 0;
dJ = 0;
        
% solve stedy state PDE
T = 500;
flag_ss = 0;
n = 1;
n_max = 10;
x0 = zeros(n_grid*n_comp,1);
while (n < n_max) && (flag_ss == 0)
    sol = modelEq(T,p,x0,[],options_simu);
    if sum(sol.xdot.^2) < 1e-6
        flag_ss = 1;
    elseif sol.status < 0
        break
    else
        x0 = sol.x(end,:);
        options_simu.sx0 = squeeze(sol.sx(end,:,:));
        T = 2*T;
    end
end
if flag_ss
    % Data set 1 - mean intensity profiles
    J = J - 0.5*sum(sum(log(2*pi*Data{1}.Sigma_Y.^2) + (Data{1}.Y-sol.y(end,1:end-1)').^2./(Data{1}.Sigma_Y.^2)));
    dJ = dJ + permute(sol.sy(end,1:end-1,:),[3,2,1])*((Data{1}.Y-sol.y(end,1:end-1)')./(Data{1}.Sigma_Y.^2));
    
    % Data set 4 - total molecule number
    J = J - 0.5*sum(sum(log(2*pi*Data{4}.Sigma_Y.^2) + (Data{4}.Y-sol.y(end,end)).^2./(Data{4}.Sigma_Y.^2)));
    dJ = dJ + permute(sol.sy(end,end,:),[3,2,1])*((Data{4}.Y-sol.y(end,end))./(Data{4}.Sigma_Y.^2));
else
    error('not converged');
end

% Data set 2 - full tip FRAP
% solve PDE for full tip bleaching
Q{3} = find(abs(options.disc.p)<0.5*pi*1.75);
x0 = squeeze(sol.x(end,:));
sx0 = squeeze(sol.sx(end,:,:));
for n = 1:n_comp
    x0((n-1)*n_grid+Q{3}) = 0;
    sx0((n-1)*n_grid+Q{3},:) = 0;
end

options_simu.sx0 = sx0;
sol_full = modelFt(options.Estdata{2}.Data(:,1),p,x0,[],options_simu);
J = J - 0.5*sum(sum(log(2*pi*Data{2}.Sigma_Y.^2) + ((Data{2}.Y-sol_full.y).^2./(Data{2}.Sigma_Y.^2))));
dJ = dJ + permute(sol_full.sy,[3,1,2])*((Data{2}.Y-sol_full.y)./(Data{2}.Sigma_Y.^2));

% Data set 2 - half tip FRAP
% solve PDE for half tip bleaching
Q{4} = find(options.disc.p<0.5*pi*1.75 & options.disc.p>0);
x0 = squeeze(sol.x(end,:));
sx0 = squeeze(sol.sx(end,:,:));
for n = 1:n_comp
    x0((n-1)*n_grid+Q{4}) = 0;
    sx0((n-1)*n_grid+Q{4},:) = 0;
end

options_simu.sx0 = sx0;
sol_half = modelHt(options.Estdata{3}.Data(:,1),p,x0,[],options_simu);
J = J - 0.5*sum(sum(log(2*pi*Data{3}.Sigma_Y.^2) + ((Data{3}.Y-sol_half.y).^2./(Data{3}.Sigma_Y.^2))));
dJ = dJ + permute(sol_half.sy,[3,1,2])*((Data{3}.Y-sol_half.y)./(Data{3}.Sigma_Y.^2));

switch nargout
    case 1
        varargout{1} = J;  
    case 2 
        varargout{1} = J;
        varargout{2} = dJ(options.grad_ind);
end
end
```
