## Supplementary material for "Mechanistic description of spatial processes using integrative modelling of noise-corrupted imaging data": mainPom1.m

```
% Main file of Pom1 example
%
% Demonstrates the use of:
% * getMultiStarts()
% 
% This example provides four models for Pom1 gradient formation. The SDD 
% and NLIC were adapted from Saunders et al. (2012). The AP was adapted 
% from Hachet et al. (2011) and the MSP from Hersch et al. (2015).

clear all;
close all;
clc;

TextSizes.DefaultAxesFontSize = 14;
TextSizes.DefaultTextFontSize = 18;
set(0,TextSizes);

% Seed random number generator
rng(0);

%% Model Definition
% The models are provided as AMICI syms files in the folder Models
addpath('Models');

%% Data
% load data provided in Saunders et. al (2011)
% absolute file paths can be used instead of relative ones
options.optionsLogPost.Estdata{1} = load('./Data/Saunders2011/FigureS5A.mat');
options.optionsLogPost.Estdata{2} = load('./Data/Saunders2011/FigureS5C.mat');
options.optionsLogPost.Estdata{3} = load('./Data/Saunders2011/FigureS5D.mat');
options.optionsLogPost.Estdata{4} = load('./Data/Saunders2011/Figure1C.mat');

%% GENREAL PARAMETERS
options.startdate = datestr(now,'yyyy-mm-dd');
options.starttime = datestr(now,'HH-MM');

% choose gradient formation model
% 'SDD' : source diffusion degradation model
% 'NLIC': non-linear clustering model
% 'AP'  : autophosphorylation model
% 'MSP' : multi-site phosphorylation model
model = 'MSP'; % 'SDD';

%% parameters
switch model
    case 'SDD'
        % Set 5 basic kinetic parameters
        parameters_est.number = 7;
        parameters_est.guess = log10(exp([-0.5,-3,5,-2,-7.184617366,-9.44,-8.69]')); %log
        parameters_est.min = [  -2,  -5,  0,   -1.5,   -5,   -5, -5]';
        parameters_est.max = [   2,   2,  4,      2,   -2,   -2, -2]';
        parameters_est.name = {'D','mu','J','w_tea','s_1','s_2','s_3'};
        Pom1p_SDD_wrap;
    case 'MSP'
        % Set 5 basic kinetic parameters
        parameters_est.number = 7;
        parameters_est.guess = log10(exp([-1.8,-9,6,-1.45,-7.133,-9.44,-8.69]'));
        parameters_est.min = [  -2, -5,  0,   -1.5,   -5,   -5, -5]';
        parameters_est.max = [   2,  2,  4,      2,   -2,   -2, -2]';
        parameters_est.name = {'D','a','J','w_tea','s_1','s_2','s_3'};
        Pom1p_MSP_wrap;
    case 'AP'
        % Set 6 basic kinetic parameters
        parameters_est.number = 9;
        parameters_est.guess = log10(exp([-2,-4,-2,-1.7, 5, 0,-6.1836,-8.7,-8]'));
        parameters_est.min =   [-2,  -5,  -5, -5,  0,   -1.5,   -5,   -5, -5]';
        parameters_est.max =   [ 2,   0,   2,  2,  4,      2,   -2,   -2, -2]';
        parameters_est.name = {'D','xi','mu','a','J','w_tea','s_1','s_2','s_3'};
        Pom1p_AP_wrap;
    case 'NLIC'
        % Set 7 basic kinetic parameters
        parameters_est.number = 12;
        parameters_est.guess = log10(exp([-2,-4,-5,-10,-10,log(0.9),4,-1.45,0.6,-8.3705,-8.5,-7.6]'));
        parameters_est.min = [ -2,  -5,  - 5, -5, -5, -8,  0,   -1.5,  1.2,  -5, -5, -5]';
        parameters_est.max = [  2,   0,    2,  2,  2,  0,  4,      2,    3,  -2, -2, -2]';
        parameters_est.name = {'D','xi','mu','a','b','e','J','w_tea','s_c','s_1','s_2','s_3'};
        Pom1p_NLIC_wrap;
end

%% OPTIMIZATION OPTIONS
% Check and assign options
options.optionsMultistart = PestoOptions();
options.optionsMultistart.obj_type = 'log-posterior';
options.optionsMultistart.comp_type = 'sequential';
options.optionsMultistart.localOptimizerOptions = optimset(options.optionsMultistart.localOptimizerOptions,...
   'Algorithm','interior-point',...
   'Display','off',...
   'GradObj','on',...
   'TolFun',1e-8,...
   'TolX',1e-8,...
   'MaxFunEvals',3000*parameters_est.number,...
   'MaxIter',600);                
options.optionsMultistart.n_starts = 5;
options.optionsMultistart.proposal = 'uniform';
options.optionsMultistart.mode = 'text';
options.optionsMultistart.save = true;
options.optionsMultistart.fh = [];

%% Set-up pde grid
% grid: -7:7 with 0.1, R = 1.75
% equidistant grid
% Changes in the grid have to be carried out in the model files as well!

n_grid = 200;
p = linspace(-7,7,200);
options.optionsLogPost.disc = struct('p',p);

%% LIKELIHOOD OPTIONS
options.optionsLogPost.counter = 'false';
options.optionsLogPost.grad_ind = (1:parameters_est.number)';
options.optionsLogPost.name = parameters_est.name;
options.optionslogPost.plot = 'true';
options.optionsLogPost.model_type = model;

% log-likelihood function
objectiveFunction = @(theta) logLikelihoodPom1(theta,options.optionsLogPost);

%% MULTI-START OPTIMIZATION
disp('Start: Multi-start optimization');

[parameters_est,fh1] = getMultiStarts(parameters_est,objectiveFunction,...
    options.optionsMultistart);

% save results
if options.optionsMultistart.save
    % create directory for estimation results
    if isdir(options.startdate)==0
        mkdir(options.startdate);
    end
    save([options.startdate,'/EstPom1P_',...
        options.optionsLogPost.model_type,'_',options.starttime,'.mat'],...
        'parameters_est','options','objectiveFunction');
end
```
