## Supplementary material for "Mechanistic description of spatial processes using integrative modelling of noise-corrupted imaging data": Pom1p_AP_syms_eq.m

```
function [model] = Pom1p_AP_syms_eq()

%% CVODES OPTIONS
% set the default absolute tolerance
% model.atol = 1e-8; 
% set the default relative tolerance
% model.rtol = 1e-8; 
% set the default maximum number of integration steps
% model.maxsteps = 1e4; 
% set the parametrisation of the problem options are 'log', 'log10' and
% 'lin' (default).
model.param = 'log10'; 
% model.noadjoint = true;

%% PDE discretization
n_grid = 200;
grid = linspace(-7,7,n_grid);
h = (n_grid/14)^2;

%% STATES
% create state syms: vector for p at each grid point (gridsize: 281 -> -7:0.05:7)
x = sym('p',[2*n_grid,1]);

%% PARAMETERS ( for these sensitivities will be computed )

% create parameter syms
syms D xi m a J w_tea s_1 s_2 s_3

% create parameter vector 
p = [D,xi,m,a,J,w_tea,s_1,s_2,s_3];

%% CONSTANTS ( for these no sensitivities will be computed )
% this part is optional and can be ommited

% create parameter syms
k = sym('k',[2*n_grid,1]);

%% SYSTEM EQUATIONS

% create symbolic variable for time
syms t

xdot = sym(zeros(size(x)));

% p boundary
xdot(1) = D*h*(x(2)-x(1)) - a*x(1) + J*exp(-grid(1)^2./(2*w_tea.^2))/(sqrt(2*pi)*w_tea);
xdot(n_grid) = D*h*(x(n_grid-1)-x(n_grid)) - a*x(n_grid) + J*exp(-grid(n_grid)^2./(2*w_tea.^2))/(sqrt(2*pi)*w_tea);

% p_p boundary
xdot(n_grid+1) = (D*xi)*h*(x(n_grid+2)-x(n_grid+1)) - m*x(n_grid+1) + a*x(1);
xdot(2*n_grid) = (D*xi)*h*(x(2*n_grid-1)-x(2*n_grid)) - m*x(2*n_grid) + a*x(n_grid);


for i = 2:n_grid-1
 xdot(i) = D*h*(x(i-1)+x(i+1)-2*x(i)) - a*x(i) + J*exp(-grid(i)^2./(2*w_tea.^2))/(sqrt(2*pi)*w_tea);
 xdot(n_grid+i) = (D*xi)*h*(x(n_grid+i-1)+x(n_grid+i+1)-2*x(n_grid+i)) - m*x(n_grid+i) + a*x(i);
end

%% INITIAL CONDITIONS

x0 = sym(zeros(size(x)));

for i = 1:n_grid
    x0(i) = k(i);
end

%% OBSERVALES

interpList = [17 18; 20 21; 23 24; 26 27; 29 30; 32 33; 34 35; 37 38; 40 41;
    43 44; 46 47; 49 50; 52 53; 54 55; 57 58; 60 61; 63 64; 66 67; 68 69;
    71 72; 74 75; 77 78; 80 81; 83 84; 86 87; 88 89; 91 92; 94 95; 97 98;
   100 101; 103 104; 106 107; 108 109; 111 112; 114 115; 117 118; 120 121;
   123 124; 125 126; 128 129; 131 132; 134 135; 137 138; 140 141; 143 144;
   146 147; 148 149; 151 152; 154 155; 157 158; 159 160; 162 163; 165 166;
   168 169; 171 172; 174 175; 177 178; 180 181; 183 184; 185 186];

weight = [0.872357142857147 0.729428571428570 0.586500000000008 0.642571428571438...
   0.314857142857137 0.186142857142853 0.872642857142856 0.914500000000007...
   0.771571428571432 0.287499999999996 0.144571428571429 0.015857142857145...
   0.057714285714284 0.729999999999997 0.786071428571432 0.458357142857143...
   0.514428571428572 0.371500000000006 0.873214285714281 0.915071428571426...
   0.786357142857144 0.657642857142860 0.699499999999996 0.201214285714288...
   0.072499999999997 0.929571428571428 0.800857142857142 0.487357142857144...
   0.344428571428574 0.386285714285715 0.072785714285708 0.114642857142855...
   0.801142857142855 0.487642857142857 0.529500000000003 0.571357142857144...
   0.243642857142855 0.114928571428566 0.801428571428572 0.843285714285717...
   0.700357142857147 0.386857142857137 0.243928571428567 0.115214285714281...
   0.157071428571423 0.014142857142857 0.700642857142863 0.571928571428569...
   0.428999999999999 0.300285714285713 0.972571428571428 0.843857142857145...
   0.885714285714284 0.742785714285723 0.784642857142856 0.300571428571429...
   0.157642857142848 0.028928571428566 0.070785714285717 0.757285714285723]';


y = sym(zeros(61,1));

% y1 = (1-weight).*(x(interpList(:,1))/x(100))+weight.*(x(interpList(:,2))/x(100));
y1 = (1-weight).*(x(interpList(:,1))+x(interpList(:,1)+n_grid))+weight.*(x(interpList(:,2))+x(interpList(:,2)+n_grid));

y2 = 2*trapz(grid,x(1:n_grid)+x(1+n_grid:2*n_grid));

y = [s_1*y1; y2];

%% SYSTEM STRUCT

model.sym.x = x;
model.sym.k = k;
model.sym.xdot = xdot;
model.sym.p = p;
model.sym.x0 = x0;
model.sym.y = y;
end
```
