## Supplementary material for "Mechanistic description of spatial processes using integrative modelling of noise-corrupted imaging data": Pom1p_MSP_syms_ft.m

```
function [model] = Pom1p_MSP_syms_ft()

%% CVODES OPTIONS

% set the default absolute tolerance
% model.atol = 1e-8; 
% set the default relative tolerance
% model.rtol = 1e-8; 
% set the default maximum number of integration steps
% model.maxsteps = 1e4; 
% set the parametrisation of the problem options are 'log', 'log10' and
% 'lin' (default).
model.param = 'log10'; 
% model.noadjoint = true;

%% PDE discretization
n_grid = 200;
grid = linspace(-7,7,n_grid);
h = (n_grid/14)^2;

%% STATES
% create state syms: vector for p at each grid point (gridsize: 281 -> -7:0.05:7)
x = sym('p',[n_grid,1]);

%% PARAMETERS ( for these sensitivities will be computed )

% create parameter syms
syms D a J w_tea s_1 s_2 s_3

% create parameter vector 
p = [D,a,J,w_tea,s_1,s_2,s_3];

%% CONSTANTS ( for these no sensitivities will be computed )
% this part is optional and can be ommited

% create parameter syms
k = sym('k',[n_grid,1]);

%% SYSTEM EQUATIONS

% create symbolic variable for time
syms t

xdot = sym(zeros(size(x)));

% p boundary
xdot(1) = D*h*(x(2)-x(1)) - a*x(1).^2 + J*exp(-grid(1)^2./(2*w_tea.^2))/(sqrt(2*pi)*w_tea);
xdot(n_grid) = D*h*(x(n_grid-1)-x(n_grid)) - a*x(n_grid).^2 + J*exp(-grid(n_grid)^2./(2*w_tea.^2))/(sqrt(2*pi)*w_tea);

for i = 2:n_grid-1
 xdot(i) = D*h*(x(i-1)+x(i+1)-2*x(i)) - a*x(i).^2 + J*exp(-grid(i)^2./(2*w_tea.^2))/(sqrt(2*pi)*w_tea);
end

%% INITIAL CONDITIONS

x0 = sym(zeros(size(x)));

for i = 1:n_grid
    x0(i) = k(i);
end

%% OBSERVALES
Q = find(abs(grid)<0.5*pi*1.75);
y = s_2*trapz(grid(Q),x(Q));

%% SYSTEM STRUCT

model.sym.x = x;
model.sym.k = k;
model.sym.xdot = xdot;
model.sym.p = p;
model.sym.x0 = x0;
model.sym.y = y;
end
```
