## Supplementary material for "Mechanistic description of spatial processes using integrative modelling of noise-corrupted imaging data": Pom1p_MSP_wrap.m

```
% clear all
% close all
% clc

%% COMPILATION

[exdir,~,~]=fileparts(which('Pom1p_MSP_wrap.m'));

% compile the model
amiwrap('Pom1p_MSP_model_hess_eq','Pom1p_MSP_syms_eq',exdir)

% add the model to the path
addpath(genpath([strrep(which('amiwrap.m'),'amiwrap.m','') 'models/Pom1p_MSP_model_hess_eq']))

% compile the full tip model
amiwrap('Pom1p_MSP_model_hess_ft','Pom1p_MSP_syms_ft',exdir)

% add the model to the path
addpath(genpath([strrep(which('amiwrap.m'),'amiwrap.m','') 'models/Pom1p_MSP_model_hess_ft']))

% compile the model
amiwrap('Pom1p_MSP_model_hess_ht','Pom1p_MSP_syms_ht',exdir)

% add the model to the path
addpath(genpath([strrep(which('amiwrap.m'),'amiwrap.m','') 'models/Pom1p_MSP_model_hess_ht']))
```
