## Supplementary material for "Mechanistic description of spatial processes using integrative modelling of noise-corrupted imaging data": Pom1p_NLIC_syms.m

```
function [model] = Pom1p_NLIC_syms()

%% CVODES OPTIONS

% set the default absolute tolerance
model.atol = 1e-10; 
% set the default relative tolerance
model.rtol = 1e-10; 
% set the default maximum number of integration steps
model.maxsteps = 1e4; 
% set the parametrisation of the problem options are 'log', 'log10' and
% 'lin' (default).
model.param = 'lin'; 
model.noadjoint = true;

%% PDE discretization
n_grid = 200;
grid = linspace(-7,7,n_grid);
h = (n_grid/14)^2;

%% STATES
% create state syms: vector for p at each grid point (gridsize: 281 -> -7:0.05:7)
x = sym('p',[2*n_grid,1]);

%% PARAMETERS ( for these sensitivities will be computed )

% create parameter syms
syms D xi m a b e J w_tea s_c s_1 s_2 s_3

% create parameter vector 
p = [D,xi,m,a,b,e,J,w_tea,s_c,s_1,s_2,s_3];

%% CONSTANTS ( for these no sensitivities will be computed )
% this part is optional and can be ommited

% create parameter syms
k = sym('k',[2*n_grid,1]);

%% SYSTEM EQUATIONS

% create symbolic variable for time
syms t

xdot = sym(zeros(size(x)));

% p boundary
xdot(1) = D*h*(x(2)-x(1)) + a*x(n_grid+1) -b*x(1)*x(n_grid+1) -m*x(1) + e*J*exp(-grid(1)^2./(2*w_tea.^2))/(sqrt(2*pi)*w_tea);
xdot(n_grid) = D*h*(x(n_grid-1)-x(n_grid)) + a*x(2*n_grid) -b*x(n_grid)*x(2*n_grid) -m*x(n_grid) + e*J*exp(-grid(n_grid)^2./(2*w_tea.^2))/(sqrt(2*pi)*w_tea);

% p_p boundary
xdot(n_grid+1) = (D*xi)*h*(x(n_grid+2)-x(n_grid+1)) - a*x(n_grid+1) + b*x(1)*x(n_grid+1) + (1-e)*J*exp(-grid(1)^2./(2*w_tea.^2))/(sqrt(2*pi)*w_tea);
xdot(2*n_grid) = (D*xi)*h*(x(2*n_grid-1)-x(2*n_grid)) - a*x(2*n_grid) + b*x(n_grid)*x(2*n_grid) + (1-e)*J*exp(-grid(n_grid)^2./(2*w_tea.^2))/(sqrt(2*pi)*w_tea);


for i = 2:n_grid-1
 xdot(i) = D*h*(x(i-1)+x(i+1)-2*x(i)) + a*x(n_grid+i) -b*x(i)*x(n_grid+i) -m*x(i) + e*J*exp(-grid(i)^2./(2*w_tea.^2))/(sqrt(2*pi)*w_tea);
 xdot(n_grid+i) = (D*xi)*h*(x(n_grid+i-1)+x(n_grid+i+1)-2*x(n_grid+i)) - a*x(n_grid+i) + b*x(i)*x(n_grid+i) + (1-e)*J*exp(-grid(i)^2./(2*w_tea.^2))/(sqrt(2*pi)*w_tea);
end

%% INITIAL CONDITIONS

x0 = sym(zeros(size(x)));

for i = 1:n_grid
    x0(i) = k(i);
end

%% OBSERVALES

y =sym('y',[n_grid,1]);

for i = 1:n_grid
    y(i) = x(i)+s_c*x(i+n_grid);
end

%% SYSTEM STRUCT

model.sym.x = x;
model.sym.k = k;
model.sym.xdot = xdot;
model.sym.p = p;
model.sym.x0 = x0;
model.sym.y = y;
end
```
