## Supplementary material for "Mechanistic description of spatial processes using integrative modelling of noise-corrupted imaging data": simulate_Pom1p_AP_model_hess_eq.m

```
% simulate_Pom1p_AP_model_hess_eq.m is the matlab interface to the cvodes mex
%   which simulates the ordinary differential equation and respective
%   sensitivities according to user specifications.
%   this routine was generated using AMICI commit # in branch unknown branch in repo unknown repository.
%
% USAGE:
% ======
% [...] = simulate_Pom1p_AP_model_hess_eq(tout,theta)
% [...] = simulate_Pom1p_AP_model_hess_eq(tout,theta,kappa,data,options)
% [status,tout,x,y,sx,sy] = simulate_Pom1p_AP_model_hess_eq(...)
%
% INPUTS:
% =======
% tout ... 1 dimensional vector of timepoints at which a solution to the ODE is desired
% theta ... 1 dimensional parameter vector of parameters for which sensitivities are desired.
%           this corresponds to the specification in model.sym.p
% kappa ... 1 dimensional parameter vector of parameters for which sensitivities are not desired.
%           this corresponds to the specification in model.sym.k
% data ... struct containing the following fields. Can have the following fields %     Y ... 2 dimensional matrix containing data.
%           columns must correspond to observables and rows to time-points
%     Sigma_Y ... 2 dimensional matrix containing standard deviation of data.
%           columns must correspond to observables and rows to time-points
%     T ... (optional) 2 dimensional matrix containing events.
%           columns must correspond to event-types and rows to possible event-times
%     Sigma_T ... (optional) 2 dimensional matrix containing standard deviation of events.
%           columns must correspond to event-types and rows to possible event-times
% options ... additional options to pass to the cvodes solver. Refer to the cvodes guide for more documentation.
%    .atol ... absolute tolerance for the solver. default is specified in the user-provided syms function.
%    .rtol ... relative tolerance for the solver. default is specified in the user-provided syms function.
%    .maxsteps    ... maximal number of integration steps. default is specified in the user-provided syms function.
%    .tstart    ... start of integration. for all timepoints before this, values will be set to initial value.
%    .sens_ind ... 1 dimensional vector of indexes for which sensitivities must be computed.
%           default value is 1:length(theta).
%    .sx0 ... user-provided sensitivity initialisation. this should be a matrix of dimension [#states x #parameters].
%        default is sensitivity initialisation based on the derivative of the state initialisation.%    .lmm    ... linear multistep method for forward problem.
%        1: Adams-Bashford
%        2: BDF (DEFAULT)
%    .iter    ... iteration method for linear multistep.
%        1: Functional
%        2: Newton (DEFAULT)
%    .linsol   ... linear solver module.
%        direct solvers:
%        1: Dense (DEFAULT)
%        2: Band (not implented)
%        3: LAPACK Dense (not implented)
%        4: LAPACK Band  (not implented)
%        5: Diag (not implented)
%        implicit krylov solvers:
%        6: SPGMR
%        7: SPBCG
%        8: SPTFQMR
%        sparse solvers:
%        9: KLU
%    .stldet   ... flag for stability limit detection. this should be turned on for stiff problems.
%        0: OFF
%        1: ON (DEFAULT)
%    .qPositiveX   ... vector of 0 or 1 of same dimension as state vector. 1 enforces positivity of states.
%    .sensi_meth   ... method for sensitivity analysis.
%        'forward': forward sensitivity analysis (DEFAULT)
%        'adjoint': adjoint sensitivity analysis 
%        'ss': steady state sensitivity analysis 
%    .adjoint   ... flag for adjoint sensitivity analysis.
%        true: on 
%        false: off (DEFAULT)
%    .ism   ... only available for sensi_meth == 1. Method for computation of forward sensitivities.
%        1: Simultaneous (DEFAULT)
%        2: Staggered
%        3: Staggered1
%    .Nd   ... only available for sensi_meth == 2. Number of Interpolation nodes for forward solution. 
%              Default is 1000. 
%    .interpType   ... only available for sensi_meth == 2. Interpolation method for forward solution.
%        1: Hermite (DEFAULT for problems without discontinuities)
%        2: Polynomial (DEFAULT for problems with discontinuities)
%    .ordering   ... online state reordering.
%        0: AMD reordering (default)
%        1: COLAMD reordering
%        2: natural reordering
%
% Outputs:
% ========
% sol.status ... flag for status of integration. generally status<0 for failed integration
% sol.t ... vector at which the solution was computed
% sol.llh ... likelihood value
% sol.chi2 ... chi2 value
% sol.sllh ... gradient of likelihood
% sol.s2llh ... hessian of likelihood
% sol.x ... time-resolved state vector
% sol.y ... time-resolved output vector
% sol.sx ... time-resolved state sensitivity vector
% sol.sy ... time-resolved output sensitivity vector
% sol.z event output
% sol.sz sensitivity of event output
function varargout = simulate_Pom1p_AP_model_hess_eq(varargin)

% DO NOT CHANGE ANYTHING IN THIS FILE UNLESS YOU ARE VERY SURE ABOUT WHAT YOU ARE DOING
% MANUAL CHANGES TO THIS FILE CAN RESULT IN WRONG SOLUTIONS AND CRASHING OF MATLAB
if(nargin<2)
    error('Not enough input arguments.');
else
    tout=varargin{1};
    phi=varargin{2};
end
if(nargin>=3)
    kappa=varargin{3};
else
    kappa=[];
end
theta = 10.^(phi(:));


if(length(theta)<9)
    error('provided parameter vector is too short');
end


pbar = ones(size(theta));
pbar(pbar==0) = 1;
xscale = [];
if(nargin>=5)
    options_ami = amioption(varargin{5});
else
    options_ami = amioption();
end
if(isempty(options_ami.sens_ind))
    options_ami.sens_ind = 1:9;
end
if(options_ami.sensi<2)
    options_ami.id = transpose([0  0  0  0  0  0  0  0  0  0  0  0  0  0  0  0  0  0  0  0  0  0  0  0  0  0  0  0  0  0  0  0  0  0  0  0  0  0  0  0  0  0  0  0  0  0  0  0  0  0  0  0  0  0  0  0  0  0  0  0  0  0  0  0  0  0  0  0  0  0  0  0  0  0  0  0  0  0  0  0  0  0  0  0  0  0  0  0  0  0  0  0  0  0  0  0  0  0  0  0  0  0  0  0  0  0  0  0  0  0  0  0  0  0  0  0  0  0  0  0  0  0  0  0  0  0  0  0  0  0  0  0  0  0  0  0  0  0  0  0  0  0  0  0  0  0  0  0  0  0  0  0  0  0  0  0  0  0  0  0  0  0  0  0  0  0  0  0  0  0  0  0  0  0  0  0  0  0  0  0  0  0  0  0  0  0  0  0  0  0  0  0  0  0  0  0  0  0  0  0  0  0  0  0  0  0  0  0  0  0  0  0  0  0  0  0  0  0  0  0  0  0  0  0  0  0  0  0  0  0  0  0  0  0  0  0  0  0  0  0  0  0  0  0  0  0  0  0  0  0  0  0  0  0  0  0  0  0  0  0  0  0  0  0  0  0  0  0  0  0  0  0  0  0  0  0  0  0  0  0  0  0  0  0  0  0  0  0  0  0  0  0  0  0  0  0  0  0  0  0  0  0  0  0  0  0  0  0  0  0  0  0  0  0  0  0  0  0  0  0  0  0  0  0  0  0  0  0  0  0  0  0  0  0  0  0  0  0  0  0  0  0  0  0  0  0  0  0  0  0  0  0  0  0  0  0  0  0  0  0  0  0  0  0  0  0  0  0  0  0  0  0  0  0  0  0  0  0  0  0  0  0  0  0  0  0  0  0  0  0  0  0  0  0  0  0  0  0  0  0]);
end
options_ami.z2event = []; % MUST NOT CHANGE THIS VALUE
chainRuleFactor = theta(options_ami.sens_ind)*log(10);

if(nargout>1)
    if(nargout>4)
        options_ami.sensi = 1;
        options_ami.sensi_meth = 'forward';
    else
        options_ami.sensi = 0;
    end
end
if(options_ami.ss>0)
    if(options_ami.sensi>1)
        error('Computation of steady state sensitivity only possible for first order sensitivities');
    end
    options_ami.sensi = 0;
end
np = length(options_ami.sens_ind); % MUST NOT CHANGE THIS VALUE
if(np == 0)
    options_ami.sensi = 0;
end
nxfull = 400;
if(isempty(options_ami.qpositivex))
    options_ami.qpositivex = zeros(nxfull,1);
else
    if(numel(options_ami.qpositivex)>=nxfull)
        options_ami.qpositivex = options_ami.qpositivex(:);
    else
        error(['Number of elements in options_ami.qpositivex does not match number of states ' num2str(nxfull) ]);
    end
end
plist = options_ami.sens_ind-1;
if(nargin>=4)
    if(isempty(varargin{4}));
        data=amidata(length(tout),61,0,options_ami.nmaxevent,length(kappa));
    else
        data=amidata(varargin{4});
    end
else
    data=amidata(length(tout),61,0,options_ami.nmaxevent,length(kappa));
end
if(data.ne>0);
    options_ami.nmaxevent = data.ne;
else
    data.ne = options_ami.nmaxevent;
end
if(isempty(kappa))
    kappa = data.condition;
end
if(isempty(tout))
    tout = data.t;
end
if(~all(tout==sort(tout)))
    error('Provided time vector is not monotonically increasing!');
end
if(not(length(tout)==length(unique(tout))))
    error('Provided time vector has non-unique entries!!');
end
if(max(options_ami.sens_ind)>9)
    error('Sensitivity index exceeds parameter dimension!')
end
if(length(kappa)<400)
    error('provided condition vector is too short');
end
if(~isempty(options_ami.sx0))
    if(size(options_ami.sx0,2)~=np)
        error('Number of rows in sx0 field does not agree with number of model parameters!');
    end
    options_ami.sx0 = bsxfun(@times,options_ami.sx0,1./(permute(theta(options_ami.sens_ind),[2,1])*log(10)));
end
sol = ami_Pom1p_AP_model_hess_eq(tout,theta(1:9),kappa(1:400),options_ami,plist,pbar,xscale,data);
if(options_ami.sensi==1)
    sol.sllh = sol.sllh.*theta(options_ami.sens_ind)*log(10);
    sol.sx = bsxfun(@times,sol.sx,permute(theta(options_ami.sens_ind),[3,2,1])*log(10));
    sol.sy = bsxfun(@times,sol.sy,permute(theta(options_ami.sens_ind),[3,2,1])*log(10));
    sol.sz = bsxfun(@times,sol.sz,permute(theta(options_ami.sens_ind),[3,2,1])*log(10));
    sol.srz = bsxfun(@times,sol.srz,permute(theta(options_ami.sens_ind),[3,2,1])*log(10));
    sol.ssigmay = bsxfun(@times,sol.ssigmay,permute(theta(options_ami.sens_ind),[3,2,1])*log(10));
    sol.ssigmayz = bsxfun(@times,sol.ssigmaz,permute(theta(options_ami.sens_ind),[3,2,1])*log(10));
end
if(options_ami.sensi_meth == 3)
    sol.dxdotdp = bsxfun(@times,sol.dxdotdp,permute(theta(options_ami.sens_ind),[2,1])*log(10));
    sol.dydp = bsxfun(@times,sol.dydp,permute(theta(options_ami.sens_ind),[2,1])*log(10));
    sol.sx = -sol.J\sol.dxdotdp;
    sol.sy = sol.dydx*sol.sx + sol.dydp;
end
if(nargout>1)
    varargout{1} = sol.status;
    varargout{2} = sol.t;
    varargout{3} = sol.x;
    varargout{4} = sol.y;
    if(nargout>4)
        varargout{5} = sol.sx;
        varargout{6} = sol.sy;
    end
else
    varargout{1} = sol;
end
end
```
