## Supplementary material for "Mechanistic description of spatial processes using integrative modelling of noise-corrupted imaging data": logLikelihoodRME.m

```
 function [varargout] = logLikelihoodRME(theta, amiData)
% compute the negative loglikelihood and gradient for RafMekErk data

    
    amiOptions.sensi_meth = 'forward';
    if (nargout > 1)
        amiOptions.sensi = 1;
        if (nargout > 2)
            amiOptions.sensi_meth = 'adjoint';
            amiOptions.sensi = 2;
        end
    end
    amiOptions.maxsteps = 1e5;
    amiOptions.atol = 1e-14;
    amiOptions.rtol = 1e-10;
    amiO = amioption(amiOptions);
    try
        % Set number of conditions
        n_u = size(amiData, 2);

        % Initialize output
        llh = 0;
        sllh = zeros(28, 1);
        s2llh = zeros(28, 28);

        for j = 1 : n_u
            % Simulate the condition
            sol = simulate_rafmekerk_pesto(amiData(j).t, theta, amiData(j).condition, amiData(j), amiO);

            % Sum up contribution from conditions
            if (sol.status ~= 0)
                sol.llh = inf;
                break;
            end
            llh = llh + sol.llh;
            if nargout > 1
                sllh = sllh + sol.sllh;
                if nargout > 2
                    s2llh = s2llh + sol.s2llh;
                end
            end
        end

    catch error_thrown

        warning(['Evaluation of likelihood failed. ',error_thrown.message]);
        varargout{1} = inf;
        varargout{2} = inf(n_theta, 1);
    end

    switch (nargout)
        case{0,1}
            varargout{1} = llh;
        case 2
            varargout{1} = llh;
            varargout{2} = sllh;
        case 3
            varargout{1} = llh;
            varargout{2} = sllh;
            varargout{3} = s2llh;
    end

end
```
