## Supplementary material for "Mechanistic description of spatial processes using integrative modelling of noise-corrupted imaging data": logLikelihoodRMELsqnonlin.m

```
 function [varargout] = logLikelihoodRMELsqnonlin(theta, amiData)
% compute the negative loglikelihood and gradient for RafMekErk data

    
    if (nargout == 2)
        amiOptions.sensi = 1;
    else
        amiOptions.sensi = 0;
    end
    amiOptions.sensi_meth = 'forward';
    amiOptions.maxsteps = 1e5;
    amiOptions.atol = 1e-14;
    amiOptions.rtol = 1e-10;
    amiO = amioption(amiOptions);
    
    try
        % Set number of conditions
        nCondition = size(amiData, 2);

        % Initialize output
        res = zeros(112, 1);
        if (nargout == 2)
            sres = zeros(112, 28);
        elseif (nargout == 3)
            llh = 0;
        end

        for iData = 1 : nCondition
            % Simulate the condition
            sol = simulate_rafmekerk_pesto(amiData(iData).t, theta, amiData(iData).condition, amiData(iData), amiO);

            % Sum up contribution from conditions
            if (sol.status ~= 0)
                error('Integration of ODE failed!');
            end
            
            thisRes = (sol.y(:) - amiData(iData).Y(:)) ./ sol.sigmay(:);
            thisRes_sigma = sqrt(log(2*pi*sol.sigmay(:).^2 - log(eps)));
            nanIndex = isnan(amiData(iData).Y(:));
            thisRes(nanIndex) = 0;
            thisRes_sigma(nanIndex) = 0;
            res = res + [thisRes; thisRes_sigma];
            
            if (nargout == 2)
                sres1 = ((1 ./ sol.sigmay(:)) * ones(1,28)) .* reshape(sol.sy, 56, 28);
                sres2 = (((sol.y(:) - amiData(iData).Y(:)) ./ sol.sigmay(:).^2) * ones(1,28)) .* reshape(sol.ssigmay, 56, 28);
                sres3 = ((1 ./ (thisRes_sigma .* sol.sigmay(:))) * ones(1,28)) .* reshape(sol.ssigmay, 56, 28);
                thisSRes = [sres1 - sres2; sres3];
                thisSRes([nanIndex; nanIndex], :) = 0;
                sres = sres + thisSRes;
            elseif (nargout == 3)
                llh = llh + sol.llh;
            end
        end

    catch error_thrown

        warning(['Evaluation of likelihood failed. ',error_thrown.message]);
        varargout{1} = inf(112,1);
        if (nargout == 2)
            varargout{2} = zeros(112,28);
        elseif (nargout == 3)
            varargout{2} = zeros(112,28);
            varargout{3} = inf;
        end
    end

    switch (nargout)
        case{0,1}
            varargout{1} = res;
        case 2
            varargout{1} = res;
            varargout{2} = sres;
        case 3
            varargout{1} = res;
            varargout{2} = [];
            varargout{3} = llh;
    end

end
```
