## Supplementary material for "Mechanistic description of spatial processes using integrative modelling of noise-corrupted imaging data": mainRafMekErkSignaling.m

```
% Hab ein bisschen auskommentiert, was genau wichtig ist

% RafMekErk example
clear akk
close all
clc
rng(0);

% Load data
load('./data.mat');
u = D.conditions;
n_u = size(u,1);
n_theta = 20;
n_sigma = 8;

% Clean up data and make Amici-readable data out of it
for j = 1 : n_u
    amiData(j) = struct(...
        't', D.t{j}, ...
        'condition', D.conditions(j,:), ...
        'Y', D.measurement{j} ...
        );
end
amiD(1) = amidata(amiData(1));
amiD(2) = amidata(amiData(2));
amiD(3) = amidata(amiData(3));

% Wrap model, if necessary
[exdir,~,~]=fileparts(which('rafmekerk_pesto_syms.m'));
% amiwrap('rafmekerk_pesto','rafmekerk_pesto_syms',exdir,1);

% 12 dynamic parameters, 8 scaling parameters, 8 sigma parameters
parameters.min = -5 * ones(28,1);
parameters.min(7) = -10;
parameters.min(9) = -7;
parameters.max = 4 * ones(28,1);
parameters.max(1:3) = 5;
parameters.max(4) = 6;
parameters.max(13:20) = 8;

parameters.number =  n_theta + n_sigma;
parameters.name = {'log_{10}(kdf_Raf)','log_{10}(kp_Raf)','log_{10}(kdp_pMek)',...
                   'log_{10}(kp_pRaf_Mek)','log_{10}(kdp_pErk)','log_{10}(kp_pMek_Erk)',...
                   'log_{10}(K_pErk_inh)','log_{10}(sust_Ras_0)','log_{10}(ts_sust_Ras)',...
                   'log_{10}(ts_trans_Ras)','log_{10}(K_Sora)','log_{10}(K_UO)',... 
                   'log_{10}(scale_pMek_20140430_gel1)','log_{10}(scale_pErk_20140430_gel1)',...
                   'log_{10}(scale_pMek_20140430_gel2)','log_{10}(scale_pErk_20140430_gel2)',...
                   'log_{10}(scale_pMek_20140505_gel1)','log_{10}(scale_pErk_20140505_gel1)',...
                   'log_{10}(scale_pMek_20140505_gel2)','log_{10}(scale_pErk_20140505_gel2)',... 
                   'log_{10}(sigma_pMek_20140430_gel1)','log_{10}(sigma_pErk_20140430_gel1)',...
                   'log_{10}(sigma_pMek_20140430_gel2)','log_{10}(sigma_pErk_20140430_gel2)',...
                   'log_{10}(sigma_pMek_20140505_gel1)','log_{10}(sigma_pErk_20140505_gel1)',...
                   'log_{10}(sigma_pMek_20140505_gel2)','log_{10}(sigma_pErk_20140505_gel2)'...
                   };

objectiveFunction = @(theta) logLikelihoodRME(theta, amiD);

%% Wichtiger Teil f�r Meigo und DHC

optionsPesto = PestoOptions();
optionsPesto.n_starts = 100; 
optionsPesto.mode = 'visual';
optionsPesto.proposal = 'latin hypercube';
optionsPesto.trace = false;
optionsPesto.obj_type = 'log-posterior';
% optionsPesto.localOptimizer = 'lsqnonlin';
optionsPesto.localOptimizer = 'fmincon';
optionsPesto.localOptimizerOptions.Algorithm = 'interior-point';
optionsPesto.localOptimizerOptions.TolX = 1e-10;
optionsPesto.localOptimizerOptions.TolFun = 1e-8;
optionsPesto.localOptimizerOptions.Display = 'iter';
optionsPesto.localOptimizerOptions.GradObj = 'on';
% optionsPesto.localOptimizerOptions.Jacobian = 'on';
optionsPesto.localOptimizerOptions.Hessian = 'on';
optionsPesto.localOptimizerOptions.MaxFunEvals = 1000;
optionsPesto.localOptimizerOptions.MaxIter = 1000;

parameters = getMultiStarts(parameters, objectiveFunction, optionsPesto);
```
