## Supplementary material for "Mechanistic description of spatial processes using integrative modelling of noise-corrupted imaging data": rafmekerk_pesto_syms.m

```
function [model] = rafmekerk_pesto_syms()

%% CVODES OPTIONS
model.param = 'log10';
%% STATES
syms pRaf pMek pErk 

x = [pRaf pMek pErk];
%% TIME
syms t

%% PARAMETERS
% Define parameters as symbolic variables:
syms kdf_Raf kp_Raf kdp_pMek kp_pRaf_Mek kdp_pErk kp_pMek_Erk K_pErk_inh ...
         sust_Ras_0 ts_sust_Ras ts_trans_Ras K_Sora K_UO ... 
         scale_pMek_20140430_gel1 scale_pErk_20140430_gel1 scale_pMek_20140430_gel2 ...
         scale_pErk_20140430_gel2 scale_pMek_20140505_gel1 scale_pErk_20140505_gel1 ...
         scale_pMek_20140505_gel2 scale_pErk_20140505_gel2 ...
         sigma_pMek_20140430_gel1 sigma_pErk_20140430_gel1 sigma_pMek_20140430_gel2 ...
         sigma_pErk_20140430_gel2 sigma_pMek_20140505_gel1 sigma_pErk_20140505_gel1 ...
         sigma_pMek_20140505_gel2 sigma_pErk_20140505_gel2
     
p = [kdf_Raf kp_Raf kdp_pMek kp_pRaf_Mek kdp_pErk kp_pMek_Erk K_pErk_inh ...
         sust_Ras_0 ts_sust_Ras ts_trans_Ras K_Sora K_UO ...
         scale_pMek_20140430_gel1 scale_pErk_20140430_gel1 scale_pMek_20140430_gel2 ...
         scale_pErk_20140430_gel2 scale_pMek_20140505_gel1 scale_pErk_20140505_gel1 ...
         scale_pMek_20140505_gel2 scale_pErk_20140505_gel2 ...
         sigma_pMek_20140430_gel1 sigma_pErk_20140430_gel1 sigma_pMek_20140430_gel2 ...
         sigma_pErk_20140430_gel2 sigma_pMek_20140505_gel1 sigma_pErk_20140505_gel1 ...
         sigma_pMek_20140505_gel2 sigma_pErk_20140505_gel2];
     
%% INPUT 
% Define input as symbolic variable

%% DOSERESPONSE
syms Sora UO
k = [Sora UO];

%% SYSTEM EQUATIONS
xdot = sym(zeros(size(x)));

RasGTP = sust_Ras_0+(1-exp(-t/ts_sust_Ras))*exp(-t/ts_trans_Ras);
xdot(1) = kp_Raf*(1-pRaf)*(RasGTP)/(1+pErk/K_pErk_inh)-kdf_Raf*pRaf;
xdot(2) = kp_pRaf_Mek*pRaf/(1+Sora/K_Sora)*(1-pMek)-kdp_pMek*pMek;
xdot(3) = kp_pMek_Erk*pMek/(1+UO/K_UO)*(1-pErk)-kdp_pErk*pErk;

%% INITIAL CONDITIONS

x0(1) = ((K_pErk_inh*(sust_Ras_0) + (K_pErk_inh^2*(sust_Ras_0)^2 + (2*K_pErk_inh^2*kdp_pErk*(sust_Ras_0)^2)/kp_pMek_Erk + (K_pErk_inh^2*kdp_pErk^2*(sust_Ras_0)^2)/kp_pMek_Erk^2 + (K_pErk_inh^2*kdp_pMek^2*kdp_pErk^2*(sust_Ras_0 + kdf_Raf/kp_Raf)^2)/(kp_pRaf_Mek^2*kp_pMek_Erk^2) + (2*K_pErk_inh^2*kdp_pMek*kdp_pErk^2*(sust_Ras_0)*(sust_Ras_0 + kdf_Raf/kp_Raf))/(kp_pRaf_Mek*kp_pMek_Erk^2) + (2*K_pErk_inh^2*kdp_pMek*kdp_pErk*(sust_Ras_0)*(sust_Ras_0 + kdf_Raf/kp_Raf))/(kp_pRaf_Mek*kp_pMek_Erk) + (4*K_pErk_inh*kdf_Raf*kdp_pMek*kdp_pErk*(sust_Ras_0))/(kp_Raf*kp_pRaf_Mek*kp_pMek_Erk))^(1/2) + (K_pErk_inh*kdp_pErk*(sust_Ras_0))/kp_pMek_Erk - (K_pErk_inh*kdp_pMek*kdp_pErk*(sust_Ras_0 + kdf_Raf/kp_Raf))/(kp_pRaf_Mek*kp_pMek_Erk)))/(2*(kdf_Raf/kp_Raf  + K_pErk_inh*sust_Ras_0 + (K_pErk_inh*kdf_Raf)/kp_Raf + (K_pErk_inh*kdf_Raf*kdp_pErk)/(kp_Raf*kp_pMek_Erk) + (K_pErk_inh*kdp_pErk*sust_Ras_0)/kp_pMek_Erk));
x0(2) = (((K_pErk_inh^2*(sust_Ras_0)^2 + (2*K_pErk_inh^2*kdp_pErk*(sust_Ras_0)^2)/kp_pMek_Erk + (K_pErk_inh^2*kdp_pErk^2*(sust_Ras_0)^2)/kp_pMek_Erk^2 + (K_pErk_inh^2*kdp_pMek^2*kdp_pErk^2*(sust_Ras_0 + kdf_Raf/kp_Raf)^2)/(kp_pRaf_Mek^2*kp_pMek_Erk^2) + (2*K_pErk_inh^2*kdp_pMek*kdp_pErk^2*(sust_Ras_0)*(sust_Ras_0 + kdf_Raf/kp_Raf))/(kp_pRaf_Mek*kp_pMek_Erk^2) + (2*K_pErk_inh^2*kdp_pMek*kdp_pErk*(sust_Ras_0)*(sust_Ras_0 + kdf_Raf/kp_Raf))/(kp_pRaf_Mek*kp_pMek_Erk) + (4*K_pErk_inh*kdf_Raf*kdp_pMek*kdp_pErk*(sust_Ras_0))/(kp_Raf*kp_pRaf_Mek*kp_pMek_Erk))^(1/2) + K_pErk_inh*sust_Ras_0 + (K_pErk_inh*kdp_pErk*sust_Ras_0)/kp_pMek_Erk - (K_pErk_inh*kdf_Raf*kdp_pMek*kdp_pErk)/(kp_Raf*kp_pRaf_Mek*kp_pMek_Erk) - (K_pErk_inh*kdp_pMek*kdp_pErk*sust_Ras_0)/(kp_pRaf_Mek*kp_pMek_Erk)))/((K_pErk_inh^2*(sust_Ras_0)^2 + (2*K_pErk_inh^2*kdp_pErk*(sust_Ras_0)^2)/kp_pMek_Erk + (K_pErk_inh^2*kdp_pErk^2*(sust_Ras_0)^2)/kp_pMek_Erk^2 + (K_pErk_inh^2*kdp_pMek^2*kdp_pErk^2*(sust_Ras_0 + kdf_Raf/kp_Raf)^2)/(kp_pRaf_Mek^2*kp_pMek_Erk^2) + (2*K_pErk_inh^2*kdp_pMek*kdp_pErk^2*(sust_Ras_0)*(sust_Ras_0 + kdf_Raf/kp_Raf))/(kp_pRaf_Mek*kp_pMek_Erk^2) + (2*K_pErk_inh^2*kdp_pMek*kdp_pErk*(sust_Ras_0)*(sust_Ras_0 + kdf_Raf/kp_Raf))/(kp_pRaf_Mek*kp_pMek_Erk) + (4*K_pErk_inh*kdf_Raf*kdp_pMek*kdp_pErk*(sust_Ras_0))/(kp_Raf*kp_pRaf_Mek*kp_pMek_Erk))^(1/2) + K_pErk_inh*sust_Ras_0 + (K_pErk_inh*kdp_pErk*(sust_Ras_0))/kp_pMek_Erk + (kdf_Raf*kdp_pMek*(2*K_pErk_inh + (K_pErk_inh*kdp_pErk)/kp_pMek_Erk + 2))/(kp_Raf*kp_pRaf_Mek) + (K_pErk_inh*kdp_pMek*(sust_Ras_0)*(kdp_pErk/kp_pMek_Erk + 2))/kp_pRaf_Mek);
x0(3) = (((K_pErk_inh^2*(sust_Ras_0)^2 + (2*K_pErk_inh^2*kdp_pErk*(sust_Ras_0)^2)/kp_pMek_Erk + (K_pErk_inh^2*kdp_pErk^2*(sust_Ras_0)^2)/kp_pMek_Erk^2 + (K_pErk_inh^2*kdp_pMek^2*kdp_pErk^2*(sust_Ras_0 + kdf_Raf/kp_Raf)^2)/(kp_pRaf_Mek^2*kp_pMek_Erk^2) + (2*K_pErk_inh^2*kdp_pMek*kdp_pErk^2*(sust_Ras_0)*(sust_Ras_0 + kdf_Raf/kp_Raf))/(kp_pRaf_Mek*kp_pMek_Erk^2) + (2*K_pErk_inh^2*kdp_pMek*kdp_pErk*(sust_Ras_0)*(sust_Ras_0 + kdf_Raf/kp_Raf))/(kp_pRaf_Mek*kp_pMek_Erk) + (4*K_pErk_inh*kdf_Raf*kdp_pMek*kdp_pErk*(sust_Ras_0))/(kp_Raf*kp_pRaf_Mek*kp_pMek_Erk))^(1/2) + K_pErk_inh*sust_Ras_0 + (K_pErk_inh*kdp_pErk*sust_Ras_0)/kp_pMek_Erk - (K_pErk_inh*kdf_Raf*kdp_pMek*kdp_pErk)/(kp_Raf*kp_pRaf_Mek*kp_pMek_Erk) - (K_pErk_inh*kdp_pMek*kdp_pErk*sust_Ras_0)/(kp_pRaf_Mek*kp_pMek_Erk)))/((kdp_pErk/kp_pMek_Erk + 1)*(K_pErk_inh^2*(sust_Ras_0)^2 + (2*K_pErk_inh^2*kdp_pErk*(sust_Ras_0)^2)/kp_pMek_Erk + (K_pErk_inh^2*kdp_pErk^2*(sust_Ras_0)^2)/kp_pMek_Erk^2 + (K_pErk_inh^2*kdp_pMek^2*kdp_pErk^2*(sust_Ras_0 + kdf_Raf/kp_Raf)^2)/(kp_pRaf_Mek^2*kp_pMek_Erk^2) + (2*K_pErk_inh^2*kdp_pMek*kdp_pErk^2*(sust_Ras_0)*(sust_Ras_0 + kdf_Raf/kp_Raf))/(kp_pRaf_Mek*kp_pMek_Erk^2) + (2*K_pErk_inh^2*kdp_pMek*kdp_pErk*(sust_Ras_0)*(sust_Ras_0 + kdf_Raf/kp_Raf))/(kp_pRaf_Mek*kp_pMek_Erk) + (4*K_pErk_inh*kdf_Raf*kdp_pMek*kdp_pErk*(sust_Ras_0))/(kp_Raf*kp_pRaf_Mek*kp_pMek_Erk))^(1/2) + K_pErk_inh*sust_Ras_0*(kdp_pErk/kp_pMek_Erk + 1)^2 + (kdf_Raf*kdp_pMek*kdp_pErk*(K_pErk_inh + (K_pErk_inh*kdp_pErk)/kp_pMek_Erk + 2))/(kp_Raf*kp_pRaf_Mek*kp_pMek_Erk) + (K_pErk_inh*kdp_pMek*kdp_pErk*(sust_Ras_0)*(kdp_pErk/kp_pMek_Erk + 1))/(kp_pRaf_Mek*kp_pMek_Erk));

%% OBSERVABLES

y = [scale_pMek_20140430_gel1*x(2);...
     scale_pErk_20140430_gel1*x(3);...
     scale_pMek_20140430_gel2*x(2);...
     scale_pErk_20140430_gel2*x(3);...
     scale_pMek_20140505_gel1*x(2);...
     scale_pErk_20140505_gel1*x(3);...
     scale_pMek_20140505_gel2*x(2);...
     scale_pErk_20140505_gel2*x(3)];

%% VARAINCES

sigma = [sigma_pMek_20140430_gel1 sigma_pErk_20140430_gel1 sigma_pMek_20140430_gel2 ...
         sigma_pErk_20140430_gel2 sigma_pMek_20140505_gel1 sigma_pErk_20140505_gel1 ...
         sigma_pMek_20140505_gel2 sigma_pErk_20140505_gel2];

%% SYSTEM STRUCT

model.sym.x = x;
model.sym.k = k;
model.sym.xdot = xdot;
model.sym.p = p;
model.sym.x0 = x0;
model.sym.y = y;
model.sym.sigma_y = sigma;

end
```
