## Supplementary material for "Mechanistic description of spatial processes using integrative modelling of noise-corrupted imaging data": mainExampleRing.m

```
% Main file of the hyper ring example
%
% Demonstrates the use of:
% * getMultiStarts()
% * getParameterProfiles()
% * getParameterSamples()
% * getParameterConfidenceIntervals()
% * getPropertyMultiStarts()
% * getPropertySamples()
%
% Demonstrates furthermore:
% * how to do chain analysis
% * how to define properties
%
% mainExampleRing.m shows how to use sampling methods in PESTO. The
% example problem is a smudged hyper-ring. Its properties can be altered in
% defineRingLLH.m and its dimension via ringDimension.
% Written by Benjamin Ballnus 2/2017
%
% Optimization is carried out as multi-start local optimization by
% getMultiStarts().
%
% Pofile likelihoods for the parameters are computed using
% getParameterProfiles().
%
% Markov chain Monte-Carlo sampling is performed by getParameterSamples() 
% and plotted in 1D and 2D.


%% Preliminaries

clear all
close all
clc


%% Problem initialization

% Seed random number generator
rng(0);

% Settings for this example
radius = 15;
sigma = 2;
logP = @(theta) simulateRingLLH(theta, radius, sigma);
ringDimension = 2;

% Set required sampling options for Parallel Tempering
parameters.number = ringDimension;
parameters.min    = -25 * ones(ringDimension,1);
parameters.max    = 25 * ones(ringDimension,1);
parameters.name   = {'X_1','X_2'};

%% Multi-start optimization and profiles

% Optimization
optionsMultistart = PestoOptions();
optionsMultistart.obj_type = 'log-posterior';
optionsMultistart.objOutNumber = 1;
optionsMultistart.n_starts = 20;
optionsMultistart.comp_type = 'sequential';
optionsMultistart.mode = 'visual';
% optionsMultistart.plot_options.add_points.par = theta_true;
% optionsMultistart.plot_options.add_points.logPost = objectiveFunction(theta_true);
% optionsMultistart.plot_options.add_points.prop = nan(properties.number,1);

% Run optimization
parameters = getMultiStarts(parameters, logP, optionsMultistart);

% Run profile calculation
parameters = getParameterProfiles(parameters, logP, optionsMultistart);


%% Sampling

% Sampling Options
optionsSampling                     = PestoSamplingOptions();
optionsSampling.nIterations         = 1e5;

% Using PT
optionsSampling.samplingAlgorithm   = 'PT';
optionsSampling.PT.nTemps           = 3;
optionsSampling.PT.exponentT        = 4;    
optionsSampling.PT.alpha            = 0.51;
optionsSampling.PT.temperatureAlpha = 0.51;
optionsSampling.PT.memoryLength     = 1;
optionsSampling.PT.regFactor        = 1e-4;
optionsSampling.PT.temperatureAdaptionScheme =  'Lacki15'; %'Vousden16';
optionsSampling.theta0              = repmat(-15*ones(ringDimension,1), 1, optionsSampling.PT.nTemps);
optionsSampling.sigma0              = 1e5*diag(ones(1,ringDimension));

% Using DRAM
% optionsSampling.samplingAlgorithm     = 'DRAM';
% optionsSampling.DRAM.nTry             = 5;
% optionsSampling.DRAM.verbosityMode    = 'debug';    
% optionsSampling.DRAM.adaptionInterval = 1;
% optionsSampling.DRAM.regFactor        = 1e-4;
% optionsSampling.theta0                = -15*ones(ringDimension,1); 
% optionsSampling.sigma0                = 1e5*diag(ones(1,ringDimension));

% Using MALA
% optionsSampling.samplingAlgorithm     = 'MALA';
% optionsSampling.MALA.regFactor        = 1e-4;
% optionsSampling.theta0                = -15*ones(ringDimension,1); 
% optionsSampling.sigma0                = 1e5*diag(ones(1,ringDimension));

% Using PHS
% optionsSampling.samplingAlgorithm     = 'PHS';
% optionsSampling.PHS.nChains           = 3;
% optionsSampling.PHS.alpha             = 0.51;
% optionsSampling.PHS.memoryLength      = 1;
% optionsSampling.PHS.regFactor         = 1e-4;
% optionsSampling.PHS.trainingTime      = ceil(opt.nIterations / 5);
% optionsSampling.theta0                = repmat([-15*ones(ringDimension,1)],1,opt.PHS.nChains); 
% optionsSampling.sigma0                = 1e5*diag(ones(1,ringDimension));

optionsMultistart.MCMC = optionsSampling;

% Perform the parameter estimation via sampling
parameters = getParameterSamples(parameters, logP, optionsMultistart);


%% Visualize the chain history and the theoretical disttribution

% Initialization
figure('Name', 'Chain diagnosis');
subplot(2,1,1); 
plot(squeeze(parameters.S.par(:,:,1))');
subplot(2,1,2); 
plotRing();

% Plotting
hold on;
plot(squeeze(parameters.S.par(1,:,1))',squeeze(parameters.S.par(2,:,1))','.');
hold off;


%% Confidence intervals
parameters = getParameterConfidenceIntervals(parameters, [0.9,0.95,0.99]);


%% Set properties
% properties (parameter function to be analyzed)
property_sum = @(par) (abs(par(1)) + abs(par(2)));
property_difference = @(par) (abs(par(1)) - abs(par(2)));
property_radius = @(par) sqrt(par(1)^2 + par(2)^2);
properties.name = {'X_1 + X_2','X_1 - X_2',...
                   'Radius'};
properties.function = {property_sum, property_difference, property_radius};
properties.min = [-40; -40; 0];
properties.max = [40; 40; 25];
properties.number = length(properties.name);


%% Properties of Multi-start results

% Clear figure in options, to make sure a new figure is created
optionsMultistart.fh = [];
properties = getPropertyMultiStarts(properties, parameters, optionsMultistart);


%% Properties of samples

% Evaluate Properties
properties = getPropertySamples(properties, parameters, optionsMultistart);
```
