## Supplementary material for "Mechanistic description of spatial processes using integrative modelling of noise-corrupted imaging data": plotRing.m

```
% Additional plotting script for examples/RingExample
%
% plotRing.m visualizes the theoretical shape of the ring (ground truth).


% Construct grid for plotting
x = linspace(-25, 25, 400);
y = linspace(-25, 25, 400);
[X,Y] = meshgrid(x,y);

% Compute output value
target = @(p1,p2) (logP([p1;p2]));
Z = arrayfun(target,X,Y);

% Plotting
contour(X,Y,Z,30,'linewidth',1.5);

% Post-processing of the plot
xlabel('X_1');
ylabel('X_2');
```
