## Supplementary material for "Mechanistic description of spatial processes using integrative modelling of noise-corrupted imaging data": simulateRingLLH.m

```
function [ llh ] = simulateRingLLH( par, radius, sigma )
% Simulation function for examples/RingExample
%
% simulateRingLLH.m provides the log-likelihood for the hyper ring example.
% 
% Parameters:
%  par: Model parameters
%  radius:radius of the ring
%  sigma: thickness of the ring
%
% Return values:
%  llh: double, value of log-likelihood / log-posterior


   squares = par.^2;
   llh = - 0.5 * ( abs(sum(squares) - radius^2) / sigma );
   
end
```
