## Supplementary material for "Mechanistic description of spatial processes using integrative modelling of noise-corrupted imaging data": runTestExamples.m

```
function runTestExamples()
% runTestExamples runs some examples for the PESTO toolbox, to test if new
% implementations cause problems

mainConversionReaction;
mainEnzymaticCatalysis;
mainTransfection;
mainErbBSignaling;
mainJakstatSignaling;
mainExampleGauss;
mainExampleRing;

end
```
