## Supplementary material for "Mechanistic description of spatial processes using integrative modelling of noise-corrupted imaging data": getFullStructs.m

```
function [parameters,properties] = getFullStructs(parameters,reducedParameters,properties,reducedProperties,options)

    % Definition of index set of optimized parameters
    freePars = setdiff(1:parameters.number, options.fixedParameters);
    
    %% Parameter struct
    % Assignment of multi-start optimisation results
    if isfield(reducedParameters, 'MS')
        % Results
        parameters.MS = reducedParameters.MS;
        % Extended parameter vector
        n_start = size(parameters.MS.par,2);
        parameters.MS.par0 = zeros(parameters.number, n_start);
        parameters.MS.par0(freePars, :) = reducedParameters.MS.par0;
        parameters.MS.par0(options.fixedParameters, :) = options.fixedParameterValues;
        parameters.MS.par = zeros(parameters.number, n_start);
        parameters.MS.par(freePars, :) = reducedParameters.MS.par;
        parameters.MS.par(options.fixedParameters, :) = options.fixedParameterValues;
        parameters.MS.gradient = zeros(parameters.number, n_start);
        parameters.MS.gradient(freePars, :) = reducedParameters.MS.gradient;
        parameters.MS.hessian = zeros(parameters.number, parameters.number, n_start);
        parameters.MS.hessian(freePars, freePars, :) = reducedParameters.MS.hessian;
    end

    % Assignment of profile calculation results
    if isfield(reducedParameters, 'P')
        % Default values or results stored in parameters.P
        for iPar = 1:parameters.number
            flag_asign_default = true;
            if isfield(parameters,'P')
                if length(parameters.P) >= iPar
                    if isfield(parameters.P(iPar),'par')
                        if ~isempty(parameters.P(iPar).par)
                            flag_asign_default = false;
                        end
                    end
                end
            end
            if flag_asign_default
                parameters.P(iPar).par = [];
                parameters.P(iPar).logPost = [];
                parameters.P(iPar).R = [];
                parameters.P(iPar).t_cpu = [];
                parameters.P(iPar).optSteps = [];
                parameters.P(iPar).intSteps = [];
                parameters.P(iPar).reOptSteps = [];
            end
        end
        % Assignment of reduced profile calculation results
        for iPar_reduced = 1:reducedParameters.number
            % Determine index of parameter in full parameter struct
            iPar = freePars(iPar_reduced);
            % Assign profile
            if length(reducedParameters.P) >= iPar_reduced
                if ~isempty(reducedParameters.P(iPar_reduced).par)
                    % Results contained in reducedParameters.P
                    parameters.P(iPar) = reducedParameters.P(iPar_reduced);
                    % Extended parameter vector
                    parameters.P(iPar).par(freePars, :) = reducedParameters.P(iPar_reduced).par;
                    parameters.P(iPar).par(options.fixedParameters, :) = options.fixedParameterValues;
                end
            end
        end
    end
    
    %% Property struct
    % Assignment of multi-start optimisation results
    if isfield(reducedProperties, 'MS')
        % Results
        properties.MS = reducedProperties.MS;
        % Extended parameter vector
        n_start = size(properties.MS.par,2);
        properties.MS.par = zeros(parameters.number, n_start);
        properties.MS.par(freePars, :) = reducedProperties.MS.par;
        properties.MS.par(options.fixedParameters, :) = options.fixedParameterValues;
    end

    % Assignment of profile calculation results
    if isfield(reducedProperties, 'P')
        for iProp = 1:properties.number
            if isempty(reducedProperties.P(iProp).par)
                % Default values or results contained in properties.P
                flag_asign_default = true;
                if isfield(properties,'P')
                    if length(properties.P) >= iProp
                        if isfield(properties.P(iProp),'par')
                            if ~isempty(properties.P(iProp).par)
                                flag_asign_default = false;
                            end
                        end
                    end
                end
                if flag_asign_default
                    properties.P(iProp).prop = [];
                    properties.P(iProp).par = [];
                    properties.P(iProp).logPost = [];
                    properties.P(iProp).R = [];
                    properties.P(iProp).exitflag = [];
                end
            else
                % Results contained in reducedProperties.P
                properties.P(iProp) = reducedProperties.P(iProp);
                % Extended parameter vector
                properties.P(iProp).par(freePars, :) = reducedProperties.P(iProp).par;
                properties.P(iProp).par(options.fixedParameters, :) = options.fixedParameterValues;
            end
        end
    end
    
end
```
