## Supplementary material for "Mechanistic description of spatial processes using integrative modelling of noise-corrupted imaging data": getMultiStarts.m

```
function [parameters,fh] = getMultiStarts(parameters, objective_function, varargin)
% getMultiStarts() computes the maximum a posterior estimate of the
% parameters of a user-supplied posterior function. Therefore, a
% multi-start local optimization is used. The parameters from the best
% value of the posterior function arethen used as the global optimum.
% To ensure that the found maximum is a global one, a sufficiently high
% number of multistarts must be done. Those starts can be initialized with
% either randomly sampled parameter values, following either a uniform
% distribution or a latin hypercube, or they can be sampled by a user
% provided initial function (provided as option.init_fun).
%
% Note: This function can exploit up to (n_start + 1) workers when running
% in 'parallel' mode.
%
% USAGE:
% * [...] = getMultiStarts(parameters,objective_function)
% * [...] = getMultiStarts(parameters,objective_function,options)
% * [parameters,fh] = getMultiStarts(...)
%
% getMultiStarts() uses the following PestoOptions members:
%  * PestoOptions::start_index
%  * PestoOptions::n_starts
%  * PestoOptions::mode
%  * PestoOptions::fh
%  * PestoOptions::fmincon
%  * PestoOptions::proposal
%  * PestoOptions::save
%  * PestoOptions::foldername
%  * PestoOptions::trace
%  * PestoOptions::comp_type
%  * PestoOptions::tempsave
%  * PestoOptions::resetobjective
%  * PestoOptions::obj_type
%  * PestoOptions::init_threshold
%  * PestoOptions::plot_options
%
% Parameters:
%   parameters: parameter struct
%   objective_function: objective function to be optimized.
%       This function should accept one input, the parameter vector.
%   varargin:
%     options: A PestoOptions object holding various options for the
%         algorithm.
%
% Required fields of parameters:
%   number: Number of parameters
%   min: Lower bound for each parameter
%   max: upper bound for each parameter
%   name = {'name1', ...}: names of the parameters
%   guess: initial guess for the parameters (Optional, will be initialized
%       empty if not provided)
%   init_fun: function to draw starting points for local optimization, must
%       have the structure init_fun(theta_0, theta_min, theta_max).
%       (Only required if proposal == 'user-supplied')
%
% Return values:
%   parameters: updated parameter object
%   fh: figure handle
%
% Generated fields of parameters:
%   MS: information about multi-start optimization
%     * par0(:,i): starting point yielding ith MAP
%     * par(:,i): ith MAP
%     * logPost(i): log-posterior for ith MAP
%     * logPost0(i): log-posterior for starting point yielding ith MAP
%     * gradient(_,i): gradient of log-posterior at ith MAP
%     * hessian(:,:,i): hessian of log-posterior at ith MAP
%     * n_objfun(i): # objective evaluations used to calculate ith MAP
%     * n_iter(i): # iterations used to calculate ith MAP
%     * t_cpu(i): CPU time for calculation of ith MAP
%     * exitflag(i): exitflag the optimizer returned for ith MAP
%     * par_trace(:,:,i): parameter trace for ith MAP
%         (if options.trace == true)
%     * fval_trace(:,i): objective function value trace for ith MAP
%         (if options.trace == true)
%     * time_trace(:,i): computation time trace for ith MAP
%         (if options.trace == true)
%
% History:
% * 2012/05/31 Jan Hasenauer
% * 2012/07/11 Jan Hasenauer
% * 2014/06/11 Jan Hasenauer
% * 2015/07/28 Fabian Froehlich
% * 2015/11/10 Fabian Froehlich
% * 2016/06/07 Paul Stapor
% * 2016/10/04 Daniel Weindl
% * 2016/12/04 Paul Stapor

global error_count

%% Check inputs
if length(varargin) >= 1
    options = handleOptionArgument(varargin{1});
else
    options = PestoOptions();
end

if isempty(options.start_index)
    options.start_index = 1:options.n_starts;
end
parameters = parametersSanityCheck(parameters);

if (strcmp(options.localOptimizer, 'fmincon') && ( ~isfield(options.localOptimizerOptions, 'MaxFunEvals') || isempty(options.localOptimizerOptions.MaxFunEvals) ) )
    options.localOptimizerOptions.MaxFunEvals = 200*parameters.number;
end

%% Initialization and figure generation
fh = [];
switch options.mode
    case 'visual'
        if isempty(options.fh)
            fh = figure('Name', 'getMultiStarts');
        else
            fh = figure(options.fh);
        end
    case 'text'
        fprintf(' \nOptimization:\n=============\n');
    case 'silent' % no output
        % Force fmincon to be silent.
        if (strcmp(options.localOptimizer, 'fmincon') || strcmp(options.localOptimizer, 'lsqnonlin'))
            options.localOptimizerOptions.Display = 'off';
        end
end

%% Definition of index set of optimized parameters
freePars = setdiff(1:parameters.number, options.fixedParameters);

%% Sampling of starting points
switch options.proposal
    case 'latin hypercube'
        % Sampling from latin hypercube
        par0 = [parameters.guess,...
            bsxfun(@plus,parameters.min,bsxfun(@times,parameters.max - parameters.min,...
            lhsdesign(options.n_starts - size(parameters.guess,2),parameters.number,'smooth','off')'))];
    case 'uniform'
        % Sampling from uniform distribution
        par0 = [parameters.guess,...
            bsxfun(@plus,parameters.min,bsxfun(@times,parameters.max - parameters.min,...
            rand(parameters.number,options.n_starts - size(parameters.guess,2))))];
    case 'user-supplied'
        % Sampling from user-supplied function
        if (~isfield(parameters, 'init_fun') || isempty(parameters.init_fun))
            if size(parameters.guess,2) < options.n_starts
                error('You did not define an initial function and do not provide enough starting points in parameters.guess. Aborting.');
            else
                par0 = parameters.guess(:,1:options.n_starts);
            end
        else
            par0 = [parameters.guess,...
                parameters.init_fun(parameters.guess,parameters.min,parameters.max,options.n_starts - size(parameters.guess,2))];
        end
end
% Correct for fixed parameters
par0 = par0(freePars,:);
parameters.MS.n_starts = options.n_starts;
parameters.MS.par0(freePars,:) = par0(:,options.start_index);
parameters.MS.par0(options.fixedParameters,:) = options.fixedParameterValues;

%% Preparation of folder
if or(options.save,options.tempsave)
    if(~exist(fullfile(pwd,options.foldername),'dir'))
        mkdir(fullfile(pwd,options.foldername))
    end
    % only save the init mat for the first start index, not every one if they are called seperately
    if(and(options.save,~isempty(find(options.start_index == 1, 1))))
        save([options.foldername '/init'],'parameters','-v7.3');
    end
end

%% Initialization
if (strcmp(options.localOptimizer, 'fmincon') || strcmp(options.localOptimizer, 'pswarm') || strcmp(options.localOptimizer, 'lsqnonlin'))
    maxOptimSteps = options.localOptimizerOptions.MaxIter;
elseif strcmp(options.localOptimizer, 'meigo-ess') || strcmp(options.localOptimizer, 'meigo-vns')
    maxOptimSteps = options.localOptimizerOptions.maxeval;
end
parameters.MS.par = nan(parameters.number,length(options.start_index));
parameters.MS.logPost0 = nan(length(options.start_index),1);
parameters.MS.logPost = nan(length(options.start_index),1);
parameters.MS.gradient = nan(length(freePars),length(options.start_index));
parameters.MS.hessian  = nan(length(freePars),length(freePars),length(options.start_index));
parameters.MS.n_objfun = nan(length(options.start_index),1);
parameters.MS.n_iter = nan(length(options.start_index),1);
parameters.MS.t_cpu = nan(length(options.start_index),1);
parameters.MS.exitflag = nan(length(options.start_index),1);
parameters.MS.AIC = nan(length(options.start_index),1);
parameters.MS.BIC = nan(length(options.start_index),1);
if(options.trace)
    parameters.MS.par_trace = nan(parameters.number,maxOptimSteps+1,length(options.start_index));
    parameters.MS.fval_trace = nan(maxOptimSteps+1,length(options.start_index));
    parameters.MS.time_trace = nan(maxOptimSteps+1,length(options.start_index));
end

% Define the negative log-posterior funtion
% (fmincon needs the neagtive log posterior for optimization)
negLogPost = setObjectiveWrapper(objective_function, options, 'negative log-posterior', [], [], true, true);

% Check, if Hessian should be used and if a Hessian function was set,
% otherwise use the third output of the objective function instead
if (strcmp(options.localOptimizer, 'fmincon') && ...
        strcmp(options.localOptimizerOptions.Hessian, 'on'))
    
    if (~isfield(options.localOptimizerOptions, 'HessFcn') ...
            || isempty(options.localOptimizerOptions.HessFcn))
        
        % this only works for box-constraints at the moment
        options.localOptimizerOptions.HessFcn = @(varargin) HessianWrap(negLogPost, varargin);
    end
end

waitbarFields1 = {'logPost', 'logPost0', 'n_objfun', 'n_iter', 't_cpu', 'exitflag'};
waitbarFields2 = {'par', 'par0', 'gradient', 'fval_trace', 'time_trace'};
waitbarFields3 = {'hessian', 'par_trace'};

%% Multi-start local optimization -- SEQUENTIAL
if strcmp(options.comp_type, 'sequential')
    
    % Matlab parallel toolbox seems to have problems with our outfun...
    if (strcmp(options.localOptimizer, 'fmincon') || strcmp(options.localOptimizer, 'lsqnonlin'))
        options.localOptimizerOptions.OutputFcn = @outfun_fmincon;
    end
    
    % initialize the waitbar
    if(strcmp(options.mode,'visual'))
        waitBar = waitbar(0, '1', 'name', 'Parameter estimation in process, please wait...', 'CreateCancelBtn', 'setappdata(gcbf, ''canceling'', 1)');
        stringTimePrediction = updateWaitBar(nan);
        waitbar(0, waitBar, stringTimePrediction);
        C = onCleanup(@() delete(waitBar));
    end
    
    % Loop: Multi-starts
    for iMS = 1 : length(options.start_index)
        % reset the objective function
        if(options.resetobjective)
            fun = functions(objective_function);
            s_start = strfind(fun.function,')')+1;
            s_end = strfind(fun.function,'(')-1;
            clear(fun.function(s_start(1):s_end(2)));
        end
        
        % Reset error count
        error_count = 0;
        
        % Test evaluation of objective function at starting point
        % Only for multi-start local, since other optimizers use a
        % different initialization
        if strcmp(options.localOptimizer, 'fmincon')
            if (strcmp(options.localOptimizerOptions.Hessian, 'on'))
                % Depending on the algorithm, the Hessian gets called
                % seperately (IP) or with the objective function (TR), so
                % different cases have to be checked.
                if strcmp(options.localOptimizerOptions.Algorithm, 'interior-point')
                    [J_0,~] = negLogPost(par0(:,iMS));
                else
                    [J_0,~,~] = negLogPost(par0(:,iMS));
                end
            elseif (strcmp(options.localOptimizerOptions.GradObj, 'on'))
                [J_0,~] = negLogPost(par0(:,iMS));
            else
                J_0 = negLogPost(par0(:,iMS));
            end
            parameters.MS.logPost0(iMS) = -J_0;
        elseif (strcmp(options.localOptimizer, 'lsqnonlin'))
            if (strcmp(options.localOptimizerOptions.Jacobian, 'on'))
                [J_0,~] = negLogPost(par0(:,iMS));
            else
                J_0 = negLogPost(par0(:,iMS));
            end
            parameters.MS.logPost0(iMS) = -sum(J_0);
            J_0 = sum(J_0);
        elseif (any(strcmp(options.localOptimizer, {'dhc','cs','bobyqa'})))
            J_0 = negLogPost(par0(:,iMS));
        end
        
        % Optimization
        startTimeLocalOptimization = cputime;
        if (isempty(J_0) || (J_0 < -options.init_threshold))
            
            %% do optimization with chosen algorithm
            
            switch options.localOptimizer
                case 'fmincon'
                    % fmincon as local optimizer
                    parameters = performOptimizationFmincon(parameters, negLogPost, iMS, par0, J_0, options);
                    
                case {'meigo-ess', 'meigo-vns'}
                    % Use the MEIGO toolbox as local / global optimizer
                    parameters = performOptimizationMeigo(parameters, negLogPost, iMS, options);
                    
                case 'pswarm'
                    % Optimization using a swarm based global optimizer PSwarm
                    parameters = performOptimizationPswarm(parameters, negLogPost, iMS, options);
                    
                case 'lsqnonlin'
                    % Optimization using dynamic hill climbin as local optimizer
                    parameters = performOptimizationLsqnonlin(parameters, negLogPost, iMS, par0, J_0, options);
                    
                case 'cs'
                    % Optimization using coordinate search as local optimizer
                    parameters = performOptimizationCs(parameters, negLogPost, iMS, par0, J_0, options);
                    
                case 'dhc'
                    % Optimization using dynamic hill climbing as local optimizer
                    parameters = performOptimizationDhc(parameters, negLogPost, iMS, par0, J_0, options);
                    
                case 'bobyqa'
                    % Optimization using bobya as local optimizer
                    parameters = performOptimizationBobyqa(parameters, negLogPost, iMS, par0, J_0, options);
            end
            
        end
        parameters.MS.t_cpu(iMS) = cputime - startTimeLocalOptimization;
        
        % Save
        if options.save
            saveResults(parameters,options,iMS)
        end
        
        % Output
        if strcmp(options.mode,'visual')
            fh = plotMultiStarts(parameters, fh, options.plot_options);
        elseif strcmp(options.mode,'text')
            fprintf('%d / %d:\t fval = %.15f, t = %.6f\n', ...
                iMS, length(options.start_index), parameters.MS.logPost(iMS), parameters.MS.t_cpu(iMS));
        end
        
        % Abort the calculation if the waitbar is cancelled
        if(strcmp(options.mode,'visual'))
            if getappdata(waitBar, 'canceling')
                parameters.MS.n_starts = iMS;
                for iWaitbarField = 1:6
                    parameters.MS.(waitbarFields1{iWaitbarField}) = ...
                        parameters.MS.(waitbarFields1{iWaitbarField})(1:iMS, :);
                end
                for iWaitbarField = 1:5
                    if (isfield(parameters.MS, waitbarFields2{iWaitbarField}))
                        parameters.MS.(waitbarFields2{iWaitbarField}) = ...
                            parameters.MS.(waitbarFields2{iWaitbarField})(:, 1:iMS);
                    end
                end
                for iWaitbarField = 1:2
                    if (isfield(parameters.MS, waitbarFields3{iWaitbarField}))
                        parameters.MS.(waitbarFields3{iWaitbarField}) = ...
                            parameters.MS.(waitbarFields3{iWaitbarField})(:, :, 1:iMS);
                    end
                end
                
                break;
            end
        end
        
        % update the waitbar
        if(strcmp(options.mode,'visual'))
            stringTimePrediction = updateWaitBar(nanmedian(parameters.MS.t_cpu(1:iMS)) * (length(options.start_index) - iMS));
            waitbar(iMS / length(options.start_index), waitBar, stringTimePrediction);
        end
    end
    
    % Order
    parameters = sortMultiStarts(parameters);
end

%% Multi-start local optimization -- PARALLEL
% TODO: adapt parallel mode to sequential mode
if strcmp(options.comp_type,'parallel')
    
    % Initialization
    par = nan(parameters.number,length(options.start_index));
    logPost0 = nan(length(options.start_index),1);
    logPost = nan(length(options.start_index),1);
    gradient = nan(parameters.number,length(options.start_index));
    hessian  = nan(parameters.number,parameters.number,length(options.start_index));
    n_objfun = nan(length(options.start_index),1);
    n_iter = nan(length(options.start_index),1);
    t_cpu = nan(length(options.start_index),1);
    exitflag = nan(length(options.start_index),1);
    
    % reset the objective function
    if(options.resetobjective)
        fun = functions(objective_function);
        s_start = strfind(fun.function,')')+1;
        s_end = strfind(fun.function,'(')-1;
        clear(fun.function(s_start(1):s_end(2)));
    end
    negLogPost = setObjectiveWrapper(objective_function, options, 'negative log-posterior', [], [], false, false);
    
    % Loop: Mutli-starts
    parfor iMS = options.start_index
        
        % Evaluation of objective function at starting point
        if (strcmp(options.localOptimizerOptions.Hessian, 'on'))
            % Depending on the algorithm, the Hessian gets called
            % seperately (IP) or with the objective function (TR), so
            % different cases have to be checked.
            if strcmp(options.localOptimizerOptions.Algorithm, 'interior-point')
                [J_0,~] = negLogPost(parameters.MS.par0(:,iMS));
            else
                [J_0,~,~] = negLogPost(parameters.MS.par0(:,iMS));
            end
        elseif (strcmp(options.localOptimizerOptions.GradObj, 'on'))
            [J_0,~] = negLogPost(parameters.MS.par0(:,iMS));
        else
            J_0 = negLogPost(parameters.MS.par0(:,iMS));
        end
        
        % Optimization
        startTimeLocalOptimization = cputime;
        if J_0 < -options.init_threshold
            % Optimization using fmincon
            [theta,J_opt,exitflag(iMS),results_fmincon,~,gradient_opt,hessian_opt] = ...
                fmincon(negLogPost,...  % negative log-posterior function
                parameters.MS.par0(:,iMS),...    % initial parameter
                parameters.constraints.A  ,parameters.constraints.b  ,... % linear inequality constraints
                parameters.constraints.Aeq,parameters.constraints.beq,... % linear equality constraints
                parameters.min,...     % lower bound
                parameters.max,...     % upper bound
                [],options.localOptimizerOptions);   % options
            
            % Assignment
            logPost(iMS) = -J_opt;
            par(:,iMS) = theta;
            gradient(:,iMS) = gradient_opt;
            if isempty(hessian_opt)
                if strcmp(options.localOptimizerOptions.Hessian,'on')
                    [~,~,hessian_opt] = objectiveWrap(theta,objective_function,options.obj_type,options.objOutNumber);
                end
            elseif max(abs(hessian_opt(:))) == 0
                if strcmp(options.localOptimizerOptions.Hessian,'on')
                    [~,~,hessian_opt] = objectiveWrap(theta,objective_function,options.obj_type,options.objOutNumber);
                end
            end
            hessian(:,:,iMS) = full(hessian_opt);
            n_objfun(iMS) = results_fmincon.funcCount;
            n_iter(iMS) = results_fmincon.iterations;
        end
        t_cpu(iMS) = cputime - startTimeLocalOptimization;
        
        % Save
        if options.save
            saveResults(parameters,options,iMS)
        end
        
        % Output
        switch options.mode
            case 'text', disp(['  ' num2str(iMS,'%d') '/' num2str(length(options.start_index),'%d')]);
            case {'silent','visual'} % no output
        end
    end
    
    % Assignment
    parameters.MS.par = par;
    parameters.MS.logPost0 = logPost0;
    parameters.MS.logPost = logPost;
    parameters.MS.gradient = gradient;
    parameters.MS.hessian  = hessian;
    parameters.MS.n_objfun = n_objfun;
    parameters.MS.n_iter = n_iter;
    parameters.MS.t_cpu = t_cpu;
    parameters.MS.exitflag = exitflag;
    parameters = sortMultiStarts(parameters);
    
    % Output
    switch options.mode
        case 'visual', fh = plotMultiStarts(parameters,fh,options.plot_options);
        case {'text','silent'} % no output
    end
    
end

%% Output
switch options.mode
    case {'visual','text'}, disp('-> Multi-start optimization FINISHED.');
    case 'silent' % no output
end

% Clear Output Function
options.localOptimizerOptions.OutputFcn = [];

%% Nested function for storing of objective function and parameter values
    function stop = outfun_fmincon(x,optimValues,state)
        
        switch state
            case 'init'
                % do nothing
            case 'interrupt'
                % do nothing
            case 'iter'
                if(options.trace)
                    parameters.MS.par_trace(:,optimValues.iteration+1,iMS) = x;
                    parameters.MS.time_trace(optimValues.iteration+1,iMS) = cputime - startTimeLocalOptimization;
                    if isfield(optimValues, 'fval')
                        parameters.MS.fval_trace(optimValues.iteration+1,iMS) = optimValues.fval;
                    else
                        parameters.MS.fval_trace(optimValues.iteration+1,iMS) = optimValues.resnorm;
                    end
                end
                if(options.tempsave)
                    if optimValues.iteration>0
                        if(mod(optimValues.iteration,10) == 0)
                            saveResults(parameters,options,iMS);
                        end
                    end
                end
            case 'done'
                % do nothing
        end
        
        if error_count <= 20
            stop = false;
        else
            warning('Too many failed objective function evaluations.')
            stop = true;
        end
        
    end
end


%% Waitbar Update
function stringTimePrediction = updateWaitBar(timePredicted)
% stringTimePrediction estimates the remaining time to display in the waitbar
%
% Parameters:
%  timePredicted: Predicted time in seconds
%
% Return values:
%  stringTimePrediction: String, Updating Message

if isnan(timePredicted)
    stringTimePrediction = 'Unknown.';
elseif (timePredicted < 60)
    stringTimePrediction = 'One minute or less...';
elseif (timePredicted >= 60 && timePredicted < 3600)
    stringTimePrediction = ['About ' num2str(round(timePredicted/60)) + 1 ' minutes'];
elseif (timePredicted >= 3600 && timePredicted < 72000)
    hours = floor(timePredicted/3600);
    minutes = round((timePredicted - 3600*hours) / 600) * 10;
    if (hours == 1)
        stringTimePrediction = ['About 1 hour'];
    else
        stringTimePrediction = ['About ' num2str(hours) ' hours'];
    end
    if (minutes == 0)
        stringTimePrediction = strcat(stringTimePrediction, '...');
    else
        stringTimePrediction = strcat(stringTimePrediction, [' and ' num2str(minutes) ' minutes...']);
    end
elseif (timePredicted >= 72000 && timePredicted < 36 * 3600)
    stringTimePrediction = 'Roughly 1 day...';
elseif (timePredicted >= 36 * 3600 && timePredicted < 2 * 365 * 24 * 3600)
    stringTimePrediction = ['About ' num2str(round(timePredicted / 24 * 3600)) ' days...'];
elseif (timePredicted >= 2 * 365 * 24 * 3600 && timePredicted < 100 * 365 * 24 * 3600)
    stringTimePrediction = ['Oh boy! Quite some years... Maybe about ' num2str(round(timePredicted / 365 * 24 * 3600)) ' of them...'];
elseif (timePredicted >= 100 * 365 * 24 * 3600 && timePredicted < 1e7 * 365 * 24 * 3600)
    stringTimePrediction = 'Well... Maybe your children, or grand-children... No, not evem them...';
else
    stringTimePrediction = 'Kingdoms will rise, civilization will decline, stars will fade - but your calculation...(!) ;)';
end
stringTimePrediction = ['Predicted remaining waiting time: ', stringTimePrediction];

end


%% Saving results
function saveResults(parameters,options,i)
% saveResults saves Multi-start results to disk
%
% Parameters:
%  parameters: Parameter struct passed to getMultiStarts
%  options: getMultiStarts options
%  i: multi-start index
dlmwrite(fullfile(pwd,options.foldername ,['MS' num2str(options.start_index(i),'%d') '__logPost.csv']),parameters.MS.logPost(i),'delimiter',',','precision',12);
dlmwrite(fullfile(pwd,options.foldername ,['MS' num2str(options.start_index(i),'%d') '__logPost0.csv']),parameters.MS.logPost0(i),'delimiter',',','precision',12);
dlmwrite(fullfile(pwd,options.foldername ,['MS' num2str(options.start_index(i),'%d') '__par.csv']),parameters.MS.par(:,i),'delimiter',',','precision',12);
dlmwrite(fullfile(pwd,options.foldername ,['MS' num2str(options.start_index(i),'%d') '__par0.csv']),parameters.MS.par0(:,i),'delimiter',',','precision',12);
dlmwrite(fullfile(pwd,options.foldername ,['MS' num2str(options.start_index(i),'%d') '__gradient.csv']),parameters.MS.gradient(:,i),'delimiter',',','precision',12);
dlmwrite(fullfile(pwd,options.foldername ,['MS' num2str(options.start_index(i),'%d') '__hessian.csv']),parameters.MS.hessian(:,:,i),'delimiter',',','precision',12);
dlmwrite(fullfile(pwd,options.foldername ,['MS' num2str(options.start_index(i),'%d') '__t_cpu.csv']),parameters.MS.t_cpu(i),'delimiter',',','precision',12);
dlmwrite(fullfile(pwd,options.foldername ,['MS' num2str(options.start_index(i),'%d') '__n_objfun.csv']),parameters.MS.n_objfun(i),'delimiter',',','precision',12);
dlmwrite(fullfile(pwd,options.foldername ,['MS' num2str(options.start_index(i),'%d') '__n_iter.csv']),parameters.MS.n_iter(i),'delimiter',',','precision',12);
dlmwrite(fullfile(pwd,options.foldername ,['MS' num2str(options.start_index(i),'%d') '__exitflag.csv']),parameters.MS.exitflag(i),'delimiter',',','precision',12);
if(options.trace)
    dlmwrite(fullfile(pwd,options.foldername ,['MS' num2str(options.start_index(i),'%d') '__par_trace.csv']),parameters.MS.par_trace(:,:,i),'delimiter',',','precision',12);
    dlmwrite(fullfile(pwd,options.foldername ,['MS' num2str(options.start_index(i),'%d') '__fval_trace.csv']),parameters.MS.fval_trace(:,i),'delimiter',',','precision',12);
    dlmwrite(fullfile(pwd,options.foldername ,['MS' num2str(options.start_index(i),'%d') '__time_trace.csv']),parameters.MS.time_trace(:,i),'delimiter',',','precision',12);
end

end
```
