## Supplementary material for "Mechanistic description of spatial processes using integrative modelling of noise-corrupted imaging data": getParameterConfidenceIntervals.m

```
function parameters = getParameterConfidenceIntervals(parameters, alpha, varargin)
% getParameterConfidenceIntervals() calculates the confidence intervals 
% for the model parameters. This is done by four approaches:
% The values of CI.local_PL and CI.PL are determined by the point on which 
% a threshold according to the confidence level alpha (calculated by a 
% chi2-distribution) is reached. local_PL computes this point by a local
% approximation around the MAP estimate using the Hessian matrix, PL uses 
% the profile likelihoods instead.
% The value of CI.local_B is computed by using the cummulative distribution
% function of a local approximation of the profile based on the Hessian
% matrix at the MAP estimate.
% The value of CI.S is calculated using samples for the model parameters
% and the according percentiles based on the confidence levels alpha.
%
% USAGE:
% * parameters = getParameterConfidenceIntervals(parameters, alpha)
%
% Parameters:
%   parameters: parameter struct
%   alpha: vector with desired confidence levels for the intervals
%   varargin: 
%    options: A PestoOptions instance
%
% Return values:
%   parameters: updated parameter struct
%
% Generated fields of parameters:
%   CI: Information about confidence levels
%     * local_PL: Threshold based approach, uses a local approximation by
%         the Hessian matrix at the MAP estimate
%         (requires parameters.MS, e.g. from getMultiStarts)
%     * PL: Threshold based approach, uses profile likelihoods 
%         (requires parameters.P, e.g. from getParameterProfiles)
%     * local_B: Mass based approach, uses a local approximation by
%         the Hessian matrix at the MAP estimate
%         (requires parameters.MS, e.g. from getMultiStarts)
%     * S: Bayesian approach, uses percentiles based on samples
%         (requires parameters.S, e.g. from getParameterSamples)
%
% History:
% * 2013/11/29 Jan Hasenauer
% * 2016/12/01 Paul Stapor

%% Checking and assigning inputs
if length(varargin) >= 1
    options = handleOptionArgument(varargin{1});
else
    options = PestoOptions();
end

% Maximum posterior index
iMAP = options.MAP_index;
if (isempty(iMAP))
    iMAP = 1;
end

% parameter index
if isempty(options.parameter_index)
    options.parameter_index = 1 : parameters.number;
end

% Initialization
parameters.CI.alpha_levels = alpha;

% Set default values for parameter profile 
if isfield(parameters,'P')
    for iPar = 1:parameters.number
        if iPar > length(parameters.P)
            parameters.P(iPar).par = [];
            parameters.P(iPar).logPost = [];
            parameters.P(iPar).R = [];
            parameters.P(iPar).t_cpu = [];
            parameters.P(iPar).optSteps = [];
            parameters.P(iPar).intSteps = [];
            parameters.P(iPar).reOptSteps = [];
        end
    end
end

% Loop: alpha levels
for k = 1:length(alpha)
    % Loop: Parameters
    for iPar = options.parameter_index
        if isfield(parameters,'MS')
            % Inversion of Hessian
            if isempty(options.fixedParameters)
                Sigma = pinv(parameters.MS.hessian(:,:,iMAP));
            else
                Sigma = nan(parameters.number);
                ind = setdiff(1:parameters.number,options.fixedParameters);
                Sigma(ind,ind) = pinv(parameters.MS.hessian(ind,ind,iMAP));
            end
            
            % Confidence intervals computed using local approximation and a
            % threshold (-> similar to PL-based confidence intervals)
            parameters.CI.local_PL(iPar,1,k) = parameters.MS.par(iPar,iMAP) - sqrt(icdf('chi2',alpha(k),1)*Sigma(iPar,iPar));
            parameters.CI.local_PL(iPar,2,k) = parameters.MS.par(iPar,iMAP) + sqrt(icdf('chi2',alpha(k),1)*Sigma(iPar,iPar));

            % Confidence intervals computed using local approximation and the
            % probability mass (-> similar to Bayesian confidence intervals)
            parameters.CI.local_B(iPar,1,k)  = icdf('norm',  (1-alpha(k))/2,parameters.MS.par(iPar,iMAP),sqrt(Sigma(iPar,iPar)));
            parameters.CI.local_B(iPar,2,k)  = icdf('norm',1-(1-alpha(k))/2,parameters.MS.par(iPar,iMAP),sqrt(Sigma(iPar,iPar)));
        end
        
        % Confidence intervals computed using profile likelihood
        if isfield(parameters,'P')
            for iPar = 1:length(parameters.P)
                if ~isempty(parameters.P(iPar).par)
                    % left bound
                    ind  = find(parameters.P(iPar).par(iPar,:) <= parameters.MS.par(iPar,iMAP));
                    j = find(parameters.P(iPar).R(ind) <= exp(-icdf('chi2',alpha(k),1)/2),1,'last');
                    if ~isempty(j)
                        parameters.CI.PL(iPar,1,k) = interp1(parameters.P(iPar).R(ind([j,j+1])),...
                            parameters.P(iPar).par(iPar,ind([j,j+1])),exp(-icdf('chi2',alpha(k),1)/2));
                    else
                        parameters.CI.PL(iPar,1,k) = -inf;
                    end
                    % right bound
                    ind  = find(parameters.P(iPar).par(iPar,:) >= parameters.MS.par(iPar,iMAP));
                    j = find(parameters.P(iPar).R(ind) <= exp(-icdf('chi2',alpha(k),1)/2),1,'first');
                    if ~isempty(j)
                        parameters.CI.PL(iPar,2,k) = interp1(parameters.P(iPar).R(ind([j-1,j])),...
                            parameters.P(iPar).par(iPar,ind([j-1,j])),exp(-icdf('chi2',alpha(k),1)/2));
                    else
                        parameters.CI.PL(iPar,2,k) = inf;
                    end
                else
                    parameters.CI.PL(iPar,[1,2],k) = nan(1,2);
                end
            end
        end
        
        % Confidence intervals computed using sample
        if isfield(parameters,'S')
            parameters.CI.S(iPar,:,k) = prctile(parameters.S.par(iPar,:,1),50 + 100*[-alpha(k)/2, alpha(k)/2]);
        end
    end
end

%% Output
switch options.mode
    case 'visual'
        plotConfidenceIntervals(parameters, alpha, [], options);
        disp('-> Calculation of confidence intervals for parameters FINISHED.');
    case 'text'
        disp('-> Calculation of confidence intervals for parameters FINISHED.');
    case 'silent' % no output
end

end
```
