## Supplementary material for "Mechanistic description of spatial processes using integrative modelling of noise-corrupted imaging data": getParameterProfiles.m

```
function [parameters,fh] = getParameterProfiles(parameters, objective_function, varargin)
% getParameterProfiles.m calculates the profiles likelihoods for the model 
% parameters, starting from the maximum a posteriori estimate. This
% calculation is done by fixing the i-th parameter and repeatedly
% reoptimizing the likelihood/posterior estimate (for all i). The initial 
% guess for the next reoptimization point is computed by extrapolation from
% the previous points to ensure a quick optimization.
%
% Note: This function can exploit up to (n_theta + 1) workers when running
% in 'parallel' mode.
%
% USAGE:
% [...] = getParameterProfiles(parameters, objective_function)
% [...] = getParameterProfiles(parameters, objective_function, options)
% [parameters, fh] = getParameterProfiles(...)
%
% getParameterProfiles() uses the following PestoOptions members:
%  * PestoOptions::calc_profiles
%  * PestoOptions::comp_type
%  * PestoOptions::dJ
%  * PestoOptions::dR_max
%  * PestoOptions::fh
%  * PestoOptions::MAP_index
%  * PestoOptions::mode
%  * PestoOptions::obj_type
%  * PestoOptions::options_getNextPoint .guess .min .max .update .mode
%  * PestoOptions::parameter_index
%  * PestoOptions::parameter_method_index
%  * PestoOptions::profile_method
%  * PestoOptions::profileReoptimizationOptions
%  * PestoOptions::plot_options
%  * PestoOptions::R_min
%  * PestoOptions::save
%
% Parameters:
%   parameters: parameter struct
%   objective_function: objective function to be optimized. 
%       This function should accept one input, the parameter vector.
%   varargin:
%     options: A PestoOptions object holding various options for the 
%         algorithm.
%
% Required fields of parameters:
%   number: Number of parameters
%   min: Lower bound for each parameter
%   max: upper bound for each parameter
%   name = {'name1', ...}: names of the parameters
%   MS: results of global optimization, obtained using for instance 
%       the routine 'getMultiStarts.m'. MS has to contain at least
%     * par: sorted list n_theta x n_starts of parameter estimates.
%          The first entry is assumed to be the best one.
%     * logPost: sorted list n_starts x 1 of of log-posterior values
%          corresponding to the parameters listed in .par.
%     * hessian: Hessian matrix (or approximation) at the optimal point
%
% Return values:
%   parameters: updated parameter struct
%   fh: figure handle
%
% Generated fields of parameters:
%   P(i): profile for i-th parameter
%     * par: MAPs along profile
%     * logPost: maximum log-posterior along profile
%     * R: ratio
%
% History:
% * 2012/05/16 Jan Hasenauer
% * 2014/06/12 Jan Hasenauer
% * 2016/10/04 Daniel Weindl
% * 2016/10/12 Paul Stapor

    %% Check and assign inputs
    if length(varargin) >= 1
        options = handleOptionArgument(varargin{1});
    else
        options = PestoOptions();
    end

    % Check if MultiStart was launched before
    if(~isfield(parameters, 'MS'))
        error('No information from optimization available. Please run getMultiStarts() before getParameterProfiles.');
    end

    % Check and assign options
    options.P.min = parameters.min;
    options.P.max = parameters.max;
    if (~isfield(options.profileReoptimizationOptions, 'MaxFunEvals') ...
                || isempty(options.profileReoptimizationOptions.MaxFunEvals)) 
        options.profileReoptimizationOptions.MaxFunEvals = 200 * parameters.number;
    end
    if (isempty(options.MAP_index))
        options.MAP_index = 1;
    end

    % Process, which profiles should be computed in which manner
    if strcmp(options.profile_method, 'default')
        if (isempty(options.profile_optim_index) && isempty(options.profile_integ_index))
            options.profile_method = 'optimization';
        elseif (~isempty(options.profile_optim_index) && isempty(options.profile_integ_index))
            options.profile_method = 'optimization';
        elseif (isempty(options.profile_optim_index) && ~isempty(options.profile_integ_index))
            options.profile_method = 'integration';
        elseif (~isempty(options.profile_optim_index) && ~isempty(options.profile_integ_index))
            options.profile_method = 'mixed';
        end
    end    
    if isempty(options.parameter_index)
        switch options.profile_method
            case 'optimization'
                if isempty(options.profile_optim_index)
                    options.profile_optim_index = 1:parameters.number;
                end
                if ~isempty(options.profile_integ_index)
                    error('Some profiles seem to be computed twice. Please redefine consistent options!');
                end

            case 'integration'
                if isempty(options.profile_integ_index)
                    options.profile_integ_index = 1:parameters.number;
                end
                if ~isempty(options.profile_optim_index)
                    error('Some profiles seem to be computed twice. Please redefine consistent options!');
                end

            case 'mixed'
                if (isempty(options.profile_optim_index) && isempty(options.profile_integ_index))
                    warning('You specified profile computation method to be "mixed", but did not specify the precise method. Doing optimization for all profiles now!');
                    options.profile_method = 'optimization';
                    options.profile_optim_index = 1:parameters.number;
                end
                if length(unique([options.profile_optim_index options.profile_integ_index])) < length([options.profile_optim_index options.profile_integ_index])
                    error('Some profiles seem to be computed twice. Please redefine consistent options!');
                end

            otherwise
                error('Unknown profile computationg method');
        end
        options.parameter_index = sort(unique([options.profile_optim_index options.profile_integ_index]));
    else
        switch options.profile_method
            case 'optimization'
                options.profile_optim_index = options.parameter_index;
                if ~isempty(options.profile_integ_index)
                    error('Some profiles seem to be computed twice. Please redefine consistent options!');
                end

            case 'integration'
                options.profile_integ_index = options.parameter_index;
                if ~isempty(options.profile_optim_index)
                    error('Some profiles seem to be computed twice. Please redefine consistent options!');
                end

            case 'mixed'
                if (length(unique([options.profile_optim_index, options.profile_integ_index])) ~= length([options.profile_optim_index, options.profile_integ_index])) ...
                        || (length([options.profile_optim_index, options.profile_integ_index]) ~= length(options.parameter_index))
                    error('Inconsistent settings for indices in profile calculation.');
                end
                options.parameter_index = sort(unique([options.profile_optim_index options.profile_integ_index]));

            otherwise
                error('Unknown profile computationg method');
        end
    end
    
    % Check that parameters for which profiles are computed are not fixed
    if ~isempty(options.profile_optim_index)
        if ~isempty(setdiff(options.fixedParameters,options.profile_optim_index))
            options.profile_optim_index = setdiff(options.profile_optim_index,options.fixedParameters);
            warning('Fixed parameters are removed from options.profile_optim_index.');
        end
    end
    if ~isempty(options.profile_integ_index)
        if ~isempty(setdiff(options.fixedParameters,options.profile_integ_index))
            options.profile_integ_index = setdiff(options.profile_integ_index,options.fixedParameters);
            warning('Fixed parameters are removed from options.profile_integ_index.');
        end
    end

    %% Initialization and figure generation
    fh = [];
    switch options.mode
        case 'visual'
            if isempty(options.fh)
                fh = figure('Name','getParameterProfiles');
            else
                fh = figure(options.fh);
            end
        case 'text'
            fprintf(' \nProfile likelihood caculation:\n===============================\n');
        case 'silent' % no output
            % Force fmincon to be silent.
            options.profileReoptimizationOptions.display = 'off';
    end

    %% Initialization of parameter struct
    for i = options.parameter_index
        parameters.P(i).par = parameters.MS.par(:,options.MAP_index);
        parameters.P(i).logPost = parameters.MS.logPost(options.MAP_index);
        parameters.P(i).R = exp(parameters.MS.logPost(options.MAP_index)-parameters.MS.logPost(1));
    end

    %% Preperation of folder
    if options.save
        [~,~,~] = mkdir(options.foldername);
        save([options.foldername '/init'],'parameters');
    end
    
    %% Profile calculation
    if options.calc_profiles
        switch options.profile_method
            case 'optimization'
                [parameters, fh] = getParProfilesByOptimization(parameters, objective_function, options,fh);

            case 'integration'
                [parameters, fh] = getParProfilesByIntegration(parameters, objective_function, options, fh);

            case 'mixed'
                if strcmp(options.comp_type,'sequential')
                    for j = options.parameter_index
                        currentOptions = options.copy();
                        if sum(j == options.profile_integ_index) == 1
                            currentOptions.profile_integ_index = j;
                            [parameters, fh] = getParProfilesByIntegration(parameters, objective_function, currentOptions, fh);
                        elseif sum(j == options.profile_integ_index) == 0
                            currentOptions.profile_optim_index = j;
                            [parameters, fh] = getParProfilesByOptimization(parameters, objective_function, currentOptions, fh);
                        else
                            error('Some really strange error for the profile calculation indices occured');
                        end
                    end

                elseif strcmp(options.comp_type,'parallel')
                    parfor j = options.parameter_index
                        currentOptions = options.copy();
                        if sum(j == options.profile_integ_index) == 1
                            currentOptions.profile_integ_index = j;
                            getParProfilesByIntegration(parameters, objective_function, currentOptions, fh);
                        elseif sum(j == options.profile_integ_index) == 0
                            currentOptions.profile_optim_index = j;
                            getParProfilesByOptimization(parameters, objective_function, currentOptions, fh);
                        else
                            error('Some really strange error for the profile calculation indices occured');
                        end
                    end

                    % Output
                    switch options.mode
                        case 'visual', fh = plotParameterProfiles(parameters,'1D',fh,options.parameter_index,options.plot_options);
                        case 'text' % no output
                        case 'silent' % no output
                    end
                end
        end

    end

    %% Output
    switch options.mode
        case {'visual','text'}, disp('-> Profile calculation for parameters FINISHED.');
        case 'silent' % no output
    end

end
```
