## Supplementary material for "Mechanistic description of spatial processes using integrative modelling of noise-corrupted imaging data": getParameterSamples.m

```
function parameters = getParameterSamples(parameters, objFkt, options)
   % getParameterSamples.m performs MCMC sampling of the posterior
   %   distribution. 
   %
   %   Note, the DRAM library routine tooparameters.minox is
   %   used internally. This function is capable of sampling with MH, AM,
   %   DRAM, MALA, PT and PHS. The sampling plotting routines should no longer
   %   be contained in here but as standalone scripts capable of using the
   %   resulting par.S.
   %
   % Parameters:
   %   parameters: parameter struct covering model options and results obtained by
   %               optimization, profiles and sampling. Optimization results
   %               can be used for initialization.
   %   objFkt: Objective function which measures the difference of model output and data
   %   options:   An options object holding various options for the
   %              sampling. Depending on the algorithm and particular flavor,
   %              different options must be set: For details, please visit
   %              PestoSamplingOptions.m
   %
   % Required fields of parameters:
   %  min: Lower parameter bounds
   %  max: Upper parameter bounds
   %  number: Number of parameters
   %
   % Return values:
   %  parameters: The provided parameters struct
   %
   % Generated fields of parameters:
   %  S: The obtained sampling results
   %
   % History:
   % * 2012/07/11 Jan Hasenauer
   % * 2015/04/29 Jan Hasenauer
   % * 2016/10/17 Benjamin Ballnus
   % * 2016/10/19 Daniel Weindl
   % * 2016/11/04 Paul Stapor
   % * 2017/02/01 Benjamin Ballnus
   
   
   
   %% Check and assign inputs, note that theta0 and sigma0 are always set manually outside this function
   options.MCMC = options.MCMC.checkDependentDefaults(parameters);
   
   %% Wrap objective function
   logPosterior = setObjectiveWrapper(objFkt, options, 'log-posterior', [], [], false, true);
   
   %% Selection of sampling procedure
   switch options.MCMC.samplingAlgorithm
      
      % DRAM
      case 'DRAM'
         parameters.S = performDRAM( logPosterior, parameters, options.MCMC );
         
         % MALA
      case 'MALA'
         parameters.S = performMALA( logPosterior, parameters, options.MCMC );
         
         % MH, AM and PT
      case 'PT'
         parameters.S = performPT( logPosterior, parameters, options.MCMC );
         
         % PHS
      case 'PHS'
         parameters.S = performPHS( logPosterior, parameters, options.MCMC );
   end
   
   %% Output
  	switch options.mode
        case 'visual'
            samplingPlottingOpt = PestoPlottingOptions();
            samplingPlottingOpt.S.plot_type = 1;
            samplingPlottingOpt.S.ind = 1;
            fh = figure('Name','plotParameterSamples - 1D');
            plotParameterSamples(parameters,'1D',fh,[],samplingPlottingOpt);
            fh = figure('Name','plotParameterSamples - 2D');
            plotParameterSamples(parameters,'2D',fh,[],samplingPlottingOpt);
            disp('-> Sampling FINISHED.');
        case 'text', disp('-> Sampling FINISHED.');
        case 'silent'
    end
   
end
```
