## Supplementary material for "Mechanistic description of spatial processes using integrative modelling of noise-corrupted imaging data": getPropertyConfidenceIntervals.m

```
function properties = getPropertyConfidenceIntervals(properties, alpha, varargin)
% getPropertyConfidenceIntervals.m calculates the confidence intervals 
% for the model properties. This is done by three approaches:
% The values of CI.local_PL and CI.PL are determined by the point on which 
% a threshold according to the confidence level alpha (calculated by a 
% chi2-distribution) is reached. local_PL computes this point by a local
% approximation around the MAP estimate using the Hessian matrix, PL uses 
% the profile likelihoods instead.
% The value of CI.local_B is computed by using the cummulative distribution
% function of a local approximation of the profile based on the Hessian
% matrix at the MAP estimate.
%
% USAGE:
% * properties = getPropertyConfidenceIntervals(properties, alpha)
%
% Parameters:
%   properties: property struct
%   alpha: vector with desired confidence levels for the intervals
%   varargin: 
%    options: A PestoOptions instance
%
% Return values:
%   properties: updated properties struct
%
% Generated fields of properties:
%   CI: Information about confidence levels
%     * local_PL: Threshold based approach, uses a local approximation by
%         the Hessian matrix at the MAP estimate
%         (requires parameters.MS, e.g. from getMultiStarts)
%     * PL: Threshold based approach, uses profile likelihoods 
%         (requires parameters.P, e.g. from getParameterProfiles)
%     * local_B: Mass based approach, uses a local approximation by
%         the Hessian matrix at the MAP estimate
%         (requires parameters.MS, e.g. from getMultiStarts)
%
% History:
% * 2013/11/29 Jan Hasenauer
% * 2016/12/01 Paul Stapor

%% Checking and assigning inputs
if length(varargin) >= 1
    options = handleOptionArgument(varargin{1});
else
    options = PestoOptions();
end

% Initialization
properties.CI.alpha_levels = alpha;

% Loop: alpha levels
for k = 1:length(alpha)
    % Loop: properties
    for i = 1:properties.number
        % Hessian
        Sigma = properties.MS.prop_Sigma(:,:,1);
        
        % Confidence intervals computed using local approximation and a
        % threshold (-> similar to PL-based confidence intervals)
        properties.CI.local_PL(i,1,k) = properties.MS.prop(i) - sqrt(icdf('chi2',alpha(k),1)*Sigma(i,i));
        properties.CI.local_PL(i,2,k) = properties.MS.prop(i) + sqrt(icdf('chi2',alpha(k),1)*Sigma(i,i));

        % Confidence intervals computed using local approximation and the
        % probability mass (-> similar to Bayesian confidence intervals)
        properties.CI.local_B(i,1,k)  = icdf('norm',  (1-alpha(k))/2,properties.MS.prop(i),sqrt(Sigma(i,i)));
        properties.CI.local_B(i,2,k)  = icdf('norm',1-(1-alpha(k))/2,properties.MS.prop(i),sqrt(Sigma(i,i)));

        % Confidence intervals computed using profile likelihood
        if isfield(properties,'P')
            if i <= length(properties.P)
                if ~isempty(properties.P(i).prop)
                    % left bound
                    ind  = find(properties.P(i).prop <= properties.MS.prop(i,1));
                    j = find(properties.P(i).R(ind) <= exp(-icdf('chi2',alpha(k),1)/2),1,'last');
                    if ~isempty(j)
                        properties.CI.PL(i,1,k) = interp1(properties.P(i).R(ind([j,j+1])),...
                            properties.P(i).prop(ind([j,j+1])),exp(-icdf('chi2',alpha(k),1)/2));
                    else
                        properties.CI.PL(i,1,k) = -inf;
                    end
                    % right bound
                    ind  = find(properties.P(i).prop >= properties.MS.prop(i,1));
                    j = find(properties.P(i).R(ind) <= exp(-icdf('chi2',alpha(k),1)/2),1,'first');
                    if ~isempty(j)
                        properties.CI.PL(i,2,k) = interp1(properties.P(i).R(ind([j-1,j])),...
                            properties.P(i).prop(ind([j-1,j])),exp(-icdf('chi2',alpha(k),1)/2));
                    else
                        properties.CI.PL(i,2,k) = inf;
                    end
                end
            end
        end
        
        % Confidence intervals computed using sample
        if isfield(properties,'S')
            properties.CI.S(i,:,k) = prctile(properties.S.prop(i,:),50 + 100*[-alpha(k)/2, alpha(k)/2]);
        end
    end
end

%% Output
switch options.mode
    case 'visual'
        plotConfidenceIntervals(properties, alpha, [], options);
        disp('-> Calculation of confidence intervals for properties FINISHED.');
    case 'text'
        disp('-> Calculation of confidence intervals for properties FINISHED.');
    case 'silent' % no output
end

end
```
