## Supplementary material for "Mechanistic description of spatial processes using integrative modelling of noise-corrupted imaging data": getPropertyMultiStarts.m

```
function [properties,fh] = getPropertyMultiStarts(properties, parameters, varargin)
% getPropertyMultiStarts.m evaluates the properties for the different
%   mutli-start results.
%
% USAGE:
% [...] = getPropertyMultiStarts(properties,parameters)
% [...] = getPropertyMultiStarts(properties,parameters,options)
% [parameters,fh] = getPropertyMultiStarts(...)
%
% getPropertyMultiStarts() uses the following PestoOptions members:
%  * PestoOptions::mode
%  * PestoOptions::fh
%  * PestoOptions::save
%  * PestoOptions::foldername
%  * PestoOptions::comp_type
%
% Parameters:
%   parameters: parameter struct containing at least:
%      MS: information about multi-start optimization
%   properties: property struct containing at least:
%     number: Number of properties
%     min: lower bound for property values       
%     max: upper bound for property values       
%     name = {'name1',...}: names of the properties 
%     function = {'function1',...}: functions to evaluate property  
%         values. These functions provide the values of the respective  
%         properties and the corresponding 1st and 2nd order
%         derivatives.
%   varargin:
%     options: A PestoOptions object holding the options for the algorithm.
%
% Return values:
%   properties: updated parameter object containing:
%     MS: properties for multi-start optimization results
%       * par(:,i): ith MAP
%       * logPost(i): log-posterior for ith MAP
%       * exitflag(i): exit flag of ith MAP
%       * prop(j,i): values of jth property for ith MAP
%       * prop_Sigma(:,:,i): covariance of properties for ith MAP
% fh: figure handle
%
% History:
% * 2015/03/03 Jan Hasenauer
% * 2016/04/10 Daniel Weindl

%% Check and assign inputs
if length(varargin) >= 1
    options = handleOptionArgument(varargin{1});
else
    options = PestoOptions();
end

properties = propertySanityCheck(properties);

%% Initialization and figure generation
fh = [];
switch options.mode
    case 'visual'
        if isempty(options.fh)
            fh = figure('Name','getPropertyMultiStarts');
        else
            fh = figure(options.fh);
        end
    case 'text'
        fprintf(' \nProperty evaluation:\n====================\n');
end

%% Initialization
properties.MS.par = parameters.MS.par;
properties.MS.logPost = parameters.MS.logPost;
properties.MS.exitflag = parameters.MS.exitflag;
properties.MS.prop = nan(properties.number,length(properties.MS.logPost));
properties.MS.prop_Sigma = nan(properties.number,properties.number,length(properties.MS.logPost));

%% Preperation of folder
if options.save
    rmdir(options.foldername,'s'); 
    mkdir(options.foldername);
    save([options.foldername '/properties_init'],'properties');
end

%% Evaluation of properties for multi-start results -- SEQUENTIAL
if strcmp(options.comp_type,'sequential')

% Loop: Multi-start results
for j = 1:length(properties.MS.logPost)
    % Loop: Properties
    G = nan(parameters.number,properties.number);
    if (~isnan(properties.MS.logPost(j)))
        for i = 1:properties.number
            try
                [properties.MS.prop(i,j),G(:,i)] = properties.function{i}(properties.MS.par(:,j));
            catch
                properties.MS.prop(i,j) = properties.function{i}(properties.MS.par(:,j));
            end
        end
        properties.MS.prop_Sigma(:,:,j) = G'*pinv(squeeze(parameters.MS.hessian(:,:,j)))*G;
    end
    
    % Save
    if options.save
        dlmwrite([options.foldername '/properties_MS' num2str(i,'%d') '__prop.csv'],properties.MS.prop(:,j),'delimiter',',','precision',12);
        dlmwrite([options.foldername '/properties_MS' num2str(i,'%d') '__prop_Sigma.csv'],properties.MS.prop_Sigma(:,:,j),'delimiter',',','precision',12);
    end
    
    % Output
    if (mod(j,100) == 0) || (j == length(properties.MS.logPost))
        str = ['Property evaluation for multi-start results completed to ' num2str(100*j/length(properties.MS.logPost),'%d') ' %'];
        switch options.mode
            case 'visual', fh = plotPropertyMultiStarts(properties,fh);
            case 'text', disp(str);
            case 'silent' % no output
        end
    end
end

end

%% Evaluation of properties for multi-start results -- PARALLEL
if strcmp(options.comp_type,'parallel')

% Initialization
prop = nan(properties.number,length(properties.MS.logPost));
prop_Sigma = nan(properties.number,properties.number,length(properties.MS.logPost));

% Create local partial copies of the parameter and the propertry struct
para_num = parameters.number;
para_hes = parameters.MS.hessian;
para_MS_par = parameters.MS.par;
prop_num = properties.number;
prop_fun = properties.function;
prop_logPost = properties.MS.logPost;
opt_save = options.save;
opt_folder = options.foldername;

% Loop: Multi-start results
parfor i = 1:length(prop_logPost)
    % Loop: Properties
    P = nan(prop_num, 1);
    G = nan(para_num, prop_num);
    if (~isnan(prop_logPost(i)))
        for j = 1 : prop_num
            try
                [P(j),G(:,j)] = prop_fun{j}(para_MS_par(:,i));
            catch
                P(j) = prop_fun{j}(para_MS_par(:,i));
            end
        end
        % Assignment
        prop(:,i) = P;
        prop_Sigma(:,:,i) = G' * pinv(squeeze(para_hes(:,:,i))) * G;
    end
    
    % Save
    if (opt_save)
        dlmwrite([opt_folder '/properties_MS' num2str(i,'%d') '__prop.csv'],prop(:,i),'delimiter',',','precision',12);
        dlmwrite([opt_folder '/properties_MS' num2str(i,'%d') '__prop_Sigma.csv'],prop_Sigma(:,:,i),'delimiter',',','precision',12);
    end
end

% Assignment
properties.MS.prop = prop;
properties.MS.prop_Sigma = prop_Sigma;

end

%% Output
switch options.mode
    case {'visual','text'}, disp('-> Property evaluation for multi-start results FINISHED.');
    case 'silent' % no output
end

end
```
