## Supplementary material for "Mechanistic description of spatial processes using integrative modelling of noise-corrupted imaging data": getPropertyProfiles.m

```
function [properties,fh] = getPropertyProfiles(properties, parameters, objective_function, varargin)
% getPropertyProfiles.m calculates the profiles of user-supplied property
% functions, starting from the maximum a posteriori estimate. This
% calculation is done by varying the value of each property function
% respectively, starting from the value of this function at the global
% optimum and by reoptimizing the likelihood/posterior estimate in each 
% variational step of the property. The initial guess for the next 
% reoptimization point is computed by extrapolation from the previous 
% points to ensure a quick optimization.
%
% Note: This function can exploit up to (n_theta + 1) workers when running
% in 'parallel' mode.
%
% USAGE:
% [...] = getPropertyProfiles(properties, parameters, objective_function)
% [...] = getPropertyProfiles(properties, parameters, objective_function, options)
% [parameters, fh] = getPropertyProfiles(...)
%
% % getPropertyProfiles() uses the following PestoOptions members:
%  * PestoOptions::boundary
%  * PestoOptions::calc_profiles
%  * PestoOptions::comp_type
%  * PestoOptions::dJ
%  * PestoOptions::dR_max
%  * PestoOptions::fh
%  * PestoOptions::fmincon
%  * PestoOptions::foldername
%  * PestoOptions::MAP_index
%  * PestoOptions::mode
%  * PestoOptions::obj_type
%  * PestoOptions::options_getNextPoint .guess .min .max .update .mode
%  * PestoOptions::plot_options
%  * PestoOptions::property_index
%  * PestoOptions::R_min
%  * PestoOptions::save
%
% Parameters:
%   properties: property struct
%   parameters: parameter struct
%   objective_function: objective function to be optimized. 
%       This function should accept one input, the parameter vector.
%   varargin:
%     options: A PestoOptions object holding various options for the 
%         algorithm.
%
% Required fields of properties:
%   number: Number of properties
%   min: Lower bound for each properties
%   max: upper bound for each properties    
%   name = {'name1', ...}: names of the properties   
%   function = {'function1', ...}: functions to evaluate property  
%       values. These functions provide the values of the respective  
%       properties and the corresponding 1st and 2nd order
%       derivatives.
%
% Required fields of parameters:
%   number: Number of parameters
%   min: Lower bound for each parameter
%   max: upper bound for each parameter    
%   name = {'name1', ...}: names of the parameters       
%   MS: results of global optimization, obtained using for instance 
%       the routine 'getMultiStarts.m'. MS has to contain at least
%     * par: sorted list n_theta x n_starts of parameter estimates.
%          The first entry is assumed to be the best one.
%     * logPost: sorted list n_starts x 1 of of log-posterior values
%          corresponding to the parameters listed in .par.
%     * hessian: Hessian matrix (or approximation) at the optimal point
% 
%
% Return values:
%   properties: updated property struct
%   fh: figure handle
%
% Generated fields of properties:
%   P(i): profile for i-th parameter
%     * prop: MAPs along profile
%     * par: MAPs along profile
%     * logPost: maximum log-posterior along profile
%     * R: ratio
%
% History:
% * 2012/03/02 Jan Hasenauer
% * 2016/04/10 Daniel Weindl
% * 2016/10/12 Paul Stapor

%% Check and assign inputs
if length(varargin) >= 1
    options = handleOptionArgument(varargin{1});
else
    options = PestoOptions();
end

% Check and assign options
%TODO
options.plot_options.mark_constraint = false;
options.P.min = parameters.min;
options.P.max = parameters.max;
options.MAP_index = 1;

% Warning if objective function gradient is not available
if ~strcmp(options.profileReoptimizationOptions.GradObj, 'on')
    warning('For efficient and reliable optimization, getPropertyProfiles.m requires gradient information.')
end

%% Initialization and figure generation
fh = [];
switch options.mode
    case 'visual'
        if isempty(options.fh)
            fh = figure('Name','getPropertyProfiles');
        else
            fh = figure(options.fh);
        end
    case 'text'
        fprintf(' \nProfile likelihood caculation:\n===============================\n');
    case 'silent' % no output
        % Force fmincon to be silent.
        options.profileReoptimizationOptions.Display = 'off';
end

% Check, if MultiStart was launched before
if(~isfield(parameters, 'MS'))
    error('No information from multi-start local optimization available. Please run getMultiStarts() before getParameterProfiles.');
end

% Check and assign options
options.P.min = properties.min;
options.P.max = properties.max;
if isempty(options.property_index)
    options.property_index = 1:properties.number;
end
if (isempty(options.MAP_index))
    options.MAP_index = 1;
end

options.profileReoptimizationOptions.algorithm = 'interior-point';
options.profileReoptimizationOptions.MaxIter = 400;
options.profileReoptimizationOptions.TolCon = 1e-4;
options.profileReoptimizationOptions.MaxFunEvals = 200*parameters.number;

%% Initialization of property struct
for i = options.property_index
    properties.P(i).prop = properties.MS.prop(i,options.MAP_index);
    properties.P(i).par = properties.MS.par(:,options.MAP_index);
    properties.P(i).logPost = properties.MS.logPost(options.MAP_index);
    properties.P(i).R = 1;
end
logPost_max = properties.MS.logPost(1);

%% Preperation of folder
if options.save
    [~,~,~] = mkdir(options.foldername);
    save([options.foldername '/init'],'properties');
end

%% Profile calculation -- SEQUENTIAL
if strcmp(options.comp_type,'sequential') && options.calc_profiles
    
    % Profile calculation
    for i = options.property_index
        % Initialization
        P_prop = properties.MS.prop(i,options.MAP_index);
        P_par = parameters.MS.par(:,options.MAP_index);
        P_logPost = parameters.MS.logPost(options.MAP_index);
        P_R = exp(parameters.MS.logPost(options.MAP_index)-parameters.MS.logPost(1));
        if isfield(parameters.MS,'exitflag')
            P_exitflag = parameters.MS.exitflag(options.MAP_index);
        else
            P_exitflag = NaN;
        end
                
        if ((P_prop <= properties.min(i)) || (properties.max(i) <= P_prop)) && ~strcmp(options.mode,'silent')
            warning(['MAP of ' num2str(i) ordstr(i) ' property not between respective minimum and maximum.']);
        end
        
        % Compute profile for in- and decreasing property
        for s = [-1,1]
            % Starting point
            prop = properties.MS.prop(i,options.MAP_index);
            theta  = parameters.MS.par(:,options.MAP_index);
            logPost = parameters.MS.logPost(options.MAP_index);
            
            computeProfile = (logPost >= (log(options.R_min) + parameters.MS.logPost(1))) && ...
                    (prop > (properties.min(i)+options.boundary_tol)) && ...
                    ((properties.max(i)-options.boundary_tol) > prop);
                
            % Sequential update    
            while computeProfile
                
                % Proposal of next profile point
                J_exp = -(log(1-options.dR_max)+options.dJ*(logPost-logPost_max)+logPost);

                % Optimization
                [theta,prop,exitflag] = ...
                    fmincon(@(theta) prop_fun(theta,properties.function{i},properties.min(i),properties.max(i),s),...
                                        theta,...
                                        parameters.constraints.A  ,parameters.constraints.b  ,... % linear inequality constraints
                                        parameters.constraints.Aeq,parameters.constraints.beq,... % linear equality constraints
                                        parameters.min,...   % lower bound
                                        parameters.max,...   % upper bound
                                        @(theta) obj_con(theta,objective_function,-J_exp,options.obj_type),...
                                        options.profileReoptimizationOptions);    % options

                % Adaptation of signs                    
                if s == +1
                    prop = -prop;
                end

                % Reoptimization at boundary
                if (prop <= properties.min(i)) || (properties.max(i) <= prop)
                    [theta,J_opt] = ...
                        fmincon(@(theta) obj(theta,objective_function,options.obj_type),...
                                            theta,...
                                            parameters.constraints.A  ,parameters.constraints.b  ,... % linear inequality constraints
                                            parameters.constraints.Aeq,parameters.constraints.beq,... % linear equality constraints
                                            parameters.min,...   % lower bound
                                            parameters.max,...   % upper bound
                                            @(theta) prop_con_fun(theta,properties.function{i},properties.min(i),properties.max(i),s),...
                                            options.profileReoptimizationOptions);    % options
                else
                    J_opt = obj(theta,objective_function,options.obj_type);
                end
                
                % Assignment of log-posterior
                logPost = -J_opt;
                
                % Sorting
                switch s
                    case -1
                        P_prop = [prop,P_prop];
                        P_par = [theta,P_par];
                        P_logPost = [logPost,P_logPost];
                        P_R = [exp(logPost - parameters.MS.logPost(1)),P_R];
                        P_exitflag = [exitflag,P_exitflag];
                    case +1
                        P_prop = [P_prop,prop];
                        P_par = [P_par,theta];
                        P_logPost = [P_logPost,logPost];
                        P_R = [P_R,exp(logPost - parameters.MS.logPost(1))];
                        P_exitflag = [P_exitflag,exitflag];
                end
                
                % Assignment
                properties.P(i).prop = P_prop;
                properties.P(i).par = P_par;
                properties.P(i).logPost = P_logPost;
                properties.P(i).R = P_R;
                properties.P(i).exitflag = P_exitflag;
                
                % Save
                if options.save
                    dlmwrite([options.foldername '/properties_P' num2str(i,'%d') '__prop.csv'],P_prop,'delimiter',',','precision',12);
                    dlmwrite([options.foldername '/properties_P' num2str(i,'%d') '__par.csv'],P_par,'delimiter',',','precision',12);
                    dlmwrite([options.foldername '/properties_P' num2str(i,'%d') '__logPost.csv'],P_logPost,'delimiter',',','precision',12);
                    dlmwrite([options.foldername '/properties_P' num2str(i,'%d') '__R.csv'],P_R,'delimiter',',','precision',12);
                    dlmwrite([options.foldername '/properties_P' num2str(i,'%d') '__exitflag.csv'],P_exitflag,'delimiter',',','precision',12);
                end
                
                % Output
                str = [num2str(i,'%d') ordstr(i) ' P: point ' num2str(length(properties.P(i).R)-1,'%d') ', R = ' ...
                    num2str(exp(- J_opt - properties.MS.logPost(1)),'%.3e')];
                switch options.mode
                    case 'visual', fh = plotPropertyProfiles(properties,'1D',fh,options.property_index,options.plot_options);
                    case 'text', disp(str);
                    case 'silent' % no output
                end
                
                % Condition for the while-loop
                computeProfile = (logPost >= (log(options.R_min) + parameters.MS.logPost(1))) && ...
                    (prop > (properties.min(i)+options.boundary_tol)) && ...
                    ((properties.max(i)-options.boundary_tol) > prop);
            end
        end
    end
elseif strcmp(options.comp_type,'parallel') && options.calc_profiles
    %% Profile calculation -- PARALLEL
    
    % Assignement of profile
    P = properties.P;
    
    % Profile calculation
    parfor i = options.property_index
        % Initialization
        P_prop = properties.MS.prop(i,options.MAP_index);
        P_par = parameters.MS.par(:,options.MAP_index);
        P_logPost = parameters.MS.logPost(options.MAP_index);
        P_R = exp(parameters.MS.logPost(options.MAP_index)-parameters.MS.logPost(1));
        if isfield(parameters.MS,'exitflag')
            P_exitflag = parameters.MS.exitflag(options.MAP_index);
        else
            P_exitflag = NaN;
        end
                
        if ((P_prop <= properties.min(i)) || (properties.max(i) <= P_prop)) && ~strcmp(options.mode,'silent')
            warning(['MAP of ' num2str(i) ordstr(i) ' property not between respective minimum and maximum.']);
        end
        
        % Compute profile for in- and decreasing property
        for s = [-1,1]
            % Starting point
            prop = properties.MS.prop(i,options.MAP_index);
            theta  = parameters.MS.par(:,options.MAP_index);
            logPost = parameters.MS.logPost(options.MAP_index);
            
            computeProfile = (logPost >= (log(options.R_min) + parameters.MS.logPost(1))) && ...
                    (prop > (properties.min(i)+options.boundary_tol)) && ...
                    ((properties.max(i)-options.boundary_tol) > prop);
                
            % Sequential update
            while computeProfile
                
                % Proposal of next profile point
                J_exp = -(log(1-options.dR_max)+options.dJ*(logPost-logPost_max)+logPost);
        
                % Optimization
                [theta,prop,exitflag] = ...
                    fmincon(@(theta) prop_fun(theta,properties.function{i},properties.min(i),properties.max(i),s),...
                    theta,...
                    parameters.constraints.A  ,parameters.constraints.b  ,... % linear inequality constraints
                    parameters.constraints.Aeq,parameters.constraints.beq,... % linear equality constraints
                    parameters.min,...   % lower bound
                    parameters.max,...   % upper bound
                    @(theta) obj_con(theta,objective_function,-J_exp,options.obj_type),...
                    options.profileReoptimizationOptions);    % options
                
                % Adaptation of signs
                if s == +1
                    prop = -prop;
                end
                
                % Reoptimization at boundary
                if (prop <= properties.min(i)) || (properties.max(i) <= prop)
                    [theta,J_opt] = ...
                        fmincon(@(theta) obj(theta,objective_function,options.obj_type),...
                        theta,...
                        parameters.constraints.A  ,parameters.constraints.b  ,... % linear inequality constraints
                        parameters.constraints.Aeq,parameters.constraints.beq,... % linear equality constraints
                        parameters.min,...   % lower bound
                        parameters.max,...   % upper bound
                        @(theta) prop_con_fun(theta,properties.function{i},properties.min(i),properties.max(i),s),...
                        options.profileReoptimizationOptions);    % options
                else
                    J_opt = obj(theta,objective_function,options.obj_type);
                end
                
                % Assignment of log-posterior
                logPost = -J_opt;
                
                % Sorting
                switch s
                    case -1
                        P_prop = [prop,P_prop];
                        P_par = [theta,P_par];
                        P_logPost = [logPost,P_logPost];
                        P_R = [exp(logPost - parameters.MS.logPost(1)),P_R];
                        P_exitflag = [exitflag,P_exitflag];
                    case +1
                        P_prop = [P_prop,prop];
                        P_par = [P_par,theta];
                        P_logPost = [P_logPost,logPost];
                        P_R = [P_R,exp(logPost - parameters.MS.logPost(1))];
                        P_exitflag = [P_exitflag,exitflag];
                end
                
                % Assignment
                P(i).prop = P_prop;
                P(i).par = P_par;
                P(i).logPost = P_logPost;
                P(i).R = P_R;
                P(i).exitflag = P_exitflag;
                
                % Save
                if options.save
                    dlmwrite([options.foldername '/property_P' num2str(i,'%d') '__prop.csv'],P_prop,'delimiter',',','precision',12);
                    dlmwrite([options.foldername '/property_P' num2str(i,'%d') '__par.csv'],P_par,'delimiter',',','precision',12);
                    dlmwrite([options.foldername '/property_P' num2str(i,'%d') '__logPost.csv'],P_logPost,'delimiter',',','precision',12);
                    dlmwrite([options.foldername '/property_P' num2str(i,'%d') '__R.csv'],P_R,'delimiter',',','precision',12);
                    dlmwrite([options.foldername '/property_P' num2str(i,'%d') '__exitflag.csv'],P_exitflag,'delimiter',',','precision',12);
                end
                
                % Condition for the while-loop
                computeProfile = (logPost >= (log(options.R_min) + parameters.MS.logPost(1))) && ...
                    (prop > (properties.min(i)+options.boundary_tol)) && ...
                    ((properties.max(i)-options.boundary_tol) > prop);
            end
        end
    end
    
    % Assignment
    properties.P = P;
    
    % Output
    switch options.mode
        case 'visual', fh = plotPropertyProfiles(properties,'1D',fh,options.property_index,options.plot_options);
        case 'text' % no output
        case 'silent' % no output
    end
    
end

%% Output
switch options.mode
    case {'visual','text'}, disp('-> Profile calculation for properties FINISHED.');
    case 'silent' % no output
end

end


%% Objetive function interface
% This function is used as interface to the user-provided objective
% function. It adapts the sign and supplies the correct number of outputs.
% Furthermore, it catches errors in the user-supplied objective function.
%   theta ... parameter vector
%   fun ... user-supplied objective function
%   type ... type of user-supplied objective function
function varargout = obj(theta,fun,type)

try
    switch nargout
        case {0,1}
            J = fun(theta);
            if isnan(J)
                error('J is NaN.')
            end
            switch type
                case 'log-posterior'          , varargout = {-J};
                case 'negative log-posterior' , varargout = { J};
            end
        case 2
            [J,G] = fun(theta);
            if max(isnan([J;G(:)]))
                error('J and/or G contain a NaN.')
            end
            switch type
                case 'log-posterior'          , varargout = {-J,-G};
                case 'negative log-posterior' , varargout = { J, G};
            end
        case 3
            [J,G,H] = fun(theta);
            if max(isnan([J;G(:);H(:)]))
                error('J, G and/or H contain a NaN.')
            end
            switch type
                case 'log-posterior'          , varargout = {-J,-G,-H};
                case 'negative log-posterior' , varargout = { J, G, H};
            end
    end
catch
    switch nargout
        case {0,1}
            varargout = {inf};
        case 2
            varargout = {inf,zeros(length(theta),1)};
        case 3
            varargout = {inf,zeros(length(theta),1),zeros(length(theta))};
    end
end

end

%% Constrained objetive function interface
% This function is used as interface to the user-provided objective
% function. It adapts the sign and supplies the correct number of outputs.
% Furthermore, it catches errors in the user-supplied objective function.
%   theta ... parameter vector
%   fun ... user-supplied objective function
%   fun_min ... minimum objective function
%   type ... type of user-supplied objective function
function varargout = obj_con(theta,fun,fun_min,type)

try
    switch nargout
        case {0,1}
            J = fun(theta);
            if isnan(J)
                error('J is NaN.')
            end
            switch type
                case 'log-posterior'          , varargout = {fun_min-J};
                case 'negative log-posterior' , varargout = {fun_min+J};
            end
        case 2
            J = fun(theta);
            if isnan(J)
                error('J is NaN.')
            end
            switch type
                case 'log-posterior'          , varargout = {fun_min-J,[]};
                case 'negative log-posterior' , varargout = {fun_min+J,[]};
            end
        case 3
            [J,G] = fun(theta);
            if max(isnan([J;G(:)]))
                error('J and/or G contain a NaN.')
            end
            switch type
                case 'log-posterior'          , varargout = {fun_min-J,[],-G};
                case 'negative log-posterior' , varargout = {fun_min+J,[], G};
            end
        case 4
            [J,G] = fun(theta);
            if max(isnan([J;G(:)]))
                error('J and/or G contain a NaN.')
            end
            switch type
                case 'log-posterior'          , varargout = {fun_min-J,[],-G,[]};
                case 'negative log-posterior' , varargout = {fun_min+J,[], G,[]};
            end
    end
catch
    switch nargout
        case {0,1}
            varargout = {inf};
        case 2
            varargout = {inf,[]};
        case 3
            varargout = {inf,[],zeros(length(theta),1)};
        case 4
            varargout = {inf,[],zeros(length(theta),1),[]};
    end
end

end

%% Property function interface
% This function is used as interface to the user-provided property
% function. It adapts the sign and supplies the correct number of outputs.
% Furthermore, it catches errors in the user-supplied objective function.
%   theta ... parameter vector
%   fun ... user-supplied property function
%   prop_min ... minumum property value of interest (= profile boundary)
%   prop_max ... maximum property value of interest (= profile boundary)
%   s ... compute profile for increasing (s = +1) and decreasing (s = -1) property
function varargout = prop_fun(theta,fun,prop_min,prop_max,s)

if s == -1
    try
        switch nargout
            case {0,1}
                prop = fun(theta);
                if prop < prop_min
                    prop = prop_min;
                end
                if isnan(prop)
                    error('prop is NaN.')
                end
                varargout = {prop};
            case 2
                [prop,propG] = fun(theta);
                if prop < prop_min
                    prop = prop_min;
                    propG = zeros(size(propG));
                end
                if max(isnan([prop;propG(:)]))
                    error('prop and/or propG contain a NaN.')
                end
                varargout = {prop,propG};
            case 3
                [prop,propG,propH] = fun(theta);
                if prop < prop_min
                    prop = prop_min;
                    propG = zeros(size(propG));
                    propH = zeros(size(propH));
                end
                if max(isnan([prop;propG(:);propH(:)]))
                    error('prop, propG and/or propH contain a NaN.')
                end
                varargout = {prop,propG,propH};
        end
    catch
        switch nargout
            case {0,1}
                varargout = {inf};
            case 2
                varargout = {inf,zeros(length(theta),1)};
            case 3
                varargout = {inf,zeros(length(theta),1),zeros(length(theta))};
        end
    end
elseif s == +1
    try
        switch nargout
            case {0,1}
                prop = fun(theta);
                if prop > prop_max
                    prop = prop_max;
                end
                if isnan(prop)
                    error('prop is NaN.')
                end
                varargout = {-prop};
            case 2
                [prop,propG] = fun(theta);
                if prop > prop_max
                    prop = prop_max;
                    propG = zeros(size(propG));
                end
                if max(isnan([prop;propG(:)]))
                    error('prop and/or propG contain a NaN.')
                end
                varargout = {-prop,-propG};
            case 3
                [prop,propG,propH] = fun(theta);
                if prop > prop_max
                    prop = prop_max;
                    propG = zeros(size(propG));
                    propH = zeros(size(propH));
                end
                if max(isnan([prop;propG(:);propH(:)]))
                    error('prop, propG and/or propH contain a NaN.')
                end
                varargout = {-prop,-propG,-propH};
        end
    catch
        switch nargout
            case {0,1}
                varargout = {inf};
            case 2
                varargout = {inf,zeros(length(theta),1)};
            case 3
                varargout = {inf,zeros(length(theta),1),zeros(length(theta))};
        end
    end    
end

end

%% Property constraint function interface
% This function is used as interface to the user-provided property
% function. It adapts the sign and supplies the correct number of outputs.
% Furthermore, it catches errors in the user-supplied objective function.
%   theta ... parameter vector
%   fun ... user-supplied property function
%   prop_min ... minumum property value of interest (= profile boundary)
%   prop_max ... maximum property value of interest (= profile boundary)
%   s ... compute profile for increasing (s = +1) and decreasing (s = -1) property
function varargout = prop_con_fun(theta,fun,prop_min,prop_max,s)

if s == -1
    try
        switch nargout
            case {0,1}
                prop = fun(theta);
                if isnan(prop)
                    error('prop is NaN.')
                end
                varargout = {prop-prop_min};
            case 2
                prop = fun(theta);
                if isnan(prop)
                    error('prop is NaN.')
                end
                varargout = {prop-prop_min,[]};
            case 3
                [prop,propG] = fun(theta);
                if max(isnan([prop;propG(:)]))
                    error('prop and/or propG contain a NaN.')
                end
                varargout = {prop-prop_min,[],propG};
            case 4
                [prop,propG] = fun(theta);
                if max(isnan([prop;propG(:)]))
                    error('prop and/or propG contain a NaN.')
                end
                varargout = {prop-prop_min,[],propG,[]};
        end
    catch
        switch nargout
            case {0,1}
                varargout = {inf};
            case 2
                varargout = {inf,[]};
            case 3
                varargout = {inf,[],zeros(length(theta),1)};
            case 4
                varargout = {inf,[],zeros(length(theta),1),[]};
        end
    end
elseif s == +1
    try
        switch nargout
            case {0,1}
                prop = fun(theta);
                if isnan(prop)
                    error('prop is NaN.')
                end
                varargout = {prop_max-prop};
            case 2
                prop = fun(theta);
                if isnan(prop)
                    error('prop is NaN.')
                end
                varargout = {prop_max-prop,[]};
            case 3
                [prop,propG] = fun(theta);
                if max(isnan([prop;propG(:)]))
                    error('prop and/or propG contain a NaN.')
                end
                varargout = {prop_max-prop,[],-propG};
            case 4
                [prop,propG] = fun(theta);
                if max(isnan([prop;propG(:)]))
                    error('prop and/or propG contain a NaN.')
                end
                varargout = {prop_max-prop,[],-propG,[]};
        end
    catch
        switch nargout
            case {0,1}
                varargout = {inf};
            case 2
                varargout = {inf,[]};
            case 3
                varargout = {inf,[],zeros(length(theta),1)};
            case 4
                varargout = {inf,[],zeros(length(theta),1),[]};
        end
    end
end

end
```
