## Supplementary material for "Mechanistic description of spatial processes using integrative modelling of noise-corrupted imaging data": getPropertySamples.m

```
function [properties,fh] = getPropertySamples(properties, parameters, varargin)
% getPropertySamples.m evaluates the properties for the sampled parameters.
%
% USAGE:
% [...] = getPropertySamples(properties,parameters)
% [...] = getPropertySamples(properties,parameters,options)
% [parameters,fh] = getPropertySamples(...)
%
% getPropertySamples() uses the following PestoOptions members:
%  * PestoOptions::property_index
%  * PestoOptions::mode
%  * PestoOptions::fh
%  * PestoOptions::save
%  * PestoOptions::foldername
%  * PestoOptions::comp_type
%  * PestoOptions::plot_options
%  * PestoOptions::MCMC.thinning
%
% Parameters:
%   properties: property struct
%   parameters: parameter struct
%   varargin:
%     options: A PestoOptions object holding various options for the 
%         algorithm.
%
% Required fields of properties:
%   number: number of parameter
%   min: lower bound for property values       
%   max: upper bound for property values       
%   name: = {'name1',...} ... names of the parameters       
%   function: = {'function1',...} ... functions to evaluate property  
%       values. These functions provide the values of the respective  
%       properties and the corresponding 1st and 2nd order
%       derivatives.
%
% Required fields of parameters:
%   S: parameter and posterior sample.
%     logPost ... log-posterior function along chain
%     par  ... parameters along chain
%   *Note* This struct is obtained using getSamples.m.
%
% Return values:
%   properties: updated parameter object
%   fh: figure handle
%
% Generated fields of properties:
%  S: properties for sampling results
%    * par(*,i): ith samples parameter vector
%    * logPost(i): log-posterior for ith samples parameter vector
%    * prop(j,i): values of jth property for ith samples parameter vector
%    * prop_Sigma(*,*,i): covariance of properties for ith samples 
%          parameter vector
%
% History:
% * 2015/04/01 Jan Hasenauer
% * 2016/10/04 Daniel Weindl

%% Check and assign inputs
if length(varargin) >= 1
    options = handleOptionArgument(varargin{1});
else
    options = PestoOptions();
end

properties = propertySanityCheck(properties);

% Check initial guess
if ~isfield(parameters,'guess')
    parameters.guess = [];
end

% Check and assign options
options.property_index = 1:properties.number;

%% Initialization and figure generation
fh = [];
switch options.mode
    case 'visual'
        if isempty(options.fh)
            fh = figure('Name','getPropertySamples');
        else
            fh = figure(options.fh);
        end
    case 'text'
        fprintf(' \nProperty evaluation:\n====================\n');
end

%% Initialization
properties.S.par = parameters.S.par;
properties.S.logPost = parameters.S.logPost;
properties.S.prop = nan(properties.number,length(properties.S.logPost));

%% Preperation of folder
if options.save
    rmdir(options.foldername,'s'); 
    mkdir(options.foldername);
    save([options.foldername '/properties_init'],'properties');
end

%% Evaluation of properties for multi-start results -- SEQUENTIAL
if strcmp(options.comp_type,'sequential')

    % Loop: Multi-start results
    for j = 1:length(properties.S.logPost)
        % Loop: Properties
        for i = options.property_index
            properties.S.prop(i,j) = properties.function{i}(properties.S.par(:,j));
        end

        % Save
        if options.save
            dlmwrite([options.foldername '/properties_S' num2str(i,'%d') '__prop.csv'],properties.S.prop(:,j),'delimiter',',','precision',12);
        end

        % Output
        if (mod(j,ceil(length(properties.S.logPost)/10)) == 0) || (j == length(properties.S.logPost))
            str = ['Property evaluation for MCMC sampling completed to ' num2str(100*j/length(properties.S.logPost),'%d') ' %'];
            switch options.mode
                case 'visual', fh = plotPropertySamples(properties,'1D',fh,options.property_index,options.plot_options);
                case 'text', disp(str);
                case 'silent' % no output
            end
        end
    end

    % Output
    switch options.mode
        % Set the correct options        
        case 'visual' 
            options.plot_options.S.plot_type = 1;
            fh = plotPropertySamples(properties,'1D',fh,options.property_index,options.plot_options);
    end

end

%% Evaluation of properties for multi-start results -- PARALLEL
if strcmp(options.comp_type, 'parallel')

    % Initialization
    prop = nan(properties.number,length(properties.S.logPost));

    % Create local partial copies of the propertry struct
    prop_num = properties.number;
    prop_fun = properties.function;
    prop_S_par = properties.S.par;
    opt_save = options.save;
    opt_ind = options.property_index;
    opt_folder = options.foldername;

    % Loop: Multi-start results
    parfor i = 1:length(properties.S.logPost)
        % Loop: Properties
        P = nan(prop_num, 1);
        for j = opt_ind
            P(j) = prop_fun{j}(prop_S_par(:,i));
        end
        prop(:,i) = P;

        % Save
        if (opt_save)
            dlmwrite([opt_folder '/properties_S' num2str(i,'%d') '__prop.csv'],prop(:,i),'delimiter',',','precision',12);
        end
    end

    % Assignment
    properties.S.prop = prop;

end

%% Output
switch options.mode
    case {'visual','text'}, disp('-> Property evaluation for samples FINISHED.');
    case 'silent' % no output
end

end
```
