## Supplementary material for "Mechanistic description of spatial processes using integrative modelling of noise-corrupted imaging data": getReducedProblem.m

```
function [newParameters,newProperties,newObjectiveFunction,newOptions] = getReducedProblem(parameters,properties,objectiveFunction,options)

    % Definition of index set of optimized parameters
    freePars = setdiff(1:parameters.number, options.fixedParameters);
    fixedPars = options.fixedParameters;
    fixedParValues = options.fixedParameterValues;
   
    %% Reduced parameter struct
    % Basic information
    newParameters.name = parameters.name(freePars);
    newParameters.min = parameters.min(freePars);
    newParameters.max = parameters.max(freePars);
    newParameters.guess = parameters.guess(freePars);
    newParameters.number = length(freePars);
    if isfield(parameters, 'constraints')
        if isfield(parameters.constraints, 'A')
            if ~isempty(parameters.constraints.A)
                newParameters.constraints.A = parameters.constraints.A(:,freePars);
            else
                newParameters.constraints.A = [];
            end
        else
            newParameters.constraints.A = [];
        end

        if isfield(parameters.constraints, 'b')
            if ~isempty(parameters.constraints.b)
                newParameters.constraints.b = parameters.constraints.b - parameters.constraints.A(:,options.fixedParameters)*options.fixedParameterValues;
            else
                newParameters.constraints.b = [];
            end
        else
            newParameters.constraints.A = [];
        end

        if isfield(parameters.constraints, 'Aeq')
            if ~isempty(parameters.constraints.Aeq)
                newParameters.constraints.Aeq = parameters.constraints.Aeq(:,freePars);
            else
                newParameters.constraints.Aeq = [];
            end
        else
            newParameters.constraints.Aeq = [];
        end

        if isfield(parameters.constraints, 'beq')
            if ~isempty(parameters.constraints.beq)
                newParameters.constraints.beq = parameters.constraints.beq - parameters.constraints.Aeq(:,options.fixedParameters)*options.fixedParameterValues;
            else
                newParameters.constraints.beq = [];
            end
        else
            newParameters.constraints.beq = [];
        end
    end

    % Multi-start optimization results
    if isfield(parameters,'MS')
        newParameters.MS.par0 = parameters.MS.par0(freePars, :);
        newParameters.MS.par = parameters.MS.par(freePars, :);
        newParameters.MS.logPost0 = parameters.MS.logPost0;
        newParameters.MS.logPost = parameters.MS.logPost;
        newParameters.MS.gradient = parameters.MS.gradient(freePars, :);
        newParameters.MS.hessian = parameters.MS.hessian(freePars, freePars,:);
        newParameters.MS.exitflag = parameters.MS.exitflag;
    end
    
    %% Objective function
    newObjectiveFunction = @(freeTheta) getReducedObjectiveFunction(freeTheta, objectiveFunction, freePars, fixedPars, fixedParValues);

    %% Property struct
    newProperties = properties;
    if ~isempty(newProperties)
        for iPro = 1:properties.number
            newProperties.function{iPro} = @(freeTheta) getReducedPropertyFunction(freeTheta, properties.function{iPro}, freePars, fixedPars, fixedParValues);
        end
    else
        newProperties = [];
    end
    
    %% Options
    newOptions = options;
    newOptions.fixedParameters = [];
    newOptions.fixedParameterValues = [];
    
end

%% 
function varargout = getReducedObjectiveFunction(freeTheta, objectiveFunction, freePars, fixedParameters, fixedParameterValues)
    
    % Construction of theta
    theta(freePars) = freeTheta;
    theta(fixedParameters) = fixedParameterValues;

    % Evaluation of objective function
    switch nargout
        case {0,1}
            J = objectiveFunction(theta);

        case 2
            [J,G] = objectiveFunction(theta);

        case 3
            [J,G,H] = objectiveFunction(theta);
    end
    
    % Assignment of output
    varargout{1} = J;
    if nargout >= 2
        varargout{2} = G(freePars);
    end
    if nargout >= 3
        varargout{3} = H(freePars,freePars);
    end
end

%% 
function varargout = getReducedPropertyFunction(freeTheta, propertyFunction, freePars, fixedParameters, fixedParameterValues)
    
    % Construction of theta
    theta(freePars) = freeTheta;
    theta(fixedParameters) = fixedParameterValues;

    % Evaluation of objective function
    switch nargout
        case {0,1}
            Prop = propertyFunction(theta);

        case 2
            [Prop,grad_Prop] = propertyFunction(theta);

        case 3
            [Prop,grad_Prop,hessian_grad_Prop] = propertyFunction(theta);
    end
    
    % Assignment of output
    varargout{1} = Prop;
    if nargout >= 2
        varargout{2} = grad_Prop(freePars);
    end
    if nargout >= 3
        varargout{3} = hessian_grad_Prop(freePars,freePars);
    end
end
```
