## Supplementary material for "Mechanistic description of spatial processes using integrative modelling of noise-corrupted imaging data": meigoDummy.m

```
function varargout = meigoDummy(theta, fun, varargin)
% Objective function wrapper for MEIGO / PSwarm / ... which need objective
% function *file*name* and cannot use function handles directly. 
% 
% Parameters:
%   theta: parameter vector
%   fun: objective function handle
%   varargin: 
%
% Return values:
% f: Objective function value

if(nargin(fun) == 1)
    switch nargout
        case 1
            [f] = fun(theta);
            varargout{1} = f;
        case 2
            [f,g] = fun(theta);
            varargout{1} = f;
            varargout{2} = g;
        case 3
            [f,g,h] = fun(theta);
            varargout{1} = f;
            varargout{2} = g;
            varargout{3} = h;
    end
else
    switch nargout
        case 1
            [f] = fun(theta, varargin);
            varargout{1} = f;
        case 2
            [f,g] = fun(theta, varargin);
            varargout{1} = f;
            varargout{2} = g;
        case 3
            [f,g,h] = fun(theta, varargin);
            varargout{1} = f;
            varargout{2} = g;
            varargout{3} = h;
    end
end

end
```
