## Supplementary material for "Mechanistic description of spatial processes using integrative modelling of noise-corrupted imaging data": plotConfidenceIntervals.m

```
function fh = plotConfidenceIntervals(pStruct, alpha, varargin)
% plotConfidenceIntervals.m visualizes confidence itervals stored in either
% the parameters or properties struct .CI
%
% USAGE:
% fh = plotParameterUncertainty(pStruct)
% fh = plotParameterUncertainty(pStruct, methods)
% fh = plotParameterUncertainty(pStruct, methods, options)
%
% plotMultiStarts() uses the following PestoPlottingOptions members:
%  * PestoPlottingOptions::P
%  * PestoPlottingOptions::S
%  * PestoPlottingOptions::MS
%  * PestoPlottingOptions::boundary
%  * PestoPlottingOptions::subplot_size_1D
%  * PestoPlottingOptions::subplot_indexing_1D
%  * PestoPlottingOptions::CL
%  * PestoPlottingOptions::hold_on
%  * PestoPlottingOptions::interval
%  * PestoPlottingOptions::bounds
%  * PestoPlottingOptions::A
%  * PestoPlottingOptions::add_points
%  * PestoPlottingOptions::labels
%  * PestoPlottingOptions::legend
%  * PestoPlottingOptions::op2D
%  * PestoPlottingOptions::fontsize
%
% Parameters:
%   pStruct: either the parameter or the property struct containing 
%       information about parameters and results of optimization (.MS) 
%       and uncertainty analysis (.P and .S). This structures is the output
%       of plotMultiStarts.m, getProfiles.m or plotSamples.m.
%  alpha: significance levels
%   varargin:
%     method: integer array, from which method confidence intervals 
%         should be plotted: 
%     options: options of plotting as instance of PestoPlottingOptions
%
% Return values:
%   fh: figure handle
%
% History:
% * 2016/11/14 Paul Stapor
% * 2017/12/27 Jan Hasenauer

%% Check and assign input

% Check which methods should be used to plot Confidence Intervals
if (length(varargin) >= 1)
    if(~isempty(varargin{1}))
        methIn = varargin{1};
        boolWarning = true;
    else
        methIn = {'local_PL', 'PL', 'local_B', 'S'};
        boolWarning = false;
    end
else
    methIn = {'local_PL', 'PL', 'local_B', 'S'};
end
methods = checkMeth(methIn, pStruct, boolWarning);
numConf = methods.num;

% Assignment of user-provided options
if (length(varargin) >= 2)
    if (~isa(varargin{2}, 'PestoOptions'))
        error('Argument 2 is not of type PestoOptions.')
    end
    allOptions = varargin{2};
    options = allOptions.plot_options.copy();
else
    allOptions = PestoOptions();
    options = PestoPlottingOptions();
end

%% Check what exactly should be plotted

if isfield(pStruct, 'function')
    type = 'Properties';
    if isempty(allOptions.property_index)
        pIndexSet = 1 : pStruct.number;
    else
        pIndexSet = allOptions.property_index;
    end
else
    type = 'Parameters';
    if isempty(allOptions.parameter_index)
        pIndexSet = 1 : pStruct.number;
    else
        pIndexSet = allOptions.parameter_index;
    end
end

% Check, if pStruct has all necessary fieds
pStruct = checkSanityOfStructs(pStruct, ['p' type(2:end)]);

numP = length(pIndexSet);
iMAP = allOptions.MAP_index;
if isempty(iMAP)
    iMAP = 1;
end

switch options.group_CI_by
    case 'parprop'
        indexSet = pIndexSet;
        indexLen = length(pIndexSet);
    case 'methods'
        indexSet = 1 : numConf;
        indexLen = numConf;
    case 'all'
        indexLen = 1;
    otherwise
        error('Call to undefinded grouping method plot plot of confidence intervals.');
end
        
%% Initialize the figure generation
sqrtPlots = ceil(sqrt(indexLen));
if ((sqrtPlots-1)*sqrtPlots >= indexLen)
    plots = [sqrtPlots-1, sqrtPlots];
else
    plots = [sqrtPlots, sqrtPlots];
end

% Open figure
fh = figure('Name', ['plotConfidenceIntervals - ' type]);
pos = nan(indexLen, 4);

%% Generate the Plots
switch options.group_CI_by
    case 'parprop'
        for iP = 1 : indexLen %indexSet
            subplot(plots(1), plots(2), iP);
            pos(iP,:) = get(gca, 'Position');
            del = 0.1 / plots(2);
            offsetLabels = 0.1;
            step = (1 - offsetLabels) / plots(2);
            pos(iP,:) = [mod((iP-1), plots(2))*step + del/2 + offsetLabels, pos(iP, 2), step-del, pos(iP, 4)];
            hold on;

            set(gca, 'YLim', [0.5, numConf + 0.5]);
            ylabel('');
            xlabel(pStruct.name{indexSet(iP)});
            if (mod(iP-1, plots(2)) == 0)
                set(gca, 'ytick', 1 : numConf, 'yticklabel', methods.name);
            else
                set(gca, 'ytick', 1 : numConf, 'yticklabel', '');
            end
            box on;

            for j = 1 : numConf
                CI = pStruct.CI.(methods.type{j});
                if (j == 1) && isnan(CI(indexSet(iP),1,1))
                    % text(pStruct.MS.par(indexSet(iP),1),j,'$\emptyset$','Interpreter','latex','HorizontalAlignment','center','VerticalAlignment','middle')
                else
                    for k = methods.numLevels : -1 : 1
                        h = methods.bars(k);
                        if (CI(indexSet(iP),1,k) == -inf)
                            CI(indexSet(iP),1,k) = pStruct.min(indexSet(iP));
                        end
                        if ((CI(indexSet(iP),2,k)) == inf)
                            CI(indexSet(iP),2,k) = pStruct.max(indexSet(iP));
                        end
                        patch([CI(indexSet(iP),1,k), CI(indexSet(iP),2,k), CI(indexSet(iP),2,k), CI(indexSet(iP),1,k)], [j-h, j-h, j+h, j+h], 'k', 'FaceColor', methods.colors(j,:,k), 'EdgeColor', 'k');
                    end
                end
                if isfield(pStruct, 'MS')
                    if (strcmp(type, 'Parameters'))
                        plot([pStruct.MS.par(indexSet(iP),1), pStruct.MS.par(indexSet(iP),1)], [j-0.4, j+0.4], 'k-', 'linewidth', 2);
                    else
                        plot([pStruct.MS.prop(indexSet(iP),1), pStruct.MS.prop(indexSet(iP),1)], [j-0.4, j+0.4], 'k-', 'linewidth', 2);
                    end
                end
            end

            if (options.draw_bounds)
                xLimits = get(gca, 'XLim');
                if (xLimits(1) <= pStruct.min(indexSet(iP)) && xLimits(2) >= pStruct.min(indexSet(iP)))
                    plot([pStruct.min(indexSet(iP)), pStruct.min(indexSet(iP))], [0.5, numConf+0.5], 'b--', 'linewidth', 2);
                    set(gca, 'XLim', [pStruct.min(indexSet(iP)), xLimits(2)]);
                end
                xLimits = get(gca, 'XLim');
                if (xLimits(1) <= pStruct.max(indexSet(iP)) && xLimits(2) >= pStruct.max(indexSet(iP)))
                    plot([pStruct.max(indexSet(iP)), pStruct.max(indexSet(iP))], [0.5, numConf+0.5], 'b--', 'linewidth', 2);
                    set(gca, 'XLim', [xLimits(1), pStruct.max(indexSet(iP))]);
                end
            end
            hold off;
        end
        
        % Relocate the images to where I want them to be. Don't try to go
        % for a more intelligent solution. I did, and it made me alomost
        % mad... The Matlab syntax for figure handles is, well... o.O'
        fig = gcf;
        fig.Children = flip(fig.Children);
        for iP = 1 : indexLen %indexSet 
            set(fig.Children(iP), 'Position', pos(iP,:));
        end
        
    case 'methods'
        for iM = indexSet
            subplot(plots(1), plots(2), iM);
            pos(iM,:) = get(gca, 'Position');
            del = 0.1 / plots(2);
            offsetLabels = 0.1;
            step = (1 - offsetLabels) / plots(2);
            pos(iM,:) = [mod((iM-1), plots(2))*step + del/2 + offsetLabels, pos(iM, 2), step-del, pos(iM, 4)];
            hold on;
            
            set(gca, 'YLim', [0.5, numP + 0.5]);
            title(methods.name{iM});
            ylabel('');
            xlabel('');
            if (iM == 1 || iM == 3)
                set(gca, 'ytick', 1 : numP, 'yticklabel', pStruct.name(pIndexSet));
            else
                set(gca, 'ytick', 1 : numP, 'yticklabel', '');
            end
            box on;

            XMin = min(pStruct.min);
            XMax = max(pStruct.max);
            ax = gca;
            iP = 1;
            for j = pIndexSet
                CI = pStruct.CI.(methods.type{iM});
                for k = methods.numLevels : -1 : 1;
                    h = methods.bars(k);
                    CI(iM,1,k) = max(CI(iM,1,k), XMin);
                    CI(iM,1,k) = min(CI(iM,1,k), XMax);
                    patch([CI(j,1,k), CI(j,2,k), CI(j,2,k), CI(j,1,k)], [iP-h, iP-h, iP+h, iP+h], 'k', 'FaceColor', methods.colors(iM,:,k), 'EdgeColor', 'k');
                end
                if isfield(pStruct, 'MS')
                    if (strcmp(type, 'Parameters'))
                        plot([pStruct.MS.par(j,iMAP), pStruct.MS.par(j,iMAP)], [j-0.4, j+0.4], 'k-', 'linewidth', 2);
                    else
                        plot([pStruct.MS.prop(j,iMAP), pStruct.MS.prop(j,iMAP)], [j-0.4, j+0.4], 'k-', 'linewidth', 2);
                    end
                end
                iP = iP + 1;
            end

            if (options.draw_bounds)
                limits = [ax.XLim(1), ax.XLim(2)];
                pIndex1 = [pStruct.min(pIndexSet(1)); pStruct.min(pIndexSet); pStruct.min(pIndexSet(end))];
                pIndex2 = [pStruct.max(pIndexSet(1)); pStruct.max(pIndexSet); pStruct.max(pIndexSet(end))];
                plot(pIndex1, 0 : numP+1, 'b--', 'linewidth', 2);
                plot(pIndex2, 0 : numP+1, 'b--', 'linewidth', 2);
                set(gca, 'XLim', limits);
            end
            hold off;
        end
        
        fig = gcf;
        fig.Children = flip(fig.Children);
        for iM = indexSet 
            set(fig.Children(iM), 'Position', pos(iM,:));
        end
        
    case 'all'
        
end

end


function methodsOut = checkMeth(methodsIn, pStruct, boolWarning)
    % checkMeth
    %
    % Parameters:
    %  methodsIn:
    %  pStruct: 
    %  boolWarning:
    %
    % Return values:
    %  methodsOut:
    tempMethType = {'PL', 'local_PL', 'local_B', 'S'};
    tempMethName = {'Profile L.', 'L.App, th.', 'L.App, mass', 'Bay., Sampl.'};

    checkMeth = zeros(1,4);
    for j = 1 : 4
        for k = 1 : length(methodsIn)
            if (strcmp(methodsIn{k}, tempMethType{j}))
                if (isfield(pStruct.CI, methodsIn{k}))
                    checkMeth(j) = 1;
                else
                    if boolWarning
                        warning(['You wanted to plot confidence intervals using the method ' methodsIn{k} ...
                        ', but you did not pass the data to do so. This confidence intervals will not be plotted.']);
                    end
                end
            end
        end
    end
    
    iMeth = 0;
    for j = 1 : 4
        if (checkMeth(j) == 1)
            iMeth = iMeth + 1;
            methodsOut.type{iMeth} = tempMethType{j};
            methodsOut.name{iMeth} = tempMethName{j};
        end
    end
    methodsOut.numLevels = length(pStruct.CI.alpha_levels);
    methodsOut.levels = pStruct.CI.alpha_levels;
    methodsOut.num = length(methodsOut.name);
    methodsOut.bars = linspace(0.3, 0.15, methodsOut.numLevels);
    colors = nan(methodsOut.num, 3, methodsOut.numLevels);
    tempColors = [0.1, 0.1, 0.7; 0.6, 0, 0; 0, 0.45, 0; 0.4, 0.3, 0];
    tempColorStep = [0.25, 0.25, 0.1; 0.1, 0.2, 0.2; 0.2, 0.15, 0.2; 0.15, 0.15, 0.2];
    colors(:,:,1) = tempColors(1:methodsOut.num,:);
    colorStep = tempColorStep(1:methodsOut.num,:);
    for j = 2 : methodsOut.numLevels
        colors(:,:,j) = min(colors(:,:,j-1) + colorStep, 1);
    end
    methodsOut.colors = colors;
end
```
