## Supplementary material for "Mechanistic description of spatial processes using integrative modelling of noise-corrupted imaging data": plotMCMCdiagnosis.m

```
function [fh] = plotMCMCdiagnosis(parameters, varargin)
% plotMCMCdiagnosis.m visualizes the Markov chains generated by getSamples.m.
%
% USAGE:
% fh = plotMCMCdiagnosis(parameters)
% fh = plotMCMCdiagnosis(parameters,type)
% fh = plotMCMCdiagnosis(parameters,type,fh)
% fh = plotMCMCdiagnosis(parameters,type,fh,I)
% fh = plotMCMCdiagnosis(parameters,type,fh,I,options)
%
% Parameters:
% parameters: parameter struct containing information about parameters
%       and results of optimization (.MS) and uncertainty analysis
%       (.S). This structures is the output of plotMultiStarts.m,
%       getProfiles.m or plotSamples.m.
% varargin:
% type: string indicating the type of visualization:
%       'parameters' (default) and 'log-posterior'
% fh: handle of figure. If no figure handle is provided, a new figure
%       is opened.
% I: index of parameters which are updated. If no index is provided
%       all parameters are updated.
% options: options of plotting as instance of PestoPlottingOptions
%
% Return values:
% fh: figure handle
%
% History:
% * 2014/06/20 Jan Hasenauer
% * 2016/10/10 Daniel Weindl

%% Check and assign inputs

% Check, if parameters has all necessary fieds
parameters = checkSanityOfStructs(parameters, 'parameters');

% Plot type
type = 'parameters';
if nargin >= 2 && ~isempty(varargin{1})
    type = varargin{1};
    if ~max(strcmp({'parameters','log-posterior'},type))
        error('The ''type'' of plot is unknown. ''type'' can only be ''parameters'' or ''log-posterior''.')
    end
end

% Figure handle
if nargin >= 3 && ~isempty(varargin{2})
    fh = figure(varargin{2});
else
    fh = figure('Name','MCMC diagnosis');
end

% Index of subplot which is updated
I = 1:parameters.number;
if nargin >= 4 && ~isempty(varargin{3})
    I = varargin{3};
end

% Options
options = PestoPlottingOptions();
options.plot_type = {'parameter','posterior'};
options.n_max = 1e4;

% Default sample plotting options
%   0 => no plot of mean
%   1 => plot of mean
options.S.plot_type = 1; 
options.S.col = [1,0,0];
options.S.ls = '';
options.S.lw = 1;
options.S.m = '.';
options.S.ms = 5;
options.S.mean_col = [0.5,0,0];
options.S.mean_lw = 1.5;
options.S.col = [0,0,1];

% Local optima
%   0 => no plot
%   1 => plot
if isfield(parameters,'MS')
    options.MS.plot_type = 1;
else
    options.MS.plot_type = 0; 
end
options.MS.col = [1,0,0];
options.MS.lw = 1.5;

% Confidence level
options.CL.plot_type = options.MS.plot_type;
options.CL.alpha = 0.95;
options.CL.type = 'point-wise'; % 'simultanous', {'point-wise','simultanous'}
options.CL.col = [1,0,0];
options.CL.lw = 1.5;

if length(varargin) >= 5
    options = setdefault(handlePlottingOptionArgument(varargin{5}), options);
end


%% Initialization
% Number of MCMC samples
j_max = length(parameters.S.logPost);

% Thinning factot
th = ceil(j_max/options.n_max);

%% Plot: Parameter chains
if strcmp(type,'parameters')
    
% Compute number of subfigure
s = round(sqrt(length(I))*[1,1]);
if prod(s) < length(I)
    s(2) = s(2) + 1;
end

% Loop: Parameter
for l = 1:length(I)
    legstr = {};
    
    % Assign parameter index
    i = I(l);
    
    % Open subplot
    subplot(s(1),s(2),l);
    
    % Hold on/off
    if options.hold_on
        hold on;
    else
        hold off;
    end

    % Plot: MCMC sample
    J = 1:th:j_max;
    plot(J,parameters.S.par(i,J),[options.S.ls options.S.m],...
        'linewidth',options.S.lw,'markersize',options.S.ms,'color',options.S.col);
    hold on;
    legstr{1} = 'MCMC sample';
    
    % Plot: MCMC sample mean
    if options.S.plot_type
        plot([0,j_max],mean(parameters.S.par(i,:))*[1,1],...
            'linewidth',options.S.mean_lw,'color',options.S.mean_col);
        legstr{end+1} = 'MCMC sample mean';
    end

    % Plot: MAP estimate
    if options.MS.plot_type
        plot([0,j_max],parameters.MS.par(i,1)*[1,1],'-',...
            'linewidth',options.MS.lw,'color',options.MS.col);
        legstr{end+1} = 'MAP';
    end
    
    % Limits
    xlim([0,j_max]);
    switch options.interval
        case 'static'
            yl = [parameters.min(i),parameters.max(i)];
        case 'dynamic'
            yl = [min(parameters.S.par(i,:)),max(parameters.S.par(i,:))];
    end
    ylim(yl);
    
    % Legend
    if l == 1
        legend(legstr,'location','SouthEast');
    end
    
    % Labels
    xlabel('sample path');
    ylabel(parameters.name(i));

end

end

%% Plot: log-Posterior chain
if strcmp(type,'log-posterior')

    % Hold on/off
    if options.hold_on
        hold on;
    else
        hold off;
    end
    
    % Plot: MCMC posterior
    J = 1:th:j_max;
    plot(J,parameters.S.logPost(J),[options.S.ls options.S.m],...
        'linewidth',options.S.lw,'markersize',options.S.ms,'color',options.S.col);
    hold on;
    legstr{1} = 'MCMC sample';
    
    % Plot: MAP estimate
    if options.MS.plot_type
        plot([0,j_max],parameters.MS.logPost(1)*[1,1],'-',...
            'linewidth',options.MS.lw,'color',options.MS.col);
        legstr{end+1} = 'MAP';
    end
    
    % Plot: Confidence level
    if options.CL.plot_type
        if max(strcmp(options.CL.type,'point-wise'))
            plot([0,j_max],[1,1]*(parameters.MS.logPost(1)-chi2inv(options.CL.alpha,1)/2),'--','linewidth',options.CL.lw,'color',options.CL.col);
            legstr{end+1} = ['point-wise ' num2str(100*options.CL.alpha) '% conf. interval'];
        end
        if max(strcmp(options.CL.type,'simultanous'))
            plot([0,j_max],[1,1]*(parameters.MS.logPost(1)-chi2inv(options.CL.alpha,parameters.number)/2),':','linewidth',options.CL.lw,'color',options.CL.col);
            legstr{end+1} = ['simultanous ' num2str(100*options.CL.alpha) '% conf. interval'];
        end
    end
    
    % Limits
    xlim([0,j_max]);
    
    % Legend
    legend(legstr,'location','SouthEast');
    
    % Labels
    xlabel('sample path');
    ylabel('log-posterior');

end
```
