## Supplementary material for "Mechanistic description of spatial processes using integrative modelling of noise-corrupted imaging data": plotMultiStartDiagnosis.m

```
% plotMultiStartRunTimes plots the distributions of comutation time,
% iterations and objective function evaluations of the result of the
% multi-start optimization stored in parameters
%
% USAGE:
% ======
% fh = plotMultiStartRunTimes(parameters)
% fh = plotMultiStartRunTimes(parameters,fh,options)
%
% INPUTS:
% =======
% parameters ... parameter struct containing information about parameters
%   and log-posterior.about
% fh ... handle of figure in which distributions are plotted. If no
%   figure handle is provided, a new figure is opened.
% options ... options of plotting
%   .title ... switches plot title off (default = 'off').
%   .col ... colors used for the different histograms, has to be a 3x3 matrix
%       (default: grey = [0.4286,0.4286,0.4286;
%                         0.5714,0.5714,0.5714;
%                         0.7143,0.7143,0.7143];).
%
% Outputs:
% ========
% fh .. figure handle

function fh = plotMultiStartDiagnosis(varargin)
    
    %% CHECK AND ASSIGN INPUTS
    %Assign parameters
    if nargin >= 1
        parameters = varargin{1};
    else
        error('plotMultiStartRunTimes requires a parameter object as input.');
    end
    
    %Open figure
    if nargin >= 2
        if ~isempty(varargin{2})
            fh = figure(varargin{2});
        else
            fh = figure;
        end
    else
        fh = figure;
    end
    
    % Options
    options.title = 'off';
    options.col = [0.2081,0.1663,0.5292;
        0.1986,0.7214,0.6310;
        0.9763,0.9831,0.0538];
    
    if nargin == 3
        options = setdefault(varargin{3},options);
    end
    
    %% SORT RESULTS
    [parameters] = sortMultiStarts(parameters);
    
    %% PLOT DISTRIBUTION COMPUTATION TIME
    subplot(3,3,9)
    cpu_time_runs = parameters.MS.t_cpu;
    cpu_time_runs(cpu_time_runs == 0) = NaN;
    getHistogram(cpu_time_runs/60,options.col(2,:),{'cpu time single start [min]'},[])
    if strcmp(options.title,'on')
        title({'CPU time per'; 'optimization'});
    end
    
    
    %% PLOT DISTRIBUTION ITERATIONS
    subplot(3,3,3)
    getHistogram(parameters.MS.n_iter,options.col(2,:),{'objective function'; 'iterations single start'},[])
    if strcmp(options.title,'on')
        title({'Number of iterations'; 'per optimization'});
    end
    
    %% PLOT DISTRIBUTION OBJECTIVE FUNCTION EVALUATIONS
    subplot(3,3,6)
    if isfield(options,'bounds')
        limits_x = [options.bounds.min(3),options.bounds.max(3)];
    else
        limits_x = [];
    end
    
    getHistogram(parameters.MS.n_objfun,options.col(3,:),{'objective function'; 'evaluations single start'},[])
    
    if strcmp(options.title,'on')
        title({'Number of objective function'; 'evaluations per optimization'});
    end
    
    %% PLOT logPost
    subplot(3,3,1)
    startidx = 1:parameters.MS.n_starts;
    
    if(isfield(parameters.MS,'fval_trace'));
        min_fval = transpose(min(parameters.MS.fval_trace));
        minLP = min(min_fval);
        plot(startidx(isnan(parameters.MS.logPost)),min_fval(isnan(parameters.MS.logPost))-minLP+1,'rx')
        hold on
    else
        minLP = min(-parameters.MS.logPost);
    end
    plot(startidx,-parameters.MS.logPost-minLP+1,'k.')
    xlim([0,parameters.MS.n_starts])
    xlabel('start index')
    ylabel('-logPost+min(logPost)+1')
    set(gca,'YScale','log')
    lim_y = get(gca,'YLim');
    lim_y(1) = 10^(-0.5);
    ylim(lim_y);
    
    
    %% PLOT FMINCON EXITFLAG
    if(isfield(parameters.MS,'exitflag'))
        subplot(3,3,4)
        eflag = parameters.MS.exitflag;
        eflag(eflag>3) = -2;
        eflag(eflag<-2) = -2;
        plot(1:parameters.MS.n_starts,eflag,'k.')
        unfinished = NaN(size(eflag));
        unfinished(isnan(parameters.MS.logPost)) = 4;
        hold on
        plot(1:parameters.MS.n_starts,unfinished,'rx')
        set(gca,'YTick',-2:4)
        set(gca,'YTickLabel',{'other','output fcn','feval/iter','gradient','change in x','change in f','not finished'})
        xlabel('start index')
        ylabel('stopping condition')
        xlim([0,parameters.MS.n_starts])
        ylim([-2.5,4.5])
    end
    
    %% PLOT FMINCON Gradient
    if(isfield(parameters.MS,'gradient'))
        subplot(3,3,7)
        ngrad  = sqrt(sum(parameters.MS.gradient.^2));
        plot(1:parameters.MS.n_starts,ngrad,'k.')
        set(gca,'YScale','log')
        xlabel('start index')
        ylabel('norm of gradient')
        xlim([0,parameters.MS.n_starts])
    end
    
    %% PLOT fval_trace
    if(isfield(parameters.MS,'fval_trace'))
        subplot(3,3,2)
        getLinePlot(parameters.MS.fval_trace,~isnan(parameters.MS.logPost))
        xlabel('iteration')
        ylabel('-logPost+min(logPost)+1')
        set(gca,'YScale','log')
    end
    
    %% PLOT diff par_trace
    if(isfield(parameters.MS,'par_trace'))
        subplot(3,3,5)
        par_step = permute(sqrt(sum(diff(parameters.MS.par_trace,1,2).^2,1)),[2,3,1]);
        getLinePlot([NaN(1,size(par_step,2));par_step],~isnan(parameters.MS.logPost))
        xlabel('iteration')
        ylabel('norm of parameter step')
        set(gca,'YScale','log')
    end
    
    
    %% PLOT time_trace
    if(isfield(parameters.MS,'time_trace'))
        subplot(3,3,8)
        getLinePlot(parameters.MS.time_trace/60,~isnan(parameters.MS.logPost))
        xlabel('iteration')
        ylabel('computation time [min]')
    end
    
end

function getHistogram(x,color,xlbl,limx)
    if(any(~isnan(x)))
        logx = log10(x);
        nbin = getBins(logx,'optimal');
        [N,X] = hist(logx,nbin);
        he = bar(X,N,1.0,'FaceColor',color);
        ylim([0,max(he.YData)+1]);
        ylabel('count');
        if(isempty(limx))
            limx = [min(logx)-0.5,max(logx)+0.5];
        end
        xlim(limx);
        xlabel(xlbl);
        xticks = get(gca,'XTick');
        while length(xticks)>5
            xticks = xticks(1:2:end);
        end
        set(gca,'XTick',xticks);
        set(gca,'XTickLabel',arrayfun(@(x) ['10^{' num2str(x) '}'],get(gca,'XTick'),'UniformOutput',false));
    end
end

function getLinePlot(y,group)
    nmaxiter = size(y,1);
    l1 = plot(1:nmaxiter,y(:,group),'k-');
    if(any(any(~isnan(y(:,not(group))))))
        hold on
        l2 = plot(transpose(1:nmaxiter),y(:,not(group)),'r-');
        legend([l1(1),l2(1)],{'finished start','unfinished start'},'Location','best')
        setLineTransparency(l2,0.2);
    end
    setLineTransparency(l1,0.4);

end

function setLineTransparency(lines,trans)
    for il = 1:length(lines)
        lines(il).Color(4) = trans;
    end
end
```
