## Supplementary material for "Mechanistic description of spatial processes using integrative modelling of noise-corrupted imaging data": plotMultiStarts.m

```
function fh = plotMultiStarts(parameters, varargin)
% plotMultiStarts plots the result of the multi-start 
% optimization stored in parameters.
%
% USAGE:
% fh = plotMultiStarts(parameters)
% fh = plotMultiStarts(parameters,fh)
% fh = plotMultiStarts(parameters,fh,options)
%
% plotMultiStarts() uses the following PestoPlottingOptions members:
%  * PestoPlottingOptions::add_points
%  * PestoPlottingOptions::title
%  * PestoPlottingOptions::draw_bounds
%
% Parameters:
% parameters: parameter struct containing information about parameters
%   and log-posterior.
% varargin:
% fh: handle of figure in which profile likelihood is plotted. If no
%   figure handle is provided, a new figure is opened.
% options: options of plotting as instance of PestoPlottingOptions
%
% Return values:
% fh: figure handle
%
% History: 
% * 2012/05/31 Jan Hasenauer
% * 2016/10/07 Daniel Weindl

%% CHECK AND ASSIGN INPUTS

% Check, if parameters has all necessary fieds
parameters = checkSanityOfStructs(parameters, 'parameters');

% Open figure
if length(varargin) >= 1 && ~isempty(varargin{1}) && isvalid(varargin{1})
    fh = figure(varargin{1});
else
    fh = figure('Name','plotMultiStarts');
end

% Options
defaultOptions = PestoPlottingOptions();
defaultOptions.add_points.par = [];
defaultOptions.add_points.logPost = [];
defaultOptions.add_points.col = [0,0.8,0];
defaultOptions.add_points.ls = '-';
defaultOptions.add_points.lw = 1;
defaultOptions.add_points.m = 'd';
defaultOptions.add_points.ms = 8;
defaultOptions.add_points.name = 'add. point';

if length(varargin) >= 2
    options = varargin{2};
    options = handlePlottingOptionArgument(options);
    options = setdefault(options, defaultOptions);
    options.add_points = setdefault(options.add_points, defaultOptions.add_points);
else
    options = defaultOptions;
end

%% SORT RESULTS
[parameters] = sortMultiStarts(parameters);

%% CLUSTERING
n_starts = length(parameters.MS.logPost);
if (n_starts > 1)
    clust = cluster(linkage(pdist(parameters.MS.logPost)),'cutoff',0.1,'criterion','distance');
else
    clust = 1;
end
uclust = unique(clust);
    
for iclust = 1:length(uclust)
    sizecluster(iclust) = sum(clust == uclust(iclust));
end

%% ASSIGN COLORS
Col = colormap(gray(n_starts+ceil(n_starts/3)));
Col = Col.^(1/3);
Col(1,:) = [1,0,0];

% sort clusters
for iclust = 1:length(uclust)
    Jclust(iclust) = max(parameters.MS.logPost(find(clust == uclust(iclust))));
end
Jclust(isnan(Jclust)) = -Inf;
[~,idx] = sort(Jclust,'descend');
uclust = uclust(idx);
sizecluster = sizecluster(idx);

if(sizecluster(1)>1)
    ColClust = [1,0,0;flipud(parula(max(sum(sizecluster>1)-1,0)))];
else
    ColClust = flipud(parula(sum(sizecluster>1)));
end

for iclust = 1:length(uclust)
    if(sizecluster(iclust)>1)
    Col(clust == uclust(iclust),:) = repmat(ColClust(sum(sizecluster(1:iclust)>1),:),[sizecluster(iclust),1]);
    end
end

%% PLOT OBJECTIVES
subplot(2,2,1);
n_finished_starts = 0;
for j = 1 : n_starts
    if ~isnan(parameters.MS.logPost(j))
        n_finished_starts = j;
    else
        break;
    end
end

plot(1:n_finished_starts,parameters.MS.logPost(1:n_finished_starts),'-','color',0.9*[1,1,1],'linewidth',2);
hold on;
for j = n_finished_starts:-1:1
    plot(j,parameters.MS.logPost(j),'o','color',Col(j,:),'linewidth',2);
    hold on;
end
if ~isempty(options.add_points.logPost)
    if length(options.add_points.logPost) == 1
        if (n_starts == 1)
            addPointsX = [0.85 1.15];
            addPointsY = options.add_points.logPost * [1 1];
        else
            addPointsX = 1 : n_starts;
            addPointsY = options.add_points.logPost * ones(size(1:n_starts));
        end
        plot(addPointsX,addPointsY,options.add_points.ls,'color',...
            options.add_points.col(1,:),'linewidth',options.add_points.lw); 
        hold on;
    else
        plot(1:length(options.add_points.logPost),options.add_points.logPost,...
            options.add_points.ls,'color',options.add_points.col(1,:),...
            'linewidth',options.add_points.lw,'marker',options.add_points.m,...
            'markersize',options.add_points.ms); hold on;        
    end
end
hold off;
xlim([1-0.2,n_starts+0.2]);
xlabel('start');
ylabel('log-likelihood');
if options.title
    title('all estimates');
end

%% PLOT TOP TEN OBJECTIVES
subplot(2,2,3);
plot(1:min(n_starts,10),parameters.MS.logPost(1:min(n_starts,10)),'-','color',0.9*[1,1,1],'linewidth',2); hold on;
for j = min(n_starts,10):-1:1
    plot(j,parameters.MS.logPost(j),'o','color',Col(j,:),'linewidth',2); hold on;
end
if ~isempty(options.add_points.logPost)
    if length(options.add_points.logPost) == 1
        plot(1:min(n_starts,10),options.add_points.logPost*ones(size(1:min(n_starts,10))),...
            options.add_points.ls,'color',options.add_points.col(1,:),...
            'linewidth',options.add_points.lw); hold on;
    else
        plot(1:min(length(options.add_points.logPost),10),options.add_points.logPost(1:min(length(options.add_points.logPost),10)),...
            options.add_points.ls,'color',options.add_points.col(1,:),...
            'linewidth',options.add_points.lw,'marker',options.add_points.m,...
            'markersize',options.add_points.ms); hold on;        
    end
end
hold off;
xlim([1-0.2,min(n_starts,10)+0.2]);
if(any(~isnan(parameters.MS.logPost(1:min(n_starts,10)))))
    ylim([min(parameters.MS.logPost(1),min(parameters.MS.logPost(1:min(n_starts,10)))-1),parameters.MS.logPost(1)+1]);
end
xlabel('start');
ylabel('log-likelihood');
if options.title
    title('top 10 estimates');
end

%% PLOT PARAMETERS
subplot(2,2,[2,4]);
for j = n_finished_starts:-1:1
    plot(parameters.MS.par(:,j)',1:parameters.number,'-o','color',Col(j,:),'linewidth',2); hold on;
end
plot(parameters.MS.par(:,1)',1:parameters.number,'r-o','linewidth',2); hold on;
if options.draw_bounds
    plot(parameters.min([1,1:parameters.number,parameters.number])',[0.99,1:parameters.number,parameters.number+0.01],'b--','linewidth',2); hold on;
    plot(parameters.max([1,1:parameters.number,parameters.number])',[0.99,1:parameters.number,parameters.number+0.01],'b--','linewidth',2); hold on;
end
if ~isempty(options.add_points.logPost)
    for j = 1:size(options.add_points.par,2)
        plot(options.add_points.par(:,j)',1:parameters.number,options.add_points.ls,...
            'color',options.add_points.col(1,:),'linewidth',options.add_points.lw,...
            'marker',options.add_points.m,'markersize',options.add_points.ms); hold on;
    end
end
hold off;
ylim([1-0.01,parameters.number+0.01]);
ylabel(' ');
xlabel('parameter value');
set(gca,'ytick',1:parameters.number,'yticklabel',parameters.name)

if options.title
    title('estimated parameters');
end
drawnow;
```
