## Supplementary material for "Mechanistic description of spatial processes using integrative modelling of noise-corrupted imaging data": plotParameterProfiles.m

```
function fh = plotParameterProfiles(parameters, varargin)
% plotParameterProfiles.m visualizes profile likelihood.
% Note: This routine provides an interface for plotUncertainty.m.
%
% USAGE:
% fh = plotParameterProfiles(parameters)
% fh = plotParameterProfiles(parameters,type)
% fh = plotParameterProfiles(parameters,type,fh)
% fh = plotParameterProfiles(parameters,type,fh,I)
% fh = plotParameterProfiles(parameters,type,fh,I,options)
%
% Parameters:
%   parameters: parameter struct containing information about parameters
%     and results of optimization (.MS) and uncertainty analysis
%     (.P and .S). This structures is the output of plotMultiStarts.m,
%     getProfiles.m or plotSamples.m.
%   varargin:
%     type: string indicating the type of visualization: '1D' or '2D'
%     fh: handle of figure. If no figure handle is provided, a new figure
%         is opened.
%     I: index of parameters which are updated. If no index is provided
%         all parameters are updated.
%     options: options of plotting as instance of PestoPlottingOptions
%
% Return values:
%   fh: figure handle
%
% History:
% * 2012/05/31 Jan Hasenauer
% * 2014/06/20 Jan Hasenauer
% * 2016/10/10 Daniel Weindl


switch length(varargin) 
    case 0
        fh = plotParameterUncertainty(parameters);
    case 1
        fh = plotParameterUncertainty(parameters, varargin{1});
    case 2
        fh = plotParameterUncertainty(parameters, varargin{1}, varargin{2});
    case 3
        fh = plotParameterUncertainty(parameters, varargin{1}, varargin{2}, varargin{3});
    case 4
        fh = plotParameterUncertainty(parameters, varargin{1}, varargin{2}, varargin{3}, varargin{4});
    otherwise
        error('Too many arguments for plotPropertyProfiles().');
end
end
```
