## Supplementary material for "Mechanistic description of spatial processes using integrative modelling of noise-corrupted imaging data": plotParameterUncertainty.m

```
function fh = plotParameterUncertainty(parameters, varargin)
% plotParameterUncertainty.m visualizes profile likelihood and MCMC samples
% stored in parameters.
%
% USAGE:
% fh = plotParameterUncertainty(parameters)
% fh = plotParameterUncertainty(parameters,type)
% fh = plotParameterUncertainty(parameters,type,fh)
% fh = plotParameterUncertainty(parameters,type,fh,I)
% fh = plotParameterUncertainty(parameters,type,fh,I,options)
%
% plotMultiStarts() uses the following PestoPlottingOptions members:
%  * PestoPlottingOptions::P
%  * PestoPlottingOptions::S
%  * PestoPlottingOptions::MS
%  * PestoPlottingOptions::boundary
%  * PestoPlottingOptions::subplot_size_1D
%  * PestoPlottingOptions::subplot_indexing_1D
%  * PestoPlottingOptions::CL
%  * PestoPlottingOptions::hold_on
%  * PestoPlottingOptions::interval
%  * PestoPlottingOptions::bounds
%  * PestoPlottingOptions::A
%  * PestoPlottingOptions::add_points
%  * PestoPlottingOptions::labels
%  * PestoPlottingOptions::legend
%  * PestoPlottingOptions::op2D
%  * PestoPlottingOptions::fontsize
%
% Parameters:
%   parameters: parameter struct containing information about parameters
%       and results of optimization (.MS) and uncertainty analysis
%       (.P and .S). This structures is the output of plotMultiStarts.m,
%       getProfiles.m or plotSamples.m.
%   varargin:
%     type: string indicating the type of visualization: '1D' or '2D'
%     fh: handle of figure. If no figure handle is provided, a new figure
%         is opened.
%     I: index of parameters which are updated. If no index is provided
%         all parameters are updated.
%     options: options of plotting as instance of PestoPlottingOptions
%
% Return values:
%   fh: figure handle
%
% History:
% * 2012/05/31 Jan Hasenauer
% * 2014/06/20 Jan Hasenauer
% * 2016/10/10 Daniel Weindl

%% Check and assign inputs
% Plot type
type = '1D';
if length(varargin) >= 1 && ~isempty(varargin{1})
    type = varargin{1};
    if ~max(strcmp({'1D','2D'},type))
        error('The ''type'' of plot is unknown.')
    end
end

% Check, if parameters has all necessary fieds
parameters = checkSanityOfStructs(parameters, 'parameters');

% Open figure
if length(varargin) >= 2 && ~isempty(varargin{2})
    fh = figure(varargin{2});
else
    fh = figure('Name','plotParameterUncertainty');
end

% Index of subplot which is updated
I = 1:parameters.number;
if length(varargin) >= 3
    if ~isempty(varargin{3})
        I = varargin{3};
        if ~isnumeric(I) || max(abs(I - round(I)) > 0)
            error('I is not an integer vector.');
        end
    end
end

% Options
% General plot options
if length(varargin) >= 4
    options = handlePlottingOptionArgument(varargin{4});
else
    options = PestoPlottingOptions();
end

if ~isfield(parameters,'P')
    options.P.plot_type = 0; 
    options.boundary.mark = 0;
end

if ~isfield(parameters,'S')
    options.S.plot_type = 0;
end

if ~isfield(parameters,'MS')
    options.MS.plot_type = 0; 
end

% Subplot arrangement
if isempty(options.subplot_size_1D)
    options.subplot_size_1D = round(sqrt(length(I))*[1,1]);
    if prod(options.subplot_size_1D) < length(I)
        options.subplot_size_1D(2) = options.subplot_size_1D(2) + 1;
    end
end
if isempty(options.subplot_indexing_1D)
    options.subplot_indexing_1D = 1:length(I);
end

%% INITALIZATION
% Maximum a posterior estimate
if (isfield(parameters, 'MS'))
    logPost_max = max(parameters.MS.logPost);
else
    logPost_max = max(parameters.S.logPost);
end

% Degrees of freedom (for chi^2 test)
dof = 1;
if max(strcmp(options.CL.type,'simultanous'))
    dof = parameters.number;
end

%% 1D Parameter distributions
if strcmp(type,'1D')

% Compute number of subfigure

% Loop: Parameter
for l = 1:length(I)
    % Initialization of legend
    legh = [];
    legs = {};

    % Assign parameter index
    i = I(l);
    
    % Open subplot
    subplot(options.subplot_size_1D(1),options.subplot_size_1D(2),options.subplot_indexing_1D(l));
    
    % Hold on/off
    if options.hold_on
        hold on;
    else
        hold off;
    end
    
    % Boundaries
    switch options.interval
        case 'dynamic'
            xl = [+inf,-inf];
            
            if isfield(parameters,'MS')
                if max(strcmp(options.CL.type,'point-wise'))
                    L = find(parameters.MS.logPost(:) > (parameters.MS.logPost(1)-chi2inv(options.CL.alpha,1)/2));
                end
                if max(strcmp(options.CL.type,'simultanous'))
                    L = find(parameters.MS.logPost(:) > (parameters.MS.logPost(1)-chi2inv(options.CL.alpha,parameters.number)/2));
                end
                xl(1) = min(xl(1),min(parameters.MS.par(i,L)));
                xl(2) = max(xl(2),max(parameters.MS.par(i,L)));
            else
                xl(1) = parameters.min(i);
                xl(2) = parameters.max(i);
            end
        
            flag_plot_P = 0;
            if options.P.plot_type >= 1
                if length(parameters.P) >= i
                    if ~isempty(parameters.P(i).par)
                        xl(1) = min(xl(1), min(parameters.P(i).par(i,:)));
                        xl(2) = max(xl(2), max(parameters.P(i).par(i,:)));
                        flag_plot_P = 1;
                    end
                end
            end
            
            if xl(1) == xl(2)
                xl(1) = xl(1) - 1e-10;
                xl(2) = xl(2) + 1e-10;
            end
        case 'static'
            if ~isempty(options.bounds)
                xl = [options.bounds.min(i),options.bounds.max(i)];
            else
                xl = [parameters.min(i),parameters.max(i)];
            end
            flag_plot_P = 0;
            if options.P.plot_type >= 1
                if length(parameters.P) >= i
                    if ~isempty(parameters.P(i).par)
                        flag_plot_P = 1;
                    end
                end
            end
    end

    % Plot: Visualizaion of MCMC samples of tempered posterior distribution
    h = [];
    switch options.S.plot_type
        case 0
            % no plot
        case 1
            % histogram
                for k = 1
                    switch options.S.bins
                        case 'optimal'
                            h = 3.49*std(parameters.S.par(i,:,k))/(length(parameters.S.par(i,:,k))^(1/3));
                            nbin = round((max(parameters.S.par(i,:,k))-min(parameters.S.par(i,:,k)))/h);
                        case 'conservative'
                            h = 2*3.49*std(parameters.S.par(i,:,k))/(length(parameters.S.par(i,:,k))^(1/3));
                            nbin = round((max(parameters.S.par(i,:,k))-min(parameters.S.par(i,:,k)))/h);
                        otherwise
                            nbin = options.S.bins;
                    end
                    [N,X] = hist(parameters.S.par(i,:,k),nbin);
                    h = bar(X,N/max(N),1,'facecolor',options.S.hist_col(k,:),'edgecolor',[0.4,0.4,0.4]);
                    hold on;
                    
                    if strcmp(options.interval, 'dynamic')
                        xl(1) = min(xl(1), min(X));
                        xl(2) = max(xl(2), max(X));
                    end
                    % bar(X,N/max(N),1,'facecolor','none','edgecolor',options.S.col(k,:)); hold on;
                end
        case 2
            % kernel-density estimate
             for k = options.S.ind:-1:1
                 x_grid = linspace(min(parameters.S.par(i,:,k)),max(parameters.S.par(i,:,k)),100);
                 [KDest] = getKernelDensityEstimate(squeeze(parameters.S.par(i,:,k)),x_grid);
                 h = plot(x_grid,KDest/max(KDest),'-','color',options.S.sp_col(k,:),'linewidth',options.S.lw); hold on;
             end
        otherwise
            error('Selected value for ''options.S.plot_type'' is not available.');
    end

    if ~isempty(h)
        legh(end+1) = h;
        legs{end+1} = options.S.name;
    end

    % Plot: Local approximation
    h = [];
    switch options.A.plot_type
        case 0
            % no plot
        case 1
            if (isfield(parameters, 'MS'))
                if isfield(parameters.MS,'hessian')
                    ind_not_nan = find(~isnan(diag(parameters.MS.hessian(:,:,1))));
                    j = find(i==ind_not_nan);
                    if ~isempty(j)
                        % Standard deviation of Gaussian approxiamtion of
                        % profile
                        Sigma = pinv(parameters.MS.hessian(ind_not_nan,ind_not_nan,1));
                        sigma = sqrt(Sigma(j,j));
                        % Get grid
                        par_grid = parameters.MS.par(i,1) + sigma*linspace(-4,4,100);
                        par_grid = par_grid(find((parameters.min(i) <= par_grid).*(par_grid <= parameters.max(i))));
                        % Calculation of objectiev function approximation
                        J = parameters.MS.gradient(i,1)*(par_grid-parameters.MS.par(i,1)) + 0.5*((par_grid-parameters.MS.par(i,1))/sigma).^2;
                        % Plot
                        h = plot(par_grid,exp(-J),'-','linewidth',options.A.lw,'color',options.A.col); hold on;
                    end
                else
                    warning('No hessian provided in .MS. Approximation in not plotted.');
                end
            else
                
            end
        case 2
            if isfield(parameters.MS,'hessian')
                % negative log-likelihood
                Sigma = pinv(parameters.MS.hessian(:,:,1));
                sigma = sqrt(Sigma(i,i));
                % Get grid
                par_grid = parameters.MS.par(i,1) + sigma*linspace(-4,4,100);
                par_grid = par_grid(find((parameters.min(i) <= par_grid).*(par_grid <= parameters.max(i))));
                % Calculation of objectiev function approximation
                % - with non-zero gradient
%                 ind_I = [1:i-1,i+1:parameters.number];
%                 dtheta_i = -parameters.MS.par(i,1)+par_grid;
%                 dtheta_ind_I = -pinv(parameters.MS.hessian(ind_I,ind_I,1))*bsxfun(@plus,parameters.MS.hessian(ind_I,i,1)*dtheta_i,parameters.MS.gradient(ind_I,1));
%                 dtheta = [dtheta_ind_I(1:i-1,:);dtheta_i;dtheta_ind_I(i:end,:)];
%                 J = nan(1,size(dtheta,2));
%                 for l = 1:size(dtheta,2)
%                     J(l) = parameters.MS.gradient(:,1)'*dtheta(:,l) + 0.5*dtheta(:,l)'*parameters.MS.hessian(:,:,1)*dtheta(:,l);
%                 end
                J = parameters.MS.gradient(i,1)*(par_grid-parameters.MS.par(i,1)) + 0.5*((par_grid-parameters.MS.par(i,1))/sigma).^2;
                % - with zero gradient
%                 J = 0.5*((par_grid-parameters.MS.par(i,1))/sigma).^2;
                % Plot
                h = plot(par_grid,J,'-','linewidth',options.A.lw,'color',options.A.col); hold on;
            else
                warning('No hessian provided in .MS. Approximation in not plotted.');
            end
        otherwise
            error('Selected value for ''options.A.plot_type'' is not available.');
    end
    if ~isempty(h)
        legh(end+1) = h;
        legs{end+1} = options.A.name;
    end

    % Plot: Profile likelihood
    h = [];
    switch options.P.plot_type * flag_plot_P
        case 0
            % no plot
        case 1
            % likelihood ratio
            h = plot(parameters.P(i).par(i,:),exp(parameters.P(i).logPost - logPost_max),'-','linewidth',options.P.lw,'color',options.P.col); hold on;
        case 2
            % negative log-likelihood
            h = plot(parameters.P(i).par(i,:),parameters.P(i).logPost,'-','linewidth',options.P.lw,'color',options.P.col); hold on;
        otherwise
            error('Selected value for ''options.P.plot_type'' is not available.');
    end
    if ~isempty(h)
        legh(end+1) = h;
        legs{end+1} = options.P.name;
    end
    
    % Plot: Additional points
    h = [];
    if ~isempty(options.add_points.par)
        % Check dimension:
        if size(options.add_points.par,1) ~= parameters.number
            warning(['The matrix options.add_points.par should possess ' num2str(parameters.number) ' rows.']);
        else
            for j = 1:size(options.add_points.par,2)
                if size(options.add_points.col,1) == size(options.add_points.par,2)
                    l = j;
                else
                    l = 1;
                end
                h = plot(options.add_points.par(i,j)*[1,1],[0,1.05],options.add_points.ls,'color',options.add_points.col(l,:),'linewidth',options.add_points.lw);
            end
        end
    end
    if ~isempty(h)
        legh(end+1) = h;
        legs{end+1} = options.add_points.name;
    end
    
    % Bounds
    if (options.P.plot_type >= 1) * flag_plot_P
        switch options.boundary.mark
            case 0
                % no plot
            case 1
                ind = find(sum( bsxfun(@gt,parameters.min+options.boundary.eps,parameters.P(i).par)...
                               +bsxfun(@gt,parameters.P(i).par,parameters.max-options.boundary.eps),1));
                if ~isempty(ind)
                    switch options.P.plot_type
                        case 1
                            % likelihood ratio
                            plot(parameters.P(i).par(i,ind),exp(parameters.P(i).logPost(ind) - logPost_max),'x','linewidth',options.P.lw,'color',options.P.col); hold on;    
                        case 2
                            % negative log-likelihood
                            plot(parameters.P(i).par(i,ind),parameters.P(i).logPost(ind),'x','linewidth',options.P.lw,'color',options.P.col); hold on;    
                    end
                end
            otherwise
                error('Selected value for ''options.boundary.mark'' is not available.');
        end
    end
    
    % Plot: Local optima
    if isfield(parameters,'MS')
        h_conv = [];
        h_nconv = [];
        if options.MS.only_optimum
            ind = 1;
        else
            ind = find(parameters.MS.logPost >= parameters.MS.logPost(1)-chi2inv(options.CL.alpha,dof)/2);
        end
        ind_conv = ind(find(min((parameters.MS.exitflag(ind) > 0)+(parameters.MS.exitflag(ind) == -3),1)));
        ind_nconv = setdiff(ind,ind_conv);
        switch options.MS.plot_type
            case 0
                % no plot
            case 1
                % likelihood ratio
                h_conv = plot(parameters.MS.par(i,ind_conv),exp(parameters.MS.logPost(ind_conv)-logPost_max),'o','linewidth',options.MS.lw,'color',options.MS.col); hold on;
                h_nconv = plot(parameters.MS.par(i,ind_nconv),exp(parameters.MS.logPost(ind_nconv)-logPost_max),'s','linewidth',options.MS.lw,'color',options.MS.col);
            case 2
                % negative log-likelihood
                h_conv = plot(parameters.MS.par(i,ind_conv),parameters.MS.logPost(ind_conv),'o','linewidth',options.MS.lw,'color',options.MS.col); hold on;
                h_nconv = plot(parameters.MS.par(i,ind_nconv),parameters.MS.logPost(ind_nconv),'s','linewidth',options.MS.lw,'color',options.MS.col); hold on;
            otherwise
                error('Selected value for ''options.MS.plot_type'' is not available.');
        end
        if ~isempty(h_conv)
            legh(end+1) = h_conv;
            legs{end+1} = options.MS.name_conv;
        end
        if ~isempty(h_nconv)
            legh(end+1) = h_nconv;
            legs{end+1} = options.MS.name_nconv;
        end
    end
    
    % Limits
    % x
    if strcmp(options.interval,'static')
        xl = [parameters.min(i),parameters.max(i)];
    end
    xlim(xl);

    % y
    switch options.P.plot_type
        case {0,1}
            % likelihood ratio
            ylim([0,1.1]);
        case 2
            % Best choice not clear => automatic assignment
    end
    
    % Plot: Confidence levels
    h = [];
    switch options.CL.plot_type
        case 0
            % no plot
        case 1
            % likelihood ratio
            if max(strcmp(options.CL.type,'point-wise'))
                plot(xl,[1,1]*exp(-chi2inv(options.CL.alpha,1)/2),'--','color',options.CL.col);
            end
            if max(strcmp(options.CL.type,'simultanous'))
                plot(xl,[1,1]*exp(-chi2inv(options.CL.alpha,parameters.number)/2),':','linewidth',options.CL.lw,'color',options.CL.col);
            end
        case 2
            % negative log-likelihood
            if max(strcmp(options.CL.type,'point-wise'))
                plot(xl,[1,1]*(parameters.MS.logPost(1)-chi2inv(options.CL.alpha,1)/2),'--','linewidth',options.CL.lw,'color',options.CL.col);
            end
            if max(strcmp(options.CL.type,'simultanous'))
                plot(xl,[1,1]*(parameters.MS.logPost(1)-chi2inv(options.CL.alpha,parameters.number)/2),':','linewidth',options.CL.lw,'color',options.CL.col);
            end
        otherwise
            error('Selected value for ''options.CL.plot_type'' is not available.');
    end
    if ~isempty(h)
        legh(end+1) = h;
        legs{end+1} = options.CL.name;
    end
    
    % Labels
    xlabel(parameters.name(i));
    if (mod(options.subplot_indexing_1D(l),options.subplot_size_1D(2)) == 1) || (length(I) == 1) || options.labels.y_always
        if isempty(options.labels.y_name)
            switch options.CL.plot_type
                case 0
                    % no plot
                    ylabel('post. prob., p');
                case 1
                    % likelihood ratio
                    ylabel('ratio, R');
                case 2
                    % negative log-likelihood
                    ylabel('log-profile, log(PL)');
            end
        else
            ylabel(options.labels.y_name);
        end
    else
        set(gca,'Ytick',[]);
    end
    set(gca,'fontsize',options.fontsize.tick);

    % Legend
    if l == 1
        if isempty(options.legend.position)
            legend(legh,legs,'color',options.legend.color,'box',options.legend.box,'orientation',options.legend.orientation);
        else
            legend(legh,legs,'color',options.legend.color,'box',options.legend.box,'orientation',options.legend.orientation,'position',options.legend.position);
        end
    end
end

end


%% 2D Parameter distributions
if strcmp(type,'2D')
    
% Loop: Parameter
for l1 = 1:length(I)
for l2 = 1:length(I)
    % Initialization of legend
    legh = [];
    legs = {};

    % Assign parameter index
    i1 = I(l1);
    i2 = I(l2);
    
    % Open subplot
    d = (1-options.op2D.b1-options.op2D.b2)/length(I);
    subplot('Position',[options.op2D.b1+(l1-1)*d,...
                        options.op2D.b1+(length(I)-l2)*d,...
                        options.op2D.r*d,options.op2D.r*d]);
    
    % Hold on/off
    if options.hold_on
        hold on;
    else
        hold off;
    end
    
    % Boundaries
    switch options.interval
        case 'dynamic'
            xl1 = [+inf,-inf];
            xl2 = [+inf,-inf];
            
            flag_plot_P_i1 = 0;
            flag_plot_P_i2 = 0;
            if options.P.plot_type >= 1
                if length(parameters.P) >= i1
                    if ~isempty(parameters.P(i1).par)
                        xl1(1) = min(xl1(1),min(parameters.P(i1).par(i1,:)));
                        xl1(2) = max(xl1(2),max(parameters.P(i1).par(i1,:)));
                        flag_plot_P_i1 = 1;
                    end
                end
                if length(parameters.P) >= i2
                    if ~isempty(parameters.P(i2).par)
                        xl2(1) = min(xl2(1),min(parameters.P(i2).par(i2,:)));
                        xl2(2) = max(xl2(2),max(parameters.P(i2).par(i2,:)));
                        flag_plot_P_i2 = 1;
                    end
                end
            end

            if options.S.plot_type >= 1
                xl1(1) = min(xl1(1),min(parameters.S.par(i1,:)));
                xl1(2) = max(xl1(2),max(parameters.S.par(i1,:)));
                xl2(1) = min(xl2(1),min(parameters.S.par(i2,:)));
                xl2(2) = max(xl2(2),max(parameters.S.par(i2,:)));
            end

        case 'static'
            if ~isempty(options.bounds)
                xl1 = [options.bounds.min(i1),options.bounds.max(i1)];
                xl2 = [options.bounds.min(i2),options.bounds.max(i2)];
            else
                xl1 = [parameters.min(i1),parameters.max(i1)];
                xl2 = [parameters.min(i2),parameters.max(i2)];
            end
    end
                    
    % Plot: MCMC samples of tempered posterior distribution
    h = [];
    switch options.S.plot_type
        case 0
            % no plot
        case 1
            % scatter plot
             for k = 1:options.S.ind
                 h = plot(parameters.S.par(i1,:,k),parameters.S.par(i2,:,k),options.S.sp_m,...
                     'color',options.S.sp_col(k,:),'markersize',options.S.sp_ms); hold on;
             end
        case 2
            % kernel-density estimate
             for k = options.S.ind:-1:1
                 x1_line = linspace(min(parameters.S.par(i1,:,k)),max(parameters.S.par(i1,:,k)),100);
                 x2_line = linspace(min(parameters.S.par(i2,:,k)),max(parameters.S.par(i2,:,k)),100);
                 [x1_grid, x2_grid] = meshgrid(x1_line, x2_line);
                 x_grid = transpose([x1_grid(:), x2_grid(:)]);
                 [KDest] = getKernelDensityEstimate([squeeze(parameters.S.par(i1,:,k)); squeeze(parameters.S.par(i2,:,k))], x_grid);
                 KDest = reshape(KDest, length(x1_line), length(x2_line));
                 [~,h] = contour(x1_line, x2_line, KDest/max(max(KDest)),'-','color',options.S.sp_col(k,:),'linewidth',options.S.lw); 
                 hold on;
             end
        otherwise
            error('Selected value for ''options.S.plot_type'' is not available.');
    end
    if ~isempty(h)
        legh(end+1) = h;
        legs{end+1} = options.S.name;
    end

    % Plot: Local approximation
    h = [];
    switch options.A.plot_type
        case 0
            % no plot
        case {1,2}
            if strcmp(options.MCMC, 'user-provided')
                if isfield(parameters, 'user')
                    userProv = 'all';
                    if ((~isfield(parameters.user, 'theta_0')) || isempty(parameters.user.theta_0))
                        userProv = 'sigmaOnly';
                    end
                    if ((~isfield(parameters.user, 'Sigma_0')) || isempty(parameters.user.Sigma_0))
                        userProv = 'no';
                    end
                else
                    userProv = 'no';
                end
            else
                userProv = 'no';
            end
            
            plot_appr = false;
            
            switch userProv
                case 'no'
                    if (isfield(parameters, 'MS') && isfield(parameters.MS, 'hessian') && (size(parameters.MS.hessian,3) >= 1))
                        Sigma = pinv(parameters.MS.hessian([i1,i2],[i1,i2],1));
                        theta_0 = parameters.MS.par([i1,i2],1);
                        plot_appr = true;
                    elseif isfield(parameters, 'MS')
                        warning('No valid values for sigma found! No plotting approximation.');
                    end
                case 'sigmaOnly'
                    if (~isfield(parameters, 'MS') || ~isfield(parameters.MS, 'par') || isempty(parameters.MS.par,3))
                        Sigma = parameters.user.Sigma_0([i1,i2],[i1,i2]);
                        theta_0 = parameters.MS.par([i1,i2],1);
                        plot_appr = true;
                    else
                        warning('No valid values for theta found! No plotting approximation.');
                    end
                case 'all'
                    Sigma = parameters.user.Sigma_0([i1,i2],[i1,i2]);
                    theta_0 = parameters.user.theta_0([i1,i2]);
                    plot_appr = true;
            end
            
            if plot_appr
                X = getEllipse(theta_0, Sigma, options.A.sigma_level);
                h = plot(X(1,:),X(2,:),'-','linewidth',options.A.lw/1.5,'color',options.A.col); hold on;
            end
            
        otherwise
            error('Selected value for ''options.A.plot_type'' is not available.');
    end
    if ~isempty(h)
        legh(end+1) = h;
        legs{end+1} = options.A.name;
    end
    
    % Plot: Local optima
    if isfield(parameters,'MS')
        h_conv = [];
        h_nconv = [];
        if options.MS.only_optimum
            ind = 1;
        else
            ind = find(parameters.MS.logPost >= parameters.MS.logPost(1)-chi2inv(options.CL.alpha,dof)/2);
        end
        ind_conv = ind(find(min((parameters.MS.exitflag(ind) > 0)+(parameters.MS.exitflag(ind) == -3),1)));
        ind_nconv = setdiff(ind,ind_conv);
        switch options.P.plot_type
            case 0
                % no plot
            case {1,2}
                h_conv = plot(parameters.MS.par(i1,ind_conv),parameters.MS.par(i2,ind_conv),'o','linewidth',options.MS.lw,'color',options.MS.col); hold on;
                h_nconv = plot(parameters.MS.par(i1,ind_nconv),parameters.MS.par(i2,ind_nconv),'s','linewidth',options.MS.lw,'color',options.MS.col); hold on;
            otherwise
                error('Selected value for ''options.MS.plot_type'' is not available.');
        end
        if ~isempty(h_conv)
            legh(end+1) = h_conv;
            legs{end+1} = options.MS.name_conv;
        end
        if ~isempty(h_nconv)
            legh(end+1) = h_nconv;
            legs{end+1} = options.MS.name_nconv;
        end
    end
    % Plot: Profile likelihood
    h = [];
    switch options.P.plot_type
        case 0
            % no plot
        case {1,2}
            if flag_plot_P_i1
                h = plot(parameters.P(i1).par(i1,:),parameters.P(i1).par(i2,:),'-','linewidth',options.P.lw,'color',options.P.col*0.8); hold on;
            end
            if flag_plot_P_i2
                h = plot(parameters.P(i2).par(i1,:),parameters.P(i2).par(i2,:),'-','linewidth',options.P.lw,'color',options.P.col*0.6); hold on;
            end
        otherwise
            error('Selected value for ''options.P.plot_type'' is not available.');
    end
    if ~isempty(h)
        legh(end+1) = h;
        legs{end+1} = options.P.name;
    end
    
    % Plot: Additional points
    h = [];
    if ~isempty(options.add_points.par)
        % Check dimension:
        if size(options.add_points.par,1) ~= parameters.number
            warning(['The matrix options.add_points.par should possess ' num2str(parameters.number) ' rows.']);
        else
            for j = 1:size(options.add_points.par,2)
                if size(options.add_points.col,1) == size(options.add_points.par,2)
                    l = j;
                else
                    l = 1;
                end
                h = plot(options.add_points.par(i1,j),options.add_points.par(i2,j),options.add_points.m,...
                    'color',options.add_points.col(l,:),'linewidth',options.add_points.lw,'markersize',options.add_points.ms);
            end
        end
    end
    if ~isempty(h)
        legh(end+1) = h;
        legs{end+1} = options.add_points.name;
    end
    
    % Bounds
    switch options.boundary.mark
        case 0
            % no plot
        case 1
            % i1
            if length(parameters.P) >= i1
                if ~isempty(parameters.P(i1).par)
                    ind = find(sum( bsxfun(@gt,parameters.min+options.boundary.eps,parameters.P(i1).par)...
                                    +bsxfun(@gt,parameters.P(i1).par,parameters.max-options.boundary.eps),1));
                    if ~isempty(ind)
                        switch options.P.plot_type
                            case {1,2}
                                plot(parameters.P(i1).par(i1,ind),parameters.P(i1).par(i2,ind),'x','linewidth',options.P.lw,'color',options.P.col*0.8); hold on;    
                        end
                    end
                end
            end
            
            % i2
            if length(parameters.P) >= i2
                if ~isempty(parameters.P(i2).par)
                    ind = find(sum( bsxfun(@gt,parameters.min+options.boundary.eps,parameters.P(i2).par)...
                                    +bsxfun(@gt,parameters.P(i2).par,parameters.max-options.boundary.eps),1));
                    if ~isempty(ind)
                        switch options.P.plot_type
                            case {1,2}
                                plot(parameters.P(i2).par(i1,ind),parameters.P(i2).par(i2,ind),'x','linewidth',options.P.lw,'color',options.P.col*0.6); hold on;    
                        end
                    end
                end
            end
        otherwise
            error('Selected value for ''options.boundary.mark'' is not available.');
    end
        
    % Limits
    if ~isinf(xl1(1))
        xlim(xl1);
    end
    if ~isinf(xl2(1))
        ylim(xl2);
    end

    % Labels
    if l2 == length(I)
        xlabel(parameters.name(i1));
    else
        set(gca,'xticklabel',[]);
    end
    if i1 == 1
        ylabel(parameters.name(i2));
    else
        set(gca,'yticklabel',[]);
    end
    set(gca,'fontsize',options.fontsize.tick);
    
    % Legend
    if (l1 == 1) && (l2 == 1)
        if isempty(options.legend.position)
            legend(legh,legs,'color',options.legend.color,'box',options.legend.box,'orientation',options.legend.orientation);
        else
            legend(legh,legs,'color',options.legend.color,'box',options.legend.box,'orientation',options.legend.orientation,'position',options.legend.position);
        end
    end

end
end

end


%% Update plot
drawnow;
```
