## Supplementary material for "Mechanistic description of spatial processes using integrative modelling of noise-corrupted imaging data": plotPropertyMultiStarts.m

```
function fh = plotPropertyMultiStarts(properties, varargin)
% plotPropertyMultiStarts plots the result of the multi-start optimization stored in properties.
%
% USAGE:
% fh = plotPropertyMultiStarts(properties)
% fh = plotPropertyMultiStarts(properties,fh)
% fh = plotPropertyMultiStarts(properties,fh,options)
%
% Parameters:
% varargin:
% properties: property struct containing information about properties
%   and log-posterior.
% fh: handle of figure in which profile likelihood is plotted. If no
%   figure handle is provided, a new figure is opened.
% options: options of plotting as instance of PestoPlottingOptions
%
% Return values:
% fh: figure handle
%
% History:
% * 2015/03/03 Jan Hasenauer

%% CHECK AND ASSIGN INPUTS
% Open figure
if length(varargin) >= 1 && ~isempty(varargin{1})
    fh = figure(varargin{1});
else
    fh = figure('Name','plotPropertyMultiStarts');
end

% Check, if properties has all necessary fieds
properties = checkSanityOfStructs(properties, 'properties');

% Options
if length(varargin) >= 2
    options = handlePlottingOptionArgument(varargin{2});
else
    options = PestoPlottingOptions();
end

%% ASSIGN COLORS
i = length(properties.MS.logPost);
Col = colormap(gray(i+ceil(i/3)));
Col = Col.^(1/3);
Col(1,:) = [1,0,0];

%% PLOT OBJECTIVES
subplot(2,2,1);
plot(1:i,properties.MS.logPost,'-','color',0.9*[1,1,1],'linewidth',2); hold on;
for j = i:-1:1
    plot(j,properties.MS.logPost(j),'o','color',Col(j,:),'linewidth',2); hold on;
end
hold off;
xlim([1-0.2,i+0.2]);
xlabel('start');
ylabel('log-likelihood');
if options.title
    title('all estimates');
end

%% PLOT TOP TEN OBJECTIVES
subplot(2,2,3);
plot(1:min(i,10),properties.MS.logPost(1:min(i,10)),'-','color',0.9*[1,1,1],'linewidth',2); hold on;
for j = min(i,10):-1:1
    plot(j,properties.MS.logPost(j),'o','color',Col(j,:),'linewidth',2); hold on;
end
hold off;
xlim([1-0.2,min(i,10)+0.2]);

ylim([min(properties.MS.logPost(1),min(properties.MS.logPost(1:min(i,10)))-1),properties.MS.logPost(1)+1]);
xlabel('start');
ylabel('log-likelihood');
if options.title
    title('top 10 estimates');
end

%% PLOT PROPERTIES
subplot(2,2,[2,4]);
for j = i:-1:1
    plot(properties.MS.prop(:,j)',1:properties.number,'-o','color',Col(j,:),'linewidth',2); hold on;
end
plot(properties.MS.prop(:,1)',1:properties.number,'r-o','linewidth',2); hold on;
if options.draw_bounds
    plot(properties.min([1,1:properties.number,properties.number])',[0.99,1:properties.number,properties.number+0.01],'b--','linewidth',2); hold on;
    plot(properties.max([1,1:properties.number,properties.number])',[0.99,1:properties.number,properties.number+0.01],'b--','linewidth',2); hold on;
end
hold off;
ylim([1-0.01,properties.number+0.01]);
ylabel(' ');
xlabel('parameters values');
set(gca,'ytick',1:properties.number,'yticklabel',properties.name)

if options.title
    title('estimated parameters');
end
drawnow;
```
