## Supplementary material for "Mechanistic description of spatial processes using integrative modelling of noise-corrupted imaging data": plotPropertyProfiles.m

```
function fh = plotPropertyProfiles(properties, varargin)
% plotPropertyProfiles.m visualizes profile likelihood of model properties.
% Note: This routine provides an interface for plotPropertyUncertainty.m.
%
% USAGE:
% fh = plotPropertyProfiles(properties)
% fh = plotPropertyProfiles(properties,type)
% fh = plotPropertyProfiles(properties,type,fh)
% fh = plotPropertyProfiles(properties,type,fh,I)
% fh = plotPropertyProfiles(properties,type,fh,I,options)
%
% Parameters:
%   properties: property struct containing information about properties
%       and results of optimization (.MS) and uncertainty analysis
%       (.P and .S).
%   varargin:
%     type: string indicating the type of visualization: '1D' or '2D'
%     fh: handle of figure. If no figure handle is provided, a new figure
%         is opened.
%     I: index of properties which are updated. If no index is provided
%         all properties are updated.
%     options: options of plotting as instance of PestoPlottingOptions
%
% Return values:
%   fh: figure handle
%
% History:
% * 2015/03/02 Jan Hasenauer
% * 2016/10/10 Daniel Weindl

switch length(varargin) 
    case 0
        fh = plotPropertyUncertainty(properties);
    case 1
        fh = plotPropertyUncertainty(properties, varargin{1});
    case 2
        fh = plotPropertyUncertainty(properties, varargin{1}, varargin{2});
    case 3
        fh = plotPropertyUncertainty(properties, varargin{1}, varargin{2}, varargin{3});
    case 4
        fh = plotPropertyUncertainty(properties, varargin{1}, varargin{2}, varargin{3}, varargin{4});
    otherwise
        error('Too many arguments for plotPropertyProfiles().');
end
end
```
