## Supplementary material for "Mechanistic description of spatial processes using integrative modelling of noise-corrupted imaging data": plotPropertyUncertainty.m

```
function fh = plotPropertyUncertainty(properties, varargin)
% plotPropertyUncertainty.m visualizes profile likelihood and MCMC samples
% stored in properties.
%
% USAGE:
% fh = plotPropertyUncertainty(properties,type)
% fh = plotPropertyUncertainty(properties,type,fh)
% fh = plotPropertyUncertainty(properties,type,fh,I)
% fh = plotPropertyUncertainty(properties,type,fh,I,options)
%
% Parameters:
%   properties: properties struct.
%   varargin:
%     type: string indicating the type of visualization: '1D'
%     fh: handle of figure. If no figure handle is provided, a new figure
%         is opened.
%     I: index of properties which are updated. If no index is provided
%         all parameters are updated.
%     options: options of plotting as instance of PestoPlottingOptions
%
% Return values:
%   fh: figure handle
%
% History:
% * 2012/05/31 Jan Hasenauer
% * 2014/06/20 Jan Hasenauer
% * 2016/10/10 Daniel Weindl

%% Check and assign inputs
% Plot type
type = '1D';
if length(varargin) >= 1 && ~isempty(varargin{1})
    type = varargin{1};
    if ~max(strcmp({'1D','2D'},type))
        error('The ''type'' of plot is unknown.')
    end
end

% Check, if properties has all necessary fieds
properties = checkSanityOfStructs(properties, 'properties');

% Open figure
if length(varargin) >= 2 && ~isempty(varargin{2})
    fh = figure(varargin{2});
else
    fh = figure('Name','plotPropertyUncertainty');
end

% Index of subplot which is updated
I = 1:properties.number;
if length(varargin) >= 3 && ~isempty(varargin{3})
    I = varargin{3};
    if ~isnumeric(I) || max(abs(I - round(I)) > 0)
        error('I is not an integer vector.');
    end
end

% Options
defaultOptions = PestoPlottingOptions();
if isfield(properties,'S')
    if isfield(properties.S,'PT');
        defaultOptions.S.PT.plot_type = defaultOptions.S.plot_type;
        defaultOptions.S.PT.ind = 1:size(properties.S.PT.prop,3);
        defaultOptions.S.PT.col = [linspace(0,1,size(properties.S.PT.prop,3))',...
                            0.2*ones(size(properties.S.PT.prop,3),1),...
                            linspace(1,0,size(properties.S.PT.prop,3))'];
    end
end

if ~isfield(properties,'MS')
    defaultOptions.MS.plot_type = 0; 
end

% Assignment of user-provided options
if length(varargin) >= 4
    options = handlePlottingOptionArgument(varargin{4});
    options = setdefault(options, defaultOptions);
else
    options = defaultOptions;
end
if ~isfield(properties,'P')
    options.P.plot_type = 0; 
end

if ~isfield(properties,'S')
    options.S.plot_type = 0; 
end


% Subplot arrangement
if ~isfield(options,'subplot_size_1D')
    options.subplot_size_1D = round(sqrt(length(I))*[1,1]);
    if prod(options.subplot_size_1D) < length(I)
        options.subplot_size_1D(2) = options.subplot_size_1D(2) + 1;
    end
end
if ~isfield(options,'subplot_indexing_1D')
    options.subplot_indexing_1D = 1:length(I);
end

%% INITALIZATION
% Maximum a posterior estimate
if isfield(properties,'MS')
    logPost_max = max(properties.MS.logPost);
end

% Degrees of freedom (for chi^2 test)
dof = 1;
if max(strcmp(options.CL.type,'simultanous'))
    dof = properties.number;
end

%% 1D Parameter distributions
if strcmp(type,'1D')
    
% Compute number of subfigure
s = round(sqrt(length(I))*[1,1]);
if prod(s) < length(I)
    s(2) = s(2) + 1;
end

% Loop: Parameter
for l = 1:length(I)
    % Initialization of legend
    legh = [];
    legs = {};

    % Assign parameter index
    i = I(l);
    
    % Open subplot
    subplot(options.subplot_size_1D(1),options.subplot_size_1D(2),options.subplot_indexing_1D(l));
        
    % Hold on/off
    if options.hold_on
        hold on;
    else
        hold off;
    end
    
    % Boundaries
    switch options.interval
        case 'dynamic'
            xl = [+inf,-inf];
            
            if isfield(properties,'MS')
                if max(strcmp(options.CL.type,'point-wise'))
                    L = find(properties.MS.logPost(:) > (properties.MS.logPost(1)-chi2inv(options.CL.alpha,1)/2));
                end
                if max(strcmp(options.CL.type,'simultanous'))
                    L = find(properties.MS.logPost(:) > (properties.MS.logPost(1)-chi2inv(options.CL.alpha,properties.numbe)/2));
                end
                xl(1) = min(xl(1),min(properties.MS.prop(i,L)));
                xl(2) = max(xl(2),max(properties.MS.prop(i,L)));
            end
            
            flag_plot_P = 0;
            if options.P.plot_type >= 1
                if length(properties.P) >= i
                    if ~isempty(properties.P(i).par)
                        xl(1) = min(xl(1),min(properties.P(i).prop));
                        xl(2) = max(xl(2),max(properties.P(i).prop));
                        flag_plot_P = 1;
                    end
                end
            end

            if options.S.plot_type >= 1
                xl(1) = min(xl(1),min(properties.S.prop(i,:)));
                xl(2) = max(xl(2),max(properties.S.prop(i,:)));
            end
            
            if xl(1) == xl(2)
                xl(1) = xl(1) - 1e-10;
                xl(2) = xl(2) + 1e-10;
            end
        case 'static'
            if isfield(options,bounds)
                xl = [options.bounds.min(i),options.bounds.max(i)];
            else
                xl = [properties.min(i),properties.max(i)];
            end
    end

    % Plot: Visualizaion of MCMC samples of posterior distribution
    h = [];
    switch options.S.plot_type
        case 0
            % no plot
        case 1
            % histogram
            S = properties.S.prop(i,~isnan(properties.S.prop(i,:)));
            switch options.S.bins
                case 'optimal'
                    b = 3.49*std(S)/(length(S)^(1/3));
                    nbin = round((max(S)-min(S))/b);
                case 'conservative'
                    b = 2*3.49*std(S)/(length(S)^(1/3));
                    nbin = round((max(S)-min(S))/b);
                otherwise
                    nbin = options.S.bins;
            end
            [N,X] = hist(S,nbin);
            h = bar(X,N/max(N),1,'facecolor',options.S.hist_col); hold on;
        case 2
            % kernel-density estimate
            x_grid = linspace(min(properties.S.prop(i,:)),max(properties.S.prop(i,:)),100);
            [KDest] = getKernelDensityEstimate(squeeze(properties.S.prop(i,:)),x_grid);
            h = plot(x_grid,KDest/max(KDest),'-','color',options.S.lin_col,'linewidth',options.S.lin_lw); hold on;
        otherwise
            error('Selected value for ''options.S.plot_type'' is not available.');
    end
    if ~isempty(h)
        legh(end+1) = h;
        legs{end+1} = options.S.name;
    end

    % Plot: Local approximation
    h = [];
    switch options.A.plot_type
        case 0
            % no plot
        case 1
            % likelihood ratio
            if isfield(properties.MS,'prop_Sigma')
                % Get grid
                prop_grid = properties.MS.prop(i,1) + sqrt(properties.MS.prop_Sigma(i,i,1))*linspace(-5,5,100);
                prop_grid = prop_grid(find((properties.min(i) <= prop_grid).*(prop_grid <= properties.max(i))));
                % Plot
                h = plot(prop_grid,exp(-0.5*((prop_grid-properties.MS.prop(i,1)).^2/properties.MS.prop_Sigma(i,i,1))),'-','linewidth',options.A.lw,'color',options.A.col); hold on;
            else
                warning('No hessian provided in .MS. Approximation in not plotted.');
            end
        case 2
            % negative log-likelihood
            if isfield(properties.MS,'prop_Sigma')
                % Get grid
                prop_grid = properties.MS.prop(i,1) + sqrt(properties.MS.prop_Sigma(i,i,1))*linspace(-5,5,100);
                prop_grid = prop_grid(find((properties.min(i) <= prop_grid).*(prop_grid <= properties.max(i))));
                % Plot
                h = plot(prop_grid,-logPost_max+0.5*((prop_grid-properties.MS.prop(i,1)).^2/properties.MS.prop_Sigma(i,i,1)),'-','linewidth',options.A.lw,'color',options.A.col); hold on;
            else
                warning('No hessian provided in .MS. Approximation in not plotted.');
            end
        otherwise
            error('Selected value for ''options.A.plot_type'' is not available.');
    end
    if ~isempty(h)
        legh(end+1) = h;
        legs{end+1} = options.A.name;
    end

    % Plot: Profile likelihood
    h = [];
    switch options.P.plot_type * flag_plot_P
        case 0
            % no plot
        case 1
            % likelihood ratio
            h = plot(properties.P(i).prop,exp(properties.P(i).logPost - logPost_max),'-','linewidth',options.P.lw,'color',options.P.col); hold on;
        case 2
            % negative log-likelihood
            h = plot(properties.P(i).prop,properties.P(i).logPost,'-','linewidth',options.P.lw,'color',options.P.col); hold on;
        otherwise
            error('Selected value for ''options.P.plot_type'' is not available.');
    end
    if ~isempty(h)
        legh(end+1) = h;
        legs{end+1} = options.P.name;
    end
    
    % Plot: Additional points
    h = [];
    if ~isempty(options.add_points) && ~isempty(options.add_points.par)
        % Check dimension:
        if size(options.add_points.prop,1) ~= properties.number
            warning(['The matrix options.add_points.par should possess ' num2str(properties.number) ' rows.']);
        else
            for j = 1:size(options.add_points.prop,2)
                if size(options.add_points.col,1) == size(options.add_points.prop,2)
                    l = j;
                else
                    l = 1;
                end
                h = plot(options.add_points.prop(i,j)*[1,1],[0,1.05],options.add_points.ls,'color',options.add_points.col(l,:),'linewidth',options.add_points.lw);
            end
        end
    end
    if ~isempty(h)
        legh(end+1) = h;
        legs{end+1} = options.add_points.name;
    end
        
    % Plot: Local optima
    h_conv = [];
    h_nconv = [];
    if options.MS.only_optimum
        ind = 1;
    else
        ind = find(properties.MS.logPost >= properties.MS.logPost(1)-chi2inv(options.CL.alpha,dof)/2);
    end
    ind_conv = ind(find(min((properties.MS.exitflag(ind) > 0)+(properties.MS.exitflag(ind) == -3),1)));
    ind_nconv = setdiff(ind,ind_conv);
    switch options.MS.plot_type
        case 0
            % no plot
        case 1
            % likelihood ratio
            h_conv = plot(properties.MS.prop(i,ind_conv),exp(properties.MS.logPost(ind_conv)-logPost_max),'o','linewidth',options.MS.lw,'color',options.MS.col); hold on;
            h_nconv = plot(properties.MS.prop(i,ind_nconv),exp(properties.MS.logPost(ind_nconv)-logPost_max),'s','linewidth',options.MS.lw,'color',options.MS.col);
        case 2
            % negative log-likelihood
            h_conv = plot(properties.MS.prop(i,ind_conv),properties.MS.logPost(ind_conv),'o','linewidth',options.MS.lw,'color',options.MS.col); hold on;
            h_nconv = plot(properties.MS.prop(i,ind_nconv),properties.MS.logPost(ind_nconv),'s','linewidth',options.MS.lw,'color',options.MS.col); hold on;
        otherwise
            error('Selected value for ''options.MS.plot_type'' is not available.');
    end
    if ~isempty(h_conv)
        legh(end+1) = h_conv;
        legs{end+1} = options.MS.name_conv;
    end
    if ~isempty(h_nconv)
        legh(end+1) = h_nconv;
        legs{end+1} = options.MS.name_nconv;
    end
        
    % Limits
%     % x
%     if strcmp(options.interval,'static')
%         xl = [properties.min(i),properties.max(i)];
%     end
%     xlim(xl);

    % y
    switch options.P.plot_type
        case {0,1}
            % likelihood ratio
            ylim([0,1.1]);
        case 2
            % Best choice not clear => automatic assignment
    end
    
    % Plot: Confidence levels
    h = [];
    switch options.CL.plot_type
        case 0
            % no plot
        case 1
            % likelihood ratio
            if max(strcmp(options.CL.type,'point-wise'))
                plot(xl,[1,1]*exp(-chi2inv(options.CL.alpha,1)/2),'--','color',options.CL.col);
            end
            if max(strcmp(options.CL.type,'simultanous'))
                plot(xl,[1,1]*exp(-chi2inv(options.CL.alpha,properties.number)/2),':','linewidth',options.CL.lw,'color',options.CL.col);
            end
        case 2
            % negative log-likelihood
            if max(strcmp(options.CL.type,'point-wise'))
                h = plot(xl,[1,1]*(properties.MS.logPost(1)-chi2inv(options.CL.alpha,1)/2),'--','linewidth',options.CL.lw,'color',options.CL.col);
            end
            if max(strcmp(options.CL.type,'simultanous'))
                h = plot(xl,[1,1]*(properties.MS.logPost(1)-chi2inv(options.CL.alpha,properties.number)/2),':','linewidth',options.CL.lw,'color',options.CL.col);
            end
        otherwise
            error('Selected value for ''options.CL.plot_type'' is not available.');
    end
    if ~isempty(h)
        legh(end+1) = h;
        legs{end+1} = options.CL.name;
    end
    
    % Labels
    xlabel(properties.name(i));
    if (mod(options.subplot_indexing_1D(l),options.subplot_size_1D(2)) == 1) || (length(I) == 1) || options.labels.y_always
        if isempty(options.labels.y_name)
            switch options.CL.plot_type
                case 0
                    % no plot
                    ylabel('post. prob., p');
                case 1
                    % likelihood ratio
                    ylabel('ratio, R');
                case 2
                    % negative log-likelihood
                    ylabel('log-profile, log(PL)');
            end
        else
            ylabel(options.labels.y_name);
        end
    else
        set(gca,'Ytick',[]);
    end
    set(gca,'fontsize',options.fontsize.tick);
    
    % Legend
    if l == 1
        if isempty(options.legend.position)
            legend(legh,legs,'color',options.legend.color,'box',options.legend.box,'orientation',options.legend.orientation);
        else
            legend(legh,legs,'color',options.legend.color,'box',options.legend.box,'orientation',options.legend.orientation,'position',options.legend.position);
        end
    end
end

end


%% 2D Parameter distributions
if strcmp(type,'2D')
    
% Loop: Parameter
for l1 = 1:length(I)
for l2 = 1:length(I)
    % Initialization of legend
    legh = [];
    legs = {};

    % Assign parameter index
    i1 = I(l1);
    i2 = I(l2);
    
    % Open subplot
%    subplot(length(I),length(I),(i2-1)*length(I)+i1);
    d = (1-options.op2D.b1-options.op2D.b2)/length(I);
    subplot('Position',[options.op2D.b1+(l1-1)*d,...
                        options.op2D.b1+(length(I)-l2)*d,...
                        options.op2D.r*d,options.op2D.r*d]);
    
    if options.hold_on
        hold on;
    else
        hold off;
    end
    
    % Boundaries
    switch options.interval
        case 'dynamic'
            xl1 = [+inf,-inf];
            xl2 = [+inf,-inf];
            
            if options.P.plot_type >= 1
                flag_plot_P_i1 = 0;
                if length(properties.P) >= i1
                    if ~isempty(properties.P(i1).prop)
                        xl1(1) = min(xl1(1),min(properties.P(i1).prop(:)));
                        xl1(2) = max(xl1(2),max(properties.P(i1).prop(:)));
                        flag_plot_P_i1 = 1;
                    end
                end
                flag_plot_P_i2 = 0;
                if length(properties.P) >= i2
                    if ~isempty(properties.P(i2).prop)
                        xl2(1) = min(xl2(1),min(properties.P(i2).prop(:)));
                        xl2(2) = max(xl2(2),max(properties.P(i2).prop(:)));
                        flag_plot_P_i2 = 1;
                    end
                end
            end

            if options.S.plot_type >= 1
                xl1(1) = min(xl1(1),min(properties.S.prop(i1,:)));
                xl1(2) = max(xl1(2),max(properties.S.prop(i1,:)));
                xl2(1) = min(xl2(1),min(properties.S.prop(i2,:)));
                xl2(2) = max(xl2(2),max(properties.S.prop(i2,:)));
            end

        case 'static'
            if isfield(options,bounds)
                xl1 = [options.bounds.min(i1),options.bounds.max(i1)];
                xl2 = [options.bounds.min(i2),options.bounds.max(i2)];
            else
                xl1 = [properties.min(i1),properties.max(i1)];
                xl2 = [properties.min(i2),properties.max(i2)];
            end
    end

    % Plot: MCMC samples
    h = [];
    switch options.S.plot_type
        case 0
            % no plot
        case 1
            % scatter plot
            h = plot(properties.S.prop(i1,:),properties.S.prop(i2,:),options.S.sp_m,...
                'color',options.S.sp_col,'markersize',options.S.sp_ms); hold on;
        otherwise
            error('Selected value for ''options.S.plot_type'' is not available.');
    end
    if ~isempty(h)
        legh(end+1) = h;
        legs{end+1} = options.S.name;
    end

    % Plot: Local approximation
    h = [];
    switch options.A.plot_type
        case 0
            % no plot
        case {1,2}
            if isfield(properties.MS,'prop_Sigma')
                % plot
                X = getEllipse(properties.MS.prop([i1,i2],1),properties.MS.prop_Sigma([i1,i2],[i1,i2],1),options.A.sigma_level);
                h = plot(X(1,:),X(2,:),'-','linewidth',options.A.lw/1.5,'color',options.A.col); hold on;
            else
                warning('No covariance matrix provided in properties.MS. Approximation in not plotted.');
            end
        otherwise
            error('Selected value for ''options.A.plot_type'' is not available.');
    end
    if ~isempty(h)
        legh(end+1) = h;
        legs{end+1} = options.A.name;
    end

    % Plot: Local optima
    h_conv = [];
    h_nconv = [];
    ind = find(properties.MS.logPost >= properties.MS.logPost(1)-chi2inv(options.CL.alpha,dof)/2);
    ind_conv = ind(find(min((properties.MS.exitflag(ind) > 0)+(properties.MS.exitflag(ind) == -3),1)));
    ind_nconv = setdiff(ind,ind_conv);
    switch options.P.plot_type
        case 0
            % no plot
        case {1,2}
            h_conv = plot(properties.MS.prop(i1,ind_conv),properties.MS.prop(i2,ind_conv),'o','linewidth',options.MS.lw,'color',options.MS.col); hold on;
            h_nconv = plot(properties.MS.prop(i1,ind_nconv),properties.MS.prop(i2,ind_nconv),'s','linewidth',options.MS.lw,'color',options.MS.col); hold on;
        otherwise
            error('Selected value for ''options.MS.plot_type'' is not available.');
    end
    if ~isempty(h_conv)
        legh(end+1) = h_conv;
        legs{end+1} = options.MS.name_conv;
    end
    if ~isempty(h_nconv)
        legh(end+1) = h_nconv;
        legs{end+1} = options.MS.name_nconv;
    end

    % Plot: Profile likelihood
    h = [];
    switch options.P.plot_type
        case 0
            % no plot
        case {1,2}
            % Calculation and visualization of property i2 along profile for property i1
            if flag_plot_P_i1
                P_prop = ones(length(properties.P(i1).prop),1);
                for k = 1:length(properties.P(i1).prop)
                    P_prop(k) = properties.function{i2}(properties.P(i1).par(:,k));
                end
                h = plot(properties.P(i1).prop,P_prop,'-','linewidth',options.P.lw,'color',options.P.col*0.8); hold on;
            end
                
            % Calculation and visualization of property i1 along profile for property i2
            if flag_plot_P_i2
                P_prop = ones(length(properties.P(i2).prop),1);
                for k = 1:length(properties.P(i2).prop)
                    P_prop(k) = properties.function{i1}(properties.P(i2).par(:,k));
                end
                h = plot(P_prop,properties.P(i2).prop,'-','linewidth',options.P.lw,'color',options.P.col*0.6); hold on;
            end
            
        otherwise
            error('Selected value for ''options.P.plot_type'' is not available.');
    end
    if ~isempty(h)
        legh(end+1) = h;
        legs{end+1} = options.P.name;
    end
    
    % Plot: Additional points
    h = [];
    if ~isempty(options.add_points.par)
        % Check dimension:
        if size(options.add_points.par,1) ~= properties.number
            warning(['The matrix options.add_points.par should possess ' num2str(properties.number) ' rows.']);
        else
            for j = 1:size(options.add_points.par,2)
                if size(options.add_points.col,1) == size(options.add_points.par,2)
                    l = j;
                else
                    l = 1;
                end
                h = plot(options.add_points.par(i1,j),options.add_points.par(i2,j),options.add_points.m,...
                    'color',options.add_points.col(l,:),'linewidth',options.add_points.lw,'markersize',options.add_points.ms);
            end
        end
    end
    if ~isempty(h)
        legh(end+1) = h;
        legs{end+1} = options.add_points.name;
    end
        
    % Limits
    if ~isinf(xl1(1))
        xlim(xl1);
    end
    if ~isinf(xl2(1))
        ylim(xl2);
    end

    % Labels
    if l2 == length(I)
        xlabel(properties.name(i1));
    else
        set(gca,'xticklabel',[]);
    end
    if i1 == 1
        ylabel(properties.name(i2));
    else
        set(gca,'yticklabel',[]);
    end
    set(gca,'fontsize',options.fontsize.tick);
    
    % Legend
    if (l1 == 1) && (l2 == 1)
        if isempty(options.legend.position)
            legend(legh,legs,'color',options.legend.color,'box',options.legend.box,'orientation',options.legend.orientation);
        else
            legend(legh,legs,'color',options.legend.color,'box',options.legend.box,'orientation',options.legend.orientation,'position',options.legend.position);
        end
    end

end
end

end


%% Update plot
drawnow;
```
