## Supplementary material for "Mechanistic description of spatial processes using integrative modelling of noise-corrupted imaging data": barrierFunction.m

```
%% Barrier function interface for optimization
%
% Last change: Paul Stapor, 06/04/17

function varargout = barrierFunction(objective, gradient, theta, borders, iteration, maxIter, method)
%% Documentation of barrierFunction
%
% This function applies barrier or penalty funcions to an objective
% function vvalue and its gradient. The precise method can be chosen by the
% string variable method.


%% Main Routine
    
    if isempty(gradient)
        gradient = zeros(size(theta));
    end
    
    % The input is passed to the different algorithms
    switch(method)
        case 'soft-barrier'
            [objective, gradient] = softBarrier(objective, gradient, theta, borders);
            
        case 'log-barrier'
            [objective, gradient] = logBarrier(objective, gradient, theta, borders, iteration, maxIter);
            
        case 'log-adaptive'           
            [objective, gradient] = logAdaptiveBarrier(objective, gradient, theta, borders, iteration, maxIter);
 
        otherwise
            error('Call to a non-existing update method');
    end
    
    switch nargout
        case 1
            varargout = {objective};
        case 2
            varargout = {objective, gradient};
        case 3
            varargout = {objective, gradient, []};
    end
end


function [objective, gradient] = softBarrier(objective, gradient, theta, borders)
% Very simple barrier function, applies a polynomial of third order to
% those parameters which are outside the borders. Since the input is only
% changed if parameter bounds are violated, it is actually a penalty
% function.
%
% IMPORTANT: The routines assumes the objective function to be a NEGATIVE
% log-posterior, i.e. it is made for minimizing.

    scale = 100;
    
    for iPar = 1 : size(borders, 1)
        % Lower bounds
        if (theta(iPar) < borders(iPar, 1))
            objective = objective + ...
                scale * (borders(iPar, 1) - theta(iPar))^3;
            gradient(iPar) = gradient(iPar) + ...
                3 * scale * (borders(iPar, 1) - theta(iPar))^2;
                
        % Upper bounds
        elseif (theta(iPar) > borders(iPar, 2))
            objective = objective + ...
                scale * (theta(iPar) - borders(iPar, 2))^3;
            gradient(iPar) = gradient(iPar) - ...
                3 * scale * (theta(iPar) - borders(iPar, 2))^2;
            
        end
    end

end


function [objective, gradient] = logBarrier(objective, gradient, theta, borders, iteration, maxIter)
% Simple log-barrier function, inspired from the interior-point algorithm. 
% Applies a negative logarithm to the diffeernce of parameters
% and the bounds. Since it is active also in the interior of the box, this
% is a classical barrier function.
%
% IMPORTANT: The routines assumes the objective function to be a NEGATIVE
% log-posterior, i.e. it is made for minimizing.

    % Scaling for barrier, which takes iteration into account
    scale = 8*(iteration/maxIter) + 2;
    scale = 10^scale;
    
    % Parabola, which determines the barrier
    parabola = - (theta - borders(:,1)) .* (theta - borders(:,2));
    parabola = parabola ./ (0.5*(borders(:,2) - borders(:,1)).^2);
    
    % Setting the values
    barrObjective = - sum(1/scale * log(parabola));
    barrGradient = - 1/scale * (2*theta - theta.*(borders(:,1)+borders(:,2))) ./ ((theta - borders(:,1)) .* (theta - borders(:,2)));
    
    objective = objective + barrObjective;
    gradient  = gradient  + barrGradient;
end
```
