## Supplementary material for "Mechanistic description of spatial processes using integrative modelling of noise-corrupted imaging data": checkSanityOfStructs.m

```
function pStruct = checkSanityOfStructs(pStruct, type)
% checkSanityOfStructs.m checks if the necessary fields of the parameters
% or the properties struct are set. If necessary fields are missing but can
% unambiguously, this will be done. Otherwise, an error will be displayed.
%
% USAGE:
% pStruct = checkSanityOfStructs(pStruct, type)
%
% Parameters:
%   pStruct: parameters or properties struct.
%   type: string, either 'parameters' or 'properties'
%
% Return values:
%   pSturct: struct containing informations about the parameters or the
%       properties
%
% History:
% * 2017/05/21 Paul Stapor


    % See which struct should be checked
    if ~(strcmp(type, 'parameters') || strcmp(type, 'properties'))
        error('The given type of struct is invalid. Only parameters and properties and be checked by this function.');
    else
        if strcmp(type, 'parameters')
            short_type = 'par';
        else
            short_type = 'prop';
        end
    end

    % The least necessary information are lower and upper bounds for parameters
    if ~isfield(pStruct, 'min')
        error(['The struct ' type ' has no lower bounds provided.']);
    end
    if ~isfield(pStruct, 'max')
        error(['The struct ' type ' has no upper bounds provided.']);
    end

    % Upper and lower bounds need to have the same length
    if ~all(size(pStruct.min) == size(pStruct.max))
        error(['The vectors for the lower and upper bounds of' type ' do not have the same size.']); 
    end

    % The number of pStruct needs to be set accordingly
    if ~isfield(pStruct, 'number')
        pStruct.number = length(pStruct.min);
    else
        if (length(pStruct.min) ~= pStruct.number)
            warning(['The size of struct ' type ' was set incorrectly in ' type '.number. This was corrected.']);
        end
    end

    % The names of the parameters have to be set correctly
    if ~isfield(pStruct, 'name')
        pStruct.name = arrayfun(@(x) [short_type num2str(x)], 1:pStruct.number, 'UniformOutput', false);
    else
        if (length(pStruct.name) ~= pStruct.number)
            error(['The names for the struct ' type ' did not have the correct length. Either provide them with correct length or leave ' type '.name empty.']);
        end
    end

end
```
