## Supplementary material for "Mechanistic description of spatial processes using integrative modelling of noise-corrupted imaging data": cmaes.m

```
function [xmin, ...      % minimum search point of last iteration
          fmin, ...      % function value of xmin
          counteval, ... % number of function evaluations done
          stopflag, ...  % stop criterion reached
          out, ...       % struct with various histories and solutions
          bestever ...   % struct containing overall best solution (for convenience)
    ] = cmaes( ...
    fitfun, ...    % name of objective/fitness function
    xstart, ...    % objective variables initial point, determines N
    insigma, ...   % initial coordinate wise standard deviation(s)
    inopts, ...    % options struct, see defopts below
    varargin ...   % arguments passed to objective function 
    )
% cmaes.m, Version 3.61.beta, last change: April, 2012 
% CMAES implements an Evolution Strategy with Covariance Matrix
% Adaptation (CMA-ES) for nonlinear function minimization.  For
% introductory comments and copyright (GPL) see end of file (type 
% 'type cmaes'). cmaes.m runs with MATLAB (Windows, Linux) and, 
% without data logging and plotting, it should run under Octave 
% (Linux, package octave-forge is needed).
%
% OPTS = CMAES returns default options. 
% OPTS = CMAES('defaults') returns default options quietly.
% OPTS = CMAES('displayoptions') displays options. 
% OPTS = CMAES('defaults', OPTS) supplements options OPTS with default 
% options. 
%
% XMIN = CMAES(FUN, X0, SIGMA[, OPTS]) locates an approximate minimum 
% XMIN of function FUN starting from column vector X0 with the initial
% coordinate wise search standard deviation SIGMA.
%
% Input arguments: 
%
%  FUN is the objective function. FUN takes as argument
%     a column vector of size of X0 and returns a scalar. An easy way to
%     implement a hard non-linear constraint is to return NaN. Then,
%     this function evaluation is not counted and a newly sampled
%     point is tried immediately.
%
%   X0 is a column vector, or a matrix, or a string. If X0 is a matrix,
%     mean(X0, 2) is taken as initial point. If X0 is a string like
%     '2*rand(10,1)-1', the string is evaluated first.
%
%   SIGMA is a scalar, or a column vector of size(X0,1), or a string
%     that can be evaluated into one of these. SIGMA determines the
%     initial coordinate wise standard deviations for the search.
%     Setting SIGMA one third of the initial search region is
%     appropriate, e.g., the initial point in [0, 6]^10 and SIGMA=2
%     means cmaes('myfun', 3*rand(10,1), 2).  If SIGMA is missing and
%     size(X0,2) > 1, SIGMA is set to sqrt(var(X0')'). That is, X0 is
%     used as a sample for estimating initial mean and variance of the
%     search distribution.
%
%   OPTS (an optional argument) is a struct holding additional input
%     options. Valid field names and a short documentation can be
%     discovered by looking at the default options (type 'cmaes'
%     without arguments, see above). Empty or missing fields in OPTS
%     invoke the default value, i.e. OPTS needs not to have all valid
%     field names.  Capitalization does not matter and unambiguous
%     abbreviations can be used for the field names. If a string is
%     given where a numerical value is needed, the string is evaluated
%     by eval, where 'N' expands to the problem dimension
%     (==size(X0,1)) and 'popsize' to the population size. 
%
% [XMIN, FMIN, COUNTEVAL, STOPFLAG, OUT, BESTEVER] = ...
%    CMAES(FITFUN, X0, SIGMA)
% returns the best (minimal) point XMIN (found in the last
% generation); function value FMIN of XMIN; the number of needed
% function evaluations COUNTEVAL; a STOPFLAG value as cell array,
% where possible entries are 'fitness', 'tolx', 'tolupx', 'tolfun',
% 'maxfunevals', 'maxiter', 'stoptoresume', 'manual',
% 'warnconditioncov', 'warnnoeffectcoord', 'warnnoeffectaxis',
% 'warnequalfunvals', 'warnequalfunvalhist', 'bug' (use
% e.g. any(strcmp(STOPFLAG, 'tolx')) or findstr(strcat(STOPFLAG,
% 'tolx')) for further processing); a record struct OUT with some
% more output, where the struct SOLUTIONS.BESTEVER contains the overall
% best evaluated point X with function value F evaluated at evaluation
% count EVALS. The last output argument BESTEVER equals 
% OUT.SOLUTIONS.BESTEVER. Moreover a history of solutions and 
% parameters is written to files according to the Log-options. 
%
% A regular manual stop can be achieved via the file signals.par. The
% program is terminated if the first two non-white sequences in any
% line of this file are 'stop' and the value of the LogFilenamePrefix
% option (by default 'outcmaes'). Also a run can be skipped.
% Given, for example, 'skip outcmaes run 2', skips the second run
% if option Restarts is at least 2, and another run will be started.
% 
% To run the code completely silently set Disp, Save, and Log options
% to 0.  With OPTS.LogModulo > 0 (1 by default) the most important
% data are written to ASCII files permitting to investigate the
% results (e.g. plot with function plotcmaesdat) even while CMAES is
% still running (which can be quite useful on expensive objective
% functions). When OPTS.SaveVariables==1 (default) everything is saved
% in file OPTS.SaveFilename (default 'variablescmaes.mat') allowing to
% resume the search afterwards by using the resume option.
%
% To find the best ever evaluated point load the variables typing
% "es=load('variablescmaes')" and investigate the variable
% es.out.solutions.bestever. 
%
% In case of a noisy objective function (uncertainties) set
% OPTS.Noise.on = 1. This option interferes presumably with some 
% termination criteria, because the step-size sigma will presumably
% not converge to zero anymore. If CMAES was provided with a 
% fifth argument (P1 in the below example, which is passed to the 
% objective function FUN), this argument is multiplied with the 
% factor given in option Noise.alphaevals, each time the detected 
% noise exceeds a threshold. This argument can be used within 
% FUN, for example, as averaging number to reduce the noise level.  
%
% OPTS.DiagonalOnly > 1 defines the number of initial iterations,
% where the covariance matrix remains diagonal and the algorithm has
% internally linear time complexity. OPTS.DiagonalOnly = 1 means
% keeping the covariance matrix always diagonal and this setting
% also exhibits linear space complexity. This can be particularly
% useful for dimension > 100. The default is OPTS.DiagonalOnly = 0. 
% 
% OPTS.CMA.active = 1 turns on "active CMA" with a negative update 
% of the covariance matrix and checks for positive definiteness. 
% OPTS.CMA.active = 2 does not check for pos. def. and is numerically
% faster. Active CMA usually speeds up the adaptation and might 
% become a default in near future. 
%
% The primary strategy parameter to play with is OPTS.PopSize, which
% can be increased from its default value.  Increasing the population
% size (by default linked to increasing parent number OPTS.ParentNumber)
% improves global search properties in exchange to speed. Speed
% decreases, as a rule, at most linearely with increasing population
% size. It is advisable to begin with the default small population
% size. The options Restarts and IncPopSize can be used for an
% automated multistart where the population size is increased by the
% factor IncPopSize (two by default) before each restart. X0 (given as
% string) is reevaluated for each restart. Stopping options
% StopFunEvals, StopIter, MaxFunEvals, and Fitness terminate the
% program, all others including MaxIter invoke another restart, where
% the iteration counter is reset to zero.
%
% Examples: 
%
%     XMIN = cmaes('myfun', 5*ones(10,1), 1.5); 
%
%   starts the search at 10D-point 5 and initially searches mainly 
%   between 5-3 and 5+3 (+- two standard deviations), but this is not 
%   a strict bound. 'myfun' is a name of a function that returns a 
%   scalar from a 10D column vector.
%
%     opts.LBounds = 0; opts.UBounds = 10; 
%     opts.Restarts = 3;  % doubles the popsize for each restart
%     X = cmaes('myfun', 10*rand(10,1), 5, opts);
%
%   searches within lower bound of 0 and upper bound of 10. Bounds can
%   also be given as column vectors. If the optimum is not located
%   on the boundary, use rather a penalty approach to handle bounds. 
%
%     opts=cmaes; 
%     opts.StopFitness=1e-10;
%     X=cmaes('myfun', 'rand(5,1)', 0.5, opts); 
%
%   stops the search, if the function value is smaller than 1e-10. The second
%   argument is re-evaluated for each restart.
%   
%     [X, F, E, STOP, OUT] = cmaes('myfun2', 'rand(5,1)', 1, [], P1, P2); 
%
%   passes two additional parameters to the function MYFUN2.
%
% See also FMINSEARCH, FMINUNC, FMINBND.

% ---------------------------------------------------------------
% CMA-ES: Evolution Strategy with Covariance Matrix Adaptation for
% nonlinear function minimization. To be used under the terms of
% either  
%   the GNU General Public License (http://www.gnu.org/copyleft/gpl.html).
% or 
%   the BSD 3-Clause License (see below). 
%
% Author (copyright): Nikolaus Hansen, 2001-2012. 
% e-mail: nikolaus.hansen AT inria.fr
%
% References: See below. 
% ---------------------------------------------------------------
%
% GENERAL PURPOSE: The CMA-ES (Evolution Strategy with Covariance
% Matrix Adaptation) is a robust search method which should be
% applied, if derivative based methods, e.g. quasi-Newton BFGS or
% conjucate gradient, (supposably) fail due to a rugged search
% landscape (e.g. noise, local optima, outlier, etc.). On smooth
% landscapes CMA-ES is roughly ten times slower than BFGS. For up to
% N=10 variables even the simplex direct search method (Nelder & Mead)
% is often faster, but far less robust than CMA-ES.  To see the
% advantage of the CMA, it will usually take at least 30*N and up to
% 300*N function evaluations, where N is the search problem dimension.
% On considerably hard problems the complete search (a single run) is
% expected to take at least 30*N^2 and up to 300*N^2 function
% evaluations.
%
% SOME MORE COMMENTS: 
% The adaptation of the covariance matrix (e.g. by the CMA) is
% equivalent to a general linear transformation of the problem
% coding. Nevertheless every problem specific knowlegde about the best
% linear transformation should be exploited before starting the
% search. That is, an appropriate a priori transformation should be
% applied to the problem. This also makes the identity matrix as
% initial covariance matrix the best choice.
%
% The strategy parameter lambda (population size, opts.PopSize) is the
% preferred strategy parameter to play with.  If results with the
% default strategy are not satisfactory, increase the population
% size. (Remark that the crucial parameter mu (opts.ParentNumber) is
% increased proportionally to lambda). This will improve the
% strategies capability of handling noise and local minima. We
% recomment successively increasing lambda by a factor of about three,
% starting with initial values between 5 and 20. Casually, population
% sizes even beyond 1000+100*N can be sensible.
%

% TODO: 
%       write dispcmaesdat for Matlab (and Octave)
%       control savemodulo and plotmodulo via signals.par 


cmaVersion = '3.62.beta'; 

% ----------- Set Defaults for Input Parameters and Options -------------
% These defaults may be edited for convenience

% Input Defaults (obsolete, these are obligatory now)
definput.fitfun = 'felli'; % frosen; fcigar; see end of file for more
definput.xstart = rand(10,1); % 0.50*ones(10,1);
definput.sigma = 0.3;

% Options defaults: Stopping criteria % (value of stop flag)
defopts.StopFitness  = '-Inf % stop if f(xmin) < stopfitness, minimization';
defopts.MaxFunEvals  = 'Inf  % maximal number of fevals';
defopts.MaxIter      = '1e3*(N+5)^2/sqrt(popsize) % maximal number of iterations';
defopts.StopFunEvals = 'Inf  % stop after resp. evaluation, possibly resume later';
defopts.StopIter     = 'Inf  % stop after resp. iteration, possibly resume later';
defopts.TolX         = '1e-11*max(insigma) % stop if x-change smaller TolX';
defopts.TolUpX       = '1e3*max(insigma) % stop if x-changes larger TolUpX';
defopts.TolFun       = '1e-12 % stop if fun-changes smaller TolFun';
defopts.TolHistFun   = '1e-13 % stop if back fun-changes smaller TolHistFun';
defopts.StopOnStagnation = 'on  % stop when fitness stagnates for a long time';
% TODO: stagnation has four parameters for the period: min = 120, const = 30N/lam, rel = 0.2, max = 2e5
% defopts.StopOnStagnation = '[120 30*N/popsize 0.2 2e5]  % [min const rel_iter max] measuring period';
defopts.StopOnWarnings = 'yes  % ''no''==''off''==0, ''on''==''yes''==1 ';
defopts.StopOnEqualFunctionValues = '2 + N/3  % number of iterations';  

% Options defaults: Other
defopts.DiffMaxChange = 'Inf  % maximal variable change(s), can be Nx1-vector';
defopts.DiffMinChange = '0    % minimal variable change(s), can be Nx1-vector';
defopts.WarnOnEqualFunctionValues = ...
    'yes  % ''no''==''off''==0, ''on''==''yes''==1 ';
defopts.LBounds = -6;%'-Inf % lower bounds, scalar or Nx1-vector'; 
defopts.UBounds = 6;%'Inf  % upper bounds, scalar or Nx1-vector'; 
defopts.EvalParallel = 'no   % objective function FUN accepts NxM matrix, with M>1?';
defopts.EvalInitialX = 'yes  % evaluation of initial solution';
defopts.Restarts     = '0    % number of restarts ';
defopts.IncPopSize   = '2    % multiplier for population size before each restart';

defopts.PopSize      = '(4 + floor(3*log(N)))  % population size, lambda'; 
defopts.ParentNumber = 'floor(popsize/2)       % AKA mu, popsize equals lambda';
defopts.RecombinationWeights = 'superlinear decrease % or linear, or equal';
defopts.DiagonalOnly = '0*(1+100*N/sqrt(popsize))+(N>=1000)  % C is diagonal for given iterations, 1==always'; 
defopts.Noise.on = '0  % uncertainty handling is off by default'; 
defopts.Noise.reevals  = '1*ceil(0.05*lambda)  % nb. of re-evaluated for uncertainty measurement';
defopts.Noise.theta = '0.5  % threshold to invoke uncertainty treatment'; % smaller: more likely to diverge
defopts.Noise.cum = '0.3  % cumulation constant for uncertainty'; 
defopts.Noise.cutoff = '2*lambda/3  % rank change cutoff for summation';
defopts.Noise.alphasigma  = '1+2/(N+10)  % factor for increasing sigma'; % smaller: slower adaptation
defopts.Noise.epsilon  = '1e-7  % additional relative perturbation before reevaluation';
defopts.Noise.minmaxevals = '[1 inf]  % min and max value of 2nd arg to fitfun, start value is 5th arg to cmaes'; 
defopts.Noise.alphaevals  = '1+2/(N+10)  % factor for increasing 2nd arg to fitfun'; 
defopts.Noise.callback = '[]  % callback function when uncertainty threshold is exceeded';
% defopts.TPA = 0; 
defopts.CMA.cs = '(mueff+2)/(N+mueff+3)  % cumulation constant for step-size'; 
   %qqq defopts.CMA.cs = (mueff^0.5)/(N^0.5+mueff^0.5) % the short time horizon version
defopts.CMA.damps = '1 + 2*max(0,sqrt((mueff-1)/(N+1))-1) + cs  % damping for step-size';
% defopts.CMA.ccum = '4/(N+4)  % cumulation constant for covariance matrix'; 
defopts.CMA.ccum = '(4 + mueff/N) / (N+4 + 2*mueff/N)  % cumulation constant for pc';
defopts.CMA.ccov1 = '2 / ((N+1.3)^2+mueff)  % learning rate for rank-one update'; 
defopts.CMA.ccovmu = '2 * (mueff-2+1/mueff) / ((N+2)^2+mueff) % learning rate for rank-mu update'; 
defopts.CMA.on     = 'yes'; 
defopts.CMA.active = '0  % active CMA, 1: neg. updates with pos. def. check, 2: neg. updates'; 

flg_future_setting = 0;  % testing for possible future variant(s)
if flg_future_setting    
  disp('in the future')

  % damps setting from Brockhoff et al 2010
  %   this damps diverges with popsize 400:
  %     defopts.CMA.damps = '2*mueff/lambda + 0.3 + cs  % damping for step-size'; 
  %   cmaeshtml('benchmarkszero', ones(20,1)*2, 5, o, 15);
  % how about:
  % defopts.CMA.damps = '2*mueff/lambda + 0.3 + 2*max(0,sqrt((mueff-1)/(N+1))-1) + cs  % damping for step-size';
  defopts.CMA.damps = '0.5 + 0.5*min(1, (0.27*lambda/mueff-1)^2) + 2*max(0,sqrt((mueff-1)/(N+1))-1) + cs  % damping for step-size';

  if 11 < 3 
      defopts.CMA.damps = '0.5 + 0.5*min(1,(lam_mirr/(0.159*lambda)-1)^2) + 2*max(0,sqrt((mueff-1)/(N+1))-1) + cs  % damping for step-size';

      defopts.mirrored_offspring = 'floor(0.5 + 0.159 * lambda)'; 
    % TODO: this should also depend on diagonal option!?  
    defopts.CMA.active = 'floor(int8(lam_mirr>0))  % active CMA 1: neg. updates with pos. def. check, 2: neg. updates'; 
  end
  
  % ccum adjusted for large mueff, better on schefelmult? 
  % TODO: this should also depend on diagonal option!?  
  defopts.CMA.ccum = '(4 + mueff/N) / (N+4 + 2*mueff/N)  % cumulation constant for pc';

  defopts.CMA.active = '1  % active CMA 1: neg. updates with pos. def. check, 2: neg. updates'; 
end
  
defopts.Resume   = 'no   % resume former run from SaveFile'; 
defopts.Science  = 'on  % off==do some additional (minor) problem capturing, NOT IN USE'; 
defopts.ReadSignals = 'on  % from file signals.par for termination, yet a stumb';
defopts.Seed = 'sum(100*clock)  % evaluated if it is a string';
defopts.DispFinal  = 'on   % display messages like initial and final message';
defopts.DispModulo = '100  % [0:Inf], disp messages after every i-th iteration';
defopts.SaveVariables = 'on   % [on|final|off][-v6] save variables to .mat file';
defopts.SaveFilename = 'variablescmaes.mat  % save all variables, see SaveVariables'; 
defopts.LogModulo = '1    % [0:Inf] if >1 record data less frequently after gen=100';
defopts.LogTime   = '25   % [0:100] max. percentage of time for recording data';
defopts.LogFilenamePrefix = 'outcmaes  % files for output data'; 
defopts.LogPlot = 'off    % plot while running using output data files';

%qqqkkk 
%defopts.varopt1 = ''; % 'for temporary and hacking purposes'; 
%defopts.varopt2 = ''; % 'for temporary and hacking purposes'; 
defopts.UserData = 'for saving data/comments associated with the run';
defopts.UserDat2 = ''; 'for saving data/comments associated with the run';

% ---------------------- Handling Input Parameters ----------------------

if nargin < 1 || isequal(fitfun, 'defaults') % pass default options
  if nargin < 1
    disp('Default options returned (type "help cmaes" for help).');
  end
  xmin = defopts;
  if nargin > 1 % supplement second argument with default options
    xmin = getoptions(xstart, defopts);
  end
  return;
end

if isequal(fitfun, 'displayoptions')
 names = fieldnames(defopts); 
 for name = names'
   disp([name{:} repmat(' ', 1, 20-length(name{:})) ': ''' defopts.(name{:}) '''']); 
 end
 return; 
end

input.fitfun = fitfun; % record used input
if isempty(fitfun)
  % fitfun = definput.fitfun; 
  % warning(['Objective function not determined, ''' fitfun ''' used']);
  error(['Objective function not determined']);
end
% if ~ischar(fitfun)
%   error('first argument FUN must be a string');
% end


if nargin < 2 
  xstart = [];
end

input.xstart = xstart;
if isempty(xstart)
  % xstart = definput.xstart;  % objective variables initial point
  % warning('Initial search point, and problem dimension, not determined');
  error('Initial search point, and problem dimension, not determined');
end

if nargin < 3 
  insigma = [];
end
if isa(insigma, 'struct')
  error(['Third argument SIGMA must be (or eval to) a scalar '...
	   'or a column vector of size(X0,1)']);
end
input.sigma = insigma;
if isempty(insigma)
  if all(size(myeval(xstart)) > 1)
    insigma = std(xstart, 0, 2); 
    if any(insigma == 0)
      error(['Initial search volume is zero, choose SIGMA or X0 appropriate']);
    end
  else
    % will be captured later
    % error(['Initial step sizes (SIGMA) not determined']);
  end
end

% Compose options opts
if nargin < 4 || isempty(inopts) % no input options available
  inopts = []; 
  opts = defopts;
else
  opts = getoptions(inopts, defopts);
end
i = strfind(opts.SaveFilename, '%'); % remove everything after comment
if ~isempty(i)
  opts.SaveFilename = opts.SaveFilename(1:i(1)-1);
end
opts.SaveFilename = deblank(opts.SaveFilename); % remove trailing white spaces

%%%%%%%%%%%%%%%%%%%%%%%%%%%%%%%%%%%%%%  
counteval = 0; countevalNaN = 0; 
irun = 0;
while irun <= myeval(opts.Restarts) % for-loop does not work with resume
  irun = irun + 1; 

% ------------------------ Initialization -------------------------------

% Handle resuming of old run
flgresume = myevalbool(opts.Resume);
xmean = myeval(xstart); 
if all(size(xmean) > 1)
   xmean = mean(xmean, 2); % in case if xstart is a population
elseif size(xmean, 2) > 1
  xmean = xmean';
end 
if ~flgresume % not resuming a former run
  % Assign settings from input parameters and options for myeval...
  N = size(xmean, 1); numberofvariables = N; 
  lambda0 = floor(myeval(opts.PopSize) * myeval(opts.IncPopSize)^(irun-1)); 
  % lambda0 = floor(myeval(opts.PopSize) * 3^floor((irun-1)/2)); 
  popsize = lambda0;
  lambda = lambda0;
  insigma = myeval(insigma);
  if all(size(insigma) == [N 2]) 
    insigma = 0.5 * (insigma(:,2) - insigma(:,1));
  end
else % flgresume is true, do resume former run
  tmp = whos('-file', opts.SaveFilename);
  for i = 1:length(tmp)
    if strcmp(tmp(i).name, 'localopts');
      error('Saved variables include variable "localopts", please remove');
    end
  end
  local.opts = opts; % keep stopping and display options
  local.varargin = varargin;
  load(opts.SaveFilename); 
  varargin = local.varargin;
  flgresume = 1;

  % Overwrite old stopping and display options
  opts.StopFitness = local.opts.StopFitness; 
  %%opts.MaxFunEvals = local.opts.MaxFunEvals;
  %%opts.MaxIter = local.opts.MaxIter; 
  opts.StopFunEvals = local.opts.StopFunEvals; 
  opts.StopIter = local.opts.StopIter;  
  opts.TolX = local.opts.TolX;
  opts.TolUpX = local.opts.TolUpX;
  opts.TolFun = local.opts.TolFun;
  opts.TolHistFun = local.opts.TolHistFun;
  opts.StopOnStagnation = local.opts.StopOnStagnation; 
  opts.StopOnWarnings = local.opts.StopOnWarnings; 
  opts.ReadSignals = local.opts.ReadSignals; 
  opts.DispFinal = local.opts.DispFinal;
  opts.LogPlot = local.opts.LogPlot;
  opts.DispModulo = local.opts.DispModulo;
  opts.SaveVariables = local.opts.SaveVariables;
  opts.LogModulo = local.opts.LogModulo;
  opts.LogTime = local.opts.LogTime;
  clear local; % otherwise local would be overwritten during load
end
  
%--------------------------------------------------------------
% Evaluate options
stopFitness = myeval(opts.StopFitness); 
stopMaxFunEvals = myeval(opts.MaxFunEvals);  
stopMaxIter = myeval(opts.MaxIter);  
stopFunEvals = myeval(opts.StopFunEvals);  
stopIter = myeval(opts.StopIter); 
if flgresume
    stopIter = stopIter + countiter
end
stopTolX = myeval(opts.TolX);
stopTolUpX = myeval(opts.TolUpX);
stopTolFun = myeval(opts.TolFun);
stopTolHistFun = myeval(opts.TolHistFun);
stopOnStagnation = myevalbool(opts.StopOnStagnation); 
stopOnWarnings = myevalbool(opts.StopOnWarnings);
flgreadsignals = myevalbool(opts.ReadSignals);
flgWarnOnEqualFunctionValues = myevalbool(opts.WarnOnEqualFunctionValues);
flgEvalParallel = myevalbool(opts.EvalParallel);
stopOnEqualFunctionValues = myeval(opts.StopOnEqualFunctionValues);
arrEqualFunvals = zeros(1, 10+N);
flgDiagonalOnly = myeval(opts.DiagonalOnly); 
flgActiveCMA = myeval(opts.CMA.active); 
noiseHandling = myevalbool(opts.Noise.on);
noiseMinMaxEvals = myeval(opts.Noise.minmaxevals);
noiseAlphaEvals = myeval(opts.Noise.alphaevals);
noiseCallback = myeval(opts.Noise.callback); 
flgdisplay = myevalbool(opts.DispFinal);
flgplotting = myevalbool(opts.LogPlot);
verbosemodulo = myeval(opts.DispModulo);
flgscience = myevalbool(opts.Science);
flgsaving = [];
strsaving = [];
if strfind(opts.SaveVariables, '-v6') 
  i = strfind(opts.SaveVariables, '%');
  if isempty(i) || i == 0 || strfind(opts.SaveVariables, '-v6') < i
    strsaving = '-v6';
    flgsaving = 1;
    flgsavingfinal = 1;
  end
end
if strncmp('final', opts.SaveVariables, 5)
  flgsaving = 0;
  flgsavingfinal = 1;
end
if isempty(flgsaving)
  flgsaving = myevalbool(opts.SaveVariables);
  flgsavingfinal = flgsaving;
end
savemodulo = myeval(opts.LogModulo);
savetime = myeval(opts.LogTime);

i = strfind(opts.LogFilenamePrefix, ' '); % remove everything after white space
if ~isempty(i)
  opts.LogFilenamePrefix = opts.LogFilenamePrefix(1:i(1)-1);
end

% TODO here silent option? set disp, save and log options to 0 

%--------------------------------------------------------------

if (isfinite(stopFunEvals) || isfinite(stopIter)) && ~flgsaving
  warning('To resume later the saving option needs to be set');
end


% Do more checking and initialization 
if flgresume % resume is on
  time.t0 = clock;
  if flgdisplay
    disp(['  resumed from ' opts.SaveFilename ]); 
  end
  if counteval >= stopMaxFunEvals 
    error(['MaxFunEvals exceeded, use StopFunEvals as stopping ' ...
	  'criterion before resume']);
  end
  if countiter >= stopMaxIter 
    error(['MaxIter exceeded, use StopIter as stopping criterion ' ...
	  'before resume']);
  end
  
else % flgresume
  % xmean = mean(myeval(xstart), 2); % evaluate xstart again, because of irun
  maxdx = myeval(opts.DiffMaxChange); % maximal sensible variable change
  mindx = myeval(opts.DiffMinChange); % minimal sensible variable change 
				      % can both also be defined as Nx1 vectors
  lbounds = myeval(opts.LBounds);		     
  ubounds = myeval(opts.UBounds);
  if length(lbounds) == 1
    lbounds = repmat(lbounds, N, 1);
  end
  if length(ubounds) == 1
    ubounds = repmat(ubounds, N, 1);
  end
  if isempty(insigma) % last chance to set insigma
    if all(lbounds > -Inf) && all(ubounds < Inf)
      if any(lbounds>=ubounds)
	error('upper bound must be greater than lower bound');
      end
      insigma = 0.3*(ubounds-lbounds);
      stopTolX = myeval(opts.TolX);  % reevaluate these
      stopTolUpX = myeval(opts.TolUpX);
    else
      error(['Initial step sizes (SIGMA) not determined']);
    end
  end

  % Check all vector sizes
  if size(xmean, 2) > 1 || size(xmean,1) ~= N
    error(['intial search point should be a column vector of size ' ...
	   num2str(N)]);
  elseif ~(all(size(insigma) == [1 1]) || all(size(insigma) == [N 1]))
    error(['input parameter SIGMA should be (or eval to) a scalar '...
	   'or a column vector of size ' num2str(N)] );
  elseif size(stopTolX, 2) > 1 || ~ismember(size(stopTolX, 1), [1 N])
    error(['option TolX should be (or eval to) a scalar '...
	   'or a column vector of size ' num2str(N)] );
  elseif size(stopTolUpX, 2) > 1 || ~ismember(size(stopTolUpX, 1), [1 N])
    error(['option TolUpX should be (or eval to) a scalar '...
	   'or a column vector of size ' num2str(N)] );
  elseif size(maxdx, 2) > 1 || ~ismember(size(maxdx, 1), [1 N])
    error(['option DiffMaxChange should be (or eval to) a scalar '...
	   'or a column vector of size ' num2str(N)] );
  elseif size(mindx, 2) > 1 || ~ismember(size(mindx, 1), [1 N])
    error(['option DiffMinChange should be (or eval to) a scalar '...
	   'or a column vector of size ' num2str(N)] );
  elseif size(lbounds, 2) > 1 || ~ismember(size(lbounds, 1), [1 N])
    error(['option lbounds should be (or eval to) a scalar '...
	   'or a column vector of size ' num2str(N)] );
  elseif size(ubounds, 2) > 1 || ~ismember(size(ubounds, 1), [1 N])
    error(['option ubounds should be (or eval to) a scalar '...
	   'or a column vector of size ' num2str(N)] );
  end
  
  % Initialize dynamic internal state parameters
  if any(insigma <= 0) 
    error(['Initial search volume (SIGMA) must be greater than zero']);
  end
  if max(insigma)/min(insigma) > 1e6
    error(['Initial search volume (SIGMA) badly conditioned']);
  end
  sigma = max(insigma);              % overall standard deviation
  pc = zeros(N,1); ps = zeros(N,1);  % evolution paths for C and sigma

  if length(insigma) == 1
    insigma = insigma * ones(N,1) ;
  end
  diagD = insigma/max(insigma);      % diagonal matrix D defines the scaling
  diagC = diagD.^2; 
  if flgDiagonalOnly ~= 1            % use at some point full covariance matrix
    B = eye(N,N);                      % B defines the coordinate system
    BD = B.*repmat(diagD',N,1);        % B*D for speed up only
    C = diag(diagC);                   % covariance matrix == BD*(BD)'
  end
  if flgDiagonalOnly
    B = 1; 
  end

  fitness.hist=NaN*ones(1,10+ceil(3*10*N/lambda)); % history of fitness values
  fitness.histsel=NaN*ones(1,10+ceil(3*10*N/lambda)); % history of fitness values
  fitness.histbest=[]; % history of fitness values
  fitness.histmedian=[]; % history of fitness values

  % Initialize boundary handling
  bnd.isactive = any(lbounds > -Inf) || any(ubounds < Inf); 
  if bnd.isactive
    if any(lbounds>ubounds)
      error('lower bound found to be greater than upper bound');
    end
    [xmean ti] = xintobounds(xmean, lbounds, ubounds); % just in case
    if any(ti)
      warning('Initial point was out of bounds, corrected');
    end
    bnd.weights = zeros(N,1);         % weights for bound penalty
    % scaling is better in axis-parallel case, worse in rotated
    bnd.flgscale = 0; % scaling will be omitted if zero 
    if bnd.flgscale ~= 0 
      bnd.scale = diagC/mean(diagC);
    else
      bnd.scale = ones(N,1);
    end
    
    idx = (lbounds > -Inf) | (ubounds < Inf);
    if length(idx) == 1
      idx = idx * ones(N,1);
    end
    bnd.isbounded = zeros(N,1);
    bnd.isbounded(find(idx)) = 1; 
    maxdx = min(maxdx, (ubounds - lbounds)/2);
    if any(sigma*sqrt(diagC) > maxdx)
      fac = min(maxdx ./ sqrt(diagC))/sigma;
      sigma = min(maxdx ./ sqrt(diagC));
      warning(['Initial SIGMA multiplied by the factor ' num2str(fac) ...
	       ', because it was larger than half' ...
	       ' of one of the boundary intervals']);
    end
    idx = (lbounds > -Inf) & (ubounds < Inf);
    dd = diagC;
    if any(5*sigma*sqrt(dd(idx)) < ubounds(idx) - lbounds(idx))
      warning(['Initial SIGMA is, in at least one coordinate, ' ...
	       'much smaller than the '...
	       'given boundary intervals. For reasonable ' ...
	       'global search performance SIGMA should be ' ...
	       'between 0.2 and 0.5 of the bounded interval in ' ...
	       'each coordinate. If all coordinates have ' ... 
	       'lower and upper bounds SIGMA can be empty']);
    end
    bnd.dfithist = 1;              % delta fit for setting weights
    bnd.aridxpoints = [];          % remember complete outside points
    bnd.arfitness = [];            % and their fitness
    bnd.validfitval = 0;
    bnd.iniphase = 1;
  end

  % ooo initial feval, for output only
  if irun == 1 
    out.solutions.bestever.x = xmean;
    out.solutions.bestever.f = Inf;  % for simpler comparison below
    out.solutions.bestever.evals = counteval;
    bestever = out.solutions.bestever;
  end
  if myevalbool(opts.EvalInitialX)
    fitness.hist(1)=feval(fitfun, xmean, varargin{:}); 
    fitness.histsel(1)=fitness.hist(1);
    counteval = counteval + 1;
    if fitness.hist(1) < out.solutions.bestever.f 
	out.solutions.bestever.x = xmean;
	out.solutions.bestever.f = fitness.hist(1);
	out.solutions.bestever.evals = counteval;
	bestever = out.solutions.bestever;
    end
  else
    fitness.hist(1)=NaN; 
    fitness.histsel(1)=NaN; 
  end
    
  % initialize random number generator
  if ischar(opts.Seed)
    randn('state', eval(opts.Seed));     % random number generator state
  else
    randn('state', opts.Seed);
  end
  %qqq
%  load(opts.SaveFilename, 'startseed');
%  randn('state', startseed);
%  disp(['SEED RELOADED FROM ' opts.SaveFilename]);
  startseed = randn('state');         % for retrieving in saved variables

  % Initialize further constants
  chiN=N^0.5*(1-1/(4*N)+1/(21*N^2));  % expectation of 
				      %   ||N(0,I)|| == norm(randn(N,1))
  
  countiter = 0;
  % Initialize records and output
  if irun == 1
    time.t0 = clock;
    
    % TODO: keep also median solution? 
    out.evals = counteval;  % should be first entry
    out.stopflag = {};

    outiter = 0;

    % Write headers to output data files 
    filenameprefix = opts.LogFilenamePrefix; 
    if savemodulo && savetime
      filenames = {};
      filenames(end+1) = {'axlen'};
      filenames(end+1) = {'fit'};
      filenames(end+1) = {'stddev'};
      filenames(end+1) = {'xmean'};
      filenames(end+1) = {'xrecentbest'};
      str = [' (startseed=' num2str(startseed(2)) ...
             ', ' num2str(clock, '%d/%02d/%d %d:%d:%2.2f') ')'];
      for namecell = filenames(:)'
        name = namecell{:};

	[fid, err] = fopen(['./' filenameprefix name '.dat'], 'w');
	if fid < 1 % err ~= 0 
	  warning(['could not open ' filenameprefix name '.dat']);
	  filenames(find(strcmp(filenames,name))) = [];
	else
%	  fprintf(fid, '%s\n', ...
%	      ['<CMAES-OUTPUT version="' cmaVersion '">']);
%	  fprintf(fid, ['  <NAME>' name '</NAME>\n']);
%	  fprintf(fid, ['  <DATE>' date() '</DATE>\n']);
%	  fprintf(fid, '  <PARAMETERS>\n');
%	  fprintf(fid, ['    dimension=' num2str(N) '\n']);
%	  fprintf(fid, '  </PARAMETERS>\n');
	  % different cases for DATA columns annotations here
%	  fprintf(fid, '  <DATA');
	  if strcmp(name, 'axlen')
	     fprintf(fid, ['%%  columns="iteration, evaluation, sigma, ' ...
		 'max axis length, min axis length, ' ...
		 'all principal axes lengths (sorted square roots ' ...
                  'of eigenvalues of C)"' str]);
	  elseif strcmp(name, 'fit')
	    fprintf(fid, ['%%  columns="iteration, evaluation, sigma, axis ratio, bestever,' ...
		' best, median, worst fitness function value,' ...
		' further objective values of best"' str]);
	  elseif strcmp(name, 'stddev')
	    fprintf(fid, ['%%  columns=["iteration, evaluation, sigma, void, void, ' ...
		'stds==sigma*sqrt(diag(C))"' str]);
	  elseif strcmp(name, 'xmean')
	    fprintf(fid, ['%%  columns="iteration, evaluation, void, ' ...
                          'void, void, xmean"' str]);
	  elseif strcmp(name, 'xrecentbest')
	    fprintf(fid, ['%%  columns="iteration, evaluation, fitness, ' ...
                          'void, void, xrecentbest"' str]);
	  end
	  fprintf(fid, '\n'); % DATA
	  if strcmp(name, 'xmean')
	    fprintf(fid, '%ld %ld 0 0 0 ', 0, counteval); 
	    % fprintf(fid, '%ld %ld 0 0 %e ', countiter, counteval, fmean); 
%qqq	    fprintf(fid, msprintf('%e ', genophenotransform(out.genopheno, xmean)) + '\n'); 
	    fprintf(fid, '%e ', xmean);
            fprintf(fid, '\n'); 
	  end
	  fclose(fid); 
          clear fid; % preventing 
	end
      end % for files
    end % savemodulo
  end % irun == 1
  
end % else flgresume 

% -------------------- Generation Loop --------------------------------
stopflag = {};
while isempty(stopflag)
  % set internal parameters
  if countiter == 0 || lambda ~= lambda_last
    if countiter > 0 && floor(log10(lambda)) ~= floor(log10(lambda_last)) ...
          && flgdisplay
      disp(['  lambda = ' num2str(lambda)]);
      lambda_hist(:,end+1) = [countiter+1; lambda];
    else
      lambda_hist = [countiter+1; lambda]; 
    end
    lambda_last = lambda;
    % Strategy internal parameter setting: Selection  
    mu = myeval(opts.ParentNumber); % number of parents/points for recombination
    if strncmp(lower(opts.RecombinationWeights), 'equal', 3)
      weights = ones(mu,1); % (mu_I,lambda)-CMA-ES
    elseif strncmp(lower(opts.RecombinationWeights), 'linear', 3)
      weights = mu+0.5-(1:mu)'; 
    elseif strncmp(lower(opts.RecombinationWeights), 'superlinear', 3)
      % use (lambda+1)/2 as reference if mu < lambda/2
      weights = log(max(mu, lambda/2) + 1/2)-log(1:mu)'; % muXone array for weighted recombination
    else
      error(['Recombination weights to be "' opts.RecombinationWeights ...
             '" is not implemented']);
    end
    mueff=sum(weights)^2/sum(weights.^2); % variance-effective size of mu
    weights = weights/sum(weights);     % normalize recombination weights array
    if mueff == lambda
      error(['Combination of values for PopSize, ParentNumber and ' ...
             ' and RecombinationWeights is not reasonable']);
    end
    
    % Strategy internal parameter setting: Adaptation
    cc = myeval(opts.CMA.ccum); % time constant for cumulation for covariance matrix
    cs = myeval(opts.CMA.cs); 

    % old way TODO: remove this at some point
    % mucov = mueff;   % size of mu used for calculating learning rate ccov
    % ccov = (1/mucov) * 2/(N+1.41)^2 ... % learning rate for covariance matrix
    %        + (1-1/mucov) * min(1,(2*mucov-1)/((N+2)^2+mucov)); 

    % new way
    if myevalbool(opts.CMA.on) 
      ccov1 = myeval(opts.CMA.ccov1); 
      ccovmu = min(1-ccov1, myeval(opts.CMA.ccovmu));
    else
      ccov1 = 0;
      ccovmu = 0;
    end
    
    % flgDiagonalOnly = -lambda*4*1/ccov; % for ccov==1 it is not needed
    % 0 : C will never be diagonal anymore
    % 1 : C will always be diagonal
    % >1: C is diagonal for first iterations, set to 0 afterwards
    if flgDiagonalOnly < 1
      flgDiagonalOnly = 0; 
    end
    if flgDiagonalOnly
      ccov1_sep = min(1, ccov1 * (N+1.5) / 3); 
      ccovmu_sep = min(1-ccov1_sep, ccovmu * (N+1.5) / 3); 
    elseif N > 98 && flgdisplay && countiter == 0
      disp('consider option DiagonalOnly for high-dimensional problems');
    end

    % ||ps|| is close to sqrt(mueff/N) for mueff large on linear fitness
    %damps = ... % damping for step size control, usually close to one 
    %    (1 + 2*max(0,sqrt((mueff-1)/(N+1))-1)) ... % limit sigma increase
    %    * max(0.3, ... % reduce damps, if max. iteration number is small
    %          1 - N/min(stopMaxIter,stopMaxFunEvals/lambda)) + cs; 
    damps = myeval(opts.CMA.damps); 
    if noiseHandling
      noiseReevals = min(myeval(opts.Noise.reevals), lambda); 
      noiseAlpha = myeval(opts.Noise.alphasigma); 
      noiseEpsilon = myeval(opts.Noise.epsilon); 
      noiseTheta = myeval(opts.Noise.theta); 
      noisecum = myeval(opts.Noise.cum);
      noiseCutOff = myeval(opts.Noise.cutoff);  % arguably of minor relevance
    else
      noiseReevals = 0; % more convenient in later coding
    end

    %qqq hacking of a different parameter setting, e.g. for ccov or damps,
    %  can be done here, but is not necessary anymore, see opts.CMA. 
    % ccov1 = 0.0*ccov1; disp(['CAVE: ccov1=' num2str(ccov1)]);
    % ccovmu = 0.0*ccovmu; disp(['CAVE: ccovmu=' num2str(ccovmu)]);
    % damps = inf*damps; disp(['CAVE: damps=' num2str(damps)]);
    % cc = 1; disp(['CAVE: cc=' num2str(cc)]);

  end    

  % Display initial message
  if countiter == 0 && flgdisplay 
    if mu == 1
      strw = '100';
    elseif mu < 8
      strw = [sprintf('%.0f', 100*weights(1)) ... 
              sprintf(' %.0f', 100*weights(2:end)')];
    else
      strw = [sprintf('%.2g ', 100*weights(1:2)') ...
              sprintf('%.2g', 100*weights(3)') '...' ...
              sprintf(' %.2g', 100*weights(end-1:end)') ']%, '];      
    end
    if irun > 1
      strrun = [', run ' num2str(irun)];
    else
      strrun = '';
    end
    disp(['  n=' num2str(N) ': (' num2str(mu) ',' ...
          num2str(lambda) ')-CMA-ES(w=[' ...
          strw ']%, ' ...
          'mu_eff=' num2str(mueff,'%.1f') ...
          ')']);
    if flgDiagonalOnly == 1
      disp('    C is diagonal');
    elseif flgDiagonalOnly
      disp(['    C is diagonal for ' num2str(floor(flgDiagonalOnly)) ' iterations']);
    end
  end

  flush;

  countiter = countiter + 1; 

  % Generate and evaluate lambda offspring
 
  fitness.raw = repmat(NaN, 1, lambda + noiseReevals);

  % parallel evaluation
  if flgEvalParallel
      arz = randn(N,lambda);

      if ~flgDiagonalOnly
        arx = repmat(xmean, 1, lambda) + sigma * (BD * arz); % Eq. (1)
      else
        arx = repmat(xmean, 1, lambda) + repmat(sigma * diagD, 1, lambda) .* arz; 
      end

      if noiseHandling 
        if noiseEpsilon == 0
          arx = [arx arx(:,1:noiseReevals)]; 
        elseif flgDiagonalOnly
          arx = [arx arx(:,1:noiseReevals) + ...
                 repmat(noiseEpsilon * sigma * diagD, 1, noiseReevals) ...
                 .* randn(N,noiseReevals)]; 
        else 
          arx = [arx arx(:,1:noiseReevals) + ...
                 noiseEpsilon * sigma * ...
                 (BD * randn(N,noiseReevals))]; 
        end
      end

      % You may handle constraints here. You may either resample
      % arz(:,k) and/or multiply it with a factor between -1 and 1
      % (the latter will decrease the overall step size) and
      % recalculate arx accordingly. Do not change arx or arz in any
      % other way.
 
      if ~bnd.isactive
        arxvalid = arx;
      else
        arxvalid = xintobounds(arx, lbounds, ubounds);
      end
      % You may handle constraints here.  You may copy and alter
      % (columns of) arxvalid(:,k) only for the evaluation of the
      % fitness function. arx and arxvalid should not be changed.
      fitness.raw = feval(fitfun, arxvalid, varargin{:}); 
      countevalNaN = countevalNaN + sum(isnan(fitness.raw));
      counteval = counteval + sum(~isnan(fitness.raw)); 
  end

  % non-parallel evaluation and remaining NaN-values
  % set also the reevaluated solution to NaN
  fitness.raw(lambda + find(isnan(fitness.raw(1:noiseReevals)))) = NaN;  
  for k=find(isnan(fitness.raw)), 
    % fitness.raw(k) = NaN; 
    tries = flgEvalParallel;  % in parallel case this is the first re-trial
    % Resample, until fitness is not NaN
    while isnan(fitness.raw(k))
      if k <= lambda  % regular samples (not the re-evaluation-samples)
        arz(:,k) = randn(N,1); % (re)sample

        if flgDiagonalOnly  
          arx(:,k) = xmean + sigma * diagD .* arz(:,k);              % Eq. (1)
        else
          arx(:,k) = xmean + sigma * (BD * arz(:,k));                % Eq. (1)
        end
      else % re-evaluation solution with index > lambda
        if flgDiagonalOnly  
          arx(:,k) = arx(:,k-lambda) + (noiseEpsilon * sigma) * diagD .* randn(N,1);
        else
          arx(:,k) = arx(:,k-lambda) + (noiseEpsilon * sigma) * (BD * randn(N,1));
        end
      end
      
      % You may handle constraints here. You may either resample
      % arz(:,k) and/or multiply it with a factor between -1 and 1
      % (the latter will decrease the overall step size) and
      % recalculate arx accordingly. Do not change arx or arz in any
      % other way.
 
      if ~bnd.isactive
        arxvalid(:,k) = arx(:,k);
      else
        arxvalid(:,k) = xintobounds(arx(:,k), lbounds, ubounds);
      end
      % You may handle constraints here.  You may copy and alter
      % (columns of) arxvalid(:,k) only for the evaluation of the
      % fitness function. arx should not be changed.
      fitness.raw(k) = feval(fitfun, arxvalid(:,k), varargin{:}); 
      tries = tries + 1;
      if isnan(fitness.raw(k))
	countevalNaN = countevalNaN + 1;
      end
      if mod(tries, 100) == 0
	warning([num2str(tries) ...
                 ' NaN objective function values at evaluation ' ...
                 num2str(counteval)]);
      end
    end
    counteval = counteval + 1; % retries due to NaN are not counted
  end

  fitness.sel = fitness.raw; 

  % ----- handle boundaries -----
  if 1 < 3 && bnd.isactive
    % Get delta fitness values
    val = myprctile(fitness.raw, [25 75]);
    % more precise would be exp(mean(log(diagC)))
    val = (val(2) - val(1)) / N / mean(diagC) / sigma^2;
    %val = (myprctile(fitness.raw, 75) - myprctile(fitness.raw, 25)) ...
    %    / N / mean(diagC) / sigma^2;
    % Catch non-sensible values 
    if ~isfinite(val)
      warning('Non-finite fitness range');
      val = max(bnd.dfithist);  
    elseif val == 0 % happens if all points are out of bounds
      val = min(bnd.dfithist(bnd.dfithist>0));  % seems not to make sense, given all solutions are out of bounds
    elseif bnd.validfitval == 0 % flag that first sensible val was found
      bnd.dfithist = [];
      bnd.validfitval = 1;
    end

    % Store delta fitness values
    if length(bnd.dfithist) < 20+(3*N)/lambda
      bnd.dfithist = [bnd.dfithist val];
    else
      bnd.dfithist = [bnd.dfithist(2:end) val];
    end

    [tx ti]  = xintobounds(xmean, lbounds, ubounds);

    % Set initial weights
    if bnd.iniphase 
      if any(ti) 
        bnd.weights(find(bnd.isbounded)) = 2.0002 * median(bnd.dfithist);
	if bnd.flgscale == 0 % scale only initial weights then
	  dd = diagC; 
	  idx = find(bnd.isbounded); 
	  dd = dd(idx) / mean(dd); %  remove mean scaling
	  bnd.weights(idx) = bnd.weights(idx) ./ dd; 
	end
	if bnd.validfitval && countiter > 2
          bnd.iniphase = 0;
	end
      end
    end

    % Increase weights
    if  1 < 3 && any(ti) % any coordinate of xmean out of bounds
      % judge distance of xmean to boundary
      tx = xmean - tx;
      idx = (ti ~= 0 & abs(tx) > 3*max(1,sqrt(N)/mueff) ... 
	     * sigma*sqrt(diagC)) ;
      % only increase if xmean is moving away
      idx = idx & (sign(tx) == sign(xmean - xold));
      if ~isempty(idx) % increase
        % the factor became 1.2 instead of 1.1, because
        % changed from max to min in version 3.52
        bnd.weights(idx) = 1.2^(min(1, mueff/10/N)) * bnd.weights(idx); 
      end
    end

    % Calculate scaling biased to unity, product is one
    if bnd.flgscale ~= 0 
      bnd.scale = exp(0.9*(log(diagC)-mean(log(diagC)))); 
    end

    % Assigned penalized fitness
    bnd.arpenalty = (bnd.weights ./ bnd.scale)' * (arxvalid - arx).^2; 

    fitness.sel = fitness.raw + bnd.arpenalty;

  end % handle boundaries
  % ----- end handle boundaries -----
  
  % compute noise measurement and reduce fitness arrays to size lambda
  if noiseHandling 
    [noiseS] = local_noisemeasurement(fitness.sel(1:lambda), ...
                                      fitness.sel(lambda+(1:noiseReevals)), ...
                                      noiseReevals, noiseTheta, noiseCutOff); 
    if countiter == 1 % TODO: improve this very rude way of initialization
      noiseSS = 0;
      noiseN = 0;  % counter for mean
    end
    noiseSS = noiseSS + noisecum * (noiseS - noiseSS); 

    % noise-handling could be done here, but the original sigma is still needed
    % disp([noiseS noiseSS noisecum])

    fitness.rawar12 = fitness.raw; % just documentary
    fitness.selar12 = fitness.sel; % just documentary
    % qqq refine fitness based on both values
    if 11 < 3  % TODO: in case of outliers this mean is counterproductive 
               % median out of three would be ok 
      fitness.raw(1:noiseReevals) = ... % not so raw anymore
          (fitness.raw(1:noiseReevals) + fitness.raw(lambda+(1:noiseReevals))) / 2; 
      fitness.sel(1:noiseReevals) = ... 
          (fitness.sel(1:noiseReevals) + fitness.sel(lambda+(1:noiseReevals))) / 2; 
    end      
    fitness.raw = fitness.raw(1:lambda); 
    fitness.sel = fitness.sel(1:lambda); 
  end
  
  % Sort by fitness 
  [fitness.raw, fitness.idx] = sort(fitness.raw); 
  [fitness.sel, fitness.idxsel] = sort(fitness.sel);  % minimization
  fitness.hist(2:end) = fitness.hist(1:end-1);    % record short history of
  fitness.hist(1) = fitness.raw(1);               % best fitness values
  if length(fitness.histbest) < 120+ceil(30*N/lambda) || ...
       (mod(countiter, 5) == 0  && length(fitness.histbest) < 2e4)  % 20 percent of 1e5 gen.
    fitness.histbest = [fitness.raw(1) fitness.histbest];          % best fitness values
    fitness.histmedian = [median(fitness.raw) fitness.histmedian]; % median fitness values
  else
    fitness.histbest(2:end) = fitness.histbest(1:end-1); 
    fitness.histmedian(2:end) = fitness.histmedian(1:end-1); 
    fitness.histbest(1) = fitness.raw(1);           % best fitness values
    fitness.histmedian(1) = median(fitness.raw);    % median fitness values
  end
  fitness.histsel(2:end) = fitness.histsel(1:end-1); % record short history of
  fitness.histsel(1) = fitness.sel(1);               % best sel fitness values

  % Calculate new xmean, this is selection and recombination 
  xold = xmean; % for speed up of Eq. (2) and (3)
  cmean = 1;  % 1/min(max((lambda-1*N)/2, 1), N);  % == 1/kappa
  xmean = (1-cmean) * xold + cmean * arx(:,fitness.idxsel(1:mu))*weights; 
  zmean = arz(:,fitness.idxsel(1:mu))*weights;%==D^-1*B'*(xmean-xold)/sigma
  if mu == 1
    fmean = fitness.sel(1);
  else
    fmean = NaN; % [] does not work in the latter assignment
    % fmean = feval(fitfun, xintobounds(xmean, lbounds, ubounds), varargin{:});
    % counteval = counteval + 1;
  end
  
  % Cumulation: update evolution paths
  ps = (1-cs)*ps + sqrt(cs*(2-cs)*mueff) * (B*zmean);          % Eq. (4)
  hsig = norm(ps)/sqrt(1-(1-cs)^(2*countiter))/chiN < 1.4 + 2/(N+1);
  if flg_future_setting
    hsig = sum(ps.^2) / (1-(1-cs)^(2*countiter)) / N < 2 + 4/(N+1); % just simplified
  end
%  hsig = norm(ps)/sqrt(1-(1-cs)^(2*countiter))/chiN < 1.4 + 2/(N+1);
%  hsig = norm(ps)/sqrt(1-(1-cs)^(2*countiter))/chiN < 1.5 + 1/(N-0.5);
%  hsig = norm(ps) < 1.5 * sqrt(N);
%  hsig = 1;

  pc = (1-cc)*pc ...
        + hsig*(sqrt(cc*(2-cc)*mueff)/sigma/cmean) * (xmean-xold);     % Eq. (2)
  if hsig == 0
    % disp([num2str(countiter) ' ' num2str(counteval) ' pc update stalled']);
  end

  % Adapt covariance matrix
  neg.ccov = 0;  % TODO: move parameter setting upwards at some point
  if ccov1 + ccovmu > 0                                                    % Eq. (3)
    if flgDiagonalOnly % internal linear(?) complexity
      diagC = (1-ccov1_sep-ccovmu_sep+(1-hsig)*ccov1_sep*cc*(2-cc)) * diagC ... % regard old matrix 
          + ccov1_sep * pc.^2 ...               % plus rank one update
          + ccovmu_sep ...                      % plus rank mu update
            * (diagC .* (arz(:,fitness.idxsel(1:mu)).^2 * weights));
%             * (repmat(diagC,1,mu) .* arz(:,fitness.idxsel(1:mu)).^2 * weights);
      diagD = sqrt(diagC); % replaces eig(C)
    else
      arpos = (arx(:,fitness.idxsel(1:mu))-repmat(xold,1,mu)) / sigma;
      % "active" CMA update: negative update, in case controlling pos. definiteness 
      if flgActiveCMA > 0
        % set parameters
        neg.mu = mu;  
        neg.mueff = mueff;
        if flgActiveCMA > 10  % flat weights with mu=lambda/2
          neg.mu = floor(lambda/2);  
          neg.mueff = neg.mu;
        end
        % neg.mu = ceil(min([N, lambda/4, mueff]));  neg.mueff = mu; % i.e. neg.mu <= N 
        % Parameter study: in 3-D lambda=50,100, 10-D lambda=200,400, 30-D lambda=1000,2000 a 
        % three times larger neg.ccov does not work. 
        %   increasing all ccov rates three times does work (probably because of the factor (1-ccovmu))
        %   in 30-D to looks fine

        neg.ccov = (1 - ccovmu) * 0.25 * neg.mueff / ((N+2)^1.5 + 2*neg.mueff);
        neg.minresidualvariance = 0.66;  % keep at least 0.66 in all directions, small popsize are most critical
        neg.alphaold = 0.5;  % where to make up for the variance loss, 0.5 means no idea what to do
                             % 1 is slightly more robust and gives a better "guaranty" for pos. def., 
                             % but does it make sense from the learning perspective for large ccovmu? 

        neg.ccovfinal = neg.ccov;

        % prepare vectors, compute negative updating matrix Cneg and checking matrix Ccheck
        arzneg = arz(:,fitness.idxsel(lambda:-1:lambda - neg.mu + 1));
        % i-th longest becomes i-th shortest
        % TODO: this is not in compliance with the paper Hansen&Ros2010, 
        %       where simply arnorms = arnorms(end:-1:1) ? 
        [arnorms idxnorms] = sort(sqrt(sum(arzneg.^2, 1))); 
        [ignore idxnorms] = sort(idxnorms);  % inverse index 
        arnormfacs = arnorms(end:-1:1) ./ arnorms; 
        % arnormfacs = arnorms(randperm(neg.mu)) ./ arnorms;
        arnorms = arnorms(end:-1:1); % for the record
        if flgActiveCMA < 20
          arzneg = arzneg .* repmat(arnormfacs(idxnorms), N, 1);  % E x*x' is N
          % arzneg = sqrt(N) * arzneg ./ repmat(sqrt(sum(arzneg.^2, 1)), N, 1);  % E x*x' is N
        end
        if flgActiveCMA < 10 && neg.mu == mu  % weighted sum
          if mod(flgActiveCMA, 10) == 1 % TODO: prevent this with a less tight but more efficient check (see below) 
            Ccheck = arzneg * diag(weights) * arzneg';  % in order to check the largest EV
          end
          artmp = BD * arzneg;
          Cneg = artmp * diag(weights) * artmp';
        else  % simple sum
          if mod(flgActiveCMA, 10) == 1
            Ccheck = (1/neg.mu) * arzneg*arzneg';  % in order to check largest EV
          end
          artmp = BD * arzneg;
          Cneg = (1/neg.mu) * artmp*artmp';

        end

        % check pos.def. and set learning rate neg.ccov accordingly, 
        % this check makes the original choice of neg.ccov extremly failsafe 
        % still assuming C == BD*BD', which is only approxim. correct 
        if mod(flgActiveCMA, 10) == 1 && 1 - neg.ccov * arnorms(idxnorms).^2 * weights < neg.minresidualvariance
          % TODO: the simple and cheap way would be to set
          %    fac = 1 - ccovmu - ccov1 OR 1 - mueff/lambda and
          %    neg.ccov = fac*(1 - neg.minresidualvariance) / (arnorms(idxnorms).^2 * weights)
          % this is the more sophisticated way: 
          % maxeigenval = eigs(arzneg * arzneg', 1, 'lm', eigsopts);  % not faster
          maxeigenval = max(eig(Ccheck));  % norm is much slower, because (norm()==max(svd())
          %disp([countiter log10([neg.ccov, maxeigenval, arnorms(idxnorms).^2 * weights, max(arnorms)^2]), ...
           %          neg.ccov * arnorms(idxnorms).^2 * weights])
          % pause
          % remove less than ??34*(1-cmu)%?? of variance in any direction
          %     1-ccovmu is the variance left from the old C
          neg.ccovfinal = min(neg.ccov, (1-ccovmu)*(1-neg.minresidualvariance)/maxeigenval); 
                                        % -ccov1 removed to avoid error message??
          if neg.ccovfinal < neg.ccov
            disp(['active CMA at iteration ' num2str(countiter) ...
                 ': max EV ==', num2str([maxeigenval, neg.ccov, neg.ccovfinal])]);
          end
        end
        % xmean = xold;  % the distribution does not degenerate!? 
        % update C
        C = (1-ccov1-ccovmu+neg.alphaold*neg.ccovfinal+(1-hsig)*ccov1*cc*(2-cc)) * C ... % regard old matrix 
            + ccov1 * pc*pc' ...     % plus rank one update
            + (ccovmu + (1-neg.alphaold)*neg.ccovfinal) ...  % plus rank mu update
              * arpos * (repmat(weights,1,N) .* arpos') ...
              - neg.ccovfinal * Cneg;                        % minus rank mu update
      else  % no active (negative) update
        C = (1-ccov1-ccovmu+(1-hsig)*ccov1*cc*(2-cc)) * C ... % regard old matrix 
            + ccov1 * pc*pc' ...     % plus rank one update
            + ccovmu ...             % plus rank mu update
              * arpos * (repmat(weights,1,N) .* arpos');
        % is now O(mu*N^2 + mu*N), was O(mu*N^2 + mu^2*N) when using diag(weights)
        %   for mu=30*N it is now 10 times faster, overall 3 times faster
      end
      diagC = diag(C);
    end
  end
  
  % the following is de-preciated and will be removed in future
  % better setting for cc makes this hack obsolete
  if 11 < 2 && ~flgscience  
    % remove momentum in ps, if ps is large and fitness is getting worse.
    % this should rarely happen. 
    % this might very well be counterproductive in dynamic environments
    if sum(ps.^2)/N > 1.5 + 10*(2/N)^.5 && ...
        fitness.histsel(1) > max(fitness.histsel(2:3))
      ps = ps * sqrt(N*(1+max(0,log(sum(ps.^2)/N))) / sum(ps.^2));
      if flgdisplay
        disp(['Momentum in ps removed at [niter neval]=' ...
              num2str([countiter counteval]) ']']);
      end
    end
  end

  % Adapt sigma
  if flg_future_setting  % according to a suggestion from Dirk Arnold (2000)
    % exp(1) is still not reasonably small enough, maybe 2/3?
    sigma = sigma * exp(min(1, (sum(ps.^2)/N - 1)/2 * cs/damps));            % Eq. (5)
  else
    % exp(1) is still not reasonably small enough
    sigma = sigma * exp(min(1, (sqrt(sum(ps.^2))/chiN - 1) * cs/damps));             % Eq. (5)
  end
  % disp([countiter norm(ps)/chiN]);
  
  if 11 < 3   % testing with optimal step-size
      if countiter == 1
          disp('*********** sigma set to const * ||x|| ******************');
      end
      sigma = 0.04 * mueff * sqrt(sum(xmean.^2)) / N; % 20D,lam=1000:25e3
      sigma = 0.3 * mueff * sqrt(sum(xmean.^2)) / N; % 20D,lam=(40,1000):17e3
                                                     %      75e3 with def (1.5)
                                                     %      35e3 with damps=0.25
  end
  if 11 < 3 
          if countiter == 1
              disp('*********** xmean set to const ******************');
          end
      xmean = ones(N,1);
  end
  
  % Update B and D from C

  if ~flgDiagonalOnly && (ccov1+ccovmu+neg.ccov) > 0 && mod(countiter, 1/(ccov1+ccovmu+neg.ccov)/N/10) < 1
    C=triu(C)+triu(C,1)'; % enforce symmetry to prevent complex numbers
    [B,tmp] = eig(C);     % eigen decomposition, B==normalized eigenvectors
                          % effort: approx. 15*N matrix-vector multiplications
    diagD = diag(tmp); 

    if any(~isfinite(diagD))
      clear idx; % prevents error under octave 
      save(['tmp' opts.SaveFilename]);
      error(['function eig returned non-finited eigenvalues, cond(C)=' ...
	     num2str(cond(C)) ]);
    end
    if any(any(~isfinite(B)))
      clear idx; % prevents error under octave
      save(['tmp' opts.SaveFilename]);
      error(['function eig returned non-finited eigenvectors, cond(C)=' ...
	     num2str(cond(C)) ]);
    end

    % limit condition of C to 1e14 + 1
    if min(diagD) <= 0
	if stopOnWarnings
	  stopflag(end+1) = {'warnconditioncov'};
	else
	  warning(['Iteration ' num2str(countiter) ...
		   ': Eigenvalue (smaller) zero']);
	  diagD(diagD<0) = 0;
	  tmp = max(diagD)/1e14;
	  C = C + tmp*eye(N,N); diagD = diagD + tmp*ones(N,1); 
	end
    end
    if max(diagD) > 1e14*min(diagD) 
	if stopOnWarnings
	  stopflag(end+1) = {'warnconditioncov'};
	else
	  warning(['Iteration ' num2str(countiter) ': condition of C ' ...
		   'at upper limit' ]);
	  tmp = max(diagD)/1e14 - min(diagD);
	  C = C + tmp*eye(N,N); diagD = diagD + tmp*ones(N,1); 
	end
    end

    diagC = diag(C); 
    diagD = sqrt(diagD); % D contains standard deviations now
    % diagD = diagD / prod(diagD)^(1/N);  C = C / prod(diagD)^(2/N);
    BD = B.*repmat(diagD',N,1); % O(n^2)
  end % if mod

  % Align/rescale order of magnitude of scales of sigma and C for nicer output
  % TODO: interference with sigmafacup: replace 1e10 with 2*sigmafacup
  % not a very usual case
  if 1 < 2 && sigma > 1e10*max(diagD) && sigma > 8e14 * max(insigma)
    fac = sigma; % / max(diagD);
    sigma = sigma/fac;
    pc = fac * pc;
    diagD = fac * diagD; 
    if ~flgDiagonalOnly
      C = fac^2 * C; % disp(fac);
      BD = B .* repmat(diagD',N,1); % O(n^2), but repmat might be inefficient todo?
    end
    diagC = fac^2 * diagC; 
  end

  if flgDiagonalOnly > 1 && countiter > flgDiagonalOnly 
    % full covariance matrix from now on 
    flgDiagonalOnly = 0; 
    B = eye(N,N);
    BD = diag(diagD);
    C = diag(diagC); % is better, because correlations are spurious anyway
  end

  if noiseHandling 
   if countiter == 1  % assign firstvarargin for noise treatment e.g. as #reevaluations
     if ~isempty(varargin) && length(varargin{1}) == 1 && isnumeric(varargin{1})
       if irun == 1
         firstvarargin = varargin{1};
       else
         varargin{1} =  firstvarargin;  % reset varargin{1}
       end
     else
       firstvarargin = 0;
     end
   end
   if noiseSS < 0 && noiseMinMaxEvals(2) > noiseMinMaxEvals(1) && firstvarargin
     varargin{1} = max(noiseMinMaxEvals(1), varargin{1} / noiseAlphaEvals^(1/4));  % still experimental
   elseif noiseSS > 0
    if ~isempty(noiseCallback)  % to be removed? 
      res = feval(noiseCallback); % should also work without output argument!?
      if ~isempty(res) && res > 1 % TODO: decide for interface of callback
                                  %       also a dynamic popsize could be done here 
        sigma = sigma * noiseAlpha;
      end
    else
      if noiseMinMaxEvals(2) > noiseMinMaxEvals(1) && firstvarargin
        varargin{1} = min(noiseMinMaxEvals(2), varargin{1} * noiseAlphaEvals);
      end
    
      sigma = sigma * noiseAlpha; 
      % lambda = ceil(0.1 * sqrt(lambda) + lambda);
      % TODO: find smallest increase of lambda with log-linear
      %       convergence in iterations
    end
    % qqq experimental: take a mean to estimate the true optimum
    noiseN = noiseN + 1;
    if noiseN == 1
      noiseX = xmean; 
    else
      noiseX = noiseX + (3/noiseN) * (xmean - noiseX); 
    end
   end
  end

  % ----- numerical error management -----
  % Adjust maximal coordinate axis deviations
  if any(sigma*sqrt(diagC) > maxdx)
    sigma = min(maxdx ./ sqrt(diagC));
    %warning(['Iteration ' num2str(countiter) ': coordinate axis std ' ...
    %         'deviation at upper limit of ' num2str(maxdx)]);
    % stopflag(end+1) = {'maxcoorddev'};
  end
  % Adjust minimal coordinate axis deviations
  if any(sigma*sqrt(diagC) < mindx)
    sigma = max(mindx ./ sqrt(diagC)) * exp(0.05+cs/damps); 
    %warning(['Iteration ' num2str(countiter) ': coordinate axis std ' ...
    %         'deviation at lower limit of ' num2str(mindx)]);
    % stopflag(end+1) = {'mincoorddev'};;
  end
  % Adjust too low coordinate axis deviations
  if any(xmean == xmean + 0.2*sigma*sqrt(diagC)) 
    if stopOnWarnings
      stopflag(end+1) = {'warnnoeffectcoord'};
    else
      warning(['Iteration ' num2str(countiter) ': coordinate axis std ' ...
                'deviation too low' ]);
      if flgDiagonalOnly
        diagC = diagC + (ccov1_sep+ccovmu_sep) * (diagC .* ...
                                                  (xmean == xmean + 0.2*sigma*sqrt(diagC)));
      else
        C = C + (ccov1+ccovmu) * diag(diagC .* ...
                                      (xmean == xmean + 0.2*sigma*sqrt(diagC)));
      end
      sigma = sigma * exp(0.05+cs/damps); 
    end
  end
  % Adjust step size in case of (numerical) precision problem 
  if flgDiagonalOnly
    tmp = 0.1*sigma*diagD; 
  else
    tmp = 0.1*sigma*BD(:,1+floor(mod(countiter,N)));
  end
  if all(xmean == xmean + tmp)
    i = 1+floor(mod(countiter,N));
    if stopOnWarnings
	stopflag(end+1) = {'warnnoeffectaxis'};
    else
      warning(['Iteration ' num2str(countiter) ...
	       ': main axis standard deviation ' ...
	       num2str(sigma*diagD(i)) ' has no effect' ]);
      sigma = sigma * exp(0.2+cs/damps); 
    end
  end
  % Adjust step size in case of equal function values (flat fitness)
  % isequalfuncvalues = 0; 
  if fitness.sel(1) == fitness.sel(1+ceil(0.1+lambda/4))
    % isequalfuncvalues = 1; 
    if stopOnEqualFunctionValues
      arrEqualFunvals = [countiter arrEqualFunvals(1:end-1)];
      % stop if this happens in more than 33%
      if arrEqualFunvals(end) > countiter - 3 * length(arrEqualFunvals)
        stopflag(end+1) = {'equalfunvals'}; 
      end
    else
      if flgWarnOnEqualFunctionValues
        warning(['Iteration ' num2str(countiter) ...
		 ': equal function values f=' num2str(fitness.sel(1)) ...
		 ' at maximal main axis sigma ' ...
        num2str(sigma*max(diagD))]);
      end
      sigma = sigma * exp(0.2+cs/damps); 
    end
  end
  % Adjust step size in case of equal function values
  if countiter > 2 && myrange([fitness.hist fitness.sel(1)]) == 0  
    if stopOnWarnings
	stopflag(end+1) = {'warnequalfunvalhist'};
    else
      warning(['Iteration ' num2str(countiter) ...
	       ': equal function values in history at maximal main ' ...
	       'axis sigma ' num2str(sigma*max(diagD))]);
      sigma = sigma * exp(0.2+cs/damps); 
    end
  end
    
  % ----- end numerical error management -----
  
  % Keep overall best solution
  out.evals = counteval;
  out.solutions.evals = counteval;
  out.solutions.mean.x = xmean;
  out.solutions.mean.f = fmean;
  out.solutions.mean.evals = counteval;
  out.solutions.recentbest.x = arxvalid(:, fitness.idx(1));
  out.solutions.recentbest.f = fitness.raw(1);
  out.solutions.recentbest.evals = counteval + fitness.idx(1) - lambda;
  out.solutions.recentworst.x = arxvalid(:, fitness.idx(end));
  out.solutions.recentworst.f = fitness.raw(end);
  out.solutions.recentworst.evals = counteval + fitness.idx(end) - lambda;
  if fitness.hist(1) < out.solutions.bestever.f
    out.solutions.bestever.x = arxvalid(:, fitness.idx(1));
    out.solutions.bestever.f = fitness.hist(1);
    out.solutions.bestever.evals = counteval + fitness.idx(1) - lambda;
    bestever = out.solutions.bestever;
  end

  % Set stop flag
  if fitness.raw(1) <= stopFitness, stopflag(end+1) = {'fitness'}; end
  if counteval >= stopMaxFunEvals, stopflag(end+1) = {'maxfunevals'}; end
  if countiter >= stopMaxIter, stopflag(end+1) = {'maxiter'}; end
  if all(sigma*(max(abs(pc), sqrt(diagC))) < stopTolX) 
    stopflag(end+1) = {'tolx'};
  end
  if any(sigma*sqrt(diagC) > stopTolUpX) 
    stopflag(end+1) = {'tolupx'};
  end
  if sigma*max(diagD) == 0  % should never happen
    stopflag(end+1) = {'bug'};
  end
  if countiter > 2 && myrange([fitness.sel fitness.hist]) <= stopTolFun 
    stopflag(end+1) = {'tolfun'};
  end
  if countiter >= length(fitness.hist) && myrange(fitness.hist) <= stopTolHistFun 
    stopflag(end+1) = {'tolhistfun'};
  end
  l = floor(length(fitness.histbest)/3);
  if 1 < 2 && stopOnStagnation && ...  % leads sometimes early stop on ftablet, fcigtab
      countiter > N * (5+100/lambda) && ...  
      length(fitness.histbest) > 100 && ... 
      median(fitness.histmedian(1:l)) >= median(fitness.histmedian(end-l:end)) && ...
      median(fitness.histbest(1:l)) >= median(fitness.histbest(end-l:end))
    stopflag(end+1) = {'stagnation'};
  end

  if counteval >= stopFunEvals || countiter >= stopIter
    stopflag(end+1) = {'stoptoresume'};
    if length(stopflag) == 1 && flgsaving == 0
      error('To resume later the saving option needs to be set');
    end
  end
  % read stopping message from file signals.par 
  if flgreadsignals
    fid = fopen('./signals.par', 'rt');  % can be performance critical 
    while fid > 0
      strline = fgetl(fid); %fgets(fid, 300);
      if strline < 0 % fgets and fgetl returns -1 at end of file
        break;
      end
      % 'stop filename' sets stopflag to manual
      str = sscanf(strline, ' %s %s', 2);
      if strcmp(str, ['stop' opts.LogFilenamePrefix]) 
        stopflag(end+1) = {'manual'};
        break;
      end
      % 'skip filename run 3' skips a run, but not the last
      str = sscanf(strline, ' %s %s %s', 3);
      if strcmp(str, ['skip' opts.LogFilenamePrefix 'run'])
        i = strfind(strline, 'run');
        if irun == sscanf(strline(i+3:end), ' %d ', 1) && irun <= myeval(opts.Restarts)
          stopflag(end+1) = {'skipped'};
        end      
      end
    end % while, break 
    if fid > 0
      fclose(fid);
      clear fid; % prevents strange error under octave
    end
  end
  
  out.stopflag = stopflag;

  % ----- output generation -----
  if verbosemodulo > 0 && isfinite(verbosemodulo)
    if countiter == 1 || mod(countiter, 10*verbosemodulo) < 1 
      disp(['Iterat, #Fevals:   Function Value    (median,worst) ' ...
	    '|Axis Ratio|' ...
	    'idx:Min SD idx:Max SD']); 
    end
    if mod(countiter, verbosemodulo) < 1 ...
	  || (verbosemodulo > 0 && isfinite(verbosemodulo) && ...
	      (countiter < 3 || ~isempty(stopflag)))
      [minstd minstdidx] = min(sigma*sqrt(diagC));
      [maxstd maxstdidx] = max(sigma*sqrt(diagC));
      % format display nicely
      disp([repmat(' ',1,4-floor(log10(countiter))) ...
	    num2str(countiter) ' , ' ...
	    repmat(' ',1,5-floor(log10(counteval))) ...
	    num2str(counteval) ' : ' ...
            num2str(fitness.hist(1), '%.13e') ...
	    ' +(' num2str(median(fitness.raw)-fitness.hist(1), '%.0e ') ...
	    ',' num2str(max(fitness.raw)-fitness.hist(1), '%.0e ') ...
	    ') | ' ...
	    num2str(max(diagD)/min(diagD), '%4.2e') ' | ' ...
	    repmat(' ',1,1-floor(log10(minstdidx))) num2str(minstdidx) ':' ...
	    num2str(minstd, ' %.1e') ' ' ...
	    repmat(' ',1,1-floor(log10(maxstdidx))) num2str(maxstdidx) ':' ...
	    num2str(maxstd, ' %.1e')]);
    end
  end

  % measure time for recording data
  if countiter < 3 
    time.c = 0.05;
    time.nonoutput = 0;
    time.recording = 0;
    time.saving  = 0.15; % first saving after 3 seconds of 100 iterations
    time.plotting = 0;
  elseif countiter > 300
    % set backward horizon, must be long enough to cover infrequent plotting etc
    % time.c = min(1, time.nonoutput/3 + 1e-9); 
    time.c = max(1e-5, 0.1/sqrt(countiter)); % mean over all or 1e-5
  end
  % get average time per iteration
  time.t1 = clock;
  time.act = max(0,etime(time.t1, time.t0));
  time.nonoutput = (1-time.c) * time.nonoutput ...
      + time.c * time.act; 

  time.recording = (1-time.c) * time.recording;  % per iteration
  time.saving  = (1-time.c) * time.saving;
  time.plotting = (1-time.c) * time.plotting;
  
  % record output data, concerning time issues
  if savemodulo && savetime && (countiter < 1e2 || ~isempty(stopflag) || ...
	countiter >= outiter + savemodulo)
    outiter = countiter; 
      % Save output data to files  
      for namecell = filenames(:)'
        name = namecell{:};

	[fid, err] = fopen(['./' filenameprefix name '.dat'], 'a');
	if fid < 1 % err ~= 0 
	  warning(['could not open ' filenameprefix name '.dat']);
	else
	  if strcmp(name, 'axlen')
	    fprintf(fid, '%d %d %e %e %e ', countiter, counteval, sigma, ...
		max(diagD), min(diagD)); 
            fprintf(fid, '%e ', sort(diagD)); 
            fprintf(fid, '\n');
	  elseif strcmp(name, 'disp') % TODO
	  elseif strcmp(name, 'fit')
	    fprintf(fid, '%ld %ld %e %e %25.18e %25.18e %25.18e %25.18e', ...
                    countiter, counteval, sigma, max(diagD)/min(diagD), ...
                    out.solutions.bestever.f, ...
                    fitness.raw(1), median(fitness.raw), fitness.raw(end)); 
            if ~isempty(varargin) && length(varargin{1}) == 1 && isnumeric(varargin{1}) && varargin{1} ~= 0
              fprintf(fid, ' %f', varargin{1});
            end                    
	    fprintf(fid, '\n');
	  elseif strcmp(name, 'stddev')
	    fprintf(fid, '%ld %ld %e 0 0 ', countiter, counteval, sigma); 
	    fprintf(fid, '%e ', sigma*sqrt(diagC)); 
            fprintf(fid, '\n');
	  elseif strcmp(name, 'xmean')
	    if isnan(fmean)
	      fprintf(fid, '%ld %ld 0 0 0 ', countiter, counteval); 
	    else
	      fprintf(fid, '%ld %ld 0 0 %e ', countiter, counteval, fmean); 
	    end
	    fprintf(fid, '%e ', xmean); 
            fprintf(fid, '\n');
	  elseif strcmp(name, 'xrecentbest')
            % TODO: fitness is inconsistent with x-value
	    fprintf(fid, '%ld %ld %25.18e 0 0 ', countiter, counteval, fitness.raw(1)); 
	    fprintf(fid, '%e ', arx(:,fitness.idx(1))); 
            fprintf(fid, '\n');
	  end
	  fclose(fid); 
	end
  end

    % get average time for recording data
    time.t2 = clock;
    time.recording = time.recording + time.c * max(0,etime(time.t2, time.t1)); 
    
    % plot
    if flgplotting && countiter > 1
      if countiter == 2
        iterplotted = 0;
      end
      if ~isempty(stopflag) || ... 
        ((time.nonoutput+time.recording) * (countiter - iterplotted) > 1 && ...
          time.plotting < 0.05 * (time.nonoutput+time.recording))
        local_plotcmaesdat(324, filenameprefix);
        iterplotted = countiter; 
        %  outplot(out); % outplot defined below
        if time.plotting == 0  % disregard opening of the window
          time.plotting = time.nonoutput+time.recording;
        else
          time.plotting = time.plotting + time.c * max(0,etime(clock, time.t2)); 
        end
      end
    end
    if countiter > 100 + 20 && savemodulo && ...
          time.recording * countiter > 0.1 && ...  % absolute time larger 0.1 second
	  time.recording > savetime * (time.nonoutput+time.recording) / 100 
      savemodulo = floor(1.02 * savemodulo) + 1;
      % disp(['++savemodulo == ' num2str(savemodulo) ' at ' num2str(countiter)]); %qqq
    end
  end % if output

  % save everything
  time.t3 = clock;
  if ~isempty(stopflag) || time.saving < 0.05 * time.nonoutput || countiter == 100
    xmin = arxvalid(:, fitness.idx(1));
    fmin = fitness.raw(1);
    if flgsaving && countiter > 2
      clear idx; % prevents error under octave
      % -v6 : non-compressed non-unicode for version 6 and earlier
      if ~isempty(strsaving) && ~isoctave
	save('-mat', strsaving, opts.SaveFilename); % for inspection and possible restart	
      else 
	save('-mat', opts.SaveFilename); % for inspection and possible restart
      end
      time.saving = time.saving + time.c * max(0,etime(clock, time.t3)); 
    end
  end
  time.t0 = clock;

  % ----- end output generation -----

end % while, end generation loop

% -------------------- Final Procedures -------------------------------

% Evaluate xmean and return best recent point in xmin
fmin = fitness.raw(1);
xmin = arxvalid(:, fitness.idx(1)); % Return best point of last generation.
if length(stopflag) > sum(strcmp(stopflag, 'stoptoresume')) % final stopping
  out.solutions.mean.f = ...
      feval(fitfun, xintobounds(xmean, lbounds, ubounds), varargin{:});
  counteval = counteval + 1;
  out.solutions.mean.evals = counteval;
  if out.solutions.mean.f < fitness.raw(1)
    fmin = out.solutions.mean.f;
    xmin = xintobounds(xmean, lbounds, ubounds); % Return xmean as best point
  end
  if out.solutions.mean.f < out.solutions.bestever.f
    out.solutions.bestever = out.solutions.mean; % Return xmean as bestever point
    out.solutions.bestever.x = xintobounds(xmean, lbounds, ubounds); 
    bestever = out.solutions.bestever;
  end
end

% Save everything and display final message
if flgsavingfinal
  clear idx; % prevents error under octave
  if ~isempty(strsaving) && ~isoctave
    save('-mat', strsaving, opts.SaveFilename); % for inspection and possible restart	
  else 
    save('-mat', opts.SaveFilename);    % for inspection and possible restart
  end
  message = [' (saved to ' opts.SaveFilename ')'];
else
  message = [];
end

if flgdisplay
  disp(['#Fevals:   f(returned x)   |    bestever.f     | stopflag' ...
        message]);
  if isoctave
    strstop = stopflag(:); 
  else
      strcat(stopflag(:), '.');
  end
  strstop = stopflag(:); %strcat(stopflag(:), '.');
  disp([repmat(' ',1,6-floor(log10(counteval))) ...
        num2str(counteval, '%6.0f') ': ' num2str(fmin, '%.11e') ' | ' ...
        num2str(out.solutions.bestever.f, '%.11e') ' | ' ...
	strstop{1:end}]);
  if N < 102
     disp(['mean solution:' sprintf(' %+.1e', xmean)]);
     disp(['std deviation:' sprintf('  %.1e', sigma*sqrt(diagC))]);
     disp(sprintf('use plotcmaesdat.m for plotting the output at any time (option LogModulo must not be zero)'));
  end
  if exist('sfile', 'var') 
    disp(['Results saved in ' sfile]); 
  end
end

  out.arstopflags{irun} = stopflag;
  if any(strcmp(stopflag, 'fitness')) ...
	|| any(strcmp(stopflag, 'maxfunevals')) ...
	|| any(strcmp(stopflag, 'stoptoresume')) ...
	|| any(strcmp(stopflag, 'manual'))
    break; 
  end
end % while irun <= Restarts

% ---------------------------------------------------------------  
% ---------------------------------------------------------------  
function [x, idx] = xintobounds(x, lbounds, ubounds)
%
% x can be a column vector or a matrix consisting of column vectors
%
  if ~isempty(lbounds)
    if length(lbounds) == 1
      idx = x < lbounds;
      x(idx) = lbounds;
    else
      arbounds = repmat(lbounds, 1, size(x,2));
      idx = x < arbounds;
      x(idx) = arbounds(idx);
    end
  else
    idx = 0;
  end
  if ~isempty(ubounds)
    if length(ubounds) == 1
      idx2 = x > ubounds;
      x(idx2) = ubounds;
    else
      arbounds = repmat(ubounds, 1, size(x,2));
      idx2 = x > arbounds;
      x(idx2) = arbounds(idx2);
    end
  else
    idx2 = 0;
  end
  idx = idx2-idx; 
  
% ---------------------------------------------------------------  
% ---------------------------------------------------------------  
function opts=getoptions(inopts, defopts)
% OPTS = GETOPTIONS(INOPTS, DEFOPTS) handles an arbitrary number of
% optional arguments to a function. The given arguments are collected
% in the struct INOPTS.  GETOPTIONS matches INOPTS with a default
% options struct DEFOPTS and returns the merge OPTS.  Empty or missing
% fields in INOPTS invoke the default value.  Fieldnames in INOPTS can
% be abbreviated.
%
% The returned struct OPTS is first assigned to DEFOPTS. Then any
% field value in OPTS is replaced by the respective field value of
% INOPTS if (1) the field unambiguously (case-insensitive) matches
% with the fieldname in INOPTS (cut down to the length of the INOPTS
% fieldname) and (2) the field is not empty.
%
% Example:
%   In the source-code of the function that needs optional
%   arguments, the last argument is the struct of optional
%   arguments:
%
%   function results = myfunction(mandatory_arg, inopts)
%     % Define four default options
%     defopts.PopulationSize = 200;
%     defopts.ParentNumber = 50;
%     defopts.MaxIterations = 1e6;
%     defopts.MaxSigma = 1;
%  
%     % merge default options with input options
%     opts = getoptions(inopts, defopts);
%
%     % Thats it! From now on the values in opts can be used
%     for i = 1:opts.PopulationSize
%       % do whatever
%       if sigma > opts.MaxSigma
%         % do whatever
%       end
%     end
%   
%   For calling the function myfunction with default options:
%   myfunction(argument1, []);
%   For calling the function myfunction with modified options:
%   opt.pop = 100; % redefine PopulationSize option
%   opt.PAR = 10;  % redefine ParentNumber option
%   opt.maxiter = 2; % opt.max=2 is ambiguous and would result in an error 
%   myfunction(argument1, opt);

%
% 04/07/19: Entries can be structs itself leading to a recursive
%           call to getoptions. 
%

if nargin < 2 || isempty(defopts) % no default options available
  opts=inopts;
  return;
elseif isempty(inopts) % empty inopts invoke default options
  opts = defopts;
  return;
elseif ~isstruct(defopts) % handle a single option value
  if isempty(inopts) 
    opts = defopts;
  elseif ~isstruct(inopts)
    opts = inopts;
  else
    error('Input options are a struct, while default options are not');
  end
  return;
elseif ~isstruct(inopts) % no valid input options
  error('The options need to be a struct or empty');
end

  opts = defopts; % start from defopts 
  % if necessary overwrite opts fields by inopts values
  defnames = fieldnames(defopts);
  idxmatched = []; % indices of defopts that already matched
  for name = fieldnames(inopts)'
    name = name{1}; % name of i-th inopts-field
    if isoctave
      for i = 1:size(defnames, 1)
        idx(i) = strncmpi(defnames(i), name, length(name));
      end
    else
	idx = strncmpi(defnames, name, length(name));
    end
    if sum(idx) > 1
      error(['option "' name '" is not an unambigous abbreviation. ' ...
	     'Use opts=RMFIELD(opts, ''' name, ...
	     ''') to remove the field from the struct.']);
    end
    if sum(idx) == 1
      defname  = defnames{find(idx)}; 
      if ismember(find(idx), idxmatched)
	error(['input options match more than ones with "' ...
	       defname '". ' ...
	       'Use opts=RMFIELD(opts, ''' name, ...
	       ''') to remove the field from the struct.']);
      end
      idxmatched = [idxmatched find(idx)];
      val = getfield(inopts, name);
      % next line can replace previous line from MATLAB version 6.5.0 on and in octave
      % val = inopts.(name);
      if isstruct(val) % valid syntax only from version 6.5.0
	opts = setfield(opts, defname, ...
	    getoptions(val, getfield(defopts, defname))); 
      elseif isstruct(getfield(defopts, defname)) 
      % next three lines can replace previous three lines from MATLAB 
      % version 6.5.0 on
      %   opts.(defname) = ...
      %      getoptions(val, defopts.(defname)); 
      % elseif isstruct(defopts.(defname)) 
	warning(['option "' name '" disregarded (must be struct)']); 
      elseif ~isempty(val) % empty value: do nothing, i.e. stick to default
	opts = setfield(opts, defnames{find(idx)}, val);
	% next line can replace previous line from MATLAB version 6.5.0 on
	% opts.(defname) = inopts.(name); 
      end
    else
      warning(['option "' name '" disregarded (unknown field name)']);
    end
  end

% ---------------------------------------------------------------  
% ---------------------------------------------------------------  
function res=myeval(s)
  if ischar(s)
    res = evalin('caller', s);
  else
    res = s;
  end
  
% ---------------------------------------------------------------  
% ---------------------------------------------------------------  
function res=myevalbool(s)
  if ~ischar(s) % s may not and cannot be empty
    res = s;
  else % evaluation string s
    if strncmpi(s, 'yes', 3) || strncmpi(s, 'on', 2) ...
	  || strncmpi(s, 'true', 4) || strncmp(s, '1 ', 2)
      res = 1;
    elseif strncmpi(s, 'no', 2) || strncmpi(s, 'off', 3) ...
	  || strncmpi(s, 'false', 5) || strncmp(s, '0 ', 2)
      res = 0;
    else
      try res = evalin('caller', s); catch
	error(['String value "' s '" cannot be evaluated']);
      end
      try res ~= 0; catch
	error(['String value "' s '" cannot be evaluated reasonably']);
      end
    end
  end
  

% ---------------------------------------------------------------  
% ---------------------------------------------------------------  
function res = isoctave
% any hack to find out whether we are running octave
  s = version;
  res = 0;
  if exist('fflush', 'builtin') && eval(s(1)) < 7
    res = 1;
  end

% ---------------------------------------------------------------  
% ---------------------------------------------------------------  
function flush
  if isoctave
    feval('fflush', stdout);
  end

% ---------------------------------------------------------------  
% ---------------------------------------------------------------  
% ----- replacements for statistic toolbox functions ------------
% ---------------------------------------------------------------  
% ---------------------------------------------------------------  
function res=myrange(x)
  res = max(x) - min(x);
  
% ---------------------------------------------------------------  
% ---------------------------------------------------------------  
function res = myprctile(inar, perc, idx)
%
% Computes the percentiles in vector perc from vector inar
% returns vector with length(res)==length(perc)
% idx: optional index-array indicating sorted order
%

N = length(inar);
flgtranspose = 0;

% sizes 
if size(perc,1) > 1
  perc = perc';
  flgtranspose = 1;
  if size(perc,1) > 1
    error('perc must not be a matrix');
  end
end
if size(inar, 1) > 1 && size(inar,2) > 1
  error('data inar must not be a matrix');
end
 
% sort inar
if nargin < 3 || isempty(idx)
  [sar idx] = sort(inar);
else
  sar = inar(idx);
end

res = [];
for p = perc
  if p <= 100*(0.5/N)
    res(end+1) = sar(1);
  elseif p >= 100*((N-0.5)/N)
    res(end+1) = sar(N);
  else
    % find largest index smaller than required percentile
    availablepercentiles = 100*((1:N)-0.5)/N;
    i = max(find(p > availablepercentiles));
    % interpolate linearly
    res(end+1) = sar(i) ...
	+ (sar(i+1)-sar(i))*(p - availablepercentiles(i)) ...
	/ (availablepercentiles(i+1) - availablepercentiles(i));

  end
end

if flgtranspose
  res = res';
end


% ---------------------------------------------------------------  
% ---------------------------------------------------------------  
% ---------------------------------------------------------------  
% ---------------------------------------------------------------  


function [s ranks rankDelta] = local_noisemeasurement(arf1, arf2, lamreev, theta, cutlimit)
% function [s ranks rankDelta] = noisemeasurement(arf1, arf2, lamreev, theta)
%
% Input: 
%   arf1, arf2 : two arrays of function values. arf1 is of size 1xlambda, 
%       arf2 may be of size 1xlamreev or 1xlambda. The first lamreev values 
%       in arf2 are (re-)evaluations of the respective solutions, i.e. 
%       arf1(1) and arf2(1) are two evaluations of "the first" solution.
%    lamreev: number of reevaluated individuals in arf2 
%    theta : parameter theta for the rank change limit, between 0 and 1, 
%       typically between 0.2 and 0.7. 
%    cutlimit (optional): output s is computed as a mean of rankchange minus 
%       threshold, where rankchange is <=2*(lambda-1). cutlimit limits 
%       abs(rankchange minus threshold) in this calculation to cutlimit. 
%       cutlimit=1 evaluates basically the sign only. cutlimit=2 could be 
%       the rank change with one solution (both evaluations of it). 
% 
% Output: 
%   s : noise measurement, s>0 means the noise measure is above the
%       acceptance threshold
%   ranks : 2xlambda array, corresponding to [arf1; arf2], of ranks 
%       of arf1 and arf2 in the set [arf1 arf2], values are in [1:2*lambda]
%   rankDelta: 1xlambda array of rank movements of arf2 compared to
%       arf1.  rankDelta(i) agrees with the number of values from
%       the set [arf1 arf2] that lie between arf1(i) and arf2(i).
%
% Note: equal function values might lead to somewhat spurious results.
%       For this case a revision is advisable. 

%%% verify input argument sizes
if size(arf1,1) ~= 1
  error('arf1 must be an 1xlambda array');
elseif size(arf2,1) ~= 1
  error('arf2 must be an 1xsomething array');
elseif size(arf1,2) < size(arf2,2)  % not really necessary, but saver
  error('arf2 must not be smaller than arf1 in length');
end
lam = size(arf1, 2);
if size(arf1,2) ~= size(arf2,2)
   arf2(end+1:lam) = arf1((size(arf2,2)+1):lam);
end
if nargin < 5
  cutlimit = inf;
end

%%% capture unusual values
if any(diff(arf1) == 0)
  % this will presumably interpreted as rank change, because
  % sort(ones(...)) returns 1,2,3,...
  warning([num2str(sum(diff(arf1)==0)) ' equal function values']);
end

%%% compute rank changes into rankDelta
% compute ranks
[ignore, idx] = sort([arf1 arf2]);
[ignore, ranks] = sort(idx);
ranks = reshape(ranks, lam, 2)';

rankDelta = ranks(1,:) - ranks(2,:) - sign(ranks(1,:) - ranks(2,:));

%%% compute rank change limits using both ranks(1,...) and ranks(2,...)
for i = 1:lamreev
  sumlim(i) = ...
      0.5 * (...
          myprctile(abs((1:2*lam-1) - (ranks(1,i) - (ranks(1,i)>ranks(2,i)))), ...
                    theta*50) ...      
          + myprctile(abs((1:2*lam-1) - (ranks(2,i) - (ranks(2,i)>ranks(1,i)))), ...
                      theta*50)); 
end

%%% compute measurement
%s = abs(rankDelta(1:lamreev)) - sumlim; % lives roughly in 0..2*lambda

%                               max: 1 rankchange in 2*lambda is always fine
s = abs(rankDelta(1:lamreev)) - max(1, sumlim); % lives roughly in 0..2*lambda

% cut-off limit
idx = abs(s) > cutlimit; 
s(idx) = sign(s(idx)) * cutlimit;
s = mean(s);

% CHANGES
% 12/04/28: (3.61) stopIter is relative to countiter after resume (thanks to Tom Holden)
% 12/04/28: (3.61) some syncing from 3.32.integer branch (cmean introduced, ...)
% 12/02/19: "future" setting of ccum, correcting for large mueff, is default now
% 11/11/15: bug-fix: max value for ccovmu_sep setting corrected
% 10/11/11: (3.52.beta) boundary handling: replace max with min in change 
%           rate formula. Active CMA: check of pos.def. improved. 
%           Plotting: value of lambda appears in the title. 
% 10/04/03: (3.51.beta) active CMA cleaned up. Equal fitness detection
%           looks into history now. 
% 10/03/08: (3.50.beta) "active CMA" revised and bug-fix of ambiguous
%           option Noise.alpha -> Noise.alphasigma. 
% 09/10/12: (3.40.beta) a slightly modified version of "active CMA", 
%           that is a negative covariance matrix update, use option
%           CMA.active. In 10;30;90-D the gain on ftablet is a factor 
%           of 1.6;2.5;4.4 (the scaling improves by sqrt(N)). On 
%           Rosenbrock the gain is about 25%. On sharp ridge the 
%           behavior is improved. Cigar is unchanged. 
% 09/08/10: local plotcmaesdat remains in backround  
% 09/08/10: bug-fix in time management for data writing, logtime was not 
%        considered properly (usually not at all). 
% 09/07/05: V3.24: stagnation termination added 
% 08/09/27: V3.23: momentum alignment is out-commented and de-preciated
% 08/09/25: V3.22: re-alignment of sigma and C was buggy
% 08/07/15: V3.20, CMA-parameters are options now. ccov and mucov were replaced
%        by ccov1 \approx ccov/mucov and ccovmu \approx (1-1/mucov)*ccov
% 08/06/30: file name xrecent was change to xrecentbest (compatible with other 
%        versions)
% 08/06/29: time stamp added to output files 
% 08/06/28: bug fixed with resume option, commentary did not work 
% 08/06/28: V3.10, uncertainty (noise) handling added (re-implemented), according
%        to reference "A Method for Handling Uncertainty..." from below.
% 08/06/28: bug fix: file xrecent was empty 
% 08/06/01: diagonalonly clean up. >1 means some iterations. 
% 08/05/05: output is written to file preventing an increasing data
%        array and ease long runs. 
% 08/03/27: DiagonalOnly<0 learns for -DiagonalOnly iterations only the 
%        diagonal with a larger learning rate. 
% 08/03 (2.60): option DiagonalOnly>=1 invokes a time- and space-linear  
%        variant with only diagonal elements of the covariance matrix 
%        updating.  This can be useful for large dimensions, say > 100. 
% 08/02: diag(weights) * ... replaced with repmat(weights,1,N) .* ...
%        in C update, implies O(mu*N^2) instead of O(mu^2*N + mu*N^2). 
% 07/09: tolhistfun as termination criterion added, "<" changed to
%        "<=" also for TolFun to allow for stopping on zero difference. 
%        Name tolfunhist clashes with option tolfun. 
% 07/07: hsig threshold made slighly smaller for large dimension, 
%        useful for lambda < lambda_default. 
% 07/06: boundary handling: scaling in the boundary handling
%        is omitted now, see bnd.flgscale. This seems not to
%        have a big impact. Using the scaling is worse on rotated
%        functions, but better on separable ones. 
% 07/05: boundary handling: weight i is not incremented anymore
%        if xmean(i) moves towards the feasible space. Increment
%        factor changed to 1.2 instead of 1.1. 
% 07/05: boundary handling code simplified not changing the algorithm
% 07/04: bug removed for saving in octave
% 06/11/10: more testing of outcome of eig, fixed max(D) to max(diag(D))
% 06/10/21: conclusive final bestever assignment in the end 
% 06/10/21: restart and incpopsize option implemented for restarts
%        with increasing population size, version 2.50. 
% 06/09/16: output argument bestever inserted again for convenience and
%        backward compatibility
% 06/08: output argument out and struct out reorganized. 
% 06/01: Possible parallel evaluation included as option EvalParallel
% 05/11: Compatibility to octave implemented, package octave-forge
%   is needed. 
% 05/09: Raise of figure and waiting for first plots improved
% 05/01: Function coordinatesystem cleaned up. 
% 05/01: Function prctile, which requires the statistics toolbox,
%        replaced by myprctile. 
% 05/01: Option warnonequalfunctionvalues included. 
% 04/12: Decrease of sigma removed. Problems on fsectorsphere can 
%        be addressed better by adding search space boundaries. 
% 04/12: Boundary handling simpyfied. 
% 04/12: Bug when stopping criteria tolx or tolupx are vectors. 
% 04/11: Three input parameters are obligatory now. 
% 04/11: Bug in boundary handling removed: Boundary weights can decrease now. 
% 04/11: Normalization for boundary weights scale changed. 
% 04/11: VerboseModulo option bug removed. Documentation improved. 
% 04/11: Condition for increasing boundary weights changed.
% 04/10: Decrease of sigma when fitness is getting consistenly
%        worse. Addresses the problems appearing on fsectorsphere for
%        large population size.
% 04/10: VerboseModulo option included. 
% 04/10: Bug for condition for increasing boundary weights removed.
% 04/07: tolx depends on initial sigma to achieve scale invariance
%        for this stopping criterion. 
% 04/06: Objective function value NaN is not counted as function
%        evaluation and invokes resampling of the search point. 
% 04/06: Error handling for eigenvalue beeing zero (never happens
%        with default parameter setting)
% 04/05: damps further tuned for large mueff 
%      o Details for stall of pc-adaptation added (variable hsig 
%        introduced). 
% 04/05: Bug in boundary handling removed: A large initial SIGMA was
%        corrected not until *after* the first iteration, which could
%        lead to a complete failure.
% 04/05: Call of function range (works with stats toolbox only) 
%        changed to myrange. 
% 04/04: Parameter cs depends on mueff now and damps \propto sqrt(mueff)
%        instead of \propto mueff. 
%      o Initial stall to adapt C (flginiphase) is removed and
%        adaptation of pc is stalled for large norm(ps) instead.
%      o Returned default options include documentation. 
%      o Resume part reorganized.
% 04/03: Stopflag becomes cell-array. 

% ---------------------------------------------------------------
%%% REFERENCES
%
% The equation numbers refer to 
% Hansen, N. and S. Kern (2004). Evaluating the CMA Evolution
% Strategy on Multimodal Test Functions.  Eighth International
% Conference on Parallel Problem Solving from Nature PPSN VIII,
% Proceedings, pp. 282-291, Berlin: Springer. 
% (http://www.bionik.tu-berlin.de/user/niko/ppsn2004hansenkern.pdf)
% 
% Further references:
% Hansen, N. and A. Ostermeier (2001). Completely Derandomized
% Self-Adaptation in Evolution Strategies. Evolutionary Computation,
% 9(2), pp. 159-195.
% (http://www.bionik.tu-berlin.de/user/niko/cmaartic.pdf).
%
% Hansen, N., S.D. Mueller and P. Koumoutsakos (2003). Reducing the
% Time Complexity of the Derandomized Evolution Strategy with
% Covariance Matrix Adaptation (CMA-ES). Evolutionary Computation,
% 11(1).  (http://mitpress.mit.edu/journals/pdf/evco_11_1_1_0.pdf).
%
% Ros, R. and N. Hansen (2008). A Simple Modification in CMA-ES
% Achieving Linear Time and Space Complexity. To appear in Tenth
% International Conference on Parallel Problem Solving from Nature
% PPSN X, Proceedings, Berlin: Springer.
%
% Hansen, N., A.S.P. Niederberger, L. Guzzella and P. Koumoutsakos
% (2009?). A Method for Handling Uncertainty in Evolutionary
% Optimization with an Application to Feedback Control of
% Combustion. To appear in IEEE Transactions on Evolutionary
% Computation.

% ---------------------------------------------------------------
%
% The BSD 3-Clause License
%
% Redistribution and use in source and binary forms, with or without
% modification, are permitted provided that the following conditions
% are met:
%
% 1. Redistributions of source code must retain the above copyright and
%    authors notice, this list of conditions and the following disclaimer.
%
% 2. Redistributions in binary form must reproduce the above copyright
%    and authors notice, this list of conditions and the following
%    disclaimer in the documentation and/or other materials provided with
%    the distribution.
%
% 3. Neither the name of the copyright holder nor the names of its
%    contributors nor the authors names may be used to endorse or promote
%    products derived from this software without specific prior written
%    permission.
%
% THIS SOFTWARE IS PROVIDED BY THE COPYRIGHT HOLDERS AND CONTRIBUTORS
% "AS IS" AND ANY EXPRESS OR IMPLIED WARRANTIES, INCLUDING, BUT NOT
% LIMITED TO, THE IMPLIED WARRANTIES OF MERCHANTABILITY AND FITNESS FOR
% A PARTICULAR PURPOSE ARE DISCLAIMED. IN NO EVENT SHALL THE COPYRIGHT
% HOLDER OR CONTRIBUTORS BE LIABLE FOR ANY DIRECT, INDIRECT,
% INCIDENTAL, SPECIAL, EXEMPLARY, OR CONSEQUENTIAL DAMAGES (INCLUDING,
% BUT NOT LIMITED TO, PROCUREMENT OF SUBSTITUTE GOODS OR SERVICES; LOSS
% OF USE, DATA, OR PROFITS; OR BUSINESS INTERRUPTION) HOWEVER CAUSED
% AND ON ANY THEORY OF LIABILITY, WHETHER IN CONTRACT, STRICT
% LIABILITY, OR TORT (INCLUDING NEGLIGENCE OR OTHERWISE) ARISING IN ANY
% WAY OUT OF THE USE OF THIS SOFTWARE, EVEN IF ADVISED OF THE
% POSSIBILITY OF SUCH DAMAGE.
```
