## Supplementary material for "Mechanistic description of spatial processes using integrative modelling of noise-corrupted imaging data": coordinateSearch.m

```
function [ x, fval, exitflag, output ] = coordinateSearch( fun, x0, lb, ub, options )
% performs a simple coordinate search with random update of search
% directions after a failure in improving the current value.
%
% Input:
% fun     : objective function to be minimized
% x0      : initial guess for parameters
% lb, ub  : bounds for parameters
% options : struct with options for the algorithm:
%   TolX              : tolerance of parameter
%   TolFun            : tolerance of objective function, currently not used
%   MaxFunEvals       : maximum number of evaluations of fun
%   MaxIter           : maximum number of iterations
%   OutputFcn         : for visual output after each iteration
%   Delta             : initial step size (rel. to 1)
%   ExpandFactor      : (default 3.5)
%   ContractFactor    : (default 0.35)
%   Barrier           : use barrier on bounds (default none)
%
% Output:
% x   : best guess for parameters
% fval: objective function at the solution, generally fval=fun(x)
% exitflag: 
%   1 : The function converged to a solution x
%   0 : Number of iterations exceeded options.MaxIter or number of function 
%       evaluations exceeded options.MaxFunEvals.
%   -1: The algorithm was terminated inappropriately
% output : struct with meta information:
%   iterations  : number of iterations
%   funcCount   : number of function evaluations
%   algorithm   : name of the algorithm
%   t_cpu       : cpu time
%
% History:
% 2017/09/27 Yannik Schaelte

    dim = length(x0);
    
    % have to stay below max values
    jIter = 0;
    funcCount = 0;
    
    [tolX,tolFun,maxIter,maxFunEvals,outputFcn,delta,expandFactor,contractFactor,barrier] = f_extractOptions(options,dim);
    if (isa(outputFcn,'function_handle'))
        visual = true;
    else
        visual = false;
    end
    
    x0 = x0(:);
    lb = lb(:);
    ub = ub(:);  
    normalize   = @(x) f_normalize(x,lb,ub);
    denormalize = @(y) f_denormalize(y,lb,ub); 
    y0      = normalize(x0);
    tolY    = tolX / norm(ub-lb);
    
    % wrap function to consider boundaries
    fun = @(y,jIter) f_wrap_fun(denormalize(y),fun,lb,ub,barrier,jIter,maxIter);
    
    % measure time
    starttime = cputime;
    
    % iteratively improved variables
    ybst = y0;
    fbst = fun(y0,jIter);
    funcCount = funcCount + 1;
    
    % search directions
    step = [eye(dim), -eye(dim)];
    
    if (visual)
        f_output(denormalize(ybst),fbst,jIter,'init',outputFcn); % create new figure and initialize
        f_output(denormalize(ybst),fbst,jIter,'iter',outputFcn); % first iteration with start point and jIter = 0
    end
    
    % update search directions if last iter was not successful
    iterSuccessful = true;
    % iterate cyclically over search directions to improve performance
    jSpinner = 1;
    jPrev    = 0;
    
    while ( delta > tolY && jIter <= maxIter && funcCount <= maxFunEvals )
        
        if (~iterSuccessful)
            U = f_createRandomOrthogonalMatrix(dim);
            step = [U,-U];
        end
    
        iterSuccessful = false;
        for j=1:2*dim
            ycur = ybst + delta*step(:,jSpinner);
            fcur = fun(ycur,jIter);
            funcCount = funcCount + 1;
            % simulate finite differences
%            surroundingValues = zeros(2*dim,1);
            if (fcur < fbst)
                ybst = ycur;
                fbst = fcur;
                
                % alternatively: expand always
                if (jSpinner == jPrev), delta = expandFactor * delta; end
                jPrev = jSpinner;
                
                iterSuccessful = true;
                break;
            end
            
%            surroundingValues(jSpinner,1) = fcur;
            
            % update coordinate index
            if jSpinner == 2*dim
                jSpinner = 1;
            else
                jSpinner = jSpinner + 1;
            end
        end
        
        % apparently bad effects
%         if (~iterSuccessful)
%             finiteDifferences = zeros(dim,1);
%             for j=1:dim
%                 finiteDifferences(j) = (surroundingValues(j)-surroundingValues(dim+j))/(2*delta);
%             end
%             U = step(:,1:dim);
%             finiteDifferences = transpose(transpose(finiteDifferences)/U);
%             ycur = ybst - delta*finiteDifferences/norm(finiteDifferences);
%             fcur = fun(ycur);
%             funcCount = funcCount + 1;
%             if (fcur < fbst)
%                 ybst = ycur;
%                 fbst = fcur;
%                 
%                 %iterSuccessful = true;
%             end
%         end
        
        if (~iterSuccessful)
            delta = contractFactor * delta;
        end
        
        jIter = jIter + 1;
        
        % update output
        if (visual)
            f_output(denormalize(ybst),fbst,jIter,'iter',outputFcn); 
        end
    
    end
    
    x    = denormalize(ybst);
    fval = fbst;
    
    if ( delta <= tolY )
        exitflag = 1;
    else
        % needed too long
        exitflag = 0;
    end
    
    output.funcCount    = funcCount;
    output.iterations   = jIter;
    output.algorithm    = 'Coordinate Search';
    output.t_cpu        = cputime - starttime;
    
    % finalize output
    if (visual)
        f_output(x,fbst,jIter,'done',outputFcn); 
    end

end

function y = f_normalize(x,lb,ub)
    y = (x-lb)./abs(ub-lb);
end

function x = f_denormalize(y,lb,ub)
    x = y.*(ub-lb) + lb;
end

function [tolX,tolFun,maxIter,maxFunEvals,outputFcn,delta,expandFactor,contractFactor,barrier] = f_extractOptions(options,dim)
% interpret options

    if (isfield(options,'TolX'))
        tolX    = options.TolX;
    else
        tolX    = 1e-6;
    end
    
    if (isfield(options,'TolFun'))
        tolFun  = options.TolFun;
    else
        tolFun  = 1e-6;
    end
    
    if (isfield(options,'MaxIter'))
        maxIter = options.MaxIter;
    else
        maxIter = 200*dim;
    end
    
    if (isfield(options,'MaxFunEvals'))
        maxFunEvals = options.MaxFunEvals;
    else
        maxFunEvals = 400*dim;
    end
    
    if (isfield(options,'OutputFcn'))
        outputFcn = options.OutputFcn;
    else
        outputFcn = nan;
    end
    
    if (isfield(options,'Delta'))
        delta          = options.Delta; % mesh size
    else
        delta          = 0.05;
    end
    
    if (isfield(options,'ExpandFactor'))
        expandFactor   = options.ExpandFactor;
    else
        expandFactor   = 3.5;
    end
    
    if (isfield(options,'ContractFactor'))
        contractFactor = options.ContractFactor;
    else
        contractFactor = 0.35;
    end
    
    if (isfield(options,'Barrier'))
        barrier        = options.Barrier;
    else
        barrier        = '';
    end
    
end

function fval = f_wrap_fun(x,fun,lb,ub,barrier,jIter,maxIter)
% set fun to inf whenever conditions not fulfilled
    if (~isequal(barrier,''))
        fval = fun(x);
        fval = barrierFunction(fval, [], x, [lb, ub], jIter, maxIter, barrier);
    else
        if (any(x>ub) || any(x<lb))
            fval = inf;
        else
            fval = fun(x);
        end        
    end
end

function U = f_createRandomOrthogonalMatrix(dim)
    M = randn(dim,dim);
    [Q,R] = qr(M);
    D = diag(R);
    D = diag(D)./abs(D);
    U = Q * D;
end

function f_output(x,fval,iter,state,outputFcn)
% short for call to output function
    optimValues.fval = fval;
    optimValues.iteration = iter;
    outputFcn(x,optimValues,state);
end
```
