## Supplementary material for "Mechanistic description of spatial processes using integrative modelling of noise-corrupted imaging data": direct.m

```
function [ret_minval,final_xatmin,history] = direct...
   (Problem,bounds,opts,varargin)
% Function   : Direct Version 4.0
% Written by : Dan Finkel
% Created on : 01/27/2003
% Last Update: 06/21/2004
% Purpose    : Direct optimization algorithm.
%
% This code comes with no guarantee or warranty of any kind
%
%  If unfamiliar with DIRECT, user is recommended to manual
%  that accompanies this code, and can be found at:
%       www4.ncsu.edu/~definkel/research/index.html
%
%
%          [x,fmin,history] = Direct(Problem,bounds,opts,varargin)
%
% Parameters:
%         IN: Problem   - Structure containing problem
%                    Problem.f              = Objective function handle
%
%                    NOTE: If you problem has no constraints (other than
%                          those on the bounds, this is the only field you
%                          need to add to Problem
%
%                    Problem.numconstraints = number of constraints
%                    Problem.constraint(i).func    = i-th constraint handle
%                    Problem.constraint(i).penalty = penalty parameter for
%                                                    i-th constraint
%                    Note: If f returns objective function AND constraints
%                          then set Problem.constraint(1).func = Problem.f
%             bounds    - an n x 2 vector of the lower and upper bounds.
%                         The first column is the lower bounds, and the second
%                         column contains the upper bounds
%             opts      - (optional) MATLAB structure.
%                    opts.ep        = Jones factor                      (default is 1e-4)
%                    opts.maxevals  = max. number of function evals     (default is 20)
%                    opts.maxits    = max. number of iterations         (default is 10)
%                    opts.maxdeep   = max. number of rect. divisions    (default is 100)
%                    opts.testflag  = 1 if globalmin known, 0 otherwise (default is 0)
%                    opts.showits   = 1 if disp. stats shown, 0 oth.
%                                      (default is 1)
%                    opts.globalmin = globalmin (if known)
%                                      (default is 0)
%                    opts.tol       = tolerance for term. if tflag=1
%                                      (default is 0.01)
%                    opts.impcons   = turns on implicit constraint capability
%                                      (default is 0)
%                                     If set to one, objective function
%                                     is expected to return a flag which represents
%                                     the feasibility of the point sampled
%             varargin  - (optional) additional arguements to be passed to
%                         objective function
%
%                NOTE: If opts.tflag == 0, maxevals, maxevals and maxdeep are ignored.
%                      DIRECT will stop when the absolute error is less
%                      than tol. Also, preallocation will not occur, and the algorithm
%                      can run slower than if opts.tflag == 1
%                NOTE: opts.maxevals is an approximate stopping condition. DIRECT will
%                      exceed this budget by a slight amount
%
%        OUT: minval    -  minimum value found
%             xatmin    - (optional) location of minimal value
%             history      - (optional) array of iteration historyory, useful for tables and plots
%                The three columns are iteration, fcn evals, and min value found.
%
%                Direct may be called by
%
%                minval = Direct(Problem,bounds);
%                          or
%                with any variation of the optional arguments

%------------------------------------------------------------------%
%
% Implementation taken from:
% D.R. Jones, C.D. Perttunen, and B.E. Stuckman. "Lipschitzian
% Optimization Without the Lipschitz Constant". Journal of
% Optimization Theory and Application, 79(1):157-181, October 1993
%
%------------------------------------------------------------------%

%-- Initialize the variables --------------------------------------%
lengths = [];c = [];fc = [];
con = [];szes = [];feas_flags=[];
om_lower     = bounds(:,1);
om_upper     = bounds(:,2);
fcncounter   = 0;
perror       = 0;
itctr        = 1;
done         = 0;
g_nargout    = nargout;
n            = size(bounds,1);

% Determine option values
if nargin<3, opts=[]; end
if (nargin>=3) & (length(opts)==0), opts=[]; end
getopts(opts, ...
 'maxits',     20,...         % maximum of iterations
 'maxevals',   10,...         % maximum # of function evaluations
 'maxdeep',    100,...        % maximum number of side divisions
 'testflag',   0,...          % terminate if within a relative tolerence of f_opt
 'globalmin',  0,...          % minimum value of function
 'ep',         1e-4,...       % global/local weight parameter.
 'tol',        0.01,...       % allowable relative error if f_reach is set
 'showits',    1,...          % print iteration stats
 'impcons',    0,...          % flag for using implicit constraint handling
 'pert',       1e-6);         % pertubation for implicit constraint handling
% 'maxflag',   0, ...         % set to 1 for max problems, 0 for min problems
% 'sizeconst', 0.5,...        % constant on rectangle size function
% 'distance',  1,...          % 1/0 for distance/volume measure of size
% 'minlength', 1e-4,...       % stop if best rectangle has all sides 1ess than this
% 'minevals',  0,...          % but must evaluate at least this many points

theglobalmin = globalmin;
tflag        = testflag;

%-- New 06/08/2004 Pre-allocate memory for storage vectors
if tflag == 0
    lengths    = zeros(n,maxevals + floor(.10*maxevals));
    c          = lengths;
    fc         = zeros(1,maxevals + floor(.10*maxevals));
    szes       = fc;
    con        = fc;
    feas_flags = fc;
end

%-- Call DIRini ---------------------------------------------------%
[thirds , lengths, c , fc, con, feas_flags minval,xatmin,perror,...
        history,szes,fcncounter,calltype] =...
        DIRini(Problem,n,bounds(:,1),bounds(:,2),...
        lengths,c,fc,con, feas_flags, szes,...
        theglobalmin,maxdeep,tflag,g_nargout, impcons, varargin{:});

ret_minval = minval;
ret_xatmin = xatmin;
%-- MAIN LOOP -----------------------------------------------------%
minval = fc(1) + con(1);
while perror > tol
   %-- Create list S of potentially optimal hyper-rectangles
   S = find_po(fc(1:fcncounter)+con(1:fcncounter),...
       lengths(:,1:fcncounter),minval,ep,szes(1:fcncounter));

   %-- Loop through the potentially optimal hrectangles -----------%
   %-- and divide -------------------------------------------------%
   for i = 1:size(S,2)
      [lengths,fc,c,con,feas_flags,szes,fcncounter,success] = ...
          DIRdivide(bounds(:,1),bounds(:,2),Problem,S(1,i),thirds,lengths,...
          fc,c,con,feas_flags,fcncounter,szes,impcons,calltype,varargin{:});
   end

   %-- update minval, xatmin --------------------------------------%
   [minval,fminindex] =  min(fc(1:fcncounter)+con(1:fcncounter));
   penminval = minval + con(fminindex);
   xatmin = (om_upper - om_lower).*c(:,fminindex) + om_lower;
   if (con(fminindex) > 0)|(feas_flags(fminindex) ~= 0)
       %--- new minval is infeasible, don't do anything
   else
       %--- update return values
       ret_minval = minval;
       ret_xatmin = xatmin;
   end

   %--see if we are done ------------------------------------------%
   if tflag == 1
      %-- Calculate error if globalmin known
      if theglobalmin ~= 0
          perror = 100*(minval - theglobalmin)/abs(theglobalmin);
      else
          perror = 100*minval;
      end
   else
      %-- Have we exceeded the maxits?
      if itctr >= maxits
         disp('Exceeded max iterations. Increase maxits')
         done = 1;
      end
      %-- Have we exceeded the maxevals?
      if fcncounter > maxevals
         disp('Exceeded max fcn evals. Increase maxevals')
         done = 1;
      end
      if done == 1
         perror = -1;
      end
   end
   if max(max(lengths)) >= maxdeep
      %-- We've exceeded the max depth
      disp('Exceeded Max depth. Increse maxdeep')
      perror = -1;
   end
   if g_nargout == 3
      %-- Store History
      maxhist = size(history,1);
      history(maxhist+1,1) = itctr;
      history(maxhist+1,2) = fcncounter;
      history(maxhist+1,3) = minval;
  end

  %-- New, 06/09/2004
  %-- Call replaceinf if impcons flag is set to 1
  if impcons == 1
      fc = replaceinf(lengths(:,1:fcncounter),c(:,1:fcncounter),...
          fc(1:fcncounter),con(1:fcncounter),...
          feas_flags(1:fcncounter),pert);
  end

  %-- show iteration stats
  if showits == 1
    if  (con(fminindex) > 0) | (feas_flags(fminindex) == 1)
        fprintf('Iter: %4i   f_min: %15.10f*    fn evals: %8i\n',...
         itctr,minval,fcncounter);
    else
        fprintf('Iter: %4i   f_min: %15.10f    fn evals: %8i\n',...
         itctr,minval,fcncounter);
    end
  end
  itctr  = itctr + 1;
end

%-- Return values
if g_nargout == 2
    %-- return x*
    final_xatmin = ret_xatmin;
elseif g_nargout == 3
    %-- return x*
    final_xatmin = ret_xatmin;

    %-- chop off 1st row of history
    history(1:size(history,1)-1,:) = history(2:size(history,1),:);
    history = history(1:size(history,1)-1,:);
end
return
%------------------------------------------------------------------%
% Function:   DIRini                                               %
% Written by: Dan Finkel                                           %
% Created on: 10/19/2002                                           %
% Purpose   : Initialization of Direct                             %
%             to eliminate storing floating points                 %
%------------------------------------------------------------------%
function [l_thirds,l_lengths,l_c,l_fc,l_con, l_feas_flags, minval,xatmin,perror,...
        history,szes,fcncounter,calltype] = DIRini(Problem,n,a,b,...
        p_lengths,p_c,p_fc,p_con, p_feas_flags, p_szes,theglobalmin,...
        maxdeep,tflag,g_nargout,impcons,varargin)

l_lengths    = p_lengths;
l_c          = p_c;
l_fc         = p_fc;
l_con        = p_con;
l_feas_flags = p_feas_flags;
szes         = p_szes;


%-- start by calculating the thirds array
%-- here we precalculate (1/3)^i which we will use frequently
l_thirds(1) = 1/3;
for i = 2:maxdeep
   l_thirds(i) = (1/3)*l_thirds(i-1);
end

%-- length array will store # of slices in each dimension for
%-- each rectangle. dimension will be rows; each rectangle
%-- will be a column

%-- first rectangle is the whole unit hyperrectangle
l_lengths(:,1) = zeros(n,1);

%01/21/04 HACK
%-- store size of hyperrectangle in vector szes
szes(1,1) = 1;

%-- first element of c is the center of the unit hyperrectangle
l_c(:,1) = ones(n,1)/2;

%-- Determine if there are constraints
calltype = DetermineFcnType(Problem,impcons);

%-- first element of f is going to be the function evaluated
%-- at the center of the unit hyper-rectangle.
%om_point   = abs(b - a).*l_c(:,1)+ a;
%l_fc(1)    = feval(f,om_point,varargin{:});
[l_fc(1),l_con(1), l_feas_flags(1)] = ...
    CallObjFcn(Problem,l_c(:,1),a,b,impcons,calltype,varargin{:});
fcncounter = 1;


%-- initialize minval and xatmin to be center of hyper-rectangle
xatmin = l_c(:,1);
minval   = l_fc(1);
if tflag == 1
    if theglobalmin ~= 0
        perror = 100*(minval - theglobalmin)/abs(theglobalmin);
    else
        perror = 100*minval;
    end
else
   perror = 2;
end

%-- initialize history
%if g_nargout == 3
    history(1,1) = 0;
    history(1,2) = 0;
    history(1,3) = 0;
%end
%------------------------------------------------------------------%
% Function   :  find_po                                            %
% Written by :  Dan Finkel                                         %
% Created on :  10/19/2002                                         %
% Purpose    :  Return list of PO hyperrectangles                  %
%------------------------------------------------------------------%
function rects = find_po(fc,lengths,minval,ep,szes)

%-- 1. Find all rects on hub
diff_szes = sum(lengths,1);
tmp_max = max(diff_szes);
j=1;
sum_lengths = sum(lengths,1);
for i =1:tmp_max+1
    tmp_idx = find(sum_lengths==i-1);
    [tmp_n, hullidx] = min(fc(tmp_idx));
    if length(hullidx) > 0
        hull(j) = tmp_idx(hullidx);
        j=j+1;
        %-- 1.5 Check for ties
        ties = find(abs(fc(tmp_idx)-tmp_n) <= 1e-13);
        if length(ties) > 1
            mod_ties = find(tmp_idx(ties) ~= hull(j-1));
            hull = [hull tmp_idx(ties(mod_ties))];
            j = length(hull)+1;
        end
    end
end
%-- 2. Compute lb and ub for rects on hub
lbound = calc_lbound(lengths,fc,hull,szes);
ubound = calc_ubound(lengths,fc,hull,szes);

%-- 3. Find indeces of hull who satisfy
%--    1st condition
maybe_po = find(lbound-ubound <= 0);

%-- 4. Find indeces of hull who satisfy
%--    2nd condition
t_len  = length(hull(maybe_po));
if minval ~= 0
    po = find((minval-fc(hull(maybe_po)))./abs(minval) +...
        szes(hull(maybe_po)).*ubound(maybe_po)./abs(minval) >= ep);
else
    po = find(fc(hull(maybe_po)) -...
        szes(hull(maybe_po)).*ubound(maybe_po) <= 0);
end
final_pos      = hull(maybe_po(po));

rects = [final_pos;szes(final_pos)];
return
%------------------------------------------------------------------%
% Function   :  calc_ubound                                        %
% Written by :  Dan Finkel                                         %
% Created on :  10/19/2002                                         %
% Purpose    :  calculate the ubound used in determing potentially %
%               optimal hrectangles                                %
%------------------------------------------------------------------%
function ub = calc_ubound(lengths,fc,hull,szes)

hull_length  = length(hull);
hull_lengths = lengths(:,hull);
for i =1:hull_length
    tmp_rects = find(sum(hull_lengths,1)<sum(lengths(:,hull(i))));
    if length(tmp_rects) > 0
        tmp_f     = fc(hull(tmp_rects));
        tmp_szes  = szes(hull(tmp_rects));
        tmp_ubs   = (tmp_f-fc(hull(i)))./(tmp_szes-szes(hull(i)));
        ub(i)        = min(tmp_ubs);
    else
        ub(i)=1.976e14;
    end
end
return
%------------------------------------------------------------------%
% Function   :  calc_lbound                                        %
% Written by :  Dan Finkel                                         %
% Created on :  10/19/2002                                         %
% Purpose    :  calculate the lbound used in determing potentially %
%               optimal hrectangles                                %
%------------------------------------------------------------------%
function lb = calc_lbound(lengths,fc,hull,szes)

hull_length  = length(hull);
hull_lengths = lengths(:,hull);
for i = 1:hull_length
    tmp_rects = find(sum(hull_lengths,1)>sum(lengths(:,hull(i))));
    if length(tmp_rects) > 0
        tmp_f     = fc(hull(tmp_rects));
        tmp_szes  = szes(hull(tmp_rects));
        tmp_lbs   = (fc(hull(i))-tmp_f)./(szes(hull(i))-tmp_szes);
        lb(i)     = max(tmp_lbs);
    else
        lb(i)     = -1.976e14;
    end
end
return
%------------------------------------------------------------------%
% Function   :  DIRdivide                                          %
% Written by :  Dan Finkel                                         %
% Created on :  10/19/2002                                         %
% Purpose    :  Divides rectangle i that is passed in              %
%------------------------------------------------------------------%
function [lengths,fc,c,con,feas_flags,szes,fcncounter,pass] = ...
    DIRdivide(a,b,Problem,index,thirds,p_lengths,p_fc,p_c,p_con,...
    p_feas_flags,p_fcncounter,p_szes,impcons,calltype,varargin)

lengths    = p_lengths;
fc         = p_fc;
c          = p_c;
szes       = p_szes;
fcncounter = p_fcncounter;
con        = p_con;
feas_flags = p_feas_flags;

%-- 1. Determine which sides are the largest
li     = lengths(:,index);
biggy  = min(li);
ls     = find(li==biggy);
lssize = length(ls);
j = 0;

%-- 2. Evaluate function in directions of biggest size
%--    to determine which direction to make divisions
oldc       = c(:,index);
delta      = thirds(biggy+1);
newc_left  = oldc(:,ones(1,lssize));
newc_right = oldc(:,ones(1,lssize));
f_left     = zeros(1,lssize);
f_right    = zeros(1,lssize);
for i = 1:lssize
    lsi               = ls(i);
    newc_left(lsi,i)  = newc_left(lsi,i) - delta;
    newc_right(lsi,i) = newc_right(lsi,i) + delta;
    [f_left(i), con_left(i), fflag_left(i)]    = CallObjFcn(Problem,newc_left(:,i),a,b,impcons,calltype,varargin{:});
    [f_right(i), con_right(i), fflag_right(i)] = CallObjFcn(Problem,newc_right(:,i),a,b,impcons,calltype,varargin{:});
    fcncounter = fcncounter + 2;
end
w = [min(f_left, f_right)' ls];

%-- 3. Sort w for division order
[V,order] = sort(w,1);

%-- 4. Make divisions in order specified by order
for i = 1:size(order,1)

   newleftindex  = p_fcncounter+2*(i-1)+1;
   newrightindex = p_fcncounter+2*(i-1)+2;
   %-- 4.1 create new rectangles identical to the old one
   oldrect = lengths(:,index);
   lengths(:,newleftindex)   = oldrect;
   lengths(:,newrightindex)  = oldrect;

   %-- old, and new rectangles have been sliced in order(i) direction
   lengths(ls(order(i,1)),newleftindex)  = lengths(ls(order(i,1)),index) + 1;
   lengths(ls(order(i,1)),newrightindex) = lengths(ls(order(i,1)),index) + 1;
   lengths(ls(order(i,1)),index)         = lengths(ls(order(i,1)),index) + 1;

   %-- add new columns to c
   c(:,newleftindex)  = newc_left(:,order(i));
   c(:,newrightindex) = newc_right(:,order(i));

   %-- add new values to fc
   fc(newleftindex)  = f_left(order(i));
   fc(newrightindex) = f_right(order(i));

   %-- add new values to con
   con(newleftindex)  = con_left(order(i));
   con(newrightindex) = con_right(order(i));

   %-- add new flag values to feas_flags
   feas_flags(newleftindex)  = fflag_left(order(i));
   feas_flags(newrightindex) = fflag_right(order(i));

   %-- 01/21/04 Dan Hack
   %-- store sizes of each rectangle
   szes(1,newleftindex)  = 1/2*norm((1/3*ones(size(lengths,1),1)).^(lengths(:,newleftindex)));
   szes(1,newrightindex) = 1/2*norm((1/3*ones(size(lengths,1),1)).^(lengths(:,newrightindex)));
end
szes(index) = 1/2*norm((1/3*ones(size(lengths,1),1)).^(lengths(:,index)));
pass = 1;

return
%------------------------------------------------------------------%
% Function   :  CallConstraints                                    %
% Written by :  Dan Finkel                                         %
% Created on :  06/07/2004                                         %
% Purpose    :  Evaluate Constraints at pointed specified          %
%------------------------------------------------------------------%
function ret_value = CallConstraints(Problem,x,a,b,varargin)

%-- Scale variable back to original space
point = abs(b - a).*x+ a;

ret_value = 0;
if isfield(Problem,'constraint')
    if ~isempty(Problem.constraint)
        for i = 1:Problem.numconstraints
            if length(Problem.constraint(i).func) == length(Problem.f)
                if double(Problem.constraint(i).func) == double(Problem.f)                    
                    %-- Dont call constraint; value was returned in obj fcn
                    con_value = 0;
                else
                    con_value = feval(Problem.constraint(i).func,point,varargin{:});
                end
            else
                con_value = feval(Problem.constraint(i).func,point,varargin{:});
            end
            if con_value > 0
                %-- Infeasible, punish with associated pen. param
                ret_value = ret_value + con_value*Problem.constraint(i).penalty;
            end
        end
    end
end
return
%------------------------------------------------------------------%
% Function   :  CallObjFcn                                         %
% Written by :  Dan Finkel                                         %
% Created on :  06/07/2004                                         %
% Purpose    :  Evaluate ObjFcn at pointed specified               %
%------------------------------------------------------------------%
function [fcn_value, con_value, feas_flag] = ...
    CallObjFcn(Problem,x,a,b,impcon,calltype,varargin)

con_value = 0;
feas_flag = 0;

%-- Scale variable back to original space
point = abs(b - a).*x+ a;

if calltype == 1
    %-- No constraints at all
    fcn_value = feval(Problem.f,point,varargin{:});
elseif calltype == 2
    %-- f returns all constraints
    [fcn_value, cons] = feval(Problem.f,point,varargin{:});
    for i = 1:length(cons)
        if cons > 0
            con_value = con_value + Problem.constraint(i).penalty*cons(i);
        end
    end
elseif calltype == 3
    %-- f returns no constraint values
    fcn_value = feval(Problem.f,point,varargin{:});
    con_value = CallConstraints(Problem,x,a,b,varargin{:});
elseif calltype == 4
    %-- f returns feas flag
    [fcn_value,feas_flag] = feval(Problem.f,point,varargin{:});
elseif calltype == 5
    %-- f returns feas flags, and there are constraints
    [fcn_value,feas_flag] = feval(Problem.f,point,varargin{:});
    con_value = CallConstraints(Problem,x,a,b,varargin{:});
end
if feas_flag == 1
    fcn_value = 10^9;
    con_value = 0;
end
return
%------------------------------------------------------------------%
% Function   :  replaceinf                                         %
% Written by :  Dan Finkel                                         %
% Created on :  06/09/2004                                         %
% Purpose    :  Assign R. Carter value to given point              %
%------------------------------------------------------------------%
function fcn_values = replaceinf(lengths,c,fc,con,flags,pert)

%-- Initialize fcn_values to original values
fcn_values = fc;

%-- Find the infeasible points
infeas_points = find(flags == 1);

%-- Find the feasible points
feas_points   = find(flags == 0);

%-- Calculate the max. value found so far
if ~isempty(feas_points)
    maxfc = max(fc(feas_points) + con(feas_points));
else
    maxfc = max(fc + con);
end

for i = 1:length(infeas_points)
    if isempty(feas_points)
        %-- no feasible points found yet
        found_points = [];found_pointsf = [];
        index = infeas_points(i);
    else
        index = infeas_points(i);

        %-- Initialize found points to be entire set
        found_points  = c(:,feas_points);
        found_pointsf = fc(feas_points) + con(feas_points);

        %-- Loop through each dimension, and find points who are close enough
        for j = 1:size(lengths,1)
            neighbors = find(abs(found_points(j,:) - c(j,index)) <= ...
                3^(-lengths(j,index)));
            if ~isempty(neighbors)
                found_points  = found_points(:,neighbors);
                found_pointsf = found_pointsf(neighbors);
            else
                found_points = [];found_pointsf = [];
                break;
            end
        end
    end

    %-- Assign Carter value to the point
    if ~isempty(found_pointsf)
        %-- assign to index the min. value found + a little bit more
        fstar = min(found_pointsf);
        if fstar ~= 0
            fcn_values(index) = fstar + pert*abs(fstar);
        else
            fcn_values(index) = fstar + pert*1;
        end
    else
        fcn_values(index) = maxfc+1;
        maxfc             = maxfc+1;
    end
end
return
%------------------------------------------------------------------%
% Function   :  DetermineFcnType                                   %
% Written by :  Dan Finkel                                         %
% Created on :  06/25/2004                                         %
% Purpose    :  Determine how constraints are handled              %
%------------------------------------------------------------------%
function retval = DetermineFcnType(Problem,impcons)

retval = 0;
if (~isfield(Problem,'constraint'))&(~impcons)
    %-- No constraints at all
    retval = 1;
end
if isfield(Problem,'constraint')
    %-- There are explicit constraints. Next determine where
    %-- they are called
    if ~isempty(Problem.constraint)
        if length(Problem.constraint(1).func) == length(Problem.f)
            %-- Constraint values may be returned from objective
            %-- function. Investigate further
            if double(Problem.constraint(1).func) == double(Problem.f)
                %-- f returns constraint values
                retval = 2;
            else
                %-- f does not return constraint values
                retval = 3;
            end
        else
            %-- f does not return constraint values
            retval = 3;
        end
    else
        if impcons
            retval = 0;
        else
            retval = 1;
        end
    end
end

if (impcons)
    if ~retval
        %-- only implicit constraints
        retval = 4;
    else
        %-- both types of constraints
        retval = 5;
    end
end
%------------------------------------------------------------------%
% GETOPTS Returns options values in an options structure
% USAGE
%   [value1,value2,...]=getopts(options,field1,default1,field2,default2,...)
% INPUTS
%   options  : a structure variable
%   field    : a field name
%   default  : a default value
% OUTPUTS
%   value    : value in the options field (if it exists) or the default value
%
% Variables with the field names will be created in the caller's workspace
% and set to the value in the option variables field (if it exists) or to the
% default value.
%
% Example called from a function:
%   getopts(options,'tol',1e-8,'maxits',100);
% where options contains the single field 'tol' with value equal to 1
% The function have two variable defined in the local workspace, tol with a
% value of 1 and maxits with a value of 100.
%
% If options contains a field name not in the list passed to getopts, a
% warning is issued.
%
%
% Many thanks to the author of this function,
% Paul Fackler
%
%
%------------------------------------------------------------------%
function varargout=getopts(options,varargin)
K=fix(nargin/2);
if nargin/2==K
  error('fields and default values must come in pairs')
end
if isa(options,'struct'), optstruct=1; else, optstruct=0; end
varargout=cell(K,1);
k=0;
ii=1;
for i=1:K
  if optstruct & isfield(options,varargin{ii})
    assignin('caller',varargin{ii},getfield(options,varargin{ii}));
    k=k+1;
  else
    assignin('caller',varargin{ii},varargin{ii+1});
  end
  ii=ii+2;
end

if optstruct & k~=size(fieldnames(options),1)
  warning('options variable contains improper fields')
end

return
%------------------------------------------------------------------%
% Versions  : 1.0 - 1st successful implemenation of DIRect
%           : 2.0 - Removed floating point arithmetic
%                   duplicated Table 5 of Jones et al.
%           : 2.1 - increased speed by storing size calcs.
%           : 2.2 - utitilized linked lists to increase speed
%           : 2.3 - rewrote ubound to increase speed
%           : 2.4 - rewrote lbound to increase speed
%           : 2.5 - removed call to calcsize
%           : 2.6 - added check_for_ties
%           : 2.7 - rewrote check_for_ties to compare fp correctly
%           : 2.8 - changed output arguments, rewrote help
%           : 3.0 - simplified input/output. Put on web.
%           : 3.1 - Performanced Tuned! Tremendous speed increase
%           : 3.2 - Removed llists; performance tuned
%                   Many thanks to Ray Muzic and Paul Fackler
%                   for their suggestions to improve this code
%           : 4.0 - Sped up code, and added 2 constraint handling
%                   mechanisms.
%------------------------------------------------------------------%
```
