## Supplementary material for "Mechanistic description of spatial processes using integrative modelling of noise-corrupted imaging data": dynamicHillClimb.m

```
function [x, fval, exitflag, output] = dynamicHillClimb(fun,x0,lb,ub,options)
% dynamic hill climbing algorithm, adapted from [DeLaMaza+Yuret, Dynamic
% Hill Climbing].
%
% Input:
% fun     : objective function to be minimized
% x0      : initial guess for parameters
% lb, ub  : bounds for parameters, no value should be inf and the
% difference ub-lb in no component be < 0
% options : struct with options for the algorithm:
%   TolX              : tolerance of parameter
%   TolFun            : tolerance of objective function
%   MaxFunEvals       : maximum number of evaluations of fun
%   MaxIter           : maximum number of iterations
%   OutputFcn         : for visual output after each iteration
%   InitialStepSize   : rel. to 1 (default 0.1)
%   ExpandFactor      : (default 2)
%   ContractFactor    : (default 0.45)
%   StuckSearchFactor : how far to expand again after got stuck (default 4)
%   Barrier           : use barrier on bounds (default extreme barrier)
%   Display           : off|iter|iter-detailed, text output (default off)
%   Mode              : 1|2, algorithm mode to use (default 1)
%
% Output:
% x   : best guess for parameters
% fval: objective function at the solution, generally fval=fun(x)
% exitflag:
%   1 : The function converged to a solution x
%   0 : Number of iterations exceeded options.MaxIter or number of function
%       evaluations exceeded options.MaxFunEvals.
%   -1: The algorithm was terminated inappropriately
% output : struct with meta information:
%   iterations  : number of iterations
%   funcCount   : number of function evaluations
%   algorithm   : name of the algorithm
%   t_cpu       : cpu time
%
% History:
% 2017/09/27 Yannik Schaelte

% number of variables
dim  = length(x0); % number of variables

% interpret options
[tolX,tolFun,maxFunEvals,maxIter,outputFcn,...
    initialStepSize,expandFactor,contractFactor,stuckSearchFactor,barrier,...
    display,mode]...
    = f_extractOptions(options,dim);
if (isa(outputFcn,'function_handle'))
    visual = true;
else
    visual = false;
end

switch mode
    case 1
        dhc_mode_1();
    case 2
        dhc_mode_2();
    case 3
        dhc_mode_3();
    otherwise
        dhc_mode_1();
end

%% dhc_mode_1
    function dhc_mode_1()
        
        % create column vectors
        lb      = lb(:);
        ub      = ub(:);
        x0      = x0(:);
        normalize   = @(x) f_normalize(x,lb,ub);
        denormalize = @(y) f_denormalize(y,lb,ub);
        y0      = normalize(x0);
        tolY    = tolX / norm(ub-lb);
        % max step size
        vmax = initialStepSize * ones(dim,1);
        
        % wrap function to consider boundaries
        fun = @(y,jIter) f_wrap_fun(denormalize(y),fun,lb,ub,barrier,jIter,maxIter);
        
        % init run variables
        smax   = [vmax;-1];        % array of max step sizes, extra value for extra vector
        [step,norms]   = f_init_step(vmax); % matrix of step vectors
        xstep  = step;             % steps before last motion
        xnorms = norms;
        gradv  = zeros(dim,1);     % gradient vector
        gradi  = -1;               % index of gradient vector, -1 indicates gradv is not set
        lastj  = -1;               % index of last step taken
        stuck  = false;            % is process stuck (in min/max)? set when step sizes are small
        % then step vectors are increased
        done   = false;            % is some finishing criterion fulfilled?
        
        nVec   = 2*dim + 2;            % maximum index in step matrix
        opp_j  = @(j) f_opp_j(j,nVec); % short for opposite index in step matrix
        
        % init meta variables
        jIter     = 0;         % number of iterations, should be <= maxIter
        exitflag  = -1;        % flag indicating exit reason
        starttime = cputime;   % to measure time difference
        funEvals  = 0;         % function evaluations
        
        % init x, fval
        ybst      = y0;
        fbst      = fun(ybst,jIter);
        funEvals  = funEvals + 1;
        
        if (visual)
            f_output(denormalize(ybst),fbst,jIter,'init',outputFcn); % create new figure and initialize
            f_output(denormalize(ybst),fbst,jIter,'iter',outputFcn); % first iteration with start point and jIter = 0
        end
        
        while (~done)
            % increase counter
            jIter = jIter + 1;
            
            if (stuck)
                % choose the smallest step, if any is smaller than the maximum size
                [v,j] = f_min(step,norms,smax,stuckSearchFactor*tolY);
            else
                % choose the largest step
                [v,j] = f_max(step,norms);
            end
            
            f_display(display,jIter,fbst,norm(v));
            
            % j == -1 indicates minimum found
            if ( j ~= -1 && jIter <= maxIter && funEvals <= maxFunEvals )
                % compute next x, fval
                ycur = ybst + v;
                fcur = fun(ycur,jIter);
                funEvals = funEvals + 1;
                
                % is better estimate?
                delta_f = fcur - fbst;
                if ( delta_f < 0 && (norm(v)>tolX/stuckSearchFactor || abs(delta_f) > tolFun) )
                    stuck = false; % we are not stuck somewhere (anymore)
                    % update x, fval
                    ybst = ycur;
                    fbst = fcur;
                    % contract opp step to not try the previous point next
                    step(:,opp_j(j)) = -contractFactor*v;
                    norms(opp_j(j)) = norm(step(:,opp_j(j)));
                    % TODO do? no.
                    % step(:,j) = expandFactor*step(:,j);
                    % if last step repeated, expand the current step
                    if ( j == lastj )
                        v = expandFactor*v;
                    end
                    % xstep always contains the steps of the last time we moved
                    xstep = step;
                    xnorms = norms;
                    % record the last step
                    lastj = j;
                    % record step
                    step(:,j) = v;
                    norms(j) = norm(v);
                    
                    % update gradient vector
                    if (gradi == -1)
                        % if gradv empty, set gradv to current step and record index
                        gradv = v;
                        gradi = min([j, opp_j(j)]);
                    elseif (gradi == min([j, opp_j(j)]))
                        % if gradv is parallel to current step, add the current vector
                        gradv = (gradv + v);
                    else
                        % else update the extra vector
                        step(:,dim+1) = gradv + v;
                        norms(dim+1) = norm(gradv + v);
                        % set extra entry in smax to max of current step smax and gradv smax
                        % (bound for norm of gradient)
                        smax(dim+1)   = max([smax(min([j,opp_j(j)])),smax(gradi)]);
                        % set the opp step to - extra step
                        step(:,dim+2) = -contractFactor*step(:,dim+1);
                        norms(dim+2) = abs(contractFactor)*norms(dim+1);
                        % update gradv
                        % TODO do every round? mixed results
                        %gradv         = step(:,j);
                        gradi =-1;%        = min([j, opp_j(j)]);
                    end
                    % else: if already stuck, increase the current step size
                else
                    if (stuck)
                        step(:,j) = expandFactor*v;
                        norms(j) = norm(step(:,j));
                        % else: if current step norm >= tolX, decrease the current step size
                    elseif ( norm(step(:,j)) >= tolY )
                        step(:,j) = contractFactor*v;
                        norms(j) = norm(step(:,j));
                        % else: (norm < tolX), set the stuck flag and set all steps to
                        % expandFactor times the last recorded steps
                    else
                        stuck = true;
                        step = expandFactor * xstep;
                        norms = abs(expandFactor)*xnorms;
                    end
                end
                
                % update output
                if (visual)
                    f_output(denormalize(ybst),fbst,jIter,'iter',outputFcn);
                end
                
            else % somehow done
                done = true;
                if (jIter <= maxIter && funEvals <= maxFunEvals)
                    % found a local minimum
                    exitflag = 1;
                else
                    % needed too long
                    exitflag = 0;
                end
            end
            
        end
        
        % assign return values
        x                   = denormalize(ybst);
        fval                = fbst;
        output.funcCount    = funEvals;
        output.iterations   = jIter;
        output.algorithm    = 'Dynamic Hill Climb';
        output.t_cpu        = cputime - starttime;
        
        % finalize output
        if (visual)
            f_output(denormalize(ybst),fbst,jIter,'done',outputFcn);
        end
    end

%% dhc_mode_2
    function dhc_mode_2()
        
        % create column vectors
        lb      = lb(:);
        ub      = ub(:);
        x0      = x0(:);
        normalize   = @(x) f_normalize(x,lb,ub);
        denormalize = @(y) f_denormalize(y,lb,ub);
        y0      = normalize(x0);
        tolY    = tolX / norm(ub-lb);
        % max step size
        vmax = initialStepSize * ones(dim,1);
        
        % wrap function to consider boundaries
        fun = @(y,jIter) f_wrap_fun(denormalize(y),fun,lb,ub,barrier,jIter,maxIter);
        
        % init run variables
        smax   = [vmax;-1;-1];     % array of max step sizes, extra value for extra vector
        [step,norms] = f_init_step_2(vmax); % matrix of step vectors
        xstep  = step;             % steps before last motion
        xnorms = norms;
        gradv  = zeros(dim,1);     % gradient vector
        gradi  = -1;               % index of gradient vector, -1 indicates gradv is not set
        lastj  = -1;               % index of last step taken
        stuck  = false;            % is process stuck (in min/max)? set when step sizes are small
        % then step vectors are increased
        done   = false;            % is some finishing criterion fulfilled?
        
        nVec   = 2*dim + 4;            % maximum index in step matrix
        opp_j  = @(j) f_opp_j(j,nVec); % short for opposite index in step matrix
        
        % init meta variables
        jIter     = 0;         % number of iterations, should be <= maxIter
        exitflag  = -1;        % flag indicating exit reason
        starttime = cputime;   % to measure time difference
        funEvals  = 0;         % function evaluations
        
        % init x, fval
        ybst      = y0;
        fbst      = fun(ybst,jIter);
        funEvals  = funEvals + 1;
        
        % create cache to remember last values
        cacheSizeMin = 4;
        cacheSizeMax = max([8,dim]); % dim?
        % arr_t remembers last step lengths
        cache_stepsize = initialStepSize;
        % arr_y remembers last positions
        cache_y = ybst;
        numSuccessfulSteps = 0;
        numCacheMaximal = 0;
        numCacheWorked = 0;
        numGradMaximal = 0;
        numGradWorked = 0;
        
        if (visual)
            % create new figure and initialize
            f_output(denormalize(ybst),fbst,jIter,'init',outputFcn);
            % first iteration with start point and jIter = 0
            f_output(denormalize(ybst),fbst,jIter,'iter',outputFcn);
        end
        
        while (~done)
            % increase counter
            jIter = jIter + 1;
            
            if (stuck)
                % choose the smallest step, if any is smaller than the maximum size
                [v,j] = f_min(step,norms,smax,stuckSearchFactor*tolY);
            else
                % choose the largest step
                [v,j] = f_max(step,norms);
            end
            
            if(j == dim+2 || j == dim+3), numCacheMaximal = numCacheMaximal + 1; end
            if(j == dim+1 || j == dim+4), numGradMaximal = numGradMaximal + 1; end
            
            f_display(display,jIter,fbst,norm(v));
            
            % j == -1 indicates minimum found
            if ( j ~= -1 && jIter <= maxIter && funEvals <= maxFunEvals )
                % compute next x, fval
                ycur = ybst + v;
                fcur = fun(ycur,jIter);
                funEvals = funEvals + 1;
                
                % is better estimate?
                delta_f = fcur - fbst;
                if ( delta_f < 0 && (norm(v)>tolX/stuckSearchFactor || abs(delta_f) > tolFun) )
                    numSuccessfulSteps = numSuccessfulSteps + 1;
                    if (j == dim+2 || j == dim+3), numCacheWorked = numCacheWorked + 1; end
                    if (j == dim+1 || j == dim+4), numGradWorked = numGradWorked + 1; end
                    stuck = false; % we are stuck nowhere (anymore)
                    % update x, fval
                    ybst = ycur;
                    fbst = fcur;
                    % contract opp step to not try the previous point next
                    step(:,opp_j(j)) = -contractFactor*v;
                    norms(opp_j(j)) = norm(step(:,opp_j(j)));
                    % TODO do? no.
                    % step(:,j) = expandFactor*step(:,j);
                    % if last step repeated, expand the current step
                    if ( j == lastj )
                        v = expandFactor*v;
                    end
                    % xstep always contains the steps of the last time we moved
                    xstep = step;
                    xnorms = norms;
                    % record the last step
                    lastj = j;
                    % record step
                    step(:,j) = v;
                    norms(j) = norm(v);
                    
                    % update gradient vector
                    if (gradi == -1)
                        % if gradv empty, set gradv to current step and record index
                        gradv = v;
                        gradi = min([j, opp_j(j)]);
                    elseif (gradi == min([j, opp_j(j)]))
                        % if gradv is parallel to current step, add the current vector
                        gradv = (gradv + v);
                    else
                        % else update the extra vector
                        step(:,dim+1) = gradv + v;
                        norms(dim+1) = norm(gradv + v);
                        % set extra entry in smax to max of current step smax and gradv smax
                        % (bound for norm of gradient)
                        smax(dim+1)   = max([smax(min([j,opp_j(j)])),smax(gradi)]);
                        % set the opp step to - extra step
                        step(:,dim+4) = -contractFactor*step(:,dim+1);
                        norms(dim+4) = abs(contractFactor)*norms(dim+1);
                        % update gradv
                        % TODO every round?
                        %gradv         = step(:,j);
                        gradi =-1;%        = min([j, opp_j(j)]);
                    end
                    % update cache
                    if (length(cache_stepsize) >= cacheSizeMax)
                        cache_stepsize = [cache_stepsize(2:cacheSizeMax),cache_stepsize(cacheSizeMax)+norm(v)];
                        cache_y = [cache_y(:,2:cacheSizeMax),ybst];
                    else
                        cache_stepsize = [cache_stepsize,cache_stepsize(end)+norm(v)];
                        cache_y = [cache_y,ybst];
                    end
                    % and update interpolating gradient
                    nCache = length(cache_stepsize);
                    if (nCache >= cacheSizeMin)
                        % prepare data
                        ynew = zeros(dim,1);
                        % create vandermonde matrix
                        % TODO: re-center and expand vectors to avoid numerical errors
                        van = ones(nCache,3);
                        van(:,2) = cache_stepsize(:).^1;
                        van(:,1) = cache_stepsize(:).^2;
                        %                         van(:,1) = cache_stepsize(:).^3;
                        predPoint = cache_stepsize(end) + (cache_stepsize(end)-cache_stepsize(end-1));
                        for k=1:dim
                            y = transpose(cache_y(k,:));
                            p = van\y;
                            ynew(k) = polyval(p,predPoint);
                        end
                        vnew = ynew - ybst;
                        %                         vnew = vnew*norm(v)/norm(vnew);
                        step(:,dim+2) = vnew;
                        step(:,dim+3) = -contractFactor*vnew;
                        normvnew = norm(vnew);
                        norms(dim+2) = normvnew;
                        norms(dim+3) = abs(contractFactor)*normvnew;
                    end
                    % else: if already stuck, increase the current step size
                else
                    if (stuck)
                        step(:,j) = expandFactor*v;
                        norms(j) = norm(step(:,j));
                        % else: if current step norm >= tolX, decrease the current step size
                    elseif ( norm(step(:,j)) >= tolY )
                        step(:,j) = contractFactor*v;
                        norms(j) = norm(step(:,j));
                        % else: (norm < tolX), set the stuck flag and set all steps to
                        % expandFactor times the last recorded steps
                    else
                        stuck = true;
                        step = expandFactor * xstep;
                        norms = abs(expandFactor)*xnorms;
                    end
                end
                
                % update output
                if (visual)
                    f_output(denormalize(ybst),fbst,jIter,'iter',outputFcn);
                end
                
            else % somehow done
                done = true;
                if (jIter <= maxIter && funEvals <= maxFunEvals)
                    % found a local minimum
                    exitflag = 1;
                else
                    % needed too long
                    exitflag = 0;
                end
            end
            
        end
        
        % assign return values
        x                   = denormalize(ybst);
        fval                = fbst;
        output.funcCount    = funEvals;
        output.iterations   = jIter;
        output.algorithm    = 'Dynamic Hill Climb';
        output.t_cpu        = cputime - starttime;
        
        % finalize output
        if (visual)
            f_output(denormalize(ybst),fbst,jIter,'done',outputFcn);
        end
        numSuccessfulSteps
        numCacheMaximal
        numCacheWorked
        numGradMaximal
        numGradWorked
    end

%% dhc_mode_3
    function dhc_mode_3()
        
        % create column vectors
        lb      = lb(:);
        ub      = ub(:);
        x0      = x0(:);
        normalize   = @(x) f_normalize(x,lb,ub);
        denormalize = @(y) f_denormalize(y,lb,ub);
        y0      = normalize(x0);
        tolY    = tolX / norm(ub-lb);
        % max step size
        vmax = initialStepSize * ones(dim,1);
        
        % wrap function to consider boundaries
        fun = @(y,jIter) f_wrap_fun(denormalize(y),fun,lb,ub,barrier,jIter,maxIter);
        
        % init run variables
        smax   = [vmax;-1;-1];     % array of max step sizes, extra value for extra vector
        [step,norms] = f_init_step_2(vmax); % matrix of step vectors
        xstep  = step;             % steps before last motion
        xnorms = norms;
        gradv  = zeros(dim,1);     % gradient vector
        gradi  = -1;               % index of gradient vector, -1 indicates gradv is not set
        lastj  = -1;               % index of last step taken
        stuck  = false;            % is process stuck (in min/max)? set when step sizes are small
        % then step vectors are increased
        done   = false;            % is some finishing criterion fulfilled?
        
        nVec   = 2*dim + 4;            % maximum index in step matrix
        opp_j  = @(j) f_opp_j(j,nVec); % short for opposite index in step matrix
        
        % init meta variables
        jIter     = 0;         % number of iterations, should be <= maxIter
        exitflag  = -1;        % flag indicating exit reason
        starttime = cputime;   % to measure time difference
        funEvals  = 0;         % function evaluations
        
        % init x, fval
        ybst      = y0;
        fbst      = fun(ybst,jIter);
        funEvals  = funEvals + 1;
        
        % create cache to remember last values
        cacheSize = 5;%max([5,dim/8]); % dim?
        % create vandermonde matrix
        van = ones(5,3);
        arr = transpose(1:5);
        van(:,2) = arr.^1;
        van(:,1) = arr.^2;
        predPoint = arr(end) + 1;
        % arr_y remembers last positions
        cache_y = ybst;
        numSuccessfulSteps = 0;
        numCacheMaximal = 0;
        numCacheWorked = 0;
        numGradMaximal = 0;
        numGradWorked = 0;
        
        if (visual)
            % create new figure and initialize
            f_output(denormalize(ybst),fbst,jIter,'init',outputFcn);
            % first iteration with start point and jIter = 0
            f_output(denormalize(ybst),fbst,jIter,'iter',outputFcn);
        end
        
        while (~done)
            % increase counter
            jIter = jIter + 1;
            
            if (stuck)
                % choose the smallest step, if any is smaller than the maximum size
                [v,j] = f_min(step,norms,smax,stuckSearchFactor*tolY);
            else
                % choose the largest step
                [v,j] = f_max(step,norms);
            end
            
            if(j == dim+2 || j == dim+3), numCacheMaximal = numCacheMaximal + 1; end
            if(j == dim+1 || j == dim+4), numGradMaximal = numGradMaximal + 1; end
            
            f_display(display,jIter,fbst,norm(v));
            
            % j == -1 indicates minimum found
            if ( j ~= -1 && jIter <= maxIter && funEvals <= maxFunEvals )
                % compute next x, fval
                ycur = ybst + v;
                fcur = fun(ycur,jIter);
                funEvals = funEvals + 1;
                
                % is better estimate?
                delta_f = fcur - fbst;
                if ( delta_f < 0 && (norm(v)>tolX/stuckSearchFactor || abs(delta_f) > tolFun) )
                    numSuccessfulSteps = numSuccessfulSteps + 1;
                    if (j == dim+2 || j == dim+3), numCacheWorked = numCacheWorked + 1; end
                    if (j == dim+1 || j == dim+4), numGradWorked = numGradWorked + 1; end
                    stuck = false; % we are stuck nowhere (anymore)
                    % update x, fval
                    ybst = ycur;
                    fbst = fcur;
                    % contract opp step to not try the previous point next
                    step(:,opp_j(j)) = -contractFactor*v;
                    norms(opp_j(j)) = norm(step(:,opp_j(j)));
                    % TODO do? no.
                    % step(:,j) = expandFactor*step(:,j);
                    % if last step repeated, expand the current step
                    if ( j == lastj )
                        v = expandFactor*v;
                    end
                    % xstep always contains the steps of the last time we moved
                    xstep = step;
                    xnorms = norms;
                    % record the last step
                    lastj = j;
                    % record step
                    step(:,j) = v;
                    norms(j) = norm(v);
                    
                    % update gradient vector
                    if (gradi == -1)
                        % if gradv empty, set gradv to current step and record index
                        gradv = v;
                        gradi = min([j, opp_j(j)]);
                    elseif (gradi == min([j, opp_j(j)]))
                        % if gradv is parallel to current step, add the current vector
                        gradv = (gradv + v);
                    else
                        % else update the extra vector
                        step(:,dim+1) = gradv + v;
                        norms(dim+1) = norm(gradv + v);
                        % set extra entry in smax to max of current step smax and gradv smax
                        % (bound for norm of gradient)
                        smax(dim+1)   = max([smax(min([j,opp_j(j)])),smax(gradi)]);
                        % set the opp step to - extra step
                        step(:,dim+4) = -contractFactor*step(:,dim+1);
                        norms(dim+4) = abs(contractFactor)*norms(dim+1);
                        % update gradv
                        % TODO every round?
                        %gradv         = step(:,j);
                        gradi =-1;%        = min([j, opp_j(j)]);
                    end
                    % update cache
                    if (size(cache_y,2) >= cacheSize)
                        cache_y = [cache_y(:,2:cacheSize),ybst];
                    else
                        cache_y = [cache_y,ybst];
                    end
                    % and update interpolating gradient
                    nCache = size(cache_y,2);
                    if (nCache >= cacheSize)
                        % prepare data
                        ynew = zeros(dim,1);
                        for k=1:dim
                            y = transpose(cache_y(k,:));
                            p = van\y;
                            ynew(k) = polyval(p,predPoint);
                        end
                        % insert difference into step array
                        vnew = ynew - ybst;
                        %vnew = vnew*norm(v)/norm(vnew); % normalize?
                        step(:,dim+2) = vnew;
                        step(:,dim+3) = -contractFactor*vnew;
                        normvnew = norm(vnew);
                        norms(dim+2) = normvnew;
                        norms(dim+3) = abs(contractFactor)*normvnew;
                    end
                    % else: if already stuck, increase the current step size
                else
                    if (stuck)
                        step(:,j) = expandFactor*v;
                        norms(j) = norm(step(:,j));
                        % else: if current step norm >= tolX, decrease the current step size
                    elseif ( norm(step(:,j)) >= tolY )
                        step(:,j) = contractFactor*v;
                        norms(j) = norm(step(:,j));
                        % else: (norm < tolX), set the stuck flag and set all steps to
                        % expandFactor times the last recorded steps
                    else
                        stuck = true;
                        step = expandFactor * xstep;
                        norms = abs(expandFactor)*xnorms;
                    end
                end
                
                % update output
                if (visual)
                    f_output(denormalize(ybst),fbst,jIter,'iter',outputFcn);
                end
                
            else % somehow done
                done = true;
                if (jIter <= maxIter && funEvals <= maxFunEvals)
                    % found a local minimum
                    exitflag = 1;
                else
                    % needed too long
                    exitflag = 0;
                end
            end
            
        end
        
        % assign return values
        x                   = denormalize(ybst);
        fval                = fbst;
        output.funcCount    = funEvals;
        output.iterations   = jIter;
        output.algorithm    = 'Dynamic Hill Climb';
        output.t_cpu        = cputime - starttime;
        
        % finalize output
        if (visual)
            f_output(denormalize(ybst),fbst,jIter,'done',outputFcn);
        end
        %         numSuccessfulSteps
        %         numCacheMaximal
        %         numCacheWorked
        %         numGradMaximal
        %         numGradWorked
    end

end

%% helper functions

function y = f_normalize(x,lb,ub)
y = (x-lb)./abs(ub-lb);
end

function x = f_denormalize(y,lb,ub)
x = y.*(ub-lb) + lb;
end

function j_opp = f_opp_j(j,nVec)
% short for opposite vector in step matrix
j_opp = nVec - (j-1);
end

function fval = f_wrap_fun(x,fun,lb,ub,barrier,jIter,maxIter)
% set fun to inf whenever conditions not fulfilled
if (~isequal(barrier,''))
    fval = fun(x);
    fval = barrierFunction(fval, [], x, [lb, ub], jIter, maxIter, barrier);
else
    % extreme barrier
    if (any(x>ub) || any(x<lb))
        fval = inf;
    else
        fval = fun(x);
    end
end
end

function [step,norms] = f_init_step(vmax)
dim  = length(vmax);

nVec = 2*dim + 2;
step = zeros(dim,nVec);
norms = zeros(1,nVec);
% 2 positions in the center reserved for gradient
for j=1:dim
    initStepSize       = abs(vmax(j));
    step(j,j)          = initStepSize;
    step(j,nVec-(j-1)) = -initStepSize;
    norms(j) = initStepSize;
    norms(nVec-(j-1)) = initStepSize;
end

end

function [step,norms] = f_init_step_2(vmax)
dim = length(vmax);

nVec = 2*dim + 4;
step = zeros(dim,nVec);
norms = zeros(1,nVec);
% 4 positions in the center reserved for gradients
for j=1:dim
    initStepSize       = abs(vmax(j));
    step(j,j)          = initStepSize;
    step(j,nVec-(j-1)) = -initStepSize;
    norms(j) = initStepSize;
    norms(nVec-(j-1))  = initStepSize;
end
end

function [v_max,j_max] = f_max(step,norms)
[~,j_max] = max(norms);
v_max = step(:,j_max);
end

function [v_min,j_min] = f_min(step,norms,smax,tolerance)
% find step with smallest norm, if one exists whose norm is smaller than
% the corresponding smax max step size
minnorm = -1;
j_min   = -1; % -1 used as indicator in calling function
v_min   = -1;
nVec    = size(step,2);
for j=1:nVec
    vnorm = norms(j);
    if ( vnorm <= tolerance && (vnorm < minnorm || minnorm < 0) && vnorm > 0) % tolerance % smax(min([j, nVec-(j-1)]))
        minnorm = vnorm;
        j_min = j;
    end
end

if (j_min ~= -1), v_min = step(:,j_min); end
end

function [tolX,tolFun,maxFunEvals,maxIter,outputFcn,...
    initialStepSize,expandFactor,contractFactor,stuckSearchFactor,barrier,...
    display,mode]...
    = f_extractOptions(options,dim)
% interpret options

if (isfield(options,'TolX') && ~isempty(options.TolX))
    tolX    = options.TolX;
else
    tolX    = 1e-8;
end

if (isfield(options,'TolFun') && ~isempty(options.TolFun))
    tolFun  = options.TolFun;
else
    tolFun  = 1e-8;
end

if (isfield(options,'MaxFunEvals') && ~isempty(options.MaxFunEvals))
    maxFunEvals = options.MaxFunEvals;
else
    maxFunEvals = 1000*dim;
end

if (isfield(options,'MaxIter') && ~isempty(options.MaxIter))
    maxIter = options.MaxIter;
else
    maxIter = 1000*dim;
end

if (isfield(options,'OutputFcn') && ~isempty(options.OutputFcn))
    outputFcn = options.OutputFcn;
else
    outputFcn = nan;
end

% adjustment parameters

if (isfield(options,'InitialStepSize') && ~isempty(options.InitialStepSize))
    initialStepSize           = options.InitialStepSize;
else
    initialStepSize           = 0.1;
end

if (isfield(options,'ExpandFactor') && ~isempty(options.ExpandFactor))
    expandFactor              = options.ExpandFactor;
else
    expandFactor              = 2.1;
end

if (isfield(options,'ContractFactor') && ~isempty(options.ContractFactor))
    contractFactor            = options.ContractFactor;
else
    contractFactor            = 0.47;
end

if (isfield(options,'StuckSearchFactor') && ~isempty(options.StuckSearchFactor))
    stuckSearchFactor         = options.StuckSearchFactor;
else
    stuckSearchFactor         = 4;
end

if (isfield(options,'Barrier'))
    barrier                   = options.Barrier;
else
    barrier                   = '';
end

if (isfield(options,'Display') && ~isempty(options.Display))
    display                   = options.Display;
else
    display                   = 'off';
end

if (isfield(options,'Mode') && ~isempty(options.Mode))
    mode = options.Mode;
else
    mode = -1;
end

end

function f_output(x,fval,iter,state,outputFcn)
% short for call to output function
optimValues.fval = fval;
optimValues.iteration = iter;
outputFcn(x,optimValues,state);
end

function f_display(display,jIter,fbst,vnorm)
if (strcmp(display,'iter') || strcmp(display,'debug'))
    if (strcmp(display,'debug'))
        show_output = true;
    else
        show_output = mod(jIter,100) == 0;
    end
    
    if (show_output), fprintf(strcat('%d\t|\t%.15f\t|\t%.15f\n'),jIter,fbst,vnorm); end
end

% function answer = f_haveAllSameLength(arr)
%     answer = true;
%     dim = size(arr,1);
%     s = norm(arr(:,1));
%     for j=2:size(arr,2)
%         s2 = norm(arr(:,j));
%         if (abs(s-s2) >= dim*eps)
%             answer = false;
%             break;
%         end
%     end
% end
%
% function U = f_createRandomOrthogonalMatrix(dim)
%     M = randn(dim,dim);
%     [Q,R] = qr(M);
%     D = diag(R);
%     D = diag(D)./abs(D);
%     U = Q * D;
% end

end
```
