## Supplementary material for "Mechanistic description of spatial processes using integrative modelling of noise-corrupted imaging data": getBins.m

```
%getBins calcualtes the number of bins in a histogramm for the vector
%values according to the option (optimal [default] or conservative)
%
% USAGE:
% ======
% nbin = getBins(values,option)
%
% INPUTS:
% =======
% values ... vector for which histogramm should be plotted.
% option ... options for the number of bins (optimal [default] and
% conservative)
%
% Outputs:
% ========
% nbins .. number of bins for histogram

function [nbin] = getBins(values,option)

switch option
    case 'optimal'
        h = 3.49*nanstd(values)/(length(values)^(1/3));
        nbin = round((max(values)-min(values))/h);
    case 'conservative'
        h = 2*3.49*nanstd(values)/(length(values)^(1/3));
        nbin = round((max(values)-min(values))/h);
    otherwise %'optimal' and the rest [default]
        h = 3.49*nanstd(values)/(length(values)^(1/3));
        nbin = round((max(values)-min(values))/h);
end

end
```
