## Supplementary material for "Mechanistic description of spatial processes using integrative modelling of noise-corrupted imaging data": getConstraints.m

```
%% Constraint generation
% This function is used to generate the linear constraints for the
% reduced system. 
%   theta ... parameter vector
%   parameter struct ...
%   I ... index set of optimized parameters
function [A,b,Aeq,beq] = getConstraints(theta,parameters,I)

% Index set of parameters which are eliminated
i = setdiff(1:parameters.number,I);

% Reduction of constraints to remaining dimensions
if ~isempty(parameters.constraints.A)
    A = parameters.constraints.A(:,I);
    if ~isempty(i)
        b = parameters.constraints.b - parameters.constraints.A(:,i)*theta(i);
    end
else
    A = [];
    b = [];
end
if ~isempty(parameters.constraints.Aeq)
    Aeq = parameters.constraints.Aeq(:,I);
    if ~isempty(i)
        beq = parameters.constraints.beq - parameters.constraints.Aeq(:,i)*theta(i);
    end
else
    Aeq = [];
    beq = [];
end

end
```
