## Supplementary material for "Mechanistic description of spatial processes using integrative modelling of noise-corrupted imaging data": getEllipse.m

```
function X = getEllipse(m,C,d)
% TODO
%
% Parameters:
% m:
% C:
% d:
%
% Return values:
% X:
phi = linspace(0,2*pi,100);

[V,L] = eig(C);
if max(diag(L)) < 1e-8
    X = m;
elseif min(diag(L)) > 1e-8
    X = bsxfun(@plus,d*sqrtm(squeeze(C))*[cos(phi);sin(phi)],m(:));
else
    [~,i] = max(diag(L));
%     V(:,i)*sqrt(L(i,i))
%     (V(:,i)*sqrt(L(i,i)))*(V(:,i)*sqrt(L(i,i)))'
%     C
    X = [m(:)-d*sqrt(L(i,i))*V(:,i),m(:)+d*sqrt(L(i,i))*V(:,i)];
end
```
