## Supplementary material for "Mechanistic description of spatial processes using integrative modelling of noise-corrupted imaging data": getFiniteDifferences.m

```
function varargout = getFiniteDifferences(theta, objectiveFunction, mode)

% This function computes the gradient and, if necessary, the Hessian of
% the objective function by a finite difference scheme, if the
% objective function itself does not provide this information.
%
% Parameters:
%   theta: parameter vector
%   objectiveFunction: the original objective function
%   mode: double, specifying if grad, Hessian or both should be computed:
%           1: gradient
%           2: gradient and Hessian
%           3: Hessian from gradient output of original obj. fun
%
% Return values (depend on mode):
%   gradient: mode == 1
%   [objective, gradient, Hessian]: mode == 2
%   Hessian: mode == 3

    persistent lastParameter;
    persistent fdStepParameter;
    persistent fdStepGradient;
    
    nPar = length(theta);
    
    switch mode
        case 1
        % Compute gradient from an objective function, which returns only
        % one value
        
            % Compute new step size if necessary
            if isempty(lastParameter)
                fdStepParameter = getStepSizeFD(theta, objectiveFunction, 1);
                lastParameter = theta;
            elseif (sum((theta - lastParameter).^2) > 0.5 * sqrt(nPar))
                fdStepParameter = getStepSizeFD(theta, objectiveFunction, 1);
                lastParameter = theta;
            end
            
            % Initialize gradient
            G = zeros(nPar, 1);
            
            % Use FD scheme
            for j = 1 : nPar
                % Set the step
                delta = zeros(nPar, 1);
                delta(j) = fdStepParameter(j);
                
                % Gradient computation
                Jplus = objectiveFunction(theta + delta);
                Jminus = objectiveFunction(theta - delta);
                G(j) = (Jplus - Jminus) / (2 * fdStepParameter(j));
            end
            
            % Assign output
            varargout{1} = G;
            
        case 2
        % Compute objective value, gradient and Hessian from an objective 
        % function, which returns only one value
            
            % Compute new step size if necessary
            if isempty(lastParameter)
                fdStepParameter = getStepSizeFD(theta, objectiveFunction, 1);
                lastParameter = theta;
            elseif (sum((theta - lastParameter).^2) > 0.5 * sqrt(nPar))
                fdStepParameter = getStepSizeFD(theta, objectiveFunction, 1);
                lastParameter = theta;
            end
            
            % Initialize gradient
            G = zeros(nPar, 1);
            H = zeros(nPar, nPar);
            
            % Objective function at given parameter vector
            J = objectiveFunction(theta);
            
            % Do FD
            for j = 1 : nPar
                % Set the step
                delta = zeros(nPar, 1);
                delta(j) = fdStepParameter(j);
                
                % Gradient and diagonal of the Hessian
                Jplus = objectiveFunction(theta + delta);
                Jminus = objectiveFunction(theta - delta);
                G(j) = (Jplus - Jminus) / (2 * fdStepParameter(j));
                H(j,j) = (Jplus + Jminus - 2*J) / (fdStepParameter(j)^2);
                
                % Rest of the Hessian
                for k = 1 : j-1
                    % Set steps
                    delta1 = delta;
                    delta2 = delta;
                    delta1(k) = fdStepParameter(k);
                    delta2(k) = -fdStepParameter(k);
                    
                    % Compute values
                    Jpp = objectiveFunction(theta + delta1);
                    Jpm = objectiveFunction(theta + delta2);
                    Jmp = objectiveFunction(theta - delta2);
                    Jmm = objectiveFunction(theta - delta1);
                    H(j, k) = (Jpp - Jpm - Jmp + Jmm) ...
                        / (4 * fdStepParameter(j) * fdStepParameter(k));
                    H(k, j) = H(j, k);
                end
            end
            
            % Assign output
            varargout{1} = J;
            varargout{2} = G;
            varargout{3} = H;
            
        case 3
        % Compute Hessian from an objective function which only provides
        % the gradient
            
            % Compute step size if necessary
            if isempty(lastParameter)
                fdStepGradient = getStepSizeFD(theta, objectiveFunction, 2);
                lastParameter = theta;
            elseif (sum((theta - lastParameter).^2) > 0.5 * sqrt(nPar))
                fdStepGradient = getStepSizeFD(theta, objectiveFunction, 2);
                lastParameter = theta;
            end
            
            % Initialize gradient
            H = zeros(nPar, nPar);
            
            % Use FD scheme
            for j = 1 : nPar
                % Set the step
                delta = zeros(nPar, 1);
                delta(j) = fdStepGradient(j);
                
                % Gradient computation
                [~, Gplus] = objectiveFunction(theta + delta);
                [~, Gminus] = objectiveFunction(theta - delta);
                H(:,j) = (Gplus - Gminus) / (2 * fdStepGradient(j));
            end
            
            % Assign output
            varargout{1} = H;
            
    end
end
```
