## Supplementary material for "Mechanistic description of spatial processes using integrative modelling of noise-corrupted imaging data": getKernelDensityEstimate.m

```
function [KDest,Sigma] = getKernelDensityEstimate(varargin)
% kde_simple.m computes the kernel density estimate KDest at X given the sample P.
% The bandwidth can be determined using different methods (see below).
%
% USAGE:
% [KDest,Sigma] = kde_simple(P,X,Sigma,options)
%
% Parameters:
% varargin:
% P: D x N - matrix, where each column defines a member of the sample.
% X: D x Ngrid - matrix, where each column defines a point
%       at which the kernel density estimate is determined.
% Sigma: kernel bandwidth. (If a kernel bandwidth is provided, it is
%       used. Otherwise, the bandwidth is computed from the data.)
% options: options for the algorithm<pre>
%   .bw_selection ... bandwidth selction methods:
%       = 'scott' ... Scott's rule
%       = 'gen scott' (default) ... generalized Scott's rule
%       = 'user' ... bandwidth is provided
%   .kernel ... type of kernel:
%       = 'normal' (default) ... multi-variate Gausian kernel
%       = 'log-normal' ... multi-variate log-normal kernel</pre>
%
% Return values:
% KDest: kernel density estimate at points defined by X.
% Sigma: covariance matrix of Gaussian kernel detemined
%           using the generalized Scott's rule.
%
% History:
% * 07/10/2010 - Jan Hasenauer
% * modified 14/01/2011 - Jan Hasenauer

%% CHECK/ASSIGN INPUTS
if nargin >= 2
    P = varargin{1};
    X = varargin{2};
else
    error('Not enough inputs!');
end

Sigma = [];
if nargin >= 3
    Sigma = varargin{3};
end
if ~isempty(Sigma)
    if (size(Sigma,1) ~= size(Sigma,2)) || (size(Sigma,2) ~= size(P,1))
        error('Dimension of data D and bandwidth Sigma does not agree.');
    end
end

% Check options
options.bw_selection = 'scott';
options.kernel = 'normal';
if nargin == 4
    options = setdefault(varargin{4},options);
end
if ~isempty(Sigma)
    options.bw_selection = 'user';
end

% Check dimension
if size(P,1) ~= size(X,1)
    error('Dimensionality of sample members and grid points must agree!');
end

%% INITIALIZATION
% Assign dimensions
D = size(P,1);
N = size(P,2);
Ngrid = size(X,2);
% Adapt coordinates dependent on kernel shape
% (We adapt the coordinates instead of the kernel
%  shape to ensure computational efficiency.)
switch options.kernel
    case 'normal'
        Xc = X;
    case 'log-normal'
        % Check positivity of grid and data points
        if (multimin(P) <= 0) || (multimin(X) <= 0)
            % error
            error('Data or grid points contain negative values or zeros. => log-normal kernels can not be used.');
        end
        P = log(P);
        Xc = log(X);
        % I do not know how the generalize Scott's rule would look in this
        % cases. This has to be checked in the future. Till then it is not
        % allowed in combination with log-normal kernels.
        options.bw_selection = 'scott';
    otherwise
        % error
        error('This option is not available.');
end
% Compute kernal shape, bandwidth, and scaling constant
if isempty(Sigma)
    switch options.bw_selection
        case 'scott'
            Sigma = diag((var(P')) * N^(-2/(D+4)));
        case 'gen scott';
            Sigma = cov(P') * N^(-2/(D+4));
        case 'user'
            % Nothing has to be done.
        otherwise
            % error
            error('This option is not available.');
    end
end
% invSigma = inv(Sigma);
c = 1/((2*pi)^(D/2)*sqrt(det(Sigma))) * 1/N;
% Initialize KDest
KDest = zeros(1,Ngrid);

%% CALCULATION OF DENSITY
for i = 1:N
    % Updata of complete density
    KDest = KDest + c*exp(-0.5*sum(bsxfun(@minus,Xc,P(:,i)).*(Sigma\bsxfun(@minus,Xc,P(:,i))),1));
end


% Adapt kde required for log-normal kernels
switch options.kernel
    case 'normal'
        % Nothing has to be done!
    case 'log-normal'
        KDest = KDest./prod(X,1);
end

end
```
