## Supplementary material for "Mechanistic description of spatial processes using integrative modelling of noise-corrupted imaging data": getNextProfilePoint.m

```
%% getNextProfilePoint is a support function for the profile calculation
%   and is called by computeProfile. It determines the length of the 
%   update step given update direction, parameter constraints,
%   log-posterior and target log posterior.
%
% USAGE:
% ======
% function [theta_next,logPost] = getNextPoint(theta,theta_min,theta_max,dtheta,logPost_target,objective_function)
%
% INPUTS:
% =======
% theta ... starting parameter   
% theta_min ... lower bound for parameters   
% theta_max ... upper bound for parameters   
% dtheta ... upper direction
% logPost_target ... target value for log-posterior
% objective_function ... log-posterior of model as function of the parameters.
%
% Outputs:
% ========
% theta_next ... parameter proposal
% logPost ... log-posterior at proposed parameter
%
% 2012/07/12 Jan Hasenauer

function [theta,J] = getNextProfilePoint(theta,theta_min,theta_max,dtheta,c,c_min,c_max,c_update,J_target,obj,constraints,update_mode,j)

    % Initialization
    % 1) modification of dtheta
    switch update_mode
        case 'multi-dimensional'
            % nothing has to be done
        case 'one-dimensional'
            dtheta([1:j-1,j+1:end]) = 0;
    end

    % 1) line search
    if dtheta(j) > 0 % increasing
        c_bound = (theta_max(j)-theta(j))/dtheta(j);
    else
        c_bound = (theta_min(j)-theta(j))/dtheta(j);
    end
    if c_bound > c_min
        c_max = min(c_max,c_bound);
        c = min(max(c,c_min),c_max);
        search = 1;
    else
        c_min = c_bound;
        c_max = c_bound;
        c = c_bound;
        search = 0;
    end
    % 2) inequality constraints
    if ~isempty(constraints.A)
        A = constraints.A;
        b = constraints.b;
    else
        A = zeros(1,length(theta));
        b = 1;
    end
    % 3) parameter projection
    theta_fun = @(c) max(min(theta + c*dtheta,theta_max),theta_min);

    % Search
    theta = theta_fun(c);
    if  min(A*theta <= b)
        J = obj(theta);
    else
        J = inf;
    end

    if search == 1
        if J > J_target % => initial c too large
            stop = 0;
            while stop == 0
                c = min(max(c/c_update,c_min),c_max);
                theta = theta_fun(c);
                if c == c_min % lower bound reached
                    stop = 1;
                    J = obj(theta);
                elseif min(A*theta <= b) % feasible
                    J = obj(theta);
                    if J <= J_target % objective smaller than target value
                        stop = 1;
                    end
                end
            end
        else % => initial c too small
            stop = 0;
            while stop == 0
                cn = min(max(c*c_update,c_min),c_max);
                thetan = theta_fun(cn);
                if min(A*theta <= b) % feasible
                    Jn = obj(thetan);
                    if Jn <= J_target % objective smaller than target value
                        c = cn;
                        theta = thetan;
                        J = Jn;
                        if cn == c_max % upper bound reached
                            stop = 1;
                        end
                    else
                        stop = 1;
                    end
                else
                    stop = 1;
                end
            end
        end
    end

end
```
