## Supplementary material for "Mechanistic description of spatial processes using integrative modelling of noise-corrupted imaging data": getParProfilesByIntegration.m

```
function [parameters, fh] = getParProfilesByIntegration(parameters, objective_function, options, varargin)
% getParProfilesByIntegration.m calculates the profiles likelihoods for the
% model parameters, starting from the maximum a posteriori estimate. 
% This is done by integrating an ODE which follows the optimal path for a
% given parameter.
%
% USAGE:
% [...] = getParameterProfiles(parameters, objective_function)
% [...] = getParameterProfiles(parameters, objective_function, options)
% [parameters, fh] = getParameterProfiles(...)
%
% getParProfilesByIntegration() uses the following PestoOptions members:
%  * PestoOptions::calc_profiles
%  * PestoOptions::comp_type
%  * PestoOptions::dJ
%  * PestoOptions::dR_max
%  * PestoOptions::fh
%  * PestoOptions::MAP_index
%  * PestoOptions::mode
%  * PestoOptions::obj_type
%  * PestoOptions::options_getNextPoint .guess .min .max .update .mode
%  * PestoOptions::parameter_index
%  * PestoOptions::profile_method
%  * PestoOptions::profileReoptimizationOptions
%  * PestoOptions::plot_options
%  * PestoOptions::R_min
%  * PestoOptions::save
%  * PestoOptions::solver .AbsTol .algorithm .eps .gamma .hessian 
%       .linSolver .MaxStep .MaxNumSteps .minCond .MinStep .nonlinSolver 
%       .RelTol .type
%
% Parameters:
%   parameters: parameter struct
%   objectiveFunction: objective function to be optimized. 
%       This function should accept one input, the parameter vector and
%       return the objective value, its gradient and, if possible, the
%       Hessian matrix.
%   options: A PestoOptions object
%   varargin:
%     fh: A figure handle. If not provided, a figure will be generated.
%
% Required fields of parameters:
%   number: Number of parameters
%   min: Lower bound for each parameter
%   max: upper bound for each parameter
%   name = {'name1', ...}: names of the parameters
%   MS: results of global optimization, obtained using for instance 
%       the routine 'getMultiStarts.m'. MS has to contain at least
%     * par: sorted list n_theta x n_starts of parameter estimates.
%          The first entry is assumed to be the best one.
%     * logPost: sorted list n_starts x 1 of of log-posterior values
%          corresponding to the parameters listed in .par.
%     * hessian: Hessian matrix (or approximation) at the optimal point
%
% Return values:
%   parameters: updated parameter struct
%   fh: figure handle
%
% Generated fields of parameters:
%   P(i): profile for i-th parameter
%     * par: MAPs along profile
%     * logPost: maximum log-posterior along profile
%     * R: ratio
%
% History:
% 2013/10/05 FF original code from thesis
% 2014/04/09 Sabrina Hross - completely reworked
% 2016/11/21 Paul Stapor
% 2017/02/02 Paul Stapor - PESTO version of the code

    %% CHECK AND ASSIGN INPUTS
    if (nargin >= 4)
        fh = varargin{1};
        options.fh = fh;
    else
        fh = [];
    end
    
    %% Check for fixed parameters
    if ~isempty(options.fixedParameters)
        error('Fixed parameters are currently not supported by getParProfilesByIntegration.');
    end
    
    %% Preperation of folder
    if options.save
        [~,~,~] = mkdir(options.foldername);
        save([options.foldername '/init'],'parameters');
    end

    %% Profile calculation -- SEQUENTIAL
    
    % Profile calculation
    if strcmp(options.comp_type, 'sequential')
        
        for j = options.profile_integ_index
            parameters = integrateProfileForParameterI(parameters, objective_function, j, options, fh);
        end
        
        
    elseif strcmp(options.comp_type, 'parallel')
        parfor j = options.profile_integ_index
            integrateProfileForParameterI(parameters, objective_function, j, options, fh);
        end
        
        % Output
        if strcmp(options.profile_method, 'integration')
            switch options.mode
                case 'visual', fh = plotParameterProfiles(parameters,'1D',fh,options.parameter_index,options.plot_options);
                case 'text' % no output
                case 'silent' % no output
            end
        end
    end
end


function parameters = integrateProfileForParameterI(parameters, objectiveFunction, iPar, options, fh)
 

    % Define global variables for communication across ODE solver
    global llhHistory;
    global yCorrection;
    global ObjFuncCounter;
    global reOptSteps;
    global intSteps;
    ObjFuncCounter = 0;
    lastCounter = 0;

    % Initial condition (used only for Profile integration)
    t0 = parameters.MS.par(iPar, options.MAP_index);
    
    % Set objective function handle
    negLogPost = setObjectiveWrapper(objectiveFunction, options, 'negative log-posterior', [], [], true, true);
    
    switch options.solver.type
        case 'CVODE'            
            cvodeOptions = CVodeSetOptions('RelTol', options.solver.RelTol, ...
                'AbsTol', options.solver.AbsTol, ...
                'MaxStep', options.solver.MaxStep, ...
                'MinStep', options.solver.MinStep, ...
                'LinearSolver', options.solver.linSolver, ...
                'MaxNumSteps', options.solver.MaxNumSteps ...
            );
        
        case {'ode45', 'ode23', 'ode23s', 'ode23t', 'ode15s', 'ode113'}
            odeMatlabOptions = odeset('RelTol', options.solver.RelTol, ...
                'AbsTol', options.solver.AbsTol, ...
                'MaxStep', options.solver.MaxStep ...
            );

        case 'ode15sDAE' 
            daeMatlabOptions = odeset('RelTol', options.solver.RelTol, ...
                'AbsTol', options.solver.AbsTol, ...
                'MStateDependence', 'strong', ...
                'MaxStep', options.solver.MaxStep, ...
                'MassSingular', 'yes', ...
                'OutputFcn', checkOptimality...
                );
    end

    % Check if user-supplied parameter function -> ToBeDone ... Later
    parameterFunction = @(theta, index) SingleParameter(theta, index);
    ySize = parameters.number;
    
    % !!! IMPORTANT: Think carefully, if this is correct for general 
    % parameter functions...
    % lambda = -(DG' * parameters.MS.gradient(:,1)) / (DG' * DG);

    %% Compute profile for in- and decreasing theta_i
    for s = [1, -1]

        % Set bound for considered parameter (ONLY integrate profiles)
        borders = [parameters.min, parameters.max];
        switch s
            case 1
                T = parameters.max(iPar);
            case -1
                T = parameters.min(iPar);
        end 

        % Starting point
        theta  = parameters.MS.par(:, options.MAP_index);
        llhHistory = parameters.MS.logPost(options.MAP_index);
        reachedEnd = 0;
        OutputFunction = @(t, y, flag) checkOptimality(t, y, flag, s, iPar, ...
            parameters.MS.logPost(1), objectiveFunction, borders, options);

        if ~strcmp(options.solver.hessian, 'user-supplied')
            approximateHessian(parameters.MS.par(:,options.MAP_index), -parameters.MS.gradient(:,options.MAP_index), parameters.MS.hessian(:,:,options.MAP_index), [], 'init');
        end
        
        % Pre-Output
        if (strcmp(options.mode, 'text'))
            fprintf('\n  |  Integrating Parameter %4i, s = %2i  |', iPar, s);
            fprintf('\n  |======================================|');
            fprintf('\n  | Running axis |  Optimality |  Ratio  |');
            fprintf('\n  |--------------|-------------|---------|');
        end
        
        % Get the computation time
        startTimeProfile = cputime;
        
        intSteps = 0;
        optSteps = 0;
        reOptSteps = 0;
        
        % Switch between different methods
        while (reachedEnd == 0)

            if (s == 1)
                theta = theta(:,end);
            else
                theta = theta(:,1);
            end
            llhHistory = llhHistory(end);

            switch options.solver.type
                case {'ode45', 'ode15s', 'ode113','ode23', 'ode23s', 'ode23t'}
                    odeMatlabOptions.OutputFcn = OutputFunction;
                    odeMatlabOptions.Events = @(t,y) getEndProfile(t, s, y, iPar, borders, negLogPost, options, parameters.MS.logPost(1));
                    if (strcmp(options.solver.type, 'ode15s'))
                        [~,y] = ode15s(@(t,y) getRhsRed(t, s, y, iPar, borders, negLogPost, parameterFunction, options),[s*theta(iPar), s*T], theta, odeMatlabOptions); 
                    elseif (strcmp(options.solver.type, 'ode45'))
                        error('ode45 is currently not implemented for profiling.');
                        % [~,y] = ode45(@(t,y) getRhsRed(t, s, y, j, borders, negLogPost, parameterFunction, options),[s*theta(j), s*T], theta, odeMatlabOptions);  
                    elseif (strcmp(options.solver.type, 'ode113'))
                        [~,y] = ode113(@(t,y) getRhsRed(t, s, y, iPar, borders, negLogPost, parameterFunction, options),[s*theta(iPar), s*T], theta, odeMatlabOptions); 
                    elseif (strcmp(options.solver.type, 'ode23s'))
                        [~,y] = ode23s(@(t,y) getRhsRed(t, s, y, iPar, borders, negLogPost, parameterFunction, options),[s*theta(iPar), s*T], theta, odeMatlabOptions);  
                    elseif (strcmp(options.solver.type, 'ode23t'))
                        [~,y] = ode23t(@(t,y) getRhsRed(t, s, y, iPar, borders, negLogPost, parameterFunction, options),[s*theta(iPar), s*T], theta, odeMatlabOptions);
                    elseif (strcmp(options.solver.type, 'ode23'))
                        [~,y] = ode23(@(t,y) getRhsRed(t, s, y, iPar, borders, negLogPost, parameterFunction, options),[s*theta(iPar), s*T], theta, odeMatlabOptions);
                    end


                    % If yCorrection is set to inf, then the ODE is too stiff, 
                    % some steps of optimization based calculation have to be done
                    if (yCorrection == inf)
                        addY = doOptimizationSteps(parameters, y, objectiveFunction, borders, iPar, s, options);
                        y = [y; addY];
                        optSteps = optSteps + 3;
                    else
                    % If reoptimization has to be done, correct the values in y by the optimized ones
                        for iLine = size(yCorrection, 2) : -1 : 1
                            % Not sure: Either all entries are nan, or none
                            % of them, so it if sufficient to test the 1st?
                            if ~isnan(yCorrection(1,iLine))
                                y(end + 1 - iLine, :) = yCorrection(:,iLine)';
                            end
                        end
                    end

                    if (s == 1)
                        theta = y';
                    else
                        theta = fliplr(y');
                    end

                case 'CVODE' 
                    error('CVODE is currently not implemented for profiling.');
%                     cvodeOptions.RootsFn = @(t,y) getEndProfile(t, s, y, j, borders, objective_function, options, parameters.MS.logPost(1));
%                     CVodeInit(@(t,y,data) getRhsRed(t, s, y, j, borders, objective_function, parameterFunction, options), options.solver.algorithm, options.solver.nonlinSolver, s*t0, theta, cvodeOptions);
% 
%                     killCounter = 0;
%                     reachedEndCVODE = 0;
%                     iTer = 1;
%                     y = nan(ySize, 100);
%                     t = nan(1, 100);
%                     while (reachedEndCVODE == 0)
%                         try
%                             if(killCounter == 1)
%                                 t_00 = t_tmp  + 1e-4;
%                                 y_00 = y_tmp;
%                                 cvodeOptions = CVodeSetOptions('RelTol', options.solver.RelTol, ...
%                                                 'AbsTol', options.solver.AbsTol, ...
%                                                 'MaxStep', options.solver.MaxStep, ...
%                                                 'MinStep', options.solver.MinStep, ...
%                                                 'LinearSolver', options.solver.linSolver, ...'StabilityLimDet', true, ...
%                                                 'MaxNumSteps', options.solver.MaxNumSteps ...'MaxOrder', 12
%                                                 );
%                                 CVodeInit(@(t,y,data) getRhsRed(t, s, y, j, borders, objective_function, parameterFunction, options), 'BDF', 'Newton', s*t_00, y_00, cvodeKillOptions);  
%                                 killCounter = 0;
%                             end
%                             [~, t_tmp, y_tmp] = CVode(s*T, 'OneStep');
%                         catch
%                             t_000 = t_tmp  + 0.025;
%                             while(s*t_000 - s*t_tmp > 0)
%                                 t_00 = t_tmp  + 5e-5;
%                                 y_00 = y_tmp;
%                                 cvodeKillOptions = CVodeSetOptions('RelTol', 1e-3, ...
%                                     'AbsTol', 1e-5, ...
%                                     'MaxStep', options.solver.MaxStep,...
%                                     'MinStep', 1e-4, ...
%                                     'LinearSolver', options.solver.linSolver, ...'StabilityLimDet', true, ...
%                                     'MaxNumSteps', options.solver.MaxNumSteps ...'MaxOrder', 12
%                                     );
%                                 CVodeInit(@(t,y,data) getRhsRed(t, s, y, j, borders, objective_function, parameterFunction, options), options.solver.algorithm, options.solver.nonlinSolver, s*t_00, y_00, cvodeOptions);  
%                                 [~, t_tmp, y_tmp] = CVode(s*(t_00 + 0.025), 'OneStep');
%                             end
%                             killCounter = 1;
%                         end
%                         y(:, iTer) = y_tmp;
%                         t(1, iTer) = t_tmp;
%                         if ((cvodeOptions.RootsFn(t_tmp, y_tmp) < 0) || ((s*T - t_tmp) < 0))
%                             reachedEndCVODE = 1;
%                         end
%                         iTer = iTer + 1;
%                         if (abs(iTer/100) < 0.001)
%                             y = [y, zeros(ySize, 100)];
%                             t = [t, zeros(1, 100)];
%                         end
%                     end
%                     y = y(1:ySize, 1:iTer-1);
%                     if (s == 1)
%                         theta = y(:,:);
%                     else
%                         theta = fliplr(y(:,:));
%                     end
%                     t = t(1:iTer-1);
% 
%                     %release CVODE Worksspace
%                     CVodeFree;               

                case 'ode15sDAE' 
                    error('ode15sDAE is currently not implemented for profiling.');
                    daeMatlabOptions.Mass = @(t, y) getMassmatrixDAE(t, s, y, iPar, negLogPost, parameterFunction);
                    daeMatlabOptions.Events = @(t,y) getEndProfile(t, s, y, iPar, borders, negLogPost, options, parameters.MS.logPost(1));
                    [~, y] = ode15s(@(t,y) getRhsDAE(t, s, y, 1, iPar, negLogPost, options), [s*t0, s*T], theta, daeMatlabOptions);

                    if (s == 1)
                        theta = y';
                    else
                        theta = fliplr(y');
                    end
            end

            %% Write results to the parameters struct
            switch s
                case 1
                    parameters.P(iPar).logPost = [parameters.P(iPar).logPost, llhHistory];
                    parameters.P(iPar).R = [parameters.P(iPar).R, exp(llhHistory - parameters.MS.logPost(1))];
                    parameters.P(iPar).par = [parameters.P(iPar).par, theta];

                    if ((parameters.P(iPar).R(end) <= options.R_min) ...
                            || parameters.P(iPar).par(iPar, end) >= borders(iPar, 2))
                        reachedEnd = 1;
                    end

                case -1
                    parameters.P(iPar).logPost = [fliplr(llhHistory), parameters.P(iPar).logPost];
                    parameters.P(iPar).R = [exp(fliplr(llhHistory) - parameters.MS.logPost(1)), parameters.P(iPar).R];
                    parameters.P(iPar).par = [theta, parameters.P(iPar).par];

                    if ((parameters.P(iPar).R(1) <= options.R_min) ...
                            || parameters.P(iPar).par(iPar, 1) <= borders(iPar, 1))
                        reachedEnd = 1;
                    end
            end

        end

        % Get computation time
        parameters.P(iPar).t_cpu(round((s+3)/2)) = cputime - startTimeProfile;
        parameters.P(iPar).optSteps(round((s+3)/2)) = optSteps;
        parameters.P(iPar).intSteps(round((s+3)/2)) = intSteps;
        parameters.P(iPar).reOptSteps(round((s+3)/2)) = reOptSteps;
        
        % Final output and storage
        if (strcmp(options.mode, 'text'))
            fprintf('\n  |======================================|\n');
            fprintf('\n  Total RHS evaluations: %i', ObjFuncCounter - lastCounter);
            fprintf('\n  Total Steps: %i\n', length(llhHistory));
        end
        lastCounter = ObjFuncCounter;

        % Save
        if (options.save)
            dlmwrite([options.foldername '/P' num2str(iPar,'%d') '__par.csv'],P_par,'delimiter',',','precision',12);
            dlmwrite([options.foldername '/P' num2str(iPar,'%d') '__logPost.csv'],P_logPost,'delimiter',',','precision',12);
            dlmwrite([options.foldername '/P' num2str(iPar,'%d') '__R.csv'],P_R,'delimiter',',','precision',12);
        end  

        % Output
        if ~strcmp(options.comp_type,'parallel')
            switch options.mode
                case 'visual', fh = plotParameterProfiles(parameters, '1D', fh, options.parameter_index, options.plot_options);
                case 'text'   % no output
                case 'silent' % no output
            end
        end        
    end 

end


function status = checkOptimality(t, y, flag, s, ind, logPostMax, objectiveFunction, borders, options)
    
    % Assume successful Check
    status = 0;
    persistent lastT;
    persistent CounterMinStep;
    global llhHistory;
    global yCorrection;
    global reOptSteps;
    global intSteps;
    
    % Initialize persistent variables in the beginning
    if strcmp(flag, 'init')
        lastT = t(1);
        CounterMinStep = 0;
        
    elseif strcmp(flag, 'done')
        lastT = [];
        clear CounterMinStep;
        
    else
        yCorrection = nan(size(y));
       logPost = setObjectiveWrapper(objectiveFunction, options, 'log-posterior', [], [], true, true);
        
        for iT = 1 : length(t)
            % Check if Minimum step size was violated
            dt = t(iT) - lastT;
            if (dt < options.solver.MinStep)
                CounterMinStep = CounterMinStep + 1;
            else
                CounterMinStep = 0;
            end
            lastT = t(iT);

            % Abort and start reoptimization, if minimum step size was violated
            if (CounterMinStep >= 10)
                display('Violated minimum step size at least 10 times in a row. Doing some optimization steps!');
                yCorrection = inf;
                status = 1;
            end

            % Check, if first optimality is violated, reoptimize if necessary
            [L, GL] = logPost(y(:,iT));
            GL(ind) = 0;
            
            % Update Hessian, if approximation method ist used
            if (~strcmp(options.solver.hessian, 'user-supplied'))
                approximateHessian(y(:,iT), -GL, [], options.solver.hessian, 'write');
            end
            
            if (sqrt(sum(GL.^2)) > options.solver.GradTol)
                isBord = 0;
                for bord = reshape(borders, [1 numel(borders)])
                    if (abs(y(ind) - bord) < 1e-8)
                        display('Lost optimal path at border!');
                        isBord = 1;
                    end
                end
                if ~isBord
                    display('Lost optimal path, doing a reoptimization!');
                    [newY, L, GL] = reoptimizePath(y(:,iT), ind, objectiveFunction, borders, options);
                    y(:,iT) = [newY(1:ind-1); y(ind,iT); newY(ind:end)];
                    yCorrection(:,iT) = y(:,iT);
                    reOptSteps = reOptSteps + 1;
                    status = 1;
                end
            end
            
            % Write ratio and increase counter
            R = exp(L - logPostMax);
            intSteps = intSteps + 1;
            
            if (strcmp(options.mode, 'text'))
                fprintf('\n  |  %11.8f | %11.7f | %7.5f |', ...
                    s*t(iT), sqrt(sum(GL.*GL)), R);
            end
            llhHistory = [llhHistory, L];
            if (R < options.R_min)
                status = 1;
            end
        end
        
    end
end


function [newY, newL, newGL] = reoptimizePath(theta, iPar, objectiveFunction, borders, options)
    
    I1 = (1 : iPar-1)';
    I2 = (iPar+1 : length(theta))';
    I = [I1; I2];
    
    % options.profileReoptimizationOptions.Display = 'off';
    negLogPostReduced = setObjectiveWrapper(objectiveFunction, options, 'negative log-posterior', iPar, theta(iPar), true, true);
    
    if (~isfield(options.profileReoptimizationOptions, 'HessFcn') ...
        || isempty(options.profileReoptimizationOptions.HessFcn))
        % this only works for box-constraints at the moment
        options.profileReoptimizationOptions.HessFcn = @(varargin) HessianWrap(negLogPostReduced, varargin);
    end 
        
    % Optimization
    [newY, newL, ~, ~, ~, newGL, newHL] = ...
        fmincon(negLogPostReduced,...
        theta(I),...
        [], [],... % linear inequality constraints
        [], [],... % linear equality constraints
        borders(I,1),...   % lower bound
        borders(I,2),...   % upper bound
        [],options.profileReoptimizationOptions);    % options
    newL = -newL;
    newGL = -newGL;
    newHL = -newHL;
    
    if ~strcmp(options.solver.hessian, 'user-supplied')
        theta = [newY(1:iPar-1); theta(iPar); newY(iPar:end)];
        tmpHL = [newHL(1:iPar-1,1:iPar-1), zeros(iPar-1,1), newHL(1:iPar-1,iPar:end);...
            zeros(1,length(theta)); newHL(iPar:end,1:iPar-1), zeros(length(theta)-iPar,1), newHL(iPar:end,iPar:end)];
        tmpHL(iPar,iPar) = inf;
        tmpGL = [newGL(1:iPar-1); inf; newGL(iPar:end)];
        approximateHessian(theta, -tmpGL, tmpHL, options.solver.hessian, 'reinit');
    end
end


function y = doOptimizationSteps(parameters, thetaFull, objectiveFunction, borders, iPar, s, options)
    
    global llhHistory;
    
    % Initialize everything
    y = [];
    theta = (thetaFull(end, :))';
    I1 = (1 : iPar-1)';
    I2 = (iPar+1 : length(theta))';
    I = [I1; I2];
    stepCounter = 1;
    
    % Initialize direction
    dtheta = zeros(length(thetaFull(end, :)),1);
    dtheta(iPar) = s * options.options_getNextPoint.guess;
        
    % dtheta = (thetaFull(end, :) - thetaFull(end-10, :))';
    borders(iPar,:) = [options.P.min(iPar), options.P.max(iPar)];
    logPost = parameters.MS.logPost(options.MAP_index);
    logPost_max = parameters.MS.logPost(1);
    negLogPost = setObjectiveWrapper(objectiveFunction, options, 'negative log-posterior', [], [], true, true);
    
    % Sequential update
    while (options.P.min(iPar) < theta(iPar)) && (theta(iPar) < options.P.max(iPar)) && ...
            (logPost >= (log(options.R_min) + logPost_max) && ...
            stepCounter < 5)
    
        % Proposal of next profile point
        [theta_next,~] = ...
            getNextProfilePoint(theta, ...
            borders(:,1), ...
            borders(:,2), ...
            dtheta/abs(dtheta(iPar)), ...
            abs(dtheta(iPar)), ...
            options.options_getNextPoint.min, ...
            options.options_getNextPoint.max, ...
            options.options_getNextPoint.update, ...
            -( log(1-options.dR_max) + options.dJ * (logPost-logPost_max) + logPost ), ...
            negLogPost,...
            parameters.constraints, ...
            options.options_getNextPoint.mode, ...
            iPar);
        negLogPostReduced = setObjectiveWrapper(objectiveFunction, options, 'negative log-posterior', iPar, theta_next(iPar), true, true);
        
        % Construction of reduced linear constraints
        [A,b,Aeq,beq] = getConstraints(theta, parameters, I);
        
        if (~isfield(options.profileReoptimizationOptions, 'HessFcn') ...
            || isempty(options.profileReoptimizationOptions.HessFcn))
            % this only works for box-constraints at the moment
            options.profileReoptimizationOptions.HessFcn = @(varargin) HessianWrap(negLogPostReduced, varargin);
        end 
        % Optimization
        [theta_I_opt, L, ~, ~, ~, newGL, newHL] = ...
            fmincon(negLogPostReduced, ...
            theta_next(I),...
            A  ,b  ,... % linear inequality constraints
            Aeq,beq,... % linear equality constraints
            parameters.min(I),...   % lower bound
            parameters.max(I),...   % upper bound
            [],...
            options.profileReoptimizationOptions);    % options
        
        % Restore full vector and determine update direction
        logPost = -L;
        dtheta = [theta_I_opt(I1); theta_next(iPar); theta_I_opt(I2-1)] - theta;
        theta = theta + dtheta;
        
        llhHistory = [llhHistory, logPost];
        R = exp(-L - parameters.MS.logPost(1));
        if (strcmp(options.mode, 'text'))
            fprintf('\n  |  %11.8f | %11.7f | %7.5f |', ...
                theta(iPar), sqrt(sum(newGL.^2)), R);
        end
        y = [y, theta];
        stepCounter = stepCounter + 1;
    end
    
    if ~strcmp(options.solver.hessian, 'user-supplied')
        tmpHL = [newHL(1:iPar-1,1:iPar-1), zeros(iPar-1,1), newHL(1:iPar-1,iPar:end);...
            zeros(1,length(theta)); newHL(iPar:end,1:iPar-1), zeros(length(theta)-iPar,1), newHL(iPar:end,iPar:end)];
        tmpHL(iPar,iPar) = inf;
        tmpGL = [newGL(1:iPar-1); inf; newGL(iPar:end)];
        approximateHessian(theta, -tmpGL, tmpHL, options.solver.hessian, 'reinit');
    end
    y = y';
    
end


%% SingleParameter is a support function for the profile integration
%  Provides function handles for the parameter function its
%  gradient and hessian for a single parameter profile
% 
% USAGE:
% =====
% [...] = SingleParameter(theta, index)
%     
% INPUTS:
% =======
% theta ... parameter values
% index ... index of parameter for which the profile should be computed
% 
% OUTPUTS:
% ========
% g   ... parameter value
% dg  ... gradient
% ddg ... hessian
% 
% 2015/11/20 Sabrina Hross
% 2017/02/20 Paul Stapor

function [varargout] = SingleParameter(theta, index)
        
    switch nargout
        case 1
          varargout{1} = theta(index, :);   
        case 2
            varargout{1} = theta(index, :);

            dg = zeros(length(theta),1); dg(index) = 1;
            varargout{2} = dg;
        case 3
            varargout{1} = theta(index,:);
            dg = zeros(length(theta),1); dg(index) = 1;
            varargout{2} = dg;
            varargout{3} = zeros(length(theta),length(theta));
    end
    
end


%% getEndProfile is a support function for the profile integration
%   and is called in integrateProfile. It gives the Rootfunction for CVODES
%   to end the profile calculation if the profile falls below the threshold
%   before the parameter bounds are reached.
%
% USAGE:
% ======
% function [R,FLAG] = getEndProfile(theta)
%
% INPUTS:
% =======
% theta ... parameter   
%
% Outputs:
% ========
% CVODE
% R ... Root function
% FLAG ... =0 if successful ~=0 if failed
% ODE15s
% R ... Root function
% isterminate ... =1 to terminate at root
% direction ... =0 find zeros independent of direction
%
% 2015/11/19 Sabrina Hross
% 2017/02/20 Paul Stapor

function [varargout] = getEndProfile(t, s, y, ind, borders, negLogPost, options, logPostMax)

    L = negLogPost(y);
    R = logPostMax - log(options.R_min) + L;

    if strcmp(options.solver.type, 'CVODE')
        if (t < borders(ind, 2)) && (t > borders(ind, 1))
            varargout{1} = R;
        else
            % display('stopped at parameter boundary')
            varargout{1} = 0;
        end
        varargout{2} = 0; % flag
    else
        withinBorders = false;
        if (s == 1)
            if (t < borders(ind, 2)) && (t > borders(ind, 1))
                withinBorders = true;
            end
        else
            if (t < s*borders(ind, 1)) && (t > s*borders(ind, 2))
                withinBorders = true;
            end
        end
        
        if withinBorders
            varargout{1} = R;
        else
            % display('stopped at parameter boundary')
            varargout{1} = 0;
        end
        varargout{2} = 1; % isterminal
        varargout{3} = 0; % locate all zeros
    end
    
end


function [dth, flag, new_Data] = getRhsRed(~, s, y, ind, borders, negLogPost, parameterFunction, options)
    
    % set parameters
    npar = length(y);
    flag = 0;
    
    global ObjFuncCounter;
    ObjFuncCounter = ObjFuncCounter + 1;
    
    switch options.solver.hessian
        case 'user-supplied'
            [~, GL, HL] = negLogPost(y);
        case {'bfgs', 'sr1', 'dfp'}
            [~, GL] = negLogPost(y);
            HL = approximateHessian(y, -GL, [], options.solver.hessian, 'read');
        otherwise
            error('Unknown type of Hessian computation.');
    end
    
    if (sum(sum(isnan(HL)))>0) || sum(isnan(GL))>0 || (sum(sum(isinf(HL)))>0) || sum(isinf(GL))>0
        disp('Warning: Undefined model output')
        flag = -1;
        dth = nan(size(y));
        new_Data = [];
        return;
    end

    % calculate hessian of parameter function
    [~, GG, ~] = parameterFunction(y, ind);

    % right handside of ODE
    try    
        % Reduce linear system by implicit funtion theorem
        A1 = [HL(1 : ind-1, :); HL(ind+1 : npar, :); s*GG'];
        b1 = [-GL(1 : ind-1) * options.solver.gamma; ...
             -GL(ind+1 : npar) * options.solver.gamma; ...
             1];
        A2 = [A1(1:end-1, 1:ind-1), A1(1:end-1, ind+1:end)];
        b2 = b1(1:end-1) - s * A1(1:end-1, ind);
        
        % Check for invertibility of the RHS
        if (rcond(A2) < options.solver.minCond) || isnan(rcond(A2))
            dth1 = pinv(A2, options.solver.eps) * b2;
        else
            dth1 = A2 \ b2;
        end
        dth = [dth1(1:ind-1); s; dth1(ind:end)];
    catch
        dth = zeros(npar + 1, 1);
        dth(ind) = s;
    end
    
    % Check, if parameter bounds are violated
    lRebuild = 0;
    for i = 1 : npar
        if (i ~= ind)
            if (y(i) <= borders(i, 1))
                if(dth(i) < 0)
                    GL(i) = 0;
                    HL(i,:) = 0;
                    HL(:,i) = 0;
                    HL(i,i) = -1;
                    lRebuild = 1;
                end
            elseif (y(i) >= borders(i, 2))
                if(dth(i) > 0)
                    GL(i) = 0;
                    HL(i,:) = 0;
                    HL(:,i) = 0;
                    HL(i,i) = -1;
                    lRebuild = 1;
                end
            end
        end
    end
    
    if (lRebuild == 1)
        try    
            % Reduce linear system by implicit funtion theorem
            A1 = [HL(1 : ind-1, :); HL(ind+1 : npar, :); s*GG'];
            b1 = [-GL(1 : ind-1) * options.solver.gamma; ...
                 -GL(ind+1 : npar) * options.solver.gamma; ...
                 1];
            A2 = [A1(1:end-1, 1:ind-1), A1(1:end-1, ind+1:end)];
            b2 = b1(1:end-1) - s * A1(1:end-1, ind);

            % Check for invertibility of the RHS
            if (rcond(A2) < options.solver.minCond) || isnan(rcond(A2))
                dth1 = pinv(A2, options.solver.eps) * b2;
            else
                dth1 = A2 \ b2;
            end
            dth = [dth1(1:ind-1); s; dth1(ind:end)];
        catch
            dth = zeros(npar + 1, 1);
            dth(ind) = s;
        end
    end
   
    new_Data = [];
end


function hessian = approximateHessian(theta, grad, hess, method, flag)
    
    persistent lastTheta;
    persistent lastGrad;
    persistent lastHess;
    
    if strcmp(flag, 'init')
        hessian = [];
        
        % Replace old by new values for next call
        lastGrad = grad;
        lastHess = hess;
        lastTheta = theta;
    elseif strcmp(flag, 'reinit')
        % Replace old by new values for next call
        lastTheta = theta;    
        [~,ind] = max(max(hess));
        ind = ind(1);
        lastHess = [hess(1:ind-1,1:ind-1), lastHess(1:ind-1,ind), hess(1:ind-1,ind+1:end); ...
            lastHess(ind,:); hess(ind+1:end,1:ind-1), lastHess(ind+1:end,ind), hess(ind+1:end,ind+1:end)];
        lastGrad = [grad(1:ind-1); lastGrad(ind); grad(ind+1:end)];

    elseif strcmp(flag, 'write')
        if (theta == lastTheta)
            hessian = lastHess;
        else
            switch method
                case 'bfgs'
                    delTheta = theta - lastTheta;
                    delGrad = grad - lastGrad;
                    u = delGrad * delGrad' / (delGrad' * delTheta);
                    v = lastHess * (delTheta * delTheta') * lastHess / (delTheta' * lastHess * delTheta);
                    hessian = lastHess + u - v;
                case 'sr1'
                    delTheta = theta - lastTheta;
                    delGrad = grad - lastGrad;
                    u = delGrad - lastHess * delTheta;
                    v = u * u' / (u' * delTheta);
                    hessian = lastHess + v;
                case 'dfp'
                    delTheta = theta - lastTheta;
                    delGrad = grad - lastGrad;
                    gamma = 1 / (delGrad' * delTheta);
                    u1 = eye(length(theta)) - gamma * (delGrad * delTheta');
                    u2 = eye(length(theta)) - gamma * (delTheta * delGrad');
                    v = gamma * (delGrad * delGrad'); 
                    hessian = u1 * lastHess * u2 + v;
            end
            
            % Replace old by new values for next call
            lastGrad = grad;
            lastHess = hessian;
            lastTheta = theta;
        end
        
    elseif strcmp(flag, 'read')
        hessian = lastHess;
    end
    
end


%% getRhsDAE is a support function for the profile integration
%   and is called in integrateProfile. It determines the right hand side of
%   of the problem in the DAE formulation.
%
% USAGE:
% ======
% function [dth,flag,new_Data] = getRhsDAE(u, s, data)
%
% INPUTS:
% =======
% u ... current state (not used but given by IDAS)
% s ... step direction (not used by IDAS, fixed input)  
% y ... current parameter set 
% yp ... parameter slope
% options ... options for profile integration
%   .solver    ... solver options determining gradient and hessian
%   calculation
%   .paramFunc ... parameter function
%
% Outputs:
% ========
% dth ... parameter proposal
% flag ... needed by IDAS
% new_Data ... needed by IDAS
%
% 2013/11/16 Sabrina Hock
% 2016/02/06 Sabrina Hross

function [dth, flag] = getRhsDAE(~, s, y, yp, ind, negLogPost, options)

    % set parameters
    npar = length(y);
    flag = 0;
    
    [~,GL] = negLogPost(y);

    if (sum(isnan(GL)) > 0 || sum(isinf(GL)) > 0)
        disp('Warning: Undefined model output')
        flag = -1;
    end

    b = [-GL(1 : ind-1) * s * options.solver.gamma; ...
             -GL(ind+1 : npar) * s * options.solver.gamma; ...
             1];

    switch options.solver.type
        case 'ode15sDAE'
            dth = b;        
        case 'IDAS'
            Mt = getMassmatrixDAE(1,s,y,options);
            dth = Mt*yp-b;
    end
end


%% getMassmatrixDAE is a support function for the profile integration
%   and is called by integrateProfileDAE. It determines the Massmatrix of
%   of the problem in the DAE formulation.
%
% USAGE:
% ======
% function [dth,flag,new_Data] = getMassmatrixDAE(u, y, data)
%
% INPUTS:
% =======
% u ... current state (not used but given by IDAS) 
% y ... current parameter set 
% options ... options for profile integration
%   .solver    ... solver options determining gradient and hessian
%   calculation
%   .paramFunc ... parameter function
%
% Outputs:
% ========
% MT ... Massmatrix
%
% 2013/11/16 Sabrina Hock
% 2016/02/06 Sabrina Hross
% 2016/11/21 Paul Stapor

function Mt = getMassmatrixDAE(c ,s, y, ind, negLogPost, parameterFunction)

    % set parameters
    npar = length(y);
    
    [~, GL, HL] = negLogPost(y);
    
    if (sum(sum(isnan(HL)))>0) || (sum(sum(isinf(HL)))>0) 
        disp('Warning: Undefined model output')
    end

    % calculate hessian of parameter function
    [~,GG,~] = parameterFunction(th, ind);
    GG=s*GG;

    A1 = [HL(1 : ind-1, :); HL(ind+1 : npar, :); s*GG'];
         
    %Massmatrix
    Mt = A1;
    
    GL1 = GL; 
    GL1(ind) = 0;
    if (isempty(c))
        display(['Jacobian Evaluation around theta = ' y']);
    else
        display(['Running axis: ' num2str(s*c) '   Optimality: ', num2str(sqrt(GL1' * GL1)), '    condition number: ', num2str(rcond(A1))]);
    end
end
```
