## Supplementary material for "Mechanistic description of spatial processes using integrative modelling of noise-corrupted imaging data": getParProfilesByOptimization.m

```
function [parameters,fh] = getParProfilesByOptimization(parameters, objective_function, options, varargin)
% getParProfilesByOptimization.m calculates the profiles likelihoods for the model 
% parameters, starting from the maximum a posteriori estimate. This
% calculation is done by fixing the i-th parameter and repeatedly
% reoptimizing the likelihood/posterior estimate (for all i). The initial 
% guess for the next reoptimization point is computed by extrapolation from
% the previous points to ensure a quick optimization.
%
% Note: This function can exploit up to (n_theta + 1) workers when running
% in 'parallel' mode.
%
% USAGE:
% [...] = getParameterProfiles(parameters, objective_function)
% [...] = getParameterProfiles(parameters, objective_function, options)
% [parameters, fh] = getParameterProfiles(...)
%
% getParProfilesByOptimization() uses the following PestoOptions members:
%  * PestoOptions::calc_profiles
%  * PestoOptions::comp_type
%  * PestoOptions::dJ
%  * PestoOptions::dR_max
%  * PestoOptions::fh
%  * PestoOptions::MAP_index
%  * PestoOptions::mode
%  * PestoOptions::obj_type
%  * PestoOptions::options_getNextPoint .guess .min .max .update .mode
%  * PestoOptions::parameter_index
%  * PestoOptions::profile_method
%  * PestoOptions::profileReoptimizationOptions
%  * PestoOptions::plot_options
%  * PestoOptions::R_min
%  * PestoOptions::save
%
% Parameters:
%   parameters: parameter struct
%   objective_function: objective function to be optimized. 
%       This function should accept one input, the parameter vector.
%   varargin:
%     options: A PestoOptions object holding various options for the 
%         algorithm.
%
% Required fields of parameters:
%   number: Number of parameters
%   min: Lower bound for each parameter
%   max: upper bound for each parameter
%   name = {'name1', ...}: names of the parameters
%   MS: results of global optimization, obtained using for instance 
%       the routine 'getMultiStarts.m'. MS has to contain at least
%     * par: sorted list n_theta x n_starts of parameter estimates.
%          The first entry is assumed to be the best one.
%     * logPost: sorted list n_starts x 1 of of log-posterior values
%          corresponding to the parameters listed in .par.
%     * hessian: Hessian matrix (or approximation) at the optimal point
%
% Return values:
%   properties: updated parameter struct
%   fh: figure handle
%
% Generated fields of parameters:
%   P(i): profile for i-th parameter
%     * par: MAPs along profile
%     * logPost: maximum log-posterior along profile
%     * R: ratio
%
% History:
% * 2012/05/16 Jan Hasenauer
% * 2014/06/12 Jan Hasenauer
% * 2016/10/04 Daniel Weindl
% * 2016/10/12 Paul Stapor

    %% No check of inputs (except figure), already done in getParameterProfiles

    % Depending on parallel computation mode, the figure will be updated
    if (nargin >= 4)
        fh = varargin{1};
        options.fh = fh;
    else
        fh = [];
    end
    
    %% Check for fixed parameters
    if ~isempty(options.fixedParameters)
        error('Fixed parameters are currently not supported by getParProfilesByOptimization.');
    end

    %% Profile calculation
    if strcmp(options.comp_type,'sequential')
        for iPar = options.profile_optim_index
            % Define the negative log-posterior function
            % (fmincon needs the neagtive log posterior for optimization)
            % with the profile parameter fixed
            parameters = optimizeProfileForParameterI(parameters, objective_function, iPar, options, fh);
        end

    elseif strcmp(options.comp_type,'parallel')
        parfor iPar = options.profile_optim_index
            % Define the negative log-posterior function
            % (fmincon needs the neagtive log posterior for optimization)
            % with the profile parameter fixed
            optimizeProfileForParameterI(parameters, objective_function, iPar, options, fh);
        end

        % Output
        if strcmp(options.profile_method, 'optimization')
            switch options.mode
                case 'visual', fh = plotParameterProfiles(parameters,'1D',fh,options.parameter_index,options.plot_options);
                case 'text' % no output
                case 'silent' % no output
            end
        end
    end

% %% REOPTIMIZE PROFILE FROM THE BORDER
% if strcmp(options.reoptimize,'true')
% disp('');
% disp('Re-optimization of profile');
% % Loop: Parameters
% for i = options.parameter_index
%     disp('');
%     % Index set
%     I1 = [1:i-1]';
%     I2 = [i+1:parameters.number]';
%     I  = [I1;I2];
%     % Likelihood function option
%     options.logPost_options.grad_ind = I(:);
%     options.logPost_options.sign = 'negative';
% 
%     %% COMPUTE OPTIMUM FOR IN-/DECREASING THETA
%     for s = [-1,1]
%         % Find set
%         if s == -1
%             ind = find(parameters.P(i).par(i,:) < parameters.MS.par(i));
%             ind = ind(2:end);
%         else
%             ind = find(parameters.P(i).par(i,:) > parameters.MS.par(i));
%             ind = ind(end-1:-1:1);
%         end
%  
%         % Loop: Index points
%         for k = ind
%             % Starting point
%             theta_next = parameters.P(i).par(:,k+s);
%             theta_next(i) = parameters.P(i).par(i,k);
%             theta_i = theta_next(i);
%             
%             % Modification of options struct
%             options_fmincon = options.fmincon;
%             if isfield(options.fmincon,'TypicalX')
%                 if ~isempty(options.fmincon.TypicalX)
%                     options_fmincon.TypicalX = options_fmincon.TypicalX(I);
%                 end
%             end
%             if isfield(options.fmincon,'FinDiffRelStep')
%                 if ~isempty(options.fmincon.FinDiffRelStep)
%                     options_fmincon.FinDiffRelStep = options_fmincon.FinDiffRelStep(I);
%                 end
%             end
%             
%             % Linear constraints
%             if ~isempty(parameters.constraints.A)
%                 A = parameters.constraints.A(:,I);
%                 b = parameters.constraints.b - 10^-10 - parameters.constraints.A(:,i)*theta_i;
%             else
%                 A = [];
%                 b = [];
%             end
%             if ~isempty(parameters.constraints.Aeq)
%                 Aeq = parameters.constraints.Aeq(:,I) - parameters.constraints.Aeq(:,i)*theta_i;
%                 beq = parameters.constraints.beq;
%             else
%                 Aeq = [];
%                 beq = [];
%             end
%             
%             % Optimize
%             [theta_next,J_opt] = ...
%                 fmincon(@(theta_I) objective_function([theta_I(I1);theta_next(i);theta_I(I2-1)],options.logPost_options),...     % negative log-likelihood function
%                                     theta_next(I),...        % initial parameter
%                                     A  ,b  ,...             % linear inequality constraints
%                                     Aeq,beq,...             % linear equality constraints
%                                     parameters.min(I),...   % lower bound
%                                     parameters.max(I),...   % upper bound
%                                     [],options_fmincon);    % options
%             logPost = -J_opt;
%             
%             % Assignment
%             if -J_opt > parameters.P(i).logPost(k);
%                 parameters.P(i).par(I,k) = theta_next;
%                 parameters.P(i).logPost(k) = logPost;
%                 parameters.P(i).R(k) = exp(logPost - parameters.MS.logPost);
%             end
%             
%             % Update plot
%             if strcmp(options.plot,'true')
%                 fh = plotP(parameters,fh,options.parameter_index,options.plot_options);
%             end
%             
%             % Output command line
%             disp([num2str(i,'%d') '-th P: point ' num2str(k,'%d') ...
%                 ' -> ' num2str(ind(end)-s,'%d')]);
%         end
%     end
%     disp('');
% end
% end
% 
%  

end

function [parameters] = optimizeProfileForParameterI(parameters, objective_function, iPar, options, fh)

    % Initialization
    logPost_max = parameters.MS.logPost(1);
    P_par = parameters.MS.par(:,options.MAP_index);
    P_logPost = parameters.MS.logPost(options.MAP_index);
    P_R = exp(parameters.MS.logPost(options.MAP_index)-parameters.MS.logPost(1));
    negLogPost = setObjectiveWrapper(objective_function, options, 'negative log-posterior', [], [], true, true);
    
    % Construction of index set
    I1 = (1 : iPar-1)';
    I2 = (iPar+1 : parameters.number)';
    I  = [I1; I2];
 
    % Compute profile for in- and decreasing theta_i
    for s = [-1,1]
        % Starting point
        theta  = parameters.MS.par(:, options.MAP_index);
        logPost = parameters.MS.logPost(options.MAP_index);

        % Lower and upper bounds for profiles
        theta_min = [parameters.min(I1); options.P.min(iPar); parameters.min(I2)];
        theta_max = [parameters.max(I1); options.P.max(iPar); parameters.max(I2)];

        % Initialize direction
        dtheta = zeros(parameters.number,1);
        dtheta(iPar) = s*options.options_getNextPoint.guess;

        % Get the computation time
        startTimeProfile = cputime;
        stepCount = 0;

        % Sequential update
        while (options.P.min(iPar) < theta(iPar)) && (theta(iPar) < options.P.max(iPar)) && ...
                (logPost >= (log(options.R_min) + parameters.MS.logPost(1)))

            % Proposal of next profile point
            [theta_next,J_exp] = ...
                getNextProfilePoint(theta,theta_min,theta_max,dtheta/abs(dtheta(iPar)),...
                abs(dtheta(iPar)),options.options_getNextPoint.min,options.options_getNextPoint.max,options.options_getNextPoint.update,...
                -(log(1-options.dR_max)+options.dJ*(logPost-logPost_max)+logPost),negLogPost,...
                parameters.constraints,options.options_getNextPoint.mode,iPar);
            negLogPostReduced = setObjectiveWrapper(objective_function, options, 'negative log-posterior', iPar, theta_next(iPar), true, true);
            
            % Check, if Hessian should be used and if a Hessian function was set,
            % otherwise use the third output of the objective function instead
            if (strcmp(options.profileReoptimizationOptions.Hessian, 'on'))
                if (~isfield(options.profileReoptimizationOptions, 'HessFcn') ...
                        || isempty(options.profileReoptimizationOptions.HessFcn))
                    % this only works for box-constraints at the moment
                    options.profileReoptimizationOptions.HessFcn = @(varargin) HessianWrap(negLogPostReduced, varargin);
                end
            end
            
            % Construction of reduced linear constraints
            [A,b,Aeq,beq] = getConstraints(theta,parameters,I);
            
            % Optimization
            try
                [theta_I_opt,J_opt] = ...
                    fmincon(negLogPostReduced,... % negative log-posterior function
                    theta_next(I),...
                    A  ,b  ,... % linear inequality constraints
                    Aeq,beq,... % linear equality constraints
                    parameters.min(I),...   % lower bound
                    parameters.max(I),...   % upper bound
                    [],options.profileReoptimizationOptions);    % options
                stepCount = stepCount + 1;
            catch errMsg
                theta_I_opt = theta_next;
                J_opt = inf;
            end

            % Restore full vector and determine update direction
            logPost = -J_opt;
            dtheta = [theta_I_opt(I1);theta_next(iPar);theta_I_opt(I2-1)] - theta;
            theta = theta + dtheta;

            % Sorting
            switch s
                case -1
                    P_par = [theta,P_par];
                    P_logPost = [logPost,P_logPost];
                    P_R = [exp(logPost - parameters.MS.logPost(1)),P_R];
                case +1
                    P_par = [P_par,theta];
                    P_logPost = [P_logPost,logPost];
                    P_R = [P_R,exp(logPost - parameters.MS.logPost(1))];
            end

            % Assignment
            parameters.P(iPar).par = P_par;
            parameters.P(iPar).logPost = P_logPost;
            parameters.P(iPar).R = P_R;

            % Save
            if options.save
                dlmwrite([options.foldername '/P' num2str(iPar,'%d') '__par.csv'],P_par,'delimiter',',','precision',12);
                dlmwrite([options.foldername '/P' num2str(iPar,'%d') '__logPost.csv'],P_logPost,'delimiter',',','precision',12);
                dlmwrite([options.foldername '/P' num2str(iPar,'%d') '__R.csv'],P_R,'delimiter',',','precision',12);
            end

            % Output
            if ~strcmp(options.comp_type,'parallel')
                switch(iPar)
                    case 1
                        ordstr = 'st';
                    case 2
                        ordstr = 'nd';
                    case 3
                        ordstr = 'rd';
                    otherwise
                        ordstr = '-th';
                end
                str = [num2str(iPar,'%d') ordstr ' P: point ' num2str(length(parameters.P(iPar).R)-1,'%d') ', R = ' ...
                    num2str(exp(- J_opt - parameters.MS.logPost(1)),'%.3e') ' (optimized) / '...
                    num2str(exp(- J_exp - parameters.MS.logPost(1)),'%.3e') ' (predicted)'];
                switch options.mode
                    case 'visual', fh = plotParameterProfiles(parameters,'1D',fh,options.parameter_index,options.plot_options);
                    case 'text', disp(str);
                    case 'silent' % no output
                end
            end
        end

        parameters.P(iPar).t_cpu(round((s+3)/2)) = cputime - startTimeProfile;
        parameters.P(iPar).optSteps(round((s+3)/2)) = stepCount;
        parameters.P(iPar).intSteps(round((s+3)/2)) = 0;
        parameters.P(iPar).reOptSteps(round((s+3)/2)) = 0;
    end

end
```
