## Supplementary material for "Mechanistic description of spatial processes using integrative modelling of noise-corrupted imaging data": getStepSizeFD.m

```
function fdStepSize = getStepSizeFD(theta, objectiveFunction, mode)

% This function computes the optimal step size for finite differences.
%
% Parameters:
%   theta: parameter vector
%   objectiveFunction: the original objective function
%   mode: double, specifying if the output of the original objective func.
%           1: only objective value is passed
%           2: gradients are computed
%
% Return values:
%   fdStepSize: vector of step sizes

    % Initialization
    nPar = length(theta);
    fdStepSize = zeros(nPar, 1);

    switch mode
        case 1
        % The original objective funcion can only compute the objective,
        % step sizes for the gradient are needed
        
            for j = 1 : nPar
                delta = zeros(nPar,1);
                gradients = zeros(1,7);
                qualityVector = zeros(1,7);

                % Compute gradient for different step sizes
                for k = 3 : 9
                    delta(j) = 10^(-k);
                    Jplus = objectiveFunction(theta + delta);
                    Jminus = objectiveFunction(theta - delta);
                    gradients(k-2) = 0.5 * 10^k * (Jplus - Jminus);
                end

                % Compare gradients for different step sizes
                for k = 2 : 6
                    qualityVector(k) = 0.5 * (abs(gradients(k-1) - gradients(k)) ...
                        + abs(gradients(k+1) - gradients(k)));
                end
                qualityVector(1) = abs(gradients(1) - gradients(2));
                qualityVector(7) = abs(gradients(5) - gradients(6));

                % Choose step sizes with best stability properties
                [maxQual, ind] = min(qualityVector);
                fdStepSize(j) = 10^(-ind-3);

                if (maxQual > 0.1)
                    warning('Could not find a good step size for finite differences. Please check the accuracy of your objetive function.');
                end
            end
        
        case 2
        % The original objective funcion can compute the gradient, step 
        % sizes for the Hessian are needed
        
            for j = 1 : nPar
                delta = zeros(nPar,1);
                hessians = zeros(nPar,7);
                qualityVector = zeros(1,7);

                % Compute gradient for different step sizes
                for k = 3 : 9
                    delta(j) = 10^(-k);
                    [~,Gplus] = objectiveFunction(theta + delta);
                    [~,Gminus] = objectiveFunction(theta - delta);
                    hessians(:,k-2) = 0.5 * 10^k * (Gplus - Gminus);
                end

                % Compare gradients for different step sizes
                for k = 2 : 6
                    qualityVector(k) = 0.5 * (sum(abs(hessians(:,k-1) - hessians(:,k)).^2) ...
                        + sum(abs(hessians(:,k+1) - hessians(:,k)).^2));
                end
                qualityVector(1) = sum(abs(hessians(:,1) - hessians(:,2)));
                qualityVector(7) = sum(abs(hessians(:,5) - hessians(:,6)));

                % Choose step sizes with best stability properties
                [maxQual, ind] = min(qualityVector);
                fdStepSize(j) = 10^(-ind-3);
                
                if (maxQual > 0.1 * nPar)
                    warning('Could not find a good step size for finite differences. Please check the accuracy of your objetive function.');
                end
            end
    end
    
end
```
