## Supplementary material for "Mechanistic description of spatial processes using integrative modelling of noise-corrupted imaging data": handleOptionArgument.m

```
function [ options ] = handleOptionArgument( options )
    %HANDLEOPTIONARGUMENT Check argument type or autocreate PestoOptions
    %from struct (deprecated)
    if isstruct(options)
        warning('Options argument is not of type PestoOptions: Initializing PestoOption instance from user-provided struct. Make sure all options are set correctly in PestoOptions(myOptionsStruct).');
        options = PestoOptions(options);
    elseif ~isa(options, 'PestoOptions')
        error('Options argument is neither of type PestoOptions nor struct.')
    end
end
```
