## Supplementary figures and images for "Mechanistic description of spatial processes using integrative modelling of noise-corrupted imaging data"

### CCL21-1.tif

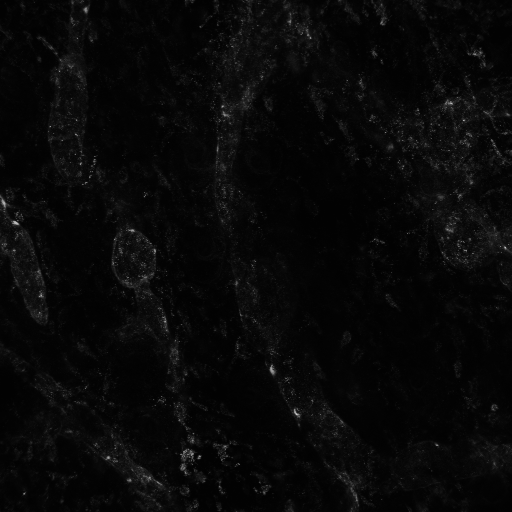

### CCL21-2.tif

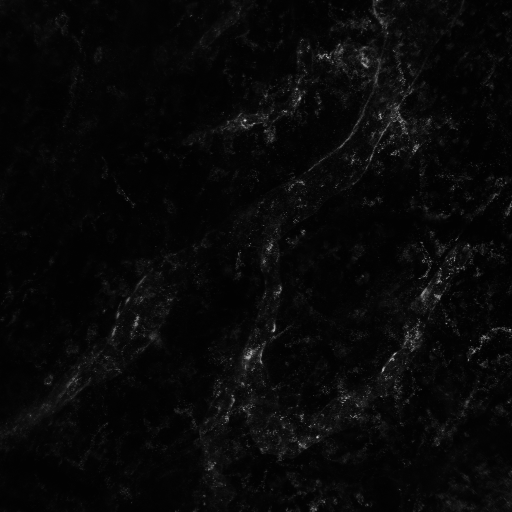

### CCL21-3.tif

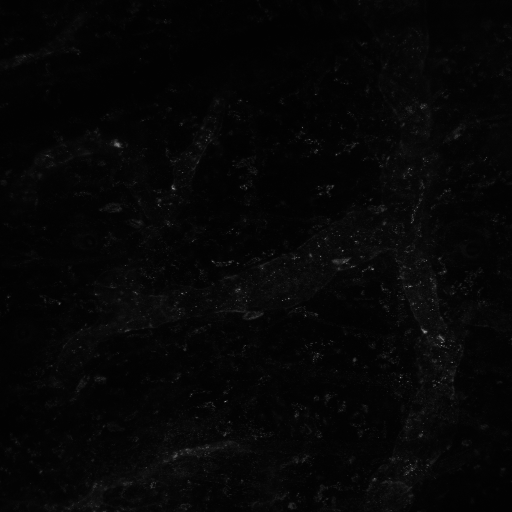

### CCL21-4.tif

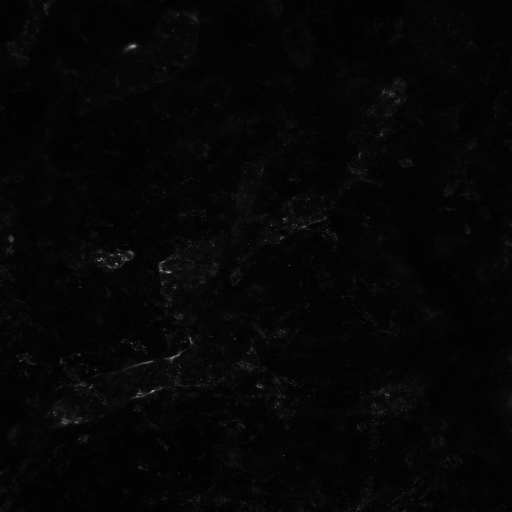

### CCL21-12.tif

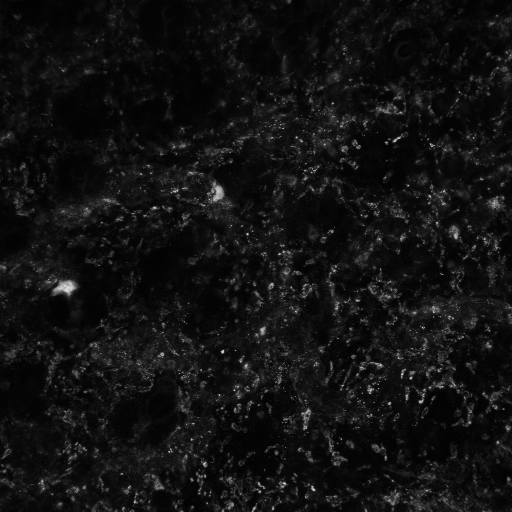

### CCL21-13.tif

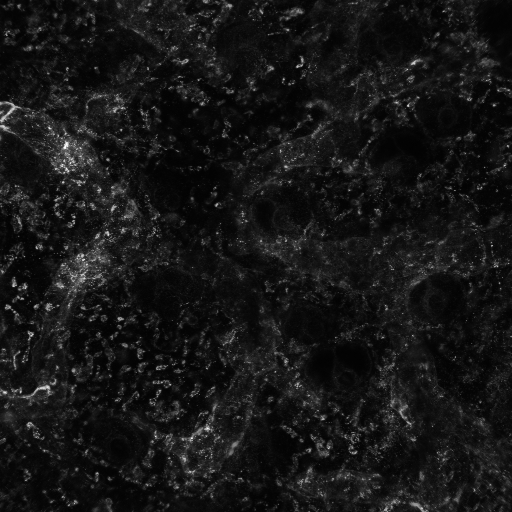

### CCL21-14.tif

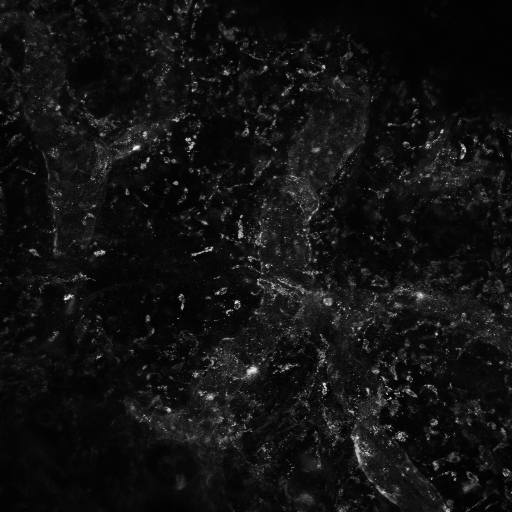

### CCL21-15.tif

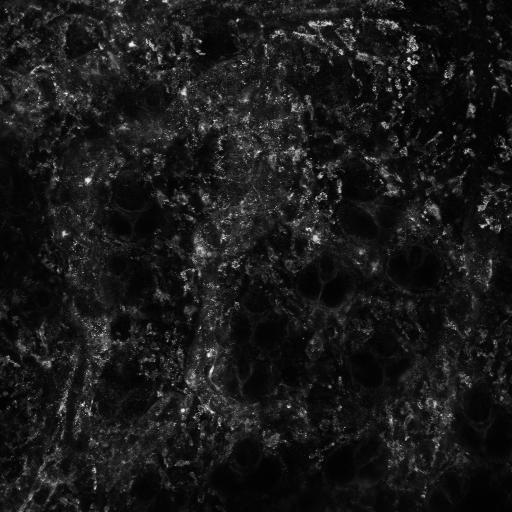

### CCL21-16.tif

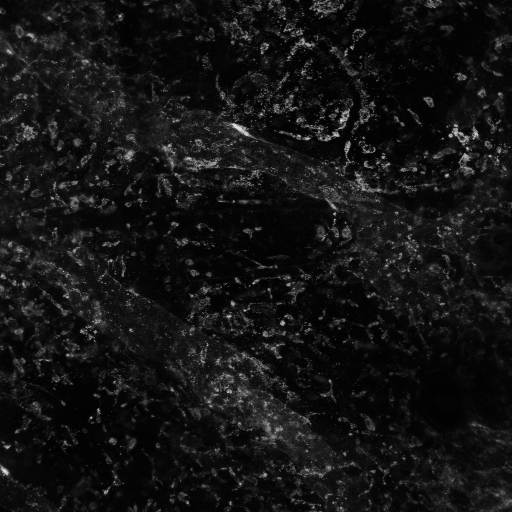

### fig_fit.jpg

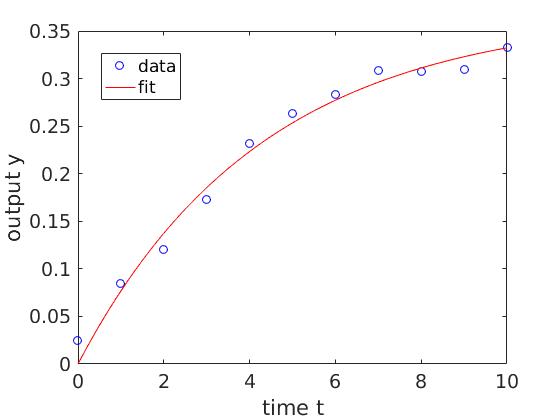

### fig_par_confint.jpg

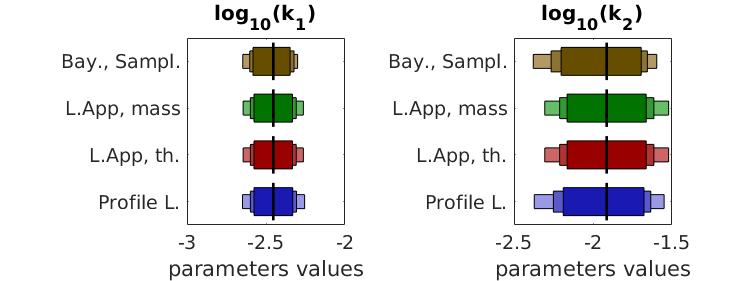

### fig_par_samples2d.jpg

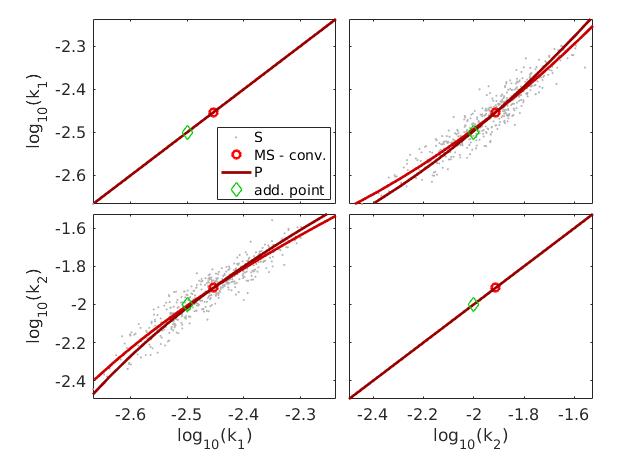

### fig_par_samples.jpg

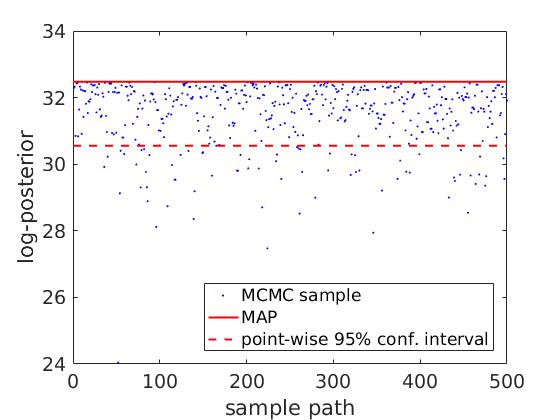

### fig_prop_samples.jpg

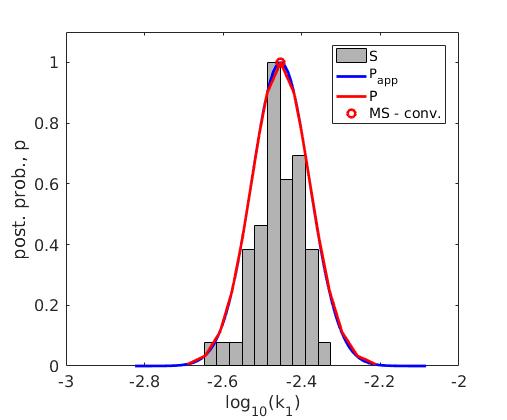

### freezeColors_pub.png

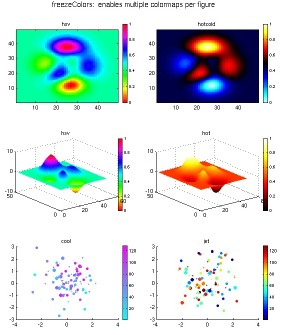

### freezeColors_pub_01.png

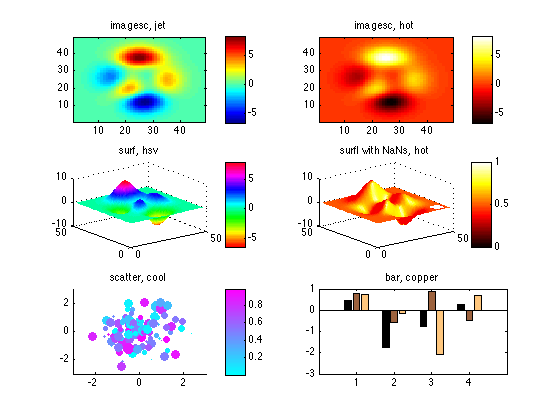

### freezeColors_pub_02.png

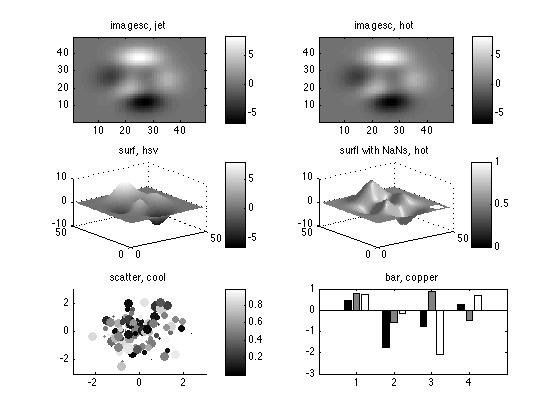

### LVMask-CCL21-1.tif

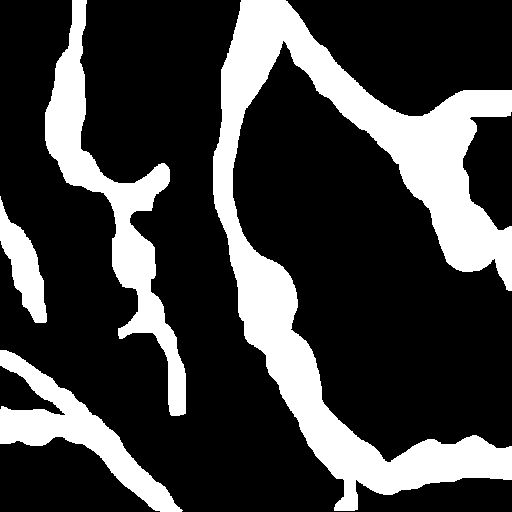

### LVMask-CCL21-2.tif

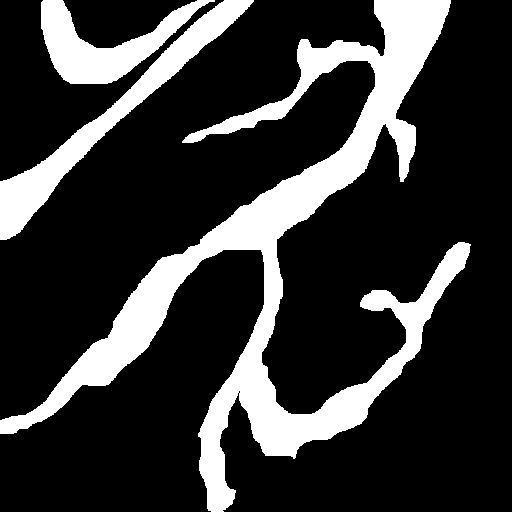

### LVMask-CCL21-3.tif

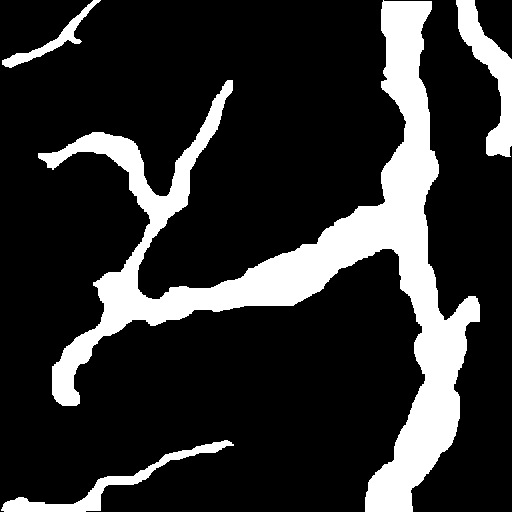

### LVMask-CCL21-4.tif

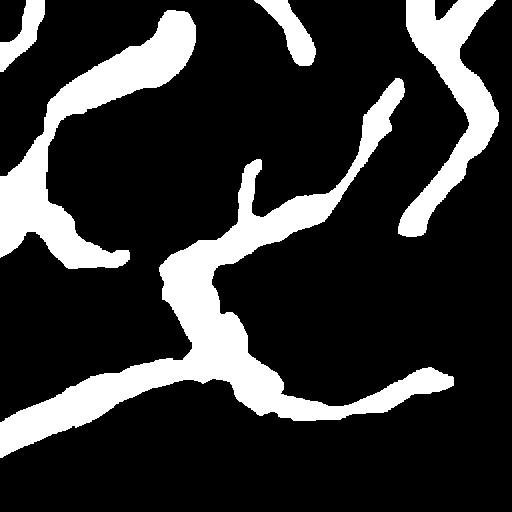

### LVMask-CCL21-12.tif

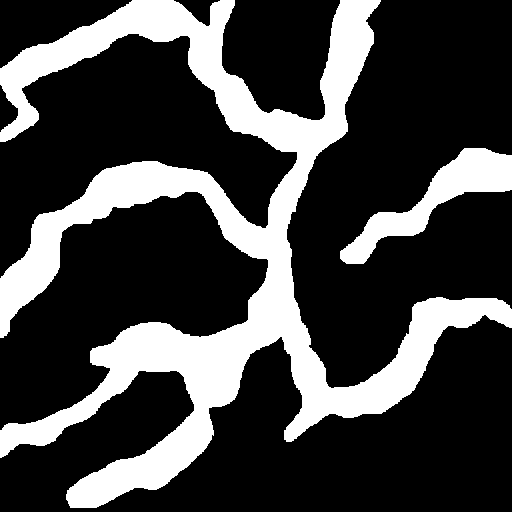

### LVMask-CCL21-13.tif

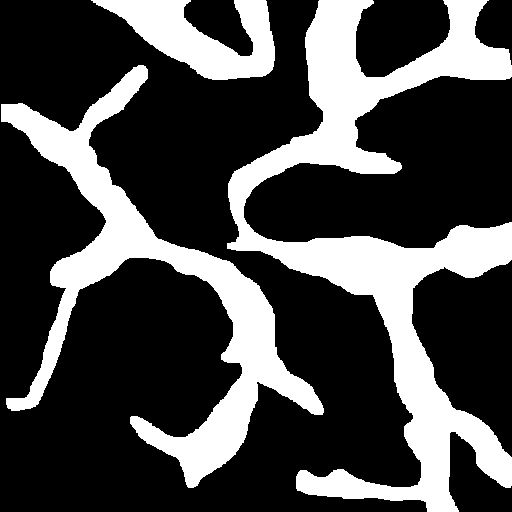

### LVMask-CCL21-14.tif

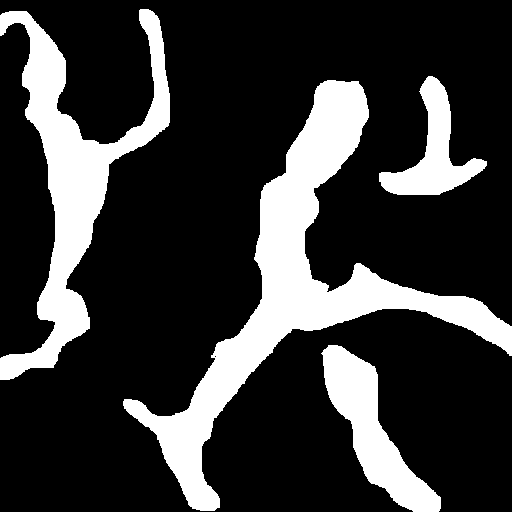

### LVMask-CCL21-15.tif

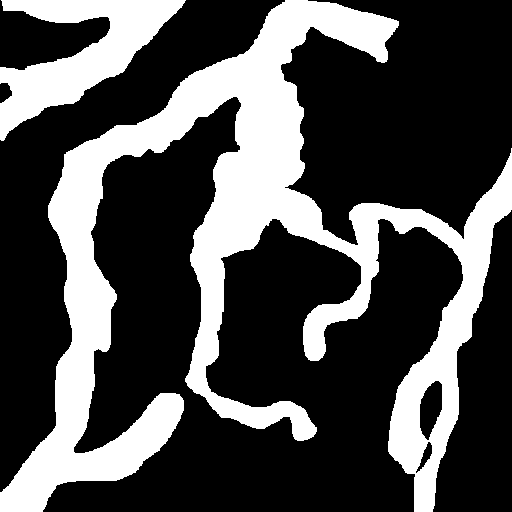

### LVMask-CCL21-16.tif

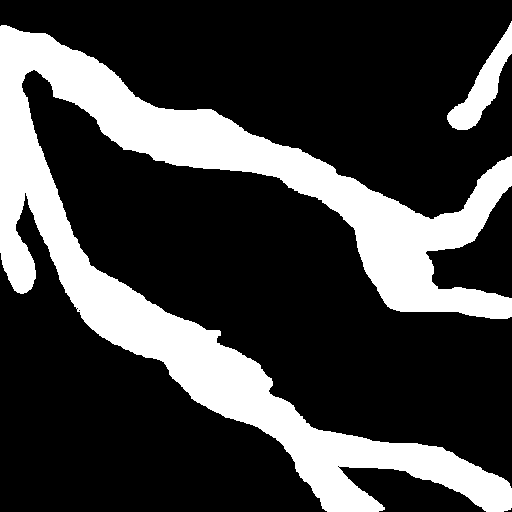
