## Supplementary Materials for "Mechanistic description of spatial processes using integrative modelling of noise-corrupted imaging data"


### 1 Two-dimensional model of CCL21 gradient formation

To describe the spatio-temporal dynamics of the concentration of soluble CCL21 ($u_1(x,t)$), of heparan sulfate ($u_2(x,t)$) and of heparan sulfate-CCL21 dimers ($u_3(x,t)$) in 2D, we consider the PDE model

$$\begin{aligned}\frac{\partial u_1(x,t)}{\partial t} - D\triangle u_1(x,t) &= \alpha \sum_l q_l(x) - k_1 u_1(x,t)u_2(x,t) + k_{-1}c(x,t) - \gamma u(x,t)\\ \frac{\partial u_2(x,t)}{\partial t} &= -k_1 u_1(x,t)u_2(x,t) + k_{-1}u_3(x,t)\\ \frac{\partial u_3(x,t)}{\partial t} &= k_1 u_1(x,t)u_2(x,t) - k_{-1}u_3(x,t)\end{aligned} \tag{1}$$

on $x \in \Omega$ with initial conditions

$$\forall x \in \Omega: \quad u_1(x,0) = 0, u_2(x,0) = s_0(x) \text{ and } x_3(x,0) = 0 \tag{2}$$

and boundary conditions

$$\forall x \in \partial\Omega: \quad \frac{\partial u_1}{\partial \nu} = 0, \tag{3}$$

for the diffusion components $u_1(x,t)$, with $\partial\Omega$ denoting the boundary of $\Omega$ and $\nu$ denoting its normal vector. For more details in the model, we refer to our previous publication (Hock *et al.*, 2013).

**Remark 1.** *A two-dimensional model appears sufficient to describe the gradient formation in mouse ear sheets, as mouse ears are rather thin.*

We consider the steady state of model (1)-(3), which is obtained by setting the time derivatives to zero. The steady state distribution of soluble CCL21, $u_{s,1}(x,t)$, fulfills the PDE

$$-D\triangle u_{s,1}(x) = \alpha \sum_l q_l(x) - \gamma u_{s,1}(x). \tag{4}$$

Division by $\gamma$ reveals that $u_{s,1}(x)$ merely depends on $D/\gamma$ and $\alpha/\gamma$. While numerical methods have to be employed to calculate the distribution of soluble CCL21 – we used the methods implemented in the MATLAB PDE toolbox –, it can be shown analytically that the solution is proportional to $\alpha/\gamma$. The distribution of soluble CCL21 shapes the steady state of heparan sulfate, $u_{s,2}(x,t)$, and of heparan sulfate-CCL21 dimers, $u_{s,3}(x,t)$, which are

$$u_{s,2}(x) = \frac{1}{\frac{u_{s,1}(x)}{K_D}+1} s_0(x) \quad \text{and} \quad u_{s,3}(x) = \frac{\frac{u_{s,1}(x)}{K_D}}{\frac{u_{s,1}(x)}{K_D}+1} s_0(x), \tag{5}$$

with dissociation constant $K_D = k_{-1}/k_1$. Indeed, the steady state distributions $u_{s,2}(x)$ and $u_{s,3}(x)$ depend merely on $s_0(x)$ and $u_{s,1}(x)/K_D$. As the dissociation constant $K_D$ has the same scaling effect as $\alpha/\gamma$, merely the effective scale parameter $\alpha/(\gamma K_D)$ needs to be considered. The measured pixel intensities

$$y_j = \int_{\Omega_j} s(u_{s,3}(x) + b)dx, \tag{6}$$

depend only on the parameters $(D/\gamma, \alpha/(\gamma K_D), s, b)$ and the overall heparan sulfate abundance $s_0(x)$ which is parameterised individually. The individual images possess different scaling constants $s$.

In addition to the pixel intensities, we consider the distance-dependent intensity of immobilised CCL21 for the 2D model. This distance-dependent intensity is obtained by averaging over all pixels which possess the same distance from the nearest lymphatic vessel. This distance-dependent intensity for the filtered and unfiltered image is depicted in Figure 1.

### 2 One-dimensional model of CCL21 gradient formation

The two-dimensional model described in the previous section can be simplified by assuming that the concentration of soluble CCL21 ($u_1(x,t)$), of heparan sulfate ($u_2(x,t)$) and of heparan sulfate-CCL21 dimers ($u_3(x,t)$) depends mostly on the distance $x$ from the next lymphatic vessel. This assumption was employed by Weber *et al.* (2013). It yields the one-dimensional

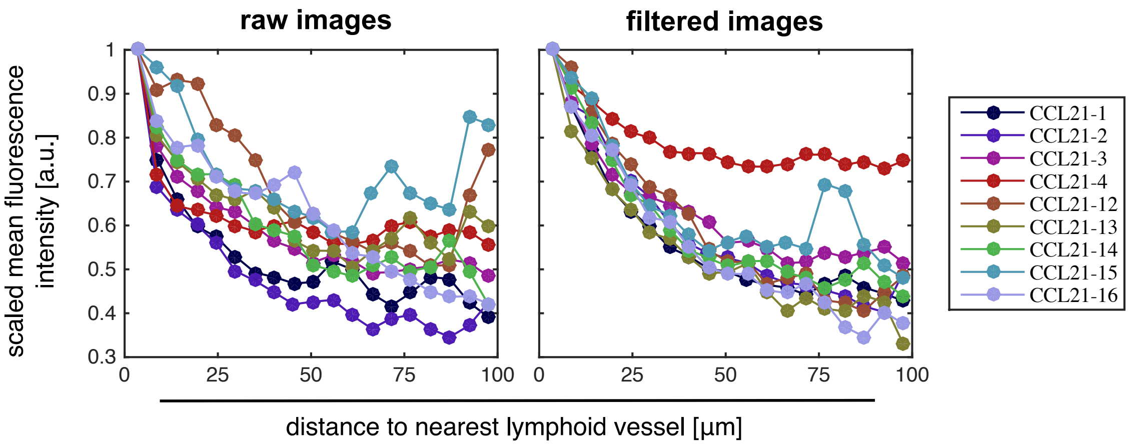


Figure 1: **Comparison for mean fluorescence intensity for raw and filter images.** (left) Mean fluorescence intensity for different distances from the nearest lymphatic vessel for the images collected by Weber *et al.* (2013). (right) Mean fluorescence intensity for different distances from the nearest lymphatic vessel for the filtered images. Maximally stable extremal region (MSER) filtering was employed to remove outliers.

PDE model

$$\begin{aligned}\frac{\partial u_1(x,t)}{\partial t} - D\triangle u_1(x,t) &= -k_1 u_1(x,t)u_2(x,t) + k_{-1}c(x,t) - \gamma u(x,t)\\ \frac{\partial u_2(x,t)}{\partial t} &= -k_1 u_1(x,t)u_2(x,t) + k_{-1}u_3(x,t)\\ \frac{\partial u_3(x,t)}{\partial t} &= k_1 u_1(x,t)u_2(x,t) - k_{-1}u_3(x,t)\end{aligned} \tag{7}$$

with $x \in [0, L]$, initial conditions

$$\forall x \in [0, L]: \quad u_1(x,0) = 0, u_2(x,0) = s_0(x) \text{ and } u_3(x,0) = 0 \tag{8}$$

and boundary conditions,

$$u_1|_{x=0} = u_{1,\mathrm{LV}} \quad \text{and} \quad \left.\frac{\partial u_1}{\partial x}\right|_{x=L} = 0. \tag{9}$$

The lymphatic vessel is localised at $x = 0$ and we consider $L \to \infty$. The concentration of soluble CCL21 in the lymphatic vessel is denoted by $u_{1,\mathrm{LV}}$. Assuming that turnover dominates diffusion, the concentration of soluble CCL21 in the lymphatic vessel is $u_{1,\mathrm{LV}} \approx \alpha/\gamma$.

We consider the steady state of model (7)-(9), which is obtained by setting the time-derivatives to zero. The steady state distribution of soluble CCL21, $u_{s,1}(x,t)$, fulfills the PDE

$$D\triangle u_{s,1}(x) = \gamma u_{s,1}(x) \tag{10}$$

with boundary conditions $u_1|_{x=0} = \frac{\alpha}{\gamma}$ and $\left.\frac{\partial u_1}{\partial x}\right|_{x=L} = 0$. This PDE possesses for $L \gg D/\gamma$, which is the case here, the analytical solution

$$u_{s,1}(x) = \frac{\alpha}{\gamma} \exp\left(-\frac{x}{\sqrt{D/\gamma}}\right). \tag{11}$$

This solution depends on two parameters, $D/\gamma$ and $\alpha/\gamma$, and is proportional to $\alpha/\gamma$. For the steady state distribution of heparan sulfate, $u_{s,2}(x,t)$, and of heparan sulfate-CCL21 dimers, $u_{s,3}(x,t)$, we find again

$$u_{s,2}(x) = \frac{1}{\frac{u_{s,1}(x)}{K_D} + 1} s_0(x) \quad \text{and} \quad u_{s,3}(x) = \frac{\frac{u_{s,1}(x)}{K_D}}{\frac{u_{s,1}(x)}{K_D} + 1} s_0(x), \tag{12}$$

yielding the distance-dependent intensity of immobilised CCL21,

$$y_j = \int_{\Omega_j} s(u_{s,3}(x) + b)dx, \tag{13}$$

which depends only on the parameters $(D/\gamma, \alpha/(\gamma K_D), s, b)$ and the overall heparan sulfate abundance $s_0(x)$.

### 3 Model parameterisation for artificial data generation

The artificial experimental data were obtained by simulating the 2D model for manually selected parameters. These parameters are provided in Table 1, along with lower and upper bounds used in the parameter estimation for the artificial data. We note that the parameters of the outlier model are not assigned during data generation, but the outliers are simulated using a more involved statistical model (see description in the manuscript).

The parameter values are provided in units of length (UL), units of concentration (UC) and units of fluorescence intensity (UI). As semi-quantitative measurements are used, the scale of the concentration is completely unknown and arbitrary. In accordance with the considered imaging data, the unit of length is $\mu m$.

### 4 Estimation results for experimental data

The parameter estimation results for the CCL21 data published by Weber *et al.* (2013) using the integrated modelling approach are reported in the Tables 2, 3 and 4. These tables provide the results for the model with constant heparan sulfate concentration (H1), the model with different heparan sulfate concentrations in lymphatic vessels and the tissue (H2) and the model with different heparan sulfate concentrations in individual lymphatic vessels and the

Table 1: **Parameters used for the generation of artificial data and the assessment of the modelling approaches.** Nominal values, lower bounds, upper bounds and units for the model parameters. The nominal values are only used for the generation of the artificial experimental data.

| | Parameter | Nominal value | Lower bound | Upper bound | Unit |
|---|---|---|---|---|---|
| dynamics | $D/\gamma$ | $8.50 \cdot 10^{+1}$ | $1.00 \cdot 10^{+0}$ | $2.20 \cdot 10^{+4}$ | UL |
| | $\alpha/(\gamma K_D)$ | $2.40 \cdot 10^{-1}$ | $6.73 \cdot 10^{-3}$ | $1.48 \cdot 10^{+2}$ | - |
| initial condition | $S_0$ | $2.10 \cdot 10^{+1}$ | $6.73 \cdot 10^{-3}$ | $1.48 \cdot 10^{+2}$ | UC |
| measurement | $b$ | $1.30 \cdot 10^{+2}$ | $2.48 \cdot 10^{-3}$ | $1.00 \cdot 10^{+0}$ | UC |
| | $\sigma$ | $1.44 \cdot 10^{-1}$ | $6.73 \cdot 10^{-3}$ | $1.00 \cdot 10^{+0}$ | - |
| | $\sigma_o$ | - | $6.73 \cdot 10^{-3}$ | $1.00 \cdot 10^{+0}$ | - |
| | $\mu_o$ | - | $3.67 \cdot 10^{-1}$ | $7.39 \cdot 10^{+1}$ | - |
| | $w = 1 - w_o$ | - | $3.67 \cdot 10^{-1}$ | $1.00 \cdot 10^{+0}$ | - |

tissue (H3), respectively. Maximum likelihood estimates were determined using multi-start local optimisation with bounds which were chosen according to the results of pre-runs.

In the Tables 2, 3 and 4 we report only the results for images 12-15 published by Weber *et al.* (2013) to enhance readability. The comparison of the maximum likelihood estimates obtained for different images is substantial and larger than the uncertainty of the parameter for the individual images as computed using profile likelihoods (results not shown). This indicates substantial biological variability between different regions of the mouse ear.

Table 2: **Maximum likelihood estimates for the model with constant heparan sulfate concentration (H1).** Lower and upper bounds used for the optimisation and results for images 12-16 (collected by (Weber *et al.*, 2013)) are reported.

| | Parameter | Lower bound | Upper bound | Maximum likelihood estimate for image | | | | | Unit |
|---|---|---|---|---|---|---|---|---|---|
| | | | | 12 | 13 | 14 | 15 | 16 | |
| dynamics | $D/\gamma$ | $1.00 \cdot 10^{+0}$ | $2.20 \cdot 10^{+4}$ | $1.30 \cdot 10^{+2}$ | $5.16 \cdot 10^{+1}$ | $2.75 \cdot 10^{+2}$ | $1.64 \cdot 10^{+2}$ | $3.24 \cdot 10^{+2}$ | UL |
| | $\alpha/(\gamma K_D)$ | $6.74 \cdot 10^{-3}$ | $1.48 \cdot 10^{+2}$ | $1.41 \cdot 10^{+0}$ | $2.04 \cdot 10^{+0}$ | $6.74 \cdot 10^{-3}$ | $6.74 \cdot 10^{-3}$ | $6.74 \cdot 10^{-3}$ | - |
| initial conditions | $S_0$ | $4.54 \cdot 10^{-5}$ | $1.00 \cdot 10^{+0}$ | $8.09 \cdot 10^{-3}$ | $7.51 \cdot 10^{-3}$ | $3.19 \cdot 10^{-1}$ | $3.26 \cdot 10^{-1}$ | $3.29 \cdot 10^{-1}$ | UC |
| measurement | $b$ | $4.54 \cdot 10^{-5}$ | $1.00 \cdot 10^{+0}$ | $2.28 \cdot 10^{-3}$ | $2.99 \cdot 10^{-3}$ | $6.03 \cdot 10^{-4}$ | $7.66 \cdot 10^{-4}$ | $4.14 \cdot 10^{-4}$ | UI |
| | $s$ | $1.00 \cdot 10^{+0}$ | $1.48 \cdot 10^{+2}$ | $9.61 \cdot 10^{+0}$ | $1.02 \cdot 10^{+1}$ | $4.11 \cdot 10^{+1}$ | $4.44 \cdot 10^{+1}$ | $3.86 \cdot 10^{+1}$ | UI/UC |
| | $\sigma$ | $2.23 \cdot 10^{-1}$ | $1.00 \cdot 10^{+0}$ | $4.56 \cdot 10^{-1}$ | $4.75 \cdot 10^{-1}$ | $4.49 \cdot 10^{-1}$ | $5.06 \cdot 10^{-1}$ | $4.24 \cdot 10^{-1}$ | - |
| | $\sigma_0$ | $3.68 \cdot 10^{-1}$ | $1.00 \cdot 10^{+0}$ | $7.45 \cdot 10^{-1}$ | $6.68 \cdot 10^{-1}$ | $7.46 \cdot 10^{-1}$ | $6.73 \cdot 10^{-1}$ | $8.00 \cdot 10^{-1}$ | - |
| | $\mu_o$ | $4.49 \cdot 10^{-1}$ | $1.22 \cdot 10^{+0}$ | $8.38 \cdot 10^{-1}$ | $8.30 \cdot 10^{-1}$ | $8.63 \cdot 10^{-1}$ | $8.22 \cdot 10^{-1}$ | $1.08 \cdot 10^{+0}$ | - |
| | $w = 1 - w_o$ | $4.49 \cdot 10^{-1}$ | $1.00 \cdot 10^{+0}$ | $6.19 \cdot 10^{-1}$ | $6.45 \cdot 10^{-1}$ | $7.05 \cdot 10^{-1}$ | $6.90 \cdot 10^{-1}$ | $5.92 \cdot 10^{-1}$ | - |

Table 3: **Maximum likelihood estimates for the model with different heparan sulfate concentrations in lymphatic vessels and the tissue (H2).** Lower and upper bounds used for the optimisation and results for images 12-16 (collected by (Weber *et al.*, 2013)) are reported.

| | Parameter | Lower bound | Upper bound | Maximum likelihood estimate for image | | | | | Unit |
|---|---|---|---|---|---|---|---|---|---|
| | | | | 12 | 13 | 14 | 15 | 16 | |
| dynamics | $D/\gamma$ | $1.00 \cdot 10^{+0}$ | $2.20 \cdot 10^{+4}$ | $7.80 \cdot 10^{+1}$ | $5.59 \cdot 10^{+1}$ | $6.21 \cdot 10^{+2}$ | $2.24 \cdot 10^{+2}$ | $8.89 \cdot 10^{+2}$ | UL |
| | $\alpha/(\gamma K_D)$ | $6.74 \cdot 10^{-3}$ | $1.48 \cdot 10^{+2}$ | $1.06 \cdot 10^{+1}$ | $4.38 \cdot 10^{+0}$ | $7.67 \cdot 10^{-2}$ | $1.84 \cdot 10^{-1}$ | $6.76 \cdot 10^{-3}$ | - |
| initial conditions | $S_T$ | $4.54 \cdot 10^{-5}$ | $1.00 \cdot 10^{+0}$ | $3.55 \cdot 10^{-3}$ | $5.13 \cdot 10^{-3}$ | $4.67 \cdot 10^{-2}$ | $2.70 \cdot 10^{-2}$ | $1.93 \cdot 10^{-1}$ | UC |
| | $S_L$ | $4.54 \cdot 10^{-5}$ | $1.00 \cdot 10^{+0}$ | $5.71 \cdot 10^{-3}$ | $6.93 \cdot 10^{-3}$ | $6.72 \cdot 10^{-2}$ | $3.42 \cdot 10^{-2}$ | $3.25 \cdot 10^{-1}$ | UC |
| measurement | $b$ | $4.54 \cdot 10^{-5}$ | $1.00 \cdot 10^{+0}$ | $2.68 \cdot 10^{-3}$ | $3.23 \cdot 10^{-3}$ | $9.27 \cdot 10^{-4}$ | $1.72 \cdot 10^{-3}$ | $2.31 \cdot 10^{-4}$ | UI |
| | $s$ | $1.00 \cdot 10^{+0}$ | $1.48 \cdot 10^{+2}$ | $7.93 \cdot 10^{+0}$ | $9.14 \cdot 10^{+0}$ | $2.35 \cdot 10^{+1}$ | $1.90 \cdot 10^{+1}$ | $5.56 \cdot 10^{+1}$ | UI/UC |
| | $\sigma$ | $2.23 \cdot 10^{-1}$ | $1.00 \cdot 10^{+0}$ | $4.48 \cdot 10^{-1}$ | $4.71 \cdot 10^{-1}$ | $4.41 \cdot 10^{-1}$ | $5.02 \cdot 10^{-1}$ | $4.04 \cdot 10^{-1}$ | - |
| | $\sigma_0$ | $3.68 \cdot 10^{-1}$ | $1.00 \cdot 10^{+0}$ | $7.46 \cdot 10^{-1}$ | $6.68 \cdot 10^{-1}$ | $7.36 \cdot 10^{-1}$ | $6.78 \cdot 10^{-1}$ | $7.99 \cdot 10^{-1}$ | - |
| | $\mu_o$ | $4.49 \cdot 10^{-1}$ | $1.22 \cdot 10^{+0}$ | $8.27 \cdot 10^{-1}$ | $8.29 \cdot 10^{-1}$ | $8.75 \cdot 10^{-1}$ | $8.07 \cdot 10^{-1}$ | $1.09 \cdot 10^{+0}$ | - |
| | $w = 1 - w_o$ | $4.49 \cdot 10^{-1}$ | $1.00 \cdot 10^{+0}$ | $6.06 \cdot 10^{-1}$ | $6.40 \cdot 10^{-1}$ | $6.99 \cdot 10^{-1}$ | $6.76 \cdot 10^{-1}$ | $5.84 \cdot 10^{-1}$ | - |

Table 4: **Maximum likelihood estimates for the model with different heparan sulfate concentrations in individual lymphatic vessels and the tissue (H3).** Lower and upper bounds used for the optimisation and results for images 12-16 (collected by (Weber *et al.*, 2013)) are reported. The different images possess different numbers of lymphatic vessels.

| | Parameter | Lower bound | Upper bound | Maximum likelihood estimate for image 12 | 13 | 14 | 15 | 16 | Unit |
|---|---|---|---|---|---|---|---|---|---|
| dynamics | $D/\gamma$ | $1.00 \cdot 10^{0}$ | $2.20 \cdot 10^{+4}$ | $7.44 \cdot 10^{+1}$ | $5.76 \cdot 10^{+1}$ | $4.45 \cdot 10^{+2}$ | $2.17 \cdot 10^{+2}$ | $8.90 \cdot 10^{+2}$ | UL |
| | $\alpha/(\gamma K_D)$ | $6.74 \cdot 10^{-3}$ | $1.48 \cdot 10^{+2}$ | $1.16 \cdot 10^{+1}$ | $4.01 \cdot 10^{0}$ | $4.73 \cdot 10^{-1}$ | $2.09 \cdot 10^{-1}$ | $6.75 \cdot 10^{-3}$ | - |
| initial conditions | $S_T$ | $4.54 \cdot 10^{-5}$ | $1.00 \cdot 10^{0}$ | $5.31 \cdot 10^{-3}$ | $9.71 \cdot 10^{-3}$ | $2.95 \cdot 10^{-2}$ | $4.08 \cdot 10^{-2}$ | $3.16 \cdot 10^{-1}$ | UC |
| | $S_{L,1}$ | $4.54 \cdot 10^{-5}$ | $2.20 \cdot 10^{+4}$ | $8.75 \cdot 10^{-3}$ | $8.54 \cdot 10^{-3}$ | $3.43 \cdot 10^{-2}$ | $6.82 \cdot 10^{-2}$ | $1.05 \cdot 10^{0}$ | UC |
| | $S_{L,2}$ | $4.54 \cdot 10^{-5}$ | $2.20 \cdot 10^{+4}$ | $7.43 \cdot 10^{-3}$ | $1.13 \cdot 10^{-2}$ | $4.89 \cdot 10^{-2}$ | $5.10 \cdot 10^{-2}$ | $5.31 \cdot 10^{-1}$ | UC |
| | $S_{L,3}$ | $4.54 \cdot 10^{-5}$ | $2.20 \cdot 10^{+4}$ | – | $1.57 \cdot 10^{-2}$ | $3.72 \cdot 10^{-2}$ | – | – | UC |
| | $S_{L,4}$ | $4.54 \cdot 10^{-5}$ | $2.20 \cdot 10^{+4}$ | – | – | $4.16 \cdot 10^{-2}$ | – | – | UC |
| measurement | $b$ | $4.54 \cdot 10^{-5}$ | $1.00 \cdot 10^{0}$ | $4.11 \cdot 10^{-3}$ | $5.88 \cdot 10^{-3}$ | $3.49 \cdot 10^{-3}$ | $2.94 \cdot 10^{-3}$ | $3.76 \cdot 10^{-4}$ | UI |
| | $s$ | $6.74 \cdot 10^{-3}$ | $1.48 \cdot 10^{+2}$ | $5.18 \cdot 10^{0}$ | $5.10 \cdot 10^{0}$ | $6.56 \cdot 10^{0}$ | $1.12 \cdot 10^{+1}$ | $3.41 \cdot 10^{+1}$ | UI/UC |
| | $\sigma$ | $6.74 \cdot 10^{-3}$ | $1.00 \cdot 10^{0}$ | $4.48 \cdot 10^{-1}$ | $4.79 \cdot 10^{-1}$ | $4.37 \cdot 10^{-1}$ | $5.02 \cdot 10^{-1}$ | $4.03 \cdot 10^{-1}$ | - |
| | $\sigma_o$ | $6.74 \cdot 10^{-3}$ | $1.00 \cdot 10^{0}$ | $7.47 \cdot 10^{-1}$ | $6.68 \cdot 10^{-1}$ | $7.42 \cdot 10^{-1}$ | $6.78 \cdot 10^{-1}$ | $7.99 \cdot 10^{-1}$ | - |
| | $\mu_o$ | $3.68 \cdot 10^{-1}$ | $1.22 \cdot 10^{0}$ | $8.24 \cdot 10^{-1}$ | $8.19 \cdot 10^{-1}$ | $8.63 \cdot 10^{-1}$ | $8.09 \cdot 10^{-1}$ | $1.09 \cdot 10^{0}$ | - |
| | $w = 1 - w_o$ | $3.68 \cdot 10^{-1}$ | $1.00 \cdot 10^{0}$ | $6.05 \cdot 10^{-1}$ | $6.57 \cdot 10^{-1}$ | $6.96 \cdot 10^{-1}$ | $6.78 \cdot 10^{-1}$ | $5.84 \cdot 10^{-1}$ | - |
